# Chromosome-scale genome assembly of the diploid oat *Avena longiglumis* reveals the landscape of repetitive sequences, genes and chromosome evolution in grasses

**DOI:** 10.1101/2022.02.09.479819

**Authors:** Qing Liu, Hongyu Yuan, Mingzhi Li, Ziwei Wang, Dongli Cui, Yushi Ye, Zongyi Sun, Xukai Tan, Trude Schwarzacher, John Seymour Heslop-Harrison

**Author notes:** Correspondence (QL);, (JSHH).

## Abstract

**Background:** Oat (*Avena sativa*, 2*n*=6*x*=42) is an important crop, and with its wild relatives including *A. longiglumis* (ALO, 2*n*=6*x*=14), has advantageous agronomic and nutritional traits. A *de-novo* chromosome-level ALO genome assembly was made to investigate diversity and structural genome variation between *Avena* species and other Poaceae in an evolutionary context, and develop genomic resources to identify the pangenome and economic traits within Pooideae.

**Results:** The 3.85 gigabase ALO genome (seven pseudo-chromosomes), contained 40,845 protein-coding genes and 87% repetitive sequences (84.21% transposable elements). An LTR retrotransposon family was abundant at all chromosome centromeres, and genes were distributed without major terminal clusters. Comparisons of synteny with *A. eriantha* and *A. strigosa* showed evolutionary translocations of terminal segments including many genes. Comparison with rice (*x*=12) and the ancestral grass karyotype showed synteny and features of chromosome evolution including fusions, translocations and insertions of syntenic blocks across Pooideae species. With a genome size 10 times larger than rice, ALO showed relatively uniform expansion along the chromosome arms, with few gene-poor regions along arms, and no major duplications nor deletions. Linked gene networks were identified (mixed-linkage glucans and cellulose synthase genes), and *CYP450* genes may be related to salt-tolerance.

**Conclusions:** The high-continuity genome assembly shows gene, chromosomal structural and copy number variation, providing a reference for the *Avena* pangenome, defining the full spectrum of diversity. Chromosomal rearrangements and genome expansion demonstrate features of evolution across the genus and grass BOP-clade, contributing to exploitation of gene and genome diversity through precision breeding.

## Background

In the Poaceae, cultivated common oat (*Avena sativa* L.; 2*n* = 6*x* = 42, AACCDD) and its wild relatives (2*x*, 4*x* and 6*x*) belong to the Aveneae tribe, which diverged from the Triticeae (syn. Hordeae) tribe (including wheat, barley and rye) around 27.8 million years ago (Mya) [1], and are part of the BOP- or BEP-clade [Bambusoideae, Oryzoideae (syn. Ehrhartoideae), and Pooideae] including half of all grasses, separating 70 Mya from the other grass lineages [2, 3, 4, 5]. Global production of oat reached 22 million tons in 2021 (http://www.fao.org/faostat/). Human studies have demonstrated the beneficial effects of consuming oats for the reduction of serum cholesterol and cardiovascular disease, associated with the soluble β-glucan component [6], and a favourable glycaemic index with a low value and slow carbohydrate breakdown. Oat has substantial concentrations of phytochemicals (e.g., avenine, avenacin and phenolic compounds) and tolerance to the harsh environments such as sandy loam soil, short growing seasons and desert climate, making the crop resilient [7]. The oil content of oat grain (6%) is high among cereals, suggesting a possible important future use, like maize, for food oils.

Due not least to the large genome size of *A. sativa* [1C genome size ∼12.5 gigabases (Gbp) [8], oat genomics has lagged behind those of other crops such as rice (*Oryza sativa*) [9], sorghum (*Sorghum bicolor*) [10] or foxtail millet (*Setaria italica*) [11] although there are increasing amounts of oat genome sequence available in databases [7, 12]. Exploitation and utilization of germplasm resources preserved in wild oat species are a pressing need for oat and related crop breeding.

The genus *Avena* contains about 25 species, including several edible species and invasive weeds, with characteristic erect culms and solitary spikelets on a panicle, and distribution throughout temperate regions of the Mediterranean Basin, Africa, Europe, Asia, Australia and the Americas [13]. Extensive chromosomal rearrangements following recurrent polyploidy events [14, 15] may have increased oat genomic variation and provided a selective advantage in the adaptation to changing growth environments. A-genome diploid *A. longiglumis* (ALO) has important traits including the high content of linoleic content in the grains [16], drought-adapted phenotypes [17], and resistance to crown rust disease [18]. The known genetic resources from wild diploid *Avena* species are limited, which impedes progress on understanding the genomic variation related to responses to biotic and abiotic stressors as well as quality traits. Beyond variation in genes and regulatory sequences, structural and copy number variation (CNV) has proved difficult to assess with short-read sequencing but its importance in control of complex traits in both farm animal [19] and plant [20] breeding is increasingly recognized and must be characterized as part of the pangenome [21].

The diploid *Avena* genomes, similar in size to the wheat group but ten times larger than rice [9] (Ouyang et al., 2007), are characterized by a high proportion of repetitive DNAs, i.e., interspersed repeats including transposable elements (TEs) and tandem repeats [22]. Along with the frequent chromosome translocations [22], the repetitive DNA makes complete genome assembly difficult. Growing evidence suggests that repetitive stretches of DNAs may cause sequencing breakage or genome assembly collapse [23]. Now, long-read sequencing technologies (eg. Oxford Nanopore Technologies, ONT) combined with genome scaffolding methods (eg. high-throughput chromatin conformation capture, Hi-C), together with Illumina short-reads used for sequence correction and optical mapping, have improved the genome contiguity and repeat annotation integrity [24]. For example, hybrid ONT/Illumina study revealed the genomic landscape among diploid *Brassica* species in unprecedented detail [25] and allowed *de novo* genome assembly of *Ensete glaucum*, identification of repeats and chromosomal rearrangements to the related genus *Musa* [26]. High-continuity assembly of large genomes with a high proportion of repeats is becoming possible [27, 28] and enables comparison between crops and their wild relatives as well as definition of the pangenome.

Here, we generate a high-quality *Avena longiglumis* ALO genome assembly by integration of Illumina, ONT and Hi-C data, aiming to uncover not only the range of genes and regulatory elements but also chromosomal rearrangements of wild oat relatives including features of genome expansion and structural variation. Besides showing structural variation within *Avena* species, we aimed to identify any intra- and inter-chromosomal rearrangements compared with the distantly related grasses rice and *Brachypodium distachyon* in the Pooideae. We aimed to identify gene families that expanded and contracted in ALO and nine grass species genomes, as well as the potential biosynthetic gene clusters that may be play a role in salt stress response and β-glucan biosynthesis of wild oat species.

## Results

### Genome assembly and annotation

The *Avena longiglumis* (ALO, 2*n* = 14) genome size was estimated to be 4.60 ± 0.11 Gbp/1C by flow cytometry (FCM, Additional file 1: Fig. S1), this value is similar to 4.7 Gbp reported by Yan et al. [8] and 3.97 Gbp we calculate by *k*-mer analysis (*k* = 17 using raw reads of Illumina data (Additional file 1: Fig. S2). The ALO genome assembly size we report here is 3.96 Gbp (Table 1), and 3.85 Gbp (97.14%) were anchored into seven pseudo-chromosomes (Fig. 1; Table 2).

**Fig. 1.**
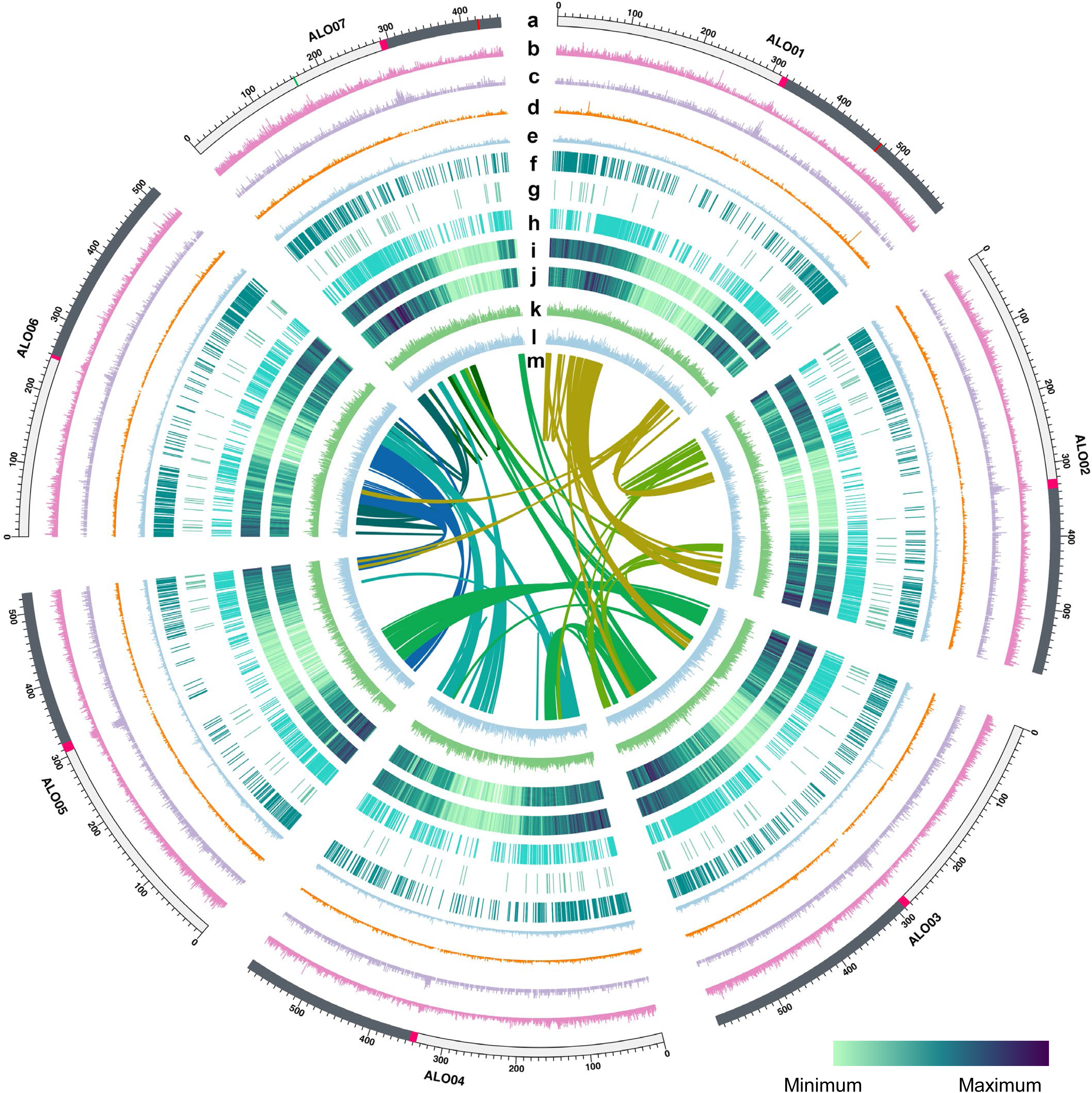
Genomic landscape of seven assemble chromosomes ALO01 to ALO07 of *Avena longiglumis* (ALO). **a** Chromosome names and sizes (100 Mbp intervals indicated) with centromere position marked in pink and major 5S (ALO07 in green) and 45S (ALO01 and ALO07 in red) rDNA sites indicated. **b** Transposable element (TE, pink) density along each chromosome. **c** LTR TE (purple) density (1 Mbp nonoverlapping windows) along each chromosome. **d** Long interspersed nuclear element (*LINE*) density (orange) along each chromosome. **e** *Helitron* density (cyan) along each chromosome. **f** Expanded gene locations in each chromosome. **g** Contracted gene locations in each chromosome. **h** Single copy orthologue gene locations in each chromosome. **i** High-confidence gene locations in each chromosome. **j** Purified selection gene locations in each chromosome (these genes with *P*-value ≤ 0.05). **k** Expression profiling of genes on each chromosome in ALO roots. **l** Expression profiling of genes on each chromosome in ALO leaves. **m** Links between syntenic genes. Orientation in outward in circles **b**, **c**, **d**, **e**, **k** and **l**.

**Table 1.**
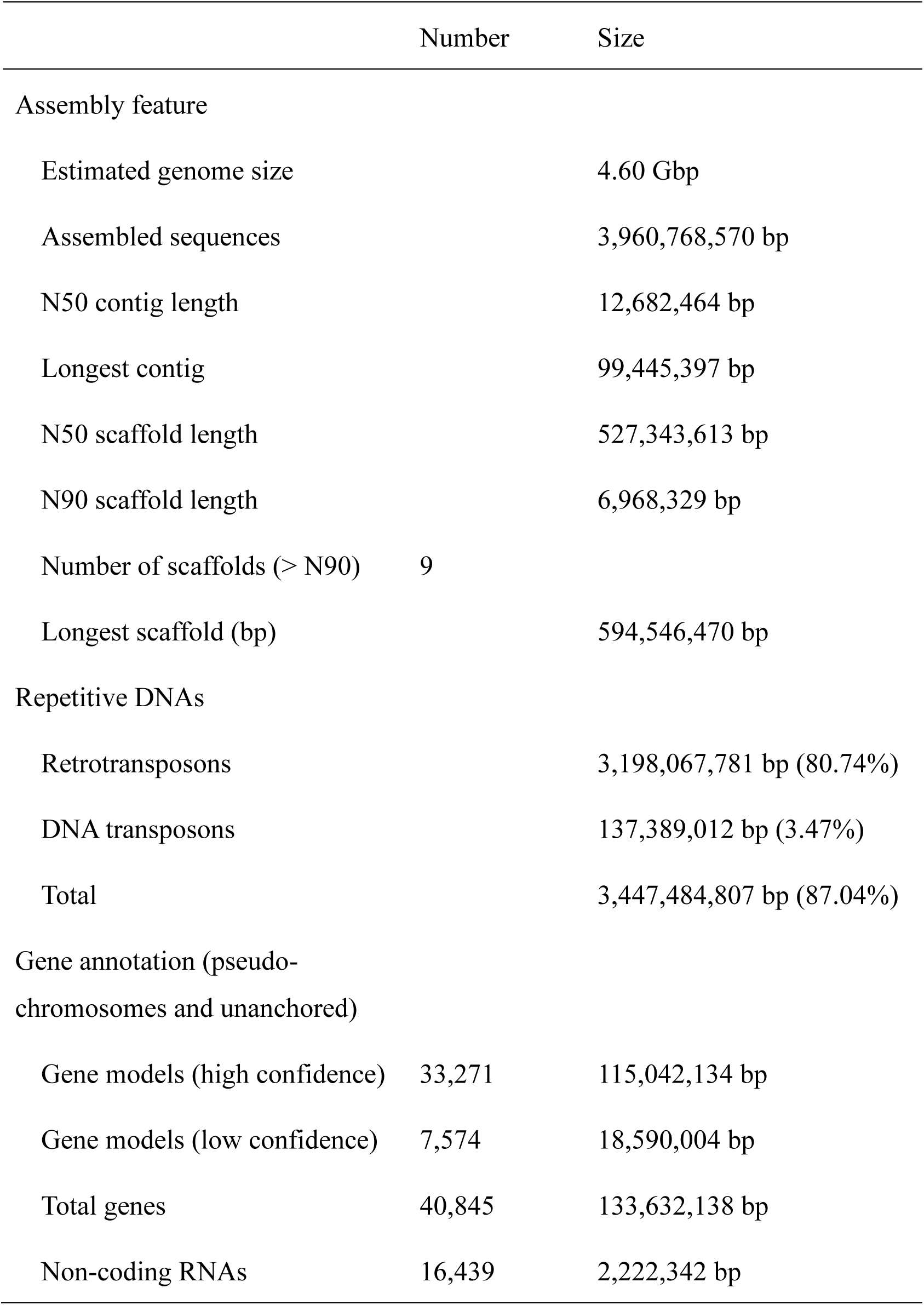
*Avena longiglumis* genome statistics and gene predictions.

|  | Number | Size |
| --- | --- | --- |
| Assembly feature |  |  |
| Estimated genome size |  | 4.60 Gbp |
| Assembled sequences |  | 3,960,768,570 bp |
| N50 contig length |  | 12,682,464 bp |
| Longest contig |  | 99,445,397 bp |
| N50 scaffold length |  | 527,343,613 bp |
| N90 scaffold length |  | 6,968,329 bp |
| Number of scaffolds (> N90) | 9 |  |
| Longest scaffold (bp) |  | 594,546,470 bp |
| Repetitive DNAs |  |  |
| Retrotransposons |  | 3,198,067,781 bp (80.74%) |
| DNA transposons |  | 137,389,012 bp (3.47%) |
| Total |  | 3,447,484,807 bp (87.04%) |
| Gene annotation (pseudo-chromosomes and unanchored) |  |  |
| Gene models (high confidence) | 33,271 | 115,042,134 bp |
| Gene models (low confidence) | 7,574 | 18,590,004 bp |
| Total genes | 40,845 | 133,632,138 bp |
| Non-coding RNAs | 16,439 | 2,222,342 bp |

**Table 2.** Pseudo-chromosome length and gene content of Avena longiglumis (ALO)

| Chromosome number | Pseudomolecular length (bp) | Arm ratio* | Gene number** | Mean length (bp) | Median length (bp) | Minimum length (bp) | Maximum length (bp) | High confidence gene number*** |
| --- | --- | --- | --- | --- | --- | --- | --- | --- |
| ALO01 | 594,546,470 | 1.24 | 6,371 | 3370.96 | 1932 | 163 | 254,605 | 5,186 |
| ALO02 | 587,543,788 | 1.14 | 5,976 | 3229.50 | 1928.5 | 165 | 158,444 | 4,814 |
| ALO03 | 587,190,583 | 1.07 | 6,417 | 3298.18 | 2069 | 150 | 105,522 | 5,323 |
| ALO04 | 583,925,327 | 1.07 | 5,846 | 3262.89 | 2018.5 | 163 | 169,716 | 4,895 |
| ALO05 | 527,343,613 | 1.39 | 4,212 | 3339.13 | 2005 | 201 | 145,603 | 3,493 |
| ALO06 | 513,337,126 | 1.08 | 5,413 | 3272.85 | 1997 | 163 | 154,734 | 4,375 |
| ALO07 | 453,691,697 | 2.01 NOR | 5,256 | 3193.56 | 1975.5 | 158 | 165,822 | 4,100 |
| Anchored genome | 3,847,578,604 |  | 39,491 | 22967.07 | 13925.5 | 1163 | 1,154,446 | 32,186 |
| Unanchored | 113,189,966 |  | 1,354 |  |  |  |  | 1,085 |

\* Arm ratio = (long arm length)/(short arm length); NOR has secondary constriction at the major Nucleolar Organizer Region on short arm.
\*\* Number of genes anchored on chromosomes.
\*\*\* Number of high confidence genes supported by transcriptome data with FPKM value larger than zero or homology. For the genes without transcriptome transcript abundance support, the alignment was performed with *A. atlantica* (identity > 95%, coverage > 95%), *A. eriantha* (identity > 90%, coverage > 90%), *Hordeum vulgare* and *Triticum aestivum* (identity > 80%, coverage > 80%) by BLASTP (https://blast.ncbi.nlm.nih.gov/Blast.cgi; *E* value = 1e-5), respectively. Those supported by alignment results of two or more species alignments were defined as high confidence genes.

Our assembly strategy is shown in Additional file 1: Fig. S3 with 67.55 × Illumina reads (150 bp paired-end), 63.82 × Nanopore and 99.55 × Hi-C sequencing data (Table 1 [summary of assembly], Additional file 1: Figs. S2c [sequence depth], d [N50], S3 [assembly strategy] and S4 [HiC contact map], Additional file 2: Tables S1 [libraries], S2 [programs] and S3 [details of all assemblies). A total of 252.78 Gbp qualified ONT read sequences (mean_qscore ≥ 7) were gained from 12 libraries (Additional file 2: Table S1). After self-correction, the 128 Gbp of read sequences generated 2,379 contigs with a first-pass assembly of 3.71 Gbp with N50 read length 11.92 megabases (Mbp). The ONT raw data and Illumina whole-genome shotgun sequencing data were used for interactive error correction three or four times, yielding a 3.96 Gbp polished assembly with a contig N50 of 12.68 Mbp and the longest contig of 99,445,397 bp (summary in Table 1, details in Table S3). We further used 394.30 Gbp Hi-C data to improve the assembly and anchor 1,974 of 2,379 scaffolds into 414 super-scaffolds (Additional file 2: Table S3). Super-scaffolds were clustered and ordered into a chromosome-scale assembly with seven chromosomes, ranging from 453.69 to 594.55 Mbp (Fig. 1, circle a, Table 2). The chromosome-scale assembly was 3.85 Gbp (97.14% of the with a super-scaffold N50 of 583.93 Mbp, 97.14% of genomic sequences (Table 2 and supplementary Table S3) were assigned to discrete chromosome locations using Hi-C assembly (Additional file 1: Fig. S4, Additional file 2: Table S3). ALO sequencing data were uploaded to National Genomic Data Center (https://bigd.big.ac.cn/bioproject/) under the accession no. PRJCA004488/CRR275304-CRR275326, CRR285670-285674. The ALO genome assembly was uploaded to https://figshare.com/s/34d0c099e42eb39a05e2.

Genome completeness was evaluated by several approaches. Of the Illumina reads, 99.94% (1,348,307,165 of 1,349,168,075) could be mapped onto the assembly after filtering chloroplast/mitochondrial/bacterial/fungal/human reads. Benchmarking Universal Single-Copy Orthologs (BUSCO) [29] identified the complete BUSCOs (C, 95.35%), complete and single-copy BUSCOs (S, 81.02%), complete and duplicated BUSCOs (D, 14.33%), fragmented BUSCOs (F, 1.16%) and missing BUSCOs (M, 3.49%), respectively (Additional file 2: Table S5a). With Conserved Core Eukaryotic Gene Mapping (CEGMA) [30], our assembly captured 243 of 248 (98.0%) conserved core eukaryotic genes from CEGMA [30], and 241 (97.18%) of these were complete [Additional file 2: Tables S4 (assembly consistency statistics), S5a (BUSCO analysis), b (CEGMA analysis)]. Assembly base accuracy was also assessed based on Illumina short read mapping. In total, 90.53% of RNA-seq reads were uniquely mapped to the genome assembly (Additional file 2: Table S5c). All of these evaluations indicate the high completeness, high continuity and high base accuracy of the genome assembly.

The long terminal repeat (LTR) Assembly Index (LAI) [31] evaluates the contiguity of intergenic and repetitive regions of genome assemblies based on the intactness of LTR retrotransposons (LTR-RTs, Additional file 2: Table S6a, b). The LAI value of the ALO genome assembly was 10.54, which was higher than that of *Aegilops tauschii* (ATA) [32], and lower than those of *A. strigosa* (AST, 11.51) [7], *Brachypodium distachyon* (BDI, 11.08) [33], *Oryza sativa* (OSA, 21.95) [9], *Sorghum bicolor* (SBI, 13.91) [10], *Setaria italica* (SIT, 17.44) [11] and *Zea mays* (ZMA, 25.57) [34] genomes (Additional file 1: Fig. S5). The LAI indicates the 3.85 Gbp genome assembly is high quality.

A heterozygosity rate of 0.48% was estimated from the frequency peaks of 17-mers from the Illumina reads following [35], with a main peak depth of 52 [Additional file 1: Fig. S2c (sequence depth), Additional file 2: Table S4a (Illumina read statistics)]. From mapping Illumina reads to the assembly, we found 3,914,721 heterozygous SNPs and 185,546 heterozygous indels (4,007,839 polymorphisms / 3,960,768,570 bp) giving a heterozygosity in single-copy regions of 0.10%, or one polymorphism per kbp (Additional file 2: Tables S4b). The extremely low value (99.9% homozygous) suggests the species (accession PI 657387) is strongly inbreeding and this is consistent with reports in other diploid *Avena* species (0.07% heterozygosity in *A. atlantica* and 0.12% in *A. eriantha*) [12].

### Identification and chromosomal distribution of transposable elements

A total of 3.45 Gbp, representing 87% of the ALO genome assembly, could be classified as repetitive DNAs using EDTA v.1.7.0 [36]. It included 3.34 Gbp (84%) as 2,205,936 complete or fragmented TEs (Additional file 2: Table S6a) and corresponds to the genome proportion in other *Avena* species [22]. The overall chromosomal distribution of TEs was relatively uniform along chromosomes (Fig. 1, circles b, c, d, except for centromeric domains), with no notable depletion in terminal or gene-rich regions (contrasting with other species with both large and small genomes such as for example wheat [32] or *Ensete* [26]) where there are abundant transposons and few genes in broad centromeric regions. In ALO, retrotransposons (Class I TEs) and their fragments were the dominant and accounted for 94.71% of the TE content (79.76% of the ALO assembly), with LTR retroelements (*Gypsy* and *Copia*) occupying 51.65% and 26.15% of the ALO assembly, respectively (Fig. 2a, Additional file 2: Table S6a). Among Class II TEs (DNA transposons), *Helitron* was the most abundant class, constituting 1.11% of the ALO assembly (Fig. 1, circle e, Additional file 2: Table S6a. There was a notable small but sharp increase in density of LTR elements at the centromere of all seven chromosomes (Fig. 1, circle c), with a uniform distribution along the rest of chromosomes. Comparison with Liu et al. [22] (their Fig. 5d) shows that a tandemly repeated sequence Ab_T105, shared widely among *Avena* species, was localized at the centromeres of all chromosomes.

**Fig. 2.**
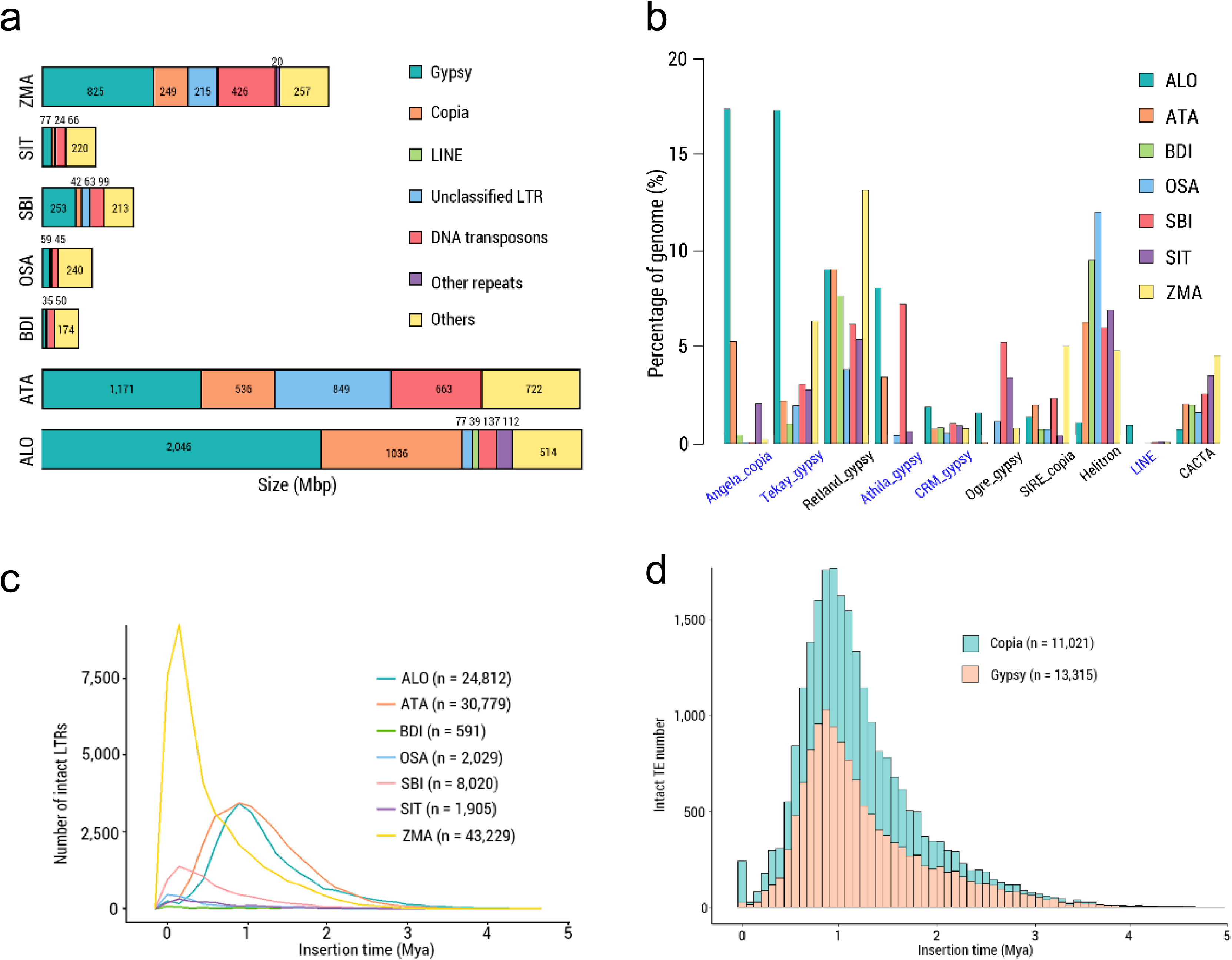
Analysis of TEs in *Avena longiglumis* (ALO) genome. **a** Genomic constituent in ALO in comparison with those in ATA, BDI, OSA, SBI, SIT and ZMA. Note that the six constituents, especially *Gypsy*, *Copia* and unclassified LTR TEs, were much more abundant in ALO than in other grasses. **b** Top 10 TE families in ALO and the percentages of these families in ATA, BDI, OSA, SBI, SIT and ZMA. Five *Gypsy* families, *Angela*, *Tekay*, *Retland*, *Athila* and *CRM*, showed increased abundance in ALO relative to those in ATA, BDI, OSA, SBI, SIT and ZMA. **c** Temporal patterns of LTR-RT insertion bursts in ALO as compared to those in ATA, BDI, OSA, SBI, SIT and ZMA. The number of intact LTR-RTs used for each species is given in parentheses. **d** Insertion bursts of *Gypsy* and *Copia* elements in ALO. The numbers of intact elements used for this analysis are provided in parentheses. *Avena longiglumis* (ALO), *Aegilops tauschii* (ATA) [32], *Brachypodium distachyon* (BDI) [33], *Oryza sativa* (OSA) [9], *Sorghum bicolor* (SBI) [10], *Setaria italica* (SIT) [11] and *Zea mays* (ZMA) [34].

Cross-genome comparisons with ATA, BDI, OSA, SIT, SBI and ZMA showed that the ALO TE content was similar to other published Poaceae genomes analysed with parallel data and software (Fig. 2a, Additional file 2: Table S6): ATA (81.71% TEs 3.5 Gbp genome size) and ZMA (81.34%, 1.9 Gbp), and a higher proportion than those of grasses with smaller genomes [< 1 Gbp [37]; BDI (35.75%, 93.0 Mbp), OSA (47.56%, 185.0 Mbp), SBI (67.83%, 495.2 Mbp), SIT (43.48%, 183.9 Mbp), but in all cases LTR-*Gypsy* elements were more frequent than LTR-*Copia* elements]. Comparison of the absolute and relative repetitive sequence composition (Fig. 2a) shows total retrotransposons (*Gypsy*, *Copia*, LINE and unclassified) were more abundant than DNA transposons in the species analysed, but, relative to other species, ALO had a lower proportion of DNA transposons (Fig. 2a, red). The ten most abundant TE families (five *Gypsy*, two *Copia*, *LINE*, and two DNA TEs) together represented about 59.66% of the ALO assembly, with the most abundant elements being from *Angela*, a family of autonomous *Copia* retrotransposons comprising 17.39% of the ALO assembly (Fig. 2b). Of the elements identified, 65% more *Copia* were intact (11,021; fragments and intact elements representing 33% of LTR-elements) compared to *Gypsy* (13,315; 65%). Two LTR-RT (intact) families, *Tekay*-*Gypsy* and *Athila*-*Copia* exhibited elevated abundance in ALO relative to other grasses (Fig. 2b, Additional file 2: Table S6c).

A unimodal distribution was found for the insertion times of all intact LTR-RTs in analysed grasses (Fig. 2c). The non-domesticated, temperate species with large genomes, ALO and ATA had the peak of amplification at around 1 Mya and a moderate proportion of older LTR-RT insertions; a younger peak, occurring approximately 0.1–0.4 Mya, was seen in OSA, SBI and ZMA (Fig. 2c). The insertion peaks overall have occurred long after the 12 Mya for ALO speciation and 20 Mya for *Avena* species separation [13, 38], but long before strong domestication pressures in 9000 to 2000 years ago [39]. Estep et al. [40] emphasized the contrasting behaviour of individual retroelement families, and indeed in ALO, bursts of both *Gypsy* and *Copia* show peaks at 1 Mya, but there is a notable additional recent burst of *Copia* elements only (Fig. 2d). Amplification of retrotransposons (*Copia*, notably *Angela*, and *Gypsy*, notably *Tekay*, Fig. 2b) contributed directly to the ALO genome expansion. Consistent with our analysis, previous studies showed that ancestral TE families followed independent evolutionary trajectories among related species, highlighting the evolution of TE populations as a key factor of genome expansion [40], and the differential dynamics of TE families within and between species.

Centromere locations were identified by multiple genomic features, some discussed in detail below. The location of the centromeric retrotransposon *Cereba* [41], SynVisio [42] results of gaps and conserved regions between *Avena* and other species assemblies, discontinuities in the Hi-C contact map (Additional file 1: Fig. S4), regions with a relatively high abundance of TEs (Fig. 1, circles b, c, d), regions of low gene density (Fig. 1, circle i, Additional file 1: Fig. S6), and examination of chromosome morphology with metacentric, sub-metacentric, and unequal-armed chromosomes, and the secondary constriction at the NOR, all gave consistent centromere core localizations on chromosomes (Additional file 1: Fig. S7, Additional file 2: Table S7a, b, c).

### Identification and chromosomal distribution of genes

Through a combination of *ab initio* prediction, homology searches, and RNA-seq-aided prediction, 40,845 protein-coding genes were identified in the ALO genome. Compared with other published Poaceae genomes, the number of genes in *A*. *longiglumis* is similar but slightly greater than that in BDI, OSA, SBI, SIT and ZMA. The mean gene and exon lengths of ALO genes were 3,272 bp and 268 bp (4.59 exons per gene), respectively (Table 2, Additional file 2: Table S8). A total of 39,558 (96.85%) protein-coding genes were assigned functions, and 86.82% of these genes exhibited homology protein domains in COG (Clusters of Orthologous Groups of proteins [43]), further 78.59% of these genes exhibited homology protein domains in Swiss-Prot database. Most of the genes were annotated with the non-redundant protein (NR) sequence database (96.44% in NCBI NR), and 93.23% of the genes were annotated in NOG (Non-supervised Orthologous Groups). The 87.09% of genes were annotated with Pfam [44, 45] annotation, 51.81% being classified according to GO terms [46], 42.32% being mapped to known plant biological pathways based on the KEGG pathway database [47], 5.88% being annotated in PlantTFDB v.5.0 [48] and 2.28% in CAZy database [49] (Additional file 2: Table S9). In addition, we predicted at least 17,712 noncoding RNAs consisting of transfer RNAs (0.0027%), microRNAs (0.0532%), and small nuclear RNAs (0.0016%) (Additional file 2: Table S10). A total of 33,271 (81.46%) high-confidence (HC) and 7,574 (18.25%) low-confidence (LC) protein-coding genes were annotated based on *de novo* prediction, homology annotation, and RNA sequence data (Table 1, Additional file 2: Table S11a, b, c).

Ribosomal DNAs (rDNAs) were collapsed in the assembly. The 45S rDNA monomer was 10,215 bp long, representing 0.30% of the genome (1,160 copies) and located on ALO07 (*A*. *longiglumis* chromosome 7) around 429,873,253 bp with a minor site on chromosome ALO01 at 476,793,000 bp (Fig. 1, circle a). The 5S rDNA monomer was 314 bp long, representing 0.005% of the genome (584 copies), with a single locus located on chromosome ALO07 around 173,926,000 bp.

Gene density along chromosomes varied, with broad regions both sides of the centromeres being depleted of genes (Fig. 1, circle i) in six of the seven chromosomes. Notably, ALO07 with the major NOR region (45S rDNA site) had a different pattern: on the long arm, there was a similar density of genes along most of the arm. Very few genes were identified between the centromere and NOR locus, while the satellite had a high gene density.

### Genome evolutionary and whole genome duplication (WGD) analysis

#### Orthologous genes in *Avena* and related grasses

We clustered the annotated genes into gene families among ALO and nine grass species (AAT, *Avena eriantha* (AER) [12], AST, ATA, BDI, OSA, SBI, SIT and ZMA) with *Arabidopsis thaliana* (ATH) [50] as the outgroup using Orthofinder v.2.3.14 [51]. A total of 1,880 single-copy genes (Additional file 2: Tables S12 and S13) were identified among 11 species, which were used for phylogenetic reconstruction (Fig. 3a, b). ALO is sister to the lineage of AAT and AST, and in turn clustered with AER, ATA, BDI and OSA, subfamily Pooideae (Fig. 3a, b, Additional file 2: Table S14). We found that ALO diverged phylogenetically from the lineage of AAT and AST at 2.34 (1.17–3.69) Mya after the divergence of *Avena* at 22.46 (15.12–29.08) Mya (Fig. 3a). The divergence times are consistent with phylogenies based on morphology or chloroplast and single-copy nuclear genes, showing when *Avena* diverged from ATA (20.04 Mya) [13] or wheat (19.90 Mya) [38]. The C-genome diploid *A. eriantha* diverged from A-genome diploid species at 9.34 (5.29–13.48) Mya.

**Fig. 3.**
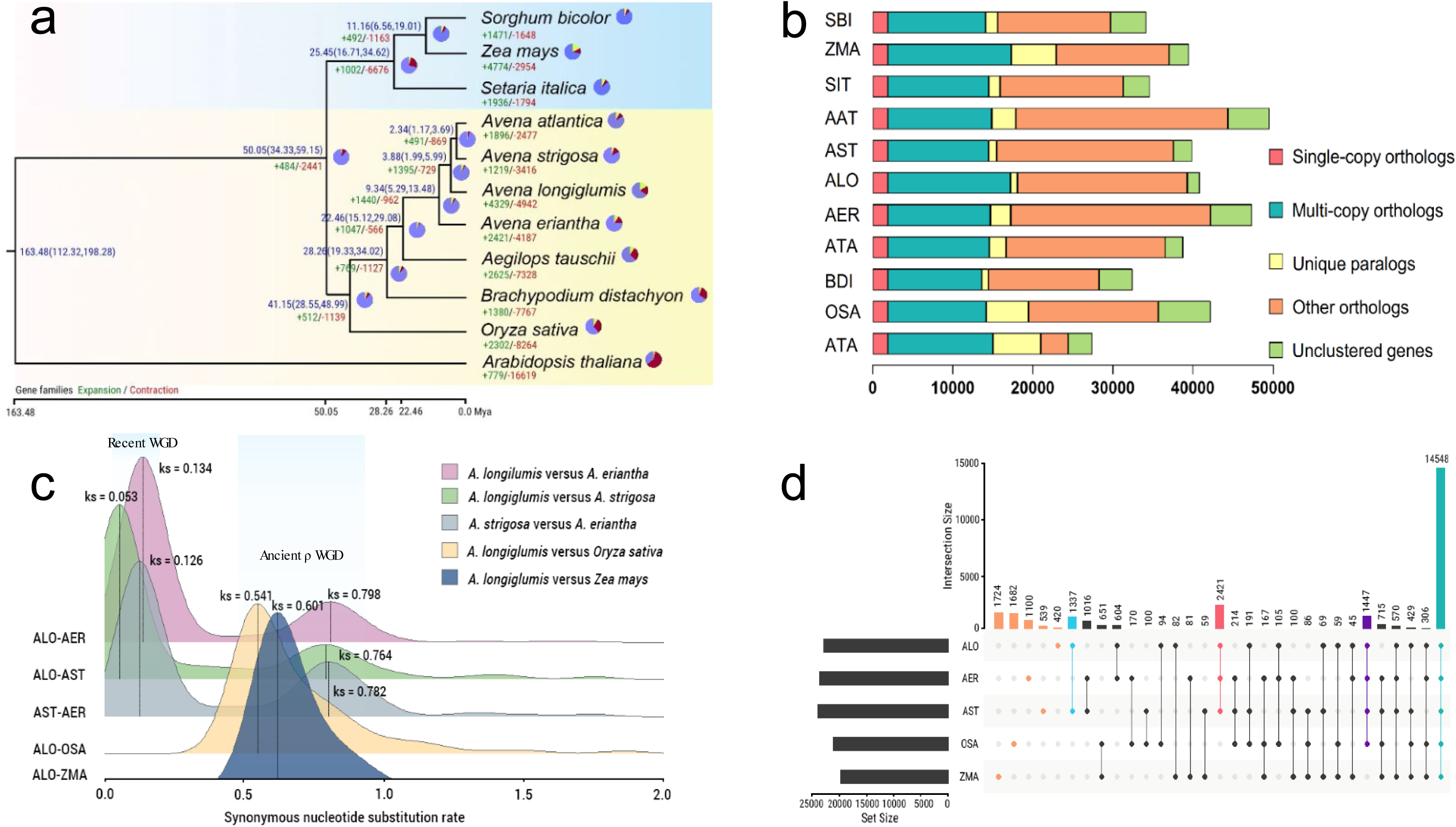
Evolution of the *Avena longiglumis* (ALO) genome. **a** Phylogenetic relationship of ALO with ten plant species. C_3_ species are shown with yellow background and C_4_ species with blue background. Divergence times are labelled in blue; gene family expansion and contraction are enumerated below the species names in green and red. **b** Gene categories are shown for all the species in Fig. 4a. **c** Distribution of ks distance between syntenic orthologous genes for ALO, AER, AST, OSA and ZMA genomes. **d** UpSetR diagram of shared orthologous gene families in five species. The number of gene families is listed for each component. *A*. *atlantica* (AAT) [12], *A*. *eriantha* (AER) [12], *A*. *longiglumis* (ALO), *A*. *strigosa* (AST) [7], *Aegilops tauschii* (ATA) [32], *Arabidopsis thaliana* (ATH) [50], *Brachypodium distachyon* (BDI) [33], *Oryza sativa* (OSA) [9], *Sorghum bicolor* (SBI) [10], *Setaria italica* (SIT) [11], *Zea mays* (ZMA) [34].

We identified homologous gene pairs in ALO, AER, AST, OSA and ZMA genomes and estimated species divergence times by analysis of synonymous nucleotide substitution rates (Fig. 3c). The results indicated that some gene pairs within *Avena* showed a peak at 0.05–0.13, probably reflecting the whole genome duplication (WGD) event in *Avena*, with an additional shallow peak at 0.8 reflecting the ancient rho (ρ) WGD event [52] (Fig. 3c). ALO comparison with OSA (0.54) and ZMA (0.61) showed a single Ks peak. Based on sequence homology among the analysed species, we assigned gene number ranging from 27,416 of ATH to 49,542 of ATA and gene number in families ranging from 24,421 of ATH to 44,363 of AAT, respectively (Additional file 2: Tables S12 and S13). A total of 2,277 gene families (845 in AAT, 355 in AST, 259 in ALO, and 818 in AER) were unique to *Avena* species (Additional file 2: Table S13). Five species (ALO, AER, AST, OSA and ZMA) were selected to identify unique and shared gene families in the Poaceae. We found a total of 14,548 shared orthologous gene families, notably with more than a third (5,423 = 2,421 + 1,337 + 1,061 + 604; Fig. 3d) gene families unique in all *Avena* species, emphasizing the novel gene pool in *Avena*.

#### Comparative genomics of gene families

Based on sequence homology, from the OrthoFinder [51] analysis, the 35,039 genes in families identified across ALO and 10 plant species genomes, we identified gene families showing expansion (1,440) and contraction (962) after the *Avena* divergence from ATA (Fig. 3a, Additional file 2: Table S14a). GO enrichment analysis revealed that the expanded genes of ALO were notably enriched (*p* < 0.05) in molecular functions associated with terpenoid biosynthesis and polysaccharide binding, cellular components such as DNA packaging and signal recognition particle, as well as biological processes associated with hormone, water, biotic and abiotic stimulus (Additional file 2: Table S15). KEGG analysis showed that the expanded genes of ALO were involved in biosynthesis of other secondary metabolites (09110), metabolism of other amino acids (09106), metabolism of terpenoids and polyketides (09109), environmental adaptation (09159), signal transduction (09132), lipid metabolism (09103) and membrane transport (09131) at hierarchy B level, while the expanded genes of AAT were involved in translation (09122), energy metabolism (09102), and environmental adaptation (09159), the expanded genes of AST were involved in replication and repair (09124), energy metabolism (09102) and folding, sorting and degradation (09123), while the expanded genes of AER were involved in energy metabolism (09102) and translation (09122) (Additional file 1: Fig. S8a, b, c and d, Additional file 2: Table S16).

We also examined the unique gene families in *Avena* species: 845 gene families were unique in AAT, 355 in ALO, 259 in AST and 818 in AER (consistent with 1,100 genes unique to AER, 420 in ALO to 539 in AST; Fig. 3d, Additional file 2: Table S13). At hierarchy B level, genes associated with energy metabolism (09102), carbohydrate metabolism (09101) and membrane transport (09131) were uniquely enriched in ALO, and carbohydrate metabolism (09101), membrane transport (09131) and amino acid metabolism (09105) were uniquely enriched in AAT, while transcription (09121) was uniquely enriched in AST (Additional file 1: Fig. S9a, b, c). Compared with the A-genome species, genes associated with folding, sorting and degradation (09123), glycan biosynthesis and metabolism (09107), transport and catabolism (09141), energy metabolism (09102) and biosynthesis of other secondary metabolites (09110) were uniquely in AER (Additional file 1: Fig. S9d). The expansion of gene families occurs during a long-term evolution and drives the evolutionary difference between wild oat species.

### Ancestral linkage group evolution

#### Intraspecific syntenic blocks in Avena longiglumis

To determine the chromosome structure in ALO, we performed an intragenomic synteny analysis. About 15 major syntenic blocks exist between pairs of ALO chromosomes based on paralogous genes (so gene-poor regions around centromeres are not represented). Examples of shared major blocks of paralogous genes between ALO01 (*A*. *longiglumis* chromosome 1) and ALO02; ALO03 and ALO05; ALO04 and ALO06 (Fig. 1, centre m, Additional file 1: Fig. S10a, d). The pairs of syntenic blocks are likely to identify the signature of the ancient ρ WGD event in the grasses. The block covered most of the gene-rich parts of the genome with no indication of major deletions following WGD. There were no regions with three or more copies, suggesting no major segmental duplications. In contrast to *Musa* (D’Hont et al. [53] their supplementary Fig. S12) and *Ensete* [26] that do not share the grass ρ WGD event, there is clear evidence for two rounds of WGD in *Avena*.

#### Avena Intergeneric chromosome rearrangements

To investigate the relationship between ALO, AST (both A-genome, Additional file 1: Fig. S10b) and AER (C-genome, Additional file 1: Fig. S10c), we conducted a synteny analysis between *Avena* genomes, all sharing the same WGD events. We found a total of 29,030, 27,116 and 21,536 pairs of collinear genes between ALO-AST, ALO-AER and AER-AST species pairs, respectively (Fig. 4a, Additional file 1: Fig. S11a, b, c, Additional file 2: Table S17). Visualization of regions of synteny between ALO, AST and AER, with SynVisio [42] shows large blocks of conservation between ALO and AST, with much more rearrangement with the more distant AER (Fig. 4a). Between ALO and AER, chromosome ALO01 was largely collinear with AER03, and ALO06 with AER02 (Fig. 4a; Additional file: Fig. S11a, b). Other AER chromosomes had multiple syntenic regions, each involving most ALO chromosomes. Whether the higher level of structural variation reflects syntenic gene clusters and the adaptation to a sandy-soil and arid environment of AER needs further investigation. Most notably, numerous evolutionary inter-chromosomal translocations represented 10% to 25% of the length of nearly all chromosomes with many in large distal domains (Fig. 4a, Additional file 1: Fig. S11a, b, c). Interestingly, these distal (sub-terminal) intragenomic evolutionary rearrangements, identified here for the first time in diploid species, are entirely consistent with distal nature and size of translocations identified using genome-specific repetitive DNA sequence probes in polyploids [22] where translocations between genomes have occurred since the polyploidy event.

**Fig. 4.**
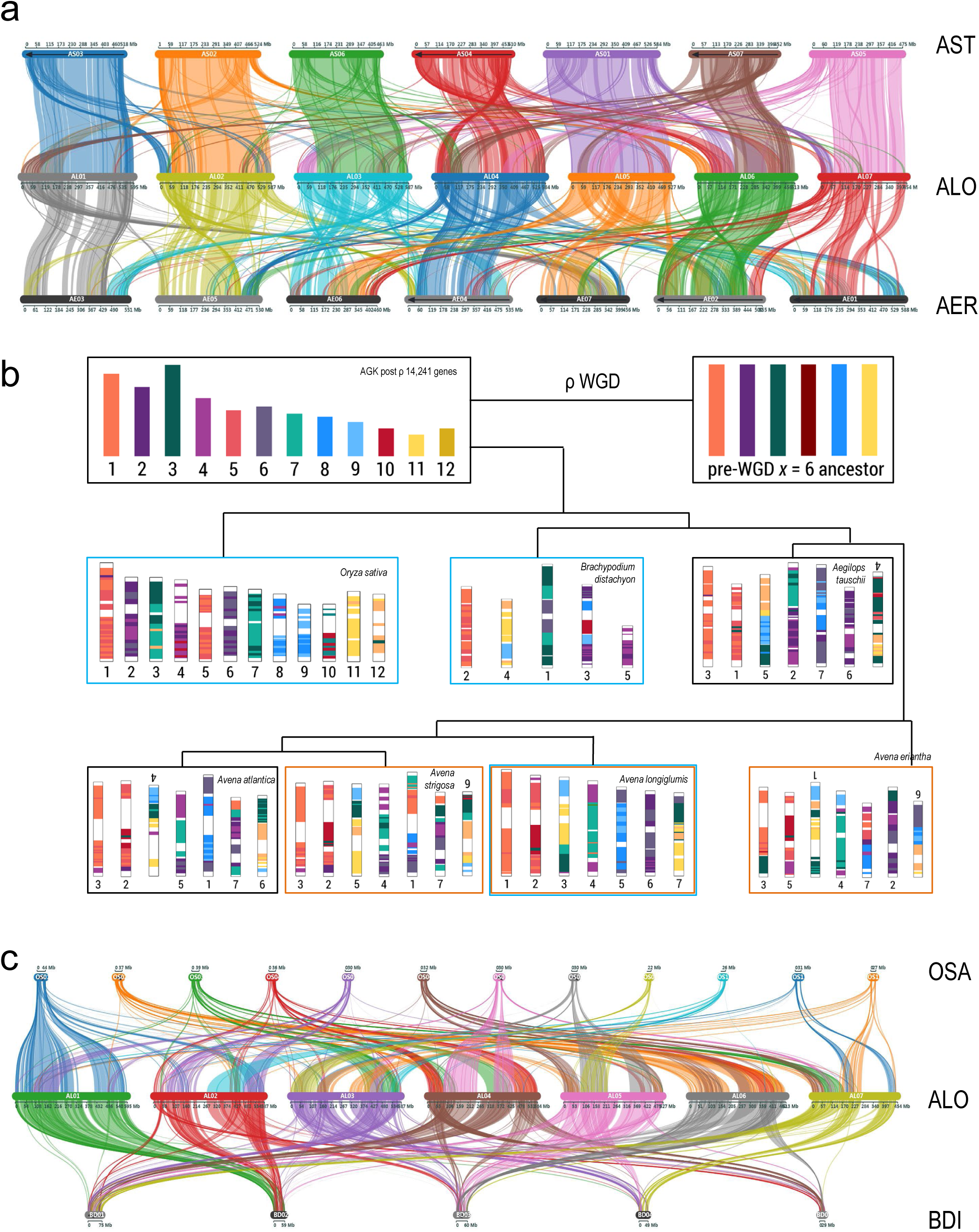
Syntenic relationships of chromosomes of the ancestral grass karyotype (AGK) and analysed species. **a** Syntenic analysis of *Avena strigosa* (AST), *A*. *longiglumis* (ALO) and *A*. *eriantha* (AER). Subterminal regions are frequently involved in interspecific evolutionary translocations. **b** Reconstruction of ancestral chromosomes for the seven species showing conservation of major syntenic blocks from the ancestral grass karyotype (AGK) with fusions and insertions leading to the reduced chromosome numbers (chromosomes numbered by published linkage groups, some are upside down to display features of evolutionary conservation). **c** Deep syntenic analysis of *Oryza sativa* (OSA), ALO and *Brachypodium distachyon* (BDI) showing detailed conservation of syntenic block and the expansions between OSA (*x* = 12,389 Mbp), BDI (*x* = 5,260 Mbp) and ALO (*x* = 7, 3,960 Mbp). Genes from the ancestral linkage groups are indicated by colours, with pairs of similar colours representing the pre-rho whole genome duplication.

**Fig. 5.**
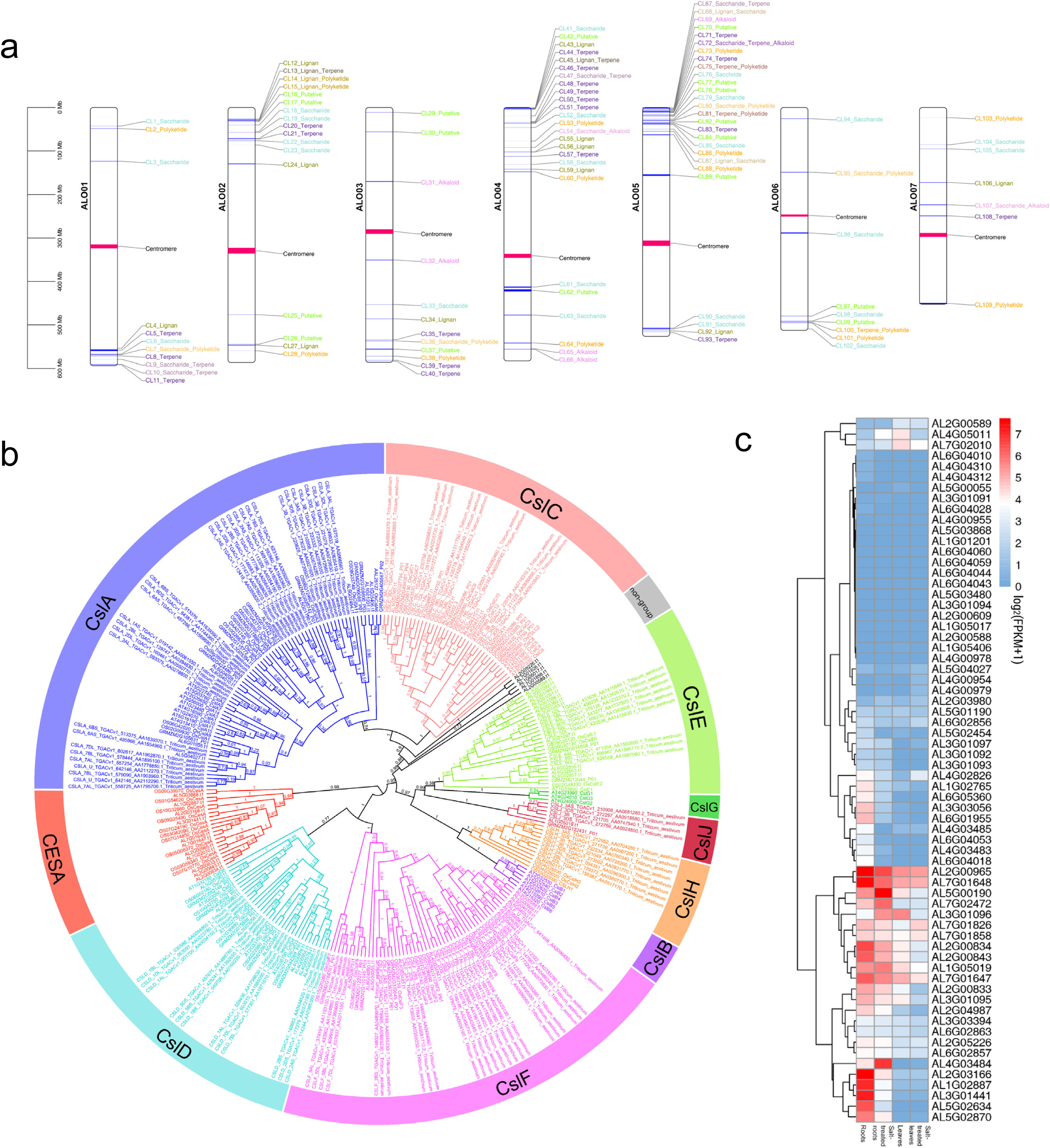
Identification of biosynthetic gene clusters (BGCs) and *cellulose synthase A* (*CesA*) and *cellulose-like* (*Csl*) gene families in ALO. **a** Total 109 BGCs identified in ALO chromosomes by plantiSMASH. The cluster types, including alkaloid, lignin, polyketide, saccharide, terpene, lignin_polyketide, lignin_saccharide, lignin_terpene, saccharide_alkaloid, saccharide_polyketide, saccharide_terpene, saccharide_terpene_alkaloid, and terpene_polyketide biosynthesis genes, labeled as different colour. The cluster position shown by blue band. The centromere position shown by pink band. A scale in the left represented length of chromosome in megabases (Mbp). **b** Maximum likelihood phylogenetic tree of CesA and CSL proteins from ALO, rice, wheat and arabidopsis. Nongroup: ALO CSL proteins are not clustered with any known CesA and Csl proteins. **c** Heat map showing hierarchical clustering of *CesA* and *Csl* gene families in roots, salt-treated roots, leaves and salt-treated leaves of ALO. Expression values were normalized by log2(FPKM + 1). Highly and weakly expressed genes were colored by red and blue boxes, respectively.

In order to understand the structural chromosomal variation including duplications and deletions, we examined the extent and nature of chromosomal rearrangements across Pooideae species. Two species with small genome sizes, OSA (*x* = 12) and BDI (*x* = 5) have been used extensively as reference genomes. Previous studies have suggested that Poaceae genomes evolved from a pre-ρ WGD ancestral grass karyotype (AGK), with 7 protochromosomes, to a post-ρ AGK with 12 protochromosomes [14]. Conserved genes for each AGK chromosome have been identified (with the proposed 12 protochromosomes showing extensive similarities with rice). Here, the AGK genes were mapped to the chromosomes of ALO and six grass species (AAT, AST, AER, ATA, BDI and OSA), and the corresponding regions of chromosomes were designed by a colour (Fig. 4b). The signature of the ancient ρ duplication (see also in Fig. 1, centre m) is shown by pairs of chromosomes with shades of similar colours. Because of divergence in gene sequences, there is some ambiguity in assignment of extant chromosome blocks to the duplicated chromosomes in the ancestor (eg. the shades of orange from AGK01 and AGK05 in ATA03) and so cannot be interpreted as clearly. Dotplots of ALO against OSA and BDI (Additional file 1: Fig. S11d, e and f) were also made as the reference for a SynVisio [42] plot of ALO with OSA and BDI (Fig. 4c).

Large segments of the ancestral chromosomes are conserved across the analysed grasses with distinct rearrangements involving translocations and fusions of syntenic blocks between the species. Some rearrangement events are shared between all *x* = 5 and *x* = 7 species (e.g., the fusion of AGK09 and AGK11; or AGK02 and AGK03; both are seen in BDI, ATA and the *Avena* species) or between the *x* = 7 species (AGK12 and AGK06 giving ALO07; Fig. 4b, Additional file 2: Table S17).

While some evolutionary events from the ancestral 12 AGK chromosomes involve fusion and rearrangement of syntenic blocks, it is notable that three events are largely characterized by insertion of one chromosome (group of syntenic genes) into another chromosome. Thus, ALO04 has AGK07 inserted into AGK04; ALO06 has much of AGK06 inserted into AGK02; while ALO05 has insertion of AGK08 into ALO06. The inheritance of fusion events is consistent with the phylogeny (Fig. 3a) and time since separation from the most recent common ancestor with the proposed ancestral grass karyotype. Given the higher number of rearrangements from the AGK, Fig. 4b suggests AER is the most derived karyotype in *Avena* and the A-genome (including ALO) is more primitive. Overall, during evolution, chromosome rearrangement has been restricted to a small number of events, presumably avoiding alteration of interactions of gene groups and promoters along chromosomes without fusion and fissions.

### Genome-wide expression analysis

As well as use for gene annotation, we analysed the relative transcription of genes in roots, salt-treated roots, leaves and salt-treated leaves of ALO to suggest key features of gene expression level variation. Differentially expressed genes (DEGs; fold change ≥ 2 and FDR ≤ 0.005), comprised 17.48% of genes (3,076 up-regulated DEGs and 3,963 down-regulated DEGs; Additional file 2: Table S18a). KEGG analysis indicated that the 1,569 up-regulated genes in salt-treated roots enriched in environmental adaptation (09159), metabolism of terpenoids and polyketides (09109), biosynthesis of other secondary metabolites (09110), signal transduction (09132) and membrane transport (09131), while the 1,627 down-regulated DEGs were enriched in salt-treated roots in biosynthesis of other secondary metabolites (09110), replication and repair (09124), metabolism of other amino acids (09106) and carbohydrate metabolism pathways (09101) (Additional file 1: Fig. S12a and b). In salt-treated leaves, there were 1,507 up-regulated DEGs enriched in terpenoid backbone biosynthesis (00900) and transcription (09121) pathway, while 2,336 down-regulated DEGs enriched in energy metabolism (09102), carbohydrate metabolism (09101), amino acid metabolism (09105), metabolism of terpenoids and polyketides (09109), membrane transport (09131), signal transduction (09132), metabolism of cofactors and vitamins (09108) and environmental adaptation (09159) pathways (Additional file 1: Fig. S12c and d).

KEGG enrichment pathways of up- and down-regulated DEGs were very similar in ALO expanded gene families, and most genes involved with salt adaptation mainly belonged to the expanded gene families. The environmental adaptation, metabolism terpenoids and polyketides, membrane transport pathways were enriched in up-regulated DEGs of salt-treated roots, while these pathways were enriched in down-regulated DEGs of salt-treated leaves, which may be related to different responses to salt stress between aboveground and underground parts of plants. Additionally, the pathways related to terpenoid synthesis were enriched in both roots and leaves, in which they may be extensively participated salt-tolerance of plants [54].

Further investigate the resilience effect of DEGs on the environmental adversity of ALO, we conducted transcriptional analyses of 4,329 expanded gene families (7,236 expanded genes, Additional file 2: Table S14). Expanded DEGs comprised 34% of total DEGs (1,352 expanded DEGs in salt-treated roots and 1,093 expanded DEGs in salt-treated leaves, Additional file 2: Table S18b, c). The number of DEGs in roots was slightly higher than in leaves.

We analysed the gene function of DEGs among expanded gene families in roots and leaves of ALO. The 599 up-regulated, expanded genes in salt-treated roots included protein kinase, cytochrome P450 (CYP450), cupin, and pathogenesis-related protein genes, and 753 down-regulated expanded genes in salt-treated roots included nucleosome histone, protein kinase, transferase and other *CYP450* families (Additional file 1: Fig. S13a). Expanded DEGs showed relatively fewer in salt-treated leaves, the 451 up-regulated genes including zinc finger protein, protein kinase, peptidase and other genes, and the 642 down-regulated DEGs in salt-treated leaves including protein kinase, CYP450 and receptor kinase (Additional file 1: Fig. S13b). For *CYP450* genes in salt-treated samples, 27 up- and 22 down-regulated DEGs were detected in salt-treated roots, and 17 down-regulated genes were detected in salt-treated leaves of ALO.

### Analysis of cytochrome P450 (*CYP450*) gene clusters

In total we identified 109 biosynthetic gene clusters (BGCs) in the ALO genome, including alkaloid, lignin, polyketide, saccharide and terpene biosynthesis genes (Fig. 5a). The result (Additional file 2: Table S19) allows assessment of each locus for its likelihood to encode genes working together in one pathway [55]. To evaluate the potential of ALO for the genetic dissection of agriculturally important traits, we focus on the evolution of triterpene synthesis, including clusters of the *CYP450* gene families, which encode proteins involved in multiple metabolic pathways with complex functions and playing important roles in defence responses to abiotic stresses. The number of *CYP450* genes in ALO (557; 1.36% of total genes) was significantly higher than in nine analysed grasses (251–454) and ATH (246; Additional file 2: Table S20a). Overall, the *CYP450* genes were relatively equally distributed on all chromosomes (average 80, between 73 and 100 observed) (Additional file 2: Table S20b). The transcript level of one *CYP450* gene (AL2G04509) was over 913-fold in salt-treated roots than roots; and AL7G05074 was 409-fold in salt-treated leaves than leaves (Additional file 1: Fig. S14, Additional file 2: Table S21).

Using the presence of at least one orthogene in the identified gene clusters as the selection criteria, we assigned 46 putative *CYP450* gene clusters (Additional file 1: Fig. S15, Additional file 2: Table S22). Five key gene clusters identified in the ALO genome were CL10 and CL95, CL37, CL98 and CL106, which included functionally characterized UDP-glycosyl transferase (AS01G000200), serine carboxy peptidase-like acyltransferase (AS01G000190), subtilisin homologue (AS01G000130), O-methyltransferase (AS01G000040) together with enzymes annotated as CYP450 (Li et al. [7] their supplementary table 7), dehydrodolichyl diphosphate synthase, aldehyde oxidase and hydrolase proteins (Additional file 1: Fig. S16a, b, c, d, e, f and g).

We examined the gene number and conserved synteny around *CYP450* gene clusters in the four *Avena* species (sharing 24 *CYP450* genes) and 10 grass species (sharing six *CYP450* genes; Additional file 2: Table S23a, b). In ten clusters (Additional file 2: Table S22), *CYP450* genes present across four *Avena* species with loss of copies and without syntenic relationship among six grass species. We identified a further 11 *CYP450* gene clusters containing terpene synthase genes (Additional file 2: Table S22) with *CYP450* genes together with terpene synthase genes, showing conservation across all species including *Avena* suggesting these gene clusters related to functional expansion of specialized terpene metabolism [55]. Tandem duplications within 18 *CYP450* gene clusters (Additional file 1: Figs. S16a, b, c, d, e, f and g) were revealed in ALO. These gene clusters, with known functionally characterized genes involved in CYP450 biosynthesis and extensive copy number variation (CNV) between species, can be taken to present the pangenome for CYP450 biosynthesis [56] (Additional file 1: Fig. S16a and b).

Overall, the analysis provides strong support for the non-random organization of *CYP450* biosynthetic genes and presence of *CYP450* gene clusters [7, 57] in *Avena* and other grasses. Indeed, the high-continuity ALO genome assembly shows the avenacin cluster (including antimicrobial terpene biosynthetic genes), in terms of both gene number and the diversity of gene families that it contains, was even stronger than the triterpene BGCs identified previously [7].

### Phylogenetic analysis of the *CesA* and *Csl* gene families

#### Identification of cellulose synthase (*CesA*) and cellulose synthase-like (*Csl*) gene families in ALO

To gain insight into whether the physical location plays a role in expansion of β-glucan biosynthesis genes, we evaluate the genomic organization of *cellulose synthase A* (*CesA*) and *cellulose synthase-like* (*Csl*) gene families (Additional file 1: Figs. S17 and S18). They encode 1,4-β-glucan synthase superfamily serving as the predominant structural polymer in primary and secondary cell walls of caryopses [58]. Dataset searches using conserved Pfam motifs PF000535 and PF03552 [44], which are specific to the glycosyltransferase GT2 superfamily [59], resulted in the identification of 11 *CesA* and 55 *Csl* genes (Additional file 2: Table S24). The maximum likelihood (ML) tree (Fig. 5b) for CesA (a single branch with 11 proteins in ALO) and Csl proteins from ALO, *Arabidopsis thaliana*, rice, wheat and maize (Additional file 1: Fig. S19) shows the ALO Csl proteins group in seven subfamilies: CslA (10 proteins), CslC (6 proteins), CslD (8 proteins), CslE (11 proteins), CslF (13 proteins) and CslH (1 protein) and CslJ (1 protein), with five proteins unclassified). The closely related CslA and CslC subfamilies were conserved across the species, as were the sister sub-families of CslD and the grass-specific CslF.

*CesA* and *Csl* genes were relatively equally distributed over all chromosomes (average 9, between 6 and 13 observed) (Additional file 2: Table S25). There were large differences in expression of *CesA* and *Csl* genes between roots, salt-treated roots, leaves and salt-treated leaves (Fig. 5c) with the fragments per kilobase of exon model per million mapped reads (FPKM) showed ratios up to 150 that are suggestive of their functional role (Additional file 2: Table S26). Some genes presenting higher levels of expression in roots than leaves, may be involved in β-glucan synthesis [60]. Among 109 gene clusters, we identified 10 metabolic gene clusters representing 2 *CesA* and 10 *Csl* gene models across six of seven chromosomes (Additional file 1: Fig. S17, Additional file 2: Table S27). Synteny analysis showed the conservion among *CesA* and *Csl* genes of gene clusters: four *Avena* species shared 1 *CesA* and 3 *Csl* genes, and 10 grass species shared two *Csl* genes (Additional file 2: Table S28a, b). In four clusters of *Csl* genes present across four *Avena* species, the loss of copies, and lack of syntenic relationships to six other grass species, suggests the opportunity to exploit the (1,4)-β-glucan biosynthesis pathway outside oats may be limited (Additional file 2: Table S24). The two gene clusters (CL32 and CL58) contained alkaloid and saccharide biosynthetic genes showed the conserved synteny relationship with other grasses. Tandem duplications within *CesA* and *Csl* gene clusterwere observed in ALO (Additional file 1: Fig. S18a). It is thus important to study the role of individual *CesA* and *Csl* in primary and secondary cell wall biosynthesis to attempt effective modification of biomass composition.

Comparing the cellulose-like synthesis clusters with homologous genomic loci in AAT genome can give important information on its evolutionary conservation or diversification (Additional file 2: Tables S28, S29 and S30). Whereas strong conservation of clusteredness across larger periods of evolutionary time may point to a selective advantage of clusteredness for these genes, diversification of *Csl* genes by co-option of glyoxalase genes may give clues to find novel variants of natural products that have been generated through directional pathway evolution (Additional file 1: Fig. S18a, b, c, d, e and f). A better understanding of the cell wall gene expression under abiotic stress is important to design strategies to produce crops in marginal lands with less β-glucan accumulation. Gene families play an important role in enhancing salt-tolerance and adaptation of ALO, which was also found in desert plants [61].

#### Identification of callose synthase (CalS) enzyme families in ALO

Callose (1,3-β-glucan), encoded by the *callose synthase* (*CalS*) or *Glucan synthase-like* (*GSL*) gene families, plays an important role in plants grown both normal and unfavourable environments [62]. Dataset searches for ALO using conserved Pfam motifs PF02364 and PF14288, identified 13 CalS or GSL proteins. With the *CalS* genes, distributed along five chromosomes (ALO01/03/05/06/07, with no notable chromosomal clusters), and five *CalS* genes were identified in the expanded gene families of ALO (Additional file 2: Table S29). The transcript level of *CalS* gene (AL6G02277) was 11-fold higher in salt-treated roots than roots, while AL3G02326 was one-fold in salt-treated leaves than leaves (Additional file 1: Fig. S20, Additional file 2: Table S30) supporting involvement in plants’ resilience to salt stress.

#### Comparative phylogenetic analysis of the CalS gene family

The 13 CalS or GSL proteins were placed in a ML tree (Additional file 1: Fig. S21) along with 23 GSL proteins from *Arabidopsis thaliana* and rice (Additional file 1: Fig. S22). The analysis grouped the proteins into eight clades, with Clades VII and VIII, and Clade II (*Arabidopsis*) and III (rice and *Avena*), being sisters. Five clades (IV to VIII) included all plant species suggesting diversification present in the common ancestors, with further duplications reflecting ancient whole genome duplication events (α, β, γ in eudicots, and τ, σ, ρ in monocots) or more recent segmental duplications.

## Discussion

The diploid wild oat, *Avena longiglumis* (ALO), with distribution around the Mediterranean Basin, is an important genetic resource for oat breeding, and a valuable reference for genomic organization and evolution in the grasses. We used a combination of Nanopore (436 Gbp), Illumina (269 Gbp) and Hi-C (331 Gbp) sequencing technologies to assemble the 3,847,578,604 bp long ALO genome into the seven pseudo-chromosomes (Fig. 1, Tables 1 and 2). The high-quality ALO genome assembly gives not only insight into the gene diversity but also into the variation in repetitive DNA content and structural variation (SV) in the genome including chromosome duplication and arrangements.

Assembled plant genome sizes range from 61 Mbp (*Genlisea tuberosa*, bladderwort [63]) to 26,454 Gbp (*Sequoia sempervirens*, coast redwood [64]). The ALO assembly falls into this range, and a combination of ONT, Illumina and Hi-C sequence approaches was essential for the high continuity chromosome level assemblies with high presence of core genes, as is for many important crop genomes, particularly cereals, with genomes larger than 2 Gbp (barley [65]; wheat [66, 67]). *Avena*, as all large genomes, includes abundant copies of TEs [22] and the long-read sequencing technology allowed examination of their organization (Figs 1 and 2). Most transposable element classes are distributed widely and rather uniformly along the ALO chromosomes (Fig. 1). In many species with smaller genomes (<3.96 Gbp of ALO), broad pericentromeric regions are reservoirs for the accumulation of a medley of (often lineage-specific) TEs [26, 68, 69], but in ALO, the overall TE density is relatively similar along the chromosomes. However, there is a high and localized abundance of an LTR retrotransposon (Fig. 1 circle c) at the centromeres of all seven ALO chromosomes. Such contrasting distributions of LTR retrotransposon clades has been found in several species [40, 70, 71, 73]. Centromeres of some species harbor arrays of tandemly repeated satellite sequences (e.g., *Arabidopsis thaliana* [74, 75], *Beta vulgaris* [76] and *Ensete glaucum* [26]), but we found no equivalent tandem repeat in ALO. However, often in genomes with centromeric satellite sequences, abundant families of retroelements are also found at the centromeres, such as the *Nanica* LINE of *Musa acuminata* [77] and *E. glaucum* [26], *Arabidopsis* retroelement domains [67] or the wheat *Quinta* and other elements [32, 78].

In a short evolutionary timescale, young TEs (< 2 Mya [79]) were frequent (Fig. 2) in ALO. LTR retrotransposons may be beneficial to their hosts by providing regulatory genetic elements [80, 81] or by disruption of genes and their promoters. While TEs are unlikely to be the only causal factor responsible for subgenome expression dominance in polyploids, methylated TEs can reduce the expression of nearby genes [82]. Further studies are needed to address whether oat A-genome dominance is determined by methylation pattern differences of retrotransposons [83], and their contribution to genetic variation in different *Avena* species.

The ρ WGD events occurred 50–70 Myr ago, after Poales separated from other monocot orders [4, 84, 85]. Based on detailed paleogenomics, using inference from *x* = 5–12 grasses in terms of gene order and content, Murat et al. [14] proposed an ancestral grass karyotype (AGK) with similarity to the extant *Oryza sativa* (2*n* = 2*x* = 24) genome including 14,241 conserved genes (Fig. 3). We delineated genome sequences between OSA, BDI, ATA and four *Avena* species, representing four tribes and different polyploidization events, confirming that the ρ Poaceae event is shared by the ancestral BOP clade and Poales (Fig. 4). Most notably, the conservation of large syntenic blocks and the orthologous relationships of the seven extant ALO chromosomes to the 12 chromosomes of OSA and AGK was evident, with defined fusion and translocation events but limited major duplications or deletions. Apart from the long-term evolutionary conservation, such regions harbour conserved sequence regions that might be synthesized as oligonucleotides for *in situ* hybridization to label linkage group 1 across all Poales grasses [86], and to use as baits (cf. https://treeoflife.kew.org/methods and Johnson et al. [87]) to identify the variation of all AGK01 genes across the group. Other chromosomes have well-defined range of fusions from the AGK or OSA reference, reducing the chromosome number from *x* = 12 to *x* = 5 or *x* = 7, but notably some chromosomes have evolutionary insertion of one ancestral chromosome into another.

The conservation of many syntenic blocks and the chromosome structure occurs despite of the huge expansion in genome size, with ALO being ten times larger than OSA (15.2 × BDI). Most notably, there is expansion in genome size throughout the chromosomes, largely involving the amplification of retroelements that are dispersed uniformly along all chromosome arms (Fig. 1) and is evidenced by the lines of synteny between ALO and the corresponding BDI and OSA syntenic blocks spreading out relatively uniformly over a much-expanded region of ALO. There are a few gene rich regions (on chromosomes ALO01, ALO04, ALO05 and ALO07) shared with BDI but not OSA that are worth further investigation. Overall, the genome structure revealed in the syntenic comparison reveals the evolutionary history of the Poales at the chromosome level, and encourages exploitation of the whole gene pool in both biodiversity studies and for plant breeding.

*In situ* hybridization using repetitive DNA probes has shown that many chromosomes in the hexaploid *A. sativa* show intergenomic translocations (i.e., between chromosomes of the diploid ancestral genomes [16, 88]), involving the terminal 10% to 25% of many chromosome arms. Such translocations have not been seen in the tribe Triticeae (syn. Hordeae, sister tribe to Aveneae). Remarkably, the three *Avena* species (ALO, AST and AER) have multiple terminal translocations between seven chromosomes of three species (Fig. 4), occurring only on one arm. Six chromosomes are involved in clear non-reciprocal translocations between ALO and AST, but no terminal regions have been lost during the translocation events. With respect to the evolutionarily more distant AER chromosomes, every ALO chromosome has a terminal translocation as well as a greater number of other rearrangements. The terminal rearrangements do not only involve repetitive DNA sequences, as is likely to be the case in maize (e.g., The P53 knob [89]) or rye (pSc250 tandem repeat sequence [73]), but also involve many genes within syntenic groups [90].

Poales species occupy differentiated environmental and ecological niches, with contrasting selective pressures so we looked at groups of syntenically conserved genes where phenotype and selection may be affected. ALO is restricted to sandy loam soils and mesic habitats in the Mediterranean desert, while AER populations thrive on shallow calcareous hills or terra rossa soil steppes around the Mediterranean Basin [91]. Notably, while there are few insertions or deletions between ALO and AST, there are many gaps, but not break in synteny along chromosome arms between ALO and AER (Fig. 4). It will be interesting to see if these regions are related to functional or selective changes in copy number and the unique paralogues of AER (Fig. 3).

The physical clustering of multiple genes from a single metabolic pathway is now established in plants [7, 92]. Clustering should favour co-inheritance of beneficial combination of alleles that confer a selective advantage together [93]. Our results show clustering of genes and regulators including terpene (Fig. 5), cellulose or phytohormone pathway enzymes. In terpene and cellulose clusters, *CYP450s* exhibit down-regulation among different tissues, while most are considered as highly tissue-specific genes. The common expression trends of homologous genes also exist in wheat and maize, implying a unique highly conserved function for each clustered gene [93]. Consistent with the model, our survey-expression data indicates some *CYP450*s are up-regulated, and others are down-regulated under salt stress, suggesting the need for detailed investigation of *CYP450* functions under salt stress [93].

The *CesA*/*Csl* gene families play a critical role in the biosynthesis of cellulose and hemicellulose. We identified 66 *CesA*/*Csl* genes which could be divided into four lineages in ALO. Orthologous genes (in different species) can be more similar than paralogous genes (of the same species), eg, the *Arabidopsis* (dicot)-specific CslB lineage was closer to grass-specific CslH lineage than CslF lineage, suggesting that CesAs and Csls diverged before the split of monocots and eudicots, c. 150 Mya [94]. This indicates that the *CesA*/*Csl* genes established their roles early in higher plant evolution, and could be a reason why there are so many *CesA*/*Csl* gene families in Poaceae. Our findings indicate that the larger *CesA*/*Csl* superfamily is the consequence of recent duplications (Fig. 3), and particularly chromosomes ALO02 and ALO06 have more *Csl* genes than other grass species.

Some expanded *CesA*/*Csl* genes may be retained simply owning to sub-functionalization where the functions of the ancestral genes were partitioned among the duplication. In the case of *CslD1* subfamily members, which are involved in root hair-deficient phenotypes of maize [95], some *CslD1* copies can be lost without any phenotypic consequences. Intragenic complementation has not been observed among alleles with mutations in different *CSLD1* domains of *Lotus japonicas* [96], suggesting that the *CSLD1* has not yet undergone the complete functional differentiation. *CslF6* and *CslH* have a functional role in the synthesis of mixed-linkage (1,3;1,4)-β-glucan (MLG [97]), and the sequence divergence of the *Csl* genes we found is likely a reflection of their functional divergence although MLG synthesis is tightly regulated and thus maximizing the yield of end-product cellulose might be difficult. For example, isoforms may utilize the same donor but a different acceptor molecule in the synthesis of the same polysaccharide, and thus, having multiple genes may be a requirement for synthesis of some types of plant polysaccharides [58].

## Conclusions

The 3.85 gigabase sequence assembly of the wild oat species *Avena longiglumis* has enabled chromosome evolution to be defined within *Avena* and diverse Poaceae species. The diversity revealed in gene and gene network will accelerate the analysis of trait genes and their control. Beyond diversity in genes and regulatory sequences, the spectrum of chromosomal structure variation and sequence copy number variation (both of genes and repetitive DNAs), can be shown by comparison with our high-continuity genome assembly, and will enable characterization of the *Avena* and broader grass pangenome. There is increasing recognition of the role of structural and copy number variation in diversity, going beyond the well-studied differences in gene alleles, networks, and transcription factors, both in plants and animals [19, 20].

Between rice (*Oryza sativa*) and *A. longiglumis* (a genome 10.18 times larger than rice), the amplification of non-coding sequences lying between genes has occurred throughout the chromosome arms: syntenic regions show relatively uniformly expansion with a few substantial gaps. The repetitive sequence component of the ALO genome—the repeatome––is characterized by both ancient and more recently amplified transposable elements (TEs) and tandem repeats occurring both along chromosome arms and at centromeres. It remains unclearly why genome size should be so different in two successful crop genera, *Avena* and *Oryza*, and whether selective pressures (dynamics of repeat replication and transposition) enhance options for evolvability. Given the high synteny observed, with presence of well-defined inter-chromosomal translocations and fusions between the species (including insertion of ancestral syntenic blocks within another), conserved nucleotide sequences and domains can be identified by major linkage blocks in grasses. We suggest that these can form the basis for synthetic pan chromosome oligonucleotide pools for *in situ* hybridization to identify major chromosomal and karyotypic rearrangements across the Poales.

The *Avena* genome assembly and analysis here, along with those of Triticeae and *Oryza* species in the BOP (Bambusoideae-Oryzoideae-Pooideae)-clade, provide insight into the extent and nature of chromosomal rearrangements and genome expansion in the pangenome, contributing to exploitation of the diversity present in the common gene pool across grasses through precision breeding using a range of approaches.

## Materials and methods

### Plant germplasm, genome sequencing and assembly

#### Plant material

The *Avena longiglumis* (ALO) (PI 657387; US Department of Agriculture at Beltsville, https://www.ars-grin.gov/, originally collected in Spain) was used for genome sequencing. After sowing, seedlings were grown in South China Botanical Garden Greenhouse at 25°C, 16 h light/8 h dark with 70% relative humidity. Four weeks later, the plants were moved outside and further grown for 4 weeks under natural day-light condition (dry season in Guangzhou).

#### Genome survey sequencing and assembly

Genomic DNA for Illumina mate-pair sequencing was extracted using the DNeasy Plant Mini Kit (Qiagen) from 8-week-old leaves of ALO seedlings. An amplification-free approach was used to prepare sequencing libraries with insert sizes of 350 bp, following the manufacture’s protocol [98]. The paired-end reads were loaded into two lanes of an Illumina HiSeq2500 platform and raw data generated reads with 2 × 150 bp length (Table 1; Additional file 2: Tables S1 and S2). ALO genome size, heterozygosity and repeat content were determined by *k*-mer (17-mer) analysis by Jellyfish v.2.2.6 [99] with the parameter “-c -m 51 -s 10G -t 50”. The output file was used as the input for GenomeScope [100] to estimate the genome size. Project data have been deposited at Genome Sequence Archive (https://ngdc.cncb.ac.cn/gsa/browse/CRA003996; Additional file 2: Tables S1 and S2a).

#### Oxford Nanopore Technology (ONT) sequencing and assembly

For ONT PromethION library construction and sequencing, genomic DNA was extracted from 3-week-old leaves of ALO seedling using the QIAGEN^®^ Genomic DNA Extraction Kit (Cat. 13323, Qiagen) according to the manufacturer protocol. DNA quantification was carried out using Qubit^®^ 3.0 Fluorometer (Invitrogen, USA). DNA purification was confirmed (OD 260/280, 1.8–2.0; OD 260/230, 2.0–2.2) and fragments in the range of 10–50 kbp recovered using a BluePippin automatic nucleic and recovery instrument (Sage Science, USA). The 3’ and 5’ overhangs were converted into blunt ends with NEBNext^®^ FFPE DNA Repair Mix (NEB, Cat. M6630) and then ‘A’ base was added to 3’ blunt ends using the A-Tailing reaction (NEBNext^®^ Ultra^TM^ II End Repair/dA-Tailing Module, NEB, Cat. E7546). The purified A-tailed DNA was ligated with adaptors from the Ligation Sequencing Kit (SQK-LSK109, Oxford Nanopore Technologies) and the NEBNext^®^ Quick Ligation Module (NEB, Cat. E6056). The purified ligation products were used as the constructed sequencing library. The DNA libraries were accurately quantified using a Qubit^®^ 3.0 Fluorometer (Cat. E33216, Invitrogen, USA) and loaded into 12 lanes of a PromethION, R9.4.1 flow cell (Oxford Nanopore Technologies, UK) for SMRT (single molecular real-time) sequencing. Sequencing results (fast5 files) were processed using the Guppy v.3.2.2 [101] (Additional file 2: Table S2b).

A total of 31.2 million passed reads (Q score ≥ 7; 252.8 Gbp) were generated with read length N50 12,682,464 bp (Additional file 2: Table S3). NextDenovo v.1.0 [102], wtdbg2.huge [103] and SMARTdenovo v.1.0.0 [104] have been used for self-correction of ONT reads. The pass reads were sent into NextDenovo v.1.0 for read correction. We tested parameters and found that using corrected reads to SMARTdenovo v.1.0.0 with the assembler parameters ‘-c 3’ and ‘-k 11’ gave good results, yielding a preliminary assembly consisting of 2,379 contigs (contig N50 11.92 Mbp). Contigs were polished three times with ONT raw data by NextPolish v.1.01 [102] and four times by the filtered Illumina whole-genome shotgun data by Fastp v.0.20.1 [105]. This procedure increased the contig N50 size to 12.68 Mbp (Additional file 2: Table S3).

#### Hi-C library preparation and sequencing

For Hi-C sequencing, 3-week-old leaves of ALO seedlings were fixed in 2% formaldehyde solution. The nuclei/chromatin was extracted from the fixed tissue and digested with DpnII (NEB, Cat. E0543L). Hi-C libraries were constructed and sequenced on the Illumina Novaseq 6000 platform to obtain 150 bp paired-end reads (Additional file 2: Table S3). Raw data were processed by trimming adaptor and removing low-quality reads (Phred quality scores < 15) by Fastp v.0.20.1 [105] with default parameters. A total of 1,453 million clean reads were kept for the mapping process. The quantity of informative Hi-C reads was estimated by Hi-C_Pro v.2.10.0 [106].

The 585 million paired-end reads (40.79% of the clean reads) were uniquely mapped to the draft assembly sequence using Bowtie2 v.2.3.2 [107] (-end-to-end --very-sensitive -L 30). The de-duplicated list of alignments of Hi-C reads to the draft ALO assembly was generated using Juicer v.1.5.7 [108]. Nine base pair-delimited resolutions (2.5, 1 Mbp, 500, 250, 100, 50, 25, 10, 5 kbp) were used to bin the reads and describe the interaction intensity of chromosome conformation. The 431 million (73.68% of unique mapped reads) valid paired-end reads were used to assemble the draft assembly into chromosome-length scaffolds with the linking information by LACHESIS [109]. Only these scaffolds >15 kbp were taken into the processes of cluster, order and orientation. The iterative round for mis-correction was set as zero time. The pseudomolecules were generated by concatenating the adjacent contigs with 100 ‘N’s [110]. Hi-C contact maps were processed by Pheatmap package for R v.3.6.3 [111] and reviewed in Juicer v.1.5.7 [108] (Additional file 1: Fig. S4, Additional file 2: Tables S4 and S5).

#### Estimation of genome size

Nuclear DNA content was estimated by flow cytometry [112]. The 20 mg of *A. brevis* (PI 657352) leaves (2C = 8.98 pg [8] served as an internal reference standard, were chopped with blades in 500 µl Otto I buffer solution (0.1 M citric acid, 0.5% v/v Tween 20 [113]). The homogenate was filtered through a 40 µm nylon mesh (BD Falcom^TM^, Cat. 352340). Nuclei were pelleted by centrifugation and resuspended in 400 µl of Otto I buffer. After 30 min incubation at room temperature, 800 µl of Otto II solution (0.4 M Na2HPO4) supplemented with 50 µg/ml RNase and 50 µg/ml propidium iodide was added. Samples were analysed by a CyFlow Space flow cytometer (Sysmex Partec GmbH, Görlitz, Germany) equipped with 533 nm laser. At least 5,000 nuclei were analysed per sample. Five plants were measured, and each plant was analysed three times on three different days. The 2C DNA content of ALO was calculated as 9.23 ± 0.20 pg (mean ± SD) by the ratio of G1 peak mean and standard value, then 1C genome size was calculated as 4,513 ± 0.099 Mbp (1 pg = 978 Mbp [112]).

Total 268.60 Gbp clean data were used for *k-*mer analysis by Kmerfreq_AR v.2.0.4 [114] (Luo et al., 2012) from SOAPec v.2.01 package (http://soap.genomics.org.cn/about.html) and Jellyfish v.2.2.6 [99] at 17-mer (Additional file 1: Fig. S2b). The genome size of *A*. *longiglumis* was estimated by the formula G = *k*-mer number / *k*-mer depth, where the *k*-mer number is the total numbers of *k*-mers, and *k*-mer depth refers to the depth of main peak. The genome size is expected to be 206,214,840,000/52 = 3.97 Gbp, which was close to the flow cytometry result. The *k*-mer (*k*=17) result indicated the heterozygosity of the ALO genome was approximately 0.48%.

#### Quality of genome assembly

The Illumina paired-end data were mapped to assembled scaffolds with Bowtie2 v.2.3.2 [107] (Langmead and Salzberg, 2012). The overall alignment rate was 99.94% with 96.90% properly paired alignments. We identified 3,830,731 heterozygous SNPs and 177,108 indels (depth ≥ 10 ×) in the ALO genome (Additional file 1: Table S4). The nanopore long reads were mapped to the assembled scaffolds using Minimap2 v.2.17 [115], and the depth of long reads was calculated using SAMtools [116] with default parameters.

The gene completeness of ALO assembly (Fig. 1) was evaluated by BUSCO v.4.0.5 [29] and CEGMA v.2.5 [30]. In BUSCO, a set of 1,375 plant-specific orthologous genes (Embryophyta_odb10) was used to search against genome assembly with parameters ‘-lineage_path embryophyta_odb10 –mode geno’ (Additional file 1: Table S5a). In CEGMA, a collection of 241 most conserved eukaryotic genes was searched against genome assembly with default parameters (Additional file 1: Table S5b). The gene completeness was defined by the proportion of completely matched proteins out of 1,375 embryophyta genes or 241 conserved eukaryotic genes. Finally, the LTR Assembly Index (LAI = 10.54) was calculated using the LTR_retriever [117].

#### RNA preparation and sequencing

Total RNA of four tissues [roots, salt-treated (the salt water of 4 mM NaCl for 48 h) roots, leaves and salt-treated leaves] were extracted using Column Plant RNAout 2.0 (Tiandz Inc., Beijing, China) according to the manufacturer’s protocol. Extracted RNA was treated with DNase (Tiandz Inc., Beijing, China) to remove genomic DNA. The RNA quality was validated using agarose gel electrophoresis, Nanodrop 2000 (Nanodrop Technologies Inc., NanoDrop 2000, Wilmington, USA), and Agilent 2100 (Agilent Technologies Inc., Pleasanton, USA) to confirm the purity, concentration and integrity, respectively. Library construction and sequencing were performed using Illumina Novaseq 6000 platform (Illumina Inc., San Diego, USA).

The clean data was generated by removing adaptor sequences, ambiguous reads (‘N’ > 10%) and low-quality reads (greater than 50% of bases in reads with a quality value Q ≤ 20) using Fastp v.0.20.1 [105]. The quality control of clean reads was filtered by FastQC v.0.11.3 (https://www.bioinformatics.babraham.ac.uk/projects/fastqc) for further genome-wide expression dominance analysis (https://ngdc.cncb.ac.cn/gsa/browse/CRA004247; Additional file 2: Tables S1 and S2).

### Genome sequence annotation

#### Repeat analysis

*De novo* repeat prediction of the ALO genome was carried out by EDTA v.1.7.0 (Extensive *de-novo* TE Annotator [36]) being composed of eight softwares (Fig. 2). LTRharvest [118], LTR_FINDER_parallel [36], LTR_retriever [117] were incorporated to identify LTR retrotransposons; Generic Repeat Finder [119] and TIR-Learner [120] were included to identify TIR transposons; HelitronScanner v.1.0 [121] identified *Helitron* transposons; RepeatModeler v.2.0.2a [122] was used to identify TEs (such as *LINEs*); Finally, RepeatMasker v.4.1.1 [123] was used to annotate fragmented TEs based on homology to structurally annotated TEs. In addition, TEsorter v.1.1.4 [124] was used to identify TE-related genes (Additional file 2: Table. S6a).

For comparison, the same protocol was applied to analyse repeat of six grass genomes, including *Aegilops tauschii* (ATA [32] Luo et al., 2017), *Brachypodium distachyon* (BDI) [33], *Oryza sativa* (OSA) [9], *Sorghum bicolor* (SBI) [10], *Setaria italica* (SIT) [11] and *Zea mays* (ZMA) [34]. For LTR-RTs, the families were clustered based on their LTR sequences. The final set of repetitive sequences in the ALO genome was obtained by integrating the *ab initio-* predicted TEs and those identified by homology through RepeatMasker (Additional file 2: Table. S6a). Intact LTR-RTs were identified and analyzed using LTR_retriever [116]. A nucleotide substitution rate (*r*) of 1.3× 10^−8^ mutations per site per year [125] was used to estimate the insertion time (*T*) of intact LTR-RTs with the formula of *T* = *K*/(*2r*) [126], where *K* is the divergence rate of 5’-LTR and 3’-LTR estimated by the Jukes-Cantor model (Additional file 2: Table. S6b, c).

Locations of centromeres were identified by multiple genomic features: (1) High abundance repetitive areas of repeat sequences on chromosome dotplots (Fig. 2, Additional file 2: Table S7a); (2) Discontinuities in the Hi-C contact map (Additional file 1: Fig. S4); (3) Location of barely (*Hordeum vulgare*) *Gypsy* LTR *Cereba* (KM948610 [41]) sequences are used to identify centromeres of wheat (*Triticum aestivum*, TAE; IWGSC, 2018), thus to identify the ALO centromeric regions, the *Cereba* sequence [42] was aligned to the ALO genome using Blastn to identify the centromere cores by Geneious Primer v.2021.1.1 (https://www.geneious.com/; Additional file 2: Table. S7b); (4) SynVisio [42] visualization of gaps and conserved regions between ALO and OSA assemblies (Additional file 2: Table. S7c); (5) Regions of low gene density along each ALO chromosome. The centromere cores are identified by the overlap regions of the high abundance repetitive areas on ALO chromosome dotplots and the low gene density areas on ALO chromosomes.

#### Gene prediction and functional annotation

Gene structure prediction depended on the application of three methods, i.e., *ab initio* prediction, homology-based prediction and RNA-seq-assisted prediction [127]. Augustus v.3.3.2 [128] was used for *de novo*-based gene predicition with default parameters to predict genes of the ALO genome. Additionally, the filtered proteins in genomes of six species ATA [31], BDI [32], *Hordeum vulgare* [65], SBI [10], *Triticum aestivum* [65] and ZMA [34] were used for homology-based prediction by GeMoMa v.1.6.1 [129] with default parameters (Additional file 2: Tables S8). Then, TransDecoder v.5.5.0 [130] were used for RNA-seq-based gene prediction. All predicted gene structures from three approaches were integrated into consensus gene models using EVidenceModeler v.1.1.1 [131]. These gene models were filtered sequentially to obtain a precise gene set, some genes whose sequences included transposable elements were removed with TransposonPSI v.2 (http://transposonpsi.sourceforge.net).

Gene functional annotation were carried out by performing BLASTP (E-value ≤ 1E-5) searches against NCBI non-redundant protein (NR) and Swiss-Prot (http://www.uniprot.org/) protein databases using BLASTP under the best match parameter [132]. NOG (Non-supervised Orthologous Groups), COG (Clusters of Orthologous Groups of proteins) [43], KEGG (Kyoto Encyclopedia of Genes and Genomes) [47], CAZy (Carbohydrate-Active enZYmes) [49], Pfam [44] annotations were performed with eggNOG v.5.0 [45]. The gene ontology (GO) IDs [46] for each gene were determined using the BLAST2GO v.1.44 [133]. Then transcription factors annotation was performed with PlantTFDB v.5.0 [48] (Additional file 2: Tables S9).

#### Identification of non-coding RNA genes

Genome-wide prediction of non-coding RNA gene set (ncRNA) was performed (Additional file 2: Tables S10). Firstly, the data set was aligned to the Rfam library v.11.0 [134] noncoding database to annotate genes encoding ribosomal RNA (rRNA), small nuclei RNA (snRNA) and microRNA (miRNA). Then the transfer RNA (tRNA) sequences were identified using tRNAscan-SE v.2.0 [135]. Meanwhile, miRNAs were predicted by miRanda v.3.0 [136], while rRNA and its subunits were predicted by RNAmmer v.1.2 [137].

#### Identification of high- and low-confidence genes

The 40,845 gene set was filtered to identify high-confidence (HC) protein-coding genes by two methods. Transcriptome raw reads were preprocessed by Fastp v.0.20.1 [105] with default parameters in order to trim adaptors and remove the low-quality RNA-seq reads (Phred quality scores < 20). The clean reads were aligned to the ALO genome by STAR aligner [138]. The initial SAM-to-BAM conversion was performed by SAMtools [116]. The mapped RNA-seq reads (in BAM file) were assembled to transcript by Stringtie v.2.0.6 [139], which was used to call the fragments per kilobase of exon model per million mapped reads (FPKM) values. Subsequently, the genes with FPKM value larger than zero were classified as HC (Additional file 2: Table S11a). For the genes without transcriptome transcript abundance support, the alignment was performed with *A*. *atlantica* (identity > 95%, coverage > 95%), *A*. *eriantha* (identity > 90%, coverage > 90%), *Hordeum vulgare* and *Triticum aestivum* (identity > 80%, coverage > 80%) by BLASTP (https://blast.ncbi.nlm.nih.gov/Blast.cgi; *E* value=1e-5), respectively. Those supported by alignment results of two or more species alignments were defined as HC genes (Additional file 2: Table S11b). Finally, the HC genes were supported by transcriptome data or homology, while the low-confidence (LC) genes were not supported by either one method (Additional file 2: Table S11c).

### Evolutionary analysis

#### Gene family identification and phylogenetic tree reconstruction

To examine evolution and divergence of the ALO genome, protein-coding gene sequences from 10 species, *Avena atlantica* (AAT) [12], *A*. *eriantha* (AER) [12], *A*. *strigosa* (AST) [7], ATA [32], *Arabidopsis thaliana* (ATH) [50], BDI [33], OSA [9], SBI [10] (McCormic et al., 2018), SIT [11] (Yang et al., 2020) and ZMA [34], were downloaded from Phytozome v.13 [140] and NCBI website (https://www.ncbi.nlm.nih.gov/) for comparative analyses (Additional file 2: Tables S12 and S13). When one gene had multiple transcripts, only the longest transcript in the coding region was kept for further analysis. Paralogs and orthologs were clustered with OrthoFinder v.2.3.14 [51] through standard mode parameters with Diamond v.0.9.24 [141]. The numbers of shared and species-specific gene families among five species (ALO, AER, AST, OSA and ZMA) were visualized by UpSetR v.1.4.0 [142] for R v.3.6.3 [111].

Single-copy of orthologous genes were extracted from the OrthoFinder [51] clustering results and MAFFT v.7.48 [143] was used to align the concatenated protein sequences to give a super-gene matrix. RAxML v.8.1.17 [144] was used to reconstruct a phylogenetic tree with the GTR+G+I model and a bootstrap value of 1000 (Fig. 3). The phylogenetic tree was visualized by FigTree v.1.4.4 (http://tree.bio.ed.ac.uk/software/figtree/). Species divergence time and the 95% confidence intervals were inferred by MCMCtree in the PAML v.4.9i [145] via the Markov Chain Monte Carlo method. A correlated rate model (clock = 3) was established, and MCMC was performed (burnin = 2,000, sample number = 20,000, sample-frequency = 2). The crown ages for Stipeae (25.5–30.1 Mya), Oryzeae (28.4–33.4 Mya) and BEP/PACMAD split (45.0–57.4 Mya) [4], obtained from the TimeTree database (http://www.timet ree.org/), were applied to calibrate the divergence times.

#### Expansion and contraction of gene families

To analyse the expanded and contracted gene families, the Computational Analysis of Gene Family Evolution software (CAFE v.4.2) [146] was run to compute changes in gene families along each lineage of the phylogenetic tree under a random birth-and-death model (Additional file 2: Table S14). The expanded and contracted gene families were localized on the chromosomes (Fig. 1, circles f and g). Although these regions may be under selective pressure and reflect major duplications or deletions at the whole-genome level, the distribution of changed gene families largely reflects overall chromosomal gene density. The clustering results and the information from the estimated divergence times were used. Using conditional likelihood as the test statistics, the corresponding *P*-value of each lineage were calculated (*P*-value ≤ 0.01). Additionally, the GO enrichment of expanded genes and KEGG enrichment of unique genes were analysed to determine their functions (Additional file 2: Tables S15 and S16). Genomic landscape of repeats, genes and expanded genes were plotted across the ALO chromosomes by TBtools v.1.092 [147].

#### Genome synteny and whole genome duplication analysis

Protein sequences within and between genomes were searched against one another to detect putative homologous genes (*E* value < 1e-5) by BLASTP. With information about homologous genes as input, MCscanX [148] were implemented to infer homologous blocks involving collinear genes within and between genomes. The maximum gap length between collinear genes along a chromosome region was set to 50 genes [149]. Then, homology dotplots were constructed by a perl script to reveal genomic correspondence. Then, homology dotplots were constructed by SynVisio [42] to reveal genomic correspondence within ALO, between three *Avena* species, between ancestral grass karyotype (AGK) and seven grass species, and between ALO, OSA and BDI (Fig. 4, Additional file 2: Table S17). Subsequently, we applied the WGDI v.0.4.7 [150] to identify the whole genome duplication events based on the high synonymous (Ks) peak of ALO versus AST, ALO versus AER, AST versus AER collinear gene pairs. Non-synonymous (Ka) and Ks substitution rates for gene pairs were calculated with KaKs_Calculator v.2.0 [151] under the YN model. The synonymous substitutions rate per site per year (*r*) equaling 1.3× 10^−8^ was applied to the recent WGD estimation [152].

### Anti-salinity and secondary metabolism gene cluster analysis

#### Genome-wide expression analysis

To investigate the expression dominance of salt-responsive gene, the FPKM values were calculated for genes in roots, leaves, salt-treated roots and salt-treated leaves. Differentially expressed genes (DEGs) were identified by DEseq2 v.3.11 [153]. We filtered the DEGs with a minimum of two-fold differential expression (|fold change| ≥ 2; false discovery rate (FDR) ≤ 0.005). The DEGs were performed KEGG enrichment by TBtools [147] (Additional file 2: Table S18).

#### Metabolic gene cluster prediction and CYP450 gene identification

The plantiSMASH v.1.3 [154] was used to identify the potential metabolic gene clusters in the ALO genome with parameter setting “run_antismash.py -c 16 --taxon plants tanxaing.gb -- outputfolder tanxiang” (Fig. 5, Additional file 2: Table S19). To identify the *CYP450* gene numbers in genomes of ALO, nine grass species (AAT, AER, AST, ATA, BDI, OSA, SBI, SIT and ZMA) and *Arabidopsis thaliana*, all proteins of each species was searched against hidden Markov model (HMM) profile of the Pfam domain (PF00067) by hmmsearch (http://hmmer.org/). Putative *CYP450* genes were further verified in the Pfam database (PF00067) to confirm the CYP450 proteins of ALO. The *CYP450* gene copy number and syntenic relationships between ALO and the nine grass species and *Arabidopsis thaliana* were visualized by TBtools v.1.092[147] (Additional file 2: Tables S20–S23).

#### CesA, Csl and CalS gene identification

To identify the *CesA*, *Csl* and *CalS* (or *GSL*) gene family members in ALO, all proteins of ALO was searched against hidden Markov model (HMM) profile of the Pfam domain [(PF00535 or PF03552 for *CesA* and *Csl*) and (PF02364 and PF14288 for *CalS*)] by hmmsearch (http://hmmer.org/). Putative *CesA*, *Csl* and *CalS* genes were further verified in the Pfam database [155] (http://pfam.xfam.org/), screened for Pfam domains [(PF00535 or PF03552 for CesA and Csl) and (PF02364 and PF14288 for CalS)] to confirm as the *CesA*, *Csl* and *CalS* proteins of ALO (Additional file 2: Table S24). The *CYP450* gene copy number and syntenic relationships between ALO and nine grass species [AAT, AER, AST, ATA, BDI, OSA, SBI, SIT and ZMA] and *Arabidopsis thaliana* were visualized by TBtools v.1.092 (Additional file 2: Tables S24–S30). Previously known *CesA* and *Csl* protein sequences were downloaded for *Arabidopsis thaliana*, rice, wheat amd maize (Kaur et al. [156] their Fig. S1). GSL protein sequences of *Arabidopsis thaliana* and rice were downloaded from RGAP http://rice.uga.edu/index.shtml) and TAIR (https://www.arabidopsis.org/). Multiple sequence alignments of CesA and Csl proteins and GSL proteins were performed by MAFFT v.7.48 [142] with default parameters (Additional file 1: Fig. S34, Additional file 2: Tables S31 and S32). A maximum likehood (ML) phylogenetic tree was constructed using FastTree v.2.1.10 with GTR model [157] and 1000 bootstrap replicates. The phylogenetic tree was visualized by FigTree v.1.4.4 (Fig. 5b).

## Supplementary information

Supplementary infromation accompanies this paper at https://doi.org/10.xxxx/syyyyy-yyy-yyyy-y.

**Additional file 1: Fig. S1.** Flow cytometric estimation of the nuclear genome size of *Avena longiglumis*. Nuclei were isolated from *A. longiglumis* (PI 657387) and *A. brevis* (CN 1979; used as an internal reference standard), stained and analyzed simultaneously. **Fig. S2.** *Avena longiglumis* spikelets and genome assembly. **Fig. S3.** Strategy for sequencing, assembly and annotation of the *Avena longiglumis* genome. **Fig. S4.** Inter-chromosomal contact matrix. The intensity of pixels represents the normalized count of Hi-C links between 100 kbp windows on ALO chromosomes on a logarithmic scale. **Fig. S5.** Evaluation of genome assemblies by LTR Assembly Index (LAI). **Fig. S6.** Gene density of 1 Mbp-sized sliding windows on seven chromosomes of *Avena longiglumis*. **Fig. S7.** Centromeric retrotransposon *Cereba* (KM948610 [41]) sequence locations on seven chromosomes of *Avena longiglumis*. Centromere area denoted by red dots. **S8.** KEGG enrichment of expanded genes in four *Avena* species. **Fig. S9.** KEGG enrichment of unique genes of four *Avena* species. **Fig. S10.** Syntenic relationships based on three *Avena* species genomes. **Fig. S11.** Syntenic relationships based on AST-ALO-AER and OSA-ALO-BDI genome homologous genes. **Fig. S12.** KEGG enrichment of up- and down-regulated DEFs in roots versus salt-treated roots and leaves versus salt-treated leaves of ALO. **Fig. S13.** Statistics of expanded DEGs (number ≥ 5) of ALO. **Fig. S14.** Heat map showing hierarchical clustering of *cytochrome P450* gene families in roots, salt-treated roots, leaves and salt-treated leaves of ALO. **Fig. S15.** Total 557 *cytochrome P450* genes located within 57 clusters identified in the ALO genome. **Fig. S16.** *Cytochrome P450* genes inserted within gene-clusters identified on ALO chromosomes. **Fig. S16a.** *Cytochrome P450* genes inserted within gene-clusters identified on ALO01. **Fig. S16b.** *Cytochrome P450* genes inserted within gene-clusters identified on ALO02. **Fig. S16c.** *Cytochrome P450* genes inserted within gene-clusters identified on ALO03. **Fig. S16d.** *Cytochrome P450* genes inserted within gene-clusters identified on ALO04. **Fig. S16e.** *Cytochrome P450* genes inserted within gene-clusters identified on ALO05. **Fig. S16f.** *Cytochrome P450* genes inserted within gene-clusters identified on ALO06. **Fig. S16g.** *Cytochrome P450* genes inserted within gene-clusters identified on ALO07. **Fig. S17.** Total 11 *CesA* and 55 *Csl* genes located within 10 clusters identified in the ALO genome. **Fig. S18.** Ten *CesA* and *Csl* gene clusters on six chromosomes of ALO. **Fig. S18a.** *CesA* and *Csl* genes inserted gene-clusters identified on ALO02. **Fig. S18b.** *Csl* genes inserted within gene-cluster identified on ALO03. **Fig. S18c.** *Csl* genes inserted within gene-cluster identified on ALO04. **Fig. S18d.** *Csl* genes inserted within gene-clusters identified on ALO05. **Fig. S18e.** *Csl* genes inserted within gene-cluster identified on ALO06. **Fig. S18f.** *Csl* genes inserted within gene-cluster identified on ALO07. **Fig. S19.** FASTA sequences of Csl proteins of ALO used for the phylogenetic analysis. **Fig. S20.** Heat map showing hierarchical clustering of the ALO *callose* gene (*CalS*, *GSL*) families in roots, salt-treated roots, leaves and salt-treated leaves. **Fig. S21.** The maximum likelihood phylogenetic tree constructed with CalS (GSL) proteins of ALO, *Arabidopsis thaliana* and *Oryza sativa*. **Fig. S22.** FASTA sequences of CalS (GSL) proteins of ALO used for the phylogenetic analysis.

**Additional file 2: Table S1.** Summary of sequencing libraries of *Avena* species included in the study. **Table S2.** Deposited data of *Avena longiglumis* (ALO) genome and software used in the study. **Table S3.** Summary of genome assembly and annotation of ALO. **Table S4.** Statistics of the ALO genome assembly consistency. **Table S5.** Evaluation of gene space completeness in the ALO genome assembly. **Table S6.** Repetitive DNA composition comparison among genomes of ALO and six grass species. **Table S7.** Size and centromere localization of the ALO pseudomolecules. **Table S8.** Gene characterization comparison among ALO and ten other plant species. **Table S9.** Statistics of gene function annotation of the ALO genome. **Table S10.** Statistics of annotated non-coding RNAs of the ALO genome. **Table S11.** Identification of high-confidence (HC) and low-confidence (LC) protein-coding genes annotated in the ALO genome. **Table S12.** Gene family categories in genomes of ALO and ten plant species. **Table S13.** Gene family statistics of ALO and ten plant species. **Table S14.** Summary of orthologous gene clusters analyzed in analysed species. **Table S15.** GO enrichment analysis of expanded genes in the ALO genome. **Table S16.** KEGG enrichment analysis of unique genes in the ALO genome. **Table S17.** The gene pair statistics of SynVisio results between post-ρ ancestral grass karyotype (AGK) and grass species and between grass species. **Table S18.** Statistics of up- and down-regulated DEGs in salt-treated roots and salt-treated leaves and expanded gene families of ALO. **Table S19.** Characterization of biosynthetic gene clusters (BGCs) in the ALO genome. **Table S20**. The *CYP450* gene copy number in analysed species and distribution along the ALO chromosomes. **Table S21.** FPKM value of *CYP450* genes in roots, leaves, salt-treated roots and salt-treated leaves of ALO. **Table S22.** Description of 46 BGCs containing 117 *CYP450* genes in the expression profiling heat map. **Table S23.** The conserved synteny among *CYP450* genes within gene clusters in ALO and nine grass species. **Table S24.** Statistics of 11 *CesA* and 55 *Csl* genes corresponding to the expanded gene families in the ALO genome. **Table S25.** Distribution of 11 *CesA* and 55 *Csl* genes in the ALO chromosomes. **Table S26.** FPKM value of *CesA* and *Csl* genes in roots, salt-treated roots, leaves and salt-treated leaves of ALO. **Table S27.** Gene description of 10 BGCs containing *CesA* and *Csl* genes in the expression profiling heat map. **Table S28.** The conserved synteny among *CesA* and *Csl* genes within gene clusters in ALO and nine grass species. **Table S29**. Statistics of 13 *CalS* (*GSL*) genes corresponding the expanded gene families in the ALO genome. **Table S30**. FPKM value of *CalS* genes in roots, salt-treated roots, leaves and salt-treated leaves of ALO.

Additional file 3: Review history.

## Acknowledgements

We thank Jun Wen, Dallas Kessler and Bockelman Harold for the seed collection, Shuyu Zhou, Haoyan Pan and Dongli Cui for the assistance with growing and maintaining the plants, and Yubo Wang, Kai Ouyang, and Huitong Tan for the statistical advice. We thank Grandomics Biosciences Co., Ltd. (Wuhan, China) for sequencing support, Huawei Elastic Cloud Server (Jiangsu, China) for supplying the computational resources.

## Review history

The review history is available as Additional file 3.

## Peer review information

XXX was the primary editor of this article and managed its editorial process and peer review in collaboration with the rest of the editorial team.

## Authors’ contributions

QL and JSHH designed the study. ZWW, YSY and XKT collected samples. ZYS and XKT sequenced DNA. MZL, ZWW and DLC performed genome assembly, polishing, validation, annotation and analysis. HYY, MZL, ZWW and ZYS performed repeat and transcriptome sequence analysis. TS and JSHH supervised genome assembly and analysis. QL, HYY and JSHH wrote the manuscript. TS and JSHH revised the manuscript. All authors read and approved the final manuscript.

## Availability of data and materials

The sequencing data used in this study have been deposited into the Genome Sequence Archive (GSA) database in BIG Data center under Accession Number PRJCA004488/CRR275304-CRR275326 and CRR285670-285674 (https://ngdc.cncb.ac.cn/gsa/browse/CRA003996 for raw data of the ALO genome; https://ngdc.cncb.ac.cn/gsa/browse/CRA004247 for raw data of ALO transcriptome). The previously reported Illumina data for were deposited into the NCBI database under Accession Number SRA: SRR6058489-SRR6058492 and from NCBI under BioProject PRJNA407595 (https://www.ncbi.nlm.nih.gov/bioproject/?term=PRJNA407595). The ALO genome assembly was uploaded to https://figshare.com/s/34d0c099e42eb39a05e2.

## Ethics approval and consent to participate

Not applicable.

## Consent for publication

Not applicable.

