## Supplementary Figures and Tables for "Chromosome-scale genome assembly of the diploid oat *Avena longiglumis* reveals the landscape of repetitive sequences, genes and chromosome evolution in grasses": Liu_et_al2022_ALO_Genome_SupplementaryFiguresTables.pdf

### Genome assembly and evolutionary genomics of wild oats

### Supplementary Figures

**Additional file 1: Fig. S1.** Flow cytometric estimation of the nuclear genome size of *Avena longiglumis*. 3

**Fig. S2.** *Avena longiglumis* spikelets and genome assembly. 4

**Fig. S3.** Strategy for sequencing, assembly and annotation of the *Avena longiglumis* genome. **Fig. S4.** Inter-chromosomal contact matrix. 5

**Fig. S5.** Evaluation of genome assemblies by LTR Assembly Index (LAI). 7

**Fig. S6.** Gene density of 1 Mbp-sized sliding windows on seven chromosomes of *Avena longiglumis*. 8

**Fig. S7.** Centromeric retrotransposon *Cereba* (KM948610 [41]) sequence locations on seven chromosomes of *Avena longiglumis*. 9

**Fig. S8.** KEGG enrichment of expanded genes in four *Avena* species. 10

**Fig. S9.** KEGG enrichment of unique genes of four *Avena* species. 11

**Fig. S10.** Syntenic relationships based on three *Avena* species genomes. 12

**Fig. S11.** Syntenic relationships based on AST-ALO-AER and OSA-ALO-BDI genome homologous genes. 13

**Fig. S12.** KEGG enrichment of up- and down-regulated DEFs in roots versus salt-treated roots and leaves versus salt-treated leaves of ALO. 14

**Fig. S13.** Statistics of expanded DEGs (number ≥ 5) of ALO. 15

**Fig. S14.** Heat map showing hierarchical clustering of *cytochrome P450* gene families in roots, salt-treated roots, leaves and salt-treated leaves of ALO. 16

**Fig. S15.** Total 557 *cytochrome P450* genes located within 57 clusters identified in the ALO genome. 17

**Fig. S16.** *Cytochrome P450* genes inserted within gene-clusters identified on ALO chromosomes. 18

***Fig. S16a.*** *Cytochrome P450 genes inserted within gene-clusters identified on ALO01. 18*

***Fig. S16b.*** *Cytochrome P450 genes inserted within gene-clusters identified on ALO02. 19*

***Fig. S16c.*** *Cytochrome P450 genes inserted within gene-clusters identified on ALO03. 20*

***Fig. S16d.*** *Cytochrome P450 genes inserted within gene-clusters identified on ALO04. 21*

***Fig. S16e.*** *Cytochrome P450 genes inserted within gene-clusters identified on ALO05. 22*

***Fig. S16f.*** *Cytochrome P450 genes inserted within gene-clusters identified on ALO06. 23*

***Fig. S16g.*** *Cytochrome P450 genes inserted within gene-clusters identified on ALO07. 24*

**Fig. S17.** Total 11 *CesA* and 55 *Csl* genes located within 10 clusters identified in the ALO genome. 25

**Fig. S18.** Ten *CesA* and *Csl* gene clusters on six chromosomes of ALO. 26

***Fig. S18a.*** *CesA and Csl genes inserted gene-clusters identified on ALO02. 26*

***Fig. S18b.*** *Csl genes inserted within gene-cluster identified on ALO03. 27*

***Fig. S18c.*** *Csl genes inserted within gene-cluster identified on ALO04. 28*

***Fig. S18d.*** *Csl genes inserted within gene-clusters identified on ALO05. 29*

***Fig. S18e.*** *Csl genes inserted within gene-cluster identified on ALO06. 30*

***Fig. S18f.*** *Csl genes inserted within gene-cluster identified on ALO07. 31*

**Fig. S19.** FASTA sequences of Csl proteins of ALO used for the phylogenetic analysis. 32

**Fig. S20.** Heat map showing hierarchical clustering of the ALO *callose* gene (*CalS*, *GSL*) families in roots, salt-treated roots, leaves and salt-treated leaves. 91

**Fig. S21.** The maximum likelihood phylogenetic tree constructed with CalS (GSL) proteins of ALO, *Arabidopsis thaliana* and *Oryza sativa*. 92

**Fig. S22.** FASTA sequences of CalS (GSL) proteins of ALO used for the phylogenetic analysis. 93

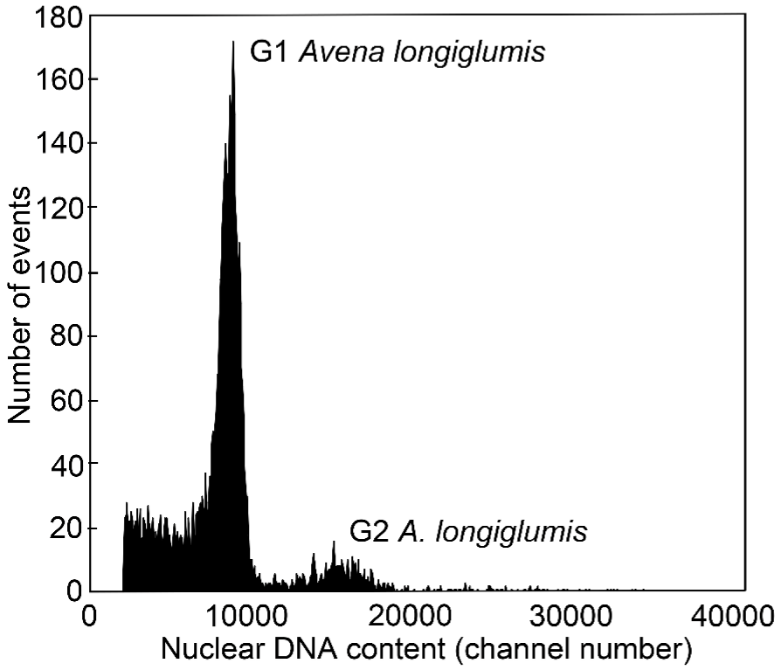

**Fig. S1.** Flow cytometric estimation of the nuclear genome size of *Avena longiglumis*. Nuclei were isolated from *A. longiglumis* (PI 657387) and *A. brevis* (CN 1979; used as an internal reference standard), stained and analyzed simultaneously. The 2C DNA content was calculated using the ratio of the G1 peak means of *A. longiglumis* and the *A. brevis* standard, giving a 2C value for *A. longiglumis* of 8.486 ± 0.074 pg DNA (mean ± SD). The 1C genome size in base pairs was calculated using the formula 1pg DNA = 0.978 $\times 10^9$ bp, giving a 1C *A. longiglumis* genome size of 4149 ± 0.036 Mbp (mean ± SD).

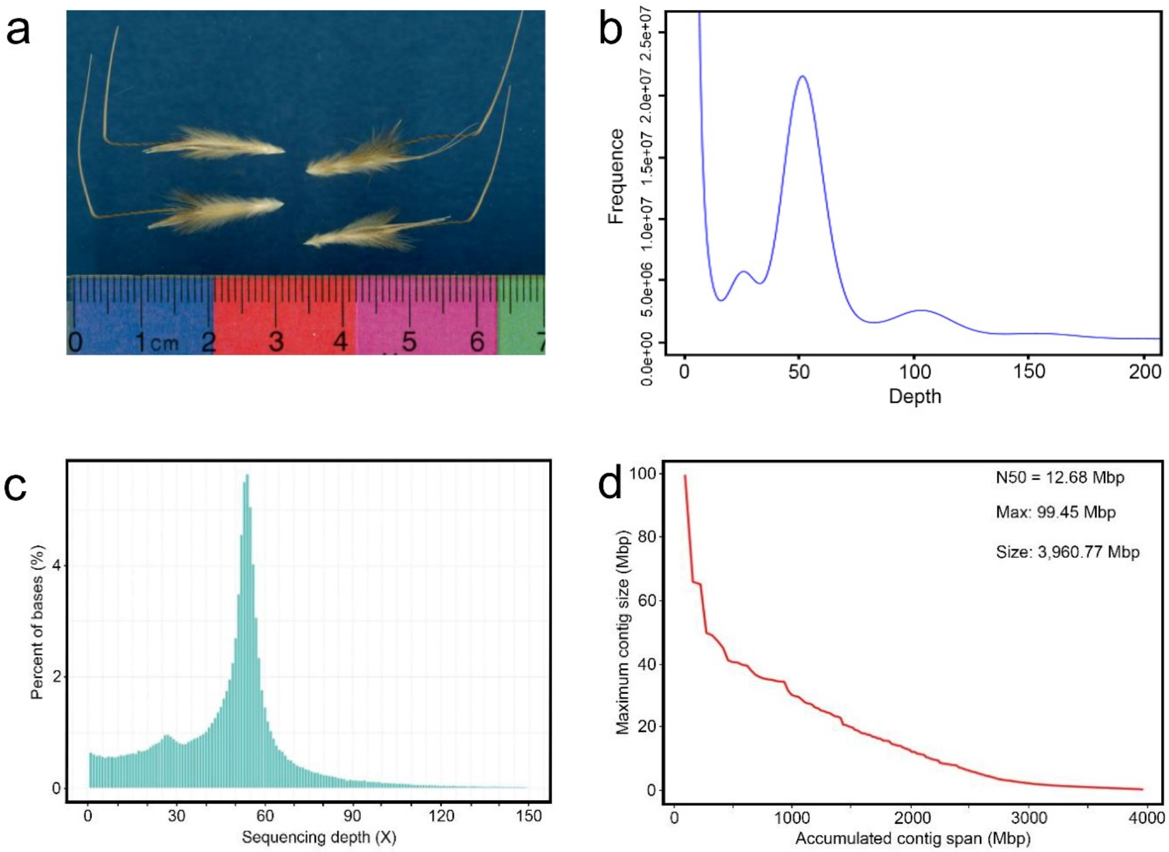

**Fig. S2.** *Avena longiglumis* spikelets and genome assembly. **a** *Avena longiglumis* spikelets. **b** *K*-mer frequency distributions. When *k*-mer = 17, a frequency peak value at 52 is observed and used to estimate the genome size. **c** Illumina genome sequencing mean depth is 67.55 ×. **d** Contig length distribution of corrected *A. longiglumis* genome (the longest contig is 99.45 Mbp).

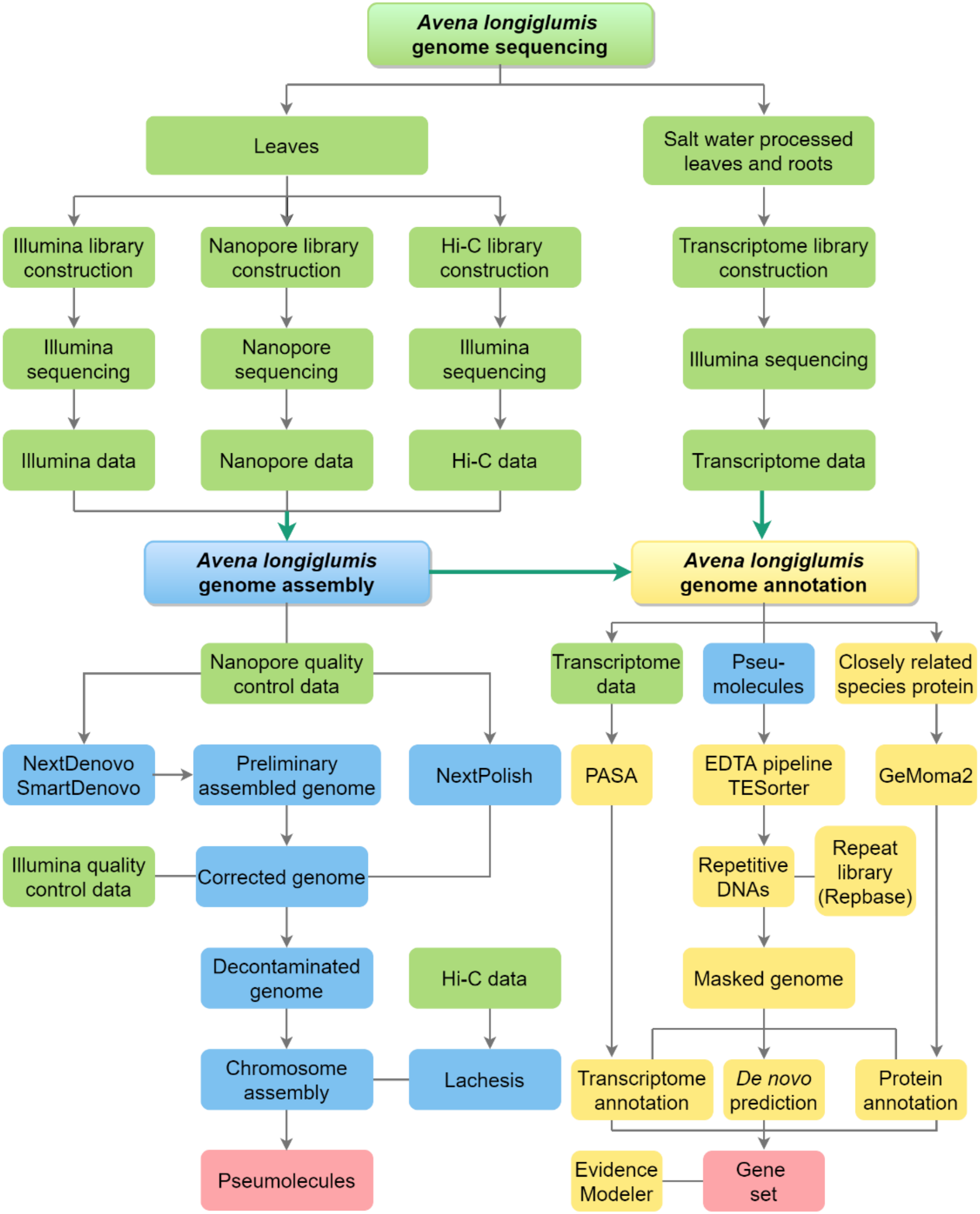

**Fig. S3.** Strategy for sequencing, assembly and annotation of the Avena longiglumis genome.

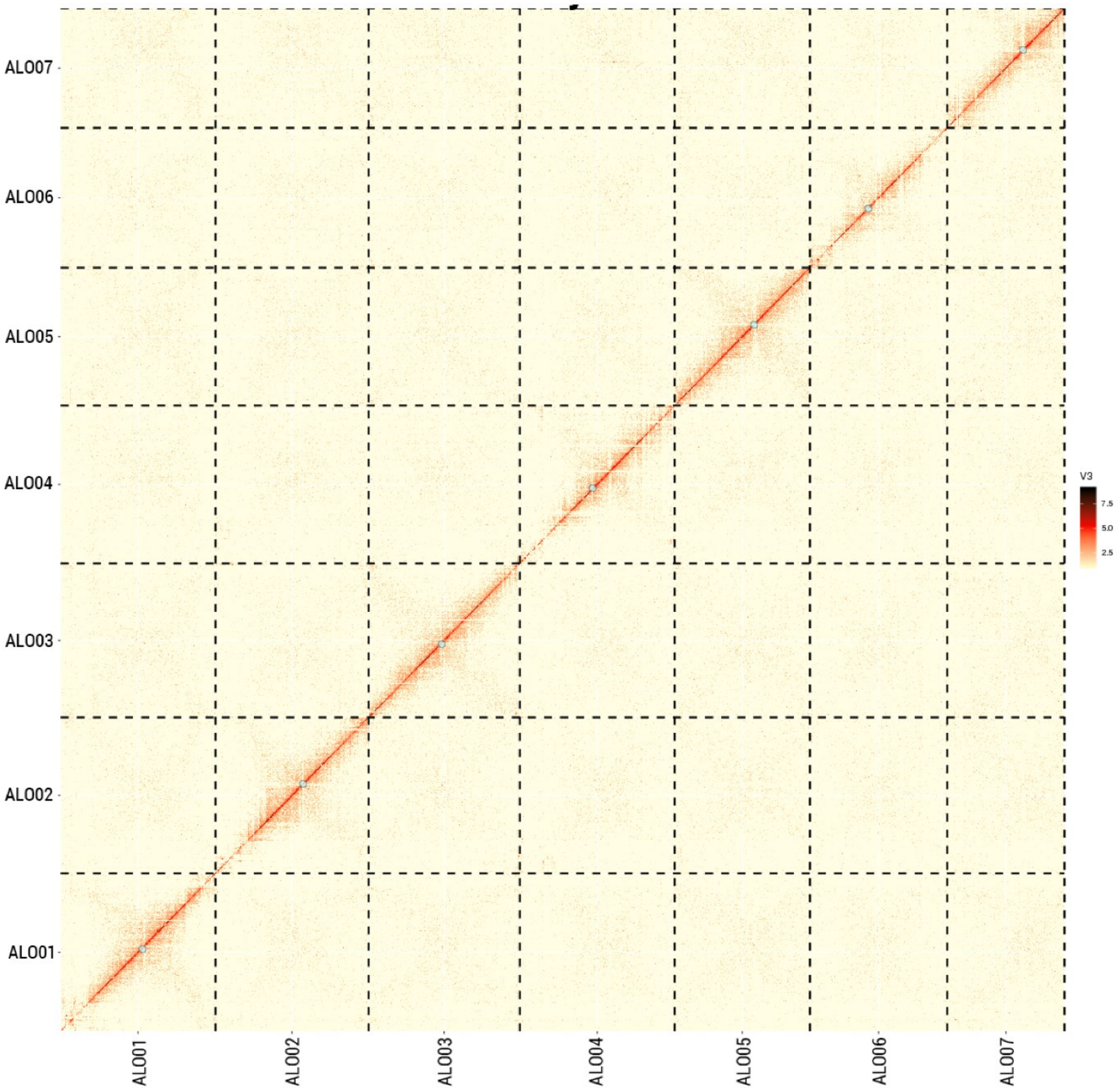

**Fig. S4.** Inter-chromosomal contact matrix. The intensity of pixels represents the normalized count of Hi-C links between 100 kbp windows on ALO chromosomes on a logarithmic scale. Green shady cirles denoted centromeric locations.

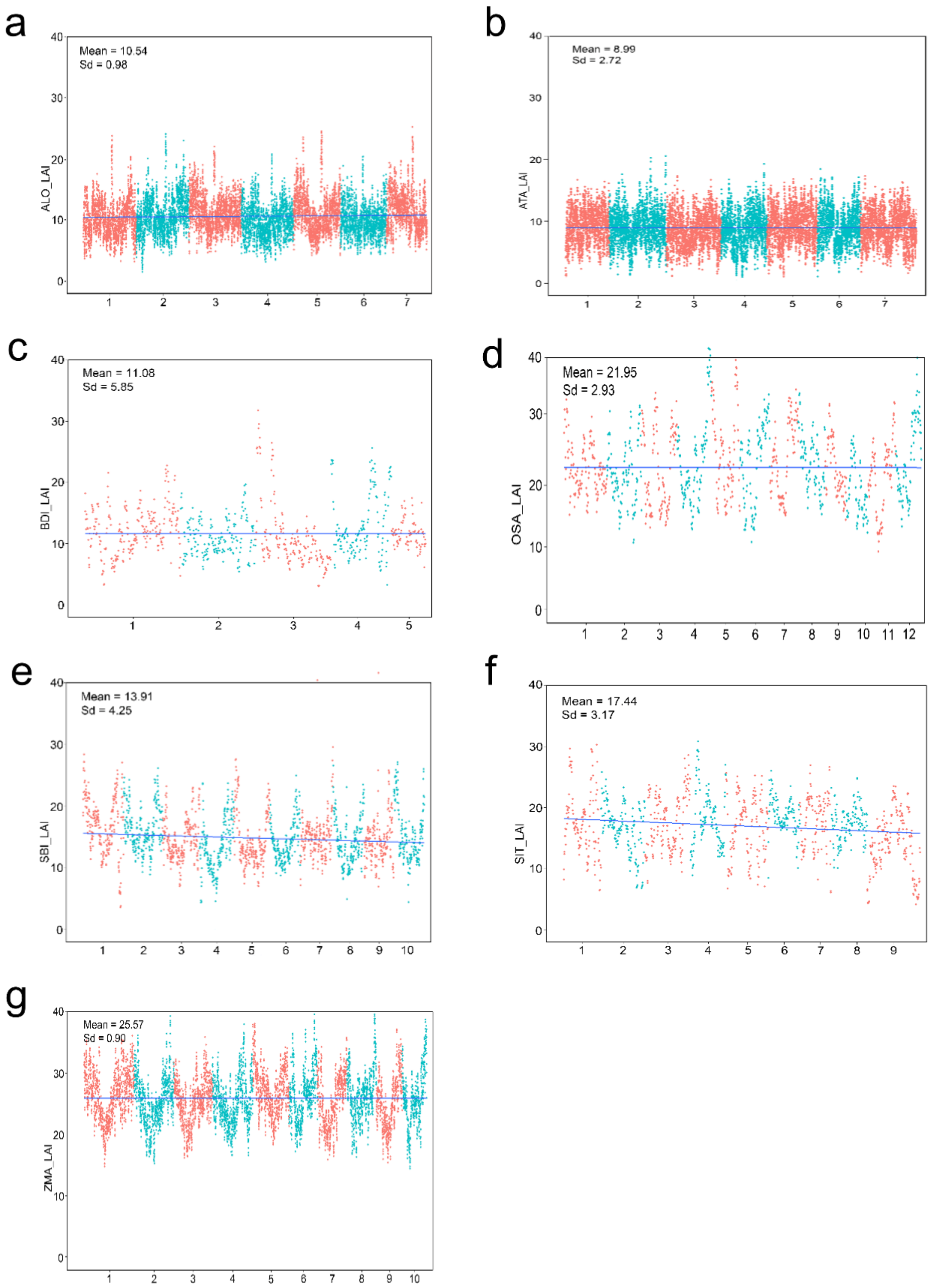

**Fig. S5.** Evaluation of genome assemblies by LTR Assembly Index (LAI). **a** *Avena longiglumis* (ALO). **b** *Aegilops tauschii* (ATA) [32]. **c** *Brachypodium distachyon* (BDI) [33]. **d** *Oryza sativa* (OSA) [9]. **e** *Sorghum bicolor* (SBI) [10]. **f** *Setaria italica* (SIT) [11]. **g** *Zea mays* (ZMA) [34]. The x-axes show the chromosomes of each genome. LAI scores, represented by the dots, were calculated using 3-Mbp-sliding windows with 300 kbp steps. The blue line indicates whole genome mean of LAI scores.

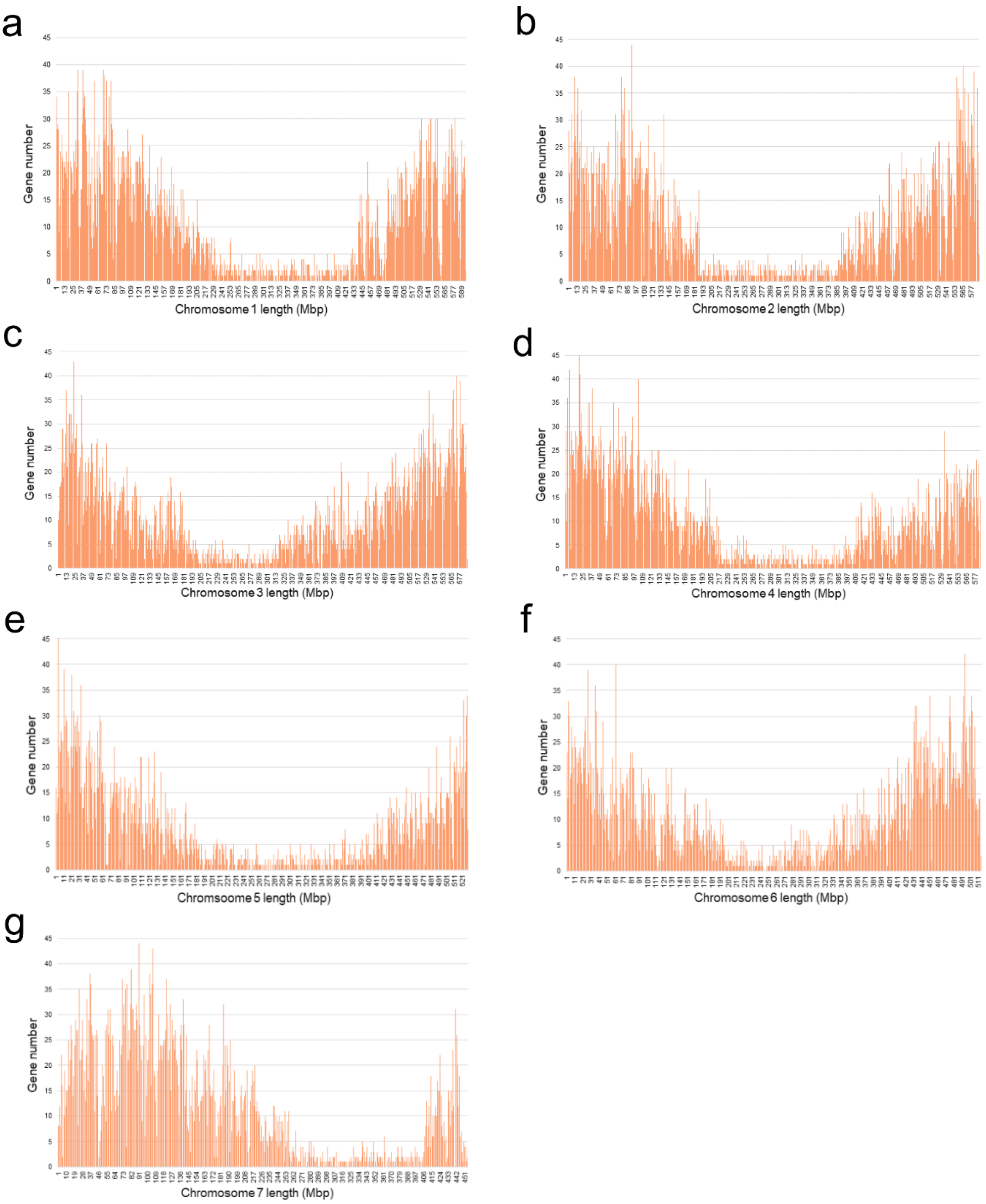

**Fig. S6.** Gene density of 1 Mbp-sized sliding windows on seven chromosomes of *Avena longiglumis*. **a** ALO01 histogram. **b** ALO02 histogram. **c** ALO03 histogram. **d** ALO04 histogram. **e** ALO05 histogram. **f** ALO06 histogram. **g** ALO07 histogram.

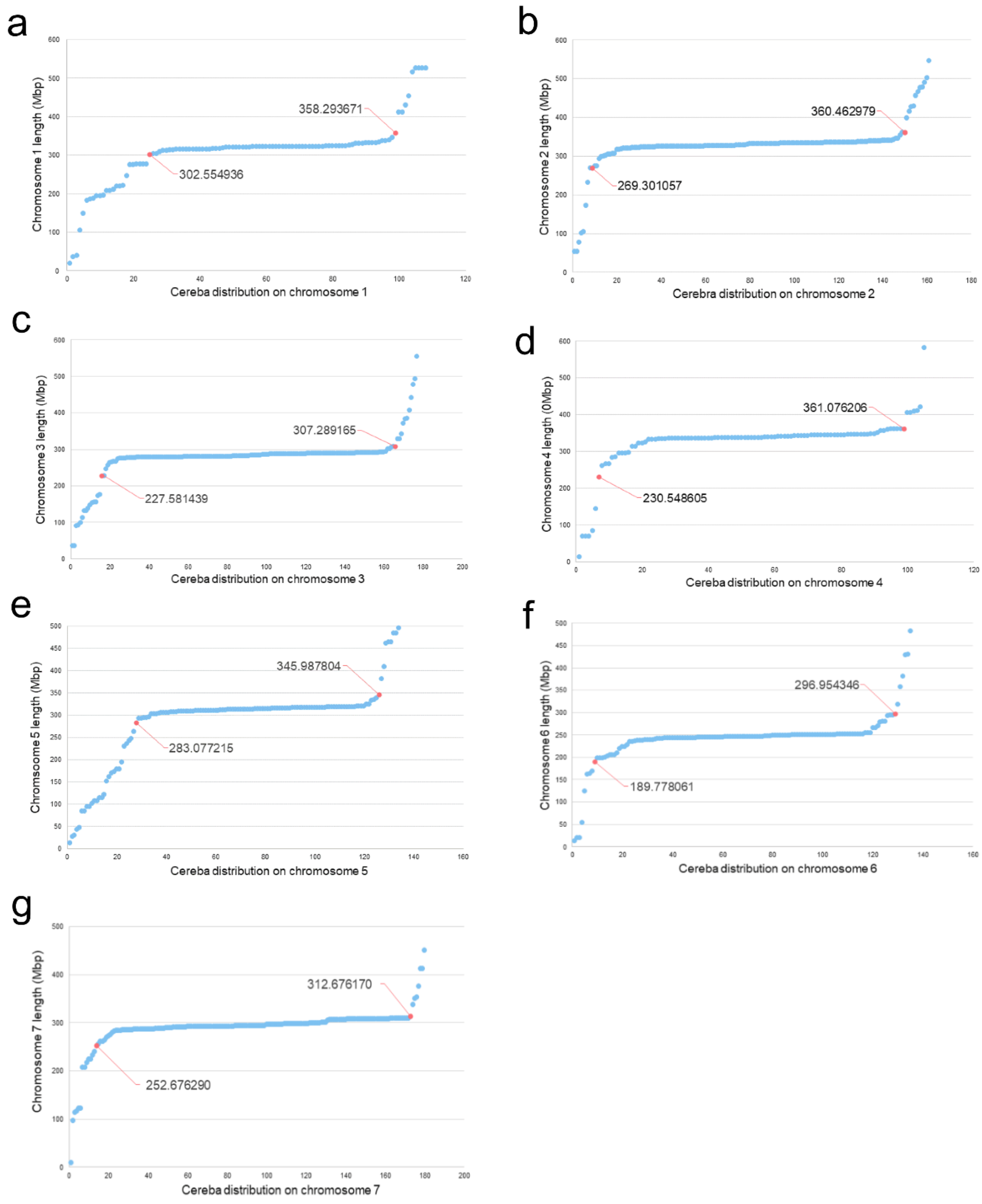

**Fig. S7.** Centromeric retrotransposon *Cereba* (KM948610 [41]) sequence locations on seven chromosomes of *Avena longiglumis*. Centromere area denoted by red dots. **a** Relatively continuous distribution of *Cereba* from 302,554,936 bp to 358,293,671 bp on chromosome 1 of *A. longiglumis* (ALO01). **b** Relatively continuous distribution of *Cereba* from 269,301,057 bp to 360,462,979 bp on ALO02. **c** Relatively continuous distribution of *Cereba* from 227,581,439 bp to 307,289,165 bp on ALO03. **d** Relatively continuous distribution of *Cereba* from 230,548,605 bp to 361,076,206 bp on ALO04. **e** Relatively continuous distribution of *Cereba* from 283,077,215 bp to 345,987,804 bp on ALO05. **f** Relatively continuous distribution of *Cereba* from 189,778,061 bp to 296,954,346 bp on ALO06. **g** Relatively continuous distribution of *Cereba* from 252,676,290 bp to 312,676,170 bp on ALO07.

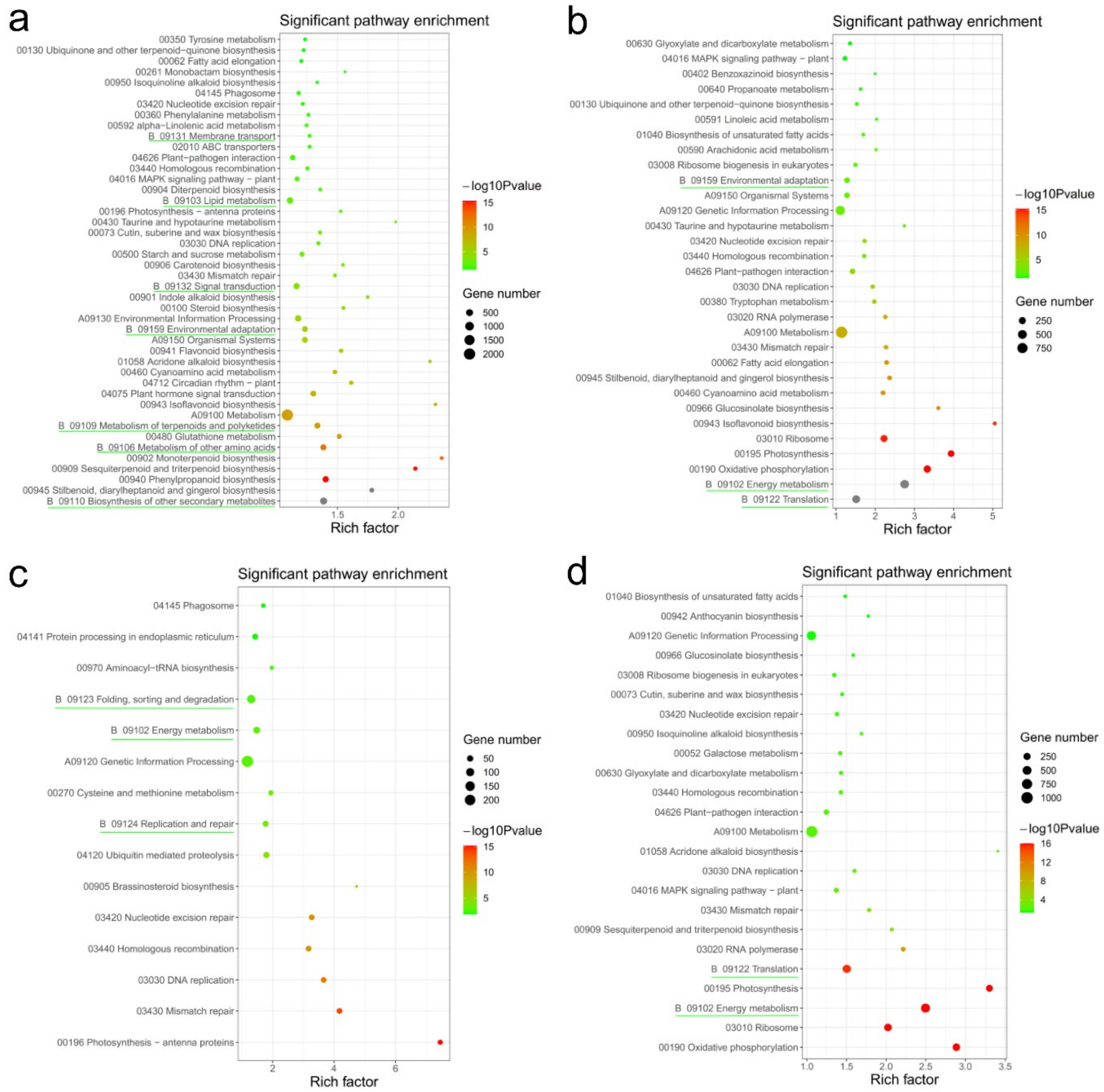

**Fig. S8.** KEGG enrichment of expanded genes in four *Avena* species. **a** ALO genome. **b** AAT genome. **c** AST genome. **d** AER genome. The corrected *p* value (Q value) is zero for black points in Fig. S8a, b and d.

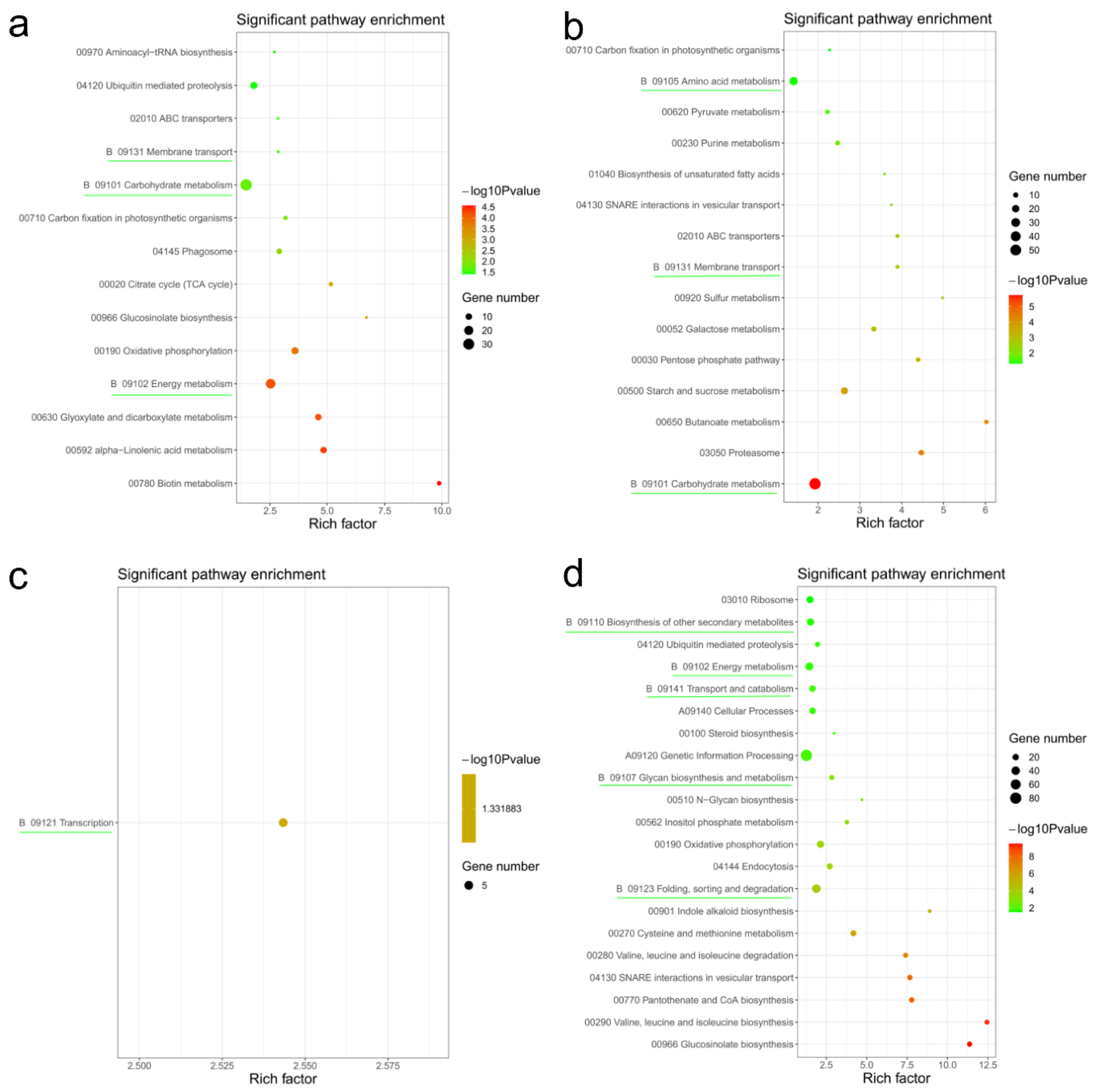

**Fig. S9.** KEGG enrichment of unique genes of four *Avena* species. **a** ALO genome. **b** AAT genome. **c** AST genome. **d** AER genome. The corrected *p* value (Q value) is zero for black points in Fig. S9b, d.

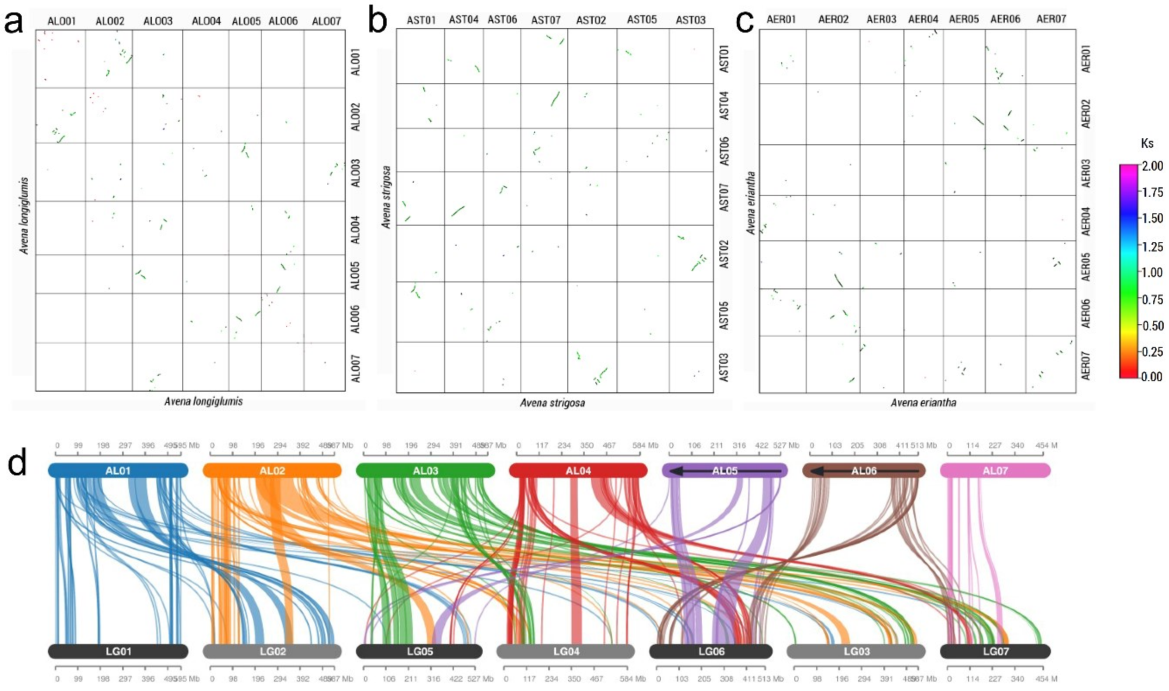

**Fig. S10.** Syntenic relationships based on three *Avena* species genomes. **a** Dotplot of ALO. **b** Dotplot of AST. **c** Dotplot of AER. **d** Syntenic analysis of the ALO genome.

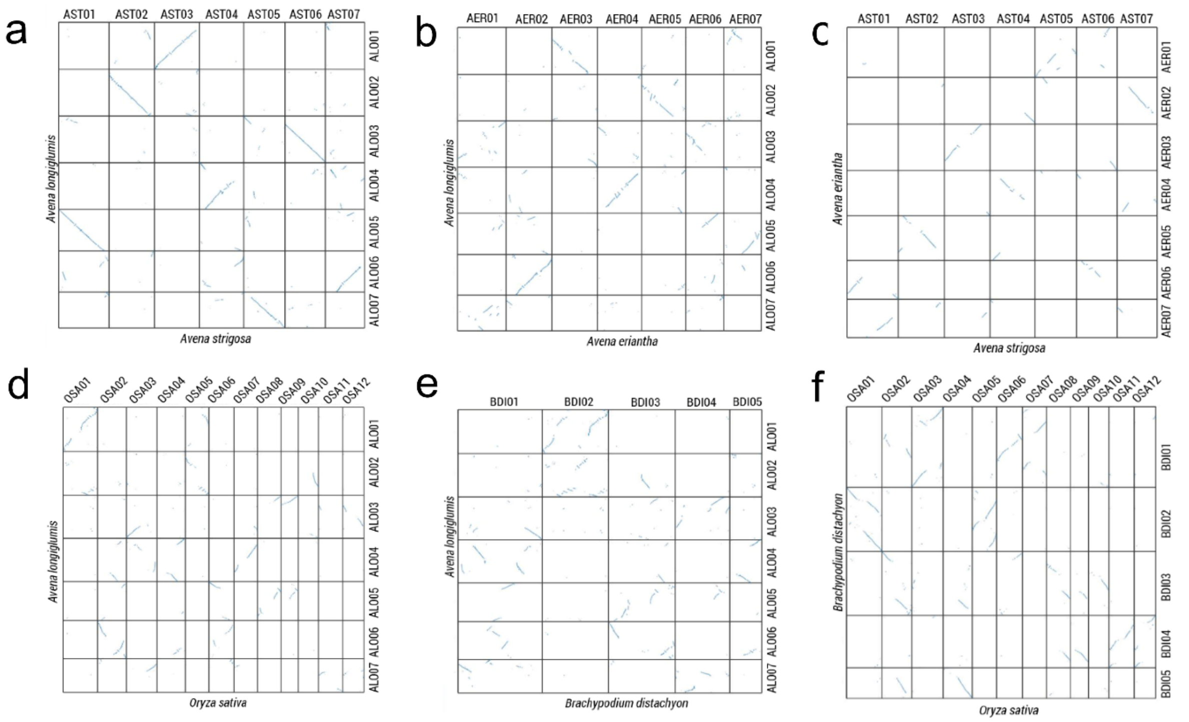

**Fig. S11.** Syntenic relationships based on AST-ALO-AER and OSA-ALO-BDI genome homologous genes. **a** Dotplot of AST-ALO genomes. **b** Dotplot of AER-ALO genomes. **c** Dotplot of AST-AER genomes. **d** Dotplot of OSA-ALO genomes. **e** Dotplot of BDI-ALO genomes. **f** Dotplot of OSA-BDI genomes.

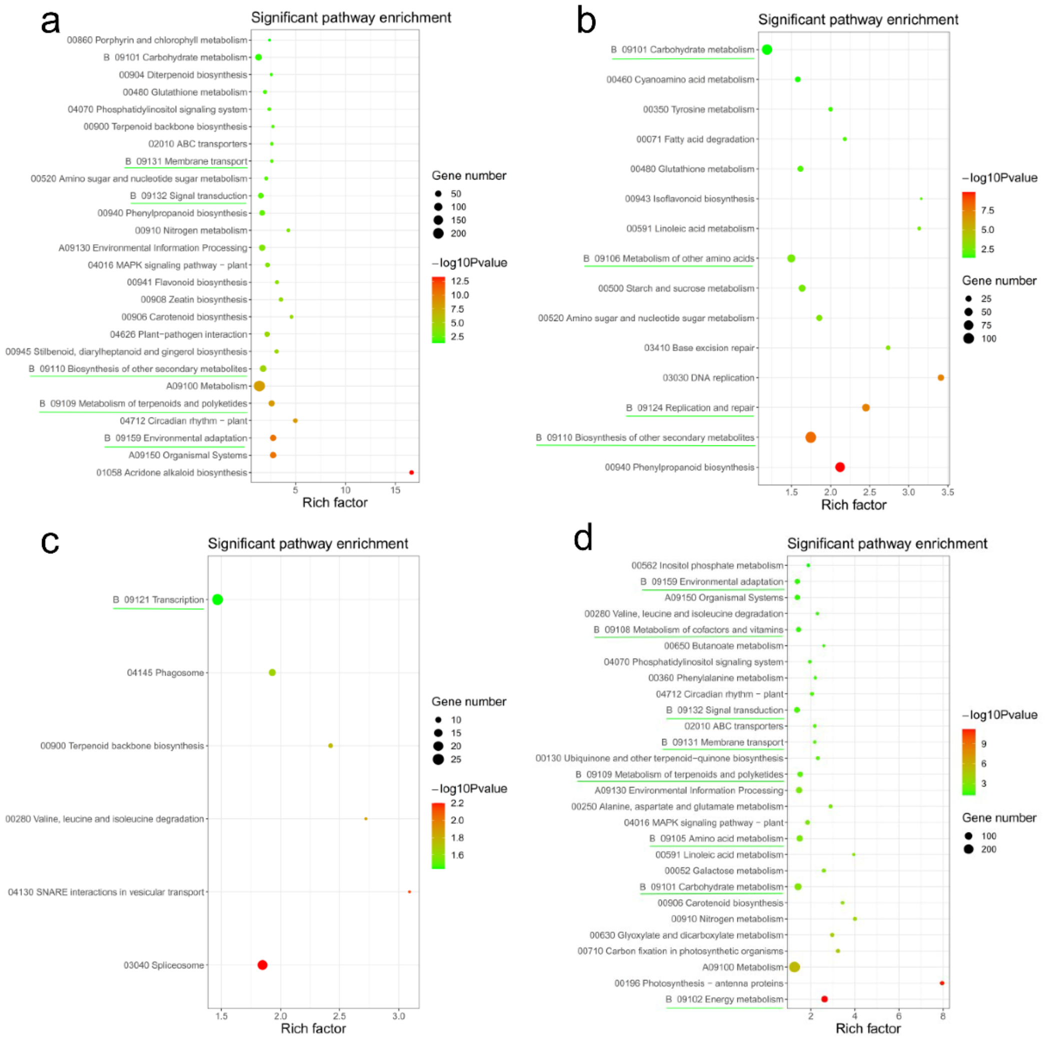

**Fig. S12.** KEGG enrichment of up- and down-regulated DEFs in roots versus salt-treated roots and leaves versus salt-treated leaves of ALO. **a** Up-regulated differentially expressed genes (DEGs) of roots versus salt-treated roots. **b** Down-regulated DEGs of of roots versus salt-treated roots. **c** Up-regulated DEGs of leaves versus salt-treated leaves. **d** Down-regulated DEGs of leaves versus salt-treated leaves.

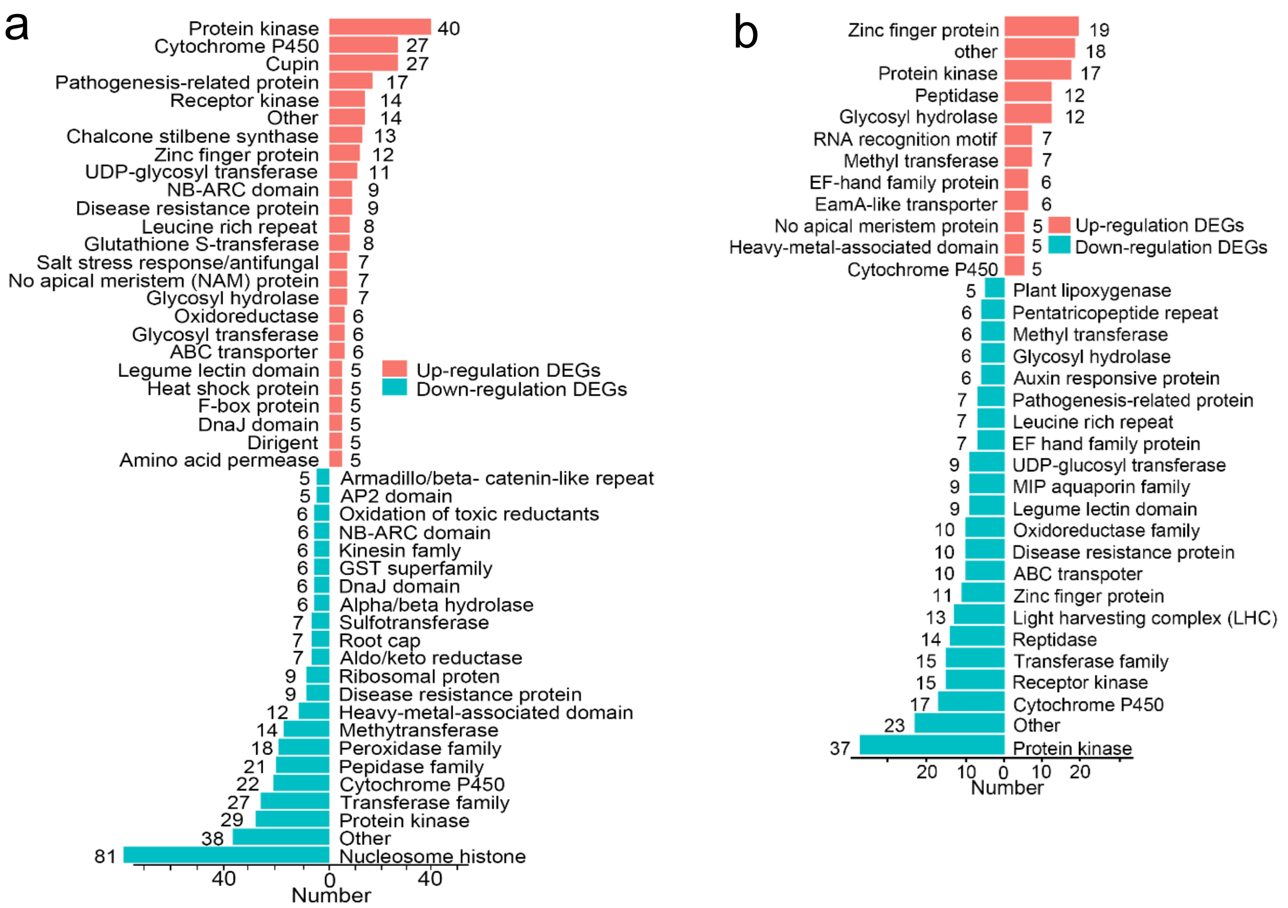

**Fig. S13.** Statistics of expanded DEGs (number ≥ 5) of ALO. **a** Expanded DEGs in roots versus salt-treated roots of ALO. **b** Expanded DEGs in leaves versus salt-treated leaves of ALO. Up-regulated and down-regulated DEG numbers denoted by red and green columns, respectively.

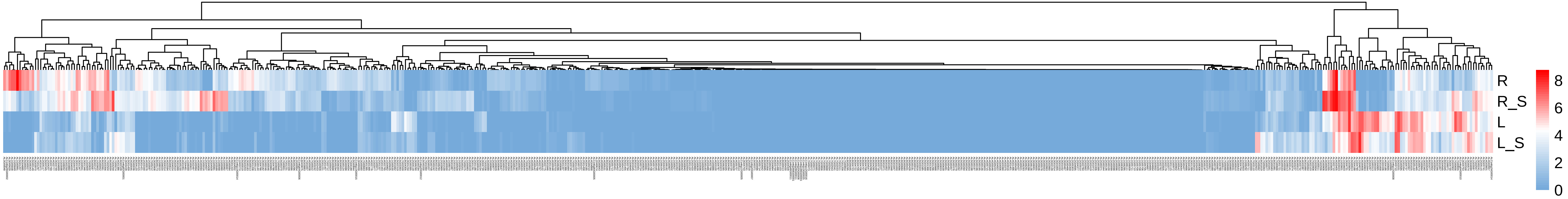

**Fig. S14.** Heat map showing hierarchical clustering of cytochrome P450 gene families in roots, salt-treated roots, leaves and salt-treated leaves of ALO. Expression values were normalized by $log_2$(FPKM + 1). Highly and weakly expressed genes were colored by red and blue boxes, respectively.

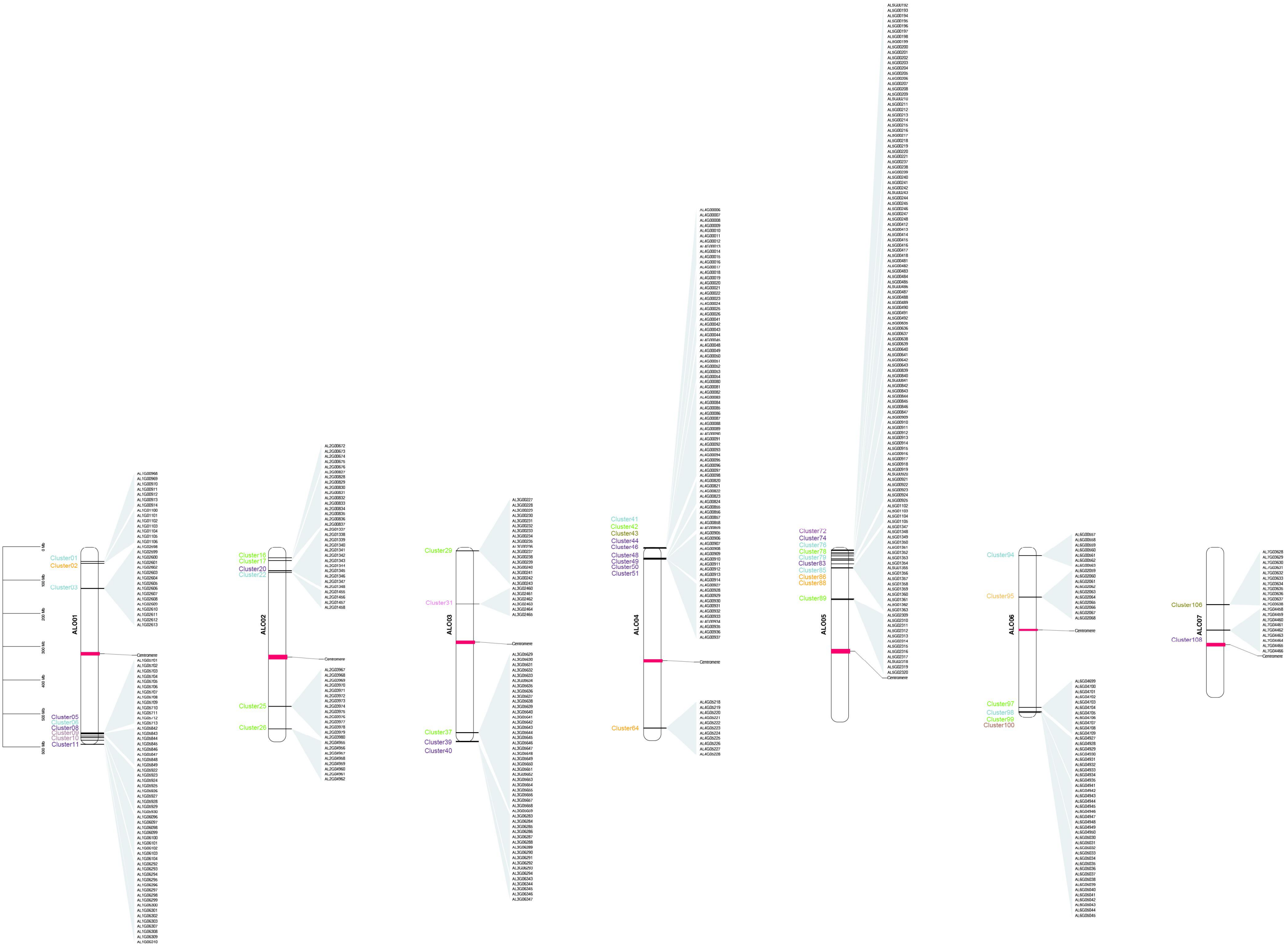

**Fig. S15.** Total 557 cytochrome P450 genes located within 57 clusters identified in the ALO genome. A scale in the left represented length of chromosome in megabases (Mbp). The cluster type labeled as the same colour as Fig 5, the position of the gene cluster (black band) scaled based on chromosome size and its physical position on the chromosome. The centromere position shown by pink band. Five key gene clusters CL10, CL37, CL95, CL98 and CL106 were identified in the ALO genome, their function characterized by UDP-glycosyl transferase (AS01G000200), serine carboxy peptidase-like acyltransferase (SAD7; AS01G000190), UDP-glycosyl transferase (AS01G000200), subtilisin homologue (AS01G000130) and O-methyltransferase (AS01G000040) together with enzymes annotated as CYP450 [7], dehydrodolichyl diphosphate synthase, aldehyde oxidase, and hydrolase proteins (Additional file 1: Figs. S16a, b, c, d, e, f and g).

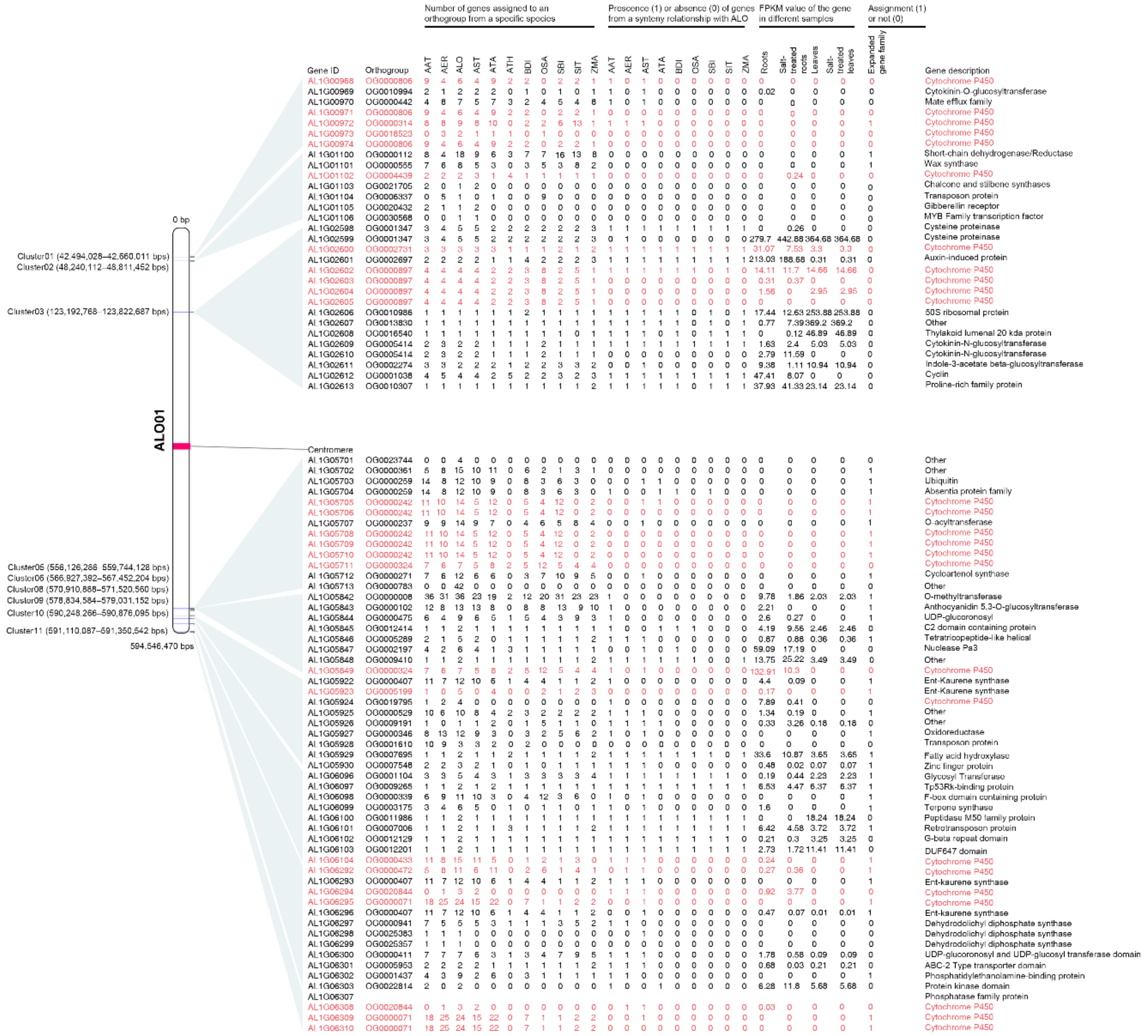

**Fig. S16.** *Cytochrome P450* genes inserted within gene-clusters identified on ALO chromosomes. We identified total 57 *cytochrome P450* gene clusters on seven chromosomes of ALO. Conserved synteny at gene clusters across 11 species indicates role of secondary metabolite clusters toward evolution of specialized metabolites in plants. The position of the gene cluster is scaled based on chromosome size and its physical position on the chromosome. FPKM values suggest role of salt stress for *cytochrome P450* gene family evolution in ALO roots and leaves. The gene description colored as pink represents genes with functions associated with sterol biosynthesis. **Fig. S16a.** *Cytochrome P450* genes inserted within gene-clusters identified on ALO01. Conserved synteny at gene clusters across 11 species indicates role of secondary metabolite clusters toward evolution of specialized metabolites in plants. The position of the gene cluster is scaled based on chromosome size and its physical position on the chromosome. FPKM values suggest role of salt stress for *cytochrome P450* gene family evolution in ALO roots and leaves. The gene description colored as pink represents genes with functions associated with sterol biosynthesis.

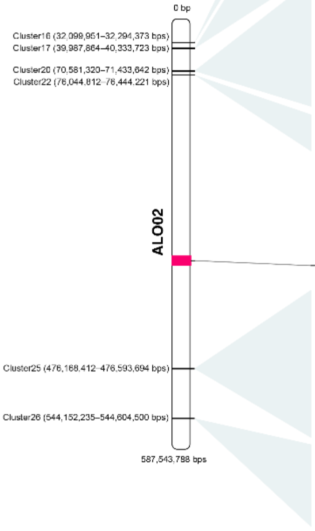

| | | Number of genes assigned to an orthogroup from a specific species | | | | | | | | | | | Presence (1) or absence (0) of genes from a synteny relationship with ALO | | | | | | | | | FPKM value of the gene in different samples | | | | Assignment (1) or not (0) | |
|---|---|---|---|---|---|---|---|---|---|---|---|---|---|---|---|---|---|---|---|---|---|---|---|---|---|---|---|
| Gene ID | Orthogroup | AAT | AER | ALO | AST | ATA | ATH | BDI | OSA | SBI | SIT | ZMA | AAT | AER | AST | ATA | BDI | OSA | SBI | SIT | ZMA | Roots | Salt-treated roots | Leaves | Salt-treated leaves | Expanded gene family | Gene description |
| AL2G00672 | OG0015299 | 1 | 1 | 2 | 1 | 1 | 0 | 1 | 1 | 1 | 1 | 1 | 0 | 0 | 0 | 0 | 0 | 0 | 0 | 0 | 0 | 1.83 | 1.28 | 1.12 | 1.12 | 0 | F-Box domain |
| AL2G00673 | OG0017289 | 1 | 1 | 3 | 1 | 0 | 0 | 1 | 1 | 1 | 1 | 0 | 0 | 0 | 0 | 0 | 0 | 0 | 0 | 0 | 0 | 0.56 | 0 | 0 | 0 | 0 | Cytochrome P450 |
| AL2G00674 | OG0017289 | 1 | 1 | 3 | 1 | 0 | 0 | 1 | 1 | 1 | 1 | 0 | 0 | 0 | 0 | 0 | 0 | 0 | 0 | 0 | 0 | 1.36 | 0 | 0 | 0 | 0 | Cytochrome P450 |
| AL2G00675 | OG0002353 | 2 | 2 | 4 | 1 | 1 | 4 | 2 | 3 | 2 | 2 | 2 | 1 | 1 | 0 | 0 | 0 | 0 | 0 | 0 | 0 | 0 | 0 | 0 | 0 | 1 | Gibberellin 3-beta-dioxygenase |
| AL2G00676 | OG0002353 | 2 | 2 | 4 | 1 | 1 | 4 | 2 | 3 | 2 | 2 | 2 | 0 | 0 | 0 | 0 | 0 | 0 | 0 | 0 | 0 | 0 | 0 | 0 | 0 | 0 | Gibberellin 3-beta-dioxygenase |
| AL2G00827 | OG0016400 | 1 | 1 | 2 | 1 | 1 | 0 | 1 | 1 | 1 | 1 | 0 | 1 | 1 | 1 | 1 | 1 | 1 | 1 | 1 | 0 | 1.11 | 0.26 | 0.02 | 0.02 | 0 | Helicase |
| AL2G00828 | OG0000430 | 0 | 1 | 44 | 12 | 0 | 0 | 0 | 0 | 0 | 1 | 0 | 0 | 0 | 0 | 0 | 0 | 0 | 0 | 0 | 0 | 0 | 0.12 | 0 | 0 | 1 | Other |
| AL2G00829 | OG0000071 | 18 | 25 | 24 | 15 | 22 | 0 | 7 | 1 | 1 | 2 | 2 | 1 | 0 | 1 | 0 | 0 | 0 | 0 | 0 | 0 | 2.4 | 0.02 | 0 | 0 | 1 | Cytochrome P450 |
| AL2G00830 | OG0002252 | 2 | 2 | 3 | 2 | 3 | 1 | 2 | 2 | 2 | 2 | 4 | 1 | 1 | 1 | 1 | 1 | 1 | 1 | 1 | 1 | 4.11 | 2.27 | 0.9 | 0.9 | 1 | Glycosyl transferase |
| AL2G00831 | OG0001068 | 6 | 4 | 6 | 7 | 1 | 0 | 4 | 2 | 3 | 3 | 0 | 1 | 0 | 1 | 0 | 0 | 0 | 0 | 0 | 0 | 0 | 0 | 0 | 0 | 0 | Carboxylate xxidase homolog 1 |
| AL2G00832 | OG0004542 | 8 | 1 | 2 | 1 | 3 | 0 | 1 | 1 | 1 | 1 | 0 | 1 | 1 | 1 | 1 | 0 | 0 | 1 | 1 | 0 | 9.66 | 0 | 0 | 0 | 0 | Proline-rich protein |
| AL2G00833 | | | | | | | | | | | | | 1 | 1 | 1 | 1 | 0 | 0 | 0 | 0 | 0 | 35.71 | 22.23 | 10.34 | 10.34 | 0 | CESA8 |
| AL2G00834 | OG0002979 | 2 | 2 | 3 | 2 | 2 | 1 | 2 | 2 | 2 | 2 | 3 | 0 | 0 | 0 | 0 | 0 | 0 | 0 | 0 | 0 | 92.21 | 34.75 | 16.33 | 16.33 | 1 | CESA2 |
| AL2G00835 | OG0011192 | 1 | 1 | 2 | 1 | 1 | 0 | 1 | 1 | 1 | 1 | 2 | 1 | 1 | 1 | 0 | 0 | 0 | 0 | 0 | 0 | 0 | 0.18 | 0 | 0 | 0 | Protease inhibitor/Seed Storage |
| AL2G00836 | OG0014283 | 1 | 1 | 2 | 1 | 0 | 1 | 1 | 1 | 1 | 1 | 1 | 1 | 1 | 1 | 0 | 0 | 0 | 0 | 0 | 0 | 5.59 | 5.13 | 34.75 | 34.75 | 0 | Fruit protein |
| AL2G00837 | OG0001684 | 2 | 2 | 3 | 2 | 1 | 1 | 2 | 3 | 3 | 2 | 8 | 1 | 1 | 1 | 1 | 0 | 0 | 1 | 1 | 1 | 32.14 | 24.83 | 19 | 19 | 1 | Zinc finger protein |
| AL2G01337 | OG0000314 | 8 | 8 | 9 | 8 | 10 | 0 | 2 | 2 | 6 | 13 | 1 | 0 | 1 | 0 | 0 | 0 | 0 | 0 | 0 | 0 | 5.33 | 0.69 | 0.01 | 0.01 | 1 | Cytochrome P450 |
| AL2G01338 | OG0018863 | 3 | 2 | 1 | 2 | 0 | 0 | 0 | 0 | 0 | 0 | 0 | 1 | 1 | 0 | 0 | 0 | 0 | 0 | 0 | 0 | 2.36 | 0.36 | 0 | 0 | 0 | Terpene synthase |
| AL2G01339 | OG0002074 | 3 | 2 | 9 | 4 | 2 | 2 | 1 | 1 | 0 | 1 | 1 | 0 | 0 | 0 | 0 | 0 | 0 | 0 | 0 | 0 | 0 | 0 | 0 | 0 | 1 | Tigr01627 family protein |
| AL2G01340 | OG0007404 | 1 | 5 | 5 | 0 | 2 | 0 | 0 | 0 | 0 | 1 | 0 | 0 | 0 | 0 | 0 | 0 | 0 | 0 | 0 | 0 | 0.09 | 0.13 | 0.09 | 0.09 | 1 | Zinc finger protein |
| AL2G01341 | OG0007404 | 1 | 5 | 5 | 0 | 2 | 0 | 0 | 0 | 0 | 1 | 0 | 0 | 0 | 0 | 0 | 0 | 0 | 0 | 0 | 0 | 0.07 | 0.07 | 0.23 | 0.23 | 1 | Retrotransposon protein |
| AL2G01342 | OG0002074 | 3 | 2 | 9 | 4 | 2 | 2 | 1 | 1 | 0 | 1 | 1 | 0 | 0 | 0 | 0 | 0 | 0 | 0 | 0 | 0 | 3.53 | 0.5 | 0.04 | 0.04 | 1 | Tigr01627 family protein |
| AL2G01343 | OG0000314 | 8 | 8 | 9 | 8 | 10 | 0 | 2 | 2 | 6 | 13 | 1 | 0 | 0 | 0 | 0 | 0 | 0 | 0 | 0 | 0 | 5.22 | 0.76 | 0.01 | 0.01 | 1 | Cytochrome P450 |
| AL2G01344 | OG0025998 | 0 | 1 | 2 | 0 | 0 | 0 | 0 | 0 | 0 | 0 | 0 | 0 | 0 | 0 | 0 | 0 | 0 | 0 | 0 | 0 | 2.14 | 0 | 0 | 0 | 0 | Terpene synthase |
| AL2G01345 | OG0000182 | 14 | 12 | 16 | 10 | 10 | 0 | 5 | 5 | 5 | 4 | 2 | 1 | 1 | 1 | 0 | 0 | 0 | 0 | 0 | 0 | 0 | 0 | 0.05 | 0.05 | 1 | Methyltransferase |
| AL2G01346 | OG0002074 | 3 | 2 | 9 | 4 | 2 | 2 | 1 | 1 | 0 | 1 | 1 | 0 | 0 | 0 | 0 | 0 | 0 | 0 | 0 | 0 | 2.57 | 0.25 | 0 | 0 | 1 | Tigr01627 family protein |
| AL2G01347 | OG0030655 | 0 | 0 | 2 | 0 | 0 | 0 | 0 | 0 | 0 | 0 | 0 | 0 | 0 | 0 | 0 | 0 | 0 | 0 | 0 | 0 | 0 | 0 | 0 | 0 | 0 | Methyltransferase |
| AL2G01348 | OG0000113 | 20 | 17 | 20 | 11 | 1 | 4 | 7 | 15 | 1 | 2 | 1 | 0 | 1 | 1 | 0 | 0 | 0 | 0 | 0 | 0 | 0 | 0 | 0 | 0 | 1 | Glutelin |
| AL2G01455 | OG0025422 | 1 | 0 | 1 | 1 | 0 | 0 | 0 | 0 | 0 | 0 | 0 | 1 | 0 | 0 | 0 | 0 | 0 | 0 | 0 | 0 | 11.4 | 1.38 | 1.43 | 1.43 | 0 | UDP-glucosyl transferase |
| AL2G01456 | OG0025458 | 1 | 0 | 1 | 1 | 0 | 0 | 0 | 0 | 0 | 0 | 0 | 1 | 0 | 0 | 0 | 0 | 0 | 0 | 0 | 0 | 6.71 | 0.49 | 19.68 | 19.68 | 0 | Cytokinin-O-glucosyltransferase |
| AL2G01457 | OG0019730 | 1 | 3 | 2 | 1 | 0 | 0 | 0 | 0 | 0 | 0 | 0 | 1 | 0 | 0 | 0 | 0 | 0 | 0 | 0 | 0 | 2.82 | 0.02 | 1.04 | 1.04 | 0 | Cytochrome P450 |
| AL2G01458 | OG0001010 | 2 | 4 | 9 | 3 | 4 | 0 | 2 | 4 | 5 | 1 | 3 | 1 | 0 | 0 | 0 | 0 | 0 | 0 | 0 | 0 | 0.05 | 0 | 0 | 0 | 1 | Cytochrome P450 |
| Gene ID | Orthogroup | | | | | | | | | | | | | | | | | | | | | | | | | | |
| AL2G03967 | OG0018505 | 0 | 1 | 1 | 0 | 1 | 1 | 1 | 1 | 1 | 1 | 1 | 1 | 1 | 1 | 1 | 1 | 1 | 1 | 1 | 1 | 153.33 | 84.22 | 78.06 | 78.06 | 0 | NADH dehydrogenase |
| AL2G03968 | OG0002782 | 2 | 2 | 2 | 2 | 2 | 3 | 2 | 2 | 2 | 2 | 2 | 0 | 1 | 0 | 1 | 1 | 1 | 1 | 1 | 0 | 45.91 | 51.99 | 33.06 | 33.06 | 0 | Transporter family protein |
| AL2G03969 | OG0016571 | 1 | 1 | 1 | 1 | 0 | 1 | 1 | 1 | 1 | 1 | 1 | 1 | 1 | 1 | 0 | 0 | 0 | 0 | 0 | 0 | 1.36 | 3.37 | 0.1 | 0.1 | 0 | Cytochrome P450 |
| AL2G03970 | OG0000982 | 4 | 3 | 4 | 3 | 3 | 1 | 3 | 3 | 6 | 3 | 4 | 1 | 1 | 1 | 0 | 0 | 0 | 0 | 0 | 0 | 27.74 | 16.26 | 2.65 | 2.65 | 0 | Fad dependent oxidoreductase |
| AL2G03971 | OG0014018 | 1 | 1 | 1 | 1 | 1 | 1 | 1 | 1 | 1 | 1 | 1 | 1 | 1 | 1 | 1 | 1 | 1 | 1 | 1 | 1 | 15.47 | 29.33 | 2.07 | 2.07 | 0 | Dehydration protein |
| AL2G03972 | OG0008930 | 1 | 1 | 1 | 1 | 1 | 1 | 1 | 2 | 2 | 1 | 1 | 1 | 1 | 1 | 1 | 1 | 1 | 1 | 1 | 1 | 2.67 | 1.77 | 1.28 | 1.28 | 0 | AMP-binding enzyme |
| AL2G03973 | OG0001379 | 2 | 3 | 4 | 2 | 2 | 2 | 3 | 3 | 3 | 3 | 5 | 1 | 1 | 1 | 0 | 1 | 1 | 1 | 1 | 1 | 135.75 | 162.29 | 77.23 | 77.23 | 1 | Cytidylyltransferase domain |
| AL2G03974 | OG0006807 | 2 | 1 | 2 | 1 | 1 | 1 | 1 | 1 | 2 | 1 | 2 | 1 | 1 | 1 | 1 | 1 | 1 | 1 | 1 | 0 | 1.14 | 2.13 | 104.58 | 104.58 | 0 | Lrgb-like family protein |
| AL2G03975 | OG0006807 | 2 | 1 | 2 | 1 | 1 | 1 | 1 | 1 | 2 | 1 | 2 | 0 | 0 | 0 | 0 | 0 | 0 | 0 | 0 | 0 | 5.97 | 1.03 | 0 | 0 | 0 | Lrgb-like family protein |
| AL2G03976 | OG0001227 | 5 | 4 | 6 | 3 | 0 | 0 | 3 | 1 | 4 | 3 | 5 | 1 | 1 | 0 | 0 | 0 | 0 | 0 | 0 | 0 | 1.14 | 2.58 | 0 | 0 | 1 | Plastocyanin-like domain |
| AL2G03977 | OG0014676 | 1 | 1 | 1 | 1 | 1 | 1 | 1 | 1 | 1 | 1 | 1 | 1 | 1 | 1 | 1 | 1 | 1 | 0 | 1 | 1 | 13.39 | 7.59 | 1.12 | 1.12 | 0 | Other |
| AL2G03978 | OG0003896 | 2 | 2 | 2 | 2 | 2 | 2 | 2 | 2 | 1 | 2 | 2 | 1 | 1 | 1 | 1 | 1 | 0 | 1 | 1 | 1 | 73.78 | 59.51 | 26.44 | 26.44 | 0 | Proteasome subunit |
| AL2G03979 | OG0011758 | 1 | 1 | 1 | 1 | 1 | 1 | 1 | 1 | 1 | 2 | 1 | 1 | 1 | 1 | 0 | 1 | 1 | 1 | 1 | 1 | 0.7 | 0.96 | 0.75 | 0.75 | 0 | PPR Repeat Domain |
| AL2G03980 | OG0001194 | 3 | 3 | 4 | 3 | 3 | 3 | 3 | 3 | 3 | 3 | 3 | 1 | 0 | 1 | 1 | 1 | 1 | 1 | 1 | 1 | 0.69 | 1.52 | 0.75 | 0.75 | 1 | CslD1 |
| AL2G04955 | OG0000008 | 36 | 31 | 36 | 23 | 19 | 2 | 12 | 20 | 31 | 23 | 23 | 0 | 1 | 1 | 1 | 1 | 1 | 1 | 1 | 1 | 21.79 | 6.17 | 0 | 0 | 1 | O-Methyltransferase |
| AL2G04956 | OG0000008 | 36 | 31 | 36 | 23 | 19 | 2 | 12 | 20 | 31 | 23 | 23 | 0 | 0 | 0 | 0 | 0 | 0 | 0 | 0 | 0 | 78.12 | 0.57 | 0 | 0 | 1 | O-Methyltransferase |
| AL2G04957 | OG0005889 | 2 | 0 | 2 | 1 | 1 | 3 | 1 | 1 | 1 | 2 | 2 | 1 | 0 | 0 | 0 | 0 | 0 | 0 | 0 | 0 | 38.99 | 25.86 | 16.21 | 16.21 | 0 | 60S ribosomal protein |
| AL2G04958 | OG0004928 | 1 | 2 | 2 | 0 | 2 | 2 | 2 | 1 | 2 | 2 | 2 | 1 | 1 | 0 | 0 | 0 | 0 | 0 | 0 | 0 | 7.86 | 3.68 | 1.58 | 1.58 | 0 | Other |
| AL2G04959 | OG0001852 | 4 | 3 | 4 | 3 | 1 | 1 | 4 | 2 | 2 | 2 | 1 | 1 | 1 | 0 | 0 | 0 | 0 | 0 | 0 | 0 | 0 | 1.06 | 0.06 | 0.06 | 0 | Mitosis pProtein dim1 |
| AL2G04960 | OG0004038 | 2 | 2 | 2 | 1 | 1 | 2 | 1 | 2 | 2 | 2 | 3 | 1 | 1 | 0 | 0 | 0 | 0 | 0 | 0 | 0 | 110.7 | 15.52 | 0.44 | 0.44 | 0 | Strubbelig-receptor family |
| AL2G04961 | OG0000008 | 36 | 31 | 36 | 23 | 19 | 2 | 12 | 20 | 31 | 23 | 23 | 0 | 0 | 0 | 0 | 1 | 0 | 0 | 0 | 0 | 52.08 | 20.45 | 0 | 0 | 1 | O-Methyltransferase |
| AL2G04962 | OG0015935 | 1 | 1 | 1 | 1 | 1 | 0 | 1 | 1 | 1 | 1 | 1 | 1 | 1 | 1 | 1 | 1 | 1 | 1 | 1 | 1 | 2.68 | 2.31 | 1.36 | 1.36 | 0 | Auxin response factor 14 |

**Fig. S16b.** *Cytochrome P450* genes inserted within gene-clusters identified on ALO02.

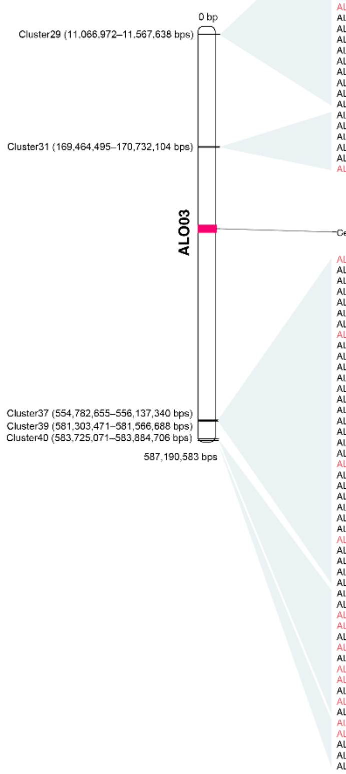

| | | Number of genes assigned to an orthogroup from a specific species | | | | | | | | | | | Presence (1) or absence (0) of genes from a synteny relationship with ALO | | | | | | | | | FPKM value of the gene in different samples | | | | Assignment (1) or not (0) | |
|---|---|---|---|---|---|---|---|---|---|---|---|---|---|---|---|---|---|---|---|---|---|---|---|---|---|---|---|
| Gene ID | Othogroup | AAT | AER | ALO | AST | ATA | ATH | BDI | OSA | SBI | SIT | ZMA | AAT | AER | AST | ATA | BDI | OSA | SBI | SIT | ZMA | Roots | Salt treated-roots | Leaves | Salt treated-leaves | Expanded gene family | Gene description |
| AL3G00227 | OG0007806 | 2 | 0 | 3 | 2 | 1 | 1 | 1 | 1 | 1 | 1 | 1 | 1 | 0 | 1 | 0 | 0 | 0 | 0 | 0 | 0 | 0 | 0 | 0 | 0 | 1 | Other |
| AL3G00228 | OG0001875 | 3 | 7 | 2 | 2 | 4 | 0 | 2 | 4 | 1 | 2 | 0 | 1 | 1 | 1 | 1 | 1 | 0 | 0 | 1 | 0 | 0.71 | 0.71 | 0.62 | 0.62 | 0 | Male sterility protein |
| AL3G00229 | OG0005987 | 2 | 2 | 3 | 3 | 3 | 0 | 3 | 0 | 0 | 0 | 0 | 1 | 1 | 1 | 1 | 0 | 0 | 0 | 0 | 0 | 0.1 | 0 | 0 | 0 | 0 | Hydrolase |
| AL3G00230 | OG0001008 | 4 | 6 | 6 | 5 | 3 | 2 | 2 | 1 | 1 | 4 | 3 | 1 | 1 | 1 | 1 | 1 | 0 | 0 | 0 | 0 | 0 | 0 | 0 | 0 | 1 | Transferase family protein |
| AL3G00231 | OG0030709 | 0 | 0 | 2 | 0 | 0 | 0 | 0 | 0 | 0 | 0 | 0 | 0 | 0 | 0 | 0 | 0 | 0 | 0 | 0 | 0 | 0 | 0 | 0 | 0 | 0 | Transferase family protein |
| AL3G00232 | OG0023305 | 1 | 1 | 1 | 1 | 0 | 0 | 0 | 0 | 0 | 0 | 0 | 0 | 0 | 0 | 0 | 0 | 0 | 0 | 0 | 0 | 0.1 | 0 | 0.37 | 0.37 | 0 | Hydrolase |
| AL3G00233 | OG0000783 | 0 | 0 | 42 | 0 | 0 | 0 | 0 | 0 | 0 | 0 | 0 | 0 | 0 | 0 | 0 | 0 | 0 | 0 | 0 | 0 | 0 | 0 | 0 | 0 | 0 | Retrotransposon protein |
| AL3G00234 | OG0025586 | 1 | 0 | 1 | 1 | 0 | 0 | 0 | 0 | 0 | 0 | 0 | 1 | 0 | 1 | 0 | 0 | 0 | 0 | 0 | 0 | 0.27 | 0.05 | 2.3 | 2.3 | 0 | Cytochrome P450 |
| AL3G00235 | OG0030709 | 0 | 0 | 2 | 0 | 0 | 0 | 0 | 0 | 0 | 0 | 0 | 0 | 0 | 0 | 0 | 0 | 0 | 0 | 0 | 0 | 0 | 0 | 0 | 0 | 0 | Transferase family protein |
| AL3G00236 | OG0004496 | 1 | 4 | 5 | 1 | 6 | 0 | 0 | 1 | 1 | 0 | 0 | 0 | 1 | 1 | 1 | 1 | 0 | 1 | 1 | 0 | 0 | 0.19 | 0 | 0 | 1 | Other |
| AL3G00237 | OG0004496 | 1 | 4 | 5 | 1 | 6 | 0 | 0 | 1 | 1 | 0 | 0 | 1 | 0 | 0 | 0 | 0 | 0 | 0 | 0 | 0 | 0.38 | 0.4 | 0.11 | 0.11 | 1 | Other |
| AL3G00238 | OG0008216 | 2 | 1 | 1 | 1 | 1 | 1 | 2 | 1 | 1 | 1 | 1 | 1 | 1 | 1 | 1 | 1 | 0 | 1 | 1 | 1 | 17.91 | 20.14 | 13.16 | 13.16 | 0 | Other |
| AL3G00239 | OG0001932 | 2 | 2 | 2 | 2 | 4 | 4 | 1 | 1 | 5 | 3 | 1 | 1 | 1 | 1 | 1 | 1 | 1 | 1 | 1 | 0 | 0 | 0 | 0 | 0 | 0 | Other |
| AL3G00240 | OG0005392 | 2 | 2 | 2 | 2 | 1 | 0 | 2 | 1 | 1 | 2 | 2 | 1 | 1 | 1 | 1 | 0 | 0 | 0 | 0 | 0 | 27.21 | 47.95 | 24.01 | 24.01 | 0 | Pattern formation protein |
| AL3G00241 | OG0002860 | 2 | 2 | 2 | 2 | 2 | 2 | 2 | 2 | 2 | 2 | 3 | 1 | 1 | 1 | 1 | 1 | 1 | 1 | 1 | 1 | 22.11 | 26.79 | 19.22 | 19.22 | 0 | NADPH reductase |
| AL3G00242 | OG0004488 | 2 | 3 | 2 | 2 | 4 | 0 | 1 | 1 | 1 | 2 | 1 | 1 | 1 | 1 | 1 | 1 | 1 | 1 | 1 | 1 | 0.33 | 0.14 | 0 | 0 | 0 | Zinc finger protein |
| AL3G00243 | OG0004488 | 2 | 3 | 2 | 2 | 4 | 0 | 1 | 1 | 1 | 2 | 1 | 0 | 0 | 0 | 0 | 0 | 0 | 0 | 0 | 0 | 0 | 0.62 | 0 | 0 | 0 | Zinc finger protein |
| AL3G02460 | OG0007005 | 1 | 1 | 1 | 1 | 1 | 0 | 1 | 2 | 2 | 2 | 3 | 1 | 1 | 1 | 1 | 1 | 1 | 1 | 1 | 1 | 0 | 0.1 | 1.06 | 1.06 | 0 | Broad complex BTB domain |
| AL3G02461 | OG0007907 | 1 | 1 | 1 | 1 | 0 | 0 | 1 | 2 | 2 | 4 | 1 | 1 | 1 | 1 | 0 | 1 | 1 | 1 | 1 | 1 | 0 | 0.05 | 1.73 | 1.73 | 0 | Strictosidine synthase |
| AL3G02462 | OG0006244 | 1 | 1 | 1 | 1 | 1 | 2 | 1 | 2 | 2 | 2 | 2 | 1 | 1 | 1 | 1 | 1 | 1 | 1 | 1 | 1 | 94.13 | 107.68 | 42.54 | 42.54 | 0 | 2-Isopropylmalate synthase |
| AL3G02463 | OG0000443 | 2 | 5 | 5 | 2 | 7 | 0 | 10 | 3 | 8 | 9 | 6 | 1 | 1 | 1 | 1 | 1 | 1 | 1 | 1 | 0 | 0 | 0 | 0 | 0 | 0 | Zinc finger protein |
| AL3G02464 | OG0000276 | 5 | 4 | 8 | 10 | 11 | 0 | 4 | 14 | 4 | 9 | 2 | 1 | 0 | 1 | 0 | 0 | 0 | 0 | 0 | 0 | 0 | 0.09 | 0 | 0 | 0 | Other |
| AL3G02465 | OG0007705 | 1 | 1 | 1 | 1 | 1 | 1 | 1 | 2 | 1 | 2 | 2 | 1 | 1 | 1 | 1 | 1 | 1 | 1 | 1 | 1 | 8.87 | 14.67 | 68.42 | 68.42 | 0 | Cytochrome P450 |
| AL3G05629 | OG0015906 | 1 | 1 | 2 | 1 | 1 | 0 | 0 | 1 | 1 | 1 | 1 | 1 | 1 | 1 | 1 | 1 | 1 | 1 | 1 | 1 | 111.34 | 21.06 | 0.32 | 0.32 | 0 | Cytochrome P450 |
| AL3G05630 | OG0007084 | 1 | 1 | 2 | 2 | 1 | 2 | 1 | 1 | 1 | 2 | 1 | 1 | 1 | 1 | 1 | 1 | 1 | 0 | 1 | 0 | 18.42 | 9.83 | 19.9 | 19.9 | 0 | ATP phosphoribosyltransferase |
| AL3G05631 | OG0009696 | 1 | 1 | 2 | 1 | 1 | 2 | 1 | 1 | 1 | 1 | 1 | 1 | 1 | 1 | 1 | 1 | 1 | 1 | 1 | 1 | 4.16 | 3.38 | 14.73 | 14.73 | 1 | Serine acetyltransferase |
| AL3G05632 | OG0000682 | 3 | 3 | 11 | 3 | 1 | 4 | 3 | 6 | 3 | 4 | 4 | 1 | 1 | 1 | 0 | 0 | 0 | 0 | 0 | 0 | 3.08 | 6.8 | 2.72 | 2.72 | 1 | Aldehyde oxidase |
| AL3G05633 | OG0000682 | 3 | 3 | 11 | 3 | 1 | 4 | 3 | 6 | 3 | 4 | 4 | 0 | 0 | 0 | 0 | 0 | 0 | 0 | 0 | 0 | 6.17 | 17.17 | 7.8 | 7.8 | 1 | Aldehyde oxidase |
| AL3G05634 | OG0000265 | 6 | 5 | 14 | 6 | 7 | 1 | 3 | 7 | 11 | 8 | 4 | 1 | 1 | 1 | 0 | 0 | 0 | 0 | 0 | 0 | 25.15 | 1.32 | 0.15 | 0.15 | 1 | Leucoanthocyanidin reductase |
| AL3G05635 | OG0000762 | 6 | 4 | 7 | 6 | 5 | 2 | 3 | 1 | 3 | 3 | 2 | 1 | 0 | 1 | 0 | 0 | 0 | 0 | 0 | 0 | 4.22 | 4.74 | 17.42 | 17.42 | 1 | Mate efflux family protein |
| AL3G05636 | OG0020698 | 1 | 2 | 2 | 1 | 0 | 0 | 0 | 0 | 0 | 0 | 0 | 1 | 1 | 1 | 0 | 0 | 0 | 0 | 0 | 0 | 0 | 0 | 0 | 0 | 0 | Cytochrome P450 |
| AL3G05637 | OG0000963 | 3 | 3 | 8 | 5 | 1 | 2 | 2 | 4 | 4 | 2 | 4 | 1 | 0 | 1 | 0 | 0 | 0 | 0 | 0 | 0 | 0 | 0 | 0 | 0 | 1 | Serine palmitoyltransferase |
| AL3G05638 | OG0001466 | 2 | 2 | 5 | 3 | 3 | 2 | 4 | 2 | 3 | 3 | 2 | 1 | 1 | 1 | 1 | 1 | 1 | 1 | 1 | 1 | 1.59 | 3.04 | 0 | 0 | 1 | AMP-binding enzyme |
| AL3G05639 | OG0001466 | 2 | 2 | 5 | 3 | 3 | 2 | 4 | 2 | 3 | 3 | 2 | 0 | 0 | 0 | 0 | 0 | 0 | 0 | 0 | 0 | 2.72 | 3.41 | 2.29 | 2.29 | 1 | AMP-binding enzyme |
| AL3G05640 | OG0006492 | 2 | 1 | 2 | 2 | 1 | 1 | 1 | 1 | 1 | 1 | 2 | 1 | 0 | 1 | 0 | 0 | 0 | 0 | 0 | 0 | 3.34 | 5.81 | 0 | 0 | 0 | Hydrolase |
| AL3G05641 | OG0009175 | 1 | 1 | 2 | 1 | 1 | 0 | 1 | 2 | 2 | 1 | 1 | 1 | 1 | 1 | 0 | 1 | 0 | 0 | 1 | 1 | 47.06 | 47.49 | 26.96 | 26.96 | 1 | Anchored protein precursor |
| AL3G05642 | OG0009063 | 1 | 1 | 2 | 1 | 1 | 1 | 1 | 1 | 1 | 1 | 2 | 1 | 1 | 1 | 1 | 1 | 1 | 0 | 0 | 1 | 9.42 | 1.34 | 13.53 | 13.53 | 1 | Other |
| AL3G05643 | OG0001023 | 1 | 26 | 10 | 0 | 0 | 0 | 0 | 0 | 0 | 0 | 0 | 0 | 0 | 0 | 0 | 0 | 0 | 0 | 0 | 0 | 0 | 0 | 0 | 0 | 1 | Other |
| AL3G05644 | | | | | | | | | | | | | 1 | 1 | 1 | 1 | 1 | 0 | 1 | 0 | 1 | 56.3 | 1.1 | 0 | 0 | 0 | Glutathione S-transferase |
| AL3G05645 | OG0012871 | 1 | 1 | 2 | 1 | 1 | 0 | 1 | 1 | 1 | 1 | 1 | 0 | 0 | 0 | 0 | 0 | 0 | 0 | 0 | 1 | 25.23 | 0.49 | 0 | 0 | 0 | Glutathione S-transferase |
| AL3G05646 | OG0008263 | 1 | 1 | 2 | 1 | 3 | 0 | 4 | 1 | 0 | 0 | 0 | 1 | 1 | 1 | 1 | 0 | 0 | 0 | 0 | 0 | 0 | 0 | 0 | 0 | 0 | Transferase family protein |
| AL3G05647 | OG0008215 | 1 | 1 | 2 | 1 | 1 | 1 | 1 | 1 | 1 | 2 | 1 | 1 | 1 | 1 | 1 | 1 | 0 | 1 | 1 | 0 | 0 | 0 | 0 | 0 | 0 | Calmodulin-binding motif family protein |
| AL3G05648 | OG0015906 | 1 | 1 | 2 | 1 | 1 | 0 | 0 | 1 | 1 | 1 | 1 | 1 | 1 | 1 | 1 | 1 | 1 | 1 | 1 | 1 | 70.73 | 10.15 | 0.18 | 0.18 | 0 | Cytochrome P450 |
| AL3G05649 | OG0007084 | 1 | 1 | 2 | 2 | 1 | 2 | 1 | 1 | 1 | 2 | 1 | 1 | 1 | 1 | 1 | 1 | 1 | 0 | 1 | 0 | 15.68 | 7.05 | 19.97 | 19.97 | 0 | ATP phosphoribosyltransferase |
| AL3G05650 | OG0009696 | 1 | 1 | 2 | 1 | 1 | 2 | 1 | 1 | 1 | 1 | 1 | 1 | 1 | 1 | 1 | 1 | 1 | 1 | 1 | 1 | 3.61 | 4.03 | 18.83 | 18.83 | 0 | Serine acetyltransferase protein |
| AL3G05651 | OG0000682 | 3 | 3 | 11 | 3 | 1 | 4 | 3 | 6 | 3 | 4 | 4 | 1 | 1 | 1 | 0 | 0 | 0 | 0 | 0 | 0 | 14.7 | 33.67 | 12.9 | 12.9 | 1 | Aldehyde oxidase |
| AL3G05652 | OG0000265 | 6 | 5 | 14 | 6 | 7 | 1 | 3 | 7 | 11 | 8 | 4 | 1 | 1 | 1 | 0 | 0 | 0 | 0 | 0 | 0 | 45.83 | 1.39 | 0.13 | 0.13 | 1 | Leucoanthocyanidin reductase |
| AL3G05653 | OG0000963 | 3 | 3 | 8 | 5 | 1 | 2 | 2 | 4 | 4 | 2 | 4 | 0 | 0 | 0 | 0 | 0 | 0 | 0 | 0 | 0 | 0 | 0 | 0 | 0 | 1 | Serine palmitoyltransferase |
| AL3G05654 | OG0026316 | 0 | 0 | 1 | 2 | 0 | 0 | 0 | 0 | 0 | 0 | 0 | 0 | 1 | 0 | 0 | 0 | 1 | 0 | 1 | 0 | 0 | 0 | 0 | 0 | 0 | AMP-binding enzyme |
| AL3G05655 | OG0020698 | 1 | 2 | 2 | 1 | 0 | 0 | 0 | 0 | 0 | 0 | 0 | 1 | 1 | 1 | 0 | 0 | 0 | 0 | 0 | 0 | 0 | 0 | 0 | 0 | 0 | Cytochrome P450 |
| AL3G05656 | OG0000762 | 6 | 4 | 7 | 6 | 5 | 2 | 3 | 1 | 3 | 3 | 2 | 0 | 0 | 0 | 0 | 0 | 0 | 0 | 0 | 0 | 4.92 | 1.14 | 32.41 | 32.41 | 1 | Mate efflux family protein |
| AL3G05657 | OG0000235 | 2 | 1 | 70 | 3 | 0 | 0 | 0 | 0 | 0 | 0 | 0 | 0 | 0 | 0 | 0 | 0 | 0 | 0 | 0 | 0 | 0 | 0 | 0 | 0 | 1 | Other |
| AL3G05658 | OG0001466 | 2 | 2 | 5 | 3 | 3 | 2 | 4 | 2 | 3 | 3 | 2 | 1 | 1 | 1 | 1 | 1 | 1 | 1 | 1 | 1 | 1.15 | 1.84 | 0 | 0 | 1 | AMP-binding enzyme |
| AL3G05659 | OG0001466 | 2 | 2 | 5 | 3 | 3 | 2 | 4 | 2 | 3 | 3 | 2 | 0 | 0 | 0 | 0 | 0 | 0 | 0 | 0 | 0 | 6.49 | 3.92 | 5.76 | 5.76 | 1 | AMP-binding enzyme |
| AL3G06283 | OG0003416 | 2 | 2 | 4 | 5 | 3 | 0 | 4 | 0 | 0 | 1 | 1 | 1 | 1 | 1 | 0 | 0 | 0 | 0 | 0 | 0 | 246.54 | 7.41 | 0 | 0 | 0 | Peroxidase precursor |
| AL3G06284 | OG0021644 | 1 | 1 | 2 | 1 | 0 | 0 | 0 | 0 | 0 | 0 | 0 | 0 | 0 | 0 | 0 | 0 | 0 | 0 | 0 | 0 | 39.42 | 2.77 | 0 | 0 | 0 | Peroxidase precursor |
| AL3G06285 | OG0000313 | 12 | 12 | 11 | 9 | 5 | 0 | 3 | 3 | 6 | 3 | 3 | 1 | 1 | 1 | 1 | 0 | 0 | 0 | 0 | 0 | 0 | 0 | 0.1 | 0.1 | 0 | Cytochrome P450 |
| AL3G06286 | OG0000589 | 7 | 6 | 8 | 9 | 6 | 0 | 1 | 3 | 4 | 2 | 3 | 1 | 0 | 1 | 0 | 0 | 0 | 0 | 0 | 0 | 0.16 | 0 | 0.29 | 0.29 | 0 | Cytochrome P450 |
| AL3G06287 | OG0022045 | 0 | 0 | 5 | 0 | 0 | 0 | 0 | 0 | 0 | 0 | 0 | 0 | 0 | 0 | 0 | 0 | 0 | 0 | 0 | 0 | 24.82 | 28.34 | 48.81 | 48.81 | 0 | Pheophorbide a oxygenase |
| AL3G06288 | OG0030776 | 0 | 0 | 2 | 0 | 0 | 0 | 0 | 0 | 0 | 0 | 0 | 0 | 0 | 0 | 1 | 0 | 0 | 0 | 0 | 0 | 0 | 0 | 0 | 0 | 0 | Cytochrome P450 |
| AL3G06289 | OG0000202 | 23 | 25 | 14 | 9 | 6 | 0 | 0 | 0 | 3 | 0 | 0 | 1 | 0 | 1 | 0 | 0 | 0 | 0 | 0 | 0 | 0 | 0 | 0 | 0 | 0 | Retrotransposon protein |
| AL3G06290 | OG0000589 | 7 | 6 | 8 | 9 | 6 | 0 | 1 | 3 | 4 | 2 | 3 | 0 | 0 | 0 | 0 | 0 | 0 | 0 | 0 | 0 | 0.05 | 0 | 3.2 | 3.2 | 0 | Cytochrome P450 |
| AL3G06291 | OG0030776 | 0 | 0 | 2 | 0 | 0 | 0 | 0 | 0 | 0 | 0 | 0 | 0 | 0 | 0 | 0 | 0 | 0 | 0 | 0 | 0 | 0 | 0 | 0 | 0 | 0 | Cytochrome P450 |
| AL3G06292 | OG0001747 | 4 | 5 | 5 | 4 | 4 | 1 | 1 | 0 | 2 | 1 | 1 | 1 | 1 | 1 | 0 | 0 | 0 | 0 | 0 | 0 | 0 | 0 | 0 | 0 | 1 | Terpene synthase |
| AL3G06293 | OG0000313 | 12 | 12 | 11 | 9 | 5 | 0 | 3 | 3 | 6 | 3 | 3 | 1 | 1 | 1 | 1 | 0 | 0 | 0 | 0 | 0 | 0 | 0 | 0 | 0 | 0 | Cytochrome P450 |
| AL3G06294 | OG0023494 | 0 | 2 | 1 | 1 | 0 | 0 | 0 | 0 | 0 | 0 | 0 | 1 | 1 | 1 | 0 | 0 | 0 | 0 | 0 | 0 | 20.24 | 30.91 | 0.03 | 0.03 | 0 | Cysteine-rich receptor-like protein |
| AL3G06343 | OG0000433 | 11 | 8 | 15 | 11 | 5 | 0 | 1 | 2 | 1 | 3 | 0 | 1 | 1 | 1 | 0 | 0 | 0 | 0 | 0 | 0 | 44.31 | 1.54 | 0 | 0 | 1 | Cytochrome P450 |
| AL3G06344 | OG0000340 | 11 | 5 | 10 | 9 | 12 | 0 | 2 | 2 | 4 | 5 | 4 | 1 | 0 | 1 | 0 | 0 | 0 | 0 | 0 | 0 | 6.03 | 0.99 | 0 | 0 | 0 | Cytochrome P450 |
| AL3G06345 | OG0022065 | 0 | 0 | 2 | 3 | 0 | 0 | 0 | 0 | 0 | 0 | 0 | 0 | 0 | 0 | 0 | 0 | 0 | 0 | 0 | 0 | 0.03 | 0 | 11.84 | 11.84 | 0 | Other |
| AL3G06346 | OG0000407 | 11 | 7 | 12 | 10 | 6 | 1 | 4 | 4 | 1 | 1 | 2 | 1 | 1 | 1 | 0 | 0 | 0 | 0 | 0 | 0 | 2.74 | 0.79 | 0 | 0 | 1 | ENT-kaurene synthase |
| AL3G06347 | OG0023336 | 1 | 1 | 1 | 1 | 0 | 0 | 0 | 0 | 0 | 0 | 0 | 1 | 0 | 1 | 0 | 0 | 0 | 0 | 0 | 0 | 0 | 0 | 0 | 0 | 0 | Peptidase |

**Fig. S16c.** *Cytochrome P450* genes inserted within gene-clusters identified on ALO03.

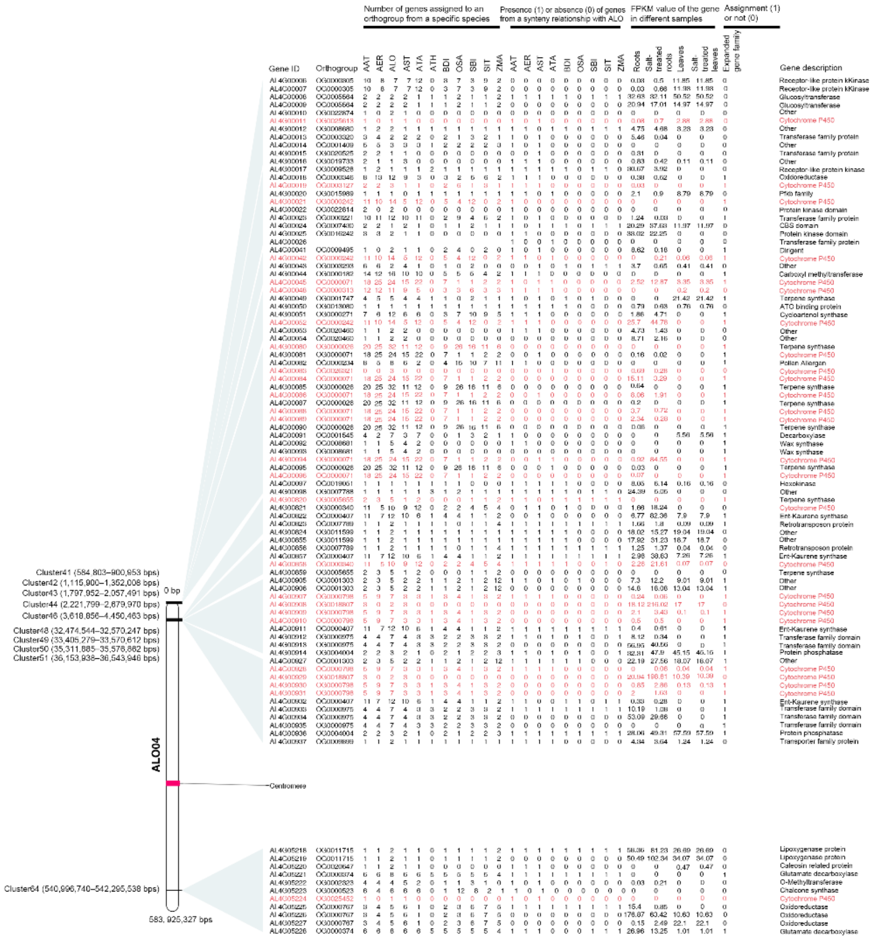

**Fig. S16d.** *Cytochrome P450* genes inserted within gene-clusters identified on ALO04.

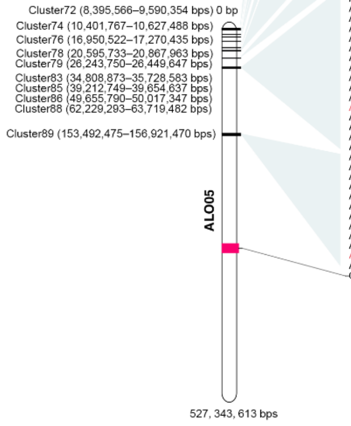

| Gene ID | Orthogroup | Number of genes assigned to an orthogroup from a specific species | | | | | | | | | | | Presence (1) or absence (0) of genes from a synteny relationship with ALO | | | | | | | | | FPKM value of the gene in different samples | | | | Assignment (1) or not (0) | Gene description |
|---|---|---|---|---|---|---|---|---|---|---|---|---|---|---|---|---|---|---|---|---|---|---|---|---|---|---|---|
| | | AAT | AER | ALO | AST | ATA | ATH | BDI | OSA | SBI | SIT | ZMA | AAT | AER | AST | ATA | BDI | OSA | SBI | SIT | ZMA | Roots | Salt-treated roots | Leaves | Salt-treated leaves | Expanded gene family | |
| AL5G00192 | OG0023734 | 0 | 0 | 4 | 0 | 0 | 0 | 0 | 0 | 0 | 0 | 0 | 0 | 0 | 0 | 0 | 0 | 0 | 0 | 0 | 0 | 0 | 0 | 0 | 0 | 0 | Ubiquitin family protein |
| AL5G00193 | OG0022884 | 1 | 0 | 2 | 1 | 0 | 0 | 0 | 0 | 0 | 0 | 0 | 1 | 0 | 1 | 0 | 0 | 0 | 0 | 0 | 0 | 0.13 | 0.28 | 0 | 0 | 0 | Cytochrome P450 |
| AL5G00194 | OG0000972 | 5 | 5 | 4 | 5 | 5 | 0 | 5 | 3 | 1 | 2 | 2 | 1 | 0 | 1 | 0 | 0 | 0 | 0 | 0 | 0 | 0.02 | 1.25 | 0.05 | 0.05 | 0 | Cytochrome P450 |
| AL5G00195 | OG0015983 | 2 | 0 | 6 | 0 | 0 | 0 | 0 | 2 | 0 | 0 | 0 | 1 | 0 | 0 | 0 | 0 | 0 | 0 | 0 | 0 | 0 | 0 | 0 | 0 | 0 | Cytochrome P450 |
| AL5G00196 | OG0021364 | 1 | 0 | 4 | 0 | 0 | 0 | 0 | 0 | 0 | 0 | 0 | 0 | 0 | 0 | 0 | 0 | 0 | 0 | 0 | 0 | 0 | 0 | 0 | 0 | 0 | Cytochrome P450 |
| AL5G00197 | OG0021364 | 1 | 0 | 4 | 0 | 0 | 0 | 0 | 0 | 0 | 0 | 0 | 0 | 0 | 0 | 0 | 0 | 0 | 0 | 0 | 0 | 0 | 0 | 0 | 0 | 0 | Cytochrome P450 |
| AL5G00198 | OG0021364 | 1 | 0 | 4 | 0 | 0 | 0 | 0 | 0 | 0 | 0 | 0 | 0 | 0 | 0 | 0 | 0 | 0 | 0 | 0 | 0 | 0 | 0 | 0 | 0 | 0 | Cytochrome P450 |
| AL5G00199 | OG0000346 | 8 | 13 | 12 | 9 | 3 | 0 | 3 | 2 | 5 | 6 | 2 | 1 | 0 | 1 | 0 | 0 | 0 | 0 | 0 | 0 | 0.42 | 0.57 | 0.62 | 0.62 | 1 | Oxidoreductase |
| AL5G00200 | OG0000972 | 5 | 5 | 4 | 5 | 5 | 0 | 5 | 3 | 1 | 2 | 2 | 0 | 1 | 0 | 0 | 0 | 0 | 0 | 0 | 0 | 0.17 | 1.98 | 0.02 | 0.02 | 0 | Cytochrome P450 |
| AL5G00201 | OG0022884 | 1 | 0 | 2 | 1 | 0 | 0 | 0 | 0 | 0 | 0 | 0 | 0 | 0 | 0 | 0 | 0 | 0 | 0 | 0 | 0 | 0 | 2.78 | 0 | 0 | 0 | Cytochrome P450 |
| AL5G00202 | OG0000346 | 8 | 13 | 12 | 9 | 3 | 0 | 3 | 2 | 5 | 6 | 2 | 0 | 0 | 0 | 0 | 0 | 0 | 0 | 0 | 0 | 0.65 | 1.15 | 1.33 | 1.33 | 1 | Oxidoreductase |
| AL5G00203 | OG0021364 | 1 | 0 | 4 | 0 | 0 | 0 | 0 | 0 | 0 | 0 | 0 | 0 | 0 | 0 | 0 | 0 | 0 | 0 | 0 | 0 | 0 | 0 | 0 | 0 | 0 | Cytochrome P450 |
| AL5G00204 | OG0015983 | 2 | 0 | 6 | 0 | 0 | 0 | 0 | 2 | 0 | 0 | 0 | 0 | 0 | 0 | 0 | 0 | 0 | 0 | 0 | 0 | 0 | 0 | 0 | 0 | 0 | Cytochrome P450 |
| AL5G00205 | OG0005758 | 2 | 1 | 2 | 2 | 2 | 2 | 1 | 1 | 1 | 1 | 1 | 1 | 1 | 1 | 1 | 1 | 0 | 1 | 1 | 1 | 84.12 | 62.28 | 33.68 | 33.68 | 0 | Other |
| AL5G00206 | OG0004791 | 1 | 1 | 1 | 1 | 1 | 1 | 1 | 1 | 2 | 1 | 8 | 1 | 1 | 1 | 1 | 1 | 1 | 1 | 1 | 1 | 3.3 | 2.9 | 1.96 | 1.96 | 0 | DNDj heat shock N-terminal domain |
| AL5G00207 | OG0019127 | 1 | 1 | 1 | 1 | 0 | 0 | 3 | 1 | 0 | 0 | 0 | 0 | 0 | 0 | 0 | 0 | 0 | 0 | 0 | 0 | 0.22 | 3.66 | 0.3 | 0.3 | 0 | Cytochrome P450 |
| AL5G00208 | OG0009045 | 2 | 2 | 2 | 2 | 1 | 0 | 1 | 0 | 1 | 1 | 1 | 1 | 1 | 1 | 0 | 0 | 0 | 0 | 0 | 0 | 8.6 | 5.8 | 0.02 | 0.02 | 0 | Anthocyanin 5-O-glucosyltransferase |
| AL5G00209 | OG0001747 | 4 | 5 | 5 | 4 | 4 | 1 | 1 | 0 | 2 | 1 | 1 | 1 | 1 | 1 | 0 | 0 | 0 | 0 | 0 | 0 | 0.07 | 0.01 | 0 | 0 | 1 | Terpene synthase |
| AL5G00210 | OG0026126 | 0 | 1 | 1 | 1 | 0 | 0 | 0 | 0 | 0 | 0 | 0 | 0 | 0 | 1 | 0 | 0 | 0 | 0 | 0 | 0 | 0.97 | 0.01 | 0 | 0 | 0 | Cytochrome P450 |
| AL5G00211 | OG0001747 | 4 | 5 | 5 | 4 | 4 | 1 | 1 | 0 | 2 | 1 | 1 | 0 | 0 | 0 | 0 | 0 | 0 | 0 | 0 | 0 | 2.17 | 1.1 | 0.19 | 0.19 | 1 | Terpene synthase |
| AL5G00212 | OG0001747 | 4 | 5 | 5 | 4 | 4 | 1 | 1 | 0 | 2 | 1 | 1 | 0 | 0 | 0 | 0 | 0 | 0 | 0 | 0 | 0 | 3.24 | 20.12 | 0.68 | 0.68 | 0 | Terpene synthase |
| AL5G00213 | OG0011663 | 1 | 1 | 1 | 1 | 2 | 1 | 1 | 1 | 1 | 1 | 1 | 1 | 1 | 1 | 0 | 1 | 1 | 1 | 1 | 1 | 1.88 | 3.35 | 2.05 | 2.05 | 0 | AMP-binding enzyme family |
| AL5G00214 | OG0002633 | 2 | 1 | 2 | 2 | 2 | 3 | 2 | 2 | 2 | 3 | 3 | 1 | 0 | 1 | 1 | 1 | 1 | 1 | 1 | 1 | 365.5 | 307.01 | 113.62 | 113.62 | 0 | 6-Phosphogluconate dehydrogenase |
| AL5G00215 | OG0004342 | 1 | 2 | 2 | 2 | 2 | 1 | 2 | 2 | 2 | 2 | 2 | 0 | 1 | 1 | 1 | 1 | 1 | 1 | 1 | 1 | 38.95 | 52.91 | 41.9 | 41.9 | 0 | G-Protein alpha subunit |
| AL5G00216 | OG0011514 | 1 | 1 | 1 | 1 | 2 | 1 | 1 | 1 | 1 | 1 | 1 | 1 | 1 | 1 | 0 | 1 | 1 | 1 | 1 | 1 | 0.11 | 0.14 | 14.75 | 14.75 | 0 | PPR repeat domain |
| AL5G00217 | OG0000740 | 4 | 2 | 9 | 7 | 6 | 0 | 3 | 3 | 3 | 4 | 2 | 1 | 1 | 1 | 0 | 0 | 0 | 0 | 0 | 0 | 0.75 | 4.15 | 0 | 0 | 1 | Potassium transporter |
| AL5G00218 | OG0000372 | 11 | 9 | 19 | 10 | 2 | 0 | 1 | 6 | 0 | 2 | 1 | 1 | 0 | 1 | 0 | 0 | 0 | 0 | 0 | 0 | 59.96 | 404.2 | 67.59 | 67.59 | 0 | Pathogenesis-related Bet VI family |
| AL5G00219 | OG0000161 | 14 | 15 | 4 | 10 | 16 | 0 | 8 | 4 | 8 | 6 | 2 | 1 | 1 | 1 | 1 | 1 | 1 | 1 | 1 | 1 | 7.36 | 7.77 | 1.78 | 1.78 | 0 | Osfbx182-F-Box domain |
| AL5G00220 | OG0001963 | 4 | 2 | 6 | 4 | 3 | 0 | 3 | 1 | 1 | 1 | 2 | 1 | 0 | 1 | 0 | 0 | 0 | 0 | 0 | 0 | 8.45 | 0.19 | 0 | 0 | 0 | Other |
| AL5G00221 | OG0003228 | 3 | 2 | 6 | 3 | 3 | 0 | 3 | 0 | 2 | 0 | 0 | 0 | 0 | 0 | 0 | 0 | 0 | 0 | 0 | 0 | 1.34 | 0.01 | 0 | 0 | 1 | Resistance protein |
| AL5G00237 | OG0000026 | 20 | 25 | 32 | 11 | 12 | 0 | 9 | 26 | 16 | 11 | 6 | 1 | 1 | 0 | 0 | 0 | 0 | 0 | 0 | 0 | 0 | 0 | 0 | 0 | 1 | Terpene synthase |
| AL5G00238 | OG0000161 | 14 | 15 | 4 | 10 | 16 | 0 | 8 | 4 | 8 | 6 | 2 | 0 | 0 | 0 | 0 | 0 | 0 | 0 | 0 | 0 | 0.05 | 0.07 | 0 | 0 | 0 | Other |
| AL5G00239 | OG0000875 | 3 | 3 | 6 | 2 | 6 | 0 | 5 | 2 | 4 | 5 | 3 | 1 | 1 | 0 | 0 | 0 | 0 | 0 | 0 | 0 | 0 | 0 | 0 | 0 | 1 | Regulatory protein |
| AL5G00240 | OG0001353 | 4 | 3 | 4 | 4 | 3 | 0 | 3 | 3 | 3 | 4 | 1 | 1 | 1 | 1 | 1 | 1 | 0 | 0 | 0 | 0 | 0 | 0 | 0 | 0 | 0 | Pectinacetylesterase domain |
| AL5G00241 | OG0000340 | 11 | 5 | 10 | 9 | 12 | 0 | 2 | 2 | 4 | 5 | 4 | 1 | 1 | 1 | 0 | 0 | 0 | 0 | 0 | 0 | 0.06 | 0 | 0 | 0 | 0 | Cytochrome P450 |
| AL5G00242 | OG0000071 | 18 | 25 | 24 | 15 | 22 | 0 | 7 | 1 | 1 | 2 | 2 | 1 | 1 | 1 | 0 | 0 | 0 | 0 | 0 | 0 | 0.72 | 1.58 | 0.09 | 0.09 | 1 | Cytochrome P450 |
| AL5G00243 | OG0000026 | 20 | 25 | 32 | 11 | 12 | 0 | 9 | 26 | 16 | 11 | 6 | 1 | 1 | 1 | 0 | 0 | 0 | 0 | 0 | 0 | 0.13 | 0.15 | 0 | 0 | 1 | Terpene synthase |
| AL5G00244 | OG0023720 | 0 | 0 | 4 | 0 | 0 | 0 | 0 | 0 | 0 | 0 | 0 | 0 | 0 | 0 | 0 | 0 | 0 | 0 | 0 | 0 | 0 | 0 | 0 | 0 | 0 | Serine/Threonine-protein kinase |
| AL5G00245 | OG0000026 | 20 | 25 | 32 | 11 | 12 | 0 | 9 | 26 | 16 | 11 | 6 | 0 | 0 | 0 | 0 | 0 | 0 | 0 | 0 | 0 | 0 | 0 | 0 | 0 | 1 | Terpene synthase |
| AL5G00246 | OG0000026 | 20 | 25 | 32 | 11 | 12 | 0 | 9 | 26 | 16 | 11 | 6 | 0 | 0 | 0 | 0 | 0 | 0 | 0 | 0 | 0 | 0.04 | 0.11 | 0 | 0 | 1 | Terpene synthase |
| AL5G00247 | OG0000026 | 20 | 25 | 32 | 11 | 12 | 0 | 9 | 26 | 16 | 11 | 6 | 0 | 0 | 0 | 0 | 0 | 0 | 0 | 0 | 0 | 0 | 0 | 0.01 | 0.01 | 1 | Terpene synthase |
| AL5G00248 | OG0000290 | 8 | 8 | 9 | 8 | 7 | 2 | 5 | 5 | 5 | 5 | 7 | 1 | 1 | 1 | 1 | 0 | 0 | 0 | 0 | 0 | 59.01 | 23.15 | 18.19 | 18.19 | 1 | Pinhead |
| AL5G00412 | OG0011621 | 1 | 1 | 1 | 1 | 1 | 0 | 2 | 1 | 1 | 1 | 2 | 1 | 1 | 1 | 0 | 1 | 1 | 1 | 1 | 1 | 4.85 | 1.02 | 0.17 | 0.17 | 0 | Starch synthase |
| AL5G00413 | OG0002731 | 3 | 3 | 3 | 3 | 3 | 1 | 1 | 1 | 2 | 1 | 2 | 1 | 1 | 1 | 0 | 0 | 0 | 0 | 0 | 0 | 0.46 | 0.04 | 0 | 0 | 0 | Cytochrome P450 |
| AL5G00414 | OG0004530 | 2 | 2 | 2 | 2 | 2 | 1 | 2 | 1 | 2 | 1 | 2 | 1 | 1 | 1 | 1 | 1 | 0 | 0 | 0 | 0 | 108.37 | 76.04 | 50.73 | 50.73 | 0 | Other |
| AL5G00415 | OG0025527 | 1 | 0 | 1 | 1 | 0 | 0 | 0 | 0 | 0 | 0 | 0 | 0 | 0 | 0 | 0 | 0 | 0 | 0 | 0 | 0 | 4.64 | 5.91 | 1.77 | 1.77 | 0 | Polymerase |
| AL5G00416 | OG0005634 | 2 | 2 | 3 | 2 | 2 | 0 | 1 | 1 | 1 | 1 | 2 | 1 | 1 | 1 | 0 | 0 | 0 | 0 | 0 | 0 | 0.03 | 0.02 | 0 | 0 | 1 | Glycosyl hydrolase family |
| AL5G00417 | OG0005634 | 2 | 2 | 3 | 2 | 2 | 0 | 1 | 1 | 1 | 1 | 2 | 0 | 0 | 0 | 0 | 0 | 0 | 0 | 0 | 0 | 0 | 0.06 | 0 | 0 | 0 | Glycosyl hydrolase family |
| AL5G00418 | OG0000877 | 4 | 2 | 6 | 5 | 10 | 0 | 3 | 6 | 1 | 2 | 0 | 0 | 0 | 1 | 0 | 0 | 0 | 0 | 0 | 0 | 1.08 | 0.72 | 200.75 | 200.75 | 1 | Other |
| AL5G00481 | OG0001678 | 1 | 1 | 1 | 1 | 1 | 11 | 1 | 3 | 1 | 3 | 5 | 1 | 1 | 1 | 1 | 1 | 1 | 1 | 1 | 1 | 5.95 | 0.39 | 0 | 0 | 0 | Other |
| AL5G00482 | OG0006083 | 2 | 2 | 2 | 2 | 1 | 1 | 1 | 1 | 1 | 1 | 2 | 1 | 1 | 1 | 1 | 1 | 1 | 1 | 1 | 0 | 3.06 | 2.33 | 0.44 | 0.44 | 0 | Brix domain containing protein |
| AL5G00483 | OG0007584 | 1 | 1 | 1 | 1 | 4 | 1 | 1 | 1 | 1 | 1 | 1 | 1 | 1 | 1 | 1 | 1 | 1 | 1 | 1 | 1 | 3 | 3.75 | 61.33 | 61.33 | 0 | Ribosome-binding factor A |
| AL5G00484 | OG0003770 | 2 | 2 | 2 | 2 | 2 | 2 | 1 | 2 | 3 | 1 | 2 | 1 | 1 | 1 | 1 | 1 | 1 | 1 | 1 | 1 | 46.14 | 8.61 | 0 | 0 | 0 | Gdp-Mannose 4,6 Dehydratase |
| AL5G00485 | OG0003770 | 2 | 2 | 2 | 2 | 2 | 2 | 1 | 2 | 3 | 1 | 2 | 0 | 0 | 0 | 0 | 0 | 0 | 0 | 0 | 0 | 78.5 | 48.08 | 29.78 | 29.78 | 0 | Gdp-Mannose 4,6 Dehydratase |
| AL5G00486 | OG0001884 | 2 | 4 | 5 | 4 | 4 | 0 | 5 | 1 | 1 | 1 | 0 | 0 | 0 | 0 | 0 | 0 | 0 | 0 | 0 | 0 | 2.03 | 0.21 | 0 | 0 | 1 | Transferase family protein |
| AL5G00487 | OG0005305 | 2 | 2 | 2 | 3 | 1 | 0 | 2 | 0 | 1 | 4 | 0 | 1 | 1 | 1 | 1 | 1 | 0 | 0 | 0 | 0 | 5.71 | 5.72 | 4.89 | 4.89 | 0 | Osfbx254-F-Box domain |
| AL5G00488 | OG0005431 | 2 | 2 | 2 | 2 | 1 | 1 | 1 | 1 | 2 | 1 | 2 | 1 | 1 | 1 | 0 | 0 | 0 | 0 | 0 | 0 | 4.39 | 5.28 | 4.45 | 4.45 | 0 | Methyltransferase |
| AL5G00489 | OG0000778 | 6 | 5 | 4 | 4 | 10 | 0 | 6 | 2 | 1 | 3 | 1 | 1 | 0 | 1 | 0 | 0 | 0 | 0 | 0 | 0 | 0 | 0 | 0 | 0 | 0 | Osfbx361-F-Box domain |
| AL5G00490 | OG0008480 | 1 | 1 | 1 | 1 | 1 | 1 | 1 | 2 | 1 | 1 | 2 | 1 | 1 | 1 | 1 | 0 | 0 | 0 | 0 | 0 | 5.48 | 2.89 | 0.09 | 0.09 | 0 | Duf630/Duf632 Domain |
| AL5G00491 | OG0000433 | 11 | 8 | 15 | 11 | 5 | 0 | 1 | 2 | 1 | 3 | 0 | 1 | 0 | 1 | 0 | 0 | 0 | 0 | 0 | 0 | 0 | 0 | 0 | 0 | 1 | Cytochrome P450 |
| AL5G00492 | OG0005855 | 2 | 2 | 2 | 2 | 2 | 1 | 1 | 1 | 1 | 1 | 1 | 1 | 1 | 1 | 1 | 1 | 0 | 1 | 1 | 1 | 17.62 | 10.68 | 10.82 | 10.82 | 0 | Bromodomain domain |
| AL5G00635 | OG0016931 | 1 | 1 | 1 | 1 | 0 | 1 | 1 | 1 | 1 | 1 | 1 | 1 | 1 | 1 | 0 | 1 | 1 | 1 | 1 | 1 | 25.33 | 29.71 | 19.66 | 19.66 | 0 | RNA recognition motif |
| AL5G00636 | OG0000159 | 9 | 10 | 14 | 12 | 12 | 1 | 6 | 9 | 2 | 9 | 4 | 0 | 0 | 0 | 0 | 0 | 0 | 0 | 0 | 0 | 1.98 | 7.41 | 8.99 | 8.99 | 1 | Transferase family domain |
| AL5G00637 | OG0018163 | 1 | 1 | 1 | 1 | 2 | 0 | 1 | 0 | 1 | 1 | 0 | 1 | 1 | 1 | 1 | 0 | 1 | 1 | 0 | 1 | 0 | 0 | 0 | 0 | 0 | Gdsl-Like lipase |
| AL5G00638 | OG0000868 | 5 | 5 | 4 | 3 | 6 | 0 | 4 | 1 | 2 | 6 | 3 | 1 | 1 | 0 | 1 | 0 | 0 | 1 | 1 | 1 | 0 | 0 | 0 | 0 | 0 | Transcription elongation factor 1 |
| AL5G00639 | OG0000710 | 7 | 11 | 9 | 6 | 2 | 0 | 3 | 0 | 1 | 3 | 2 | 1 | 1 | 1 | 0 | 0 | 0 | 0 | 0 | 0 | 0 | 0 | 0 | 0 | 1 | Cytochrome P450 |
| AL5G00640 | OG0005058 | 2 | 2 | 2 | 2 | 1 | 2 | 1 | 1 | 2 | 2 | 1 | 1 | 1 | 1 | 0 | 1 | 0 | 0 | 0 | 0 | 0 | 0 | 0 | 0 | 0 | Other |
| AL5G00641 | OG0023099 | 1 | 1 | 1 | 1 | 0 | 0 | 0 | 0 | 0 | 0 | 0 | 1 | 1 | 1 | 0 | 0 | 0 | 0 | 0 | 0 | 0.05 | 0.04 | 0 | 0 | 0 | Xylosyltransferase |
| AL5G00642 | OG0013775 | 1 | 1 | 1 | 1 | 1 | 1 | 1 | 1 | 1 | 1 | 1 | 1 | 1 | 1 | 1 | 1 | 1 | 1 | 1 | 1 | 3.61 | 8.13 | 88.56 | 88.56 | 0 | Starch synthase |
| AL5G00643 | OG0013687 | 1 | 1 | 1 | 1 | 1 | 0 | 1 | 2 | 1 | 1 | 1 | 1 | 1 | 1 | 1 | 1 | 0 | 1 | 1 | 1 | 20.19 | 0.65 | 0 | 0 | 0 | Other |
| AL5G00839 | OG0001144 | 3 | 1 | 6 | 4 | 4 | 2 | 4 | 3 | 3 | 3 | 2 | 1 | 1 | 1 | 0 | 0 | 0 | 0 | 0 | 0 | 4.42 | 5.63 | 1.25 | 1.25 | 1 | Cycloartenol synthase |
| AL5G00840 | OG0000242 | 11 | 10 | 14 | 5 | 12 | 0 | 5 | 4 | 12 | 0 | 2 | 1 | 1 | 0 | 0 | 0 | 0 | 0 | 0 | 0 | 0.05 | 0.24 | 0 | 0 | 1 | Cytochrome P450 |
| AL5G00841 | OG0000242 | 11 | 10 | 14 | 5 | 12 | 0 | 5 | 4 | 12 | 0 | 2 | 0 | 0 | 0 | 0 | 0 | 0 | 0 | 0 | 0 | 0 | 0 | 0 | 0 | 1 | Cytochrome P450 |
| AL5G00842 | OG0002010 | 3 | 2 | 4 | 3 | 2 | 1 | 2 | 2 | 2 | 2 | 3 | 1 | 1 | 1 | 1 | 1 | 1 | 1 | 1 | 1 | 0.2 | 0 | 0 | 0 | 1 | AP2 domain containing protein |
| AL5G00843 | OG0005230 | 2 | 2 | 5 | 3 | 1 | 0 | 1 | 0 | 1 | 2 | 1 | 0 | 0 | 1 | 0 | 0 | 0 | 0 | 0 | 0 | 1.28 | 0.02 | 0 | 0 | 1 | Cytochrome P450 |
| AL5G00844 | OG0005230 | 2 | 2 | 5 | 3 | 1 | 0 | 1 | 0 | 1 | 2 | 1 | 0 | 0 | 0 | 0 | 0 | 0 | 0 | 0 | 0 | 4.02 | 0.24 | 0 | 0 | 1 | Cytochrome P450 |
| AL5G00845 | OG0023729 | 0 | 0 | 3 | 0 | 0 | 0 | 0 | 0 | 0 | 0 | 1 | 0 | 0 | 0 | 0 | 0 | 0 | 0 | 0 | 0 | 0 | 0 | 0 | 0 | 0 | Transcription initiation factor |
| AL5G00846 | OG0005230 | 2 | 2 | 5 | 3 | 1 | 0 | 1 | 0 | 1 | 2 | 1 | 0 | 0 | 0 | 0 | 0 | 0 | 0 | 0 | 0 | 2.29 | 0.01 | 0 | 0 | 0 | Cytochrome P450 |
| AL5G00847 | OG0014008 | 1 | 1 | 1 | 1 | 1 | 0 | 1 | 1 | 1 | 1 | 2 | 1 | 1 | 1 | 1 | 1 | 1 | 1 | 1 | 1 | 15.61 | 26.21 | 2.24 | 2.24 | 0 | Other |
| AL5G00909 | OG0000091 | 15 | 12 | 17 | 13 | 19 | 4 | 7 | 8 | 6 | 6 | 2 | 1 | 0 | 1 | 0 | 0 | 0 | 0 | 0 | 0 | 0 | 0 | 0 | 0 | 1 | Flavonol synthase |
| AL5G00910 | OG0002738 | 1 | 2 | 4 | 2 | 2 | 2 | 2 | 2 | 2 | 2 | 2 | 1 | 1 | 1 | 1 | 1 | 1 | 0 | 1 | 1 | 1.65 | 0.34 | 0 | 0 | 1 | Serine threonine kinase |
| AL5G00911 | OG0030858 | 0 | 0 | 1 | 1 | 0 | 0 | 0 | 0 | 0 | 0 | 0 | 0 | 0 | 0 | 0 | 0 | 0 | 0 | 0 | 0 | 0.37 | 0.09 | 0 | 0 | 0 | Serine threonine kinase |
| AL5G00912 | OG0002738 | 1 | 2 | 4 | 2 | 2 | 2 | 2 | 2 | 2 | 2 | 2 | 0 | 0 | 0 | 0 | 0 | 0 | 0 | 0 | 0 | 3.66 | 0.69 | 0 | 0 | 1 | Serine threonine kinase |
| **AL5G00913** | **OG0001594** | **3** | **2** | **4** | **4** | **1** | **0** | **1** | **3** | **5** | **3** | **3** | **1** | **1** | **1** | **1** | **1** | **0** | **1** | **1** | **1** | **61.32** | **92.82** | **42.26** | **42.26** | **0** | **Cytochrome P450** |
| AL5G00914 | OG0001847 | 6 | 15 | 2 | 3 | 0 | 0 | 1 | 0 | 0 | 0 | 0 | 0 | 0 | 0 | 0 | 0 | 0 | 0 | 0 | 0 | 4.74 | 8.17 | 9.22 | 9.22 | 0 | Retrotransposon protein |
| AL5G00915 | OG0001594 | 3 | 2 | 4 | 4 | 1 | 0 | 1 | 3 | 5 | 3 | 3 | 0 | 0 | 0 | 0 | 0 | 0 | 0 | 0 | 0 | 2.6 | 0.76 | 0.03 | 0.03 | 0 | Cytochrome P450 |
| AL5G00916 | OG0001594 | 3 | 2 | 4 | 4 | 1 | 0 | 1 | 3 | 5 | 3 | 3 | 0 | 0 | 0 | 0 | 0 | 0 | 0 | 0 | 0 | 0 | 0 | 1.49 | 1.49 | 0 | Cytochrome P450 |
| AL5G00917 | OG0001594 | 3 | 2 | 4 | 4 | 1 | 0 | 1 | 3 | 5 | 3 | 3 | 0 | 0 | 0 | 0 | 0 | 0 | 0 | 0 | 0 | 0.41 | 0.6 | 0 | 0 | 0 | Cytochrome P450 |
| AL5G00918 | OG0002151 | 2 | 1 | 3 | 2 | 2 | 2 | 3 | 3 | 2 | 3 | 3 | 1 | 1 | 1 | 1 | 1 | 0 | 1 | 1 | 1 | 44.56 | 41.16 | 33.44 | 33.44 | 1 | Osmkk5 |
| AL5G00919 | OG0012182 | 1 | 1 | 1 | 1 | 2 | 1 | 1 | 1 | 1 | 1 | 1 | 1 | 1 | 1 | 0 | 0 | 1 | 0 | 1 | 0 | 8.23 | 9.34 | 6.73 | 6.73 | 0 | Zinc ion binding protein |
| AL5G00920 | OG0005204 | 1 | 1 | 1 | 1 | 2 | 0 | 1 | 7 | 2 | 2 | 0 | 1 | 1 | 1 | 1 | 1 | 0 | 1 | 1 | 0 | 0.91 | 0.88 | 0.53 | 0.53 | 0 | Glutamate receptor |
| AL5G00921 | OG0003074 | 1 | 3 | 1 | 1 | 3 | 4 | 1 | 3 | 2 | 2 | 2 | 1 | 1 | 1 | 1 | 1 | 1 | 1 | 1 | 1 | 0 | 0 | 0 | 0 | 0 | Glucosyltransferase |
| AL5G00922 | OG0001335 | 6 | 3 | 3 | 7 | 3 | 0 | 3 | 1 | 2 | 1 | 3 | 1 | 1 | 1 | 1 | 0 | 0 | 0 | 0 | 0 | 1.41 | 0.2 | 0 | 0 | 0 | Cytochrome P450 |
| AL5G00923 | OG0001230 | 3 | 3 | 3 | 3 | 2 | 4 | 4 | 3 | 4 | 3 | 2 | 1 | 1 | 1 | 1 | 1 | 1 | 1 | 1 | 1 | 463.09 | 207.28 | 131.31 | 131.31 | 0 | 60S ribosomal protein |
| AL5G00924 | OG0004969 | 2 | 2 | 2 | 2 | 1 | 1 | 2 | 2 | 2 | 1 | 1 | 1 | 1 | 1 | 1 | 1 | 0 | 1 | 1 | 0 | 3.19 | 2.38 | 2.24 | 2.24 | 0 | DNA binding protein |
| AL5G00925 | OG0018567 | 0 | 1 | 1 | 0 | 1 | 0 | 2 | 1 | 1 | 1 | 1 | 1 | 1 | 1 | 1 | 1 | 1 | 1 | 1 | 1 | 0.1 | 0 | 0.06 | 0.06 | 0 | Hypro1 |
| AL5G01102 | OG0026347 | 0 | 0 | 2 | 0 | 0 | 0 | 0 | 0 | 1 | 0 | 0 | 0 | 0 | 0 | 0 | 0 | 0 | 0 | 0 | 0 | 0 | 0 | 0 | 0 | 0 | Chalcone synthase |
| AL5G01103 | OG0000424 | 6 | 8 | 12 | 6 | 8 | 0 | 1 | 7 | 4 | 4 | 2 | 0 | 1 | 0 | 0 | 0 | 0 | 0 | 0 | 0 | 0 | 0.04 | 0 | 0 | 1 | Cytochrome P450 |
| AL5G01104 | OG0000801 | 6 | 5 | 9 | 6 | 5 | 0 | 4 | 3 | 1 | 1 | 1 | 0 | 0 | 0 | 0 | 0 | 0 | 0 | 0 | 0 | 0 | 0 | 0 | 0 | 1 | Chalcone synthase |
| AL5G01105 | OG0026347 | 0 | 0 | 2 | 0 | 0 | 0 | 0 | 0 | 1 | 0 | 0 | 0 | 0 | 0 | 0 | 0 | 0 | 0 | 0 | 0 | 0 | 0 | 0 | 0 | 0 | Chalcone synthase |
| AL5G01347 | OG0004940 | 2 | 2 | 2 | 2 | 0 | 0 | 2 | 2 | 2 | 2 | 2 | 1 | 1 | 1 | 0 | 1 | 1 | 1 | 1 | 1 | 0.23 | 76.94 | 0 | 0 | 0 | Odorant1 |
| AL5G01348 | OG0013040 | 1 | 1 | 1 | 1 | 1 | 1 | 1 | 1 | 1 | 1 | 1 | 1 | 1 | 1 | 0 | 0 | 0 | 0 | 0 | 0 | 188.07 | 193.43 | 137.28 | 137.28 | 0 | Delta14-sterol reductase |
| AL5G01349 | OG0012813 | 1 | 1 | 1 | 1 | 1 | 1 | 1 | 1 | 1 | 1 | 1 | 1 | 1 | 1 | 1 | 1 | 1 | 1 | 1 | 1 | 37.03 | 51.05 | 17.39 | 17.39 | 0 | NAD binding domain |
| AL5G01350 | OG0015981 | 1 | 1 | 1 | 1 | 1 | 0 | 1 | 1 | 1 | 1 | 1 | 1 | 1 | 1 | 1 | 1 | 1 | 1 | 1 | 1 | 49.57 | 88.29 | 12.44 | 12.44 | 0 | Phosphate 5-kinase family protein |
| AL5G01351 | OG0000531 | 5 | 4 | 7 | 5 | 4 | 4 | 4 | 5 | 4 | 3 | 6 | 1 | 1 | 1 | 1 | 1 | 1 | 1 | 1 | 1 | 75.43 | 196.1 | 29.04 | 29.04 | 1 | Other |
| AL5G01352 | OG0000599 | 6 | 4 | 7 | 6 | 4 | 0 | 4 | 4 | 7 | 3 | 3 | 0 | 0 | 1 | 0 | 0 | 0 | 1 | 1 | 0 | 0.03 | 0 | 0.39 | 0.39 | 1 | 3-Ketoacyl-Coa synthase |
| AL5G01353 | OG0000599 | 6 | 4 | 7 | 6 | 4 | 0 | 4 | 4 | 7 | 3 | 3 | 0 | 0 | 0 | 0 | 0 | 0 | 0 | 0 | 0 | 0 | 0 | 1.51 | 1.51 | 1 | 3-Ketoacyl-Coa synthase |
| AL5G01354 | OG0021839 | 1 | 0 | 3 | 1 | 0 | 0 | 0 | 0 | 0 | 0 | 0 | 0 | 0 | 1 | 0 | 0 | 0 | 0 | 0 | 0 | 0 | 0 | 0 | 0 | 0 | 3-Ketoacyl-Coa synthase |
| AL5G01355 | OG0021839 | 1 | 0 | 3 | 1 | 0 | 0 | 0 | 0 | 0 | 0 | 0 | 0 | 0 | 0 | 0 | 0 | 0 | 0 | 0 | 0 | 0 | 0 | 0 | 0 | 0 | 3-Ketoacyl-Coa synthase |
| AL5G01356 | OG0021839 | 1 | 0 | 3 | 1 | 0 | 0 | 0 | 0 | 0 | 0 | 0 | 0 | 0 | 0 | 0 | 0 | 0 | 0 | 0 | 0 | 0 | 0 | 0 | 0 | 0 | 3-Ketoacyl-Coa synthase |
| AL5G01357 | OG0014973 | 2 | 1 | 6 | 1 | 0 | 0 | 0 | 1 | 0 | 0 | 0 | 0 | 0 | 1 | 0 | 0 | 0 | 1 | 1 | 0 | 0 | 0 | 0 | 0 | 0 | 3-Ketoacyl-Coa synthase |
| AL5G01358 | OG0020581 | 2 | 0 | 2 | 2 | 0 | 0 | 0 | 0 | 0 | 0 | 1 | 0 | 0 | 1 | 0 | 0 | 0 | 0 | 0 | 0 | 0 | 0 | 0 | 0 | 0 | 3-Ketoacyl-Coa synthase |
| AL5G01359 | OG0006532 | 8 | 0 | 6 | 1 | 0 | 0 | 0 | 0 | 0 | 0 | 0 | 0 | 0 | 0 | 0 | 0 | 0 | 0 | 0 | 0 | 0 | 0 | 0 | 0 | 1 | Cytochrome P450 |
| AL5G01360 | OG0000072 | 1 | 0 | 113 | 0 | 3 | 0 | 0 | 0 | 0 | 0 | 0 | 0 | 0 | 0 | 0 | 0 | 0 | 0 | 0 | 0 | 0 | 0 | 0 | 0 | 0 | Retrotransposon protein |
| AL5G01361 | OG0014973 | 2 | 1 | 6 | 1 | 0 | 0 | 0 | 1 | 0 | 0 | 0 | 0 | 0 | 0 | 0 | 0 | 0 | 0 | 0 | 0 | 0 | 0 | 0 | 0 | 0 | 3-Ketoacyl-Coa synthase |
| AL5G01362 | OG0014973 | 2 | 1 | 6 | 1 | 0 | 0 | 0 | 1 | 0 | 0 | 0 | 0 | 0 | 0 | 0 | 0 | 1 | 0 | 0 | 0 | 0 | 0 | 0 | 0 | 0 | 3-Ketoacyl-Coa synthase |
| AL5G01363 | OG0014973 | 2 | 1 | 6 | 1 | 0 | 0 | 0 | 1 | 0 | 0 | 0 | 0 | 0 | 0 | 0 | 0 | 0 | 0 | 0 | 0 | 0 | 0 | 0 | 0 | 0 | 3-Ketoacyl-Coa synthase |
| AL5G02309 | OG0017336 | 1 | 1 | 1 | 1 | 0 | 1 | 1 | 1 | 1 | 1 | 1 | 1 | 1 | 1 | 0 | 1 | 0 | 1 | 1 | 1 | 0 | 1.65 | 0.42 | 0.42 | 0 | Oxidoreductase |
| AL5G02310 | OG0005675 | 2 | 1 | 2 | 2 | 1 | 1 | 2 | 3 | 1 | 1 | 1 | 1 | 0 | 1 | 0 | 0 | 0 | 0 | 0 | 0 | 31.85 | 44.52 | 23.93 | 23.93 | 0 | Retrotransposon protein |
| AL5G02311 | OG0020687 | 1 | 1 | 1 | 1 | 0 | 0 | 1 | 1 | 0 | 0 | 0 | 1 | 1 | 1 | 1 | 1 | 1 | 1 | 1 | 1 | 1.52 | 0.56 | 0 | 0 | 0 | Other |
| AL5G02312 | OG0000665 | 5 | 5 | 5 | 4 | 3 | 1 | 4 | 3 | 5 | 5 | 5 | 1 | 1 | 1 | 1 | 1 | 1 | 1 | 1 | 1 | 94.54 | 109.44 | 49.77 | 49.77 | 0 | Gibberellin receptor Gid1L2 |
| AL5G02313 | OG0018535 | 0 | 1 | 1 | 1 | 1 | 0 | 1 | 1 | 1 | 1 | 1 | 0 | 0 | 1 | 1 | 1 | 1 | 1 | 1 | 1 | 9.9 | 12.61 | 5.24 | 5.24 | 0 | Gibberellin receptor Gid1L2 |
| AL5G02314 | OG0016995 | 1 | 1 | 1 | 1 | 1 | 0 | 1 | 1 | 1 | 1 | 1 | 0 | 0 | 1 | 1 | 1 | 0 | 0 | 1 | 1 | 6.64 | 71.51 | 0.77 | 0.77 | 0 | Gibberellin receptor Gid1L2 |
| AL5G02315 | OG0006850 | 2 | 2 | 2 | 2 | 1 | 0 | 1 | 1 | 1 | 3 | 0 | 0 | 0 | 0 | 0 | 0 | 0 | 0 | 0 | 0 | 7.41 | 7.33 | 1.5 | 1.5 | 0 | Gibberellin receptor Gid1L2 |
| AL5G02316 | OG0002904 | 2 | 2 | 2 | 2 | 2 | 1 | 2 | 2 | 3 | 4 | 1 | 1 | 1 | 1 | 1 | 1 | 1 | 1 | 1 | 1 | 16.93 | 29.87 | 22.38 | 22.38 | 0 | Protein transport protein |
| AL5G02317 | OG0002337 | 3 | 2 | 3 | 3 | 4 | 1 | 1 | 2 | 2 | 2 | 2 | 1 | 1 | 1 | 1 | 1 | 1 | 1 | 1 | 1 | 13.38 | 51.01 | 0.05 | 0.05 | 0 | AP2 Domain |
| AL5G02318 | OG0001381 | 3 | 3 | 3 | 3 | 2 | 0 | 1 | 4 | 4 | 7 | 2 | 1 | 1 | 1 | 1 | 1 | 1 | 1 | 1 | 1 | 3.46 | 0.52 | 0.03 | 0.03 | 0 | Alpha-Amylase precursor |
| AL5G02319 | OG0005074 | 2 | 2 | 2 | 2 | 1 | 2 | 1 | 2 | 1 | 1 | 2 | 1 | 1 | 1 | 1 | 1 | 1 | 1 | 1 | 1 | 0.18 | 2.3 | 6 | 6 | 0 | Cytochrome P450 |
| AL5G02320 | OG0005985 | 2 | 2 | 2 | 2 | 1 | 2 | 1 | 1 | 1 | 1 | 1 | 1 | 1 | 1 | 0 | 1 | 1 | 1 | 1 | 1 | 8.16 | 4.7 | 4.18 | 4.18 | 0 | Fucosyltransferase |
| Centromere | | | | | | | | | | | | | | | | | | | | | | | | | | | |

**Fig. S16e.** *Cytochrome P450* genes inserted within gene-clusters identified on ALO05.

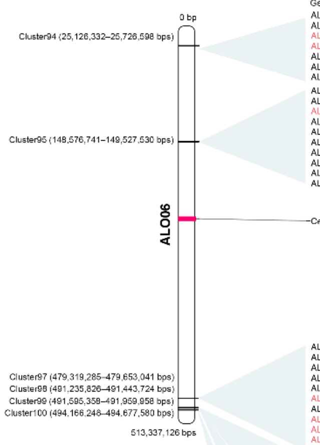

| | | Number of genes assigned to an orthogroup from a specific species | | | | | | | | | | | Presence (1) or absence (0) of genes from a synteny relationship with ALO | | | | | | | | | FPKM value of the gene in different samples | | | | Assignment (1) or not (0) | |
|---|---|---|---|---|---|---|---|---|---|---|---|---|---|---|---|---|---|---|---|---|---|---|---|---|---|---|---|
| Gene ID | Orthogroup | AAT | AER | ALO | AST | ATA | ATH | BDI | OSA | SBI | SIT | ZMA | AAT | AER | AST | ATA | BDI | OSA | SBI | SIT | ZMA | Roots | Salt-treated roots | Leaves | Salt-treated leaves | Expanded gene family | Gene description |
| AL6G00557 | OG0018340 | 1 | 0 | 6 | 2 | 0 | 0 | 0 | 0 | 0 | 0 | 0 | 0 | 0 | 0 | 0 | 0 | 0 | 0 | 0 | 0 | 0 | 0 | 0 | 0 | 0 | Nodulin |
| AL6G00558 | OG0014768 | 1 | 2 | 1 | 1 | 1 | 2 | 0 | 1 | 1 | 1 | 0 | 1 | 1 | 1 | 0 | 0 | 0 | 0 | 0 | 0 | 63.12 | 81.46 | 108.26 | 108.26 | 0 | Dj-1 family protein |
| AL6G00559 | OG0020641 | 1 | 2 | 1 | 1 | 1 | 0 | 0 | 0 | 0 | 0 | 0 | 1 | 1 | 1 | 0 | 0 | 0 | 0 | 0 | 0 | 5.52 | 48.41 | 3.88 | 3.88 | 0 | Cytochrome P450 |
| AL6G00560 | OG0000313 | 12 | 12 | 11 | 9 | 5 | 0 | 3 | 3 | 6 | 3 | 3 | 1 | 1 | 1 | 0 | 0 | 0 | 0 | 0 | 0 | 0 | 0 | 0 | 0 | 0 | Cytochrome P450 |
| AL6G00561 | OG0018178 | 1 | 4 | 1 | 1 | 1 | 0 | 1 | 0 | 0 | 0 | 0 | 1 | 1 | 1 | 0 | 0 | 0 | 0 | 0 | 0 | 0 | 0 | 0 | 0 | 0 | Cytokinin-O-glucosyltransferase |
| AL6G00562 | OG0016811 | 1 | 0 | 1 | 1 | 1 | 1 | 1 | 1 | 1 | 1 | 1 | 1 | 0 | 1 | 0 | 0 | 0 | 0 | 0 | 0 | 0.08 | 1.12 | 1066.81 | 1066.81 | 0 | Erythronate-4-Phosphate dehydrogenase |
| AL6G00563 | OG0000824 | 3 | 3 | 3 | 3 | 10 | 2 | 3 | 2 | 6 | 4 | 2 | 1 | 1 | 1 | 0 | 0 | 0 | 0 | 0 | 0 | 484.25 | 447.27 | 351.51 | 351.51 | 0 | Skp1-like protein |
| AL6G02059 | OG0008610 | 1 | 1 | 1 | 1 | 1 | 1 | 1 | 1 | 1 | 2 | 2 | 1 | 1 | 1 | 1 | 1 | 1 | 1 | 1 | 1 | 0.25 | 0.81 | 15.16 | 15.16 | 0 | FAD binding domain |
| AL6G02060 | OG0005921 | 1 | 1 | 1 | 1 | 1 | 3 | 1 | 2 | 1 | 2 | 2 | 1 | 1 | 1 | 1 | 1 | 1 | 1 | 1 | 1 | 3.77 | 6.65 | 5.47 | 5.47 | 0 | Agenet domain |
| AL6G02061 | OG0003278 | 2 | 2 | 2 | 2 | 1 | 0 | 2 | 4 | 2 | 2 | 3 | 1 | 1 | 1 | 1 | 1 | 1 | 1 | 1 | 1 | 24.13 | 46.62 | 0.14 | 0.14 | 0 | Cytochrome P450 |
| AL6G02062 | OG0002818 | 2 | 2 | 2 | 2 | 2 | 4 | 1 | 2 | 2 | 2 | 2 | 1 | 1 | 1 | 1 | 0 | 1 | 1 | 1 | 1 | 0.51 | 0 | 0 | 0 | 0 | Other |
| AL6G02063 | OG0003660 | 2 | 2 | 2 | 1 | 2 | 2 | 2 | 2 | 2 | 2 | 2 | 1 | 1 | 1 | 1 | 1 | 1 | 1 | 1 | 1 | 93.27 | 97.86 | 91.01 | 91.01 | 0 | 26S protease regulatory subunit |
| AL6G02064 | OG0001260 | 3 | 3 | 3 | 3 | 2 | 4 | 3 | 3 | 3 | 3 | 3 | 1 | 1 | 1 | 1 | 1 | 1 | 1 | 1 | 1 | 160.41 | 233.47 | 25.47 | 25.47 | 0 | 3-Ketoacyl-Coa synthase |
| AL6G02065 | OG0021753 | 1 | 0 | 2 | 0 | 0 | 0 | 0 | 0 | 0 | 0 | 2 | 0 | 0 | 0 | 0 | 0 | 0 | 0 | 0 | 0 | 0 | 0 | 0 | 0 | 0 | Putative lysosomal pro-X carboxypeptidase |
| AL6G02066 | OG0000751 | 7 | 6 | 7 | 6 | 4 | 0 | 3 | 4 | 1 | 3 | 1 | 1 | 1 | 1 | 1 | 1 | 1 | 1 | 1 | 1 | 7.15 | 16.95 | 0 | 0 | 0 | UDP-glucoronosyl and UDP-glucosyl transferase |
| AL6G02067 | OG0000751 | 7 | 6 | 7 | 6 | 4 | 0 | 3 | 4 | 1 | 3 | 1 | 0 | 0 | 0 | 0 | 1 | 0 | 0 | 0 | 0 | 0.01 | 0.51 | 12.33 | 12.33 | 0 | UDP-glucoronosyl and UDP-glucosyl transferase |
| AL6G02068 | OG0000751 | 7 | 6 | 7 | 6 | 4 | 0 | 3 | 4 | 1 | 3 | 1 | 0 | 0 | 0 | 0 | 0 | 0 | 0 | 0 | 0 | 19.04 | 32.34 | 10.53 | 10.53 | 0 | UDP-glucoronosyl and UDP-glucosyl transferase |
| AL6G04699 | OG0005415 | 2 | 2 | 3 | 3 | 1 | 0 | 1 | 2 | 1 | 1 | 1 | 1 | 1 | 1 | 0 | 0 | 0 | 0 | 0 | 0 | 0 | 0 | 0 | 0 | 0 | Oxoacyl-reductase |
| AL6G04700 | OG0011298 | 1 | 1 | 2 | 1 | 1 | 1 | 1 | 1 | 1 | 1 | 1 | 1 | 1 | 1 | 1 | 1 | 1 | 1 | 1 | 1 | 0.55 | 0.41 | 81.2 | 81.2 | 0 | Pheophorbide A oxygenase |
| AL6G04701 | OG0011227 | 1 | 1 | 2 | 1 | 1 | 1 | 1 | 1 | 1 | 1 | 1 | 1 | 1 | 1 | 0 | 1 | 1 | 1 | 1 | 1 | 135.83 | 210.67 | 124.91 | 124.91 | 0 | Tumour differentially expressed protein, Putative, Expresse |
| AL6G04702 | OG0023139 | 1 | 0 | 2 | 1 | 0 | 0 | 0 | 0 | 0 | 0 | 0 | 1 | 0 | 1 | 0 | 0 | 0 | 0 | 0 | 0 | 3.99 | 15.66 | 6.41 | 6.41 | 0 | Other |
| AL6G04703 | OG0010980 | 1 | 1 | 2 | 1 | 1 | 1 | 1 | 1 | 1 | 1 | 1 | 1 | 1 | 1 | 1 | 1 | 1 | 1 | 1 | 1 | 7.47 | 8.2 | 4.66 | 4.66 | 0 | Zinc finger protein |
| AL6G04704 | OG0001010 | 2 | 4 | 9 | 3 | 4 | 0 | 2 | 4 | 5 | 1 | 3 | 1 | 1 | 1 | 0 | 0 | 0 | 0 | 0 | 0 | 0.44 | 0.42 | 0 | 0 | 1 | Cytochrome P450 |
| AL6G04705 | OG0001466 | 2 | 2 | 5 | 3 | 3 | 2 | 4 | 2 | 3 | 3 | 2 | 1 | 0 | 1 | 0 | 0 | 0 | 0 | 0 | 0 | 0.06 | 0.04 | 0 | 0 | 0 | AMP-binding domain containing protein |
| AL6G04706 | OG0001010 | 2 | 4 | 9 | 3 | 4 | 0 | 2 | 4 | 5 | 1 | 3 | 0 | 0 | 0 | 0 | 0 | 0 | 0 | 0 | 0 | 0 | 0 | 0 | 0 | 1 | Cytochrome P450 |
| AL6G04707 | OG0001010 | 2 | 4 | 9 | 3 | 4 | 0 | 2 | 4 | 5 | 1 | 3 | 0 | 0 | 0 | 0 | 0 | 0 | 0 | 0 | 0 | 0 | 0 | 0 | 0 | 1 | Cytochrome P450 |
| AL6G04708 | OG0001010 | 2 | 4 | 9 | 3 | 4 | 0 | 2 | 4 | 5 | 1 | 3 | 0 | 0 | 0 | 0 | 0 | 0 | 0 | 0 | 0 | 0 | 0 | 0 | 0 | 1 | Cytochrome P450 |
| AL6G04709 | OG0001010 | 2 | 4 | 9 | 3 | 4 | 0 | 2 | 4 | 5 | 1 | 3 | 0 | 0 | 0 | 0 | 0 | 0 | 0 | 0 | 0 | 0 | 0 | 0 | 0 | 1 | Transposon protein |
| AL6G04927 | OG0002988 | 2 | 2 | 3 | 2 | 2 | 1 | 2 | 2 | 2 | 3 | 2 | 1 | 1 | 1 | 1 | 1 | 1 | 1 | 1 | 1 | 6.26 | 5.26 | 1.54 | 1.54 | 1 | Ribonucleoside-diphosphate reductase large subunit |
| AL6G04928 | OG0000921 | 5 | 4 | 6 | 6 | 2 | 0 | 3 | 2 | 3 | 5 | 2 | 1 | 0 | 1 | 0 | 0 | 0 | 0 | 0 | 0 | 0 | 0 | 0 | 0 | 1 | Cytokinin-O-glucosyltransferase |
| AL6G04929 | OG0000361 | 5 | 8 | 15 | 10 | 11 | 0 | 6 | 2 | 1 | 3 | 1 | 1 | 1 | 0 | 0 | 0 | 0 | 0 | 0 | 0 | 0 | 0 | 0 | 0 | 1 | Putative subtilisin homologue |
| AL6G04930 | OG0004433 | 4 | 2 | 5 | 3 | 1 | 0 | 1 | 1 | 0 | 1 | 1 | 1 | 0 | 1 | 0 | 0 | 0 | 0 | 0 | 0 | 0.05 | 0.07 | 0 | 0 | 1 | Putative subtilisin homologue |
| AL6G04931 | OG0002024 | 2 | 2 | 3 | 3 | 2 | 3 | 2 | 2 | 2 | 2 | 3 | 1 | 1 | 1 | 1 | 1 | 1 | 1 | 1 | 1 | 14.39 | 14.22 | 7.04 | 7.04 | 0 | Other |
| AL6G04932 | OG0023243 | 1 | 1 | 1 | 0 | 0 | 0 | 1 | 0 | 0 | 0 | 0 | 0 | 0 | 1 | 0 | 0 | 0 | 0 | 0 | 0 | 18.94 | 2.32 | 0 | 0 | 0 | Cytochrome P450 |
| AL6G04933 | OG0000340 | 11 | 5 | 10 | 9 | 12 | 0 | 2 | 2 | 4 | 5 | 4 | 1 | 1 | 0 | 0 | 0 | 0 | 0 | 0 | 0 | 5.06 | 0.71 | 0 | 0 | 0 | Cytochrome P450 |
| AL6G04934 | OG0000472 | 5 | 8 | 11 | 6 | 11 | 0 | 2 | 6 | 1 | 4 | 1 | 1 | 0 | 1 | 0 | 0 | 0 | 0 | 0 | 0 | 3.53 | 0.67 | 0 | 0 | 1 | Cytochrome P450 |
| AL6G04935 | OG0025622 | 1 | 0 | 2 | 0 | 0 | 0 | 0 | 0 | 0 | 0 | 0 | 1 | 0 | 0 | 0 | 0 | 0 | 0 | 0 | 0 | 0 | 0 | 0 | 0 | 0 | Osfbx120-F-box domain |
| AL6G04941 | OG0004594 | 2 | 2 | 4 | 2 | 1 | 1 | 1 | 1 | 1 | 2 | 2 | 0 | 0 | 0 | 1 | 1 | 1 | 1 | 1 | 1 | 19.45 | 33.63 | 28.6 | 28.6 | 1 | Dehydrogenase |
| AL6G04942 | OG0000313 | 12 | 12 | 11 | 9 | 5 | 0 | 3 | 3 | 6 | 3 | 3 | 1 | 0 | 1 | 0 | 0 | 0 | 0 | 0 | 0 | 0 | 0 | 0.07 | 0.07 | 0 | Cytochrome P450 |
| AL6G04943 | OG0000052 | 19 | 16 | 29 | 13 | 11 | 0 | 6 | 9 | 9 | 11 | 11 | 1 | 1 | 1 | 0 | 0 | 0 | 0 | 0 | 0 | 0 | 0 | 0 | 0 | 1 | Sam dependent carboxyl methyltransferase |
| AL6G04944 | OG0004594 | 2 | 2 | 4 | 2 | 1 | 1 | 1 | 1 | 1 | 2 | 2 | 0 | 0 | 0 | 0 | 0 | 0 | 0 | 0 | 0 | 13.23 | 10.04 | 2.53 | 2.53 | 1 | Dehydrogenase |
| AL6G04945 | OG0010685 | 1 | 1 | 2 | 1 | 1 | 1 | 1 | 1 | 1 | 1 | 1 | 1 | 1 | 1 | 1 | 1 | 1 | 1 | 1 | 1 | 4.86 | 4.48 | 18.33 | 18.33 | 0 | Elmo/Ced-12 family protein |
| AL6G04946 | OG0008553 | 1 | 1 | 2 | 1 | 1 | 1 | 1 | 1 | 1 | 1 | 2 | 1 | 1 | 1 | 0 | 1 | 0 | 0 | 1 | 0 | 18.63 | 21.78 | 22.01 | 22.01 | 1 | Hydrolase |
| AL6G04947 | OG0010457 | 1 | 1 | 1 | 1 | 1 | 1 | 1 | 1 | 1 | 2 | 1 | 1 | 1 | 1 | 1 | 1 | 1 | 0 | 1 | 0 | 8.36 | 9.72 | 36.54 | 36.54 | 0 | Acetyltransferase |
| AL6G04948 | OG0010384 | 1 | 1 | 2 | 1 | 1 | 0 | 1 | 1 | 1 | 1 | 2 | 1 | 1 | 1 | 0 | 1 | 1 | 1 | 1 | 1 | 65.07 | 70.31 | 24.24 | 24.24 | 0 | Zinc finger domain |
| AL6G04949 | OG0000159 | 9 | 10 | 14 | 12 | 12 | 1 | 6 | 9 | 2 | 9 | 4 | 1 | 1 | 1 | 0 | 0 | 0 | 0 | 0 | 0 | 0.14 | 0 | 0 | 0 | 1 | Transferase family protein |
| AL6G04950 | OG0010218 | 1 | 1 | 2 | 1 | 1 | 1 | 1 | 1 | 1 | 1 | 1 | 1 | 1 | 1 | 1 | 1 | 1 | 1 | 1 | 1 | 4.52 | 0.96 | 0 | 0 | 1 | Other |
| AL6G05030 | OG0013570 | 1 | 1 | 1 | 1 | 1 | 0 | 1 | 1 | 1 | 1 | 2 | 1 | 1 | 1 | 1 | 1 | 1 | 1 | 1 | 1 | 47.6 | 75.22 | 8.15 | 8.15 | 0 | Pb1 domain |
| AL6G05031 | OG0000628 | 3 | 6 | 7 | 2 | 5 | 0 | 3 | 9 | 4 | 5 | 3 | 1 | 1 | 0 | 1 | 1 | 1 | 1 | 1 | 1 | 0.05 | 0 | 0 | 0 | 1 | Dehydrogenase |
| AL6G05032 | OG0000628 | 3 | 6 | 7 | 2 | 5 | 0 | 3 | 9 | 4 | 5 | 3 | 0 | 0 | 0 | 0 | 0 | 0 | 0 | 0 | 0 | 0.06 | 0 | 0 | 0 | 1 | Dehydrogenase |
| AL6G05033 | OG0026345 | 0 | 0 | 3 | 0 | 0 | 0 | 0 | 0 | 0 | 0 | 0 | 1 | 0 | 1 | 0 | 0 | 0 | 0 | 0 | 0 | 0 | 0 | 0 | 0 | 0 | G-beta repeat domain |
| AL6G05034 | OG0006938 | 2 | 3 | 2 | 1 | 0 | 0 | 0 | 3 | 2 | 2 | 0 | 0 | 0 | 0 | 0 | 0 | 0 | 0 | 0 | 0 | 0 | 0 | 0 | 0 | 0 | Chalcone synthase |
| AL6G05035 | | | | | | | | | | | | | 0 | 0 | 0 | 0 | 0 | 0 | 0 | 0 | 0 | 0.11 | 0.3 | 0 | 0 | 0 | Oxidoreductase |
| AL6G05036 | OG0006545 | 2 | 1 | 2 | 2 | 2 | 1 | 1 | 1 | 1 | 1 | 1 | 0 | 0 | 0 | 0 | 0 | 0 | 0 | 0 | 0 | 6.68 | 5.25 | 1.95 | 1.95 | 0 | G-beta repeat domain |
| AL6G05037 | OG0000628 | 3 | 6 | 7 | 2 | 5 | 0 | 3 | 9 | 4 | 5 | 3 | 0 | 0 | 0 | 0 | 0 | 0 | 0 | 0 | 0 | 0.92 | 0.76 | 0 | 0 | 1 | Dehydrogenase |
| AL6G05038 | OG0000628 | 3 | 6 | 7 | 2 | 5 | 0 | 3 | 9 | 4 | 5 | 3 | 0 | 0 | 0 | 0 | 0 | 0 | 0 | 0 | 0 | 0 | 0 | 0 | 0 | 1 | Dehydrogenase |
| AL6G05039 | OG0000628 | 3 | 6 | 7 | 2 | 5 | 0 | 3 | 9 | 4 | 5 | 3 | 0 | 0 | 0 | 0 | 0 | 0 | 0 | 0 | 0 | 6.65 | 4.63 | 0 | 0 | 0 | Dehydrogenase |
| AL6G05040 | OG0006450 | 1 | 1 | 2 | 1 | 2 | 3 | 1 | 1 | 1 | 1 | 1 | 1 | 1 | 1 | 1 | 1 | 1 | 1 | 1 | 1 | 7.2 | 10.02 | 4.8 | 4.8 | 0 | Translation ilnitiation factor |
| AL6G05041 | OG0009998 | 1 | 1 | 2 | 1 | 1 | 0 | 1 | 1 | 1 | 1 | 2 | 1 | 1 | 1 | 1 | 1 | 1 | 1 | 1 | 1 | 2.88 | 1.46 | 0 | 0 | 0 | Osfbx66-F-box domain |
| AL6G05042 | OG0019916 | 0 | 2 | 2 | 1 | 2 | 0 | 0 | 0 | 0 | 0 | 0 | 0 | 0 | 0 | 1 | 1 | 1 | 1 | 0 | 0 | 0 | 0.05 | 0 | 0 | 0 | Osfbx66-F-box domain |
| AL6G05043 | OG0006691 | 2 | 1 | 4 | 2 | 4 | 0 | 0 | 1 | 0 | 1 | 0 | 0 | 0 | 1 | 0 | 0 | 0 | 0 | 1 | 0 | 0.42 | 0.25 | 0.17 | 0.17 | 1 | Disease resistance protein |
| AL6G05044 | OG0003175 | 3 | 4 | 6 | 5 | 0 | 1 | 0 | 1 | 1 | 0 | 1 | 0 | 0 | 1 | 0 | 0 | 0 | 0 | 0 | 0 | 0 | 0 | 0 | 0 | 1 | Terpene synthase |
| AL6G05045 | OG0020844 | 0 | 1 | 3 | 2 | 0 | 0 | 0 | 0 | 0 | 0 | 0 | 0 | 0 | 1 | 0 | 0 | 0 | 0 | 0 | 0 | 0 | 0 | 0 | 0 | 0 | Cytochrome P450 |

**Fig. S16f.** *Cytochrome P450* genes inserted within gene-clusters identified on ALO06.

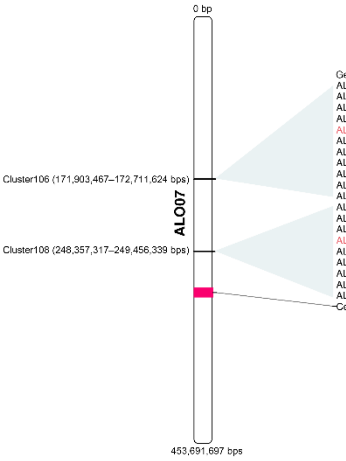

| Gene ID | Orthogroup | Number of genes assigned to an orthogroup from a specific species | | | | | | | | | | | Presence (1) or absence (0) of genes from a synteny relationship with ALO | | | | | | | | | FPKM value of the gene in different samples | | | | Assignment (1) or not (0) | Gene description |
|---|---|---|---|---|---|---|---|---|---|---|---|---|---|---|---|---|---|---|---|---|---|---|---|---|---|---|---|
| | | AAT | AER | ALO | AST | ATA | ATH | BDI | OSA | SBI | SIT | ZMA | AAT | AER | AST | ATA | BDI | OSA | SBI | SIT | ZMA | Roots | Salt-treated roots | Leaves | Salt-treated leaves | Expanded gene family | |
| AL7G03628 | OG0010563 | 1 | 1 | 1 | 1 | 1 | 1 | 1 | 1 | 2 | 1 | 1 | 1 | 1 | 1 | 1 | 1 | 0 | 0 | 0 | 0 | 5.52 | 5.51 | 5.81 | 5.81 | 0 | G-beta repeat domain |
| AL7G03629 | OG0000392 | 9 | 8 | 9 | 7 | 10 | 0 | 4 | 7 | 2 | 3 | 1 | 1 | 0 | 1 | 0 | 1 | 0 | 1 | 1 | 0 | 348.76 | 636.71 | 0.17 | 0.17 | 1 | Methyltransferase |
| AL7G03630 | OG0000392 | 9 | 8 | 9 | 7 | 10 | 0 | 4 | 7 | 2 | 3 | 1 | 0 | 0 | 0 | 1 | 0 | 0 | 0 | 0 | 0 | 377.9 | 810.69 | 0 | 0 | 1 | Methyltransferase |
| AL7G03631 | OG0002214 | 3 | 3 | 3 | 3 | 2 | 0 | 3 | 4 | 2 | 1 | 1 | 1 | 1 | 1 | 0 | 0 | 0 | 0 | 0 | 0 | 153.01 | 18.93 | 15.72 | 15.72 | 0 | Dirigent |
| AL7G03632 | OG0000324 | 7 | 6 | 7 | 5 | 8 | 2 | 5 | 12 | 5 | 4 | 4 | 1 | 1 | 1 | 0 | 0 | 0 | 0 | 0 | 0 | 50.91 | 8.68 | 7.61 | 7.61 | 0 | Cytochrome P450 |
| AL7G03633 | OG0000392 | 9 | 8 | 9 | 7 | 10 | 0 | 4 | 7 | 2 | 3 | 1 | 0 | 0 | 0 | 0 | 0 | 0 | 0 | 0 | 0 | 248.27 | 332.74 | 0.41 | 0.41 | 1 | Methyltransferase |
| AL7G03634 | OG0000072 | 1 | 0 | 113 | 0 | 3 | 0 | 0 | 0 | 0 | 0 | 0 | 0 | 0 | 0 | 0 | 0 | 0 | 0 | 0 | 0 | 0.04 | 0 | 0.03 | 0.03 | 0 | Other |
| AL7G03635 | OG0000392 | 9 | 8 | 9 | 7 | 10 | 0 | 4 | 7 | 2 | 3 | 1 | 0 | 0 | 0 | 0 | 0 | 0 | 0 | 0 | 0 | 0.05 | 0 | 0 | 0 | 1 | Methyltransferase |
| AL7G03636 | OG0000392 | 9 | 8 | 9 | 7 | 10 | 0 | 4 | 7 | 2 | 3 | 1 | 0 | 0 | 1 | 0 | 0 | 0 | 0 | 0 | 0 | 32.29 | 63.6 | 0 | 0 | 0 | Methyltransferase |
| AL7G03637 | OG0002898 | 4 | 3 | 4 | 4 | 4 | 0 | 1 | 1 | 1 | 1 | 0 | 1 | 1 | 1 | 0 | 1 | 0 | 0 | 1 | 0 | 0.04 | 1.37 | 0 | 0 | 0 | Other |
| AL7G03638 | OG0009509 | 1 | 1 | 1 | 1 | 3 | 1 | 1 | 1 | 2 | 1 | 0 | 1 | 1 | 1 | 1 | 1 | 0 | 0 | 0 | 0 | 42.39 | 37.52 | 2.03 | 2.03 | 0 | Cycloeucalenol cycloisomerase |
| AL7G04458 | OG0013614 | 1 | 1 | 1 | 1 | 1 | 1 | 1 | 1 | 1 | 1 | 1 | 1 | 0 | 1 | 1 | 1 | 0 | 0 | 1 | 0 | 3.96 | 2.86 | 1.95 | 1.95 | 0 | Other |
| AL7G04459 | OG0013528 | 1 | 1 | 1 | 1 | 1 | 1 | 1 | 1 | 1 | 1 | 1 | 1 | 0 | 1 | 1 | 1 | 0 | 1 | 1 | 0 | 10.46 | 6.76 | 180.92 | 180.92 | 0 | Phytoene synthase |
| AL7G04460 | OG0022989 | 1 | 1 | 1 | 1 | 0 | 0 | 0 | 0 | 0 | 0 | 0 | 1 | 1 | 1 | 0 | 0 | 0 | 0 | 0 | 0 | 4.78 | 4.09 | 2.72 | 2.72 | 0 | Other |
| AL7G04461 | OG0007399 | 1 | 2 | 1 | 1 | 2 | 0 | 1 | 2 | 1 | 2 | 1 | 1 | 1 | 1 | 0 | 0 | 0 | 0 | 0 | 0 | 2.14 | 26.57 | 0.32 | 0.32 | 0 | Cytochrome P450 |
| AL7G04462 | OG0006568 | 1 | 1 | 1 | 0 | 1 | 0 | 1 | 1 | 2 | 5 | 2 | 1 | 0 | 0 | 1 | 0 | 1 | 0 | 0 | 0 | 0 | 1.98 | 0 | 0 | 0 | Late embryogenesis abundant protein |
| AL7G04463 | | | | | | | | | | | | | 0 | 0 | 0 | 0 | 0 | 0 | 0 | 0 | 0 | 0 | 0 | 0 | 0 | 0 | Other |
| AL7G04464 | OG0000577 | 6 | 5 | 3 | 8 | 2 | 0 | 4 | 2 | 13 | 5 | 1 | 1 | 1 | 1 | 0 | 0 | 0 | 0 | 0 | 0 | 1.69 | 1.04 | 0.26 | 0.26 | 0 | Leucine rich repeat family protein |
| AL7G04465 | OG0003175 | 3 | 4 | 6 | 5 | 0 | 1 | 0 | 1 | 1 | 0 | 1 | 1 | 1 | 1 | 0 | 0 | 0 | 0 | 0 | 0 | 0 | 0 | 0 | 0 | 1 | Terpene synthase |
| AL7G04466 | OG0003175 | 3 | 4 | 6 | 5 | 0 | 1 | 0 | 1 | 1 | 0 | 1 | 0 | 0 | 0 | 0 | 0 | 0 | 0 | 0 | 0 | 0.71 | 0 | 0 | 0 | 0 | Terpene synthase |

**Fig. S16g.** *Cytochrome P450* genes inserted within gene-clusters identified on ALO07.

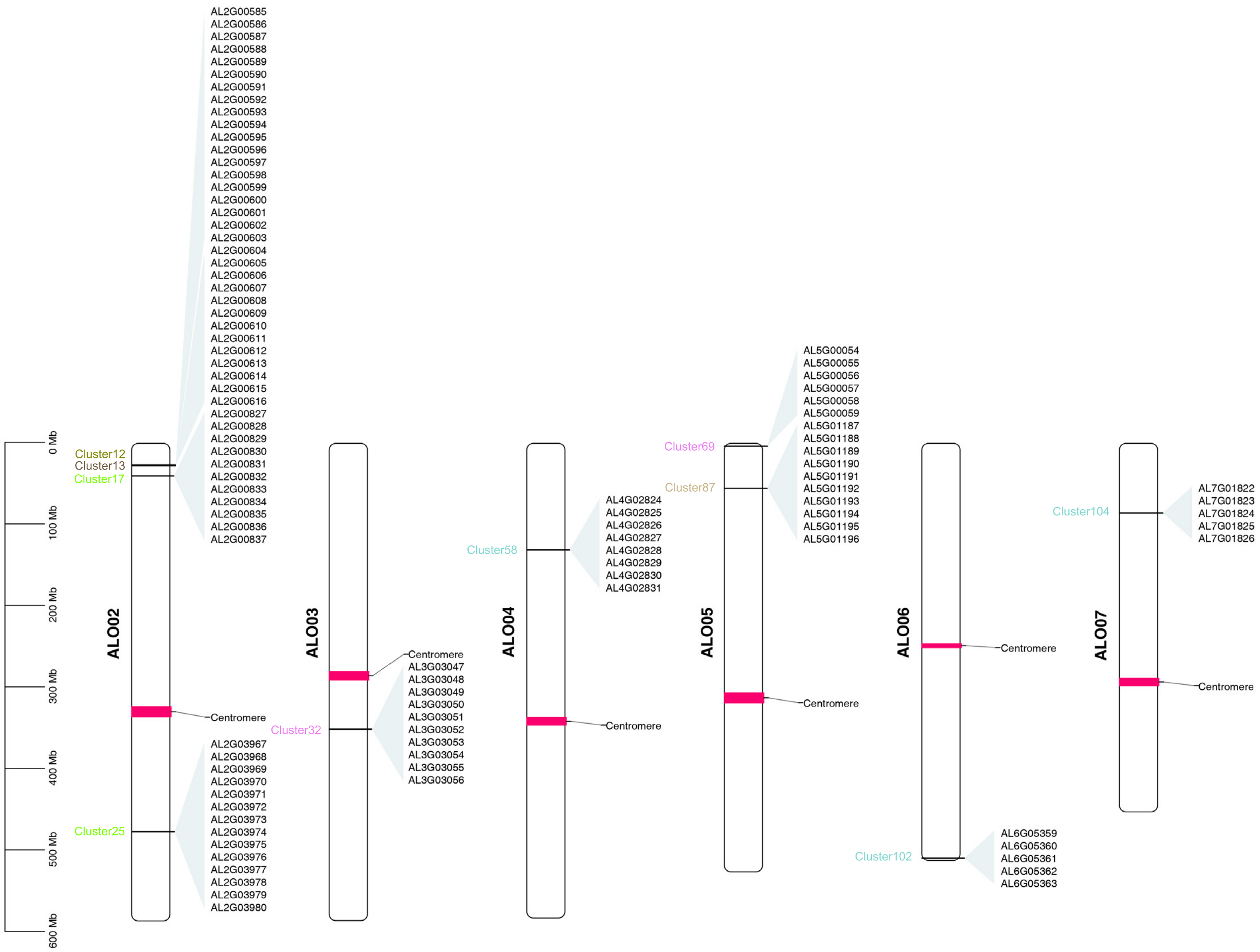

**Fig. S17.** Total 11 *CesA* and 55 *Csl* genes located within 10 clusters identified in the ALO genome. A scale in the left represented length of chromosome in megabases. The cluster type labeled as the same colours as Fig. 5a, the position of gene cluster (black band) scaled based on chromosome size and its physical position on the chromosome. The centromere position shown by pink band

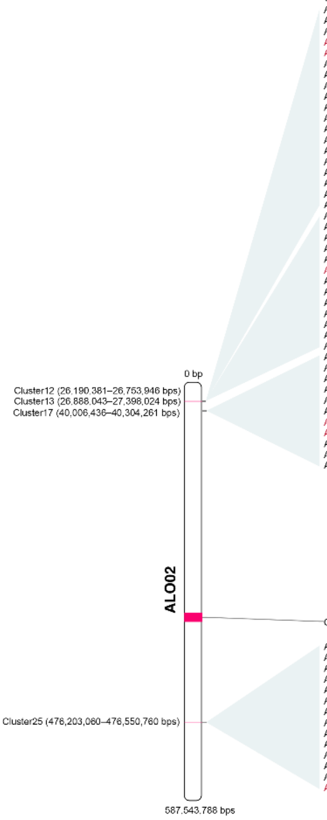

| | | Number of genes assigned to an orthogroup from a specific species | | | | | | | | | | | Presence (1) or absence (0) of genes from a synteny relationship with ALO | | | | | | | | | FPKM value of the gene in different samples | | | | Assignment (1) or not (0) | |
|---|---|---|---|---|---|---|---|---|---|---|---|---|---|---|---|---|---|---|---|---|---|---|---|---|---|---|---|
| Gene ID | Orthogroup | AAT | AER | ALO | AST | ATA | ATH | BDI | OSA | SBI | SIT | ZMA | AAT | AER | AST | ATA | BDI | OSA | SBI | SIT | ZMA | Roots | Salt-treated roots | Leaves | Salt-treated leaves | Expaned gene family | Gene description |
| AL2G00585 | OG0006787 | 2 | 1 | 4 | 1 | 1 | 1 | 1 | 1 | 1 | 1 | 1 | 1 | 1 | 1 | 1 | 1 | 0 | 0 | 0 | 0 | 11.41 | 11.4 | 10.61 | 16.52 | 1 | YL1 nuclear protein C-terminal domain |
| AL2G00586 | OG0017372 | 1 | 1 | 2 | 1 | 1 | 0 | 1 | 0 | 1 | 1 | 1 | 1 | 1 | 1 | 1 | 1 | 0 | 0 | 0 | 0 | 2.03 | 2.01 | 1.4 | 1.77 | 0 | Other |
| AL2G00587 | OG0003073 | 3 | 2 | 5 | 7 | 4 | 0 | 1 | 1 | 0 | 0 | 0 | 1 | 1 | 1 | 0 | 0 | 0 | 0 | 0 | 0 | 0.03 | 0 | 0 | 0 | 0 | Jacalin-like lectin domain containing protein |
| AL2G00588 | OG0025675 | 1 | 0 | 2 | 0 | 0 | 0 | 0 | 0 | 0 | 0 | 0 | 1 | 0 | 0 | 0 | 0 | 0 | 0 | 0 | 0 | 0 | 0 | 0 | 0 | 0 | CslF9 |
| AL2G00589 | OG0029677 | 1 | 0 | 1 | 0 | 0 | 0 | 0 | 0 | 0 | 0 | 0 | 0 | 0 | 0 | 0 | 0 | 0 | 0 | 0 | 0 | 0.72 | 0.81 | 5.93 | 5.15 | 0 | CslF8 |
| AL2G00590 | OG0011683 | 1 | 1 | 2 | 1 | 1 | 1 | 1 | 1 | 1 | 1 | 1 | 1 | 1 | 1 | 1 | 0 | 0 | 1 | 1 | 0 | 26.91 | 23.11 | 32.8 | 39.81 | 0 | Glyoxalase family |
| AL2G00591 | OG0014656 | 1 | 1 | 2 | 1 | 1 | 1 | 1 | 1 | 1 | 0 | 1 | 1 | 1 | 1 | 1 | 0 | 0 | 1 | 0 | 0 | 14.41 | 15.74 | 12.88 | 6.38 | 0 | FAD dependent oxidoreductase domain |
| AL2G00592 | OG0005130 | 2 | 2 | 4 | 2 | 2 | 0 | 1 | 1 | 1 | 2 | 1 | 1 | 1 | 1 | 1 | 1 | 0 | 0 | 0 | 0 | 7.22 | 1.28 | 4.84 | 1.75 | 1 | Dehydrogenase |
| AL2G00593 | OG0016610 | 2 | 2 | 4 | 1 | 0 | 0 | 0 | 0 | 0 | 1 | 0 | 1 | 1 | 0 | 0 | 0 | 0 | 0 | 0 | 0 | 0 | 0 | 0 | 0 | 0 | Leucoanthocyanidin dioxygenase |
| AL2G00594 | OG0019069 | 1 | 2 | 4 | 0 | 0 | 0 | 1 | 0 | 0 | 0 | 0 | 0 | 0 | 1 | 0 | 0 | 0 | 0 | 0 | 1 | 0.24 | 1.28 | 0.25 | 0.03 | 0 | NBS-LRR disease resistance protein |
| AL2G00595 | OG0018303 | 2 | 4 | 3 | 0 | 0 | 0 | 0 | 0 | 0 | 0 | 0 | 0 | 0 | 0 | 0 | 0 | 0 | 0 | 0 | 0 | 2.36 | 2.37 | 1.28 | 5.97 | 0 | NB-ARC domain |
| AL2G00596 | OG0003102 | 0 | 1 | 3 | 2 | 5 | 1 | 2 | 2 | 4 | 2 | 1 | 0 | 0 | 0 | 0 | 0 | 0 | 0 | 0 | 0 | 0 | 0 | 0 | 0 | 1 | MYB family transcription factor |
| AL2G00597 | OG0016610 | 2 | 2 | 4 | 1 | 0 | 0 | 0 | 0 | 0 | 1 | 0 | 0 | 0 | 0 | 0 | 0 | 0 | 0 | 0 | 0 | 0.12 | 0.13 | 0 | 0 | 0 | Leucoanthocyanidin dioxygenase |
| AL2G00598 | OG0020529 | 1 | 1 | 2 | 1 | 1 | 0 | 0 | 0 | 0 | 0 | 0 | 1 | 1 | 1 | 0 | 0 | 0 | 0 | 0 | 0 | 0.73 | 2.19 | 1.85 | 2.27 | 0 | G-patch domain |
| AL2G00599 | OG0000087 | 12 | 10 | 15 | 11 | 11 | 8 | 8 | 9 | 9 | 8 | 9 | 1 | 1 | 1 | 1 | 0 | 0 | 0 | 0 | 0 | 330.83 | 167.28 | 11.94 | 40.57 | 1 | Aquaporin protein |
| AL2G00600 | OG0001782 | 3 | 1 | 5 | 1 | 9 | 0 | 3 | 0 | 2 | 2 | 2 | 1 | 1 | 0 | 1 | 1 | 0 | 0 | 0 | 0 | 0 | 0 | 0 | 0 | 1 | Flavonol synthase |
| AL2G00601 | OG0001782 | 3 | 1 | 5 | 1 | 9 | 0 | 3 | 0 | 2 | 2 | 2 | 0 | 0 | 0 | 0 | 0 | 0 | 0 | 0 | 0 | 0.44 | 0.59 | 0.33 | 0.82 | 1 | Flavonol synthase |
| AL2G00602 | OG0030648 | 0 | 0 | 2 | 0 | 0 | 0 | 0 | 0 | 0 | 0 | 0 | 0 | 0 | 0 | 0 | 0 | 0 | 0 | 0 | 0 | 0.23 | 0.47 | 2.14 | 9.92 | 0 | Other |
| AL2G00603 | OG0003292 | 1 | 1 | 2 | 1 | 2 | 2 | 2 | 2 | 2 | 4 | 3 | 1 | 1 | 1 | 1 | 1 | 0 | 0 | 0 | 0 | 81.7 | 2.71 | 0.12 | 0 | 1 | Zinc finger protein |
| AL2G00604 | OG0006787 | 2 | 1 | 4 | 1 | 1 | 1 | 1 | 1 | 1 | 1 | 1 | 1 | 1 | 1 | 1 | 1 | 0 | 0 | 0 | 0 | 3.4 | 2.67 | 4.37 | 4.42 | 1 | YL1 nuclear protein C-terminal domain |
| AL2G00605 | OG0017372 | 1 | 1 | 2 | 1 | 1 | 0 | 1 | 0 | 1 | 1 | 1 | 1 | 1 | 1 | 1 | 1 | 0 | 0 | 0 | 0 | 2.76 | 3.11 | 2.23 | 1.62 | 0 | Other |
| AL2G00606 | OG0003073 | 3 | 2 | 5 | 7 | 4 | 0 | 1 | 1 | 0 | 0 | 0 | 1 | 1 | 1 | 0 | 0 | 0 | 0 | 0 | 0 | 6.71 | 0 | 0 | 0 | 0 | Dirigent |
| AL2G00607 | OG0003073 | 3 | 2 | 5 | 7 | 4 | 0 | 1 | 1 | 0 | 0 | 0 | 0 | 0 | 1 | 0 | 0 | 0 | 0 | 0 | 0 | 0.09 | 0 | 0 | 0 | 0 | Jacalin-like lectin domain |
| AL2G00608 | OG0003073 | 3 | 2 | 5 | 7 | 4 | 0 | 1 | 1 | 0 | 0 | 0 | 0 | 0 | 0 | 0 | 0 | 0 | 0 | 0 | 0 | 1.83 | 0 | 0 | 0 | 0 | Jacalin-like lectin domain |
| AL2G00609 | OG0025675 | 1 | 0 | 2 | 0 | 0 | 0 | 0 | 0 | 0 | 0 | 0 | 1 | 0 | 0 | 0 | 0 | 0 | 0 | 0 | 0 | 0 | 0 | 0 | 0 | 0 | CslF9 |
| AL2G00610 | OG0011683 | 1 | 1 | 2 | 1 | 1 | 1 | 1 | 1 | 1 | 1 | 1 | 1 | 1 | 1 | 1 | 1 | 0 | 0 | 0 | 0 | 22.14 | 20.25 | 25.54 | 29.27 | 0 | Glyoxalase family |
| AL2G00611 | OG0014656 | 1 | 1 | 2 | 1 | 1 | 1 | 1 | 1 | 1 | 0 | 1 | 1 | 1 | 1 | 1 | 1 | 0 | 0 | 1 | 1 | 7.43 | 12.33 | 9.21 | 2.22 | 0 | FAD dependent oxidoreductase domain |
| AL2G00612 | OG0005130 | 2 | 2 | 4 | 2 | 2 | 0 | 1 | 1 | 1 | 2 | 1 | 1 | 1 | 1 | 1 | 0 | 0 | 0 | 0 | 0 | 4.39 | 1.01 | 1.57 | 0.45 | 1 | Dehydrogenase |
| AL2G00613 | | | | | | | | | | | | | 0 | 0 | 0 | 0 | 0 | 0 | 0 | 0 | 0 | 0 | 0 | 0 | 0 | 0 | Ent-kaurene synthase |
| AL2G00614 | OG0005199 | 1 | 0 | 5 | 0 | 4 | 0 | 0 | 2 | 1 | 2 | 3 | 0 | 0 | 0 | 0 | 0 | 0 | 0 | 0 | 0 | 0 | 0 | 0 | 0 | 1 | Ent-kaurene synthase |
| AL2G00615 | OG0016610 | 2 | 2 | 4 | 1 | 0 | 0 | 0 | 0 | 0 | 1 | 0 | 1 | 1 | 1 | 0 | 0 | 0 | 0 | 0 | 0 | 0.07 | 0 | 0 | 0 | 0 | Leucoanthocyanidin dioxygenase |
| AL2G00616 | OG0022927 | 1 | 0 | 2 | 1 | 0 | 0 | 0 | 0 | 0 | 0 | 0 | 1 | 0 | 1 | 0 | 0 | 0 | 0 | 0 | 0 | 0 | 0 | 0 | 0 | 0 | Other |
| AL2G00827 | OG0016400 | 1 | 1 | 2 | 1 | 1 | 0 | 1 | 1 | 1 | 1 | 0 | 1 | 1 | 1 | 1 | 1 | 1 | 1 | 1 | 0 | 1.11 | 0.26 | 0.02 | 0 | 0 | Helicase conserved C-terminal domain |
| AL2G00828 | OG0000430 | 0 | 1 | 44 | 12 | 0 | 0 | 0 | 0 | 0 | 1 | 0 | 0 | 0 | 0 | 0 | 0 | 0 | 0 | 0 | 0 | 0 | 0.12 | 0 | 0 | 1 | Other |
| AL2G00829 | OG0000071 | 18 | 25 | 24 | 15 | 22 | 0 | 7 | 1 | 1 | 2 | 2 | 1 | 0 | 1 | 0 | 0 | 0 | 0 | 0 | 0 | 2.4 | 0.02 | 0 | 0 | 1 | Cytochrome P450 |
| AL2G00830 | OG0002252 | 2 | 2 | 3 | 2 | 3 | 1 | 2 | 2 | 2 | 2 | 4 | 1 | 1 | 1 | 1 | 1 | 1 | 1 | 1 | 0 1 | 4.11 | 2.27 | 0.9 | 0.18 | 1 | Glycosyl transferase |
| AL2G00831 | OG0001068 | 6 | 4 | 6 | 7 | 1 | 0 | 4 | 2 | 3 | 3 | 0 | 1 | 0 | 1 | 0 | 0 | 0 | 0 | 0 | 0 | 0 | 0 | 0 | 0 | 0 | 1-aminocyclopropane-1-carboxylate oxidase |
| AL2G00832 | OG0004542 | 8 | 1 | 2 | 1 | 3 | 0 | 1 | 1 | 1 | 1 | 0 | 1 | 1 | 1 | 1 | 0 | 0 | 1 | 1 | 0 | 9.66 | 0 | | 0 | 0 | Proline-rich protein |
| AL2G00833 | | | | | | | | | | | | | 1 | 1 | 1 | 1 | 0 | 0 | 0 | 0 | 0 | 35.7 | 22.23 | 10.34 | 1.58 | 0 | CESA8 cellulose synthase |
| AL2G00834 | OG0002979 | 2 | 2 | 3 | 2 | 2 | 1 | 2 | 2 | 2 | 2 | 3 | 0 | 0 | 0 | 0 | 0 | 0 | 0 | 0 | 0 | 92.21 | 34.75 | 16.33 | 6.69 | 1 | CESA2 cellulose synthase |
| AL2G00835 | OG0011192 | 1 | 1 | 2 | 1 | 1 | 0 | 1 | 1 | 1 | 1 | 2 | 1 | 1 | 1 | 0 | 0 | 0 | 0 | 0 | 0 | 0 | 0.18 | 0 | 0 | 0 | LTP family/seed storage |
| AL2G00836 | OG0014283 | 1 | 1 | 2 | 1 | 0 | 1 | 1 1 | 1 | 1 | 1 | 1 | 1 | 1 | 1 | 0 | 0 | 0 | 0 | 0 | 0 | 5.59 | 5.13 | 34.75 | 20.74 | 0 | Fruit protein PKIWI502 |
| AL2G00837 | OG0001684 | 2 | 2 | 3 | 2 | 1 | 1 | 2 1 | 3 | 3 | 2 | 8 | 1 | 1 | 1 | 1 | 0 | 0 | 1 | 1 | 1 | 32.14 | 24.83 | 19 | 16.53 | 1 | Zinc finger protein |

| Gene ID | Orthogroup | AAT | AER | ALO | AST | ATA | ATH | BDI | OSA | SBI | SIT | ZMA | AAT | AER | AST | ATA | BDI | OSA | SBI | SIT | ZMA | Roots | Salt-treated roots | Leaves | Salt-treated leaves | Expaned gene family | Gene description |
|---|---|---|---|---|---|---|---|---|---|---|---|---|---|---|---|---|---|---|---|---|---|---|---|---|---|---|---|
| AL2G03967 | OG0018505 | 0 | 1 | 1 | 0 | 1 | 1 | 1 | 1 | 1 | 1 | 1 | 1 | 1 | 1 | 1 | 1 | 1 | 1 | 1 | 1 | 153.33 | 84.22 | 78.06 | 148.95 | 0 | NADH dehydrogenase |
| AL2G03968 | OG0002782 | 2 | 2 | 2 | 2 | 2 | 3 | 2 | 2 | 2 | 2 | 2 | 0 | 1 | 0 | 1 | 1 | 1 | 1 | 1 | 0 | 45.91 | 51.99 | 33.06 | 21.61 | 0 | Transporter family protein |
| AL2G03969 | OG0016571 | 1 | 1 | 1 | 1 | 0 | 1 | 1 | 1 | 1 | 1 | 1 | 1 | 1 | 1 | 0 | 0 | 0 | 0 | 0 | 0 | 1.36 | 3.37 | 0.1 | 0 | 0 | Cytochrome P450 |
| AL2G03970 | OG0000982 | 4 | 3 | 4 | 3 | 3 | 1 | 3 | 3 | 6 | 3 | 4 | 1 | 1 | 1 | 0 | 0 | 0 | 0 | 0 | 0 | 27.74 | 16.26 | 2.65 | 0.12 | 0 | FAD dependent oxidoreductase domain |
| AL2G03971 | OG0014018 | 1 | 1 | 1 | 1 | 1 | 1 | 1 | 1 | 1 | 1 | 1 | 1 | 1 | 1 | 1 | 1 | 1 | 1 | 1 | 1 | 15.47 | 29.33 | 2.07 | 3.35 | 0 | Dehydration protein |
| AL2G03972 | OG0008930 | 1 | 1 | 1 | 1 | 1 | 1 | 1 | 2 | 2 | 1 | 1 | 1 | 1 | 1 | 1 | 1 | 1 | 1 | 1 | 1 | 2.67 | 1.77 | 1.28 | 0.89 | 0 | AMP-binding enzyme |
| AL2G03973 | OG0001379 | 2 | 3 | 4 | 2 | 2 | 2 | 3 | 3 | 3 | 3 | 5 | 1 | 1 | 1 | 0 | 1 | 1 | 1 | 1 | 1 | 135.75 | 162.29 | 77.23 | 94.3 | 1 | Cytidylyltransferase domain |
| AL2G03974 | OG0006807 | 2 | 1 | 2 | 1 | 1 | 1 | 1 | 1 | 2 | 1 | 2 | 1 | 1 | 1 | 1 | 1 | 1 | 1 | 1 | 0 | 1.14 | 2.13 | 34.58 | 61.87 | 0 | IrgB-like family |
| AL2G03975 | OG0006807 | 2 | 1 | 2 | 1 | 1 | 1 | 1 | 1 | 2 | 1 | 2 | 0 | 0 | 0 | 0 | 0 | 0 | 0 | 0 | 0 | 5.97 | 1.03 | 0 | 0 | 0 | IrgB-like family |
| AL2G03976 | OG0001227 | 5 | 4 | 6 | 3 | 0 | 0 | 3 | 1 | 4 | 3 | 5 | 1 | 1 | 0 | 0 | 0 | 0 | 0 | 0 | 0 | 1.14 | 2.58 | 0 | 0 | 1 | Plastocyanin-like domain |
| AL2G03977 | OG0014676 | 1 | 1 | 1 | 1 | 1 | 1 | 1 | 1 | 1 | 1 | 1 | 1 | 1 | 1 | 1 | 1 | 1 | 0 | 1 | 1 | 13.39 | 7.59 | 1.12 | 2.42 | 0 | Other |
| AL2G03978 | OG0003896 | 2 | 2 | 2 | 2 | 2 | 2 | 2 | 2 | 1 | 2 | 2 | 1 | 1 | 1 | 1 | 1 | 0 | 1 | 1 | 1 | 73.78 | 59.51 | 26.44 | 35.02 | 0 | Proteasome subunit |
| AL2G03979 | OG0011758 | 1 | 1 | 1 | 1 | 1 | 1 | 1 | 1 | 1 | 2 | 1 | 1 | 1 | 1 | 0 | 1 | 1 | 1 | 1 | 1 | 0.7 | 0.96 | 0.75 | 0.26 | 0 | PPR repeat domain |
| AL2G03980 | OG0001194 | 3 | 3 | 4 | 3 | 3 | 3 | 3 | 3 | 3 | 3 | 3 | 1 | 0 | 1 | 1 | 1 | 1 | 1 | 1 | 1 | 0.69 | 1.52 | 0.75 | 1.38 | 1 | CslD1 |

**Fig. S18.** Ten *CesA* and *Csl* gene clusters on six chromosomes of ALO. Conserved synteny at gene clusters across 11 species suggest role of secondary metabolite clusters toward evolution of specialized metabolites in plants. The position of the gene cluster is scaled based on chromosome size and its physical position on the chromosome. FPKM values suggest role of salt stress for *Csl* gene family evolution in ALO roots and leaves. The gene description colored as pink represents genes with functions associated with mixed-linkage (1,3;1,4)-β-glucan biosynthesis. **Fig. S18a.** *CesA* and *Csl* genes inserted gene-clusters identified on ALO02.

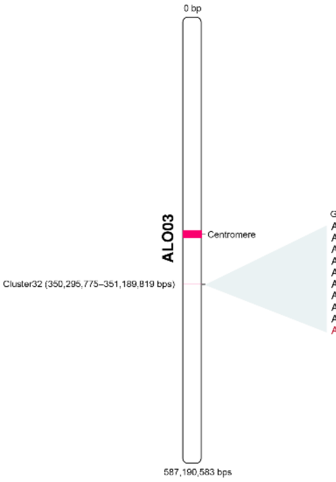

| | | Number of genes assigned to an orthogroup from a specific species | | | | | | | | | | | Prescence (1) or absence (0) of genes from a synteny relationship with ALO | | | | | | | | | | FPKM value of the gene in different samples | | | | Assignment (1) or not (0) |
|---|---|---|---|---|---|---|---|---|---|---|---|---|---|---|---|---|---|---|---|---|---|---|---|---|---|---|---|
| Gene ID | Orthogroup | AAT | AER | ALO | AST | ATA | ATH | BDI | OSA | SBI | SIT | ZMA | AAT | AER | AST | ATA | BDI | OSA | SBI | SIT | ZMA | Roots | Salt-treated roots | Leaves | Salt-treated rlaeves | Expanded gene family | Gene description |
| AL3G03047 | OG0004634 | 2 | 2 | 2 | 2 | 2 | 1 | 2 | 2 | 1 | 1 | 2 | 1 | 1 | 1 | 1 | 1 | 1 | 0 | 0 | 1 | 14.95 | 13.26 | 4.08 | 5.75 | 0 | Polygalacturonase |
| AL3G03048 | OG0001177 | 6 | 2 | 21 | 4 | 1 | 0 | 0 | 0 | 0 | 0 | 0 | 0 | 0 | 0 | 0 | 0 | 0 | 0 | 0 | 0 | 0 | 0 | 0 | 0 | 1 | Retrotransposon protein |
| AL3G03049 | OG0025291 | 1 | 0 | 1 | 1 | 0 | 0 | 0 | 0 | 0 | 0 | 0 | 0 | 1 | 1 | 1 | 1 | 1 | 0 | 1 | 1 | 0.19 | 0.19 | 7.1 | 3.78 | 0 | Pathogenesis-related protein |
| AL3G03050 | OG0000372 | 11 | 9 | 19 | 10 | 2 | 0 | 1 | 6 | 0 | 2 | 1 | 0 | 0 | 0 | 0 | 0 | 0 | 0 | 0 | 0 | 0 | 0 | 0 | 0 | 1 | Pathogenesis-related protein |
| AL3G03051 | OG0000372 | 11 | 9 | 19 | 10 | 2 | 0 | 1 | 6 | 0 | 2 | 1 | 0 | 0 | 0 | 0 | 0 | 1 | 0 | 0 | 0 | 20.09 | 0.51 | 0 | 0 | 1 | Pathogenesis-related protein |
| AL3G03052 | OG0000372 | 11 | 9 | 19 | 10 | 2 | 0 | 1 | 6 | 0 | 2 | 1 | 1 | 0 | 0 | 0 | 0 | 0 | 0 | 0 | 0 | 16.58 | 190.94 | 19.74 | 5.4 | 1 | Pathogenesis-related protein |
| AL3G03053 | OG0000072 | 1 | 0 | 113 | 0 | 3 | 0 | 0 | 0 | 0 | 0 | 0 | 0 | 0 | 0 | 0 | 0 | 0 | 0 | 0 | 0 | 0 | 0 | 0 | 0 | 0 | Retrotransposon protein |
| AL3G03054 | OG0000372 | 11 | 9 | 19 | 10 | 2 | 0 | 1 | 6 | 0 | 2 | 1 | 0 | 0 | 0 | 0 | 0 | 0 | 0 | 0 | 0 | 28.55 | 176.77 | 44.25 | 11 | 1 | Pathogenesis-related protein |
| AL3G03055 | OG0000372 | 11 | 9 | 19 | 10 | 2 | 0 | 1 | 6 | 0 | 2 | 1 | 0 | 0 | 0 | 0 | 0 | 0 | 0 | 0 | 0 | 64.37 | 252.24 | 15.2 | 2.65 | 1 | Pathogenesis-related protein |
| AL3G03056 | OG0012597 | 1 | 1 | 1 | 1 | 1 | 1 | 1 | 1 | 1 | 1 | 1 | 1 | 1 | 1 | 1 | 1 | 1 | 1 | 1 | 1 | 24.03 | 2.03 | 0.25 | 0.01 | 0 | CslD4 |

**Fig. S18b.** *Csl* genes inserted within gene-cluster identified on ALO03.

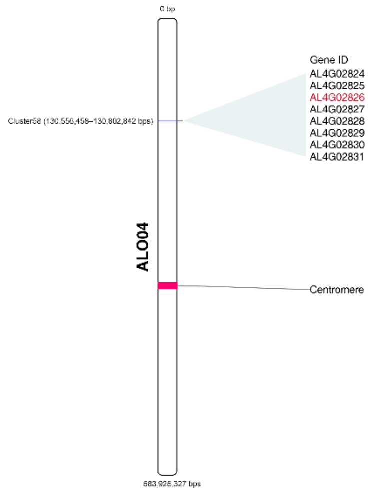

| Gene ID | Orthogroup | Number of genes asinged to an orthogroup from a specific species: AAT | AER | ALO | AST | ATA | ATH | BDI | OSA | SBI | SIT | ZMA | Prescence (1) or absence (0) of genes from a synteny relationship with ALO: AAT | AER | AST | ATA | BDI | OSA | SBI | SIT | ZMA | FPKM value of the gene in different samples: Roots | Salt-treated roots | Leaves | Salt-treated leaves | Assignment (1) or not (0): Expanded gene family | Gene description |
|---|---|---|---|---|---|---|---|---|---|---|---|---|---|---|---|---|---|---|---|---|---|---|---|---|---|---|---|
| AL4G02824 | OG0003823 | 2 | 2 | 3 | 2 | 1 | 1 | 2 | 4 | 1 | 1 | 2 | 1 | 1 | 1 | 1 | 1 | 1 | 1 | 1 | 0 | 2.73 | 3.29 | 0.21 | 4.32 | 1 | Zinc finger protein |
| AL4G02825 | OG0014479 | 1 | 1 | 1 | 1 | 1 | 1 | 1 | 1 | 1 | 1 | 1 | 1 | 1 | 1 | 1 | 1 | 1 | 1 | 1 | 1 | 15.74 | 22.14 | 11.52 | 19.87 | 0 | Other |
| AL4G02826 | OG0014581 | 1 | 1 | 1 | 1 | 2 | 0 | 1 | 1 | 1 | 1 | 1 | 1 | 1 | 1 | 1 | 1 | 1 | 1 | 1 | 1 | 14.96 | 3.86 | 0.59 | 7.44 | 0 | CslA7 |
| AL4G02827 | OG0000449 | 6 | 5 | 6 | 4 | 7 | 0 | 11 | 11 | 3 | 2 | 1 | 1 | 1 | 1 | 0 | 0 | 0 | 0 | 0 | 0 | 2.99 | 0.88 | 0.14 | 0.76 | 0 | Other |
| AL4G02828 | OG0023313 | 1 | 1 | 1 | 1 | 0 | 0 | 0 | 0 | 0 | 0 | 0 | 1 | 1 | 1 | 1 | 1 | 1 | 1 | 1 | 1 | 12.12 | 0.61 | 0 | 0 | 0 | Ribonuclease T2 family |
| AL4G02829 | OG0000530 | 7 | 7 | 23 | 9 | 1 | 0 | 0 | 4 | 0 | 0 | 0 | 0 | 0 | 0 | 0 | 0 | 0 | 0 | 0 | 0 | 0 | 0 | 0 | 0 | 1 | Other |
| AL4G02830 | OG0002366 | 2 | 3 | 4 | 2 | 1 | 3 | 2 | 2 | 2 | 1 | 3 | 0 | 0 | 0 | 0 | 0 | 0 | 0 | 0 | 0 | 0 | 0 | 0 | 0 | 0 | Gibberellin 2-beta-dioxygenase |
| AL4G02831 | OG0030684 | 0 | 0 | 2 | 0 | 0 | 0 | 0 | 0 | 0 | 0 | 0 | 0 | 0 | 0 | 0 | 0 | 0 | 0 | 0 | 0 | 0 | 0 | 0 | 0 | 0 | Gibberellin 2-beta-dioxygenase |

**Fig. S18c.** *Csl* genes inserted within gene-cluster identified on ALO04.

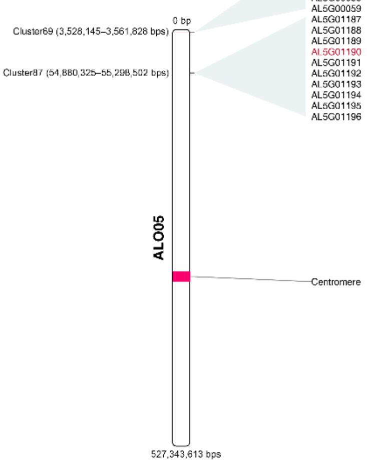

| | | Number of genes assigned to an orthogroup from a specific species | | | | | | | | | | | Presence (1) or absence (0) of genes from a synteny relationship with ALO | | | | | | | | | | FPKM value of the gene in different samples | | | | Assignment (1) or not (0) |
|---|---|---|---|---|---|---|---|---|---|---|---|---|---|---|---|---|---|---|---|---|---|---|---|---|---|---|---|
| Gene ID | Orthogroup | AAT | AER | ALO | AST | ATA | ATH | BDI | OSA | SBI | SIT | ZMA | AAT | AER | AST | ATA | BDI | OSA | SBI | SIT | ZMA | Roots | Salt-treated roots | Leaves | Salt-treated leaves | Expanded gene family | Gene description |
| AL5G00054 | OG0023206 | 1 | 1 | 1 | 1 | 0 | 0 | 0 | 0 | 0 | 0 | 0 | 1 | 1 | 1 | 0 | 0 | 0 | 0 | 0 | 0 | 0.17 | 0.31 | 0.19 | 0 | 0 | DNA methyltransferase protein |
| AL5G00055 | OG0003005 | 2 | 2 | 2 | 1 | 2 | 0 | 3 | 1 | 6 | 2 | 2 | 1 | 1 | 1 | 0 | 0 | 1 | 0 | 0 | 0 | 0 | 0 | 0.35 | 0 | 0 | CslF3 |
| AL5G00056 | OG0000182 | 14 | 12 | 16 | 10 | 10 | 0 | 5 | 5 | 5 | 4 | 2 | 0 | 0 | 0 | 0 | 0 | 0 | 0 | 0 | 0 | 5.28 | 4.01 | 5.08 | 44.33 | 1 | SAM dependent carboxyl methyltransferase |
| AL5G00057 | OG0000466 | 5 | 7 | 5 | 6 | 12 | 0 | 6 | 5 | 3 | 4 | 2 | 0 | 0 | 0 | 0 | 0 | 0 | 0 | 0 | 0 | 1.46 | 1.97 | 1.83 | 0.98 | 0 | Other |
| AL5G00058 | OG0004236 | 3 | 4 | 3 | 3 | 1 | 0 | 1 | 1 | 2 | 1 | 1 | 1 | 1 | 1 | 0 | 0 | 0 | 0 | 0 | 0 | 0.13 | 0.23 | 0 | 0 | 0 | Pathogenesis-related protein |
| AL5G00059 | OG0004222 | 1 | 1 | 1 | 1 | 4 | 2 | 1 | 2 | 1 | 2 | 4 | 1 | 1 | 1 | 1 | 0 | 1 | 1 | 1 | 1 | 11.64 | 19.58 | 15.05 | 14.38 | 0 | AAA family ATPase |
| AL5G01187 | OG0010371 | 1 | 1 | 1 | 1 | 2 | 1 | 1 | 1 | 1 | 1 | 1 | 1 | 1 | 1 | 0 | 1 | 1 | 1 | 1 | 1 | 10.15 | 6.19 | 0.13 | 0.8 | 0 | GDSL-like lipase/acylhydrolase |
| AL5G01188 | OG0013041 | 1 | 1 | 1 | 1 | 1 | 0 | 1 | 2 | 1 | 1 | 1 | 1 | 1 | 1 | 1 | 1 | 1 | 1 | 1 | 1 | 0 | 0 | 0 | 0 | 0 | Soluble starch synthase |
| AL5G01189 | OG0006495 | 2 | 2 | 2 | 2 | 1 | 1 | 1 | 1 | 1 | 1 | 1 | 1 | 1 | 1 | 0 | 0 | 0 | 0 | 0 | 0 | 17.51 | 8.91 | 4.65 | 9.88 | 0 | Other |
| AL5G01190 | OG0001102 | 3 | 3 | 2 | 3 | 4 | 5 | 3 | 3 | 3 | 3 | 3 | 1 | 1 | 1 | 0 | 1 | 1 | 1 | 1 | 1 | 2.4 | 1.66 | 2.21 | 1.4 | 0 | CslA3 |
| AL5G01191 | OG0000370 | 5 | 5 | 6 | 5 | 5 | 11 | 5 | 4 | 5 | 5 | 5 | 1 | 0 | 1 | 1 | 1 | 0 | 1 | 1 | 1 | 0.37 | 2.08 | 7.27 | 3.8 | 1 | Myosin |
| AL5G01192 | OG0020301 | 3 | 0 | 1 | 1 | 0 | 0 | 1 | 0 | 0 | 0 | 0 | 1 | 0 | 1 | 0 | 0 | 0 | 0 | 0 | 0 | 0 | 3.85 | 590.04 | 515.95 | 0 | Dirigent |
| AL5G01193 | OG0019177 | 1 | 1 | 1 | 1 | 1 | 0 | 1 | 1 | 0 | 1 | 0 | 1 | 1 | 1 | 1 | 1 | 1 | 0 | 1 | 0 | 1.12 | 1.65 | 0.32 | 0.22 | 0 | Pentatricopeptide |
| AL5G01194 | OG0001390 | 5 | 5 | 6 | 5 | 2 | 0 | 2 | 3 | 2 | 1 | 1 | 1 | 1 | 1 | 1 | 1 | 0 | 0 | 0 | 0 | 19.22 | 45.37 | 5.07 | 4.46 | 0 | Cytokinin-O-glucosyltransferase |
| AL5G01195 | OG0001793 | 3 | 4 | 5 | 3 | 3 | 2 | 2 | 1 | 2 | 1 | 2 | 1 | 1 | 1 | 1 | 1 | 1 | 1 | 1 | 1 | 77.56 | 54.92 | 36.87 | 46.36 | 1 | Other |
| AL5G01196 | OG0017258 | 1 | 1 | 1 | 1 | 1 | 0 | 1 | 0 | 1 | 2 | 1 | 1 | 1 | 1 | 1 | 1 | 1 | 0 | 1 | 0 | 1.57 | 1.32 | 3.19 | 3.88 | 0 | Other |

**Fig. S18d.** *Csl* genes inserted within gene-clusters identified on ALO05.

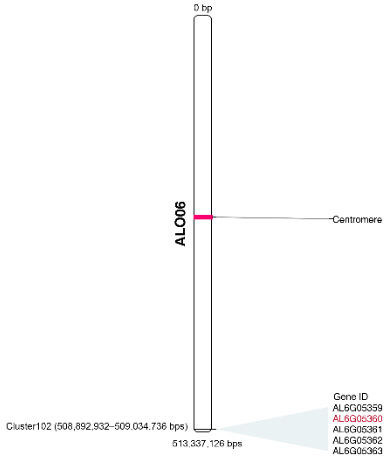

| | | Number of genes assigned to an orthogroup from a specific species | | | | | | | | | | | Presence (1) or absence (0) of genes from a synteny relationship with ALO | | | | | | | | | FPKM value of the gene in different samples | | | | Assignment (1) or not (0) | |
|---|---|---|---|---|---|---|---|---|---|---|---|---|---|---|---|---|---|---|---|---|---|---|---|---|---|---|---|
| Gene ID | Orthogroup | ATA | AER | ALO | AST | ATA | ATH | BDI | OSA | SBI | SIT | ZMA | ATA | AER | AST | ATA | BDI | OSA | SBI | SIT | ZMA | Roots | Salt-treated roots | Leaves | Salt-treated leaves | Expanded gene family | Gene description |
| AL6G05359 | OG0019175 | 2 | 2 | 2 | 2 | 0 | 0 | 0 | 0 | 0 | 0 | 0 | 0 | 0 | 0 | 0 | 0 | 0 | 0 | 0 | 0 | 0.76 | 0.18 | 0 | 0 | 0 | Jacalin-like lectin domain |
| AL6G05360 | OG0001474 | 4 | 2 | 7 | 6 | 2 | 1 | 2 | 2 | 2 | 2 | 1 | 1 | 0 | 1 | 0 | 0 | 0 | 0 | 0 | 0 | 16.57 | 2.84 | 0.02 | 0 | 0 | CslE6 |
| AL6G05361 | OG0023429 | 1 | 1 | 1 | 1 | 0 | 0 | 0 | 0 | 0 | 0 | 0 | 1 | 0 | 1 | 0 | 0 | 0 | 0 | 0 | 0 | 43.91 | 10.21 | 0 | 0 | 0 | Glycosyl transferase |
| AL6G05362 | OG0000346 | 8 | 13 | 12 | 9 | 3 | 0 | 3 | 2 | 5 | 6 | 2 | 1 | 0 | 1 | 0 | 0 | 0 | 0 | 0 | 0 | 0.1 | 0.15 | 0 | 0 | 0 | Oxidoreductase |
| AL6G05363 | OG0000947 | 4 | 3 | 6 | 5 | 1 | 1 | 1 | 6 | 7 | 2 | 2 | 1 | 0 | 1 | 0 | 0 | 0 | 0 | 0 | 0 | 7.17 | 1.79 | 0 | 0 | 0 | UDP-glucoronosyl/UDP-glucosyl transferase |

**Fig. S18e.** *Csl* genes inserted within gene-cluster identified on ALO06.

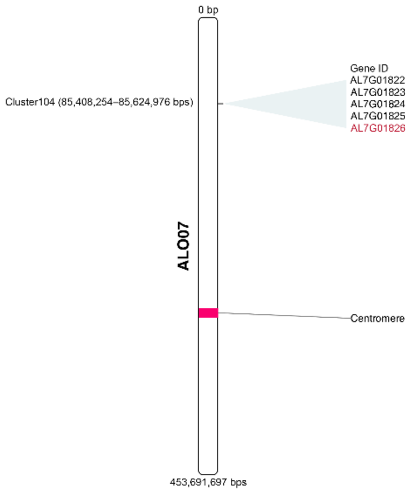

| Gene ID | Orthogroup | Number of genes assigned to an orthogroup from a specific species: AAT | AER | ALO | AST | ATA | ATH | BDI | OSA | SBI | SIT | ZMA | Presence (1) or absence (0) of genes from a synteny relationship with ALO: AAT | AER | AST | ATA | BDI | OSA | SBI | SIT | ZMA | FPKM value of the gene in different samples: Roots | Salt-treated roots | Leaves | Salt-treated leaves | Assignment (1) or not (0): Expanded gene family | Gene description |
|---|---|---|---|---|---|---|---|---|---|---|---|---|---|---|---|---|---|---|---|---|---|---|---|---|---|---|---|
| AL7G01822 | OG0012666 | 1 | 1 | 1 | 1 | 1 | 0 | 1 | 1 | 1 | 1 | 2 | 1 | 1 | 1 | 1 | 1 | 1 | 1 | 1 | 1 | 48.81 | 17.25 | 3.68 | 7.45 | 0 | Protein kinase family |
| AL7G01823 | OG0001091 | 6 | 7 | 5 | 6 | 4 | 0 | 5 | 1 | 0 | 1 | 0 | 1 | 1 | 1 | 0 | 0 | 0 | 0 | 0 | 0 | 12.64 | 7.25 | 1.83 | 1.39 | 0 | Gibberellin receptor GID1L2 |
| AL7G01824 | OG0000258 | 6 | 4 | 6 | 6 | 8 | 2 | 6 | 5 | 10 | 12 | 8 | 1 | 1 | 1 | 1 | 0 | 1 | 1 | 1 | 0 | 23.3 | 72.35 | 0.54 | 1.01 | 0 | Glucosyl transferase |
| AL7G01825 | OG0018443 | 1 | 1 | 1 | 0 | 0 | 0 | 1 | 1 | 1 | 1 | 2 | 1 | 1 | 0 | 0 | 0 | 0 | 0 | 0 | 0 | 1.43 | 1.59 | 0 | 0 | 0 | Membrane protein |
| AL7G01826 | OG0009492 | 1 | 1 | 2 | 2 | 1 | 1 | 1 | 1 | 1 | 1 | 1 | 1 | 1 | 1 | 1 | 0 | 1 | 1 | 1 | 1 | 28.25 | 21.27 | 8.97 | 22.76 | 0 | Glycosyl transferase |

**Fig. S18f.** *Csl* genes inserted within gene-cluster identified on ALO07.

>AL1G02887.t1
MESGAHHPCAACGDDAHAACRACSYALCRACLDEDVAEGRTACARCAGEYAALDTAHGKESAVEE
EEVLENQLAAEGLRGRVTMATQLTDRQDQVSHARTVSSMSGVGSELNDESGKPIWKNRVDSWKEK
KNEKKASAKKAAAKAQVPPVEEQIMDEKDLTDAYEPLSKIIPISKNKLTPYRAVIIMRLVVLGLFFHYRIT
NPVDSAFGLWLTSVICEIWFGFSWILDQFPKWCPVNRETYVDRLIARYGDGEDSGLAPVDFFVSTVD
PLKEPPLITANTVLSILAVDYPVEKVSCYVSDDGASMLTFESLAETAEFARRWVPFCKMYSIEPRTPEF
YFSQKIDYLKDKIHPSFVKERRAMKRDYEEYKVRINALVAKAQKTPEEGWIMPDGTQWPGNNSRDHP
GMIQVFLGETGARDYDGNELPRLVYVSREKRPGYQHHKKAGAMNALVRVSAVLTNAPYILNLDCDHY
VNNSKAVREAMCFMMDPSVGRDVCYVQFPQRFDGIDRSDRYANRNVVFFDVNMKGLDGIQGPVYV
GTGCCFYRQALYGYGPPSLPALPKSSACSWCCCCCPKKKVERTEKEMHRDSRREDLESAIFNLREI
DNYDEYERSMLISQMSFEKSFGQSSVFIESTLMENGGLPESADPSTLIKEAIHVISCGYEEKTEWGKE
LGWIYGSVTEDILTGFKMHCRGWRSIYCMPIRPAFKGSAPINLSDRLHQVLRWALGSVEIFFSRHCPL
WYGYGGGRLKWLQRLSYINTIVYPFTSVPLVAYCCLPAICLLTGKFIIPILSNAATIWFLGLFTSIILTSVL
ELRWSGIGIEDWWRNEQFWVIGGVSAHLFAVFQGILKMVIGLDTNFTVTSKAAEDGDFGELYVFKWT
TVLIPPTTILVLNLVGVVAGFSDALNSGYESWGPLFGKVFFAMWVIMHLYPFLKGLMGRQNRTPTIVIL
WSVLLASVFSLLWVKIDPFITNAETTNTGACSSIDC
>AL1G05017.t1
MGFAYYAFPMLCYATVPQLCFFHGGDATSFPEASSLCFAALFVSSLLQHLIEVSVAKRELAVRTWWN
EQRFWALNAVMGQLFAFLAVALDLVGAGGQAVDFGLTSKASDHEDLYRDSVFDFSGCSTVLLPATTL
LTARQRGGPASHPRPPAPAILLLPSSRRPCPPPTCPRLATTWPVLPPQRPVRSPPLSAEPPAPRLCR
SSTSTVSLASLALPPRLGPPPASRPVAPPRAQPSSAFAAFGPTSPSERLRPSSSTLMPLPRLLPSGTP
PPVGLILSPPASPTATAAIPPSAPPSLPPTDPPPPSPPRRRLAPVDPSARGPQRRPLCHGLPAASAGV
LQRTHTRPPASAARDLRRRRPLCPSVAEVLQPLPVLAVLLRSQMPR
>AL1G05019.t1
MAAPLQDAAPLQLHTVEVDQPLAAVNRLLAGLHLALAATAIAHRGVHDILLAADLVLLFLWLLSQAPML
RPVSRAAFPGRLSRPALPAVDVMVVTADPDKEPAAKVMNTVISAMALDYPGGRLTVYLSDDAGSPLT
LLAARKAYAFARAWVPFCRKYGVQCPCPERFFAGNDQLDGGGGVDHDRELAGDKIRVKNMYETFR
KGIDEATKDAALSGSCTRAARQDHDAHVEIIADKQDSNSDGDEDAMPSLVYVSREKRPAWAHHFKA
GALNVLLRVSSLMSNAPYVLVLDCDMYCNARSSALEAMCFHLDRRRDGLAFVQFPQMFHNLSGNDI
YANELRSIFSTRWKGLDGLRGPILSGTGFCLRRDAVYGARPSSSQDPFSSMDVGELKARFGHSNGH
LASLRRTAGSTTIVARDGVLPQDAALVASCEYETGTRWGEEVGFLYQSVVEDYFTGYRQLYCRGWT
SVYCYPAPSRPAPFLGSVPTNLNDVLVQNKRWMSGMLAVGLSRHCPLGSALAVSLPQSMGFAYYAF
MALYAFPVLCYATVPQLRFFHAGGASSFPQASSLWFTAVFVSSLLQHLIEVSVAKRSLAVRTWWNEQ
RFWALNAVTGQLFACLAVALDLVGAGGRAVDFDLTSKASDDGGLYRDGEFDFAGCSTVLLPATTLCL
LNDAALVGGVWKMVTSGGSWGDTVSVELFLLCYVAALSYPLLQGMFLRRDPARVPAWITAMSVGM
AATLLSLFG
>AL2G00588.t1
MFFDATLLGLNGIQGPSFIGTGCMFRRAALYGADPPRWRHDDAKAKVLHMPTIFGKSTAFINSVPAAA
NHQERSLASPTTIVDEIELASVMTGKYEDGTEWGNDVGWVYNIATEDVVTGFRLHRAGWRSTYCAM
EPNAFRGTGPINLTARLYQILRWSGGSLEMFFSRFCPLLAGRRIHPVQRIAYFNMTMTIYPFYIQRLFQ
TFAVFVAVVIATVEMIGMVEIKWAGLTLLDWFRSEQLYIIGTTGVYPTAMLHVLLRFHGLKGVSFKVTA
KKLTGGARERLAELYDVKWVPLLVPTVVVMAVNVVSIGVAAGKAIAGRWSSAQVAWAATGLLFNMW
MLLLLYPFALGIMGRWSKRPYILFVLLAAVRATTASVYVAIARSM
>AL2G00589.t1
MTIYPVSRFFLVMYDLYPVMWLLHGQFYIQRPFQTFAVFVAVVIATAELIGMVEIKWAGLTLLDWFRSE
QLYIIGTTGVYPTAMLHVLLRFLGLKGAYYRVDPEHEEKANEVFEIMARKQVRQMIYDAKNIAISHYYA
EQGKRKLKSHIVAEKLGDKVKVEDLLAVAANDGVDIPDVEVWQQARKKKDPKPNEDPYGSTIENPLR
SVAEGDDPDVPEQLNDNGGEGGVHARRSEVSAALGWCSAHWASPRSKCVMPSWKLEAPASPSLE
LGRQPWRVASIPGGVDELAGVKIGGDGGFRRALGEDGGALWLARMEDDADDNSGEVFLTGSARRR
QYFSPAQRDATDAQEEEEVGVLQPEFAGARSGRSGDQSSSMACEVCGAGQRTEVRTVAAA>>AL2
G00609.t1
MFFDATLLGLNGIQGPSFIGTGCMFRRAALYGADPPRWRHDDAKAKVLHMPTIFGKSTAFINSVPAAA
NHQERSLASPTTIVDEIELASVMTGKYEDGTEWGNDVGWVYNIATEDVVTGFRLHRAGWRSTYCAM
EPNAFRGTGPINLTARLYQILRWSGGSLEMFFSRFCPLLAGRRIHPVQRIAYFNMTMTIYPFYIQRLFQ
TFAVFVAVVIATVEMIGMVEIKWAGLTLLDWFRSEQLYIIGTTGVYPTAMLHVLLRFLGLKGVSFKVTAK
KLTGGARERLAKLYDVQWVPLLVPTVVVMAVNVVSIGVAAGKAIAGR
>AL2G00833.t1
MDGDAKSGRRGAAQVCQICGDGVGVTAEGEVFAACDVCAFPVCRPCYEYERKDGTQACPQCKTKY
KRHKGSPAIRGEEGDDTDGDEGSDFNYPASGTEDQKQKVADRMRSWRMSTGGGGNVGHPKYDS
GEIGLSKYDSGEIPRGYIPSVTNSQISGEIPGASPDHHMMSPTGNIGKRVPFPYVNHSPNPSREFSGS
VGNVAWKDRVDGWKMKQDKGAVPMTNGSSIAPSEGRGAADDASTEYNMEDALLNDETRQPLSRK
VPIPSSRINPYRMVICLRLVVLSIFLHYRITNPVRNAYPLWLLSVICEIWFAMSWILDQFPKWNPINRETY

LDRLALRYDREGEPSQLAAVDIFVSTRPLKEPPIVTANTVLSILAVDYPVDKLSCYVSDDGASMLTFDA
LAETSEFARNGYHLLRSMKLNPELLSGTSPRKLIT
>AL2G00834.t1
MKREYEEFKIRINGLVAKAIKVPEEGWIMQDGTPWPGNNTRDHPGMIQVFLGHSGGLDTEGNELPRL
VYVSREKRPGFQHHKKAGAMNALVRVSAVLTNGQYMLNLDCDHYINNSKAIREAMCFLMDPNLGRG
VCYVQFPQRFDGIDTNDRYANRNTVFFDINLRGLDGIQGPVYVGTGCVFNRTAIYGYEPPIKPKKGGF
LSSLCGGKKKASKSKRKSSDKKKSNKHVDSSVPVFNLEDIEEGVEGAGFDDEKSVLMSQMSLEKRF
GQSAAFVASTLMEYGGVPQSSTPESLLKEAIHVISCGYEDKSEWGTEIGWIYGSVTEDILTGFKMHAR
GWRSIYCMPKRPAFKGSAPINLSDRLNQVLRWALGSVEILFSRHCPLWYGYGGRLKFLERFAYINTTI
YPLTSIPLLVYCILPAICLLTGKFIMPEISNLASIWFISLFISIFATGILEMRWSGVGIDEWWRNEQFWVIG
GISAHLFAVFQGLLKVLAGIDTNFTVTSKANDEDGDFAELYMFKWTTLLIPPTTILIINMVGVVAGTSYAI
NSGYQSWGPLFGKLFFAFWVIVHLYPFLKGLMGRTNRTPTIVIVWAILLASIFSLLWVRIDPFTTRVAG
PNIQTCGINC
>AL2G00843.t1
MMSPTGNIGKRVPFPYVNHSPNPSREFSGSVGNVAWKDRVDGWKMKQDKGAVPMTNGSSIAPSE
GRGAADDASTEYNMEDALLNDETRQPLSRKVPIPSSRINPYRMVICLRLVVLSIFLHYRITNPVRNAYP
LWLLSVICEIWFAMSWILDQFPKWNPINRETYLDRLALRYDREGEPSQLAAVDIFVSTVDPLKEPPIVT
ANTVLSILAVDYPVDKLSCYVSDDGASMLTFDALAETSEFARKWVPFVKKYEIEPRAPEWYFSQKIDY
LKDKVQPSFVKERRAMKREYEEFKIRINGLVAKAIKVPEEGWIMQDGTPWPGNNTRDHPGMIQVFLG
HSGGLDTEGNELPRLVYVSREKRPGFQHHKKAGAMNALVRVSAVLTNGQYMLNLDCDHYINNSKAI
REAMCFLMDPNLGRGVCYVQFPQRFDGIDTNDRYANRNTVFFDINLRGLDGIQGPVYVGTGCVFNR
TAIYGYEPPIKPKKGGFLSSLCGGKKKASKSKRKSSDKKKSNKHVDSSVPVFNLEDIEEGVEGAGFDD
EKSVLMSQMSLEKRFGQSAAFVASTLMEYGGVPQSSTPESLLKEAIHVISCGYEDKSEWGTEIGWIY
GSVTEDILTGFKMHARGWRSIYCMPKRPAFKGSAPINLSDRLNQVLRWALGSVEILFSRHCPLWYGY
GGRLKFLERFAYINTTIYPLTSIPLLVYCILPAICLLTGKFIMPEISNLASIWFISLFISIFATGILEMRWSGV
GIDEWWRNEQFWVIGGISAHLFAVFQGLLKVLAGIDTNFTVTSKANDEDGDFAELYMFKWTTLLIPPT
TILIINMVGVVAGTSYAINSGYQSWGPLFGKLFFAFWVIVHLYPFLKGLMGRTNRTPTIVIVWAILLASIF
SLLWVRIDPFTTRVAGPNIQTCGINC
>AL2G00965.t1
MSASGGMVAGSHNRNEFVTIRRDGDAPAPGKQAKGAAVCQICGDTVGVSATGDVFVACNECAFPV
CRPCYEYERKEGNQCCPQCKTRYKRLKGSPRVQGDEEEEDVDDLDNEFNYKQDNGKGPEWQLQG
PGEDIDLSSSSRHEPHHRIPRLTSGHQISGEIPDASPDRHSIRSPTSSYVDPSVPVPVRIVDPSKDLNS
YGINSVDWKERVESWRVKQDKSMMQVANKYPDARGGGDMEGTGSNGEDMQMVDDARLPMSRIV
PIPANQLNLYRIVIILRLIILCFFFQYRVSHPVRDAYGLWLVSVICEIWFALSWLLDQFPKWYPINRETYL
DRLALRYDREGEPSQLAPIDIFVSTVDPLKEPPLITANTVLSILAVDYPVDKVSCYVSDDGSAMLSFESL
SETAEFARKWVPFCKKHSIEPRAPEFYFAQKIDYLKDKIQPSFVKERRSMKREYEEFKIRINALVAKAQ
KVPEEGWTMADGTAWPGNNPRDHPGMIQVFLGHSGGLDTEGNELPRLVYVSREKRPGFQHHKKA
GAMNALIRVSAVLTNGAYLLNVDCDHYFNSSKALREAMCFMMDPALGRKTCYVQFPQRFDGIDLHD
RYANRNIVFFDINMKGLDGIQGPMYVGTGCCFNRQALYGYDPVLTEADLEPNIIVKSFCGGRKKKNKS
YMDNKNRMMKRTESSAPIFNMEDIEEGIEGYDDERSMLMSQKKLEKRFGQSPIFTASTFMTQGGIPP
STNPASLLKEAIHVISCGYEDKTEWGKEIGWIYGSVTEDILTGFKMHTRGWISIYCMPSRPCFKGSAPI
NLSDRLNQVLRWALGSVEILFSRHCPIWYNHGGRLKFLERMAYINTIVYPITSIPLIAYCVLPAICLLTNK
FIIPEISNYAGMFFILMFASIMATGILELQWSGVGIEDWWRNEQFWVIGGTSAHLFAVFQGLLKVLAGID
TNFTVTSKANDEDGDFAELYVFKWTSLLIPPTTVLVINMVGMVAGISYAINSGYQSWGPLFGKLFFSIW
VILHLYPFLKGLMGKQNRTPTIVIVWSILLASIFSLLWVKIDPFISDTQKAVAMGQCGVNC>>AL2G0316
6.t1
MDTGEPKAKACRACGDDVGTREDGSPFVACAECGFPVCRPCYEYERSDGTQCCPQCNTRYKRHK
GCPRAEGDQEDGEMDDLEDEFQAKSPKKPHDPVPFDVYSENGEQPPHKWRPGGPAMSSFGGSVA
GKDLETEREMEGSMEWKDRIDKWKTKQEKRGKLNSDEDDDEKNEDEYMLLAEARQPLWRKLPVPS
SMINPYRIVIVVRLVVLCFFLRFRIMTPANDAIPLWLVSVICELWFALSWILDQLPKWSPVTRETYLDRL
ALRYDREGEPSRLSPIDFFVSTVDPLKEPPIITANTVLSILAVDYPVDRNSCYVSDDGASMLCFDTLSE
TAEFARRWVPFCKKFAIEPRAPEFYFSQKIDYLKDKVQPTFVKERRAMKREYEEFKVRINGLVAKAEK
KPEEGWVMQDGTPWPGNNTRDHPGMIQVYLGSQGALDVEGHELPRLVYVSREKRPGHNHHKKAG
AMNALVRVSAVLTNAPFILNLDCDHYVNNSKAVREAMCFLMDPQLGKKLCYVQFPQRFDGIDLHDRY
ANRNVVFFDINMKGLDGIQGPVYVGTGCVFNRQALYGYDPPRAEKRPKMTCDCWPSWCCCCCCFG
GGNNKHRKSKDKKGGDGEEEPRRGLLGFYKKRGSKKDKLGGGPKKGSYRKHQRGYELEEIEEGIE
GYDELDRSSLMSQKSFQKRFGQSPVFIASTLVEDGGLPQGAAADPASLIKEAIHVISCGYEEKTEWGK
EIGWIYGSVTEDILTGFKMHCRGWKSVYCTPTRPAFKGSAPINLSDRLHQVLRWALGSVEIFMSRHC
PLWYAYGGRLKWLERFAYTNTIVYPFTSIPLIAYCTIPAVCLLTGKFIIPTLNNLASIWFIALFMSIIATGVL
ELRWSGVSIEDWWRNEQFWVIGGVSAHLFAVFQGFLKVLGGVDTNFTVTSKAAADEADAFGDLYLF
KWTTLLIPPTTLIIINMVGIVAGVSDAVNNGYGSWGPLFGKLFFSFWVIVHLYPFLKGLMGRQNRTPTIV
VLWSVLLASIFSLVWVRIDPFIAKSKGPILKPCGVQC

>AL2G03980.t1
MGSKGILKNSGSGKPPTVPTSGPQVVFGRRTESGRFISYSRDDLDSEISSVDFQDYHVHLPMTPDNQ
PMEDGKADEQYVSSSLFTGGFNSVTRAHVMDKQGPDTEMGRRGPKGSTCMVEGCDSKIMRNGRG
EDILPCECDFKICVDCFTDAVKGGGSVCPGCKELYKYTEWEEVLSNSNNEFTRALSLPHGPGGKMER
RLSLVKQSTMNTNQSGEFDHNRWLFETKGTYGYGNAIWPDDNVDDDAGNGMQGHPKELMTKPWR
PLTRKLKIPAGVISPYRLLVLIRLVALAFFLMWRIKHVNNDAIWLWGMSIVCELWFAFSWVLDQLPKLC
PINRATDLSVLKEKFETPTPNNPTGKSDLPGIDIFVSTADPEKEPVLVTANTILSILAVDYPVDKLACYVS
DDGGALLTFEAMAEAASFANLWVPFCRKHEIEPRNPDSYFNLKRDPFKNKVKADFVKDRRRVKREY
DEFKVRVNGLPDAIRRRSDAYHAREEIQAMNLQREKIKAGGDEQFQPVKIPKATWMADSTHWPGTW
LHSSQDHARGDHAGIIQVMLKPPRDMPTYGNIEKSSLDFSAVDTRLPMLVYMSREKRPGYDHNKKA
GAMNALVRASAIMSNGPFILNLDCDHYVYNSKAFREGMCFMMDRGGDRLCYVQFPQRFEGIDPSDR
YANHNTVFFDINMRALDGLQGPVYVGTGCLFRRIALYGFDPPRSKDHSPGFCGCCLPRHRKTSASNA
NPEETMALRMGDFDGDSMNLATFPKKFGNSSFLIDSIPVAEFQGRPLADHPSVKNGRPPGALTIPREI
LDASIVAEAISVVSCWYEEKTEWGIRVGWIYGSVTEDVVTGYRMHNRGWKSVYCVTQRDAFRGTAPI
NLTDRLHQVLRWATGSVEIFFSRNNALLASSKMKVLQRIAYLNVGIYPFTSIFLIVYCFLPALSLFSGQFI
VQTLNVTFLVYLLIITITLCLLAMLEIKWSGIALEEWWRNEQFWLIGGTSAHLAAVMQGLLKVIAGIEISF
TLTSKTVSDDVDDEFADLYEVKWTSLMIPPLTIIMVNLVAIAVGFSRTIYSTIPQWSKLLGGVFFSFWVL
AHLYPFAKGLMGRRGRTPTIVYVWAGLVSITISLLWIAINPPSSAANSQLGGGSFTFP
>AL3G01091.t1
METSESRRLFATESFGGRAVYRLHAVTVAAGILLVLYYRATHVPAAGEGRAAWLGMLAAELWYSAY
WVLTQSVRWSPVRRRPFIERLAARFGERLPCVDILVCTADPKSEPPSLVISTILSLMAYDYPAEKLSVY
LSDDGGSVLTFYAMWEASLFAKHWIPFCKRYNIEPRSPTAYFLESDGLQDLCTPKEWSFIKDTYDEM
TERIDTTVMLEKIPEEIKAKHKGFDEWNLEINSKDHQPIVQILIDGKDQNAVDNEGNVLPTLVYVAREKR
PQLNHNFKAGAMNSLIRVSSVISNSPVIMNVDCDMYSNNKDTIRDALCFFLDEEMGHKIGFVQYPQNY
NNMTKNDLYGNSMKAINKVELRGLDSAGGPLYIGTGCFHRREILCGRRFTEDYKEDWNKGIKYITQE
NIYDTEEKAKSLATCTCEKDTQWGDEIGLKYGSPVEDVITGLAIHCRGWGSVCHNPTRPAFLGVGPS
TLAQTLLQHKRWSEGNFTNFLSKYCPFLFGHGKIKLHHQMGYCIYGLWAPNSLATLYYVTIPSLSLLK
GTLLFPEANKDATSYRYKSIKFWDMISTLYCRDRATGAGARTANESAVEMAEENANTTVNNKDANSS
TQADEDRPKKRYRSDDSIATMLGDKLDNFTAAFKADVLEPPPKPASPEEIWAVLDGIPDLEDD
>AL3G01092.t1
METTPESRRLFATESLGGRGPHRLHAVTVAAGILLVLYYRATHVPAAGEGMLAAELWYSAYWVITQS
VRWSPVHRRPFKDRLAARYGERLPCVDIFVCTADPQSEPPSLVISTVLSLMAYDYPTEKLSVYLSDDG
GSILTFYAMWEASSFAKHWIPFCKRYTIEPRSPAIYFSESDGLQDLCTPKEWSLIKDMYDEMTQRIDTI
VMSGKIPEETKANHKGFDEWNIEIYSKDHQPIVQILIDGKDQNAVDTEGNVLPTLVYMARERGLNTTIT
SKLGP>>AL3G01093.t1
MDFLKRAHFSTQMRIRVSSVISNSPIIMNADCDMYSNNKDAIRESLCFFLDEEMGHKIGFVQYPQNYN
NMTKNNIYGNSLQVINKELT
>AL3G01094.t1
MAYDYPTEKLSVYLSDDGGSILTFYAMWEASSFAKHWIPFCKRYSVEPRSPAIYFSESDGLQDLCTPK
EWSLINDMYDEMTQRIDTIVMSGKIPEETKANHKGFDEWNIEIYSKNHQPIVQILIDGKDQNAVDTEGN
VLPTLVYMAREKRPQHHHNFKAGAMNALIRVSSVISNSPIIMNADCDIYSNNKDAIRESLCFFLDEEMG
HKIGFVQYPQNYNNMTKNNIYGNSLQVINKVELRGLDSVGGPLYIGTGCFHRREILCGRRFTKDYKED
WNGGIKDQTQESIHEIEEKAKFLATCTYELGTQWGDEIGLKYGCPVEDAITGLAIHCRGWGSICNLPT
RPAFLGVGPTTLAQTLLQHKRWTEGNFSIFLSKYCSFLFGHGKIGLRLQMGYCIYGLWAPNSLATLYY
VTIPSLALLKGTPIFPEMTSPWIAPFIYVFFVKNLYSLYEALSCGDTLKG
>AL3G01095.t1
METTPESRRLFATESLGGRAVYRLHAVTVAAGILLVLYYRATHVPAAGEGRAAWLGMLAAELWYSAY
WVVTQSVRWSPVHRRPFKDRLAARYGERLPCVDIFVCTADPQSEPPSLVISTVLSLMAYDYPAEKLS
VYLSDDGGSVLTFYAMWEASLFAKHWIPLCKRYNIEPRSPAVYFSESDGLEDLCTPKEWSLIKDMYD
EMTQRIDTVVMSGKIPEETKANHKGFDEWNLEINSKDHQPIVKILIDGKDQNAVDTEGNVLPTLVYMA
REKRPQHHHNFKAGAMNALIRVSSVISNSPIIMNVDCDMYSNNKDAIRESLCFFLDEEMGRKIGFVQY
PQNYNNMTKNNIYGNSLQVINKVELRGLDSVGGPLYIGTGCFHRREILCGRRFTKDYKEDWNGGIKD
QTQESIHEIEEKAKSLATCTYELGTQWGDEIGIKYGCPVEDVITGLAIHCRGWGSICNLPTRPAFLGVC
PATLAQTLLQHKRWSEGNFSIFLSKYCAFLFGHGKIGLRLQMGYCIYGLWAPNSLATLYYVTIPSLALL
KGTPLFPEMTSPWITPFIYVFFVKNLYSLYEALSCGDTLKGWWNGQRMWLVKRITSYLFGVIDTLRKL
LGLSQMSFAVSSKVSDEDVSKRYEQEIMEFGSSTPEIVIIATTTLLNLVCMVGGLRQITIVGLNIFSPQLII
CGMLVITSIPFYEAMFMRKDKGSIPSSVTLSSVGFVVLALLVVILVPIV
>AL3G01096.t1
METESRRLFATESIGGSRAVYKVHAITVAAGILLVLYYRATHIPAAGEGRAAWLGMFAAELWYSAYWV
ITQSVRWNPVRRRPFRDRLAARYGERLPCVDIFVCTADPQSEPPSLVISTILSIMAYDYPAEKLSVYLS
DDGGSILTFYAMWEASSFAKHWLPFCKRYNIEPRSPAAYFSESHGLQDLCNKKEWSLIKDMYEEMTE
RIDTAVMSGKIPEEIKANHKGFDEWDLEINSKDHQPIIQILIDGKDENAVDNEGNVIPTLVYMAREKRPQ

HHHNFKAGAMNALIRVSSVISNSPIIMNVDCDMYSNNKDTIRDALCFFLDEEMGHRIGFVQYPQTYNN
MTKNDIYGNSMHVISKVELRGFDSVGGPLYIGTGCFHRREILCGRRFTEEYKEDWDGGIKEKKQEHV
YEVEKKAKSLATCTYEHDTQWGDEIGLKYGCPVEDVITGLAIHCRGWESVLNSPTRPAFVGVGPTTL
AQTLLQHKRWSEGNLIIFLSKYCPFLFGHGKIKLRHQMGYCIYGLWAPNSLATLYYVTIPSVALLKGTP
LFPEITSPWITPFIYVFCVKNIYSLYEALSCGDTLKGWWNGQRMWIVKRITSYLFGVIDTLRKLLGLSQ
MTFAVSSKVSDEDVSKRYELEIMEFGSSTPEYVIISTIALLNLVCLVGGLRHFMTGGWNALLNAFSPQL
ILCGMLVIINIPLYEAMFLRKDKGSIPFSVTLASIGFVVSAFLVDILAPIV
>AL3G01097.t1
MGGERAQEREMMESRQLFATERVAGRAVYRLHAITVAVGILLVLYYRATHAPAAGEGRAAWLGMLA
AELWYAAYWVVTQSVRWSPLRHRPFRDRLAARYGERLPCVDIFICTADPESEPPSLVISTVLSVMAY
NYPTEKISVYLSDDGGSVLTFYALWEASLFAKHWLPFCKRYNIEPRSPAAYFSESDGHKALCTPKEW
SFIRDMYEEMTERTDTAAISGKIPEEIKENHKGFHEWNQKITSKNHQPIVKLLIDGKSGNAVDNEGNVL
PTLVYMSREKRPQYHHNFKAGAMNALIRVSSVVSNSPIIMNVDCDMYSNNCNTIRDALCFFLDEEMG
HKIGFVQYPQNFNNLTKNNIYGNSHQVTNQVLMGGMDSVGGPMYVGTGCFHRREILCGRRFTEDYK
EDWKRGIKEKTQESIDEIEEKAMSLAACMYEHGTQWGDDIGLKYGYPAEDIITGLAIHCRGWMSVHN
NPPRAAFLGVAPTTLAQTLLQHKRWSDGAFSIFLSKYCSFLFGHGKIKLRHQMGYSIYGLWAPNSLPT
LYYVIIPSLALLKGTSLFPEITNPWITPFMYVFCVKNMYSLYEALSCGDTLKGWWNEQRMWLVRRITS
YLYGFIDTARKLLGLSKMAFAVTSKVSEEDESKRYEQEIMEFGSASPEYVIIATVALLNLICLVGGLSQI
MTGGWNIILNSFFPQLILCGMLVITNIPFYEAMFVRKDKGRIPLSVTLASIGFAVLALLAPIV
>AL3G01441.t1
MEAGAGLVAGSHNRNELVLIRGHEDHKPLRALSGQVCEICGDEVGLTVDGDLFVACNECGFPVCRP
CYEYERREGTQNCPQCKTRYKRLKGSPRVQGDEDEEDIDDLEHEFNIDDEKKHKQHGAIQNSQITEA
MLHGKMSYGRASEDGGDGNHTPMIPPIITGNRSMPVSGEFPISAGHGHSEFTSSLHKRIHPYPISEPG
SAKWDERKEVSWKERMDDWKSKHGGADPDDMDADVPLNDEARQPLSRKVSIASSKVNPYRMVIIL
RLFVLCVFLRYRILNPVPEAIPLWLTSIVCEIWFAVSWILDQFPKWYPIDRETYLDRLSLRYEREGEPSM
LSAVDLFVSTVDPLKEPPLVTANTVLSILAVDYPVDKVSCYVSDDGASMLSFESLSETAEFARKWVPF
CKKFSIEPRAPEFYFSRKVDYLKDKVQPTFVQERRAMKREYEEFKVRINALVSKAQKVPDEGWIMKD
GTPWPGNNTRDHPGMIQVFLGHSGGLDTEGNELPRLVYVSREKRPGFQHHKKAGAMNALIRVSAVL
TNAPFMLNLDCDHYINNSKAIREAMCFLMDPQVGRKVCYVQFPQRFDGIDAHDRYANRNTVFFDINM
KGLDGIQGPVYVGTGCVFRRQALYGYNPPSGPKRPKMVTCDCCPCFGRKKRKNAMDGLPEGVADG
GMDGDKEMMMSQMNFEKRFGQSAAFVTSTFMEEGGVPPSSSPAALLKEAIHVISCGYEDKTDWGL
ELGWIYGSITEDILTGFKMHCRGWRSIYCMPKLAAFKGSAPINLSDRLNQVLRWALGSVEIFFSRHSPL
LYGYKGGNLKWLERFAYINTTIYPFTSLPLLAYCTLPAVCLLTGKFIMPPISTFASLFFISLFISIFATGILE
LRWSGVSIEEWWRNEQFWVIGGVSAHLFAVIQGLLKVLAGIDTNFTVTSKATGDEDDEFAELYAFKW
TTLLIPPTTLLVINIIGVVAGISDAINNGYQSWGPLFGKLFFAFWVIVHLYPFLKGLMGRQNRTPTIVIIWS
VLLASIFSLLWVRIDPFTVKAKGPDVKQCGINC
>AL3G03056.t1
MSRRLSLPAGSPVTVTVSPLGGNQAGSGGGSPGDGVVRRASGITSPVSRHSIGSSTATLQVSPVRR
SGGGSRYASRDDGSAEFVQYTVHIPPTPDRTTVASAASTDAPAAEEEVRPQRSYVSATIFTGGLNCA
TRGHVLADGAARPAASANLSCKMRGCDMPAFLRAGHPPCDCGFMICQECHTDCVAAAGNCPGCKE
PYSAGSDTDDSVDEDDDEAFSSSEERDQMPMTSMAKRFSMVHSIKMPTPSTNGKPADFDHARWLF
ETKGTYGYGNALWPKNGHGSGGAGATTGFVGIEEPPNFGARCRRPLTRKTSVSQAILSPYRMLIAIR
LVALGFFLTWRIRHPNPDAMWLWALSVTCEVWFAFSWLLDSLPKLCPVNRACDLDVLADRFECPTA
RNPKGRSDLPGIDVFVSTADPEKEPPLVTANTILSILAADYPVEKLACYLSDDGGALLTFEALAETASFA
RTWVPFCRKHGVEPRCPEAYFGQKRDFLKNKVRLDFVRERRKVKREYDEFKVRVNSLPEAIRRRSD
AYNAGEELRTRRRQQEEAVAAGGTLGASPLAETVAVKATWMSDGSQWHGTWLTGATDHARGDHA
GIIQAMLAPPTSEPVLGGEPAESGALIDTTGVDIRLPMLVYVSREKRPGYDHNKKAGAMNALVRSSAI
MSNGPFILNLDCDHYVHNSAALREGMCFMLDRGGDRVCYVQFPQRFEGIDPNDRYANHNLVFFDVA
MRAMDGLQGPMYVGTGCIFRRTALYGFSPPRATEHHGWLGRKKIKLFLRKKPSMGKKTDRESEHES
MLPPLEDDDHNQLSDIESSTLMPKRFGNSGTFVSSISVAEYQGRLLQDMPGVHQGRPAGALAVPRE
PLDAATVAEAISVISCFYEDKTEWGRRIGWIYGSVTEDVVTGYRMHNRGWRSVYCVTRRDAFRGTAP
INLTDRLHQVLRWATGSVEIFFSRNNAMFATRRMKLLQRIAYFNVGMYPFTSVFLLVYCVLPAVSLFT
GKFIVSHLSATFLVFLLIITITLCLLALLEIKWSGITLHEWWRNEQFWVIGGTSAHPAAVLQGLLKVIAGV
DISFTLTSKPGGADDGEEDTFAELYEVRWSFLMVPPVTIMMLNAVAMAVATARTLYSEFPQWSKLLG
GAFFSFWVLCHLYPFAKGLLGRRGRVPTIVFVWSGLICMIVSLLWVYICPPAGVKEGIGGFSFP
>AL4G00954.t1
MHVMAAGTSAHLYAVVQGLLKVMAGIEISFTLTAKAAAEDNEDIYADLYVVKWSSLLIPPITIGMLNIIAI
AFAFARTIYSDNPRWGKFIGGGFFSFWVLAHLYPFAKGLMGRRGKTPTIVFIWSGLISITISLLWVAISP
PDANATGGVHGAGFQFP
>AL4G00955.t1
MSNTQGKKGTKNAGGAPAPRPSSGQGLKFARRTASGRYLSLSREELDYTEGELAADYANYTVQIPP
TPDNQPMKDGSEPTTVAMKAEEQYVSNSLFTGGFNSVTRAHLMDKVIDSEVTHPQMAGVKAARCA

MPACDGKTMRNERGEDMDPCECRFKICRDCYLDAQKDGCICPGCKEHYKIGDYADDDPNDVSSSG
KLHLQGAGAGLLARNGEFDHNRWLFESSGTYGYGNAFMPKAGMHDDDDDPDEEGHGGGGGGGG
LPEQKPFKPLTRKIPMPASIISPYRIFIVIRMFVLIFYLTWRIRNPNMEALWLWGMSIVCELWFAFSWLL
DMLPKVNPINRSTDLAVLKEKFETPSPSNPHGRSDLPGLDVFVSTADPEKEPVLTTANTILSILAVDYP
VEKLACYVSDDGGALLTFESMAEAASFANIWVPFCKKHDIEPRNPDSYFSIKGDPTKGKRRSDFVKD
RRKVKREYDEFKVRINGLPDSIRRRSDAFNAREDMKMLKHLRESGADPSEQPKVKKATWMADGTH
WPGTWAVSAPDHAKGNHAGILQVMLRPPSPDPLYGMHDEDQLLDYSDVDIRLPMLVYMSREKRPG
YDHNKKAGAMNALAVREAMCFMMDRGGERICYIQFPQRFEGIDPSDRYANHNTVFFDGNMRALDGL
QGPMYVGTGCMFRRFALYGFDPPRTAEYTGWTFRKKKVTMFQQDPESDTQSLKTEDFDAELTAQL
VPRRFGNSSAMLASIPVAEFQARPIADHPAILHGRPPGSLTVPRPPLEPPAVAEAVSVISCWYEDKTE
WGDRVGWIYGSVTEDVVTGYRMHNRGWRSVYWISKRDAFLGTAPINMTDRLHQVLRWATGSVEIFF
SRNNAFLASRKLMFLQRVAYLNVGIYPFTSIFLVTYCFIPALSLFSGFFIVQTLNVAFLCYLLTITVTLIAL
GILEVKWSGVELEDWWRNEQFWLISGTRTPTPVND
>AL4G00978.t1
MSNTQGKKGTKNSGGAPAPRPSSGQGLKFARRTASGRYLSLSREELDYTEGELAADYANYTVQIPP
TPDNQPMKDGSEPTTVAMKAEEQYVSNSLFTGGFNSVTRAHLMDKVIDSEVTHPQMAGVKAARCA
MPACDGKTMRNERGEDMDPCECRFKICRDCYLDAQKDGCICPGCKEHYKIGDYADDDPNDVSSSG
KLHLQGAGAGLLARNGEFDHNRWLFESSGTYGYGNAFMPKAGMHDDDDDPDEEGHGGGGGGGL
PEQKPFKPLTRKIPMPASIISPYRIFIVIRMFVLIFYLTWRIRNPNMEALWLWGMSIVCELWFAFSWLLD
MLPKVNPINRSTDLAVLKEKFETPSPSNPHGRSDLPGLDVFVSTADPEKEPVLTTANTILSILAVDYPV
EKLACYVSDDGGALLTFESMAEAASFANIWVPFCKKHDIEPRNPDSYFSIKGDPTKGKRRSDFVKDR
RKVKREYDEFKVRINGLPDSIRRRSDAFNAREDMKMLKHLRESGADPSEQPKVKKATWMADGTHW
PGTWAVSAPDHAKGNHAGILQVMLRPPSPDPLYGMHDEDQLLDYSDVDIRLPMLVYMSREKRPGYD
HNKKAGAMNALVRCSAVMSNFPFILNFDCDHYINNTQAVREAMCFMMDRGGERICYIQFPQRFEGID
PSDRYANHNTVFFDGNMRALDGLQGPMYVGTGCMFRRFALYGFDPPRTAEYTGWTFRKKKVTMFQ
QDPESDTQSLKTEDFDAELTAQLVPRRFGNSSAMLASIPVAEFQARPIADHPAILHGRPPGSLTVPRP
PLEPPAVAEAVSVISCWYEDKTEWGDRVGWIYGSVTEDVVTGYRMHNRGWRSVYWISKRDAFLGT
APINMTDRLHQVLRWATGSVEIFFSRNNAFLASRKLMFLQRVAYLNVGIYPFTSIFLVTYCFIPALSLFS
GFFIVQTLNVAFLCYLLTITVTLIALGILEVKWSGVELEDWWRNEQFWLISGTRTPTPVND
>AL4G00979.t1
MHVMAAGTSAHLYAVVQGLLKVMAGIEISFTLTAKAAAEDNEDIYADLYVVKWSSLLIPPITIGMLNIIAI
AFAFARTIYSDNPRWGKFIGGGFFSFWVLAHLYPFAKGLMGRRGKTPTIVFIWSGLISITISLLWVAISP
PDANATGGVHGAGFQFP
>AL4G03483.t1
MGSLAAGNGTAGHASNGGGALADHALALEMNGAAGNGHRKAGAGAERAAPVLQANGGGKTKKKKI
SPKDKYWVASDDGEVAAAIQDGGEDGRRPLLYRNFRVKAVLLYPYRLLSLIRLVAIVLFFIWRVRHPY
ADGMWLWWISMVGDLWFGVTWLLNQVAKLSPIKRVPDLALLKQQFDQADGNSTLPILDVFINTVDPI
NEPMIYTMNSILSILAADYPVDKHACYLSDDGGSIIHYDGLLETAKFAALWVPFCRKHSIEPRAPESYFS
VKTRPYTGNAPEEFISDQKHIFREYDEFKCRLDALFTVIPKRSDGYNHSDTKEGAKATWMADGTQWP
GTWIEPAKNHKRGDHDGIVKVMLKHPGHEPELGLPASTNNLLDFSSVDLRLPMLVYISREKRPGCDH
QKKAGAMNVQLRVSALLTNAPFIINFDCDHYVNNSKAFRAGICFMLDRREGDNTAFVQFPQRFDDVD
PTDRYCNHNRVFFDATLLGLNGIQGPSFVGTGCMFRRFALYGTDPPRWRSDDVKIVDSSKKFGSSLS
FLSSIPASANQERSIVSPPALEESVMADLADVMTCAYEDGTEFGREVGWVYNIATEDVVTGFRLHRN
GWRSMYCRMEPDAFAGTAPINLTERLYQILRWSGGSLEMFFSRYCPLLAGGRLHPMQRIAYANMTA
YPVSSIFLVFYLLFPVIWIFRGQFYIQKPFPTYVLYLVIVIVMTELIGVIEIKWAGLTMLDWIRNEQFYIVGA
TAVYPTAVLHMVLKLFGFKGVSFKLTAKQVASGTSDKFAELYAVQWAPLLIPTMVVIAVNVCAIGASIG
KAIFGGWSLMQMADAGLGLLFNAWILVLIYPFALGMIGWWSKRPYVLFIMLIVAFIVIAMMDIAIQSMRS
GFVRFHFKSSGGATFPTSWGL
>AL4G03484.t1
MAAAVTRRLHALRVEGPDGHAVADVESERGASEEADTPVAKRVADAKDDVWVAADEGDVPGTGRP
LLFRTMKVKGSILHPYRFLILARLIAVVAFFAWRMKHRNHDGGAWLWATSMVADTWFGFSWLLNQLP
KLNPVKRVPDLAALADQHGSSGGEAANANLPGIDVFVTTVDPVDEPILYTVNTILSILAADYPVDRYAC
YLSDDGGTLVHYEAMAEVAGFAALWVPFCRMHCVEPRSPENYFGMKTRPYVGRMAGEFVRDHRR
VRREYEEFKVRIGSLASTIRQRSDAYNSSSGKGVGARATWMADGTQWPGTWIEQAENHRRGQHAG
IVQVILNHPSCKPQLGSPASTNNPFDFSNVDTRLPMLVYISREKRPGYNHQKKAGAMNVMLRVSALL
SNAPFVVNFDCDHYINNSQALRAPMCFMLDPRDGQNTAFVQFPQRFDDVDPTDRYANHNRVFFDG
TMLSLNGLQGPSYLGTGTVFRRVALYGMEPPRYRAEKIKLVAGKADEFGSSTSFVNSMPDAAIQERSI
TPVLVDEGLGDDLAALMTCAYEDGSSWGRDVGWVYNIATEDVVTGFRMHSQGWRSMYCSMEPAA
FRGTAPINLTERLYQVLRWSGGSLEMFFSHSNALLAGGRLHPLQRVAYLNMSTYPIVTVFILAYNLFPV
MWLFSEQFYIQRPFGEYIAYLVGVIAMIHLIGMFEVRWAGITLLDWCRNEQFYMIGATGVYPTAVLYM
ALKLVTGKGLHFRLTSKQTEACSGDKFADLYTVRWVPLLIPTVAVLVVNVAAVGAAIGKAATWGFFTD

QARHVVLGMVFNVWILVLLYPFALGIMGQWGKRPVILFVMLVMAIGAVGIVYFTLHTPYPAEWSEVAA
SLGKTSLTGGPSG
>AL4G03485.t1
MASAAGSSGGNASLADPLLASAKKLVGAKGKYWEAADKEARRAAKESGGEDGRPLLFRTYKVKGAL
LHPYRALIFIRLIAVLLFFVWRVKHNKSDIMWFWTMSVVGDVWFGFSWLLNQLPKFNPVKTIPDIAALR
RQYDLPDGTSSLPGIDVFVTTADPIDEPILYTMNCVLSILASDYPVDRCACYLSDDSGALIQYEALVETA
KFATLWVPFCRKHCIEPRAPESYFELEAPLYTGRAPEEFTDDHTSVRREYAEYKERLESLSSAISKRS
DVYNSMKTDQDDVKATWMANGTQWPGTWIDTTETHRKGHHVGIVKVVLDHPTRGHNLGSQACTHS
LNFTNIDERIPMLVYIARGKNPSYDHNKKAGALNAQLRASGLLSNAQFIINFDCDHYINNSQALRAAICF
MLDQREGDNTAFVQFPQRFDNVDPSDRYGNHNRVFFDGTMLALNGLQGPSYLGTGCMFRRIALYGI
DPPHCREDNIVVDANRFGSSTLFQDSVSKAINQERSTIPPTLSETILTEMETVVSASHDKETGWGKGV
GYIYEIATEDIVTGFRIHGQGWRSMYCTMERDAFCGIAPINLTERLHQIVRWSGGSLEMFFSRNNPLIG
GGRLQTLQRVSYLNMTVYPATSLFILLYSLSPVMWLIPDEVYIQRPFTKYVVYLLVIILMIHIIGWLEIKW
AGITWLDYWRNEQFFMIGSTSAYPAAVLHMVVNLLTKKGIHFRVTSKQTAADTNDKFADLYDMRWVP
MLIPTAVVLICNVSAIGVAMGKTIVYMGAWTIAQKTHAALGLLFNVWIMVLLYPFALAVMGRWAKRPVI
LVVLLPVAFVIVCLVYITVHILLGGLVAF
>AL4G05011.t1
MASAKKLQERVALPRTAWILADFLILFLLLALLSRRAASLWEEHGPTSTSTTWLPALALACEAWFTFV
WLLNMNGKWSPVRFDTYPENLPDEQELPAVDMFVTTADPALEPPVITTNTVLSLLAVDYPDAGGKLA
CYVSDDGCSPVTCYALREAAEFAAIWVPFCRKHGVAVRAPFMYFSSAPEVAKGTADHEFLESWAFM
KSEYEKLVTRIENAGEGSILRDGDFAEFIEAERRNHPTIVKVLWDNSKAKTGEGFPHLVYVSREKSPE
HYHNFKAGAMNVLTRVSGVMTNSPIMLNVDCDMFANNPQVVLHAMCLLLGFADETQSGFVQAPQKF
YGGLKDDPFGNQMEVIYKKVGGGVAGIQGLFYGGTGCFHRRKVIYGVPPPDVVVKHERSGSPSYKE
LQTKFGSSKELIESSRDIISGDVLLARPTESISSRVEVAKQVGACSYEAGTCWGQEIGWVYGSMTEDIL
TGQRIHATGWKSALLDTTPPAFLGCAPTGGPASLTQFKRWATGLLEILISRNSPILGAIFRRLQLRQCL
AYLIIHVWPVRAPFELCYALLGPFCLLTNHSFLPKVSDEGFRVTLALFMAYNMYNLMEYKECGLSARA
WWNNHRMQRITSASAWLLAFLTVLLKTVGLSETVFEVTRKESSSRSSSTADGGAASTGEADDPGLFT
FDSSPVFVPATALSILNVVAVAVGAWRALAGTATGVRGGPGMGEFVCCGWMVLCLWPFVRGLVSR
GRYGIPWSVKVKAGLIVSLFVHLCTRN
>AL5G00055.t1
MYCLQIHGCSVTNLQAASYTYARQAAMSPVTAVDNAGKLDEPLLVANGNRRSDQAAAAAMAKANGL
KKKKKKEEGGANKEEEQYWVDIDQQPAGDDVLLDLEDGQPLLLLFRNKKVKRRILYPYRMLIFARLIA
VILFFIWRLKNNNSDIIWLWAMSVAGDVWFGFSWLLNQLPKVNPVKSIPDLDALKQKCDMGEGISSLP
GIDVFVTTANPVDEPILYTMNCILSILAIDYPVDKNSCYLSDDAGALVSYEALVETAKFATLWVPFCRKY
CIEPRAPESYFEVKATPYAGSSVEDFITDHKYVHRKYEELKMTIEMLPETIRERSNAYKNMHMNATLM
EDGTQWPGTWIDPTENHKKGHHASIVQIVVNKPSHTPHLGLPVGANDTLDFSTIDVRLPMLVYVSRE
KHPSYDHQKKAGAMNTQLRVSALLSNAPFIINFDCDHYINNSQSFRAAVCFMLDPREGENTAFVQFP
QRFDNVDPTDRCISKYILSTNLPSKNMGYVL
>AL5G00190.t1
MASNGGGGGVLRHSNSSRLSRMSYSGDDARHDRASAPVGDRPMVTFARRTHSGRYVSYSRDDLD
SELANADLGNGNGGFSPDREEFLSYHVHIPVTPDNQPMDPAISARVEEQYVSNSLFTGGFNSVTRAH
LMDKVIESEASHPQMAGAKGSSCAVPGCDAKVMSDERGQDILPCECDFKICADCFGDAVKNAGALC
PGCKESYKSTEMEDLVGPGQPTLSLPPPPGAGGAPASRMERRLSMVRSQKALTRSQTGDWDHNR
WLFETKGTYGYGNAIWPKENDGQGDSNNNGSSGAGGGGLGGGGTNGQPAEFTSKPWRPLTRKLKI
PAAVLSPYRLLVLIRLAVLALFLHWRVTHTNPDAMWLWGMSVVCEIWFGFSWILDQLPKLCPINRATD
LAVLKDKFETPTPSNPNGRSDLPGLDLFVSTADPEKEPPLVTANTILSILAADYPVEKLACYVSDDGGA
LLTFEAMAEAASFANMWVPFCRKHVIEPRNPDSYFALKRDPYKNKVRSDFVKDRRRIKREYDEFKVRI
NGLPDSIRRRSDAYHAREEIKAMKRQREAALDDAVEAVKIAKATWMADGTHWPGTWIQPSAEHMKG
DHAGIIQVMLKPPSDDPLYGSNGDEGTPLDFTDIDIRLPMLVYVSREKRPGYDHNKKAGAMNALVRS
SAVMSNGPFILNLDCDHYVYNSQAFREGMCFMMDRGGDRLAYVQFPQRFEGIDPSDRYANHNTVFF
DVNMRALDGLMGPVYVGTGCLFRRVALYGFDPPRSTEHGGCCSCCFPQKRKIKSSNNEETQALRM
ADFDDEEMNMSTFPKRFGNSNFLINSIPIAEFQGRPLADHPGVKNGRPPGALTVPRDLLDASTVAEAI
SVISCWYEDKTEWGQRVGWIYGSVTEDVVTGYRMHNRGWKSVYCVTKRDAFRGTAPINLTDRLHQ
VLRWATGSVEIFFSRNNALLASRRMKLLQRIAYLNVGIYPFTSIFLIVYCFLPALSLVSGQFIVQTLDVTF
LVYLLVITLTLIMLAVLEIKWSGINLEEWWRNEQFWLIGGTSAHLAAVLQGLLKVIAGIEISFTLTSKSGG
DDENDEYADLYIVKWTSLMIPPIVIMMLNLFAIAVGFSRTIYSEIPQWSKLLGGVFFSFWVLAHLYPFAK
GLMGRRGRTPTIVFVWSGLVAITISLLWVAISPPSQNSQIGGSFTFP
>AL5G02634.t1
MAPPSTYVSSDSSWWGGKEEHGTPVVVKMDNPYSLVEIDGPGMDSAEKARRKNAKQFKWVLLLRA
HRAVGCVAWLAGGFWALMGAVNRRVRRSRDADAEPDAEASGRGRHMLRFLRAFLLLSLAMLCFET
VAYLKGWHFPRNLPEEYLRHLPDHVLLNLPEHLRHLPENLRIPERREIKGWLHGAYVAWLSFRIDYIA
WAIQKLSGFCIVLFMVQSVDRIILCLGCFWIKIRGLKPRLPAAPTKPRSNDDLEDGDDLDAYFPMVLVQ

MPMCNEKEVYETSILHVCQIDWPRDRMLIQVLDDSDDVTCQMLIKAEVTKWSQKGVNIIYRHRLSRTG
YKAGNLKSAMGCEYVKDYEFVALFDADFQPNPDFLKLTVPHFKGNPELALVQARWIFVNKDENLLTR
LQNINLCFHFEVEQQVNGVYLNFFGFNGTAGVWRIEALEDSGGWMDRTTVEDMDIAVRAHLQGWKF
IYLNDVKVLCELPESYTAYRKQQHRWHSGPMQLFRVCLPAIFKSKIPFWKKANLVMLFFLLRKLILPFY
SFTLFCVILPLTMFVPEAQLPVWVICYVPMLMSVLNILPAPKSAPFIIPYLLFENTMSVTKFNAMVSGLF
QLGSSYEWVVTKKAGRTSSESDMLALAEEADTLARPAGAKLVRGVSEGGLEAWAKMHEHDKKDLQ
LQADAQDEEDATSPPAKKNSKPKAPNRIFKKELALASLLLIAATRSLLSAQGLHFYFLLFQGVTFLVVG
LDLIGEQVS
>AL5G02870.t1
MAPAVAGGGRVRSNEAPAASAAAAATGNPCACGFQVCACTGTAAVASAASSVDMDIMATGRIGPLN
DESWVGVELGEDGETDESGAAVDDRPVFRTEKIKAVLLYPYRVLIFVRLIAFTLFVIWRISHKNPDAM
WLWVTSICGEFWFGFSWLLDQLPKLNPINRVPDLAVLRQRFDRPDGTSTLPGLDIFVTTADPFKEPIL
STANSVLSILAADYPVDRNTCYVSDDSGMLLTYEALAEASKFATLWVPFCRKHGIEPRGPESYFELKS
HPYMGRAQDEFVNDRRRVRKEYDEFKARINSLDHDIRQRNDGYNAANAHREGEPRPTWMADGTQ
WEGTWVDASENHRKGDHAGIVKVLLNHPSHSRQYGPPASADNPLDFSGVDVRVPMLVYVSREKRP
GHNHQKKAGAMNALTRASALLSNAPFILNLDCDHYINNSQALRSGICFMLGRDSDTVAFVQFPQRFE
GVDPTDLYANHNRIFFDGSLRALDGMQGPIYVGTGCLFRRITVYAFDPPRINVGGPCFPMLGGMFAK
TKYQKPGLEMTMAKAKAAPVPAKGKHGFLPLPKKTYGKSDAFVDSIPLASHPSPYVAAYNTAEGIVTD
EATMAEAVNVTAAAFEKKTGWGKEIGWVYDTVTEDVVTGYRMHIKGWRSRYCSIYPHAFIGTAPINLT
ERLFQVLRWSTGSLEIFFSKNNPLFGSTYLHPLQRIAYINITTYPFTAIFLIFYTTVPALSFVTGHFIVQRP
TTMFYVYLGIVLATLLIIAVLEVKWAGVTVFEWFRNGQFWMTASMSAYLQAVCQVLIKVIFQKDISFKLT
SKLPAGDEKKDPYADLYVVRWTPLMIVPIIVIFVNIIGSAVAFAKVLDGEWTHWLKVAGGVFFNFWVLF
HLYPFAKGILGKHGKTPVVVLVWWAFTFVITAVLYINIPHMHSPGGKHTKVAHGHHGQKFLGWP>>AL
5G03480.t1
MSNDNTNYTVFMPPTPDNQPGAAQAGASGSGGSTKPDNHPLPPYSSGSKLVNRRGGDDAGAASG
GGSKMERRLSTAPVASPSKSLLVRNQAGDFDHNRWLFATKGTYGIGNAYWPQDDNPYNNEDGTGM
GGGGEGGVKMEDLIDKPWKPLSRKVPIDPSILSPYRLLVLVRFVSLFLFLIWRASNPNPDAMWLWGIS
IVCEYWFAFSWILDQMPKLNPINRAADLAALREKFESKTPSNPTGRSDLPGLDVFISTADPYKEPPLVT
ANTLLSILATDYPVEKLFVYISDDGGALLTFEAMAEACAYAKVWVPFCRKHSIEPRNPDAYFNQKGDP
TKGKKRPDFVKDRRWIKRAYDEYKVRINDLPDAIRRRANAMNARERKIAREKAAASSETPALPIKATW
MADGTHWPGTWLDSAPDHGKGDHASIVQVMIKNPHYDVVYGDAGDHPYLDFTNVDVRIPMFVYMS
REKRQGYDHNKKAGAMNAMVRASAILSNGPFMLNFDCDHYVYNCQAIREAMCYMLDRGGDRICYIQ
FPQRFEGIDPSDRYANHNTVFFDGNMRALDGLQGPMYVGTGCLFRRHAIYGFNPPRATEYHGLVGQ
TKVPVDPNARSGVDGDGDELSPLTDHPDHEAPQRFGKSKLFVESIAVSEYQGRPLQDHPSVRNGRP
PGALLMPRPPLDAATVAEAVSVISCWYEDTTEWGQRVGWIYGSVTEDVVTGYRMHNRGWRSVYCIT
KRDAFRGTAPINLTDRLHQVLRWATGSVEIFFSKNNALCASRRLMFLQRMSYLNVGIYPFTSLFLIMYC
LLPALSLFSGQFIVATLDPTFLCYLLLISITLILLCLLEVKWSGIGLEEWWRNEQFWVIGGTSAHLAAVLQ
GLLKVTAGIEISFTLTAKAAAEDDDDPYAELYLIKWTSLFIPPLAIIGINIIAMVVGLSRCVYAEIPQYSKLL
GGGFFSFWVLAHYYPFAKGLMGRRGRTPTLVYVWAGLISITISLLWITISPPDDNISQGGILV
>AL5G03868.t1
MDDESLQKAPPNTPEPVLRGASQSDGGDDDEGPPPPEPLSSKVPIPSALLSLYRAAVTIRAVLLALFL
RYRVTHPVQHDSYALWLTSVACECCLALSWLAAQLTKLSPTHRATRLHLLEKEEEKDGGKQHLASVD
VFVTAADAAREPPLATANTVLSVLAADYPAGRLACYVSDDGADALLFEALHETARLARAWVPFCRAH
GVEPRAPGPYFARGVDYIRDKVAPSFVRERRKVKREYEEFKVRLNRLAAATARARKVPEGGWVMSD
GTPWPGNDPRDHPAMVQLLLGHAAGDDAEASEELLPRLVYVSREKRPGFQHHGKAGAMNALLRVS
AVLTNGAYVLNLDYDHYVNNSGALREAMCFMMDPAEAGNRTCFVQFPLRPTVAAAADDDAGHDSV
FFNIDMKCLDGIQGPVYVGSGCCFNRKALYGFDPAVADDADDLEAGRRKWCCFGARESRKARLKRT
MSVVPLLDSEDEDEETERARKRRLHSYRAALERHFGLSPSFIVSVFGEQGTGGDSPRPAAMAVSALP
TEAIINVISCAYEERTRWGREFGWIYGAGASGGVATATGFRMHARGWESAYCVPARPAIRSYAGRSP
SEMLAGASRQAAADMGILLSRHCPVWAAGSGMKPLQRLAYVNRVAYPLASLPLTVYCALSAACLLSG
KSIFPDDVGYDAGVLLVLLLTSVIASVALELKWSGVSLREWWRDEKLWVVLGMSAGLAAVFLGIIRAC
TGIDVAFSTDTTAASVSNQEAEDDDDEEMPEKRLVQWSNLLVPPISVLLGNLAGAVVAFSYGVDHGY
ESWGPLAWRLLFAAWVVAHLQGFLRGLLARRERAPTIAVLWSVLFVSILSLLWVNVQTYYAPAAVTH
QPIF>>AL6G02856.t1
MAGSSFSAVAGGKPPLFATEKPKRMVAYRLYAGTIFAGILLIWVYRATHIPPRDSSSLGWRASLRLLV
AEVFFGLYWSPHPLRAVEPHPSHHLQGPPHGKDLYRDLTDRVNSVVHSGKIPEVPECNCRDFSEWN
ETITSEDHPSIVQILIDRNKRKAVDVDGNVLPQLVYMAREKRPQEKHHFKAGSLNALIRASSVISNSPVI
LNVDCDMYSNNSESIRDALCFFLDVEQAENADELERMSKELVTCTYEHDTLWGIEKGVRYGCPLEDV
ITGLQIQCRGWRSVYYNPVRKGFLGMAPTSLGQILIQHKRWSEGFLQISFSNYSPFILGHGKIKLGLQM
GYGLWLLGLNSFPTFYYAIIPSLCFLGGVSVFPEPSRVIAICDTAVEWWNAQRMWFFRRTTSYLLAAID
TIGEMLGVSESGFDLTVKVNDSEALERYKKGKMEFGPISGMFVILTTIALFNLVCMVVGLGRVLWREG
AEGLGPLFLQAVLCVAIVAINAPVYEALFFRKDSGSLPSFVTLISMCFVSSLCLQAI

>AL6G02857.t1
MAGSSFSAVAGGKPPLFATEKPKRMVAYRLYAGTIFAGILLIWVYRATHIPPRDSSSLGWRASLGLLV
AEVFFGLYWVLTLSVRWNPILRTTFKDRLTERYGEDQLPGVDIFVCTADPALEPPMLVISTVLSVMAY
DYPPEKLNVYLSDDAGSAATFYALCEASEFAKHWIPFCKNYKVEPRSPAAYFAEAATPHDASSPQEL
LRMKDLYRDLTDRVNSVVHSGKIPEVPECNCRDFSEWNETITSEDHPSIVQILIDRNKRKAVDVDGNV
LPQLVYMAREKRPQEKHHFKAGSLNALIRASSVISNSPVILNVDCDMYSNNSESIRDALCFFLDVEQG
QDIGFVQYPQNFDNVVHNDIYGNPINVVNELDHPCLDGWGGMCYYGTGCFHRRETLCGQIYSKDYK
EDWARGVGTAENADELERMSKELVTCTYEHDTLWGIEKGVRYGCPLEDVITGLQIQCRGWRSVYYN
PVRKGFLGMAPTSLGQILIQHKRWSEGFLQISFSNYSPFILGHGKIKLGLQMGYAVCGFWALNSFPTF
YYAIIPSLCFLGGVSVFPEITSPWIIPFGYVAVAAYSCSLAESLQCGDTAVEWWNAQRMWFFRRTTSY
LLAAIDTIGEMLGVSESGFDLTVKVNDSEALERYKKGKMEFGPISGMFVILTTIALFNLVCMVVGLGRV
LWREGAEGLGPLFLQAVLCVAIVAINAPVYEALFFRKDSGSLPSFVTLISMCFVSSLCLQAI
>AL6G02863.t1
MAGSSFSAVAGGKPPLFATEKPKRMVAYRLYAGTIFAGILLIWVYRATHIPPRDSSSLGWRASLGLLV
AEVLFGLYWVLTLSVRWNPILRTTFNDRLTERYGEDQLPGVDIFVCTADPALEPPMLVISTVLSVMAY
DYPPEKLNVYLSDDAGSAATFYALCEASEFAKHWIPFCKNYKVEPRSPAAYFAEAATPHDASSPQEL
LRMKDLYRDLTDRVNSVVHSGKIPEVPECNCRDFSEWNEAITSEDHPSIVQILIDRNKRKAVDVDGNV
LPQLVYMAREKRPQEKHHFKAGSLNALIRASSVISNSPVILNVDCDMYSNNSESIRDALCFFLDVEQG
QDIGFVQYPQNFDNVVHNDIYGNPINVVNELDHPCLDGWGGMCYYGTGCFHRRETLCGQIYSKDYK
EDWARGVGTAENADELERMSKELVTCTYEHDTLWGIEKGVRYGCPLEDVITGLQIQCRGWRSVYYN
PVRKGFLGMAPTSLGQILIQHKRWSEGFLQISFSNYSPFILGHGKIKLGLQMGYGLWLLGPEQLSHLLL
CHHPFSLLPRWCLCFP
>AL6G04018.t1
MSSPAAAAGRRLADPLLAASNGDGEIVVSAKDKYWVPSDEKEVRLAASQDCGGGGTPPLLYRTFRV
KGFLINLYRLLTLVRVMVVIMFFTWRMEHRDSDSMLLWWISVVGDVWFGVTWLMNQVTKLKPRKCV
PKLSMLREQFDSSDGSSDLPILDVFINTVDPVDEPVLYTMNSILSILATDYPAEKYSTYFSDDGGSLIHY
EGLLETTKFAASWVPFCRKHCVEPRAPESYFGRNVRPYTGNAPSDFLDDHRRIRAEYGEFKARLDGL
STVIEQRSEALNGANTKEGHVNATWMADGTQWQGTWIEPAKGHKKGHHPAVFQVMLNQPSDDPQL
GMPASSDHPLDFSAVDVRLPTLVYIAREKRPAYDHQKKAGAMNVQLRVSALLSNAPFIINFDGDHYIN
NSQAFRAAMCFMLDPRDGDNTAFVQFPQRFDDVDPTDRYCNHNRMFFDATLLGLNGIQGPSFVGT
GCMFRRVALYGADPPQWRPDDAAKAKVLQRPNVFGKSTAFINSMPAAANQERSIASPATLQETASV
DAELASVMTCAYEAGTEWGNDVGWVYNIATEDVVTGFRLHRNGWRSTYCAMEPDAFRGTAPINLTE
RLYQILRWSGGSLEMFFSRFCPLLAGRRLHPMQRIAYINMTTYPVSTFFLVMYDLYPVMWLVHGQFYI
QRPFQTFALFVAVIIATVELIGMVEIKWAGLTLLDWIRNEQFYIIGTTGVYPTALLHILLRALGLKGVSFKL
TAKKLTGGARERLAELYDVQWVPLLVPTVVVMAVNVAAIGAAAGKAIAGRWSAAQVAGASTGLVFNV
WMLLLLYPFALGIMGRWSKRPYILFALLVAAVAVITSVYATLASSVPYFHSGYFFQLN
>AL6G04043.t1
MLRWSGGSLEMFFSRFCPLLVGDRRLRLMQRIAYINMTTYPVATFFIVMYNLCPVMWLFQDQLYIQR
PFQTYTLFLAVAIDTVDLIGMVEIKWAGLTLLDWFPNEQMYIIGTTGVYPMAMMHILLRFLGLKGVSFKL
TAKKLMGCTTK
>AL6G04044.t1
MSRYTGRSPKEFLDDHRRMRTEYDKFKARLEVLPSVIARQSEALSLANPDVGCENAPWMADGTQW
QGTWIEPASGHRKGDHPAILQVVLKQPSSKPQLGMPSSSDHPLNLSAIDVRLPMLVYISREKRPGYD
HQKKAGAMNVQLRVSALLSNAPFVINFNGDHYINNLQAFRAAMCFMLDPRAGQDTAFVQFLQRFDD
VDTTDRYDNHNCMFFDATLLGLNGIQGPSYVSTGCMFRRVSRGPTTVAAGWRRRCLAQARHLWKID
SVLELGSGGC
>AL6G04053.t1
MSSPAAAAGRRLADPLLAASNGDGEIVVSAKDKYWVPSDEKEVRLAASQDCGGGGTPPLLYRTFRV
KGFLINLYRLLTLVRLMVVILFFTWRMEHRDSDSMWLWWISVVGDLWFGVTWLMNQVTKLKPRKCV
PKLSMLREQFDLSDGSSDLPILDVFINTVDPVDEPVLYTMNSILSILATDYPAEKYSTYFSDDGGSLVH
YEGLLETAKFAASWVPFCRKHCVEPRAPESYFGRNVRPYTGNAPSDFLDDHRRIRAEYGEFKARLD
GLSTVIEQRSEALNGANTKEGHVNATWMADGTQWQGTWIEPAKGHKKGHHPAVFQVMLNQPSDDP
QLGMPASSDHPLDFSAVDVRLPTLVYIAREKRPAYDHQKKAGAMNVQLRVSALLSNAPFIINFDGDHY
INNSQAFRAAMCFMLDPRDGDNTAFVQFPQRFDDVDPTDRYCNHNRMFFDATLLGLNGIQGPSFVG
TGCMFRRVALYGADPPQWRPDDAAKAKVLQRPNVFGKSTAFINSMPAAANQERSIASPATLQETAS
VDAELASVMTCAYEAGTEWGNDVGWVYNIATEDVVTGFRLHRNGWRSTYCAMEPDAFRGTAPINLT
ERLYQILRWSGGSLEMFFSRFCPLLAGRRLHPMQRIAYINMTTYPVSTFFLVMYDLYPVMWLVHGQF
YIQRPFQTFALFVAVIIATVELIGMVEIKWAGLTLLDWIRNEQFYIIGTTGVYPTALLHILLRALGLKGVSF
KLTAKKLTGGARERLAELYDVQWVPLLVPTVVVMAVNVAAIGAAAGKAIAGRWSAAQVAGASTGLVF
NVWMLLLLYPFALGIMGRWSKRPYILFALLVAAVAVITSVYATLASSVPYFHSGYFFQLS
>AL6G04059.t1
MLYTMNSILSILSTDYPVEKYATYFFDDGGSLVHYEGLLETTQFAALWVLFCQKHCVEPRAPESYFWT

KMSRYTEKSPKEFLDDHRRMRTEYDKFKARLEVLPSTVVLNQPSSKPQLGMPASSDHPLNLSAIDVR
LPMLVYISRDKRPGYNHQNKAGAMNAQLHVSALLSNAPFVINFDGDHYINNSQAFRAAMCFMLDPRA
GQDTAFVLFPQRFDDVDRMDRYCNHNRMFFDATLLGLNGIQGPSYVGTGCMFRRVAVYGADPPRW
RPNGGAGVLRRPDIFGKSTVFLNSVLVAANQECPAATAALTTLEETELADVMACVYEEGTKWGNKVG
WVYNIATEDVVTGFRLHGIGWRSSYRSMVPDAFRGTMFFSRFCPLLVGDRRLRLMQRVAYINMTTY
PVAAFFIVMYNLCPVIWLFQDQLYIQRPFQAYTLFLTVAINTVDLHGGD
>AL6G04060.t1
MVVIAINVATISAAVGKAIAEKWPAAKVAQATTGLLFNVWMLLLLYPFALGVMGRWSKRTYVLFMLLV
MIAVASVYIIAVLPFGISFALLTCKVVYIFPAN
>AL6G05360.t1
METETETERRLFATEILGGRTVYRLHAVTVAVGIVLVLYYRASHIPAAGEGRAAWVGMLAAELWYSAY
WVITQSVRWSPVRRHPFRDRLAARHGEMLPCVDIFVCTADPRSEPPSLVISTILSLMAYDYPAEKLSV
YLSDDGCSIFTFYAMWEATSFAKHWIPFCKRYNIEPRSPAAYFSDSDGPQEPCTPKDWSFIKDMYDE
MNERINSAIMSGKIPEEIKANHKGFCEWNLKINSKDHQSIVEILVDGKDQNAVDNEGNVLPTLVYMARE
KRPQHNHNFKAGAMNALIRVSSVISNSPIIMNVDCDMYSNNKDTIRDSLCFFLDEEMGHKIGFVQYPQ
NYNNITKNNIYGNSLQAIKKVEMGGLDSVGGPPYIGTGCFHRREILCGMIFTKDYKVDWDRGVKERIQ
ENIVYNIEEKAKSLTTCTYEHSTQWGNEIGVKYGCPVEDVITGLAIHCRGWESVYYNPIRPAFLGVAPA
TLSQTLLQTKRWSEGNFTIFLSKYCPFLFGHGKIKLRHQMGYCIYALWAPNSLATLYYVTIPSLNLIKIS
PLFPEITSPWITPFVFVFCLTNLYNLHEALSCGDTLKGWWNVQRMWMVKRITSYLFGVIDTLRKLLGL
SHMTFAVSSKVSDEDVSKRYEQEIMELGSSTPEYVIIATIALLNLVCLVGGIFQIMTCSWNVFLNVFSP
QLILCGMLVIINIPFYEAMFFRKDKGRMPFPVTLASTGFLVLAFLIEIIIPII
>AL7G01647.t1
MEASAGLVAGSHNRNELVVIRRDGEPGPRQLKQQSHRACQICGDDVGLTPDGEPFVACNECAFPVC
RDCYEYERREGTQNCPQCKTRFKRLKGCARVPGDEEEEGADDLEDEFNWRDRDDSQYAAESMLH
AHMTYGRGGDIDPHPFQPNPNVPLLTNGQMVDDIPPEQHALVPSFVGGGGKRIHPLPYADPSLPVQ
PRSMDPSKDLAAYGYGSVAWKERMESWKQKQDRLQQTRNDGGGKDWDGDGDDGDLPLMDEAR
QPLSRKIPIPSSQINPYRMIIVIRLVIVCLFFHYRVTHPVNDAFVLWLISVICEIWFAVSWILDQFPKWLPI
ERETYLDRLTLRFDKEGQPSQLAPVDFFVSTVDPAKEPPLVTANTILSILAIDYPVDKISCYVSDDGAA
MLTFEGLSETSEFAKKWVPFCKKYSIEPRAPEWYFQQKIDYLKDKVAPNFVRDRRAMKREYEEFKVR
INALVAKAQKVPEEGWTMQDGTPWPGNNVRDHPGMIQVFLGQSGGLDVEGNELPRLVYVSREKRP
GYNHHKKAGAMNALVRVSAVLTNAPYMLNLDCDHYVNNSKAVKEAMCFMMDPLVGKKCCYVQFPQ
RFDSIDRHDRYANKNVVFFDINMKGLDGIQGPIYVGTGCCFRRQALYGYDAPKTKKPPSRTCNCWPK
WCVCCFCFGNRKNKKKVTKPKTEKKKRFFFKKEENQSPAYALSEIDEAAAGAETQKAGIVNQVKLEK
KFGQSSVFVASTLLENGGTLRSASPASLLKEAIHVIGCGYEDKTDWGKEIGWIYGSVTEDILTGFKMH
CHGWRSIYCIPKRAAFKGSAPLNLSDRLNQVLRWALGSIEIFFSNHCPLWYGYGGGLKFLERFSYINSI
VYPWTSIPLLAYCTLPAICLLTGKFITPELSNLASIWYMSLFVCIFATGILEMRWARVAVDDWWRNEQF
WVIGGVSAHLFAVFQGLLKVIAGVDTSFTVTTKAGDDEEFSELYTFKWTTLLIPPTTLLLLNFIGVVAGI
SNAINNGYESWGPLFGKLFFAFWVIVHLYPFLKGLLGRQNRTPTIVIVWSILLASIISLLWVRVNPFLAK
TDGPTLEECGLDCA
>AL7G01648.t1
MLAHHTYAHQRRRPSTPTPPTPTPPHPPLPTSVPLHLHPLSALTPGAWFLDASLFCLSVPFLLLSSVL
DVPSLFAGKRIHPLPYADPSLPVQPRSMDPSKDLAAYGYGSVAWKERMESWKQKQDRLQQTRNDG
SKDWDGDGDDGDLPLMDEARQPLSRKIPIPSSQINPYRMIIVIRLVIVCLFFHYRVTHPVNDAFVLWLIS
VICEIWFAVSWILDQFPKWLPIERETYLDRLTLRFDKEGQPSQLAPVDFFVSTVDPAKEPPLVTANTILS
ILAIDYPVDKISCYVSDDGAAMLTFEGLSETSEFAKKWVPFCKKYSIEPRAPEWYFQQKIDYLKDKVAP
NFVRDRRAMKREYEEFKVRINALVAKAQKVPEEGWTMQDGTPWPGNNVRDHPGMIQVFLGQSGGL
DVEGNELPRLVYVSREKRPGYNHHKKAGAMNALVRVSAVLTNAPYMLNLDCDHYVNNSKAVKEAM
CFMMDPLVGKKCCYVQFPQRFDSIDRHDRYANKNVVFFDINMKGLDGIQGPIYVGTGCCFRRQARY
GYDAPKAKKPPSRTCNCWPKWCVCCFCFGNRKNKKKVTKPKTEKKKRFFFKKEENQSPAYALSEID
EAAAGAETQKAGIVNQVKLEKKFGQSSVFVASTLLENGGTLRSASPASLLKEAIHVIGCGYEDKTDWG
KEIGWIYGSVTEDILTGFKMHCHGWRSIYCIPKRAAFKGSAPLNLSDRLNQVLRWALGSIEIFFSNHCP
LWYGYGGGLKFLERFSYINSIVYPWTSIPLLAYCTLPAICLLTGKFITPELSNLASIWYMSLFICIFATGIL
EMRWARVAVDDWWRNEQFWVIGGVSAHLFAVFQGLLKVIAGVDTSFTVTTKAGDDEEFSELYTFKW
TTLLIPPTTLLLLNFIGVVAGISNAINNGYESWGPLFGKLFFAFWVIVHLYPFLKGLLGRQNRTPTIVIVW
SILLASIISLLWVRVNPFLAKTDGPTLEECGLDCA
>AL7G02010.t1
MAGAMVGSSSSGGGTAAKSGWHGGGQVCQICGDGVGRAADGDLFIACEVCGYPVCRPCYEYERK
DGALACPQCKTKYKHHKGSPPARGDESEDVDVDDATDLNYPPSGNQGQSHKIPEKMLTWRTNPGIS
DDVCHANKYGGGEIGLSKYDSGEIPHGCIPSFAHSQVSGEIPGASPDHMMSPAGNVGKHGHPFPYA
NHSSNPSREFSGSLGNVAWKERVDGWKMKDKGAIPMTNGTSIAPSEGHGNGDIDACTDYNTEDALL
NDEARQPLSRKVPISSSRINPYRMVIVLRLIVLCIFLHYRITNPVRNAYPLWLLSVMCEIWFAFSWILDQ
FPKWSPINRETYLDRLALRYDRDGELSQLAAVDIFVSTVDPMKEPPLVTANTVLSILAVDYPVDKVSCY

VSDDGAAMLTFDALAETSEFARKWVPFCKKYNVEPRAPEWYFSQKIDFLKDKVQTSFVKDRRAMKR
EYEEFKVRVNGLVAKAEKVPEEGWIMQDGTPWPGNNTRDHPGMIQVFLGHGGGLDSTGNELPRLV
YVSREKRHGFDHHKKAGAMNALVRVSAVLTNGQYMLNLDCDHYINNSKAIREAMCFLMDPNLGRNI
CYVQFPQRFDGIDTNDRYANKNTVFFDINLRGLDGIQGPVYVGTGCVFNRTALYGYEPPIKYKKPGFF
SSLCGGRKKTSKSSNTNNKKSHKHVDSSVPVFNLEDIEGVEDSGLDDENSLLMSQMSLEKRFGQSS
VFVASTLMEYGGVPQSATQETLLKEAIHVISCGYEDKSDWGREIGWIYGSVTEDILTGFKMHARGWR
SIYCMPNPPAFKGSAPINLSDRLDQVLRWALGSVEILFSRHCPIWYGYGGRLKFLERLAYINTTIYPLTS
IPLLIYCILPAVCLLTGKFIIPQISNIASIWFILLFISIFATGILEMRWSGVGIDEWWRNEQFWVIGGISAHLF
AVFQGLLKVLANVDTSFTVTSKASDEDSDFAELYMFKWTTLLIPPTTILIINMVGVVAGTSYAINSGYQS
WGPLFGKLFFAFWVIIHLYPFLKGLMGRQNRMPTIVIVWAILLASIFSLLWVRIDPFTTRVTGPDIQLCGI
NC
>AL1G01201.t1
MQQDGGHHHRTAAAGDVFALSVDRLAYSFSPVVVDVLEQLLLAWSFLAVPLLRGAVTLCMAMSVMV
LAEKVFLGTISAAANVLRRLRPGTGRNMHGSVDPAMLRDEEAGGGSAAFPMVLVQIPMYNEREVYQ
LSIGAACRLTWPADRLIVQVLDDSTDAAIKELVGAECERWARSGVDVRYEARHDRAGHKAGNLTEG
MRHGYARRCEFVAIFDADFQPAPGFLADTVPLLLRDPALALVQTRWEFVNADECLLTRMQEMSMDY
HFKVEQQAGSSLCSFFGYNGSAGVWRRQAIDESGGWDDRTTAEDMDLALRAALLGWKFVYVGAIK
VKSELPSSLKAYRSQQHRWSCGPALLFKKMFWEILAAKEVSIWKKFYIIYNFFVARRIVSTFFTFFFFTV
LLPAKLLFTQVRISTWQMIYIPTATTLLNAVGTPRRSLHLVIPWITFENAMALHRFKAILIGLFEAGRANE
WIVTYKSGNVGQKVKSVTPGDYLKQRFHGLELLMGAFLLMSGCYDCMYGHGYFYLFALPQSVMYFM
VGFELLGVSVCS
>AL1G02765.t1
MAPWWGQEARGGGVSGGGTPVVVKMQTPDWAISEVPPPGSPGAAGGKDGRGKNARQITWVLLLK
AHRAAGKLTGAASAALSVAAAARRRVAAGRTDSDSDADAAPRAPGEESPALRARFYGFLRAFLLFS
MLLLAVDVAAHAQGWHLPAAVPDLEDVQGLFAAAYGSWMRARVAYLGPALQFLTNACVVLFMIQTA
DRLILCLGCFWIKLRGIKPVPNVAAAAGKGADDVEAGLQDFPMVLVQIPMCNEKEVYQQSIGAVCNLD
WPRSNFLVQVLDDSDDVVTSALIKEEVEKWQREGVRILYRHRVIRDGYKAGNLKSAMNCSYVKDYEF
VVIFDADFQPQEDFLKLTVPHFKGKEDVGLVQARWSFVNKDENLLTRLQNINLCFHFEVEQQVNGAF
LNFFGFNGTAGVWRIKALEDSGGWMERTTVEDMDIAVRAHLKGWKFLYLNDVECQCELPESYEAYR
KQQHRWHSGPMQLFRLCFMDIIKSKIGFWKKFNLIFLFFLLRKLILPFYSFTLFCVILPMTMFAPEAELPI
WVVCYIPATMSLLNILPAPKSFPFIVPYLLFENTMSVTKFNAMISGLFQLGSAYEWVVTKKSGRSSEGD
LVALIENEKQPKHLRVGSAPNLDSLAKEESRLKVDPKKKKHNRLYRKELALSFLLLTAAARSLLSVQGI
HFYFLLFQGVSFLLVGLDLIGEQVTE
>AL1G05406.t1
MDGAVWTDGGAQAASARETVANLKLIAGQLLEGERFFGGDTIGYLDIALGGITQWMDVFHEIARVPLL
TEEEHPALAGGGHHQTAAALLTAAGDMFALSVDRLAADTFSPVVAAVLEQLLLAWSFLAVPLLRDAV
TLCMAMSVMVLAEKVFLGTVSAAASVLRRRGPGKGRSVRVGIDTALLRDEEAGGGNAAFPMVLVQIP
MYNEREVYQLSIGAACRLTWPADRLIVQVLDDSTDAAIKELVRAECERWSRSGVDVRYEARQDRAG
HKAGNLTEGMRHAYARRCEFVAIFDADFQPAPGFLADTVPLLLRDPALALVQTRWEFGTYLNLIKAMA
FSYTVHGPCWHSVNAETRGCKFVVNADECLLTRMQEMSMDYHFNVEQQAGSSLCSFFGYNGIVYP
LAGVWRRQAIDESGGWDDRTTAEDMDLALRAALLGWKFVYVGAIKVKSELPSSLKAYRSQQHRWS
CGPALLFKKMFWEILAAKEVLIWKKFYIIYNFFVARRIVSTFFTFFFFTVLLPAKLLFPQVRISTWQMIYIP
TATTLLNAVGTPRRSLHLVIPWITFENAMALHRFKAILIGLFEAGRANEWIVTYKSGNVGQKVKLVTPG
DYLKQRFHGLELLMGAFLLMSGCYDCMYGHGYFYLFALPQSVMYFMIGFELLGWASVSAVD
>AL2G04987.t1
MAPSFWGREARVGGGMGNGGGTPVVVKMENPNWSISEVEPDVDPGSPAGLAAGKAGRGKNARQI
TWVLLLKAHRAAGRLTGAASAALAVAAAARRRVAAGRTDSDAAPGESTALRTRFYLLIRAFLLLSVLLL
AVEVAAYFQGWHLQMPELPAGFSAEDGLVAIDGLFAAAYAGWMRVRLQYLAPPLQFLTNSCVVLFMI
QSVDRLVLCLGCLWIKLKGIKPVPLAADKDDVEAGAEDFPMVLVQMPMCNEREVYQQSIGAICNLDW
PRSNFLVQVLDDSDDATTSALIKEEVEKWQREGVHIVYRHRVIRDGYKAGNLKSAMNCSYVKDYEYV
VIFDADFQPQADFLKRAMPHFKGKDDVGLVQARWSFVNNDENLLTRLQNINLCFHFEVEQQVNGTFL
NFFGFNGTAGVWRIKALEDSGGWMERTTVEDMDIAVRAHLKGWKFLYLNDVECQCELPESYEAYRK
QQHRWHSGPMQLFRLCFVDIIKSKIGFWKKCNLIFLFFLLRKLILPFYSFTLFCVILPMTMFVPEAELPA
WVVCYVPVTMSIMSILPAPKSFPFIVPYLLFENTMSVTKFNAMVSGLFQLGSAYEWVVTKKSGRSSEG
DLVALVEKHSAPQRVGSAPNLDALAKEESLPKKDAKKKQKHNRIYRKELALSFLLLTAAARSVLSAQG
VHFYFLLFQGISFLVMGLDLIGEQVE>>AL2G05226.t1
METGGEAGGGKPEYSIIVPTYNERLNVAIIVYLIFKHLPDAKFEIIIVDDGSPDGTQDIVKQLQQVYGED
RVLLRARPRKLGLGTAYIHGLKHASGDFVVIMDADLSHHPKYLPSFIRKQKETGADIVTGTRYVNNGG
VHGWNLMRKLTSRGANVLAQTLLRPGASDLTGSFRLYKRSVLEDVISSCVSKGYVFQMEMIVRATRK
GYHIEEVPITFVDRVFGISKLGGSEIVGYLKGLVYLLLTT
>AL3G03394.t1
MAALGADAVATAWVALRARGVAPALRAAVWACLVMSAMLLVEAACMSLVSLVAVQLIRWRPERRYK

WEPMAGAATGRDVEDPPASADGCEFPRVLVQIPMYNEKEVYKLSIAAVCALTWPPDRIIIQVLDDSTD
PIIKELVELECQDWASKKIDIKYEVRNNRKGYKAGALKKGMEHAYAQQCDFVAIFDADFQPESDFLLK
TVPFLVHNPEIALVQTRWEFVNYDVCLMTRIQKMSLDYHFKVEQESGSFMHAFFGFNGTAGVWRVS
AINESGGWKDRTTVEDMDLAVRAGLKGWEFLYVGDIRVKSELPSTFKAYRHQQHRWTCGAANLFRK
MGWEIITNKGVSIWKKYHLLYSFFFVRRVIAPILTFLFYCVVIPLSAMVPEVSIPVWGLVYIPTAITIMNAI
RNPRSLHLMPFWILFENVMSMHRMRAALTGLLETAHANDWVVTEKVGDVVNDDLNVPFLEPLKPTE
RVERIYIPELLLALYLLICASYDLAFGSQTYYMYIYLQAFAFIVLGFGFVGTKTPCS>>AL4G02826.t1
MEAGEIGGALLFLLAAAAALAVAVSTGAVDFSRALTVGGRLDFQDTISWFIGVFDGSSSSESAGGVSP
ADVYELWVRVRGGVIAPVLQVAVWACMVMSVMLVVEALYNCVVSLGVKVVGWRPEWRFKWEPFA
GDDDEEKGTAAHYPTVLVQIPMYNELEVYKLSIGAVCELNWPKDRIIVQVLDDSTDPYTKNLVELECE
NWASKGMNIKYATRSSRKGFKAGALKKGMEYDYAKQCEYVTIFDADFQPEQDFLLRTVPFLMHNPE
VALVQARWAFVNDTESMLTRVQKMFFDYHFKVEQEAGSATFSFFSFNGTAGVWRAAAIKDAGGWK
DRTTVEDMDLAVRATLKGWKFIYVGDIRVKSELPSTYKAYCRQQFRWSCGGAHLFRQVAKDILAAKD
VSILKKFHMLYSFFFVRRVVAPTVTCILYNIILPISVMIPELLIPIWGIAYIPMVLTVVTAIRHPKNLHILPFWI
LFENVMTIHRMRAALAGLFELSEFNEWVVTKKVGNNFEDNEVPLLQKTRKRLRDRVNFREIVFAAFLF
FCASYNLVFHGKTSYYFNLYLQGLAFVFLGLNFTGTCTWCQ
>AL4G04310.t1
MSVFLLAAEVAAYSNGRGALAVFFGSFYASWVRVRAAYLAPPLQFLADTCVVLFLVQSADRLVQSVG
CFYILLKRIKLKPVSPALPDAEDPDAGYYPMVLVQIPMCNEKEVYQQSIAAVCNLDWPRSNILVQVLDD
SDDPITQVLIKQEVEKWRQNGARIVYRHRVLREGYKAGNLKSAMSCSYVKDYEYVAIFDADFQPYPD
FLKRTVPHFKDNEELGLVQARWSFVNKDENLLTRLQYINLCFHFEVEQQVNGVFINFFGFNGTAGVW
RIKALEDSGGWMERTTVEDMDIAVRAHLKGWKFVFLNDVECQCELPESYEAYRKQQHRWHSGPMQ
LFRLCLPDIIRCKIAFWKKANLILLFFLLRKLILPFYSFTLFCIILPMTMFVPEAELPNWVVCYIPALMSFLN
IVPAPKSFPFIIPYLLFENTMSVTKFNAMISGLFQLGSAYEWVVTKKSGRSSEGDLIALAPKGLKQQKA
DSVPVIDATIKEQPKPKKESKKYNRIYKKELALSLLLLTAAARSLLSKQGTHFYFLLFQGISFLLVGLDLI
GEDVK
>AL4G04312.t1
MAPWSGLWAGRAGGDAYRGTPVVVKMENPNWSISEIDGGGDDGEDFLPAAGGRRRRGKNAKQIT
WVLLLKAHRAAGCLAWLASAAVALGCAARRRVTAGRTDSDATEVECEGEEEAAAHAVPRSRFYAFIK
ACLLMSVFLLAAEVAAYSNGRGALAVFFGSFYASWVRVRAAYLAPPLQFLADTCVVLFLVQSADRLV
QSVGCFYILLKRIKLKPVSPALPDAEDPDAGYYPMVLVQIPMCNEKEVYQQSIAAVCNLDWPRSNILV
QVLDDSDDPITQVLIKQEVEKWRQNGARIVYRHRVLREGYKAGNLKSAMSCSYVKDYEYVAIFDADF
QPYPDFLKRTVPHFKDNEELGLVQARWSFVNKDENLLTRLQYINLCFHFEVEQQVNGVFINFFGFNG
TAGVWRIKALEDSGGWMERTTVEDMDIAVRAHLKGWKFVFLNDVECQCELPESYEAYRKQQHRWH
SGPMQLFRLCLPDIIRCKIAFWKKANLILLFFLLRKLILPFYSFTLFCIILPMTMFVPEAELPNWVVCYIPA
LMSFLNIVPAPKSFPFIIPYLLFENTMSVTKFNAMISGLFQLGSAYEWVVTKKSGRSSEGDLIALAPKGL
KQQKADSVPVIDATIKEQPKPKKESKKYNRIYKKELALSLLLLTAAARSLLSKQGTHFYFLLFQGISFLL
VGLDLIGEDVK
>AL5G01190.t1
MAGEGPASSAAARLLDGSLLLLRRWRASGGGELLLWGAWDAVRAGAVAPALAAASGACLAMSAAL
LAEAVFMAAASLARRRPERRYSADPLGACDGEDQESGLLGYPMVLVQIPMYNEREVYKLSIGAACGL
SWPSERLIVQVLDDSTDPTVKDLVELECKFWAKKGKNVKYEVRNNRRGYKAGALKEGMLRAYVQQC
DFLAVFDADFQPEADFLIRTIPYLARNPQIALVQARWEFVNPNECLMTRIQKMTLDYHFKVEQEAGSS
TFAFFGFNGTAGVWRISAIKEAGGWDDRTTVEDMDLAVRAGLKGWKFVYVGDVKVKSELPSSLKAY
RRQQHRWTCGAANLFRKTGKEILLTKEVSFWRKLYLLYSFFFVRKVVAHVVPFVLYCVVIPLSVLIPEV
TIPAWGVVYIPTTITLLYAIRNPSSIHFIPFWILFENVMSFHRTKATFIGLLELGSVNEWVVTEKLGNSSN
TKPVPQILEIKRPRCRFWDRWTISEILFAVFLFTCATYNLVHGPDLYFIYIYLQAITFLIVGTGFCGASNS
>AL5G02454.t1
MDAIAHGEEVVARMWEELPVRVDWATVPAQCAYAWAQARAFVVVPAIRLLLVLSLAMTVMILLEKLF
VAAVYFTVKAFGLKPERRYRWRPITAAAAGDGDEEACLEGGGGGAGFPVVLVQIPMYNEREVYKLSI
GAACALEWPSDRLVIQVLDDSTDPVVKDLVEIECKRWLGKGVNIKYEVRGNRKGYKAGALKEGLKHS
YVHECEYIAMFDADFQPESDFLLRTVPFLVHNADIALVQTRWKFVNSDECLLTRFQEMSLDYHFKFEQ
EAGSVVYSFFGFNGTAGVWRISAIDDAGGWKDRTTVEDMDLAVRTTLQGWKFVYVGDVKVRSELPS
TFKAYRFQQHRWSCGPANLFKKMMVEILENKKVSFWSKLHLLYDFFFVGKIAAHTVTFIYYCFAIPVSV
FFPEIQIPLWGMVYVPTVITLCKAIGSPSSFHLVILWVLFENVMSLHRIKAATTGLLDAGRVNEWVVTEK
LGDANKAKSATDGLEAVKVIDVELTTPLVPKLKKWRLRFWDKYNCSEIFVGTYIIICGCYDLFYANKGY
YIYLFVQGLAFLVVGFEYIGTRPPTAE
>AL5G04027.t1
MDAAEQLEAMWRQVRGPVIVPLLRASVAVCLAMCVILFVEKVYMAVVITAVRLLGRRPERQWRWEA
VQEDDADPECGGGNSAYPMVLVQIPMYNEREVYQLSIGAACGLSWPSDRIVIQVLDDSTDHVIKELV
QMEVQRWARKGVNIKYEIRDNRRGYKAGALKEGMKHGYVRDCDFVAIFDADFQPEADFLRRAMPFF

IHNPEIALVQGRWRFVNADECLMTRMQEMSLDYHFKVEQEVGSSTYAFFGFNGTAGVWRISALNEA
GGWKDRTTVEDMDLAVRASLKGWKFVYLGDLMKEPGAELVNVTFR
>AL6G01955.t1
MDAAVGLPDAWSQVRAPLIVPLLRLAVAACLLMSVLLFLERLYMGVVIVGVKLLRRTPERRYKCDPLS
EDDDPELGSAAFPVVLVQIPMFNEREVYHLSIGAVCGLSWPSDRLVVQVLDDSTDPVIKEMVRMECE
RWAHKGINITYQIREDRKGYKAGALKQGMKHGYVRECEYMVIFDADFQPDPDFLHRTIPYLYHNPDIA
LVQARWRFVNADECLMTRMQEMSLDYHFKVEQEVSSSVCAFFGFNGTAGVWRISAVNEAGGWND
RTTVEDMDLAIRASLKGWKFVYLGDVQVKSELPSTFKAFRFQQHRWSCGPANLFRKMLLEIVTNKKV
TIWKKFHVIYNFFLVRKIVAHIVTFTFYCIVIPATIFVPEVHIPKWGCIYIPSIITLLNSVGTPRSFHLLFFWIL
FENVMSLHRTKATLIGLLEAGRVNEWVVTEKLGNVLKMKSSNKGSAKKSFMRMWERLNVPELGVSA
FLFSCGWYDLAFGGKDNFFIYLFFQSIAFFVVGIGYVGTIVPQS
>AL6G04010.t1
MLTMARAAWAGVRYTVVVPLLHVAVYLCAAMSLMLFVERLYMGIVGAVLWLNRRRRRRGADAHDD
DKKDDSDDLEAGAGAEKPMVLVQIPMFNEKQVYRLSIGAACGLWWPSEKLVIQVLDDSTDGAVRALV
EAECRRWASKGVRVHYENRSNRNGYKAGAMREGLKKGYAKDCQFVAVFDADFQPDADFLRRTVPL
LQADPAVALVQARWRFVNADECILTRIQEMSLDYHFSVEQEVGSACHAFFGFNGTAGVWRVQALAD
AGGWKDRTTVEDMDLAVRASMRGWKFVFTGDVKVRNELPSTFKAYRYQQHRWSCGPANLMRKMF
WEILTSRQVSTWKKLHVLYGFFFVRKVVAHLVTFLFYCVVIPACVLVQGDVRLPKYVAMYVPAIITLLN
AVCTPRSWHLLVFWILFENVMSMHRSKATIIGLLEASRANEWVVTEKLGSTTTTTVATKKKNQNQGF
HTAEIVLGICLLYCAIYDIIFGHDHFYVYLLMQSAAAFVIGLDYVGSQ
>AL6G04028.t1
MLTMARAAWAGVRYTVVVPLLHVAVYLCAAMSLMLFVERLYMGIVGAVLWLNRRRRRRGADAHDD
DKKDDSDDLEAGAGAEKPMVLVQIPMFNEKQVYRLSIGAACGLWWPSEKLVIQVLDDSTDGAVRALV
EAECRRWASKGVRVHYENRSNRNGYKAGAMREGLKKGYAKDCQFVAVFDADFQPDADFLRRTVPL
LQADPAVALVQARWRFVNADECILTRIQEMSLDYHFSVEQEVGSACHAFFGFNGTAGVWRVQALAD
AGGWKDRTTVEDMDLAVRASMRGWKFVFTGDVKVRNELPSTFKAYRYQQHRWSCGPANLMRKMF
WEILTSRQVSTWKKLHVLYGFFFVRKVVAHLVTFLFYCVVIPACVLVQGDVRLPKYVAMYVPAIITLLN
AVCTPRSWHLLVFWILFENVMSMHRSKATIIGLLEASRANEWVVTEKLGSTTTTTVATKKKKNQNQGF
HTAEIVLGICLLYCAIYDIIFGHDHFYVYLLMQSAAAFVIGLDYVGSQ
>AL7G01826.t1
METGGEAGGGKPEYSIIVPTYNERLNVAIIVYLIFKHLQDAKFEIIIVDDGSPDGTQDIVKQLQQVYGEE
RVLLRARPRKLGLGTAYVHGLKHASGDFVVIMDADLSHHPKYLPSFIRKQKLTGADIVTGTRYVNNGG
VHGWNLMRKLTSRGANVLAQTLLRPGASDLTGSFRLYKRSVLEDVISSCVSKGYVFQMEMIVRATRK
GYHIEEVPITFVDRVFGISKLGGSEIVGYLKGLVYLLLTT
>AL7G01858.t1
MAGAAWPLPFAADLLPASLSLTLLLASLVAVLGLGIAAVFFEHIRKIGCMHSLERTAVSDAFFEDPGSL
KKVPCPSIFDPSEKYISLIMPAYNEEYRLPEALTETLNYLKQRSVADKSFSYEVLIVDDGSTDRTSKVAF
EYVKKHNIDNVRVLLLGRNHGKGEAVRKGMLHARGELLLMLDADGATQVTDLAKLEAQIHALSEKVG
SSLAPSASSQRLSDVEIAVFGSRAHLEKQALATRKWYRNFLMKGFHLVVLLTAGPGIRDTQCGFKMF
TRSAARKLFTNIRLKRWCFDVEIVYLCKRLRIPMTEVSVSWTEIPGSKVRMTSILHMVFELLLIRVGYGL
GIWKIYS
>AL7G02472.t1
MAPWSGLWGGRTGGDAYRGTPVVVKMENPNWSISEISPDDDDEDFLVDAGATGGGRRKGGRGKN
AKQITWVLLLKAHRAAGCLASLASTAVALGAAARRRVADGRTDADAGAAGAAGESPVLRSRFYAFIR
AFLVLSMLLLAVELAAHFNGWNLAASALALPIIGVESLYGSWLRLRAAYLAPLLQFLTDACVVLFLIQSA
DRLIQCLGCFYINLKRIKPKLKSAALPDAEDPDAGYYPMVLVQIPMCNEKEVYQQSIAAVCNLDWPRS
NLLVQVLDDSDDPTTQSLIKEEVAKWQQNGARILYRHRVLRDGYKAGNLKSAMSCSYVKDYEFVAIF
DADFQPNPDFLKRSVPHFKDNDELGLVQARWSFVNKDENLLTRLQNINLCFHFEVEQQVNGVFLNFF
GFNGTAGVWRIKALEDSGGWMERTTVEDMDIAVRAHLHGWKFIFLNDVECQCELPESYEAYRKQQH
RWHSGPMQLFRLCLPDIIKSKISVWKKANLIFLFFLLRKLILPFYSFTLFCIILPMTMFVPEAELPDWVVC
YIPALMSLLNILPSPKSFPFIIPYLLFENTMSVTKFNAMISGLFQLGSAYEWVVTKKSGRSSEGDLIALAV
VPPKELRHQKTGSVPNLDLVIMKDPSSPPTMNDAKKKKKKHNRIYKKELVLSLLLLTAAARSLLTKQGI
HFYFLLFQGISFLLVGLDLIGEQVE
>GRMZM2G015886_P01
MSRRLSLPAGSQVTVTVSPTRGKAESPGDGVIKRSGGRIGLTSPAPRHSLGGAGSSTIPTVQLSPVR
RSGGSRYASLDGGAGADDSAEFVHYTVHIPPTPERTVAASADSVDAPAPTAYDEDGGAAGVRAQRS
YISGTIFTGGLNQATRGHVLNTSANSAAVAASANMSCKMRGCDMPAFLSSGAGGGPCDCGFMICRE
CYADCVAAAGNCPGCKEPYSAGSDTDDDGEDDEAVSSSEERDQLPLTSMAKRFSLIHSMKMPSNN
GGGGGKPAEFDHARWLFETKGTYGYGNALWPKDGHGGGGGGGGFSGFEEPPNFGSRCRRPLTR
KTSISQAILSPYRLLIAIRLVALGFFLTWRIRHPNPEAVWLWALSVTCEVWFAFSWLLDSLPKLCPIHRA
ADLDVLAERFELPTARNPKGHSDLPGIDVFVSTADPEKEPPLVTANTILSILAADYPVEKLACYLSDDG
GALLTFEALAETASFARTWVPFCRKHGVEPRCPEAYFGQKRDFLRNKVRVDFVRERRKVKREYDEF

KVRVNSLPEAIRRRSDAYNAGEELRARRRQQEEAMAAGTILGALPEAAGAVKATWMSDGSQWPGT
WLTSAPDHSRGDHAGIIQAMLAPPTSEPVLGAEPAESGGLIDTTGVDIRLPMLVYVSREKRPGYDHNK
KAGAMNALVRTSAIMSNGPFILNLDCDHYVHNSAALREGMCFMLDRGGDRVCYVQFPQRFEGIDPN
DRYANHNLVFFDVAMRAMDGLQGPMYVGTGCVFRRTALYGFSPPRATEHHGWLGRRKIKLLLRKPT
MGKKTDRENNSDKEMMLPPIEDDAFQQLDDIESSALLPRRFGSSATFVASIPVAEYQGRLLQDTPGA
HQGRPAGALAVPREPLDADTVAEAISVISCFYEDKTEWGRRIGWIYGSVTEDVVTGYRMHNRGWRS
VYCVTRRDAFRGTAPINLTDRLHQVLRWATGSVEIFFSRNNALFASPRMKFLQRVAYFNVGMYPFTSI
FLLVYCVLPAVSLFSGKFIVQSLNATFLALLLVITITLCLLALLEIKWSGITLHEWWRNEQFWVIGGTSAH
PAAVLQGLLKVIAGVDISFTLTSKPGTGDDGEEDAFAELYEVRWSFLMVPPVTIMMVNAVAVAVASAR
TLYSEFPQWSKLLGGAFFSFWVLCHLYPFAKGLLGRRGRVPTIVFVWSGLISMTISLLWVYISPPAGA
RELIGGGGFSFP
>OS12G36890_OsCslD4
MSRRLSLPAGAPVTVAVSPVRSPGGDAVVRRGSGLTSPVPRHSLGSSTATLQVSPVRRSGGSRYLG
ASRDGGADESAEFVHYTVHIPPTPDRATASVASEAEAAAEAEEVHRPQRSYISGTIFTGGLNCATRG
HVLNFSGEGGATAASRAAASGNMSCKMRGCDMPAFLNGGRPPCDCGFMICKECYAECAAGNCPG
CKEAFSAGSDTDESDSVTDDDDDEAVSSSEERDQLPLTSMARKFSVVHSMKVPGAAANGNGKPAE
FDHARWLFETKGTYGYGNALWPKDGHAHSGAGFVAADEPPNFGARCRRPLTRKTSVSQAILSPYRL
LIAIRLVALGFFLAWRIRHPNPEAVWLWAMSVACEVWFAFSWLLDSLPKLCPVHRAADLAVLAERFES
PTARNPKGRSDLPGIDVFVTSADPEKEPPLVTANTILSILAADYPVEKLACYLSDDGGALLSFEALAET
ASFARTWVPFCRKHGVEPRCPEAYFGQKRDFLKNKVRVDFVRERRKVKREYDEFKVRVNSLPEAIR
RRSDAYNAGEELRARRRQQEEAAAAAAAGNGELGAAAVETAAVKATWMSDGSHWPGTWTCPAAD
HARGDHAGIIQAMLAPPTSEPVMGGEAAECGGLIDTTGVDVRLPMLVYVSREKRPGYDHNKKAGAM
NALVRTSAIMSNGPFILNLDCDHYVHNSSALREGMCFMLDRGGDRVCFVQFPQRFEGVDPSDRYAN
HNLVFFDVSMRAMDGLQGPMYVGTGCVFRRTALYGFSPPRATEHHGWLGRRKIKLFLTKKKSMGKK
TDRAEDDTEMMLPPIEDDDGGADIEASAMLPKRFGGSATFVASIPVAEYQGRLLQDTPGCHHGRPA
GALAVPREPLDAATVAEAIGVISCFYEEKTEWGRRIGWIYGSVTEDVVTGYRMHNRGWRSVYCVTPR
RDAFRGTAPINLTDRLHQVLRWATGSVEIFFSRNNALFASPRMKLLQRVAYFNAGMYPFTSVFLLAYC
LLPAVSLFSGKFIVQRLSATFLAFLLVITLTLCLLALLEIKWSGITLHEWWRNEQFWVIGGTSAHPAAVL
QGLLKVIAGVDISFTLTSKPGNGGGDGGVGGEGNDDEAFAELYEVRWSYLMVPPVTIMMVNAVAIAV
AAARTLYSEFPQWSKLLGGAFFSFWVLCHLYPFAKGLLGRRGRVPTIVFVWSGLISMIISLLWVYINPP
AGARERIGGGGFSFP
>CSLD_1BL_TGACv1_030586_AA0094860.1_Triticum_aestivum
MGSKGILKNSGSSRMPPHGPSKPPTAPTSAPQVVFGRRTESGRFISYSRDDLDSEISSVDFQDYHVH
IPMTPDNQPMEEDGTKADEQYVSSSLFTGGFNSVTRAHVMDKQGPDSDIGRSGPKGSICMVEGCDS
KIMRNGRGEDILPCECDFKICVDCFTDAVKGGGGVCPGCKELYKHTEWEEVLSNSSNELTRALSLPH
GPGGKMERRLSLVKQGTMNNQSGEFDHNRWLFETKGTYGYGNAIWPDDNVDDDGRNGVPGHPKE
LMSKPWRPLTRKLQIPAAVISPYRLLVLIRLVALAFFLMWRIKHQNDDAIWLWGMSIVCELWFAFSWVL
DQLPKLCPINRATDLSVLKEKFETPTPSNPTGKSDLPGIDIFVSTADPEKEPVLVTANTILSILAVDYPVD
KLACYVSDDGGALLTFEAMAEAASFANFWVPFCRKHDIEPRNPDSYFNLKRDPFKNKVKADFVKDRR
RIKREYDEFKVRVNGLPDSIRRRSDAYHAREEIQAMNLQREKIKAGGDEQFEPVKIPKATWMADSTH
WPGTWIHSSQDHARGDHAGIIQVMLKPPSDMPMYGNIEKSPLDFSEVDTRLPMLVYMSREKRPGYD
HNKKAGAMNALVRASAIMSNGPFILNLDCDHYVYNSKAFREGMCFMMDRGGDRLCYVQFPQRFEGI
DPSDRYANHNTVFFDINMRALDGLQGPVYVGTGCLFRRIALYGFDPPRSKDHSPGFCGCCLPRRRK
ASASNANPEETMALRMGDFDGDSMNLATFPKKFGNSSFLIDSIPVAEFQGRPLADHPSVKNGRPPGA
LTIPREILDASIVAEAISVVSCWYEEKTEWGTRVGWIYGSVTEDVVTGYRMHNRGWKSVYCVTQRDA
FRGTAPINLTDRLHQVLRWATGSVEIFFSRNNALFASSKMKVLQRIAYLNVGIYPFTSIFLIVYCFLPALS
LFSGQFIVQTLNVTFLTYLLIITITLCLLAMLEIKWSGIALEEWWRNEQFWLIGGTSAHLAAVMQGLLKV
VAGIEISFTLTSKQVGDDIDDEFAELYEVKWTSLMIPPLTIIMVNLVAIAVGFSRTIYSTDIDDEFAELYEV
KWTSLMIPPLTIIMVNLVAIAVGFSRTIYSTIPQWSKLLGGVFFSFWVLAHLYPFAKGLMGRRGRTPTIV
YVWAGLVSITISLLWIAINPPSSAANQQLGGSFSFP
>AT1G02730_CslD5
MVKSAASQSPSPVTITVTPCKGSGDRSLGLTSPIPRASVITNQNSPLSSRATRRTSISSGNRRSNGDE
GRYCSMSVEDLTAETTNSECVLSYTVHIPPTPDHQTVFASQESEEDEMLKGNSNQKSFLSGTIFTGG
FKSVTRGHVIDCSMDRADPEKKSGQICWLKGCDEKVVHGRCECGFRICRDCYFDCITSGGGNCPGC
KEPYRDINDDPETEEEDEEDEAKPLPQMGESKLDKRLSVVKSFKAQNQAGDFDHTRWLFETKGTYG
YGNAVWPKDGYGIGSGGGGNGYETPPEFGERSKRPLTRKVSVSAAIISPYRLLIALRLVALGLFLTWR
VRHPNREAMWLWGMSTTCELWFALSWLLDQLPKLCPVNRLTDLGVLKERFESPNLRNPKGRSDLP
GIDVFVSTADPEKEPPLVTANTILSILAVDYPVEKLACYLSDDGGALLTFEALAQTASFASTWVPFCRK
HNIEPRNPEAYFGQKRNFLKNKVRLDFVRERRRVKREYDEFKVRINSLPEAIRRRSDAYNVHEELRAK
KKQMEMMMGNNPQETVIVPKATWMSDGSHWPGTWSSGETDNSRGDHAGIIQAMLAPPNAEPVYG
AEADAENLIDTTDVDIRLPMLVYVSREKRPGYDHNKKAGAMNALVRTSAIMSNGPFILNLDCDHYIYNS
MALREGMCFMLDRGGDRICYVQFPQRFEGIDPNDRYANHNTVFFDVSMRALDGLQGPMYVGTGCIF

RRTALYGFSPPRATEHHGWLGRRKVKISLRRPKAMMKKDDEVSLPINGEYNEEENDDGDIESLLLPK
RFGNSNSFVASIPVAEYQGRLIQDLQGKGKNSRPAGSLAVPREPLDAATVAEAISVISCFYEDKTEWG
KRVGWIYGSVTEDVVTGYRMHNRGWRSIYCVTKRDAFRGTAPINLTDRLHQVLRWATGSVEIFFSRN
NAIFATRRMKFLQRVAYFNVGMYPFTSLFLIVYCILPAISLFSGQFIVQSLDITFLIYLLSITLTLCMLSLLEI
KWSGITLHEWWRNEQFWVIGGTSAHPAAVLQGLLKVIAGVDISFTLTSKSSAPEDGDDEFADLYVVK
WSFLMVPPLTIMMVNMIAIAVGLARTLYSPFPQWSKLVGGVFFSFWVLCHLYPFAKGLMGRRGRVPT
IVFVWSGLLSIIVSLLWVYINPPSGKQDYMQFQFP
>GRMZM2G052149_P01
MASTGGGGGLRHSNSSRLSRMSFSGGGSDDARAQAAAAPGGDRPMVTFARRTRSGRYVSYSRDD
LDSELGAAGVDMSPDRDEQFASYHVHIPATPDNQPMDPAISARVEEQYVSNSLFTGGFNSVTRAHL
MDKVIDSEASHPQMAGARGSSCAVNGCDANVMSDERGDDILPCECDFKICAECFADAVKNAGAICP
GCKEPYKNTELDDVVGAAADATAGGRPTLSLPPPPGAAASRMERRLSIMRSQKAMTRSQTGDWDH
NRWLFETKGTYGYGNAIWPKENEVDAAGGLGGGGADGQPAEFTTKPWRPLTRKLSIPAGVLSPYRL
LILIRMAVLGLFLTWRIKHKNEDAIWLWGMSVVCELWFGFSWLLDQLPKLCPVNRATDLAVLKDKFET
PTPSNPTGRSDLPGLDIFVSTADPEKEPPLVTANTILSILAADYPVEKLSCYVSDDGGALLTFEAMAEA
ASFANMWVPFCRKHNIEPRNPDSYFNLKKDPYKNKVRQDFVKDRRRVKREYDEFKVRINGLPDSIRR
RSDAYHAREEIKAMKRQREAALDDAVEPVKIPKATWMADGTHWPGTWIQPSAEHTRGDHAGIIQVM
LKPPSDDPLYGSTGDEGRPLDFTEVDIRLPMLVYVSREKRPGYDHNKKAGAMNALVRSSAVMSNGP
FILNLDCDHYVYNSQAFREGMCFMMDRGGDRIGYVQFPQRFEGIDPSDRYANHNTVFFDVNMRALD
GLMGPVYVGTGCLFRRVALYGFDPPRSKEHGGCCSCCFPQRRKIKASAAAPEETRALRMADFDEDE
MNMSSFPKKFGNSSFLIDSIPIAEFQGRPLADHPGVKNGRPPGALTVPRDLLDASTVAEAVSVISCWY
EDKTEWGHRVGWIYGSVTEDVVTGYRMHNRGWKSVYCVTKRDAFRGTAPINLTDRLHQVLRWATG
SVEIFFSRNNALLASRRMKFLQRIAYLNVGIYPFTSIFLIVYCFLPALSLFSGQFIVKTLNVTFLTYLLVITL
TLCLLAVLEIKWSGISLEEWWRNEQFWLIGGTSAHLAAVLQGLLKVVAGIEISFTLTSKSGGDDVDDEF
ADLYIVKWTSLMIPPIVIMMVNLIGIAVGFSRTIYSEIPQWSKLLGGVFFSFWVLAHLYPFAKGLMGRRG
RTPTIVFVWAGLLSITISLLWVAINPPSQNQQIGGSFTFP
>OS06G02180_OsCslD2
MASSGGGGLRHSNSSRLSRMSYSGEDGRAQAPGGGGDRPMVTFARRTHSGRYVSYSRDDLDSEL
GNSGDMSPESGQEFLNYHVTIPATPDNQPMDPAISARVEEQYVSNSLFTGGFNSVTRAHLMDKVIES
EASHPQMAGAKGSSCAINGCDAKVMSDERGDDILPCECDFKICADCFADAVKNGGACPGCKDPYKA
TELDDVVGARPTLSLPPPPGGLPASRMERRLSIMRSQKAMTRSQTGDWDHNRWLFETKGTYGYGN
AIWPKENEVDNGGGGGGGGGLGGGDGQPAEFTSKPWRPLTRKLKIPAGVLSPYRLLILIRMAVLGLF
LAWRIKHKNEDAMWLWGMSVVCELWFGLSWLLDQLPKLCPVNRATDLAVLKDKFETPTPSNPNGR
SDLPGLDIFVSTADPEKEPPLVTANTILSILAADYPVEKLSCYVSDDGGALLTFEAMAEAASFANMWVP
FCRKHDIEPRNPESYFNLKRDPYKNKVRSDFVKDRRRVKREYDEFKVRINSLPDSIRRRSDAYHARE
EIKAMKRQREAALDDVVEAVKIPKATWMADGTHWPGTWIQPSAEHARGDHAGIIQVMLKPPSDDPLY
GTSGEEGRPLDFTEVDIRLPMLVYVSREKRPGYDHNKKAGAMNALVRSSAVMSNGPFILNLDCDHYV
YNSQAFREGMCFMMDRGGDRIGYVQFPQRFEGIDPSDRYANHNTVFFDVNMRALDGIMGPVYVGT
GCLFRRIALYGFDPPRSKEHSGCCSCCFPQRRKVKTSTVASEERQALRMADFDDEEMNMSQFPKKF
GNSNFLINSIPIAEFQGRPLADHPGVKNGRPPGALTVPRDLLDASTVAEAISVISCWYEDKTEWGQRV
GWIYGSVTEDVVTGYRMHNRGWKSVYCVTKRDAFRGTAPINLTDRLHQVLRWATGSVEIFFSRNNA
LLASRKMKFLQRIAYLNVGIYPFTSIFLIVYCFLPALSLFSGQFIVRTLNVTFLTYLLVITLTMCMLAVLEIK
WSGISLEEWWRNEQFWLIGGTSAHLAAVLQGLLKVIAGIEISFTLTSKSGGDEADDEFADLYIVKWTSL
MIPPIVIMMVNLIAIAVGFSRTIYSEIPQWSKLLGGVFFSFWVLAHLYPFAKGLMGRRGRTPTIVFVWSG
LLAITISLLWVAINPPSQNSQIGGSFTFP
>OS02G49330_OsCslE2
MATAAALALHTQFRPPRSPRRLRQHLALPSGVLIRSPVRASAASASASAPAQREAAAAGVPWGCEIE
SLESAVSLERWLTDSGLPEQRLGIQRVDVGERGLVALKNIRKGEKLLFVPPSLVITADSEWGCPEVGN
VLKRNSVPDWPLIATYLISEASLESSSRWSSYIAALPRQPYSLLYWTRPELDAYLVASPIRERAIQRITD
VVGTYNDLRDRIFSKHSDLFPEEVYNLETFRWSFGILFSRLVRLPSMDGRVALVPWADMLNHSPEVE
TFLDYDKSSGGIVFTTDRSYQPGEQVFISYGKKSSGELLLSYGFVPKEGTNPNDSVELLVSLNKSDKC
YKEKLQALKRNGLSEFESFPLRVTGWPVELMAYAFLVVSPPEMSQRFEEMAVAASNKSPSKPGLNY
PELEEQALQFILDCCESNIAKYTKFLEGSSGSLQLSTNSKQANRTLLLKQLARDLCISERRILYRTQYVV
FVADEAQTVEEKEIDKSDYNLSRQATDARRLEVVGGSMAGSGGGVVSGGRQRGPPLFATEKPGRM
AMAAYRVSAATVFAGVLLIWLYRATHLPPGGGDGVRRWAWLGMLAAELWFGFYWVLTLSVRWCPV
YRRTFKDRLAQRILIANSYSEDELPSVDIFVCTADPTAEPPMLVISTVLSVMAYDYLPEKLNIYLSDDAG
SVLTFYVLCEASEFAKHWIPFCKKYKVEPRSPAAYFAKVASPPDGCGPKEWFTMKELYKDMTDRVN
SVVNSGRIPEVPRCHSRGFSQWNENFTSSDHPSIVQIRVSSVISNSPIIMNVDCDMYSNNSESIRDAL
CFFLDEEQGQDIGFVQYPQNFENVVHNDIYGHPINVVNELDHPCLDGWGGMCYYGTGCFHRREALC
GRIYSQEYKEDWTRVAGRTEDANELEEMGRSLVTCTYEHNTIWGIEKGVRYGCPLEDVTTGLQIQCR
GWRSVYYNPKRKGFLGMTPTSLGQILVLYKRWTEGFLQISLSRYSPFLLGHGKIKLGLQMGYSVCGF
WAVNSFPTLYYVTIPSLCFLNGISLFPEKTSPWFIPFAYVMVAAYSCSLAESLQCGDSAVEWWNAQR

MWLIRRITSYLLATIDTFRRILGISESGFNLTVKVTDLQALERYKKGMMEFGSFSAMFVILTTVALLNLA
CMVGVKCTASTSTCEMGAI
>GRMZM2G436299_P01
MASKGILKNSGSRRMPPQGAGSRPPTAPTSAPTVVFGRRTDSGRFISYSRDDLDSEISSVDFQDYHV
HIPMTPDNQPMDEDDSGTARAEDQRYVSGSLFTGGFNTVTRAHVMDKPDGDAAGRRGGGGAKGP
SACMVEGCDARAMRDARGDDVLPCECDFRVCVDCFTDAVKAGAGVCPGCKEPYKNTEWEGLGAG
AAEEVTRALSLPRGPAGANGHRMERRLSLVKQANVNQSGEFDHNLWLFQTKGTYGYGNAIWPQDE
ADDDTDGGAPAGHPKELLTKPWRPLTRKLRIPAAVISPYRLLVLIRLVALAFFLMWRIKHQNEDAIWLW
GMSIVCELWFAFSWVLDQLPKLCPINRATDLSVLKEKFETPTPNNPTGKSDLPGVDIFVSTADPEKEP
VLVTANTILSILAADYPVEKLACYLSDDGGALLTFEAMAEAASFANLWVPFCRKHDIEPRNPDSYFNLR
RDPFKNKVKPDFVKDRRRVKREYDEFKVRVNGLPDAIRRRSDAYHAREEIQAMNLQREKLKGGGDE
PPFEPVKIPKATWMADGTHWPGTWLQPSQDHARGDHAGIIQVMLKPPSDMPTTMYDAASKTPLDLA
GVDTRLPMLVYMSREKRPGYDHNKKAGAMNALVRASAIMSNGPFILNLDCDHYIYNSKALREGMCF
MMDRGGDRLCYVQFPQRFEGIDPSDRYANHNTVFFDVNMRALDGLQGPVYVGTGCLFRRIALYGFD
PPRSKDHSPGFCSCCLPRRRKPSAASREETMALRMGGFDGDSMDLATFPKKFGNSSFLIDSIPVAEF
QGRPLADHPSVKNGRPPGALTIPREMLDASIVAEAISVVSCWYEEKTEWGIRVGWIYGSVTEDVVTG
YRMHNRGWKSVYCVTQRDAFRGTAPINLTDRLHQVLRWATGSVEIFFSRNNALFASSKMKVLQRIAY
LNVGIYPFTSIFLIVYCFLPALSLFSGQFIVQTLNVTFLTYLLIITVTLCLLAMLEIKWSGIALEEWWRNEQ
FWLIGGTSAHLAAVLQGLLKVVAGIEISFTLTSKQVGDDVEDEFAELYIVKWTSLMIPPLTIIMINLVAIAV
GFSRTIYSTIPQWSKLLGGVFFSFWVLAHLYPFAKGLMGRRGRTPTIVYVWSGLVSITISLLWIAIKPPS
QAANSQFGGSFSFP
>GRMZM2G044269_P01
MSNPPPKKAIRNPGGGAGGPAGGPRGPAGNTVKFARRTSSGRYVSLSREDIDMEGELAADYTNYTV
QIPPTPDNQPMMDPASVAMKAEEQYVSNSLFTGGFNSVTRAHLMDKVIESEVTHPQMAGSKGSRCA
MPACDGKVMRNERGEDIDPCECRFKICRDCYLDAQKEGCLCPGCKEHYKIGEYADDDPNDASSGKH
YLPGPGGGMMNNSKSLLARNQNGEFDHNRWLFESSGTYGYGNAFWPKGGMYDDDLDDEGGPGG
GGGDGMLPEQKPFKPLTRKIPMPTSIISPYRIFIVIRMFVLIFYLTWRVRNPNMEALWLWGMSIVCELW
FAFSWLLDMLPKVNPVNRSTDLAVLKEKFETPSPSNPHGRSDLPGLDVFVSTADPDKEPVLTTATTIL
SILAADYPVEKLACYVSDDGGALLTFEAMAEAASFANIWVPFCKKHDIEPRQPDSYFSIKGDPTKGKR
RSDFVKDRRKVKREFDEFKVRINGLPDSIRRRSDAFNAREDMKMLKHLRETGADPAEQPKVKKATW
MADGTHWPGTWAVSAPDHAKGNHAGILQVMLKPPSPDPLYGMHDEEQLIDFSDVDIRLPMLVYMSR
EKRPGYDHNKKAGAMNALVRCSAVMSNGPFILNFDCDHYINYAQAIREAMCFVMDRGGERIAYIQFP
QRFEGIDPSDRYANNNTVFFDGNMRALDGLQGPMYVGTGCMFRRFALYGFDPPRTTEYTGWLFKK
KKVTTFGKADQGETDTQSLNSKGAEDFDAELTSMLVPRRFGNSSALMASIPVAEFQARPLADHPAVL
HGRPPGSLTVPRPPLDPPTVAEAVSVISCWYEDKTEWGDRVGWIYGSVTEDVVSGYRMHNRGWRS
VYCIPKRDAFLGTAPINLTDRLHQVLRWATGSVEIFFSRNNAFLASRRLMFLQRVAYLNVGIYPFTSIFL
LVYCFIPALSLFSGFFIVQTLNVAFLCYLLTITVTLIALGVLEVKWSGIELEDWWRNEQFWLISGTSAHLY
AVVQGLLKVMAGIEISFTLTAKAAVDDNEDIYADLYVVKWSSLLIPPITIGMINVIAIAFAFARTVYSDNPR
WGKFIGGGFFSFWVLAHLYPFAKGLMGRRGKTPTIVFVWSGLISITISLLWVAISPPEASAGGRGAGF
QFP
>CSLD_1AL_TGACv1_001700_AA0034150.2_Triticum_aestivum
MGSKGILKNSGSSRVPPHGPSKPPTAPTSAPQVVFGRRTESGRFISYSRDDLDSEISSVDFQDYHVHI
PMTPDNQPMEEDGTKADEQYVSSSLFTGGFNSVTRAHVMDKQGPDSDMGRSGPKGSICMVEGCD
SKIMRNGRGEDILPCECDFKICVDCFTDAVKGGGGVCPGCKELYKHTEWEEVLSNSSNELTRALSLP
HGPGGKMERRLSLVKQGTMNNQSGEFDHNRWLFETKGTYGYGNAIWPDDNVDDDGRNGVPGHPK
ELMSKPWRPLTRKLQIPAAVISPYRLLVLIRLVALAFFLMWRIKHQNDDAIWLWGMSIVCELWFALSW
VLDQLPKLCPINRATDLSVLKEKFETPTPSNPTGKSDLPGIDIFVSTADPEKEPVLVTANTILSILAVDYP
VDKLACYVSDDGGALLTFEAMAEAASFANFWVPFCRKHDIEPRNPDSYFNLKRDPFKNKVKADFVKD
RRRIKREYDEFKVRVNGLPDSIRRRSDAYHAREEIQAMNLQREKIKAGGDEQFEPVKIPKATWMADS
THWPGTWIHPSQDHARGDHAGIIQVMLKPPSDMPMYGNIEKSPLDFSGVDTRLPMLVYMSREKRPG
YDHNKKAGAMNALVRASAIMSNGPFILNLDCDHYVYNSKAFREGMCFMMDRGGDRLCYVQFPQRF
EGIDPSDRYANHNTVFFDINMRALDGLQGPVYVGTGCLFRRIALYGFDPPRSKDHSPGFCGCCLPRR
RKASASNANPEETMALRMGDFDGDSMNLATFPKKFGNSSFLIDSIPVAEFQGRPLADHPSVKNGRPP
GALTIPREILDASIVAEAISVVSCWYEEKTEWGTRVGWIYGSVTEDVVTGYRMHNRGWKSVYCVTQR
DAFRGTAPINLTDRLHQVLRWATGSVEIFFSRNNALFASSKMKVLQRIAYLNVGIYPFTSIFLIVYCFLP
ALSLFSGQFIVQTLNVTFLTYLLIITVTLCLLAMLEIKWSGIALEEWWRNEQFWLIGGTSAHLAAVMQGL
LKVVAGIEISFTLTSKQVGDDIDDEFAELYEVKWTSLMIPPLTIIMVNLVAIAVGFSRTIYSTIPQWSKLLG
GVFFSFWVLAHLYPFAKGLMGRRGRTPTIVYVWAGLVSITISLLWIAINPPSTAANQQLGGSFSFP
>AT3G03050_CslD3
MASNNHFMNSRSNLSTNSDAAEAERHQQPVSNSVTFARRTPSGRYVNYSRDDLDSELGSVDLTGY
SVHIPPTPDNQPMDPSISQKVEEQYVSNSLFTGGFNSVTRAHLMEKVIDTETSHPQMAGAKGSSCAV
PGCDVKVMSDERGQDLLPCECDFKICRDCFMDAVKTGGMCPGCKEPYRNTDLADFADNNKQQRP

MLPPPAGGSKMDRRLSLMKSTKSGLMRSQTGDFDHNRWLFETSGTYGFGNAFWTKDGNFGSDKD
GNGHGMGPQDLMSRPWRPLTRKLQIPAAVISPYRLLILIRIVVLALFLMWRIKHKNPDAIWLWGMSVV
CELWFALSWLLDQLPKLCPINRATDLNVLKEKFETPTPSNPTGKSDLPGLDMFVSTADPEKEPPLVTS
NTILSILAADYPVEKLACYVSDDGGALLTFEAMAEAASFANMWVPFCRKHNIEPRNPDSYFSLKRDPY
KNKVKADFVKDRRRVKREYDEFKVRINSLPDSIRRRSDAYHAREEIKAMKLQRQNRDEEIVEPVKIPK
ATWMADGTHWPGTWINSGPDHSRSDHAGIIQVMLKPPSDEPLHGVSEGFLDLTDVDIRLPLLVYVSR
EKRPGYDHNKKAGAMNALVRASAIMSNGPFILNLDCDHYIYNSQALREGMCFMMDRGGDRLCYVQF
PQRFEGIDPSDRYANHNTVFFDVNMRALDGLMGPVYVGTGCLFRRIALYGFDPPRAKEHHPGFCSC
CFSRKKKKSRVPEENRSLRMGGDSDDDEEMNLSLVPKKFGNSTFLIDSIPVAEFQGRPLADHPAVQN
GRPPGALTIPRELLDASTVAEAIAVISCWYEDKTEWGSRIGWIYGSVTEDVVTGYRMHNRGWKSVYC
VTKRDAFRGTAPINLTDRLHQVLRWATGSVEIFFSRNNAFFASPRMKILQRIAYLNVGIYPFTSFFLIVY
CFLPALSLFSGQFIVQTLNVTFLVYLLIISITLCLLALLEIKWSGISLEEWWRNEQFWLIGGTSAHLAAVIQ
GLLKVVAGIEISFTLTSKSGGEDVDDEFADLYIVKWTSLMIPPITIMMVNLIAIAVGFSRTIYSVIPQWSKL
IGGVFFSFWVLAHLYPFAKGLMGRRGRTPTIVYVWSGLVAITISLLWVAINPPAGSTQIGGSFTFP
>OS10G42750_OsCslD1
MASKGILKNGGKPPTAPSSAAPTVVFGRRTDSGRFISYSRDDLDSEISSVDFQDYHVHIPMTPDNQP
MDPAAGDEQQYVSSSLFTGGFNSVTRAHVMEKQASSARATVSACMVQGCGSKIMRNGRGADILPC
ECDFKICVDCFTDAVKGGGGVCPGCKEPYKHAEWEEVVSASNHDAINRALSLPHGHGHGPKMERRL
SLVKQNGGAPGEFDHNRWLFETKGTYGYGNAIWPEDDGVAGHPKELMSKPWRPLTRKLRIQAAVIS
PYRLLVLIRLVALGLFLMWRIKHQNEDAIWLWGMSIVCELWFALSWVLDQLPKLCPINRATDLSVLKDK
FETPTPSNPTGKSDLPGIDIFVSTADPEKEPVLVTANTILSILAADYPVDKLACYVSDDGGALLTFEAMA
EAASFANLWVPFCRKHEIEPRNPDSYFNLKRDPFKNKVKGDFVKDRRRVKREYDEFKVRVNGLPDAI
RRRSDAYHAREEIQAMNLQREKMKAGGDEQQLEPIKIPKATWMADGTHWPGTWLQASPEHARGDH
AGIIQVMLKPPSPSPSSSGGDMEKRVDLSGVDTRLPMLVYVSREKRPGYDHNKKAGAMNALVRASAI
MSNGPFILNLDCDHYVYNSKAFREGMCFMMDRGGDRLCYVQFPQRFEGIDPSDRYANHNTVFFDV
NMRALDGLQGPVYVGTGCLFRRIALYGFDPPRSKDHTTPWSCCLPRRRRTRSQPQPQEEEEETMAL
RMDMDGAMNMASFPKKFGNSSFLIDSIPVAEFQGRPLADHPSVKNGRPPGALTIPRETLDASIVAEAI
SVVSCWYEEKTEWGTRVGWIYGSVTEDVVTGYRMHNRGWKSVYCVTHRDAFRGTAPINLTDRLHQ
VLRWATGSVEIFFSRNNALFASSKMKVLQRIAYLNVGIYPFTSVFLIVYCFLPALSLFSGQFIVQTLNVT
FLTYLLIITITLCLLAMLEIKWSGIALEEWWRNEQFWLIGGTSAHLAAVLQGLLKVIAGIEISFTLTSKQLG
DDVDDEFAELYAVKWTSLMIPPLTIIMINLVAIAVGFSRTIYSTIPQWSKLLGGVFFSFWVLAHLYPFAK
GLMGRRGRTPTIVYVWSGLVAITISLLWIAIKPPSAQANSQLGGSFSFP
>CSLD_2BS_TGACv1_148683_AA0494520.1_Triticum_aestivum
MSKAPRNPGGGSAGAPKSSSGQPVKFARRTPSGRYLSLSREDIDMEGEMGPDYANYTVHIPPTPDN
QPMKDGSEPTAVAMKAEEQYVSNSLFTGGFNSVTRAHLMDRVIDSDVKHPQMAGAKATRCAMPAC
DGKVMRNERGEEVDPCECRFKICRDCYLDAQKDGCLCPGCKEHYKIGDYADDDPHDVSAGKSLLAR
NQNGEFDHNRWLFESSGTYGYGNAFMPKGGMYEDDLDEDGAGGDGGMPADLSQKPFKPLTRKIP
MPTSIISPYRIFIVIRFFVLIFYLTWRIRNPNMEALWLWGMSIVCELWFAFSWLLDMLPKVNPINRSTDL
AVLKEKFETHSPSNPHGRSDLPGLDVFVSTADPEKEPVLTTANTILSILAVDYPVEKLACYVSDDGGAL
LTFEAMAEAASFANIWVPFCKKHDIEPRNPDSYFALKGDPTKGKRRSDFVKDRRKVKREYDEFKVRI
NGLPDSIRRRSDAFNAREDMKMLKHLRETGADPSEQPKVKKATWMADGTHWPGTWAVSSPDHAK
GNHAGILQVMLRPPSPDPLYGMHDEDQLVDYSDVDTRLPMLVYMSREKRPGYDHNKKAGAMNALV
RCSAVMSNAPFILNFDCDHYINNTQAIREAMCFMMDRGGERICYIQFPQRFEGIDPSDRYANHNTVFF
DGNMRALDGLQGPMYVGTGCMFRRFALYGFDPPRTAEYTGWLFKKKKVTNFKDPESDTQQLKAED
FDAELTAQLVPRRFGNSSAMLASIPIAEFQARPIADHPAVLHGRPPGTLTVPRPPLDPPTVAEAVSVIS
CWYEDKTEWGDRVGWIYGSVTEDVVTGYRMHNRGWRSVYWISKRDAFLGTAPINMTDRLHQVLR
WATGSVEIFFSRNNAFLASRKLMFLQRVAYLNVGIYPFTSIFLLTYCFIPALSLFSGFFIVQTLNVAFLFY
LLTITVTLIALGILEVKWSGIELEDWWRNEQFWLISGISAHLYAVVQGLLKVMAGIEISFTLTAKAAAEDN
EDIYADLYVVKWSSLLIPPITIGMLNIIAIAFAFARTIYSDNPRWGKFIGGGFFSFWVLAHLNPFAKGLMG
RRGKTPTIIFVWSGLISITISLLWVALSPPEANSTGGARGGGFQFP
>CSLD_2DS_TGACv1_177279_AA0572180.1_Triticum_aestivum
MSKAPRNPGGGSAGAPKSSSGQPVKFARRTPSGRYLSLSREDIDMEGEMGPDYANYTVHIPPTPDN
QPMKDGAERTAVAMKAEEQYVSNSLFTGGFNSVTRAHLMDRVIDSDVKHPQMAGARPARCAMPAC
DGKVMRNERGEEIEPCECRFKICRDCYLDAQKDGCLCPGCKEHYKIGDYADDDTHDVSAGKSLLAR
NQNGEFDHNRWLFESSGTYGYGNAFMPKGGMYEDDLDEDGAAGDDGMQDMNQKPFKPLTRKIPM
PASIISPYRIFIVIRFFVLIFYLTWRIRNPNMEALWLWGMSIVCELWFAFSWLLDMLPKVNPINRSTDLAV
LKEKFETPSPSNPHGRSDLPGLDVFVSTADPEKEPVLTTANTILSILAVDYPVEKLACYVSDDGGALLT
FEAMAEAASFANIWVPFCKKHDIEPRNPDSYFALKGDPTKGKRRSDFVKDRRKVKREYDEFKVRING
LPDSIRRRSDAFNAREDMKMLKHLRETGADPSEQPKVKKATWMADGTHWPGTWAVSSPDHAKGN
HAGILQVMLRPPSPDPLYGMHDEDQLIDYSDVDTRLPMLVYMSREKRPGYDHNKKAGAMNALVRCS
AVMSNAPFILNFDCDHYINNNQAVREAMCFMMDRGGERICYIQFPQRFEGIDPSDRYANHNTVFFDG
NMRALDGLQGPMYVGTGCMFRRFALYGFDPPRTAEYTGWLFKKKKVTNFKDPESDTQKLKAEDFD

AELTAQLVPRRFGNSSAMLASIPIAEFQARPIADHPAVLHGRPPGTLTVPRPPLDPPTVAEAVSVISCW
YEDKTEWGDRVGWIYGSVTEDVVTGYRMHNRGWRSVYWISKRDAFLGTAPINMTDRLHQVLRWAT
GSVEIFFSRNNAFLASRKLMFLQRVAYLNVGIYPFTSIFLLTYCFIPALSLFSGFFIVQTLNVAFLFYLLTI
TVTLIALGILEVKWSGIELEDWWRNEQFWLISGISAHLYAVVQGLLKVMAGIEISFTLTAKAAAEDNEDI
YADLYVVKWSSLLIPPITIGMLNIIAIAFAFARTIYSDNPRWGKFIGGGFFSFWVLAHLNPFAKGLMGRR
GKTPTIIFVWSGLISITISLLWVALSPPEANSTGGARGGGFQFP
>CSLD_2AS_TGACv1_114244_AA0365360.1_Triticum_aestivum
MSKAPRNPGGGSAGAPKSSSGQPVKFARRTPSGRYLSLSREDIDMEGEMGPDYANYTVHIPPTPDN
QPMKDGAEPTAVAMKAEEQYVSNSLFTGGFNSVTRAHLMDRVIDSDVKHPQMAGAKATRCAMPAC
DGKVMRNERGEEIDPCECRFKICRDCYLDAQKDGCLCPGCKEHYKIGDYADDDPHDVSSGKSLLAR
NQNGEFDHNRWLFESSGTYGYGNAFMPKGGMYEDDLDEDGVGGDGGMQDMNQKPFKPLTRKIPM
PTSIISPYRIFIVIRFFVLIFYLTWRIRNPNMEALWLWGMSIVCELWFAFSWLLDMLPKVNPINRSTDLAV
LKEKFETPSPSNPHGRSDLPGLDIFVSTADPEKEPVLTTANTILSILAVDYPVEKLACYVSDDGGALLTF
EAMAEAASFANIWVPFCKKHDIEPRNPDSYFALKGDPTKGKRRSDFVKDRRKVKREYDEFKVRINGL
PDSIRRRSDAFNAREDMKMLKHLRETGADPSEQPKVKKATWMADGTHWPGTWAVSSPDHAKGNH
AGILQVMLRPPSPDPLYGMHDEDQLIDYSDVDTRLPMLVYMSREKRPGYDHNKKAGAMNALVRCSA
VMSNAPFILNFDCDHYINNNQAVREAMCFMMDRGGERICFIQFPQRFEGIDPSDRYANHNTVFFDGN
MRALDGLQGPMYVGTGCMFRRFALYGFDPPRTAEYTGWLFKKKKVTNFKDPDSDTQQLKAEDFDA
ELTAQLVPRRFGNSSAMLASIPIAEFQARPIADHPAVLHGRPPGTLTVPRPPLDPPTVAEAVSVISCWY
EDKTEWGDRVGWIYGSVTEDVVTGYRMHNRGWRSVYWISKRDAFLGTAPINMTDRLHQVLRWATG
SVEIFFSRNNAFLASRKLMFLQRVAYLNVGIYPFTSIFLLTYCFIPALSLFSGFFIVQTLNVAFLFYLLTITI
TLIALGILEVKWSGIELEDWWRNEQFWLISGISAHLYAVVQGLLKVMAGIEISFTLTAKAAADDNEDIYA
DLYVVKWSSLLIPPITIGMLNIIAIAFAFARTIYSENPRWGKFIGGGFFSFWVLAHLNPFAKGLMGRRGK
TPTIIFVWSGLISITISLLWVALSPPEANSTGGARSGGFQFP
>OS08G25710_OsCslD3
MSTGPGKKAIRNAGGVGGGAGPSAGGPRGPAGQAVKFARRTSSGRYVSLSREDIDMEGELAADYT
NYTVQIPPTPDNQPMLNGAEPASVAMKAEEQYVSNSLFTGGFNSATRAHLMDKVIESSVSHPQMAG
AKGSRCAMPACDGSAMRNERGEDVDPCECHFKICRDCYLDAQKDGCICPGCKEHYKIGEYADDDP
HDGKLHLPGPGGGGNKSLLARNQNGEFDHNRWLFESSGTYGYGNAFWPKGGMYDDDLDDDVDKL
GGDGGGGGGGGPLPEQKPFKPLTRKIPMPTSVISPYRIFIVIRMFVLLFYLTWRIRNPNMEALWLWGM
SIVCELWFAFSWLLDMLPKVNPVNRSTDLAVLKEKFETPSPSNPHGRSDLPGLDVFVSTADPEKEPV
LTTATTILSILAVDYPVEKLACYVSDDGGALLTFEAMAEAASFANVWVPFCKKHDIEPRNPDSYFSVKG
DPTKGKRRNDFVKDRRRVKREFDEFKVRINGLPDSIRRRSDAFNAREDMKMLKHLRETGADPSEQP
KVKKATWMADGSHWPGTWAASAPDHAKGNHAGILQVMLKPPSPDPLYGMHDDDQMIDFSDVDIRL
PMLVYMSREKRPGYDHNKKAGAMNALVRCSAVMSNGPFMLNFDCDHYINNAQAVREAMCFFMDR
GGERIAYIQFPQRFEGIDPSDRYANNNTVFFDGNMRALDGLQGPMYVGTGCMFRRFAVYGFDPPRT
AEYTGWLFTKKKVTTFKDPESDTQTLKAEDFDAELTSHLVPRRFGNSSPFMASIPVAEFQARPLADH
PAVLHGRPSGALTVPRPPLDPPTVAEAVSVISCWYEDKTEWGDRVGWIYGSVTEDVVTGYRMHNRG
WRSVYCITKRDAFLGTAPINLTDRLHQVLRWATGSVEIFFSRNNAFLASRKLMLLQRISYLNVGIYPFT
SIFLLVYCFIPALSLFSGFFIVQKLDIAFLCYLLTMTITLVALGILEGLLKVMAGIEISFTLTAKAAADDNEDI
YADLYIVKWSSLLIPPITIGMVNIIAIAFAFARTIYSDNPRWGKFIGGGFFSFWVLAHLNPFAKGLMGRR
GKTPTIVFVWSGLLSITVSLLWVAISPPEANSNGGARGGGFQFP
>AT4G38190_CslD4
MASTPPQTSKKVRNNSGSGQTVKFARRTSSGRYVSLSRDNIELSGELSGDYSNYTVHIPPTPDNQP
MATKAEEQYVSNSLFTGGFNSVTRAHLMDKVIDSDVTHPQMAGAKGSSCAMPACDGNVMKDERGK
DVMPCECRFKICRDCFMDAQKETGLCPGCKEQYKIGDLDDDTPDYSSGALPLPAPGKDQRGNNNN
MSMMKRNQNGEFDHNRWLFETQGTYGYGNAYWPQDEMYGDDMDEGMRGGMVETADKPWRPLS
RRIPIPAAIISPYRLLIVIRFVVLCFFLTWRIRNPNEDAIWLWLMSIICELWFGFSWILDQIPKLCPINRSTD
LEVLRDKFDMPSPSNPTGRSDLPGIDLFVSTADPEKEPPLVTANTILSILAVDYPVEKVSCYLSDDGGA
LLSFEAMAEAASFADLWVPFCRKHNIEPRNPDSYFSLKIDPTKNKSRIDFVKDRRKIKREYDEFKVRIN
GLPDSIRRRSDAFNAREEMKALKQMRESGGDPTEPVKVPKATWMADGTHWPGTWAASTREHSKG
DHAGILQVMLKPPSSDPLIGNSDDKVIDFSDTDTRLPMFVYVSREKRPGYDHNKKAGAMNALVRASAI
LSNGPFILNLDCDHYIYNCKAVREGMCFMMDRGGEDICYIQFPQRFEGIDPSDRYANNNTVFFDGNM
RALDGVQGPVYVGTGTMFRRFALYGFDPPNPDKLLEKKESETEALTTSDFDPDLDVTQLPKRFGNST
LLAESIPIAEFQGRPLADHPAVKYGRPPGALRVPRDPLDATTVAESVSVISCWYEDKTEWGDRVGWI
YGSVTEDVVTGYRMHNRGWRSVYCITKRDSFRGSAPINLTDRLHQVLRWATGSVEIFFSRNNAILAS
KRLKFLQRLAYLNVGIYPFTSLFLILYCFLPAFSLFSGQFIVRTLSISFLVYLLMITICLIGLAVLEVKWSGI
GLEEWWRNEQWWLISGTSSHLYAVVQGVLKVIAGIEISFTLTTKSGGDDNEDIYADLYIVKWSSLMIPP
IVIAMVNIIAIVVAFIRTIYQAVPQWSKLIGGAFFSFWVLAHLYPFAKGLMGRRGKTPTIVFVWAGLIAITI
SLLWTAINPNTGPAAAAEGVGGGGFQFP
>OS07G14850_OsCesA

MEASAGLVAGSHNRNELVVIRRDGGGGGGVGGRRAAEAKAACQICGDDVGEGPDGEPFVACNECA
FPVCRNCYDYERREGSQACPQCKTRFKRLKGCPRVAGDEEEDGVDDLEGEFGLDGREDDPQYIAE
SMLRANMSYGRGGDLQPFQPIPNVPLLTNGQMVDDIPPEQHALVPSYMGGGGGGGKRIHPLPFADP
SVPVQPRSMDPSKDLAAYGYGSVAWKERMEGWKQKQERMQQLRSEGGGDWDGDGDADLPLMD
EARQPLSRKVPISSSRINPYRMIIIIRLVVLGFFFHYRVMHPVNDAFALWLISVICEIWFAMSWILDQFPK
WLPIERETYLDRLSLRFDKEGQPSQLAPVDFFVSTVDPSKEPPLVTANTVLSILSVDYPVEKVSCYVS
DDGAAMLTFEALSETSEFAKKWVPFCKKFNIEPRAPEWYFQQKIDYLKDKVAASFVRERRAMKRDYE
EFKVRINALVAKAQKVPEEGWTMQDGSPWPGNNVRDHPGMIQVFLGQSGGRDVEGNELPRLVYVS
REKRPGYNHHKKAGAMNALVRVSAVLSNAPYLLNLDCDHYINNSKAIREAMCFMMDPLVGKKVCYV
QFPQRFDGIDRHDRYANRNVVFFDINMKGLDGIQGPIYVGTGCVFRRQALYGYDAPKTKKPPSRTCN
CWPKWCCCCCCGNRHTKKKTTKPKPEKKKRLFFKKAENQSPAYALGEIEEGAPGAETDKAGIVNQQ
KLEKKFGQSSVFVASTLLENGGTLKSASPASLLKEAIHVISCGYEDKTDWGKEIGWIYGSITEDILTGFK
MHCHGWRSIYCIPKRPAFKGSAPLNLSDRLHQVLRWALGSVEIFFSKHCPLWYGYGGGLKFLERFSY
INSIVYPWTSIPLLAYCTLPAICLLTGKFITPELTNVASLWFMSLFICIFVTGILEMRWSGVAIDDWWRNE
QFWVIGGVSSHLFAVFQGLLKVLAGVDTSFTVTSKAGDDEEFSELYTFKWTTLLIPPTTLLLLNFIGVV
AGVSNAINNGYESWGPLFGKLFFAFWVIVHLYPFLKGLVGRQNRTPTIVIVWSILLASIFSLLWVRIDPF
LAKNNGPLLEECGLDCN
>OS03G62090_OsCesA
MEASAGLVAGSHNRNELVVIRRDGEPGPKPVKHTNGQVCQICGDDVGLTPDGEPFVACNECAFPVC
RDCYEYERREGTQNCPQCKTRFKRLKGCARVPGDEEEEDVDDLENEFNWRDKTDSQYVAESMLH
GHMSYGRGGDLDGVPQHFQPIPNVPLLTNGEMADDIPPEQHALVPSFMGGGGKRIHPLPYADPNLP
VQPRSMDPSKDLAAYGYGSVAWKERMESWKQKQERLHQMRNDGGGKDWDGDGDDADLPLMDE
ARQPLSRKIPISSSLVNPYRMIIIIRLVVLGFFFHYRVMHPVPDAFALWLISVICEIWFAMSWILDQFPKW
FPIERETYLDRLTLRFDKEGQQSQLAPVDFFVSTVDPMKEPPLVTANTVLSILAVDYPVDKVSCYVSD
DGAAMLTFEALSETSEFAKKWVPFCKRYSLEPRAPEWYFQQKIDYLKDKVAPNFVRERRAMKREYE
EFKVRINALVAKAQKVPEEGWTMQDGTPWPGNNVRDHPGMIQVFLGQSGGHDVEGNELPRLVYVS
REKRPGYNHHKKAGAMNALVRVSAVLTNAPYMLNLDCDHYINNSKAIKEAMCFMMDPLVGKKVCYV
QFPQRFDGIDRHDRYANRNVVFFDINMKGLDGIQGPIYVGTGCVFRRQALYGYDAPKSKKPPSRTCN
CWPKWCICCCCFGNRTNKKKTAKPKTEKKKRLFFKRAENQSPAYALGEIDEGAPGAENEKAGIVNQ
QKLEKKFGQSSVFVASTLLENGGTLKSASPASLLKEAIHVISCGYEDKTDWGKEIGWIYGSVTEDILTG
FKMHCHGWRSIYCIPKRAAFKGSAPLNLSDRLHQVLRWALGSIEIFFSNHCPLWYGYGGGLKCLERF
SYINSIVYPWTSIPLLAYCTLPAICLLTGKFITPELTNIASLWFMSLFICIFATGILEMRWSGVGIDDWWR
NEQFWVIGGVSSHLFAVFQGLLKVIAGIDTSFTVTSKGGDDEEFSELYTFKWTTLLIPPTTLLLLNFIGV
VAGVSNAINNGYESWGPLFGKLFFAFWVIVHLYPFLKGLVGRQNRTPTIVIVWSILLASIFSLLWVRIDP
FLAKNDGPLLEECGLDCN
>OS07G10770_OsCesA
MDGDADAVKSGRHGSGQACQICGDGVGTTAEGDVFAACDVCGFPVCRPCYEYERKDGTQACPQC
KTKYKRHKGSPAIRGEEGEDTDADDVSDYNYPASGSADQKQKIADRMRSWRMNAGGGGDVGRPK
YDSGEIGLTKYDSGEIPRGYIPSVTNSQISGEIPGASPDHHMMSPTGNIGKRAPFPYVNHSPNPSREF
SGSIGNVAWKERVDGWKLKQDKGAIPMTNGTSIAPSEGRGVGDIDASTDYNMEDALLNDETRQPLS
RKVPLPSSRINPYRMVIVLRLVVLSIFLHYRITNPVRNAYPLWLLSVICEIWFALSWILDQFPKWFPINRE
TYLDRLALRYDREGEPSQLAAVDIFVSTVDPMKEPPLVTANTVLSILAVDYPVDKVSCYVSDDGAAML
TFDALAETSEFARKWVPFVKKYNIEPRAPEWYFSQKIDYLKDKVHPSFVKDRRAMKREYEEFKVRIN
GLVAKAQKVPEEGWIMQDGTPWPGNNTRDHPGMIQVFLGHSGGLDTEGNELPRLVYVSREKRPGF
QHHKKAGAMNALVRVSAVLTNGQYMLNLDCDHYINNSKALREAMCFLMDPNLGRSVCYVQFPQRF
DGIDRNDRYANRNTVFFDINLRGLDGIQGPVYVGTGCVFNRTALYGYEPPIKQKKKGSFLSSLCGGR
KKASKSKKKSSDKKKSNKHVDSAVPVFNLEDIEEGVEGAGFDDEKSLLMSQMSLEKRFGQSAAFVAS
TLMEYGGVPQSATPESLLKEAIHVISCGYEDKTEWGTEIGWIYGSVTEDILTGFKMHARGWRSIYCMP
KRPAFKGSAPINLSDRLNQVLRWALGSVEILFSRHCPIWYGYGGRLKFLERFAYINTTIYPLTSIPLLIYC
VLPAICLLTGKFIIPEISNFASIWFISLFISIFATGILEMRWSGVGIDEWWRNEQFWVIGGISAHLFAVFQG
LLKVLAGIDTNFTVTSKASDEDGDFAELYMFKWTTLLIPPTTILIINLVGVVAGISYAINSGYQSWGPLFG
KLFFAFWVIVHLYPFLKGLMGRQNRTPTIVVVWAILLASIFSLLWVRIDPFTTRVTGPDTQTCGINC
>OS05G08370_OsCesA
MAANAGMVAGSRNRNEFVMIRPDGDAPPPAKPGKSVNGQVCQICGDTVGVSATGDVFVACNECAF
PVCRPCYEYERKEGNQCCPQCKTRYKRHKGSPRVQGDEEEEDVDDLDNEFNYKHGNGKGPEWQI
QRQGEDVDLSSSSRHEQHRIPRLTSGQQISGEIPDASPDRHSIRSGTSSYVDPSVPVPVRIVDPSKDL
NSYGINSVDWQERVASWRNKQDKNMMQVANKYPEARGGDMEGTGSNGEDMQMVDDARLPLSRIV
PIPSNQLNLYRIVIILRLIILMFFFQYRVTHPVRDAYGLWLVSVICEIWFALSWLLDQFPKWYPINRETYL
DRLALRYDREGEPSQLAPIDVFVSTVDPLKEPPLITANTVLSILAVDYPVDKVSCYVSDDGSAMLTFEA
LSETAEFARKWVPFCKKHNIEPRAPEFYFAQKIDYLKDKIQPSFVKERRAMKREYEEFKVRINALVAKA
QKVPEEGWTMADGTAWPGNNPRDHPGMIQVFLGHSGGLDTDGNELPRLVYVSREKRPGFQHHKK
AGAMNALIRVSAVLTNGAYLLNVDCDHYFNSSKALREAMCFMMDPALGRKTCYVQFPQRFDGIDLH

DRYANRNIVFFDINMKGLDGIQGPVYVGTGCCFNRQALYGYDPVLTEADLEPNIVVKSCCGGRKKKS
KSYMDSKNRMMKRTESSAPIFNMEDIEEGIEGYEDERSVLMSQKRLEKRFGQSPIFIASTFMTQGGIP
PSTNPASLLKEAIHVISCGYEDKTEWGKEIGWIYGSVTEDILTGFKMHARGWISIYCMPPRPCFKGSAP
INLSDRLNQVLRWALGSVEILLSRHCPIWYGYNGRLKLLERLAYINTIVYPITSIPLIAYCVLPAICLLTNK
FIIPEISNYAGMFFILLFASIFATGILELRWSGVGIEDWWRNEQFWVIGGTSAHLFAVFQGLLKVLAGIDT
NFTVTSKASDEDGDFAELYVFKWTSLLIPPTTVLVINLVGMVAGISYAINSGYQSWGPLFGKLFFSIWVI
LHLYPFLKGLMGRQNRTPTIVIVWSILLASIFSLLWVKIDPFISPTQKAVALGQCGVNC
>OS03G59340_OsCesA
MDGAKSGKQCHVCQICGDGVGTAADGELFTACDVCGFPVCRPCYEYERKDGSQACPQCKTKYKRH
KGSPPILGDESDDVDADDASDVNYPTSGNQDHKHKIAERMLTWRMNSGRNDDIVHSKYDSGEIGHP
KYDSGEIPRIYIPSLTHSQISGEIPGASPDHMMSPVGNIGRRGHPFPYVNHSPNPSREFSGSLGNVAW
KERVDGWKMKDKGAIPMANGTSIAPSEGRGVGDIDASTDYNMEDALLNDETRQPLSRKVPISSSRIN
PYRMVIVLRLIVLCIFLHYRITNPVRNAYPLWLLSVICEIWFALSWILDQFPKWSPINRETYLDRLALRYD
REGEPSQLAPVDIFVSTVDPMKEPPLVTANTVLSILAVDYPVDKVSCYVSDDGAAMLTFDALAETSEF
ARKWVPFCKKYSIEPRAPEWYFAQKIDYLKDKVQASFVKDRRAMKREYEEFKVRVNALVAKAQKVPE
EGWIMQDGTPWPGNNTRDHPGMIQVFLGHSGGLDTEGNELPRLVYVSREKRPGFQHHKKAGAMN
ALVRVSAVLTNGQYLLNLDCDHYINNSKALREAMCFLMDPNLGRRVCYVQFPQRFDGIDRNDRYAN
RNTVFFDINLRGLDGLQGPVYVGTGCVFNRTALYGYEPPIKQKRPGYFSSLCGGRKKTKKSKEKSTE
KKKSHKHVDSSVPVFNLEDIEEGIEGSGFDDEKSLLMSQMSLEKRFGQSSVFVASTLMEYGGVPQSA
TPESLLKEAIHVISCGYEDKSDWGTEIGWIYGSVTEDILTGFKMHARGWRSIYCMPKRPAFKGSAPINL
SDRLNQVLRWALGSVEILFSRHCPIWYGYGGRLKFLERFAYINTTIYPLTSIPLLLYCILPAICLLTGKFII
PEISNFASIWFISLFLSIFATGILEMRWSGVGIDEWWRNEQFWVIGGISAHLFAVFQGLLKVLAGIDTSF
TVTSKASDEEGDFAELYMFKWTTLLIPPTTILIINLVGVVAGISYAINSGYQSWGPLFGKLFFAFWVIVHL
YPFLKGLMGRQNRTPTIVVVWAILLASIFSLLWVRIDPFTTRVTGPDTQKCGINC
>OS10G32980_OsCesA
MDTASVTGGEHKGKEKTCRVCGEEVAAREDGKPFVACAECGFPVCKPCYEYERSEGTQCCPQCNT
RYKRHKGCPRVEGDEDDGGDMDDFEEEFQIKSPTKQKPPHEPVNFDVYSENGEQPAQKWRPGGP
ALSSFTGSVAGKDLEQEREMEGGMEWKDRIDKWKTKQEKRGKLNRDDSDDDDDKNDDEYMLLAEA
RQPLWRKVPIPSSKINPYRIVIVLRLVVLCFFLKFRITTPAMDAVPLWLASVICELWFALSWILDQLPKW
SPVTRETYLDRLALRYERDGEPCRLAPIDFFVSTVDPLKEPPIITANTVLSILAVDYPVDRVSCYVSDDG
ASMLLFDTLSETAEFARRWVPFCKKFTIEPRAPEFYFSQKIDYLKDKVQPTFVKERRAMKREYEEFKV
RINALVAKAQKKPEEGWVMQDGTPWPGNNTRDHPGMIQVYLGSQGALDVEGSELPRLVYVSREKR
PGYNHHKKAGAMNSLVRVSAVLTNAPFILNLDCDHYVNNSKAVREAMCFLMDKQLGKKLCYVQFPQ
RFDGIDRHDRYANRNTVFFDINMKGLDGIQGPVYVGTGTVFNRQALYGYDPPRPEKRPKMTCDCWP
SWCCCCCCFGGGKRGKSHKNKKGGGGGEGGGLDEPRRGLLGFYKKRSKKDKLGGGAASLAGGK
KGYRKHQRGFELEEIEEGLEGYDELERSSLMSQKSFEKRFGQSPVFIASTLVEDGGLPQGAAADPAA
LIKEAIHVISCGYEEKTEWGKEIGWIYGSVTEDILTGFKMHCRGWKSVYCTPARAAFKGSAPINLSDRL
HQVLRWALGSVEIFMSRHCPLWYAYGGRLKWLERFAYTNTIVYPFTSIPLLAYCTIPAVCLLTGKFIIPT
LNNLASIWFIALFLSIIATGVLELRWSGVSIEDWWRNEQFWVIGGVSAHLFAVFQGLLKVLGGVDTNFT
VTSKAAADETDAFGELYLFKWTTLLVPPTTLIIINMVGIVAGVSDAVNNGYGSWGPLFGKLFFSFWVIL
HLYPFLKGLMGRQNRTPTIVVLWSILLASIFSLVWVRIDPFIPKPKGPVLKPCGVSC
>OS09G25490_OsCesA
MEASAGLVAGSHNRNELVLIRGHEEPKPLRALSGQVCEICGDEVGRTVDGDLFVACNECGFPVCRP
CYEYERREGTQNCPQCKTRYKRLKGSPRVPGDEDEEDIDDLEHEFNIDDEKQKQLQQDQDGMQNS
HITEAMLHGKMSYGRGPDDGDGNSTPLPPIITGARSVPVSGEFPISNSHGHGEFSSSLHKRIHPYPVS
EPGSAKWDEKKEVSWKERMDDWKSKQGIVAGGAPDPDDYDADVPLNDEARQPLSRKVSIASSKVN
PYRMVIILRLVVLGFFLRYRILHPVPDAIPLWLTSIICEIWFAVSWILDQFPKWYPIDRETYLDRLSLRYE
REGEPSLLSAVDLFVSTVDPLKEPPLVTANTVLSILAVDYPVDKVSCYVSDDGASMLTFESLSETAEFA
RKWVPFCKKFSIEPRAPEFYFSQKVDYLKDKVHPNFVQERRAMKREYEEFKVRINALVAKAQKVPAE
GWIMKDGTPWPGNNTRDHPGMIQVFLGHSGGHDTEGNELPRLVYVSREKRPGFQHHKKAGAMNAL
IRVSAVLTNAPFMLNLDCDHYINNSKAIREAMCFLMDPQVGRKVCYVQFPQRFDGIDVHDRYANRNT
VFFDINMKGLDGIQGPVYVGTGCVFRRQALYGYNPPKGPKRPKMVTCDCCPCFGRKKRKHGKDGL
PEAVAADGGMDSDKEMLMSQMNFEKRFGQSAAFVTSTLMEEGGVPPSSSPAALLKEAIHVISCGYE
DKTDWGLELGWIYGSITEDILTGFKMHCRGWRSVYCMPKRAAFKGSAPINLSDRLNQVLRWALGSV
EIFFSRHSPLLYGYKNGNLKWLERFSYINTTIYPFTSLPLLAYCTLPAVCLLTGKFIMPPISTFASLFFIAL
FISIFATGILEMRWSGVSIEEWWRNEQFWVIGGVSAHLFAVVQGLLKVLAGIDTNFTVTSKATGDEDD
EFAELYAFKWTTLLIPPTTLLILNIIGVVAGVSDAINNGSEAWGPLFGKLFFAFWVIVHLYPFLKGLMGR
QNRTPTIVVIWSVLLASIFSLLWVRIDPFTIKARGPDVRQCGINC
>AT2G33100_CslD1
MASSPPKKTLNSQSSSLSRPPQAVKFGRRTSSGRIVSLSRDDDMDVSGDYSGQNDYINYTVLMPPT
PDNQPAGSSGSTSESKGDANRGGGGGDGPKMGNKLERRLSVMKSNNKSMLLRSQTGDFDHNRWL
FESKGKYGIGNAFWSEEDDTYDGGVSKSDFLDKPWKPLTRKVQIPAKILSPYRLLIVIRLVIVFFFLWW

RITNPNEDAMWLWGLSIVCEIWFAFSWILDILPKLNPINRATDLAALHDKFEQPSPSNPTGRSDLPGVD
VFVSTADPEKEPPLVTANTLLSILAVDYPIEKLSAYISDDGGAILTFEAMAEAVRFAEYWVPFCRKHDIE
PRNPDSYFSIKKDPTKNKKRQDFVKDRRWIKREYDEFKVRINGLPEQIKKRAEQFNMREELKEKRIAR
EKNGGVLPPDGVEVVKATWMADGTHWPGTWFEPKPDHSKGDHAGILQIMSKVPDLEPVMGGPNEG
ALDFTGIDIRVPMFAYVSREKRPGFDHNKKAGAMNGMVRASAILSNGAFILNLDCDHYIYNSKAIKEG
MCFMMDRGGDRICYIQFPQRFEGIDPSDRYANHNTVFFDGNMRALDGLQGPVYVGTGCMFRRYAL
YGFNPPRANEYSGVFGQEKAPAMHVRTQSQASQTSQASDLESDTQPLNDDPDLGLPKKFGNSTMF
TDTIPVAEYQGRPLADHMSVKNGRPPGALLLPRPPLDAPTVAEAIAVISCWYEDNTEWGDRIGWIYGS
VTEDVVTGYRMHNRGWRSVYCITKRDAFRGTAPINLTDRLHQVLRWATGSVEIFFSKNNAMFATRRL
KFLQRVAYLNVGIYPFTSIFLVVYCFLPALCLFSGKFIVQSLDIHFLSYLLCITVTLTLISLLEVKWSGIGLE
EWWRNEQFWLIGGTSAHLAAVVQGLLKVIAGIEISFTLTSKASGEDEDDIFADLYIVKWTGLFIMPLTIII
VNLVAIVIGASRTIYSVIPQWGKLMGGIFFSLWVLTHMYPFAKGLMGRRGKVPTIVYVWSGLVSITVSL
LWITISPPDDVSGSGGISV
>CSLD_5BS_TGACv1_425241_AA1392650.1_Triticum_aestivum
MSRRLSLPAGSPVTVTVSPTKGKGAGGGSPGDGVVRRGSGLTSPVPRHSIGSSTATLQVSPVRRSG
GSRYASRDGADASAEFVHYTVHIPPTPDRTTASASTDVPAAEEEGEVLPQRSYVSGTIFTGGLNCAT
RAHVLSNSADGARPAASANMSCKMRGCDMPAFLNAGRGGHPPCDCGFMICEECYMDCVAAAGNC
PGCKEAYSAGSDTDDSVDEDDDDAISSSEERDQMPMTSMSKRFSMVHSIKMPMPSSNGKPADFDH
ARWLFETKGTYGYGNALWPKNEHGGGGNNAGATSGFVGIEEPPNFGARCRRPLTRKTSVSQAILSP
YRMLIAIRLVALGFFLAWRIRHPNPDAMWLWALSVTCEVWFAFSWLLDSLPKLCPVNRSCDLDVLAD
RFELPTARNPKGRSDLPGIDVFVSTADPEKEPPLVTANTILSILAADYPVEKLACYLSDDGGALLTFEAL
AETASFARTWVPFCRKHGVEPRCPESYFGQKRDFLKNKVRLDFVRERRKVKREYDEFKVRVNSLTE
AIRRRSDAYNAGEELRARRRLQEEAVAAGGALGTAPLAETGAVKATWMSDGSQWPGTWLTGATDH
ARGNHAGIIQAMLAPPTSEPVLGGEPAESGALIDTTGVDIRLPMLVYVSREKRPGYDHNKKAGAMNAL
VRTSAIMSNGPFILNLDCDHYVHNSAALREGMCYMLDRGGDRVCYVQFPQRFEGIDPNDRYANHNL
VFFDVAMRAMDGLQGPMYVGTGCIFRRTALYGFSPPRATEHHGWLGRKKIKLFLRKPTMGKKTDRE
SEHESMLPPIEDDDHNQLGDGVRGDLLLPQQRTVRHPADEAPAARGLLQRGHVPVHLHVPHRLLRA
PGRLPLHRQVHRPAPERHVPRLPAHHHHHAVPAGAAGDQVVRDHAARVVAQRAVLGDRRHQRAP
GCGAAGPPQGDRRRGHLLHAHVQAGRRRRRGGGHVRGAVRGAVELPDGAPRDHHDAERGGAGG
GDGEDAVQRVPAVEQAAGRRLLQLLGAVPPLPL
>GRMZM2G061764_P01
MHEGLGRWGLDSLKYGCHSNYTVRMPPTPDNQPYSVGGGAPLSVCAGGTKPDDLPLPPSKLVNRR
GGACADDGLAGASGKMDRRLSTARVPAPSKSLLVRSQTGDFDHNRWLFETRGTYGIGNAYWPQDS
SAYADDEDGGVGSDPVKMEDLVDKPWKPLSRKVPIPPGILSPYRLLVLVRFISLFLFLIWRATNPNLDA
LWLWGISIVCEFWFAFSWLLDQMPKLNPINRAVDLSALREKFESPTPSNPTGRSDLPGLDVFISTADP
YKEPPLTTANSLLSILGTEYPVEKLFVYISDDGGALLTFEAMAEACEFAKVWVPFCRKHSIEPRNPDAY
FNQKGDPTKGKKRPDFVKDRRWIKREYDEFKVRINGLADLIRRRANAMNARERKIARDKAAAASSDA
PVADASTVKATWMADGTHWPGTWLDSAPDHAKGDHASIVQVMIKNPHYDVVHGDAGSHPYLDFTG
VDVRIPMFVYLSREKRPGYDHNKKAGAMNAMVRASAILSNGPFMLNFDCDHYIFNCMAIREAMCYML
DRGGDRICYIQFPQRFEGIDPSDRYANHNTVFFDGNMRALDGLQGPMYVGTGCLFRRYAIYGFNPP
RTNEYRGIYGQVKVPIDPHGHHAPGAAEELRPLSEHPDHEAPQRFGKSKMFIETIAVAEYQGRPLQD
HPSVQNGRPPGALLMPRPPLDAATVAESVAMISCWYEDGTEWGQRVGWIYGSVTEDVVTGYRMHN
RGWRSVYCITRRDAFRGTAPINLTDRLHQVLRWATGSVEIFFSKNNALLASQRLKFLQRLSYLNVGIY
PFTSLFLIMYCLLPALSLFSGQFIVATLDPTFLCYLLLITITLMLLCLLEVKWSGIGLEEWWRNEQFWVIG
GTSAHLAAVLQGLLKVIAGIEISFTLTAKAAAEDDDDPFAELYLVKWTSLFIPPLAVIGINIIALVVGVSRA
VYAEIPQYSKLLGGGFFSFWVLAHYYPFAKGLMGRRGRTPTLVYVWAGLISITVSLLWITISPPDDRIT
QGGVDV
>CSLD_1DL_TGACv1_063091_AA0223780.1_Triticum_aestivum
MGSKGILKNSGSSRVPPHGPSKPPTAPTSAPQVVFGRRTESGRFISYSRDDLDSEISSVDFQDYHVHI
PMTPDNQPMEEDGTKADEQYVSSSLFTGGFNSVTRAHVMDKQGPDSDMGRSGPKGSICMVEGCD
SKIMRNGRGEDILPCECDFKICVDCFTDAVKGGRGVCPGCKELYKHTEWEEVLSNSSNELTRALSLP
HGPGGKMERRLSLVKQGTMNNQSGEFDHNRWLFETKGTYGYGNAIWPDDNVDDDGRNGVPGHPK
ELMSKPWRPLTRKLQIPAAVISPYRLLVLIRLVALAFFLMWRIKHQNDDAIWLWGMSIVCELWFALSW
VLDQLPKLCPINRATDLSVLKEKFETPTPSNPTGKSDLPGIDIFVSTADPEKEPVLVTANTILSILAVDYP
VDKLACYVSDDGGALLTFEAMAEAASFANFWVPFCRKHDIEPRNPDSYFNLKRDPFKNKVKADFVKD
RRRIKREYDEFKVRVNGLPDSIRRRSDAYHAREEIQAMNLQREKIKAGGDEQFEPVKIPKATWMADS
THWPGTWIHSSQDHARGDHAGIIQVMLKPPSDMPMYGNIEKSPLDFSEVDTRLPMLVYMSREKRPG
YDHNKKAGAMNALVRASAIMSNGPFILNLDCDHYVYNSKAFREGMCFMMDRGGDRLCYVQFPQRF
EGIDPSDRYANHNTVFFDINMRALDGLQGPVYVGTGCLFRRIALYGFDPPRSKDHSPGFCGCCLPRR
RKASASNANPEETMALRMGDFDGDSMNLATFPKKFGNSSFLIDSIPVAEFQGRPLADHPSVKNGRPP
GALTIPREILDASIVAEAISVVSCWYEEKTEWGTRVGWIYGSVTEDVVTGYRMHNRGWKSVYCVTQR
DAFRGTAPINLTDRLHQVLRWATGSVEIFFSRNNALFASSKMKVLQRIAYLNVGIYPFTSIFLIVYCFLP

ALSLFSGQFIVQTLNVTFLTYLLIITVTLCLLAMLEIKWSGIALEEWWRNEQFWLIGGTSAHLAAVMQGL
LKNSTK
>OS06G22980_OsCslD5
MSVDYANYTVLMPPTPDNQPSGGAPPAAPSAGGARPGDLPLPPYGSSSSSRLVNRRGGGDDGAK
MDRRLSTARVPAPSSNKSLLVRSQTGDFDHNRWLFETKGTYGIGNAYWPQDNVYGDDGGGGAVK
MEDLVEKPWKPLSRKVPIPPGILSPYRLLVLVRFVALFLFLVWRVTNPNMDALWLWGISIVCEFWFAF
SWLLDQMPKLNPINRAADLAALKEKFESPSPTNPTGRSDLPGLDVFISTADPYKEPTLVTANTLLSILA
TEYPVEKLFVYISDDGGALLTFESMAEACAFAKVWVPFCRKHSIEPRNPDSYFTQKGDPTKGKKRPD
FVKDRRWIKREYDEFKIRVNSLPDLIRRRANALNARERKLARDKQAAGDADALASVKAATWMADGTH
WPGTWLDPSPDHAKGDHASIVQVMIKNPHHDVVYGEAGDHPYLDMTDVDMRIPMFAYLSREKRAG
YDHNKKAGAMNAMVRASAILSNGPFMLNFDCDHYIYNCQAIREAMCYMLDRGGDRICYIQFPQRFEG
IDPSDRYANHNTVFFDGNMRALDGLQGPMYVGTGCLFRRYAIYGFNPPRAIEYRGTYGQTKVPIDPR
QGSEAMPGAGGGRSGGGSVGGDHELQALSTAHPDHEAPQKFGKSKMFIESIAVAEYQGRPLQDHP
SVLNGRPPGALLMPRPPLDAATVAESVSVISCWYEDNTEWGQRVGWIYGSVTEDVVTGYRMHNRG
WRSVYCITRRDAFRGTAPINLTDRLHQVLRWATGSVEIFFSKNNAVLASRRLKFLQRMAYLNVGIYPF
TSLFLIMYCLLPALSLFSGQFIVATLDPTFLSYLLLITITLMLLCLLEVKWSGIGLEEWWRNEQFWVIGGT
SAHLAAVLQGLLKVVAGIEISFTLTAKAAAEDDDDPFAELYLIKWTSLFIPPLAVIGINIIALVVGVSRTVY
AEIPQYSKLLGGGFFSFWVLAHYYPFAKGLMGRRGRTPTIVYVWAGLISITVSLLWITISPPDDSVAQG
GIDV
>CSLD_7DL_TGACv1_603510_AA1985050.1_Triticum_aestivum
MASDHTNYTVFMPPTPDNQPGAAPTPASGGSTKPENLPLPRYTSGSKLVNRRSGDDGAAGGAKMD
RWLSTEQVASPSKSLLVRSQTGEFDHNRWLFETQGTYGIGNAYWPQDDNDDGAGMGGGSVKMED
LVDKPWKPLSRKVAIPPGILSPYRLLVLVRFVALFLFLVWRATNPNPDAMWLWGISIVCEYWFALSWL
LDQMPKLNPINRAADLAALREKFESKTPSNPTGRSDLPGLDVFISTADPYKEPPLVTANTLLSILATDY
PVEKLFVYISDDGGALLTFEAMAEACAYAKVWVPFCRKHSIEPRNPEAYFTQKGDPTKGKKRPDFVK
DRRWIKREYDEYKVRINDLPEAIKRRAKAMNAHERKIARETAAASSDAAPPPVKATWMADGTHWPGT
WLDSAPDHGKGDHASIIQVMIKNPHHDVVYGDADDHAYLDFTNVDVRIPMFVYLSREKRPGYDHNKK
AGAMNAMVRASAVLSNGPFMLNFDCDHYVYNCQAIREAMCYMLDRGGDRICYIQFPQRFEGIDPSD
RYANHNTVFFDGNMRALDGLQGPMYVGTGCLFRRYAIYGFNPPRAVEYHGLVGQTRVPIDPHARSG
DGIADELRPLSDHPDHEAPQRFGKSKMFIESIAVAEYQGRPLADHPSVRNGRPAGALLMPRPPLDAA
TVAEAVSVISCWYEDNTEWGLRVGWIYGSVTEDVVTGYRMQNRGWRSVYCITKRDAFRGTAPINLT
DRLHQVLRWATGSVEIFFSKNNAMLASRRLMFLQRMSYINVGIYPFTSLFLIMYCLLPALSLFSGQFIV
ATLDPTFLCYLLLITVTLVLLCLLEVKWSGIGLEEWWRNEQFWVIGGTSAHLAAVLQGLLKVAAGIEISF
TLTAKAAAEDDDDPFAELYLIKWTSLFIPPLAIIGINIIAMVVGVSRCVYAEIPQYSKLLGGGFFSFWVLA
HYYPFAKGLMGRRGRTPTIVYVWAGLISITVSLLWITISPPDDRVSQSGIEV
>CSLD_7BL_TGACv1_577301_AA1871610.1_Triticum_aestivum
MASDHTNYTVFMPPTPDNQPGAAPAPASGGSTKPDNLPLPRYTSGSKLVNRRSGDDGAAGGAKMD
RGLSTEHVASPSKSLLVRSQTGEFDHNRWLFETQGTYGIGNAYWPQDENDDGAGMGGGSVKMEDL
VDKPWKPLSRKVAIPPGILSPYRLLVLVRFVALFLFLIWRATNPNPDAMWLWGISIVCEYWFALSWLLD
QMPKLNPINRAADLAALREKFESKTPSNPTGRSDLPGLDVFISTADPYKEPPLVTANTLLSILATDYPV
EKLFVYISDDGGALLTFEAMAEACAYAKVWVPFCRKHSIEPRNPEAYFTQKGDPTKGKKRPDFVKDR
RWIKREYDEYKVRINDLPEAIKRRAKAMNAHERKIARETAAASSDAAPPPVKATWMADGTHWPGTW
LDSAPDHGKGDHASIVQVMIKNPHHDVVYGEADDHAYLDFTNVDVRIPMFVYLSREKRPGYDHNKKA
GAMNAMVRASAVLSNGPFMLNFDCDHYVYNCQAIREAMCYMLDRGGDRICYIQFPQRFEGIDPSDR
YANHNTVFFDGNMRALDGLQGPMYVGTGCLFRRYAIYGFNPPRAVEYHGLVGQTRVPIDPHARSGD
GVADELRPLSDHPDHEAPQRFGKSKMFIESIAVAEYQGRPLADHPSVRNGRPAGALLMPRPPLDAAT
VAEAVSVISCWYEDNTEWGLRVGWIYGSVTEDVVTGYRMHNRGWRSVYCITKRDAFRGTAPINLTD
RLHQVLRWATGSVEIFFSKNNAMLASRRLMFLQRMSYINVGIYPFTSLFLIMYCLLPALSLFSGQFIVAT
LDPTFLCYLLLITVTLVLLCLLEVKWSGIGLEEWWRNEQFWVIGGTSAHLAAVLQGLLKVAAGIEISFTL
TAKAAAEDDDDPFAELYLIKWTSLFIPPLAIIGINIIAMVVGVSRCVYAEIPQYSKLLGGGFFSFWVLAHY
YPFAKGLMGRRGRTPTIVYVWAGLISITVSLLWITISPPDDRVSQSGIEV
>CSLD_7AL_TGACv1_559436_AA1799630.1_Triticum_aestivum
MASDHTNYTVFMPPTPDNQPGAASAPASGGPTKPDNLPLPRSSGSKLVNRRSGDDGAAGGGKMD
RRLSPVQVASPSKSLLVRSQTGEFDHNRWLFETQGTYGIGNAYWPQEDNDDGAGMGGGSVKMED
LVDKPWKPLSRKVAIPPGILSPYRLLVLVRFVALFLFLVWRATNPNPDAMWLWGISIVCEYWFALSWL
LDQMPKLNPINRAADLAALREKFESKTPSNPTGRSDLPGLDVFISTADPYKEPPLVTANTLLSILATDY
PVEKLFVYISDDGGALLTFEAMAEACAYAKVWVPFCRKHSIEPRNPEAYFTQKGDPTKGKKRPDFVK
DRRWIKREYDEYKVRINDLPEAIKRRAKAMNAHERKIARETAAASSDAAPPPVKATWMADGTHWPGT
WLDSAPDHGKGDHASIVQVMIKNPHHDVVYGDADDHAYLDFTNVDVRIPMFVYLSREKRPGYDHNK
KAGAMNAMVRASAVLSNGPFMLNFDCDHYVYNCQAIREAMCYMLDRGGDRICYIQFPQRFEGIDPS
DRYANHNTVFFDGNMRALDGLQGPMYVGTGCLFRRYAIYGFNPPRAVEYHGLVGQTRVPIDPNARS
GDGVADELRPLSDHPDHEAPQRFGKSKMFIESIAVAEYQGRPLADHPSVRNGRPPGALLMPRPPLD

AATVAEAVSVISCWYEDNTEWGLRVGWIYGSVTEDVVTGYRMHNRGWRSVYCITKRDAFRGTAPIN
LTDRLHQVLRWATGSVEIFFSKNNALLASRRLMFLQRMSYINVGIYPFTSLFLIMYCLLPALSLFSGQFI
VATLDPTFLCYLLLITITLVLLCLLEVKWSGIGLEEWWRNEQFWVIGGTSAHLAAVLQGLLKVAAGIEIS
FTLTAKAAAEDDDDPFAELYLIKWTSLFIPPLAIIGINIIAMVVGVSRCVYAEIPQYSKLLGGGFFSFWVL
AHYYPFAKGLMGRRGRTPTIVYVWAGLISITVSLLWITISPPDDRVSQSGIEV
>OS01G54620_OsCesA
MMESGVPPCAACGDDAHAACRACSYALCKACLDEDAAEGRTTCARCGGEYGAPDPAHGQGAVVE
EEVEESHEPAAGGVRERVTMASQLSDHQDEGVHARTMSTHARTISSVSGVGSELNDESGKPIWKNR
VESWKEKKKEKKASAKKAAAKAQAPPVEEQIMDEKDLTDAYEPLSRIIPISKNKLTPYRAVIIMRLVVLG
LFFHYRITNPVYSAFGLWMTSVICEIWFGFSWILDQFPKWCPINRETYVDRLIARYGDGEDSGLAPVD
FFVSTVDPLKEPPLITANTVLSILAVDYPVEKISCYVSDDGSAMLTFESLAETAEFARRWVPFCKKYSIE
PRAPEFYFSQKIDYLKDKIHPSFVKERRAMKRDYEEYKVRINALVAKAQKTPEEGWIMQDGTPWPGN
NPRDHPGMIQVFLGETGARDFDGNELPRLVYVSREKRPGYQHHKKAGAMNALVRVSAVLTNAPYIL
NLDCDHYVNNSKAVREAMCFMMDPSVGRDVCYVQFPQRFDGIDRSDRYANRNVVFFDVNMKGLD
GLQGPVYVGTGCCFYRQALYGYGPPSLPALPKSSVCSWCCCCCPKKKAEKSEKEMHRDSRREDLE
SAIFNLREIDNYDEYERSMLISQMSFEKSFGLSSVFIESTLMENGGVPESANPSTLIKEAIHVISCGYEE
KTEWGKEIGWIYGSVTEDILTGFKMHCRGWRSIYCMPIRPAFKGSAPINLSDRLHQVLRWALGSVEIF
LSRHCPLWYGYGGGRLKWLQRLSYINTIVYPFTSLPLIAYCCLPAICLLTGKFIIPTLSNAATIWFLGLFIS
IIVTSVLELRWSGIGIEDWWRNEQFWVIGGVSAHLFAVFQGILKMIAGLDTNFTVTAKATDDTEFGELY
VFKWTTVLIPPTSILVLNLVGVVAGFSDALNSGYESWGPLFGKVFFAMWVIMHLYPFLKGLMGRQNR
TPTIVVLWSVLLASVFSLLWVKIDPFIGSSETTTTNSCANFDC
>CSLD_5DS_TGACv1_457675_AA1488780.1_Triticum_aestivum
MSRRLSLPASSPVTVTVSPTRGKGAGGGSPGDGVVRRGSGLTSPVPRHSIGSSTATLQVSPVRRSG
GSRYASRDGADASAEFVHYTVHIPPTPDRNTASASTDAPVAEEEGEVLPQRSYVSGTIFTGGLNCTT
RAHVLSNSADGARPAASVNMSCKMRGCDMPAFLNAGRGGRPPCDCGFMICEECYMDCVAAAGNC
PGCKEAYSAGSDTDDSVDEDDDDAISSSEERDQMPMTSMSKRFSMVHSIKMPMPSSNGNGGGKPA
DFDHARWLFETKGTYGYGNALWPKNEHGGGGNTAGATSGFVGIEEPPNFGARCRRPLTRKTSVSQ
AILSPYRMLIAIRLVALGFFLAWRIRHPNPDAMWLWALSVTCEVWFAFSWLLDSLPKLCPVNRSCDLD
VLADRFELPTARNPKGRSDLPGIDVFVSTADPEKEPPLVTANTILSILAADYPVEKLACYLSDDGGALL
TFEALAETASFARTWVPFCRKHGVEPRCPESYFGQKRDFLKNKVRLDFVRERRKVKREYDEFKVRV
NSLTEAIRRRSDAYNAGEELRARRRLQEEAVAAGGALGAAPLAETGAVKGTWMSDGSQWPGTWLT
GATDHARGDHAGIIQAMLAPPTSEPVLGGVPAESGALIDTTGVDIRLPMLVYVSREKRPGYDHNKKA
GAMNALVRTSAIMSNGPFILNLDCDHYVHNSAALREGMCYMLDRGGDRVCYVQFPQRFEGIDPNDR
YANHNLVFFDVAMRAMDGLQGPMYVGTGCIFRRTALYGFSPPRATEHHGWLGRKKIKLFLRKPTTG
KKTDRESEHESMLPPIEDDDHNQLGDIESSALMPKRFGSSATFVSSIPVAEYQGRLLQDMPGVHQGR
PAGALAVPREPLDAATVGEAISVISCFYEEKTEWGRRIGWIYGSVTEDVVTGYRMHNRGWRSVYCVT
RRDAFRGTAPINLTDRLHQVLRWATGSVEIFFSRNNALFATRRMKLLQRVAYFNVGMASSR
>AT1G32180_CslD6
MMDGESPLRHPRISHVSNSGSDFGSSSDYNKYLVQIPPTPDNNPGPASLSIVLLEIDSNQESVPSVSG
DIVSGSSGKDNEPDLTDVRINVGEEEEDDTLLSKISYSLTRVVKISPIIIALYRILIVVRVVSLALFLFWRIR
NPNNKALWLWLLSVICELWFAFSWLLDQIPKLFPVNHATDIEALKATFETPNPDNPTGKSDLPGIDVFV
STADAEKEPPLVTANTILSILSVDYPVEKLSVYISDDGGSLVTFEAIAEAASFAKIWVPFCRKHKIEPRNP
ESYFGLKRDPYKDKVRHDFVRERRYVKRAYDEFKVRVNALPHSIRRRSDAFNSKEEIKALEKWKHWK
VKVEEDQIKEPRPALVAPKATWMSDGTHWPGTWAVSGPHHSRGDHASVIQVLLDPPGDEPVEGKG
GEGRALDLEGVDIRLPMLVYVSREKRPGYDHNKKAGAMNALVRASAIMSNGPFILNLDCDHYVYNSR
AFRDGICFMMDHDGDRVSYVQFPQRFEGIDPSDRYANKNTVFFDINLRALDGIQGPMYVGTGCLFRR
TALYGFNPPDVFVVEEEPSGSYCFPLIKKRSPATVASEPEYYTDEEDRFDIGLIRKQFGSSSMLVNSV
KVAEFEGRPLATVHSSRLGRPPGSLTGSRKPLDFATVNEAVNVISCWYEDKTEWGFNVGWIYGSVT
EDVVTGFRMHEKGWRSFYCVTEPDAFRGSAPINLTDRLHQVLRWATGSVEIFFSRNNAIFAGPKLKL
LQRIAYLNVGIYPFTSIFILTYCFLPPLSLFSGHFVVETLTGSFLIYLLIITLSLCGLAVLEVKWSGISLEEW
WRNEQFWLIGGTSAHLVAVLQGILKVIAGVEISFTLTSKSSTGGDDEDDEFADLYLFKWTALMIPPLTIII
LNIVAILFAVCRTVFSANPQWSNLLGGTFFASWVLLHMYPFAKGLMGRGGKTPTVVYVWSGLIAICLS
LLYITIKNSEIDGGSFMLV
>OS08G06380_OsCslF6
MAPAVAGGGGRRNNEGVNGNAAAPACVCGFPVCACAGAAAVASAASSADMDIVAAGQIGAVNDES
WVAVDLSDSDDAPAAGDVQGALDDRPVFRTEKIKGVLLHPYRVLIFVRLIAFTLFVIWRIEHKNPDAM
WLWVTSIAGEFWFGFSWLLDQLPKLNPINRVPDLAVLRRRFDHADGTSSLPGLDIFVTTADPIKEPILS
TANSILSILAADYPVDRNTCYLSDDSGMLLTYEAMAEAAKFATLWVPFCRKHAIEPRGPESYFELKSH
PYMGRAQEEFVNDRRRVRKEYDDFKARINGLEHDIKQRSDSYNAAAGVKDGEPRATWMADGSQWE
GTWIEQSENHRKGDHAGIVLVLLNHPSHARQLGPPASADNPLDFSGVDVRLPMLVYVAREKRPGCN
HQKKAGAMNALTRASAVLSNSPFILNLDCDHYINNSQALRAGICFMLGRDSDTVAFVQFPQRFEGVD
PTDLYANHNRIFFDGTLRALDGLQGPIYVGTGCLFRRITLYGFEPPRINVGGPCFPRLGGMFAKNRYQ

KPGFEMTKPGAKPVAPPPAATVAKGKHGFLPMPKKAYGKSDAFADTIPRASHPSPYAAEAAVAADEA
AIAEAVMVTAAAYEKKTGWGSDIGWVYGTVTEDVVTGYRMHIKGWRSRYCSIYPHAFIGTAPINLTER
LFQVLRWSTGSLEIFFSRNNPLFGSTFLHPLQRVAYINITTYPFTALFLIFYTTVPALSFVTGHFIVQRPT
TMFYVYLAIVLGTLLILAVLEVKWAGVTVFEWFRNGQFWMTASCSAYLAAVLQVVTKVVFRRDISFKL
TSKLPAGDEKKDPYADLYVVRWTWLMITPIIIILVNIIGSAVAFAKVLDGEWTHWLKVAGGVFFNFWVL
FHLYPFAKGILGKHGKTPVVVLVWWAFTFVITAVLYINIPHIHGPGRHGAASPSHGHHSAHGTKKYDF
TYAWP
>GRMZM2G122277_P01
MAPGGRRSNGETPTGQQQQQQQADGRRGCACGGFPVCACAGAAAVASAASSADMDRVAVAATE
GQIGAVNDESWVAVDLSDDGLSSAADPGAVALEERPVFRTEKIKGVLLHPYRVLIFVRLIAFTLFVIWRI
SHRNPDALWLWVTSIAGEFWFGFSWLLDQLPKLNPINRVPDLGALRQRFDRADGTSRLPGLDIFVTT
ADPFKEPILSTANSILSILAADYPVERNTCYLSDDSGMLLTYEAMAEAAKFATVWVPFCRKHGIEPRGP
ESYFELKSHPYMGRSQEDFVNDRRRVRRDYDEFKARINGLENDIRQRSDAYNAARGLKDGEPRATW
MADGTQWEGTWVEPSENHRKGDHAGIVLVLLNHPSHSRQLGPPASADNPLDLSMVDVRLPMLVYV
SREKRPGHNHQKKAGAMNALTRCSAVLSNSPFILNLDCDHYINNSQALRAGICFMLGRDSDTVAFVQ
FPQRFEGVDPTDLYANHNRIFFDGTLRALDGMQGPIYVGTGCLFRRITLYGFDPPRINVGGPCFPSLG
GMFAKTKYEKPGLELTTKAAVAKGKHGFLPMPKKSYGKSDAFADTIPMASHPSPFAAAAAVVAEEATI
AEAVAVCAAAYEKKTGWGSDIGWVYGTVTEDVVTGYRMHIKGWRSRYCSIYPHAFIGTAPINLTERLF
QVLRWSTGSLEIFFSRNNPLFGSTFLHPLQRVAYINITTYPFTAIFLIFYTTVPALSFVTGHFIVQRPTTM
FYVYLAIVLGTLLILAVLEVKWAGVTVFEWFRNGQFWMTASCSAYLAAVCQVLVKVVFRRDISFKLTS
KQPAGDEKKDPYADLYVVRWTWLMVTPIIIILVNIIGSAVAFAKVLDGEWTHWLKVAGGVFFNFWVLF
HLYPFAKGILGRHGKTPVVVLVWWAFTFVITAVLYINIPHIHGPGGKHGGAIGKHGAAHHGKKFDGYY
LWP
>GRMZM2G110145_P01
MAAGQQQASGGAKHGCVCGFPVCACAGAAAVASAASSADMDRVAVAATEGQIGAVNDESWIAVDL
SDDGLSADGADPGVALEDRPVFRTEKIKGVLLHPYRVLIFVRLIAFTLFVIWRISHRNPDALWLWVTSIA
GEFWFGFSWLLDQLPKLNPINRVPDLAALRQRFDRAGGGAGGGTSLLPGLDVFVTTADPFKEPILST
ANSVLSILAADYPVERNTCYLSDDSGMLLTYEAMAEAAKFATVWVPFCRKHGIEPRGPESYFDLKSH
PYMGRSQEDFVNDRRRVRKDYDEFKARINGLDHDIKQRSDAYNAARGLKDGEPRATWMADGTQWE
GTWVEPSENHRKGDHAGIVLVLLNHPSHSRQLGPPASADNPLDLSMVDVRLPMLVYVSREKRPGHN
HQKKAGAMNALTRCSAVLSNSPFILNLDCDHYINNSQALRAGICFMLGRDSDTVAFVQFPQRFEGVD
PTDLYANHNRIFFDGTLRALDGMQGPIYVGTGCLFRRITLYGFDPPRINVGGPCFPALGGMFAKAKYE
KPGLELTTTKAAVAKGKHGFLPMPKKSYGKSDAFADTIPMASHPSPFAAASAASVVADEATIAEAVAV
CAAAYEKKTGWGSDIGWVYGTVTEDVVTGYRMHIKGWRSRYCSIYPHAFIGTAPINLTERLFQVLRW
STGSLEIFFSRNNPLFGSTFLHPLQRVAYINITTYPFTAIFLIFYTTVPALSFVTGHFIVQRPTTMFYVYLA
IVLGTLLILAVLEVKWAGVTVFEWFRNGQFWMTASCSAYLAAVCQVLVKVVFRRDISFKLTSKQPAGD
EKKDPYADLYVVRWTWLMVTPIIIILVNIIGSAVAFAKVLDGEWTHWLKVAGGVFFNFWVLFHLYPFAK
GILGRHGKTPVVVLVWWAFTFVITAVLYINIPHIHGPGGKHGGAIGRHGGDAHHHGKKFDGYYLWP
>CSLA_3B_TGACv1_223332_AA0780350.1_Triticum_aestivum
MLLLKIDIAKAFDTVSWEYILELLQRMNFPAHWRDRIALLLSSVSSAYLLKGDPGPAILHQRGLRQGDP
LSAILFILVIVPLHRMLEAAQQAGTIAPLPAGAARLRVTLYADDAIFFANPVRQEIDTIMQLLQGFGEAAG
LRGNPQKSSAATLNYGSIDLIDVLKNFSGTRVGFPIRYLGLPLCIGRLPLCTRVGFPIRYLGWLLGKAN
SCIAPPLAVASHVLVRCVLSALPAFAMAVLRIPKRFYKDVDKARWRFLWVHDHEVTGGRCKVNWRLV
TSPVDHGGLGIPSMERFARALCLRWLWLAWTDPARPWARMGTPCDDKDRALFASATTVTVGDGNR
VLFWHCSWLGEQPVRQDYPNLFRRSTRKNRMMADAIRDDRWIMDLRRSGAGEEVMAMAATAGLW
AEVPVRLDWATVAAQCALAGEQARAFLVVPAVRLLVLLSLAMTVMILLEKLFVAAVCYAAKALGHRPE
RRYKWGPVAASACKTGGDDEEDGIVGVGSGSGSAAFPVVLVQIHMYNEREDLVKIECQRWKSKGV
NIRYEVRQNRKGYKAGALKEGLIRDYVRECEFIAMFDADFQPESDFLLRTVPFLVHNPDIALVQTRWK
FVNSDECLLTRFLEMSLDYHFKFEQEAGSIVYSFFGFNGTAGVWRISAINDAGGWKERTMVEDMDLV
VRTALLGLKFVYIGAVKVKSELPSTFKAYRFQQHRWSCGPANLFKKMLVEILQNKKVSFWSKLHLLYD
FFFVGKIIAHIVTFIYYCFATPVSVFFPEIQIPLWGVVYVPTVITLCKALGSPSSFHLVILWVLFDNVMSLH
RIKATITGLLDARRVNEWVVTEKLGDTNKTEPAMEGLNDVQVIDVELSTPLVPKLEKRYNCSEIFVGTC
IIICGCYDVLYANKGCYIYLFIQGVAFLVIGFEYIGTRPPGAE
>GRMZM2G103972_P01
MAAAVTRRVGLHVEATNGGADDESRRNSSAADHSPVAKRINDAAANAKRNDVWVAAQEGEMPAAA
GNSSQPPLLFRTMKVKGSILHPYRFVILLRLVAIVAFFIWRIRNRNRDGVWLWAMSMVGDVWFGFSW
VLNQLPKLNPIKRVPDLAAIRDQYEQPSASGGESNNKLPGIDVFVTTVDPVDEPILYTVNSVLSILATDY
PVEKYACYLSDDGGTLVHYEAMLEVASFARLWAPFCRKHSVEPRAPESYFGVKRRQPYTGSVQGEF
TSDHRRMRREYEEFKVRIDSLFSTVCQRSQAYNRKHAKDDEAGMVMKATWMADGTQWPGTWIEQ
AENHRKGHHAGIVKVVLNHPGHKPELGSPASIDNPFDFSNTDTRLPMLVYMSREKRTGYNHQKKAG
AMNAMLRVSALLSNAPFLINFDCDHYVNNSQAFRASMCFMLDPRDGRNTAFVQFPQRFDGVDPTDR
YANHNRVFFDGTMLSLNGLQGPSYLGTGTMFRRAALYGMEPPRWRTTGSVKVIDDDDDHKGKEYG

RSTLFRNAVLDDAANQERSITPVFLDDDETTTISSEVASLMTCAYEDGTTWGRDVGWVYNIATEDVVT
GFRMHRQGWRSMYCSVEPAAFRGTAPINLTERLLQVLRWSGGSLEMFFSHSNAFLAGARMHPLQR
VAYLNMSTYPVVTVFILAYNLFPLMWLVSERYYIQRPFGTYVLYLVATIAMIHVIGMFEVRWAGITLLD
WCRNEQFYMIGATGVYPTAVLYMALKLVTGKSIHFRLTSKQTEACSGGDKFADLYVVRWVPLLVPTIA
VLAVNVAAVGVAVGKAATWGLLTQQAQHALLGMVFNVWILVLLYPFALGVMGRWGKRPAILFGVLV
MAIGAVAVVYISFRATYYPTGWSEMETTSALGKAESFF
>GRMZM2G113432_P01
MAPKAPANNGAVVVAGGHHGGSTDVGGVAGGATTTTVPTSVAANGNGKQHGTRKQRRPAAVVVS
PMDKYWTPVDDKEAAEAVDDGGEDGRRPLLFRTYKVKGILLHPYRLLTLLRLIAIILFFIWRIRHPHADG
MWLWWISIVGDFWFGVTWLLNQVAKLNPTKRVPDLSLLRQQFDLPDGNSNLPRLDVFINTVDPINEP
MIYTMNSILSILAVDYPIDRTATYLSDDGGSIIHYEGLLETANFATLWVPFCRKHSIEPRAPESYFAVKS
RPYTGNVPDEFADDHRRMSKEYDEFKVRLDALFTKIPERSDAHNAEAKEGVKATWMADGTQWPGT
WFDPAENHKKGQHAGIVKVMLNHPGDEPRFGGPASAETPLDFSAVDVRLPMLVYISREKSPSHDHQ
KKAGAMNVQLRISALLTNAPFIINFDGDHYVNNSQAFRAAMCFMLDRRDGENTAFVQFPQRFDDVDP
TDRYCNHNRVFFDATLLGLNGIQGPSYVGTGCMFRRIAVYGIDPPRWRTDAFKLVDNPSKFGSSMLF
INSIPSAANQEWSMASPPAHEESVMEELNNVMKCAYEEGTEFGKEIGWVYNIATEDVVTGFRVHRTG
WRSMYCRMEPDAFRGTAPINLTERLCQILRWSGGSLEMFFSHCPLLAGRRLNLMQRIAYTNMTAYPI
SSVFLVFYLLFPVIWIFRGEFYIQKPFPTYVLYLVVIIAMTELIGMVEIKWAGLTLLDWIRNEQFYIIGATA
VYPLATLHIVLKLVLRGNGVSFKLTAKQATSAVNEKYAEMYVVQWTPLLIPTIAVIAVNVGAIGAAIGKA
VVGGWSLLQMADASLGLVFNAWILLLIYPFALGVMGRWSKRPYILFVLFMIGFAVVAAVVVAIHAART
GSVRFHSGHSGGASFPTSWGF
>CSLF_2AS_TGACv1_112322_AA0335280.1_Triticum_aestivum
MGSLAAANGAGHASNGAGVADQALALENGTGNGHKAGVANRATPPLQANGGSKVAKKISPKDKYW
VAADEGEMAAAIADGGEDGRRPLLYRTFKVKGILLHPYRLLSLIRLVAIVLFFVWRVRHPYADGMWLW
WISMVGDLWFGVTWLLNQVAKLNPVKRVPNLTLLEQQFDLPDGNSNLPCLDVFINTVDPINEPMIYTM
NSIISILAADYPVDKHACYLSDDGGSIIHYDGLLETAKFAALWVPFCRKHSIEPRAPESYFSLNTRPYTG
NAPQDFVNDRRHMCREYDEFKERLDALFTLIPKRSDVYNHAAAKEGAKATWMADGTQWPGTWIDP
AENHKKGQHVGIVKVMLKHPSYEPELGLGASTNSPLDFSAIDVRLPMLVYISREKSPSCDHQKKAGA
MNVQLRVSALLTNAPFIINFDGDHYVNNSKAFRAGICFMLDRREGDNTAFVQFPQRFDDVDPTDRYC
NHNRVFFDATLLGLNGIQGPSYVGTGCMFRRVALYGVDPPRWRPDNVKIVDSSTKFGSSASFISSILP
AADQERSIMSPPALEEPVMADLAHVMTCAYEDGTEWGREVGWVYNIATEDVVTGFRLHRNGWRSM
YCRMEPDAFAGTAPINLTERLYQILRWSGGSLEMFFSRNCPLLAGRRLHPMQRIAYANMTAYPVSSV
FLVFYLLFPVIWIFRGQFYIQKPFPTYVLYLVIVIALTELIGMVEIKWAGLTLLDWIRNEQFYIIGATAVYPT
AVFHIVLKLFGLKGVSFKLTAKQVASSTSDKFAELYAVQWAPMLIPTMVVIAVNVCAIGASIGKAIVGG
WSLMQMADAGLGLVFNAWILVLIYPFALGMIGRWSKRPYILFILFVIAFILIALVDIAIQAMRSGFVRFHF
KSSGGATFPTSWGL
>OS07G36740_OsCslF4
MSAAAVTRRINAGGLRVEVTNGNGAAGVYVAAAAAPCSPAAKRVNDGGGKDDVWVAVDEADVSGP
SGGDGVRPTLFRTYKVKGSILHPYRFLILVRLIAIVAFFAWRVRHKNRDGAWLWTMSMAGDVWFGFS
WALNQLPKLNPIKRVADLAALADRQQHGTSGGGELPGVDVFVTTVDPVDEPILYTVNSILSILAADYPV
DRYACYLSDDGGTLVHYEAMVEVAKFAELWVPFCRKHCVEPRAPESYFAMKTQAYRGGVAGELMS
DRRRVRREYEEFKVRIDSLFSTIRKRSDAYNRAKDGKDDGENATWMADGTHWPGTWFEPAENHRK
GQHAGIVQVLLNHPTSKPRFGVAASVDNPLDFSGVDVRLPMLVYISREKRPGYNHQKKAGAMNALLR
VSALLSNAPFIINFDCDHYVNNSQAFRAPMCFMLDRRGGGDDVAFVQFPQRFDDVDPTDRYANHNR
VFFDGTTLSLNGLQGPSYLGTGTMFRRAALYGLEPPRWGAAGSQIKAMDNANKFGASSTLVSSMLD
GANQERSITPPVAIDGSVARDLAAVTACGYDLGTSWGRDAGWVYDIATEDVATGFRMHQQGWRSV
YTSMEPAAFRGTAPINLTERLYQILRWSGGSLEMFFSHSNALLAGRRLHPLQRIAYLNMSTYPIVTVFI
FFYNLFPVMWLISEQYYIQQPFGEYLLYLVAIIAMIHVIGMFEVKWSGITVLDWCRNEQFYMIGSTGVY
PTAVLYMALKLFTGKGIHFRLTSKQTTASSGDKFADLYTVRWVPLLIPTIVVLAVNVGAVGVAVGKAAA
WGLLTEQGRFAVLGMVFNVWILALLYPFALGIMGQRGKRPAVLFVATVMAVAAVAIMYAAFGAPYQA
GLSGVAASLGKAASLTGPSG
>OS07G36690_OsCslF2
MAATAASTMSAAAAVTRRINAALRVDATSGDVAAGADGQNGRRSPVAKRVNDGGGGKDDVWVAVD
EKDVCGARGGDGAARPPLFRTYKVKGSILHPYRFLILLRLIAIVAFFAWRVRHKNRDGVWLWTMSMV
GDVWFGFSWVLNQLPKLSPIKRVPDLAALADRHSGDLPGVDVFVTTVDPVDEPILYTVNTILSILAADY
PVDRYACYLSDDGGTLVHYEAMVEVAKFAELWVPFCRKHCVEPRSPENYFAMKTQAYKGGVPGEL
MSDHRRVRREYEEFKVRIDSLSSTIRQRSDVYNAKHAGENATWMADGTHWPGTWFEPADNHQRGK
HAGIVQVLLNHPSCKPRLGLAASAENPVDFSGVDVRLPMLVYISREKRPGYNHQKKAGAMNVMLRV
SALLSNAPFVINFDGDHYVNNSQAFRAPMCFMLDGRGRGGENTAFVQFPQRFDDVDPTDRYANHN
RVFFDGTMLSLNGLQGPSYLGTGTMFRRVALYGVEPPRWGAAASQIKAMDIANKFGSSTSFVGTML
DGANQERSITPLAVLDESVAGDLAALTACAYEDGTSWGRDVGWVYNIATEDVVTGFRMHRQGWRS
VYASVEPAAFRGTAPINLTERLYQILRWSGGSLEMFFSHSNALLAGRRLHPLQRVAYLNMSTYPIVTV

FIFFYNLFPVMWLISEQYYIQRPFGEYLLYLVAVIAMIHVIGMFEVKWAGITLLDWCRNEQFYMIGSTGV
YPTAVLYMALKLVTGKGIYFRLTSKQTTASSGDKFADLYTVRWVPLLIPTIVIIVVNVAAVGVAVGKAAA
WGPLTEPGWLAVLGMVFNVWILVLLYPFALGVMGQWGKRPAVLFVAMAMAVAAVAAMYVAFGAPY
QAELSGGAASLGKAAASLTGPSG
>OS07G36630_OsCslF8
MAANGGGGGAGGCSNGGGGGAVNGAAANGGGGGGGGSKGATTRRAKVSPMDRYWVPTDEKEM
AAAVADGGEDGRRPLLFRTFTVRGILLHPYRLLTLVRLVAIVLFFIWRIRHPYADGMFFWWISVIGDFW
FGVSWLLNQVAKLKPIRRVPDLNLLQQQFDLPDGNSNLPGLDVFINTVDPINEPMIYTMNAILSILAADY
PVDKHACYLSDDGGSIIHYDGLLETAKFAALWVPFCRKHSIEPRAPESYFAVKSRPYAGSAPEDFLSD
HRYMRREYDEFKVRLDALFTVIPKRSDAYNQAHAEEGVKATWMADGTEWPGTWIDPSENHKKGNH
AGIVQVMLNHPSNQPQLGLPASTDSPVDFSNVDVRLPMLVYIAREKRPGYDHQKKAGAMNVQLRVS
ALLTNAPFIINFDGDHYVNNSKAFRAGICFMLDRREGDNTAFVQFPQRFDDVDPTDRYCNHNRVFFD
ATLLGLNGIQGPSYVGTGCMFRRVALYGVDPPRWRPDDGNIVDSSKKFGNLDSFISSIPIAANQERSII
SPPALEESILQELSDAMACAYEDGTDWGKDVGWVYNIATEDVVTGFRLHRTGWRSMYCRMEPDAF
RGTAPINLTERLYQILRWSGGSLEMFFSHNCPLLAGRRLNFMQRIAYINMTGYPVTSVFLLFYLLFPVI
WIFRGIFYIQKPFPTYVLYLVIVIFMSEMIGMVEIKWAGLTLLDWIRNEQFYIIGATAVYPLAVLHIVLKCF
GLKGVSFKLTAKQVASSTSEKFAELYDVQWAPLLFPTIVVIAVNICAIGAAIGKALFGGWSLMQMGDAS
LGLVFNVWILLLIYPFALGIMGRWSKRPYILFVLIVISFVIIALADIAIQAMRSGSVRLHFRRSGGANFPTS
WGF
>CSLA_2DS_TGACv1_177798_AA0584795.1_Triticum_aestivum
MAAWNPETHGSGAIIVGADDCETTVEDEMAAGRDANTKLFHRVANGRKLKNFIPAISVEGITITDQAA
KEEAFFEAYSELLGRCGSREHTLDLDYLGIESINLEDQDLVFQEEEVWKVVRDMPSDRALGPNGFIGV
FFQKAWAIVKRDVMAALNKLFLNNGRGFGRLNQALITLIPKNHEACQIKDFRPICLVHSIPKLASKLLAT
RLCPRMGELVHAKQSAFIKGRNIHDNFLQVRQLARKLYKRKTKSVMLKLDISRAFDSLSWPFLFEVLR
VKGFSRTWRFWIATLLTTASSRVVVNGCVGKKFMHACGLRQGDSISPLLFVIAMDVLSAMILKARETN
AVSKIPGCAPIQRLSLYVDDVVMFIKPSWTDLWFVQEALRVFGEASGLKVNFSKSSAVMIRSEEEEEV
LVRKAMPWKMETFPIKYLGLQLGIKQLTRSEWQPVVDQALKMMPGWQRGPVTRPGRLPLVNQVVR
ARPIHHLIVAEAPKRALDRVDKGCRAFFWAGSEEIQGGQCAVAWRGVYRPKQMGGLGVVDLHKHGI
ALRLSLSQTSFSEQSRSSCTIQKLLLFKLSGPSNLNGTVSLLTRIQKMFFDYHFKVEQEAGSATFAFFS
FNGTAGVWRTAAIKEAGGWKDRTTVEDMDLAIRATLKGWKFIYVGDIRVKSELPSSYKAYCRQQFR
WACGGANLFRKVAVDILTSKDVSVIKKFYMLYSFLFVRRVVAPAVACILSNIIVPLSVMIPELYLPVWGV
AYIPTVLLVVTAIRNPKNIHLLPFWILFESVMTIHRTRAALVGLFELTEFDEWLVTKKTGNNFEDNKVPLL
QKTRKRLRDRVNFPEILFSAFLFFCASYNLVFPGKTSYYFNLYFQGLAFAFLGLNFTGTCTCFQ
>CSLF_2AS_TGACv1_112790_AA0345230.1_Triticum_aestivum
MTTSPATHDGAATGLSEPLLPNRNGVHAGALVVTPVVANGHGGGDKLKGDLKAKDKYWKDVDQPD
DVAAAPDLENGGGRPLLFSNRRVKNIILYPYRVLILIRVIAVILFVGWRIKHNNSDVMWFWVMSVVADV
WFSLSWLSYQLPKYNPIKMIPDLATLRKQFDTPGRSSQLPGIDVIVTTASATDEPILYTMNGVLSILAAD
YHIGRCNCYLSDDSGSLVLYEALVETAKFAALWVPFCRKHQIEPRAPESYFELEGTLCGGASHKEFIQ
DYKHVRTQYDEFKKHLDMLPNTIRQRSDIYSRTGTKDEDATVTWMADGTQWPGTWLDPTEKHRPG
HHAGIVKIVQSHPEHVVPLGVQESNDNPLNFDDVDMRLPMLVYVAREKSPGVEHNKKAGALNAELRI
SALLSNAPFFINFDCDHYINNSEALRAAICFMLDPREGDNTGFVQFPQRFDNVDPTDRYGNHNRVFF
DGAMYGLNGQQGPTYLGTGCMFRRLALYGIDPPCWRDEDIIVDSNRFGNSLLFLNSVLAAIKQEEGV
TLQPPLDDSFLEEVTKVVSSSYDDSTDWGRGIGYIYNMATEDIVTGFRIHGQGWRSMYATMEREAFR
GTAPINLTERLRQIVRWSGGSLEMFFSHISPLFAGRRLSLVQRLSYINFTIYPLTSLFILMYAFCPVMWL
LPTEILVQRPYTRYIVYLIIVIAMIHVIGMFEIMWAGITWLDWWRNEQFFMIGSVTAYPTAVLHMVVNLLT
KKGIHFRVTTKQPVADTDDKYAEMYEVHWVPMMVPAVVILFSNILAIGVAIGKSVLYMGTWSAAQRR
HGALGLLFNLWIMVLLYPFALAIIGRWAKRTGILFILLPIAFLSTALMYIGIHTFLLHFFPSMLI
>CSLF_2AS_TGACv1_112322_AA0335290.1_Triticum_aestivum
MVSPATGGGRGGNAGLAEPLLATNDDSDGAKHVFGAKAKHWVPADEKEMAASRECGGEDGRPLL
YRTFKVRGFLVNTYRFLNLARLTAVIVFFAWRVQHPDSDAMWLWWISVVGDFWFGLSWWLNQVPKL
NPTICIPTIPLLRQQFDLPDGGSNLPVLDVFISTVDPVEEPMLHTMNSILSILATDYPVDKYATYLSDDG
GSLLHYDGLVETAKFAALWVPFCRKHHVEPRAPESYFGMKVRPYKGNLPEEFLDDHRRLRREYEEF
KTRLDALFTVIPQRSEAHGREDAKGGGGAKATWMADGTQWPGTWTEPAEGHRKGDHAGIIQVMLS
QPSGEPQLGAPASSDDNPLDFSAVDVRLPMLVYVSREKRPGYDHQKKAGALNVQLRVSALLSNAPFI
INFDCDHYINNSQAFRAAMCFMMDRRDGDNVAFVQFPQRFDDVDPTDRYANHNRMFFDATMLGMN
GIQGPSYVGTGSMFRRVALYGADPPRWRPDDVKVLENPNKFGKSMTFINSIPVAANQERSVMSPVS
LDEPATTELADVMTCAYEDGTEWGDGVGWVYDMATEDAVTGFRLHRTGWRSMYCDMEPPAFCGT
APINMTERMYQILRWSGGSLEVFFSRFCPLLAGRRLHPMQRVAYTNMTFYPLSALFVVCYHLLPLMW
VFNGRFYIQKPYPTYVMYVLVIIVSNEVIGMVEIVWAGLTLLDWFRNEQFYMICATGVYPTAVLHVVLR
SLGLKGMSFKMTAKQLATGARERFAELYNVQWAPLLIPTLVVIAVNVVAIGAAVGKAITWGWSAGQV
VEAASGLMFNVWILLMFYPFALGVIGRWGKRPYVLFAMFVAAFAAIAAVYVAVQAALAGNLLYFQLGH
WSIGGAVSLPSRRV

>CSLF_2BS_TGACv1_148027_AA0489970.1_Triticum_aestivum
MTTSPATAAGAATGLSEPLLSNGNGVHAGALVVTPVVANGHGGDKLKGDLKAKDKYWKDVDQPDD
VAAAPDLENGGGRPLLFSNRRVKNIILYPYRVLILIRVIAVILFVGWRIKHNNSDVMWFWVMSVVADVW
FSLSWLSYQLPKYNPIKMIPDLATLRKQFDTPGSSSQLPGIDVIVTTASATDEPILYTMNCVLSILAADY
HIGRCNCYLSDDSGSLVLYEALVETAKFAALWVPFCRKHQTEPRAPARYFELEGPLCGGASHKEFIQ
DYKHVRMQYEEFKKHLDMLPNTIRQRSDIYSKTGTKDEDAKVTWMADGTQWPGTWVDPAEKHRAG
HHAGIVKIVQSHPEHVVPLGVQESNDNPLNFDDVDMRLPMLVYVAREKSPGVEHNKKAGALNAELRI
SALLSNAPFFINFDCDHYINNSEALRAAVCFMLDPREGDNTGFVQFPQRFDNVDPTDRYGNHNRVFF
DGAMYGLNGQQGPTYLGTGCMFRRLALYGIDPPCWRAEDMIVDSNRFGNSLPFLNSVLAAIKQEEG
VTLPPPLDDSFLEEMTKVVSSSYDDSTDWGRGIGYLYNMATEDIVTGFRIHGQGWRSMYVTMEREA
FRGTAPINLTERLRQIVRWSGGSLEMFFSHISPLFAGRRLSLVQRLSYINFTIYPLTSLFILMYAFCPVM
WLLPTEILIQRPYTRYIVYLLIVIAMIHVIGMFEIMWAGITWLDWWRNEQFFMIGSVTAYPTAVLHMVVN
LLTKKGIHFRVTTKQPVADTDDKYAEMYEVHWVPMMVPAVVVLFSNILAIGVAIGKSVLYMGTWSAAQ
KRHGALGLLFNMWIMVLLYPFALAIIGRWAKRTGILFILLPIAFLSTALMYIGIHTFLLHFFPSMLI
>CSLF_2BS_TGACv1_147667_AA0486240.1_Triticum_aestivum
MVSPATSGGRGGNAGLADPLLATNDDSDGARHVFGAKAKYWAPADEKEMTASRECSGEDGRPLLY
RTFKVKGFLVNTYRFLNLARLTAVIVFFAWRVQHPDSDAMWLWWISVVGDFWFGLSWWLNQVPKLN
PTICIPTIPLLRQQFDLPDGGSNLPVLDVFISTVDPVEEPMLHTMNSILSILATDYPVDKYATYLSDDGG
SLLHYDGLVETAKFAALWVPFCRKHHVEPRAPESYFGMKIRPYTGNLPEEFLDDHRRLRREYEEFKT
RLDALFTVIPQRSEAHGREDAKGGGGKATWMADGTQWPGTWTEPAEGHRKGDHAGIIQVMLSQPS
SEPQLGEPASSDDGPLDFSAVDVRLPMLVYVSREKRPGYDHQKKAGALNVQLRVSALLSNAPFIINF
DCDHYINNSQAFRAAMCFMMDRRDGDNVAFVQFPQRFDDVDPTDRYANHNRMFFDATMLGMNGI
QGPSYVGTGSMFRRVALYGADPPRWRPDDVKVLENPNKFGKSMTFINSIPVAANQERSVMSPVSLD
EPATTELADVMTCAYEDGTEWGDGVGWVYDMATEDAVTGFRLHRTGWRSMYCDMEPPAFCGTAP
INMTERMYQILRWSGGSLEVFFSRFCPLLAGRRLHPMQRVAYTNMTFYPLSALFVVCYHLLPLMWVF
NGRFYIQKPYPTYVMYVLIIIISNEVIGMVEIVWAGLTLLDWFRNEQFYMICATGVYPTAVLHVVLRSLG
LKGISFKMTAKQLATGARERFAELYDVQWAPLLIPTLVVIAVNVVAIGAAVGKAITWGWSAGQVVEAA
SGLMFNVWILLMFYPFALGVIGRWGKKPYVLFAMFVAAFAAIAAVYVAVQAALAGNLPYFQLGHWSIG
GAVSLPSRRV
>GRMZM2G367267_P01
MTMTSPQALSAGAVTVEDDGGLAVPLLADGSGATKTEEECGAEAEGKYWVAADEAERQAVTDCGA
EDGRALLFRTYKLRGAILHPYRALILVRLVAVLLFFVWRIRNSASNVMWFWATSVAGDAWFGFSWLL
NQLPKFSPVKSVPDLAALRRHYDLLPADDGAASKLPGVDVFVTTADSVDEPVLYTMNSILSILATDYPA
DRLACYVSDDSGALVLYEALVEAAKFARLWVPFCRKHCVEPRAPERYFETEPQGGRASQEFVNDYK
RVQMEYDEFKVRLGNLPDTIRKRSGTGSMRASEGDAQGTWMADGMQWPGTWMDPTEKHRKGHH
AGIVKVVLDHHPSRGHHGPRAGAGAGAENKQSADDFGAAAGLRLPMLVYVSREKDPNYDHNKKAG
ALNAQLRVSALLSNAQFVINFDCDHYVNNSQALRAAVCLMLDQREGGDTAFVQFPQRFDNVDPTDR
YGNHNRVFFDGTMLALNGLQGPSYLGTGCMFRRIALYGVDPPHCRQQQLESVAPEPASKYGKSTAL
IHSVSEAMGERERLTTPPPVPPLDVEMVVAASYDGGTDWGKGVGYIYGIATEDIVTGFRIHGKGWRS
MYCTMRRDAFRGTAPINLTERLHQIVRWSGGSLEMFFSRNNPLVGGQRLKLLQRVSYLNMTVYPVT
SLFILLYALCPVMWLVPEEVHIQRPFTRYVVYLLITILMIHMIGWLEMKWSGVAWLDHWRNEQFFMIGS
TSAYPIALWHMAKKLLTRKGIHFRVTSKQTTAGTDDKFADLYEMRWTPMLVPTAFVLVANVGAVGVA
MGKALVYMGVWTVAQKTHAALGLLFNVWIMLLLYPFALAIMGRWAKRPIILLVLLPAVFAVVGTIYVAL
HFLLANVIPI
>CSLF_2DS_TGACv1_177329_AA0573830.1_Triticum_aestivum
MVSPAASGGGNAGLADPLLATNDNSEGARHVFGAKAKYWVPADEKEIAASRECGGEDGRPLLYRTF
KVKGMLVNTYRFLNLARLTAVIVFFAWRVQHPDSDAMWLWWISVVGDFWFGLSWWLNQVPKLNPTI
CIPTIPLLRQQFDLPDGGSNLPVLDVFISTVDPVEEPMLHTMNSILSILATDYPVDKYATYLSDDGGSLL
HYDGLVETAKFAALWVPFCRKHHVEPRAPESYFGVKIRPYMGNLPEEFLDDHGRLRREYEEFKTRLD
ALFTLIPQRSEAHGREDAKGGGGKATWMADGTQWPGTWTEPAEGHRKGDHAGIIQVMLSQPSSEP
QLGEPASSDHSPLDFSAVDVRLPMLVYVSREKRPGYDHQKKAGALNVQLRVSALLSNAPFIINFDCD
HYINNSQAFRAAMCFMMDRRDGDNVAFVQFPQRFDDVDPTDRYANHNRMFFDATMLGMNGIQGPS
YVGTGSMFRRVALYGADPPRWRPDDVKVLENPNKFGKSMTFINSIPVAANQERSVMSPVSLDEPAT
TELADVMTCAYEDGTEWGDGVGWVYDMATEDAVTGFRLHRTGWRSMYCDMEPPAFCGTAPINMT
ERMYQILRWSGGSLEVFFSRFCPLLAGRRLHPMQRVAYTNMTFYPLSALFVVCYHLLPLMWVFNGQ
FYIQKPYPTYVMYVLIIIVSNEVIGMVEIVWAGLTLLDWFRNEQFYMICATGVYPTAVLHVVLRSLGLKG
MSFKMTAKQLATGARERFAELYDVQWAPLLIPTLVVIAVNVVAIGVAVGKAITWGWSAGQVVEAASG
LMFNVWILLMFYPFALGVIGRWGKRPFVLFAMFVAAFAAIAAVYVAVQAALAGNLPYFQLGHRSIGGA
VSLASRRV
>CSLF_2DS_TGACv1_178985_AA0603230.1_Triticum_aestivum
MTTSPATDAGAATGLSEPLLSNRNGVHAGALVVTPVAANGHGGKAKDKYWKDVDQPGDMAVAPDL
ENGGGRPLLFSNRRVKNIILYPYRVLILIRVIAVILFVGWRIKNNNSDVMWFWVISVVADVWFSLSWLS

YQLPKYNPIKMIPDLATLRKQFDTPGRSSQLPGIDVIVTTASATDEPILYTMNCVLSILAADYHIGRCNC
YLSDDSGSLVLYEALVETAKFAALWVPFCRKHQIDPRAPESYFELEGPLCGGASHKEFIQDYKHVCT
QYEEFKKHLDMLPNTIRQRADIYSKTGTKDEDAKVTWMADGTQWPGTWLDPAEKHRAGHHAGIVKI
VQSHPEHVVPLGVHESNDSSLNFDGVDMRLPMLVYVAREKCPGVEHNKKAGALNAELRISALLSNAP
FFINFDCDHYINNSEALHAAVCFMLDPREGDNTGFVQFPQRFDNVDPTDRYGNHNRVFFDGAMYGL
NGQQGPTYLGTGCMFRRLALYGIDPPCWRAEDIIVDSNRFGNSLPFLNSVLAAIKQEEGVTLPPTLDD
SFLEEMTKVVSSSYDDSTDWGRGIGYIYNMATEDIVTGFRIHGQGWRSMYATMEREAFRGTAPINLT
ERLRQIVRWSGGSLEMFFSHISPLFAGRRLSLVQRLSYINFTIYPLTSLFILMYAFCPVMWLLPTEILVQ
RPYTRYIVYLIIVIVMIHVIGMFEIMWAGITWLDWWRNEQFFMIGSVTAYPTAVLHMVVNILTKKGIHFR
VTTKQPVADTDDKYAEMYEVHWVPMMIPAVVVLFSNILAIGVAIGKSILYMGTWSAAQKRHGALGLLF
NLWIMVLLYPFALAIIGRWAKRTGILFILLPIAFLSTSLMYIGVHTFLLHFFPSMLI
>OS07G36750_OsCslF3
MASPASVAGGGEDSNGCSSLIDPLLVSRTSSIGGAERKAAGGGGGGAKGKHWAAADKGERRAAKE
CGGEDGRRPLLFRSYRVKGSLLHPYRALIFARLIAVLLFFGWRIRHNNSDIMWFWTMSVAGDVWFGF
SWLLNQLPKFNPVKTIPDLTALRQYCDLADGSYRLPGIDVFVTTADPIDEPVLYTMNCVLSILAADYPV
DRSACYLSDDSGALILYEALVETAKFATLWVPFCRKHCIEPRSPESYFELEAPSYTGSAPEEFKNDSRI
VHLEYDEFKVRLEALPETIRKRSDVYNSMKTDQGAPNATWMANGTQWPGTWIEPIENHRKGHHAGI
VKVVLDHPIRGHNLSLKDSTGNNLNFNATDVRIPMLVYVSRGKNPNYDHNKKAGALNAQLRASALLS
NAQFIINFDCDHYINNSQAFRAAICFMLDQREGDNTAFVQFPQRFDNVDPKDRYGNHNRVFFDGTML
ALNGLQGPSYLGTGCMFRRLALYGIDPPHWRQDNITPEASKFGNSILLLESVLEALNQDRFATPSPVN
DIFVNELEMVVSASFDKETDWGKGVGYIYDIATEDIVTGFRIHGQGWRSMYCTMEHDAFCGTAPINLT
ERLHQIVRWSGGSLEMFFSHNNPLIGGRRLQPLQRVSYLNMTIYPVTSLFILLYAISPVMWLIPDEVYIQ
RPFTRYVVYLLVIILMIHMIGWLEIKWAGITWLDYWRNEQFFMIGSTSAYPTAVLHMVVNLLTKKGIHFR
VTSKQTTADTNDKFADLYEMRWVPMLIPTMVVLVANIGAIGVAIGKTAVYMGVWTIAQKRHAAMGLLF
NMWVMFLLYPFALAIMGRWAKRSIILVVLLPIIFVIVALVYVATHILLANIIPF
>CSLF_2AL_TGACv1_094713_AA0301960.1_Triticum_aestivum
MMAAAVTRRSNALRVDVPDGDAVAVSVVADSPVAKRGLGAKEDVWVAVDEGGMSGDGNRPLLFRT
MKVKGSILHPYRFLMLMRLVAVVIFFKWRMEHKNHDGVWLWTVSMTADVWFGFSWLLNQLPKLNPI
KRVPDLAALADRHDDATLPGIDVFVTTVDPVDEPVLYTVNTILSILAADYPVDNYACYLSDDGGTLVHY
EAMVQVASFAALWVPFCRKHCVEPRSPESYFGIKTHSYAGGMAGEFMRDRRRVRREYEEFKVRIDS
LSTTIRQRSDAYNSKNKGGVSATWMADGTQWPGTWVEQAENHRRGQHAGIVQVLLDHPSCEPQLG
SPASTDNPFDFSNVDTRLPMLVYISREKRPGYDNQKKAGAMNVMLRVSALLSNAPFVINFDCDHYIN
NSQALRAPMCFMLDPRDGQNTAFVQFPQRFDDVDPTDRYANHNRVFFDGTMLALNGLQGPTYLGT
GTMFRRVSLYGIEPPRYRAENTKLVRKAGEFGYSTSFVNSVPDAAIQDRSITPVLVDEGLRKDLTTLM
TCAYEDGSSWGRDAGWVYNIATEDVVTGFRIHRQGWRSMYCSMEPAAFRGTAPINLTERLYQVLR
WSGGSLEVFFSHSNALIASRRLHLLQRIAYLNMSTYPIVTVFILSYNFFPVMWLFSEQLYIQRPFGTYM
AYLVAIIAMVHLIGMFEVRWSGITLLDWFRNEQFYMIGATGVYPTAVLYMLLKLVTGKGIYFRLTSKQT
EACSNDKFADLYTVRWVPLLIPTTAVIIVNVAAVGAAIGKAATWGFFTDEARHALLGMVFNMGILVLLY
PFALGIMGKWGKRPIILFIVLVMAISAVGLLYVMLHAPYTGEWSQVAVSLGKASLTGPSGSG
>CSLF_2DL_TGACv1_160109_AA0546890.1_Triticum_aestivum
MAAAVTRRSNALRVDVPGGEAVAVSVAADSPVAKRGLGAKDDVWVAADEGGIMSGDGNRPLLFRT
MKVKGSILHPYRFLMLVRLVAVVAFFKWHVEHKNQDSVWLWTASMTADPWFGFSWLLNQLPKLNPI
KRVPDLADRHDDATLPRIDVFVTTVDPVDEPVLYTVNTILSILAADYPIDNYACYISDDGGTLVHYEAMV
QVASFAALWVPFCRKHCVEPRSPESYFGIKTRSYIGGMAGEFMRDHRRVRREYEEFKVRIDSLSTTI
RQRSDAYNSSNKGGVSATWMADGTHWPGTWVEQAENHRRGQHAGIVQVLLDHPSCKPQLGSPAS
TDNPFDFSNIDTRLPMLVYISREKRPGYDNQKKAGAMNVMLRVSVLLSNAPFVINFDCDHYINNSQAL
RAPMCFMLDPHDGQNTAFVQFPQRFDDVDPTDRYANHNRVFFDGTMLALNGLQGPTYLGTGTMFR
RVSLYGIEPPRYRAENTKLVRKTGEFGYSTSFVNSVPDAAIQDRSITPVLVDEHLRKDLATLMTCAYE
DGSSWGRDAGWVYNIATEDVVTGFRIHRQGWRSMYCSMEPAAFRGTAPINLTERLYQVLRWSGGS
LEVFFSHSNALIASRRLHPLQRIAYLNMSTHPIVTVFILSYNFFPVMWLFSEQLYIQRPFGMYMGYLVAI
IAMVHLIGMFEVRWSGITLLDWFRNEQFYMIGATGVYPTAVLYMLLKLATGKGIYFRLTSKQTEACSN
DKFADLYTVRWVPLLIPTTAVIIVNVAAVGAAIGKAATWGFFTDEARHALLGMVFNMGILVLLYPFALGI
MGKWAKRPIILFIVLVMAISVVGLLYVSLHAPYTGEWSQVAVSLGKASLTGPSGSG
>OS07G36700_OsCslF1
MSAAAAVTSWTNGCWSPAATRVNDGGKDDVWVAVDEADVSGARGSDGGGRPPLFQTYKVKGSIL
HPYRFLILARLIAIVAFFAWRIRHKNRDGAWLWTMSMVGDVWFGFSWVLNQLPKQSPIKRVPDIAALA
DRHSGDLPGVDVFVTTVDPVDEPILYTVNTILSILAADYPVDRYACYLSDDGGTLVHYEAMVEVAKFA
ELWVPFCRKHCVEPRSPENYFAMKTQAYKGGVPGELMSDHRRVRREYEEFKVRIDSLSSTIRQRSD
VYNAKHAGENATWMADGTHWPGTWFEPADNHQRGKHAGIVQVLLNHPSCKPRLGLAASAENPVDF
SGVDVRLPMLVYISREKRPGYNHQKKAGAMNVMLRVSALLSNAPFVINFDGDHYVNNSQAFRAPMC
FMLDGRGRGGENTAFVQFPQRFDDVDPTDRYANHNRVFFDGTMLSLNGLQGPSYLGTGTMFRRVA
LYGVEPPRWGAAASQIKAMDIANKFGSSTSFVGTMLDGANQERSITPLAVLDESVAGDLAALTACAY

EDGTSWGRDVGWVYNIATEDVVTGFRMHRQGWRSVYASVEPAAFRGTAPINLTERLYQILRWSGG
SLEMFFSHSNALLAGRRLHPLQRVAYLNMSTYPIVTVFIFFYNLFPVMWLISEQYYIQRPFGEYLLYLV
AVIAMIHVIGMFEVKWAGITLLDWCRNEQFYMIGSTGVYPTAVLYMALKLVTGKGIYFRLTSKQTAASS
GDKFADLYTVRWVPLLIPTIVIMVVNVAAVGVAVGKAAAWGPLTEPGWLAVLGMVFNVWILVLLYPFA
LGVMGQWGKRPAVLFVAMAMAVAAVAAMYVAFGAPYQAELSGVAASLGKVAAASLTGPSG
>OS06G39970_OsCesA
MDGESPEIMPVECPDPEPASSESGDDHDIPEPLSSRLSVPSGELNLYRAAVALRLVLLAAFFRYRVTR
PVADAHALWVTSVACELWLAASWLIAQLPKLSPANRVTYLDRLASRYEKGGEASRLAGVDVFVAAAD
AAREPPLATANTVLSVLAADYPAGGVACYVHDDGADMLVFESLFEAAGFARRWIPFCRRHGVEPRA
PELYFARGVDYLRDRAAPSFVKDRRAMKREYEEFKVRMNHLAARARKVPEEGWIMSDGTPWPGNN
SRDHPAMIQVLLGHPGDRDVDGGELPRLFYVSREKRPGFRHHGKAGAMNALLRVSAVLTNGAYVLN
LDCDHCVNNSSALREAMCFMMDPVAGNRTCFVQFALRDSGGGDSVFFDIEMKCLDGIQGPVYVGS
GCCFSRKALYGFEPAAAADDGDDMDTAADWRRMCCFGRGKRMNAMRRSMSAVPLLDSEDDSDEQ
EEEEAAGRRRRLRAYRAALERHFGQSPAFIASAFEEQGRRRGGDGGSPDATVAPARSLLKEAIHVVS
CAFEERTRWGKEIGWMYGGGVATGFRMHARGWSSAYCSPARPAFRRYARASPADVLAGASRRAV
AAMGILLSRRHSPVWAGRRLGLLQRLGYVARASYPLASLPLTVYCALPAVCLLTGKSTFPSDVSYYD
GVLLILLLFSVAASVALELRWSRVPLRAWWRDEKLWMVTATSASLAAVFQGILSACTGIDVAFSTETA
ASPPKRPAAGNDDGEEEAALASEITMRWTNLLVAPTSVVVANLAGVVAAVAYGVDHGYYQSWGALG
AKLALAGWVVAHLQGFLRGLLAPRDRAPPTIAVLWSVVFVSVASLLWVHAASFSAPTAAPTTEQPIL
>GRMZM2G339645_P01
MASPAPAGDAVYAANGGLTDPLLVSANGHGAAARKAGHGARGRYWVASDKAERRAAKESGGEDG
RALLFRKYKVKGALLHPYRLLIIIRLVAVLAFFAWRIRHNKSDIMWFWTMSIVGDVWFGFSWLLNQLPK
FNPVKTIPDLAALQRHFGYPDGGASRLPGIDVFVTTADPIDEPILYTMNCVLSILSVDYPVDRLACYLSD
DSGALVLYEALAEVGKFAPLWVPFCRKYSIEPRAPESYFEHVAPPQAGRVTQEFLNDYRRVQMEYD
EFKARLDNLPDAIRKRSDVYNSVRDAGGAQKATWMANGTQWPGTWIDPAENHRKGHHAPIAKVVLN
HPSRGQHPITESNPSIATTDERLPMLVYVSREKNPGYDHNKKAGALNAQLRASALLSNAQLIINFDCD
HYINNSQALSSAVCFMLDQRDGDNTAFVQFPQRFDNVDPTDRYGNHNRVFFDGTMLALNGLQGPS
YLGTGCMFRRLALYGIDPPHCRAENITAEASRFGNSTIFLDSVSKALKNDRTITPPPIDDTFLAELERVV
TCSYDKGTDWGKGVGYIYDIATEDIVTGFRIHGQGWRSMYCTMEHDAFCGVAPINLTERLHQIVRWS
GGSLEMFFSHNNPFIGGRRIQPLQRVSYLNMTVYPVTSVFILIYALSPVMWLIPDEVYIQRPFTRYVVY
LLVIIVMIHMIGWLEIKWAGVTWLDYWRNEQFFMIGSTSAYPMAVLHMAVNLLTKKGIHFRVTSKQTAA
DDNDKFADLYDFRWVPMLIPTMAVLICNVGAIGVALGKTVVYIGTWTAAKKMHAALGLLFNIWIMFLLY
PFALAIMGRWAKRPIILVVLLPVVFALVALLYVGIHILLAGLIPF
>CSLF_U_TGACv1_641498_AA2096480.1_Triticum_aestivum
MPSPAAVGGGRLADPLLAADVVVGAKDKYWVPADEREILASQKSGAGEDGRAPLLYRTFRVKGPLIN
LYRLLTLVRVIVVILFFTWRMRHRDSDAMWLWWISVVGDLWFGVTWLLNQITKLRPRKCVPSISVLRD
HLDQPDGGSDLPLLDVFINTVDPVDEPMLYTMNSILSILATDYPVEKYATYFSDDGGSLVHYEGLQLA
AEFAASWVPFCRKHCVEPRAPESYFWAKMRGEYAGSAPKEFLDDHRRMRAAYEEFKARLDGLSAAI
EQRSEACNRANGKDKEECANATWMADGSTQWQGTWIKPAKGHRKGHHPAILQVMLDQPSKDPEL
GMAASSDHPLDFSAVDARLPMLVYIAREKRPGYDHQKKAGAMNVQLRVSALLSNAPFIINFDGDHYV
NNSQAFRAAICFMLDPRDGADTAFVQFPQRFDDVDPTDRYCNHNRMFFDATLLGLNGIQGPSFVGT
GCMFRRVALYSADPPRWRPDDAKEAKASRYRPNMFGKSTSFINSMPAAANQERSVPSPATVGEAEL
ADAMTCAYEDGTEWGNDVGWVYNIATEDVVTGFRLHRTGWRSTYCAMEPDAFRGTAPINLTERLY
QILRWSGGSLEMFFSRFCPLLAGRRLHPMQRVAYINMTTYPVSTFFICMYYLYPVMWLFQGEFYIQR
PFQTFALFVVVIIATVELIGMVEIRWAGLTLLDWVRNEQFYIIGTTGVYPMAMLHILLRSLGIKGVSFKLT
AKKLTGGARERLAELYDVQWVPLLVPTVVVMAVNVAAIGAAAGKAIVGRWSAAQVAGAASGLVFNV
WMLLLLYPFALGIMGRWSKRPYILFIVLVTAVAATASMYVALAGSLPYLHSGIKLV
>CSLF_2BS_TGACv1_148608_AA0494060.1_Triticum_aestivum
MASAVGAGGANAGLADPLLASRDGGAKKPVGAKGKHWVAADKDQRRAAKESGGEEGRPLLFRTYK
VKGTLLHPYRALIFIRLIAVLLFFVWRIKHNKSDIMWFWTMSVVGDVWFGFSWLLNQLPKFNPVKTIPD
MVALRRQYDLPDGTSTLPGIDVFVTTADPIDEPILYTMNCVLSILASDYPVDRCACYLSDDSGALIQYE
ALIETAKFATLWVPFCRKHCIEPRAPESYFELEAPLYTGSASEEFKNDHSSVHREYDEFKEHLDSLSS
AISKRSDAYNSMKTGEGDAKATWMANGTQWPGSWIDTTEIHRKGHHAGIVKVVLDHSVRGHNLGSQ
ASTHNLNFANTDVRLPMLVYISRGKNPSYDHNKKAGALNAQLRASALLSNAQFIINFDCDHYINNSQA
LRAAMCFMLDQRQGDSTAFVQFPQRFDNVDPSDRYGNHNRVFFDGTMLALNGLQGPSYLGTGCMF
RRIALYGIDPPEWRHDNIVVDDKRFGSSIPFLDSVSKAINQERSTIPPPISETLVAEMERVVSASHDKAT
GWGKGVGYIYDIATEDIVTGFRIHGQGWRSMYCTMERDAFCGIAPINLTERLHQIVRWSGGSLEMFF
SLNNPLIGGRRIQSLQRVSYLNMTVYPVTSLFILLYALSPVMWLIPDEVYIQRPFTKYVVFLLVIILMIHVI
GWLEIKWAGVTWLDYWRNEQFFMIGSTSAYPAAVLHMVVNLLTKKGIHFRVTSKQTAADTNDKFADL
YDMRWVPMLIPTTVVLIANVGAIGVAMGKTIVYMGAWTIAQKTHAALGLLFNVWIMVLLYPFALAIMGR
WAKRPVILLVLLPVAFTIVCLVYVAVHILLLSYLTF
>CSLF_2DS_TGACv1_177641_AA0581710.2_Triticum_aestivum

MASAAGAGGANAGLADPLLASAKKPVGAKGKHWVAADKDQRRAAKESGGEDGRPLLFRTYKVKGT
LLHPYRALIFIRLIAVLLFFVWRIKHNKSDIMWFWTLSVVGDVWFGFSWLLNQLPKFNPVKTIPDMVAL
KRQYDLPDGTSTLPGIDVFVTTADPIDEPILYTMNCVLSILASDYPVDRCACYLSDDSGALIQYEALVET
AKFATLWVPFCRKHCIEPRAPESYFELEAPLYTGSAPEDFKNDHSSVHREYDEFKEHLDSISSAISKR
SDAYNSMKTEEGDAKATWMANGTQWPGSWIDTTEIHRKGHHAGIVKVVLGHSIRGHNLGSQASTNN
LNFASTDVRLPMLVYISRGKNPSYDHNKKAGALNAQLRASALLSNAQFIINFDCDHYINNSQALRAAM
CFMLDQRQGDSTAFVQFPQRFDNVDPSDRYGNHNRVFFDGTMLALNGLQGPSYLGTGCMFRRIAL
YGIDPPEWRHDNIVVDDKRFGSSIPFLESVSKAINQERSTIPPPISETLVAEMERVVSASHDKATGWG
KGVGYIYDIATEDIVTGFRIHGQGWRSMYCTMERDAFCGIAPINLTERLHQIVRWSGGSLEMFFSLNN
PLIGGRRIQALQRVSYLNMTVYPVTSLFILLYALSPVMWLIPDEVYIQRPFTKYVVFLLVIILMIHVIGWLE
IKWAGVTWLDYWRNEQFFMIGSTSAYPAAVLHMVVNLLTKKGIHFRVTSKQTAADTNDKFADLYDMR
WVPMLIPTTVVLIANVGAIGVAMGKTIVYMGAWTIAQKTHAALGLLFNVWIMVLLYPFALAIMGRWAKR
PVILLVLLPVAFTIVCLVYVAVHILLLSYLTF
>CSLF_2AS_TGACv1_113659_AA0359050.1_Triticum_aestivum
MASAAGAGGANAGLADPLLASAKKPVGAKGKHWVAADKDQRRAAKESGGEDGRPLLFRTYKVKGT
LLHPYRALIFIRLIAVLLFFVWRIKHNKSDVMWFWTMSVVGDVWFGFSWLLNQLPKFNPVKTIPDMVA
LRRQYDLPDGTSTLPGIDVFVTTADPIDEPILYTMNCVLSILASDYPVDRCACYLSDDSGALIQYEALVE
TAKFATLWVPFCRKHCIEPRAPESFFEQEAPLYTGSAPEEFKNDHNSVYIEYDEFKECLDSLSSAISKR
SDAYNSMKTEEGDANATWMANGTQWPGSWIDTTEIHRKGHHAGIVKVVLDHSIRGHNLGSQASTHN
LNFASTDVRLPMLVYISRGKNPSYDHNKKAGALNAQLRASALLSNAQFIINFDCDHYINNSQALRAAM
CFMLDQRQGDSTAFVQFPQRFDNVDPSDRYGNHNRVFFDGTMLALNGLQGPSYLGTGCMFRRIAL
YGIDPPEWRHANIVVDDKRFGSSIPFLESVSKAINQERSTIPPPISETLVAEMERVVSASHDKATGWGK
GVGYIYDIATEDIVTGFRIHGQGWRSMYCTMERDAFCGIAPINLTERLHQIVRWSGGSLEMFFSLNNP
LIGGRRIQSLQRVSYLNMTVYPVTSLFILLYALSPVMWLIPDEVYIQRPFTKYVVFLLVIILMIHVIGWLEI
KWAGVTWLDYWRNEQFFMIGSTSAYPAAVLHMVVNLLTKKGIHFRVTSKQTAADTNDKFADLYDMR
WVPMLIPTTVVLIANVGAIGVAMGKTIVYMGAWTIAQKTHAALGLLFNVWIMVLLYPFALAIMGRWAKR
PVILVVLLPVAFTIVCLVYVAVHILLLSYLTF
>CSLF_2DL_TGACv1_159781_AA0542640.1_Triticum_aestivum
MAAAVTRRVNALRVEVPDGNADTANAPAAKRILDAKDDVWVSADDGTSAGNGNQPLLFRTMKVKGS
ILHPYRFLILVRLVAVAAFFAWRLEHRNHDGTWLWATSMVADAWFGFSWLLNQLTKLNPIKRVPDLAT
LADQHGEAILPGIDVFVTTADPVDEPVLYTVNTVLSILAADYPIDKYACYLSDDGGTLVHYEAMTQVAS
FAALWAPFCRKHCVEPRSPENYFGMKAQPYAGSMPGDFTRDRRRVRREYDEFMVRIDSLSTTIRQR
SDAYNNGDGVHATRMADGAPWPGTWIEQAENHRRAQHAGIVQVILEHPGCKPQLGSSASTDNPFD
FNNVDMRLPMLVYISREKRLGYDNQKKAGAMNAMLRISALLSNAPFIINFDCDHYINNSKALRAPMCF
MLDPRDGQNTAFVQFPQRFDDVDPTDRYANHNRVFFDGTMLALNGLQGPSYLGTGTMFRRVTLYG
MEPPRYRVENIKLVDNAHEFGNSTSFTNSMPDGAIQERSITPVLVDEGLINDLATLITCAYEDGSSWG
RDIGWVYNIATEDVVTGFRIHRQGWRSMYCSMEPAAFRGTAPINLTERLYQVLRWSGGSLEVFFSHN
NALIAGRRLHPLQRIAYFNMSTYPIVTVFILAYNFFPVMWLFSEQLYIQRPFGTYIAYLVAVIAMMHVIG
MFEVKWAGITLLDWCRNEQFYLIAATGVYPTAVLYMALKLVTGKGMHFRLTSKQTEACSRDKFANLY
TVRWVPLLIPTTAVLVVNVAAVGAAIGKAAAWGFSTDQARHVLLGMVFNVGTLMLLYPFALGIMGKR
GKTPVILFVLLLMAIAAVGLLYVTLYAPYPQESLTFLSW
>GRMZM2G164761_P01
MSTTYTTKSNGTTSDDNKWPKDHVKKSASVDRPPLVRTTKLSTITIKLYRLMIILRMGIFVLFFKWRIGT
ALVMISSTGTDDKSTVLGMWMVSMAGELWFALMWVLDQVPKMQPVRRVVYLAALDEPMLPAMDVF
VTTVDTEKEPPLVTVNTILSILAADYPAEKLTCYVSDDGGALLTRDAVAEAARFSALWVPFCRKHAVEP
RNPEAYFSPGASNGFKAWRADYKGTAWPELARDRRRVRREYEELRLRIDALQAGGRAAVDAVAAD
RSCWRRGAAEDHAGAVELLVDNPGPGSTPRLGVSGTVDGVSNLLDLSSVDVRVPALVYMCREKRR
GRVNHGKAGALNALLRASAVLSNAPFILNLDCDHYVNNSQALRAGVCHMLDGEGGNGNDVAFVQFP
QRFDGVDPADRYANHNRVFFDCTELGLDGLQGPIYVGTGCVFRRSALYGVDPPLWRPQGDDAGKG
AANGIETGKLGVSTPFLRSVYAVLTNQSDQWDTVSISSPPCSFDAAAIGEATALVSCGYEDRTAWGR
DIGWIYGTVTEDVATGFCMHRRGWRSSYCATAPDAFRGTAPINLTDRLYQVLRWAAGSLEIFFSRNN
ALLAGRRLHPLQRLAYLNTTVYPFTSIFLIAYCGLFPAIPLVTGNGATTGAFFSIIIRPPSATYIAFVAALM
LTLAVVAVLEVRWSGISLGDWWRNQQFWMVSATSAYLAAAVQVALKIAAGKEISFKLTSKQRATSTV
ASVKDRFAELYAVKWTVLMVPTAVVLAVNLTSIVAAMEGGSWRDGPMAVFALAFNAYVVVHLYPFAL
GLMGRWSNTLSPLLLLIAFFTVRLLCIVLYVQML
>OS07G24190_OsCesA
MDEARQPLSRKVPIPSSQINPYRMVIIIRLVVLGFFFHYRVMHPVPDAFALWLISVICEIWFAMSWILDQ
FPKWFPIERETYLDRLTLRFDKEGQTSQLAPIDFFVSTVDPLKEPPLVTANTVLSILAVDYPVDKVSCY
VSDDGAAMLTFEALSETSEFAKKWVPFCKKYSIEPRAPEWYFQQKIDYLKDKVAPYFVRERRAMKRE
YEEFKVRINALVAKAQKVPEEGWTMQDGTPWPGNNVRDHPGMIQVFLGQSGGHDIEGNELPRLVYV
SREKRPGYNHHKKAGAMNALVRVSAVLTNAPYMLNLDCDHYINNSKAIKEAMCFMMDPLVGKKVCY
VQFPQRFDGIDRHDRYANRNVVFFDINMKGLDGIQGPIYVGTGCVFRRQALYGYDAPKTKKPPSRTC

NCWPKWCICCCCFGDRKSKKKTTKPKTEKKKRSFFKRAENQSPAYALGEIEEGAPGAENEKAGIVN
QQKLEKKFGQSSVFVASTLLENGGTLKSASPASLLKEAIHVISCGYEDKTDWGKEIGWIYGSVTEDILT
GFKMHCHGWRSIYCIPKLPAFKGSAPLNLSDRLHQVLRWALGSVEIFFSNHCPLWYGYGGGLKCLER
FSYINSIVYPFTSIPLLAYCTLPAICLLTGKFITPELTNVASLWFMSLFICIFATGILEMRWSGVGIDDWW
RNEQFWVIGGVSSHLFALFQGLLKVIAGIDTSFTVTSKGGDDEEFSELYTFKWTTLLIPPTTLLLLNFIG
VVAGVSNAINNGYESWGPLFGKLFFAFWVIVHLYPFLKGLVGRQNRTPTIVIVWSILLASIFSLLWVRID
PFLAKNDGPLLEECGLDCN
>CSLF_7AL_TGACv1_557532_AA1782680.1_Triticum_aestivum
MPLRVEALVATDTASAAAAEGRRAKDDVWVAEEGDMSGASAGRPLLFRTMKVKGSILHPYRFLILVR
LVAIVAFFAWRVEHRNHDGTWLWATSMVADAWFGFSWLLNQLPKLNPIKRVPDLVALADRHGEAILP
GIDVFVTTVDPVDEPVLYTVNTILSILAADYPVDKYACYLSDDGGTLVHYEAMLQVASFAALWVPFCR
KHCVEPRSPENYFGMKTRPYVGGMAGEFMSDHRRVRREYGEFKVRIDSLSTTIRRRSDAYNKGDD
GVHATWMADGTQWAGTWIEQADNHRRGHHAGIVQVMLDHPSCKPQLGSSVSTNSPIDLSNVDTRL
PMLVYISREKHPGYDNQKKAGAMNVMLRVSALLSNAPFVINFDCDHYINNSRALRAPMCFMLDPRDG
QNTAFVQFPQRFDNVDPTDRYSNHNRVFFDGTMLSLNGLQGPTYLGTGTMFRRVALYGMEPPRYK
AENIKLVGKAAELGNSTPFLKSIPDGAIQERSITPVLVDEALTSDLATLMTCAYEDRTSWGRDVGWVY
NIATEDVVTGFRIHRQGWRSMYCSMEPAAFRGTAPINLTERLYQVLRWSGGSLEAFFSHSNALIASR
RLHPLQRIAYLNMSIYPIATMFILAYSFFPVMWLFSEESYYIQRPFGTFIMYLVAVIAMMHVIGMFEVKW
AGITLQDWWRNEQFYMITATGVYPTAVLYMALKLIRGKGIYFRLTSKQTEACSGEKFADLYTVRWVPL
LIPTVAVLVVNIAAIGAAIGKAATWGFFTDQAWHAVLGMVFNVGTLVLLYPFALGIMGQWGKRPGILLV
MLVMAIGTVGLLYVTLQQDGHRMSFLTRPSG
>OS07G36610_OsCslF9
MSLLPPSCSAFLPSSSLYLRLAAELVLSAMLLTLVRLIAVVLFLAWRLKHRDSDAMWLWWISIAGDFW
FGVTWLLNQASKLNPVKRVPDLSLLRRRFDDGGLPGIDVFINTVDPVDEPMLYTMNSILSILATDYPAD
RHAAYLSDDGASLAHYEGLIETARFAALWVPFCRKHRVEPRAPESYFAAKAAPYAGPALPEEFFGDR
RLVRREYEEFKARLDALFTDIPQRSEASVGNANTKGAKATLMADGTPWPGTWTEPAENHKKGQHAG
IVKVMLSHPGEEPQLGMPASSGHPLDFSAVDVRLPILVYIAREKRPGYDHQKKAGAMNAQLRVSALL
SNAPFIFNFDGDHYINNSQAFRAALCFMLDCRHGDDTAFVQFPQRFDDVDPTDRYCNHNRVFFDATL
LGLNGVQGPSYVGTGCMFRRVALYGADPPRWRPEDDDAKALGCPGRYGNSMPFINTIPAAASQER
SIASPAAASLDETAAMAEVEEVMTCAYEDGTEWGDGVGWVYDIATEDVVTGFRLHRKGWRSMYCA
MEPDAFRGTAPINLTERLYQILRWSGGSLEMFFSRNCPLLAGCRLRPMQRVAYANMTAYPVSALFMV
VYDLLPVIWLSHHGEFHIQKPFSTYVAYLVAVIAMIEVIGLVEIKWAGLTLLDWWRNEQFYMIGATGVY
LAAVLHIVLKRLLGLKGVRFKLTAKQLAGGARERFAELYDVHWSPLLAPTVVVMAVNVTAIGAAAGKA
VVGGWTPAQVAGASAGLVFNVWVLVLLYPFALGIMGRWSKRPCALFALLVAACAAVAAGFVAVHAV
LAAGSAAPSWLGWSRGATAILPSSWRLKRGF
>CSLF_7DL_TGACv1_602590_AA1961740.1_Triticum_aestivum
MALRVEALVATDTAAAEGRRAKDDVWVAAEEGDMSGASAGRPLLFRTMKVKGSILHPYRFLILVRLV
AIIAFFAWRVEHRNHDGMWLWATSMVADAWFGFSWLLNQLPKLNPIKRVPDLAALADLHGEAVLPGI
DVFVTTVDPVDEPVMYTVNTILSILAADYPVDKYACYLSDDGGSLVHYEAMIQIVHFAALWVPFCRKH
CIEPRSPENYFGMKTRPYVGGMAGEFMSDHRRVRREYGEFKVIIDSLSTTIRRRSDAYNKRDDGVHA
TWMADGTQWAGTWIEQADNHRRGQHAGIVQVMLDHPSCKPQLGSSARTNNPIDLSNVDTRLPMLV
YISREKHPGYDNQKKAGAMNVMLRVSALLSNAPFVINFDCDHYINNSQALRAPMCFMLDPRDGQNT
AFVQFPQRFDDVDPTDRYSNHNRVFFDGTMLSLNGLQGPTYLGTGTMFRRVALYGMEPPCYRAENI
KLVGKAAELGNSTPFLNSIPDGAIQERSITPVLVDEGLSNDIATLMTCTYEDGSSWGRDVGWVYNIAT
EDVVTGFRIHRQGWRSMYCSMEPAAFRGTAPINLTERLYQVLRWSGGSLEVFFSHSNALIASRRLNP
LQRIAYLNMSIYPIATMFILAYSFFPVMWLFSEQSYYIQRPFGTFIMYLVVVIAMMHVIGMFEVKWAGIT
LQDWWRNEQFYMIAATGVYPTAVLYMALKLIRGKGIYFRLTSKQTEACSDEKFADLYTVRWVPLLIPT
VAVLIVNVTAVGAAIGKAATWGFFTDQAWHAVLGMVFNVGTLVLLYPFALGIMGQWGKRPGILLVML
VMAIGTVGLLYVTLQQDGHRMSFLTRPSG
>OS10G20260_OsCslF7
MPPSAGLATESLPAATCPAKKDAYAAAASPESETKLAAGDERAPLVRTTRISTTTIKLYRLTIFVRIAIFV
LFFKWRITYAARAISSTDAGGIGMSKAATFWTASIAGELWFAFMWVLDQLPKTMPVRRAVDVTALND
DTLLPAMDVFVTTADPDKEPPLATANTVLSILAAGYPAGKVTCYVSDDAGAEVTRGAVVEAARFAAL
WVPFCRKHGVEPRNPEAYFNGGEGGGGGGKARVVARGSYKGRAWPELVRDRRRVRREYEEMRL
RIDALQAADARRRRCGAADDHAGVVQVLIDSAGSAPQLGVADGSKLIDLASVDVRLPALVYVCREKR
RGRAHHRKAGAMNALLRASAVLSNAPFILNLDCDHYVNNSQALRAGICFMIERRGGGAEDAGDVAFV
QFPQRFDGVDPGDRYANHNRVFFDCTELGLDGLQGPIYVGTGCLFRRVALYGVDPPRWRSPGGGV
AADPAKFGESAPFLASVRAEQSHSRDDGDAIAEASALVSCAYEDGTAWGRDVGWVYGTVTEDVAT
GFCMHRRGWRSAYYAAAPDAFRGTAPINLADRLHQVLRWAAGSLEIFFSRNNALLAGGRRRLHPLQ
RAAYLNTTVYPFTSLFLMAYCLFPAIPLIAGGGGWNAAPTPTYVAFLAALMVTLAAVAVLETRWSGIAL
GEWWRNEQFWMVSATSAYLAAVAQVALKVATGKEISFKLTSKHLASSATPVAGKDRQYAELYAVRW

TALMAPTAAALAVNVASMAAAGGGGRWWWWDAPSAAAAAAAALPVAFNVWVVVHLYPFALGLMG
RRSKAVRPILFLFAVVAYLAVRFLCLLLQFHTA
>AT4G15320_CslB6
MADSSSSLLPLCERISHKSYILRIVDLTILVLLFSLLWYRILHMCENNTIWLVAFLCESCFSFMWLIITCIK
WSPAEDKPYPNRLDERVHDLPSVDMFVPTADPVREPPIIVVNTVLSLLAVNYPANKLACYVSDDGCS
PLTYFSLKEASKFVKIWAPFCKKYNVRVRAPFRYFLNPLVATDDSVFSKDWKMMKIYKVFYYVYFCIN
MKREYVKLCRKVEDATGDSHWLDADDDFEAFSNTKPNDHSTIVKVLLKLFLKTTVRVFVQFSKVMYIL
KLIIVVWENKGGVGDEKEVPHLVYISREKRPNYLHHYKTGAMNFLVNDFYLTHLSFFDILIYLKINVNDC
RAVSFCYYDKNMMSLIYNFKQLRVSGLMTNAPYMLNVDCDMYANEPDVVRQAMCVFLQNSKNSNH
CAFVQFPQNFYDSYTNELVVLQHYMKRGVAGIQGPIYIGSGCFHTRRVMYGLSSDDLEDDGSLSSVA
SREFLSEDSLVRKYGSSKELVKSVVDALQRKSNPQKSLANLVEAAQEVGHCHYEYQTSWGNLGWLY
DSVAEDTNTSIGIHLRGWTSSFISPDPPAFLGSTPSVGPEAIVQHRRWATGSIEVLFNKQSPLIGFRRKI
KFRQRLAYFWVLMCIRSIPELVYCLLPAYCLLNNSALFPKGPCLGIIVTLVGMHCLYTLWQFMILGFSV
KSCWLFSIQDIILKLLGISKIGFIVAKKNMPETRSGYESKSKPSQGEDDGLKLELAGFLVRLQRSSYSH
GGGGGSALAETCGCAMIVFGENDGSQEGLELPKLNSKRGISSIVEKTFVACILPFVRNHKVNEKKNHH
INIDRPNLL
>CSLF_5BL_TGACv1_409916_AA1366600.2_Triticum_aestivum
MSMTYISKKHDYAATLDEKEQPKDQKSASVESLLVRTTKLTTVTIKLYRIMVFVRMAIFVLFFKWRISTA
LAMISDGATTVRAMWTMPIAGELWFALMWVLDQLPKMQPVRRTVFATALEEPLLPTMDVFVTTADPE
KEPPLVTVNTILSILAADYPPDKLTCYVSDDGGALLTREAVAHAARFARLWVPFCRKHGVEPRNPEAY
FCPGVKARVVSRAAYMGRSWPELARDRRRVRREYEELRLRIDALHAGDVRRQPWSRGTPEYHAGV
VEVLVGPPGSTPELGVSDLLDLTSVDVRVPAVVYMCREKRHGRVHHRKAGAMNALLRTSAVLSNAP
FILNLDCDHYVSNSQALRAGVCLMLDRGGSNVAFVQFPQRFDGVDPADRYANHNRVFFDCTELGLD
GLQGPIYVGTGCMFRRAALYNADPPLWRPHGGDRDAGKDVATEADKFGISTPFLVSVRAALNLNRS
EQWNTTSPPRSFDGAAVGEATALVSCGYEDRTAWGRDIGWIYGTVTEDVATGFCMHRRGWRSAYC
ATAPDAFRGTAPINLTDRLHQVLRWAAGSLEIFFSRNNALLAGARLHPLQRLAYLNTTVYPFTSIFLLLY
CLLPAIPLVTRSASTSAFSVNTPPSATYIGFVAALMLTLAMVAALEVRWSGITLGEWWRNEQFWMVS
ATSAYAAAVVQVALKVAAGKEIAFKLTSKQRAPSAGGGVVKRQVRGAVRREMDGADGSDGGGADG
ERGVHGSSGTRGTVEERPRGGARDGVQRVGGGASPPVRPWSHGPLEQDVEPPALARRSVHSSIT
MFCPPFAYALIWLLFG
>GRMZM2G454081_P04
MVQWWGREASGGRGTSVVVKMESPEWTIPAESEVAAGGKAGRGKNARQITWVLLLKAHRTAGKLT
GAASAVLSIAGAAWRRVAAGRTDEDEDEDAGAAPPGESPALRSRFYGFLRASLVLSVLLLAADVAAH
LQGWHLAVDVPDLLAVEGLFAAGYASWARVRLEYLAPALQFLANACVVLFLVQSADRLILCLGCLWIK
HRGIRPVPKAGGKGSDDVEAGTGNFPMVLVQIPMRNEKEVYQQSIGAVCGLDWPRPNFLVQVLDDS
DDAATSTLIKEEVEKWQREGVRILYRHRLIRDGYKAGNLKSAMNCSYVKDYEFVVIFDADFQPQPDFL
KRTVPHFKGNEDVGLVQARWSFVNKDENLLTRLQNINLCFHFEVEQQVNGIFLNFFGFNGTAGVWRI
KALEDSGGWMERTTVEDMDIAVRAHLKGWKFLFLNDVECQCELPESYEAYRKQQHRWHSGPMQS
DSGRKFNLIFLFFLLRKLILPFYSFTLFCIILPMTMFIPEAELPAWVVCYIPATMSLLNILPAPKSFPFIVPY
LLFENTMSVTKFNAMVSGLFQLGSAHEWVVTKKSGRSSEGDLVVEKQPRQQRAGSASNLGSLAKEP
SSSLRKDSQRKKHNRIYRKELALSFLLLTAAARSLISVQGIHFYFLLFQGDCAKGNPEKGKKMFCMSA
SVECLRALFVNARAQITCGSRTVIFQPTCPAVMVERDRVSGELGKGFRNNPSPFTNGASNSERRLCL
ENSYSRSCSQGGDRCVNRCRYGCCFRSANIFIRRLGLHLVSLPSLPVFQSPCLLSLAGSVRMQSDLR
TV
>CSLF_5DL_TGACv1_433902_AA1424880.1_Triticum_aestivum
MSMTYITKKHDYAATLDEKEPSEDQKSASVKNLLVRTTKLTTVTIKLYRLMVFVRLTIFVLFFKWRVST
ALTVISDGTTTARAMWTMSIAGELWFALMWVLDQLPKMQPVRRTVYVTALEEPRLPTMDVFVTTTDP
EKEPPLVTVNTILSILAADYPPDKLTCYVSDDGGALLTREAVAHAACFARLWVPFCRKHGVEPRNPEA
YFCPGVKARVVSRADYMGRSWPELARDRRRVRREYEELRLRIDALHAGDVRPQQWSRGTAENHAG
VVEVLVGPPGSTPELGVSDLLDLSSVDVRVPAVVYMCREKRHGRVHHRKAGAMNALLRTSAVLSNA
PFILNLDCDHYVNNSQALRAGVCLMLDRGGSNVAFVQFPQRFDGVDPADRYANHNRVFFDCTELGL
DGLQGPIYVGTGCMFRRAALYNADPPLWRPHGGDRDAGKDVATEADKFGISTPFLGSVRAALGLNR
SEQWNTTTKPPRSFDGAAVGEATALVSCGYEDRTAWGRDIGWIYGTVTEDVATGFCMHRRGWRSA
YCATAPDAFRGTAPINLTDRLHQVLRWAAGSLEIFFSRNNALLAGARLHPLQRLAYLNTTVYPFTSIFL
LLYCLLPAIPLVTRSASASAFSVTMPPSGTYMGFVAALMLTLAMVAVLEVRWSGITLGEWWRNEQFW
MVSATSAYAAAVVQVALKVSAGKEIAFKLTSKQRASSPGGGVKERFAELYAVRWTVLMVPTAVVLAV
NVMSMAAAVQEGRWRKGPAAVLAMAFNAWVVVHLHPFALGLMGRWSKTLSPLLLLVVGFTVLSLCF
VLHLHML
>CSLF_5AL_TGACv1_374191_AA1193100.1_Triticum_aestivum
MSMTYITKKHDYVASLDGKESPEHEKSASVERLLVRTTKLTTVTIKLYRLVVFVRMIIFVLFFKWRSSTA
LAMISDGTTTVRAMWTMSIAGELWFALMWVLDQLPKMQPVRRTVYATALEESLLPAMDVFVTTADPE
KEPPLVTVNTILSILAADYPPDKLTCYVSDDGGALLTREAVAQAAWFARLWVPFCRKHGVEPRNPEA

YFCPGVKARVVSRADYRAKSWPELARDRRRVRREYEELRLRIDALHAGDVRRQQWSRGTAEDHAG
VVEVLVGPPGSTPELGVSDLLDLGSVDVRVPAVVYMCREKRHGRVHHRKAGAMNALLRTSAVLSNA
PFILNLDCDHYVNNSQALRAGVCLMLDRGGSNVAFVQFPQRFDGVDPADRYANHNRVFFDCTELGL
DGLQGPIYVGTGCMFRRAALYNADPPLWRPHGDRDAGKDVAAEADKFGISTPFLGSVRAALNLNQS
EQWNTTSPPRSFDGAAVGEATALVSCGYEDRTAWGRDIGWIYGTVTEDVATGFCMHRRGWSSAYC
ATAPDAFRGTAPINLTDRLHQVLRWAAGSLEIFFSRNNALLAGPRLHPLQRLAYLNTTVYPFTSIFLLLY
CLLPAIPLVTRNASTSAFSVNTPPSATYIAFVAALMLTLAMVAVLEVRWSGITLGDWWRNEQFWMVS
ATSAYAAAVVQVALKVAAGKEIAFKLTSKHRASNSGGGVVKDRFAELYAVRWTVLMVPTAVVLAVNV
TSMAAAVQEGRWRKGPAAVLAMAFNAWVVVHLYPFALGLMGRWSKTLSPLLLLVVVFTVLSLCFVL
HLHML
>CSLH_2BL_TGACv1_129372_AA0380770.1_Triticum_aestivum
MHRGEDSLSGLYKCTLAFVACGCGWSCGVVLLASLLLLVASYLSATAMAGGKKLQERVALGRSAWM
LADFVILFLVLALVARRAASLGERGGTWLAALVCEAWFAFVWILNMNGKWSPVRFDTYPENLSHRME
ELPAVDMFVTTADPALEPPLITVNTVLSLLALDYPHVGKLACYVSDDGCSPLTCYSLREAAKFASLWV
PFCKRHDVGVRAPFMYFSSAPEVDTGTVDHEFLESWALMKSEYEKLASRIENADEVSILRDGGDEFA
EFIDAERGNHPTIVKVLWDNSKNKTGEGFPHLVYLSREKSPRHRHNFKAGAMNVLTRVSAVMTNAPI
MLNVDCDMFANNPQVALHAMCLLLGFDDEIHSGFVQAPQKFYGGLKDDPFGNQMQVITKKIGGGLA
GIQGTFYGGTGCFHRRKVIYGMPPPDTVKHETRGSPSYKELQAKFGSSKELIESSRNIISGDLLARPT
VDISSRVEMAKQVGDCNYEAGTCWGQEIGWVYGSMTEDILTGQRIQAAGWESALLDTDPPAFLGCA
PTGGPASLTQFKRWATGLLEILISRNSPILGTIFRRLQLRQCLAYLIVNAWPMRAPFEMCYALLGPFCL
LTNQSFLPTTSNEGFRIPAALFLSYHVYHLMEYKECGLSVRAWWNNHRMQRITSASAWLLAFLTVILK
TLGLSETVFEVTRKESSTSSDGGTGTDEADTGLFTFDSAPVFIPVTALSMLNIVALAVAAWRAVVGTA
AGVHGGPGVGEFVCCGWMVLCFWPFMRGLVSSGKYGIPWSVRVKAGLIVAAFVHLCTRN
>CSLF_2DS_TGACv1_179076_AA0604160.1_Triticum_aestivum
MAAAVTRRANALRAEAPDGNAESGRASLAADSPAAKRAVDAKDDVWVAADEGDTSGAIAGDGNRP
PLFRTFKVKGSILHPYRFMILVRLVAIVAFFAWRVKHKNHDGVWLWATSMVADVWFGFSWLLNQLPK
LNPVKRVPDLAALADHSGDANLPGIDIFVTTVDPVDEPLLYTVNTILSILATDYPVDKYACYLSDDGGTL
VHYEAMIEVANFAVLWVPFCRKYCVEPRSPENYFGMKTQPYAGSMAGEFMRDHRRVRREYDEFKV
RVDSLSTTIRQRSDAYNSSKKGDGVRATWMADGTQWPGTWIEQVENHRRGQHAGIVQVILGHPSC
KPQLGSPASADNPLDFSNVDTRLPMLVYMSREKRPGYNHQKKAGAMNVMLRVSAMLSNAPFVVNF
DGDHYINNSQALRAPMCFMLDPRDGQNTAFVQFPQRFDDVDPTDRYANHNRVFFDGTMLSLNGLQ
GPSYLGTGTMFRRVALYGMEPPRYRAENIKLAGKVNEFGSSTSFINSMPDGAIQERSITPVLVDEALS
NDLATLMTCAYEDGSSWGRDVGWVYNIATEDVVTGFRMHRQGWRSMYCSMEPAAFRGTAPINLTE
RLYQVLRWSGGSLEMFFSHSNALMAGRRLHPLQRVAYLNMSTYPIVTVFILAYNLFPVLWLFSEQFYI
QRPFGTYIMYLVAVIAMIHVIGMFEVKWAGITLLDWCRNEQFYMIGATGVYPTAVLYMALKLVTGKGIY
FRLTSKQTDACSNDKXXXXXXXXXXXXXXXXXXXXXXXXGCRCCSRPSWCSS
>AT4G24010_CslG3
METHRKNSVVGNILHTCHPCRRTIPYRIYAIFHTCGIIALMYHHVHSLVTANNTLITCLLLLSDIVLAFMW
ATTTSLRLNPVHRTECPEKYAAKPEDFPKLDVFICTADPYKEPPMMVVNTALSVMAYEYPSDKISVYV
SDDGGSSLTFFALIEAAKFSKQWLPFCKKNNVQDRSPEVYFSSESHSRSDEAENLKTNILKCEVEQM
MYEDMKSRVEHVVESGKVETAFITCDQFRGVFDLWTDKFSRHDHPTIIQVLQNSETDMDNTRKYIMP
NLIYVSREKSKVSPHHFKAGALNTLLRVSGVMTNSPIILTLDCDMYSNDPATLVRALCYLTDPEIKSGL
GYVQFPQKFLGISKNDIYACENKRLFIINMVGFDGLMGPTHVGTGCFFNRRAFYGPPYMLILPEINELK
PYRIADKSIKAQDVLSLAHNVAGCIYEYNTNWGSKIGFRYGSLVEDYYTGFMLHCEGWRSVFCNPKK
AAFYGDSPKCLVDLVGQQIRWAVGLFEMSFSKYSPITYGIKSLDLLMGLGYCNSPFKPFWSIPLTVYG
LLPQLALISGVSVFPKASDPWFWLYIILFFGAYAQDLSDFLLEGGTYRKWWNDQRMLMIKGLSSFFFG
FIEFILKTLNLSTPKFNVTSKANDDDEQRKRYEQEIFDFGTSSSMFLPLTTVAIVNLLAFVWGLYGILFC
GGELYLELMLVSFAVVNCLPIYGAMVLRKDDGKLSKRTCFLAGNLHVGSYCVKLLRPQVTSPLRLIHN
NNTSGWFKRKKHNMNESV
>CSLJ_3AS_TGACv1_210908_AA0681280.2_Triticum_aestivum
MAARPSQDAPLQLHTVQTDQPLATVNRLLAALHVALAAAAIAHRGAHVMLAPDLVLLFLWALSQAPM
WRPVSRAAFPSRLSRPALPAVDVMVVTADPDKEPAAKVMNTVVSAMALDYPGGRLSVYLSDDAGSP
RTLLAARKAYALARAWVPFCRKYGVRCPCPDRFFAGDDQLDLGDHHRQELDDDRLRIKNVYETFKE
GVEEVMNDATLSQSWTKADHHAHVEIITDEQGQDSSHSNSGDGDGDEDAMPLLVYVSREKRRSST
HHFKAGALNVLLRVSSLMSNSPYVMVLDCDMYCNSRSSLLEAMCFHLDGRRRADLAFVQFPQMFHN
LSSSDIYANELRPIFWVRKKTNRPCIASVIFSEFSSNLGACMVQTRWKGLDGLRGPILSGTGFCVRRD
AVYGAGPGSSQEQFSGVEVGELKRRFGVSNGHIASLRRSGTGSTIVAAGDVLPQDAELVASCDYET
GTEWGEDVGFLYQSVVEDYFTGYRQLYCRGWTSVYCFPATGSRPPFLGSVPTNLNDALVQNKRWM
SGLLAVGLSRHCPLASAAAISVPQSMGFAYYAFTPLYAFPLLCYATVPQLCFLRGATSFPEAASTLWF
AAVFASSSLQHLVEVSVAKRGLAARTWWNEQRFWALNAVTGQIFACLGVALSLVGAGGRAVDFDLT
SKASGDRLYRDGVFDFAGCSALLLPATTLCLLNAAALVGGVWKMVGRGGNVSGTGELFLLCYVAAL
SYPLLQGMFLRRDPARVPARITAVSVAIVATLLSLFG

>GRMZM2G142685_P01
MAPPPSSWWGSEEQRGTPVVVKMDNPYSLVEIDGPGMPPSDKARGKNAKQFTWVLLLRAHRAVG
CVAWLAGGFWGVLGAVNRRVRRSRDADDEPDAEASGRGRVMLRFLRAFLLLSLAMLAFETVAHLK
GWQFPQHLMPGNLQELEEQLQHLPEHLRHLPENLRQLPDHLRVPERQEIQGWLHRAYVAWLEFRV
DYIAWAIQKLSTFCILLFMVQSVDRIVQCLACFWIKIRGIKPRIPASAGGKPRGGTTGRKRVDDVENGD
ADDDRYFPMVLVQMPMCNEKEVYETSISHVCQMDWPRDRLLIQVLDDSDDEVCQMLIKAEVTKWSQ
RGVNVIYRHRLSRTGYKAGNLKSAMACDYVKDYEFVAIFDADFQPNPDFLKLTVPHFKENPELGLVQ
ARWSFVNKDENLLTRLQNINLCFHFEVEQQVNGVYLNFFGFNGTAGVWRIKALEDSGGWMERTTVE
DMDIAVRAHLNGWKFIFLNDVKVLCELPESYEAYRKQQHRWHSGPMQLFRLCIPTVFRSKIPFWKKA
NLVMLFFLLRKLVLPFYSFTLFCVILPLTMFVPEAELPIWVICYIPVLMSILNILPAPKSFPFIIPYLLFENT
MSVTKFNAMVSGLFQLGSSYEWIVTKKAGRTSSASDILALAEADAHAHAPAPAAKLVRGVSEGGLQE
WGRLREQEAAEWANKEDAAAALAAAAAPATPKKSSNKAKKPPNRIFKKELALACLLLTAATRSLLSKQ
GLHFYFLLFQGVTFLAVGLDLIGEQVS
>OS04G35020_OsCslH2
MAVVAAAAATGSTTRSGGGGGEGTRSGRKKPPPPPLQERVPLGRRAAWAWRLAGLAVLLLLLALLA
LRLLRHHGGAGGDGGVWRVALVCEAWFAALCALNVSAKWSPVRFVTRPENLVAEGRTPSTTAAEY
GELPAVDMLVTTADPALEPPLVTVNTVLSLLALDYPRAGERLACYVSDDGCSPLTCHALREAAGFAAA
WVPFCRRYGVAVRAPFRYFSSSSSPESGGPADRKFLDDWTFMKDEYDKLVRRIKNTDERSLLRHGG
GEFFAEFLNVERRNHPTIVKTRVSAVMTNAPIMLNMDCDMFVNNPQAVLHAMCLLLGFDDEASSGFV
QAPQRFYDALKDDPFGNQMECFFKRFISGVQGVQGAFYAGTGCFHRRKAVYGVPPNFNGAEREDTI
GSSSYKELHTRFGNSEELNESARNIIWDLSSKPMVDISSRIEVAKAVSACNYDIGTCWGQEVGWVYG
SLTEDILTGQRIHAMGWRSVLMVTEPPAFMGSAPIGGPACLTQFKRWATGQSEIIISRNNPILATMFKR
LKFRQCLAYLIVLGWPLRAPFELCYGLLGPYCILTNQSFLPKASEDGFSVPLALFISYNTYNFMEYMAC
GLSARAWWNNHRMQRIISVSAWTLAFLTVLLKSLGLSETVFEVTGKDKSMSDDDDNTDGADPGRFT
FDSLPVFIPVTALAMLNIVAVTVGACRVAFGTAEGVPCAPGIGEFMCCGWLVLCFFPFVRGIVWGKGS
YGIPWSVKLKASLLVAMFVTFCKRN
>GRMZM2G122431_P01
MEDGAEKATTPDSSSSQAAPPALSAVYVNRRLVAGNRAMAAVHVALVATVIGQRALALLSAGTSSAS
PASRSLQVQHVAMALADLTLLFVWALSQSGLWRPVTRAAFPGRLLAAVPRGALPRVDVLVVTADPD
KEPPLGVVNTVVSAMALDYPGGKLSVYLSDDAGSPLTLLAARKAYAFAARAWVPFCRRHSVQCPWP
DRYFAGDDDDADGDRRREELAEERARVRKLYEKLKADIEAAKNDETISGSWSKDKRQNHDAYVEIIE
DGDGVEEIPALVYVAREKRRAWPHHFKAGALNALLRVSGVVSNAPYVLVLDCDMACNSRASALDAM
CFLLDRRPPPDSLAFVQFPQLFHNLSHKDIYANELRYIFGTRWFGLDGVRGPPLSGSGFYVRRDALY
GATPTADFMPDATAVAELKTRFGHSDRLVASLRSPGVPPEAEAMMSLAALASCAYEAGTAWGAGVG
FMYQSVVEDYFTGFQRFFARGWTSAYCYPEPRPAFLGSVPTNLNDVLVQNKRWMSGMLAVGVSRR
HSPLACRPLLRASLLQAMAYAYFGFAALCAVPVLCYATLPQLCLLRGVPLFPCPAATAAAFASSLLQH
MAEVCVSRRGRLDLRTWWNEQRFWVLNALTAQLFGCVSAAQELLGARALDFDLTSKAAVDGSLYQ
DGVFDFTGCSALLLPATTLSVLNAAAIVAGTWKMSSSSSSSSGGFHFAPQLFLMCYGAALSYPLLEG
MFLRRDPARVPPRITALSVALAAVLLAAMLG
>CSLC_3DS_TGACv1_271926_AA0910940.1_Triticum_aestivum
MASSWWGDKEEHGTPVVVKMDNPYSLVEIDGPGMDSSEKARRSKNAKQFKWVLLLRAHRAVGCVA
WLAGGFWGLLGAVNRRVRRSRDADAEPDAEASGRGRHMLGFLRAFLLLSLAMLAFETAAYLKGWH
YFPRDLPEHYLRQLPEHLQNLPEHLRHLPENLRHLPENLRHLPDGLRMPEQQEIQGWLHRAYVAWL
AFRIDYIAWAIEKLSGFCIVLFMVQSIDRILLCLGCFWIKLRGIKPGLKAAANKRGSKYADDDDLEDGDD
LGAYFPMVLLQMPMCNEKEVYETSISHVCQIDWPRDRMLVQVLDDSDDETCQMLIRAEVTKWNQRG
VNIIYRHRLSRTGYKAGNLKSAMSCEYVKDYEFVAIFDADFQPNPDFLKLTVPHFKGNPELGLVQARW
SFVNKDENLLTRLQNINLCFHFEVEQQVNGIYLNFFGFNGTAGVWRIEALEDSGGWMERTTVEDMDI
AVRAHLQGWKFIYLNDVKVLCELPESYQAYRKQQHRWHSGPMQLFRLCLPAIIKSKIPLWKKANLVM
LFFLLRKLILPFYSFTLFCVILPLTMFVPEAELPIWVICYVPMIMSVLNILPAPKSFPFVIPYLLFENTMSVT
KFNAMVSGLFQLGSSYEWVVTKKAGRTSSESDIFAMAEETDTATRPAPRLVRGVSEAGLEAWAKTH
QLDNKDLQLKAQAEEVTSLAAAIKKTSKAKPPNRIFKKELALAFLLLIAATRSLLSAQGLHFYFLLFQGV
TFLVVGLDLIGEQVS
>GRMZM2G135286_P01
MAPPSSWWGGEDQDGTPVVVKMDNPYSLVEIDGLGVPPADKARGKNAKQFTWVLLLRAHRAVGCV
AWLAGGFWGVLGAVNRRVRRSRDADDEPDAEASGRGRAMLRFLRAFLLLSLAMLAFETVAHLKGW
QFPQHLPGNLQELEEQLQHLPEHLRHLPENLRQLPDHLRMPERQEIQGWLHRAYVAWLEFRVDYIA
WAIQKLSSFCILLFMVQSVDRIVQCLACFWIKIRGIKPRVPASGGKPRGTTGRKSADAENGFADGDAD
GYFPMVLVQMPMCNEKEVYETSISHVCQIDWPRDRLLIQVLDDSDDEVCRMLIKAEVTKWSQRGVN
VIYRHRLSRTGYKAGNLKSAMACDYVKDYEFVAIFDADFQPNPDFLKLTVPHFKENPELGLVQARWS
FVNKDENLLTRLQNINLCFHFEVEQQVNGVYLNFFGFNGTAGVWRIKALEDSGGWMERTTVEDMDIA
VRAHLNGWKFIFLNDVKVLCELPESYQAYRKQQHRWHSGPMQLFRLCIPAVFRSKIPFWKKANLVML
FFLLRKLVLPFYSFTLFCVILPLTMFVPEAELPIWVICYIPVLMSLLNILPAPKSFPFVIPYLLFENTMSVTK

FNAMVSGLFQLGSSYEWIVTKKAGRTSSASDILALAEADSHAPLPPPAKLVRRVSEGGLQEWSRLRE
QETAEWANKEEAAAALAAAAPATPKMTRKTKKPNRIFKKELALACLLLTAATRSLLSKQGLHFYFLLFQ
GVTFLAVGLDLIGEQVS
>AT4G15290_CslB5
MADSSSSLHPLCERISHKSYVLRAVDLTILGLLYSLLLYRILHISENDNVWLLAFFCESCFSLVWLIFTCL
KWSPAEDIPYINTLNERVHDLPSLDMFVPTADTVRESPIITVNTVLSLLAVNYPANKLACYVSDDGCSP
LTYFSLKEASKFVKIWAPFCKKYNVRVRAPFRYFLNPLVATDDSVFSKDWKMMKREYVKLCRKVEDA
TGDSHWLDADDDFEAFSNTKPNDHSTIVKVVWENKGGVGDEKEVPHLVYISREKRPNYLHHYKTGA
MNFLLRVSGLMTNAPYTLNVDCDMYANEPDVVRQAMCVFLQNSKNSNHCAFVQFPQKFYDSYTNEL
AVLQSILGRGVAGIQGPFYIGTGCFHTRRVMYGLSSDDLEDNGNISQVATREFLAEDSLVRKYGNSKE
LVKSVVDALQRKSNPQKSLANLIEAAQEVGHCHYEYQTSWGNLGWMYDSVAEDINTSVGIHLRGWT
SSFISPDPPAFIGSTPTLGLEAIVQQRRWATGAIEVLFNKQSPFMGMFHGKIKFRQRLAYFWALMCLR
SIPELIYCLLPAYCLLHDSALFPKGPCLCTIVTLVGMHCLYSLWQFMSLGFSVQSWYVVQSLWRIIATS
SWLFSIQDIILKLLGISQIGFVIAKKTIPETKSVYESKPSQGEDDVPKLNLGKFEFDSSGLFIPGTFIMLVN
LAALAGYLVRLQRSSCSHGGGGSGLAEACGCILVVMLFLPFLKGLFEHGKYSIPLSTLSKAAFLTVLFV
FFCVGK
>AT2G32620_CslB4
MADSSCSLPPLYENISYKSYILRAVDLTILGLLFSLLLHRILYMSQNGIIWLVAFLCESCFSFVWLLSTCT
KWSPAETKPYPDRLDERVYDLPSVDMFVPTADPVREPPIMVVNTVLSLLAVNYPANKLACYVSDDGC
SPLTYFSLKEASKFAKIWVPFCKKYNLKVRAPFRYFLNPFAATEGSEFSRDWEMTKREYEKLCRKVE
DATGDSHLLGTDNELEAFSNTKPNDHSTIIKVVWENKGGVGDEKEVPHIVYISREKRPNYLHHYKAGA
MNFLARVSGLMTNAPYMLNVDCDMYANEADVVRQAMCIFLQKSQNQNHCAFVQFPQEFYDSNTIKL
TVIKSYMGRGIAGIQGPINVGSGCFHSRRVMYGLSPDELEDNGSLSSVATRELLAEDSLSSGFGNSKE
MVTSVVEALQRKPNPQNILTNSIEAAQEVGHCDYESQTSWGKTIGWLYDSMSEDMNTSIGIHSRGWT
SSYIAPDPPAFLGSMPPGGLEAMIQQRRWATGSIEVLFNKQSPLLGLFCRKLRFRQRVAYLCVSICVR
SIPELIYCLLPAYCLLHNSALFPKGLCLGITMLLAGMHCLYTLWEFMCLGHSIQSWYVSQSFWRIVATS
SWLFSIFDIILKLLGLSKNVFLVSKKTMPVETMSGSGIGPSQREDDGPNSGKTEFDGSLYFLPGTFIVL
VNLAALVGVFVGLQRSSYSHGGGGSGLGEACACILVVMLFFPFLKGLFAKGKYGIPLSTLSKAGFLAV
SFVVFSVGN
>AT2G32610_CslB3
MADSSFSLPPLCERISYTNYFLRAVYLTVLGLFFSLLLHRIRHTSEYDNVWLVAFFCESCFFLVCLLITC
LKWSPADTKPFPDRLDERVHDLPSVDMFVPTADPVREPPIMVVDTVLSLLAVNYPANKLACYVSDDG
CSPLTYFSLKEASKFAKIWVPFCKKYNTRVRAPSRYFLKPISVATEDYEFNRDWEKTKREYEKLRRKV
EDATGDSHMLDVEDDFEAFSNTKPNDHSTLVKVVWENKGGVGDEKEIPHIIYISREKRPNYVHNQKC
GAMNFLARVSGLMTNAPYILNVDCDMYANDADVVRQAMCILLQESLNMKHCAFVQFRQEFYDSSTE
LIVVLQSHLGRGIAGIQGPIYIGSGCVHTRRVMYGLSPDDFEVDGSLSSVATREFLVKDSLARRFGNS
KEMMKSVVDAIQRNPNPQNILTNSIEAAREVGHCQYEYQTSWGNTIGWLYDSVAEDLNTSIGIHSRG
WTSSYISPDTPAFLGSMPAGVPEALLQQRRWATGWIEILFNKQSPLRGLFSKKIRFRQRLAYLCIITCL
RSIPELIYCLLPAYCLLHNSTLFPKGLYLGITVTLVGIHCLYTLWEFMSLGYSVQSWLVSQSVWRIVATS
SWLFSIFDITLKLLGISETVFIITKKTVAGTKSALGSGPSQGEDVGPNSDLFKFEFDGSLCFLPGTFIVLV
NIAALAVFSVGLQRSSYSHEGGGSGLAEACGCVLVMMLFLPFLMGLFKKGKYGTPLSTLSIAGFLAVL
FVVFSV
>CSLE_5DL_TGACv1_433536_AA1415840.1_Triticum_aestivum
MVAIGRRTGQQHGHWRLAAESPPYLGPRDGEEHEAVRDGDSRGPGGVQAPRRHGGRRILLLLYYR
ATRVPAAGEGRAAWLGMLAAELWYAAYWVVTQSVRWSPVRRRPFIDRLAARHGETLPCVDIFVCTA
DPYSEPPSLVVSTILSLMAYNYPPEKLSVYLSDDGGSILTFYGMWEASLFAKHWLPFCKRYNIEPRSP
AAYFSESDGHQELCNPKEWSLIKDMFDKMTERIDTVVMSGKVPEEIKASHKGFYEWNQEITSKNHQP
IVQILIDSKDQNAVDNEGKVLPTLVYMAREKRPQHHHNFKAGAMNALIRVSSVISNSPIIMNVDCDMYS
NNNDAVRDALCFFLDEEMGHKIGFVQYPQNYYNLSKNNIYGNSLHVINEVEMGGMDSLGGPMYIGT
GCFHRREILCGRKFTEDYQEDWNAGIKDKLQESIDETEEKAKSLAACTYEHGTQWADEIGVKYGCVV
EDVNTGLAIHCRGWDSVYNNPKRPAFMGVGPTTLAQTILQHKRWSEGIFSIFLSKYNVFLFAHGKTKL
RHQMGYHIYGLWAPNSLATLYYVIIPSLALLKGTSLFPEITSPWIAPFVYVFCVKNMYSLYEALLSGDTL
KGWWNGQRMWLVKRITSYLFGVLDNLRKLLGLSKINFVVTPKVSDEDESKRYEQEIMEFGSSDPEYV
IIAAIALLNLVCLMGGLSKVMKGGWNVHLDALFPQLILCGMLVITSIPFYEAMFLRKDKGRIPFPVTLASI
GFVMLALLAKIV
>AT2G32540_CslB2
MAESSSPLPPLCERISHKSYFLRAVDLTILGLLLSLLLYRILHVNQKDTVWIVAFLCETCFTFVWLLITNIK
WSPADYKTYPERLDERVHELPPVDMFVTTADPVREPPLIVVNTVLSLLAVNYPANKLACYVSDDGCS
PLTYFSLKEASKFAKIWVPFCKKYNVRVRAPFMYFRNSPEAAEGSEFSKDWEMTKREYEKLSQKVED
ATGSSHWLDAEDDFEAFLNTKSNDHSTIVKVVWENKGGVGDEKEVPHVVYISREKRPNHFHHYKAG
AMNFLVRVSGLMTNAPYMLNVDCDMYVNEADVVRQAMCIFLQKSMDSNHCAFVQYPQDFYDSNVG
ELTVLQLYLGRGIAGIQGPQYAGSGCFHTRRVMYGLSLDDLGDDGSLSSIATRKYLAEESLTREFGNS

KEMVKSVVDALQRKPFPQKNLKDSLETAQEMGHCHYEYQTSWGKNIGWLYDSTTEDVNTSIGIHSR
GWTSSYIFPDPPAFLGCMPQGGPEVMVQQRRWATGLLEILFNKQSPLIGMFCRKIRFRQSLAYLYVF
SWGLRSIPELFYCLLPAYCLLHNSALFPKGVYLGIIITLVGIHCLYTLWEFMNLGFSIQSWYVTQSFGRI
KTTCSWLFSVLDVILKLLGISKTVFIVTKKTMPETKSGSGSKKSQREVDCPNQDSGKFEFDGSLYFLP
GTFIVLVNLAALAGCLVGLQSRGGGGSGLAEACGCILVVILFLPFLKGMFEKGKYGIPFSTLSKAAFLA
ALFVVLSVGN
>AT2G32530_CslB1
MADSSSSLPPLCEKISYKNYFLRVVDLTILGFLFSLLLYRILLMNQNNSVWVVAFLCESFFSFIWLLITSI
KWSPASYKSYPERLDERVHDLPSVDMFVTTADPVREPPILVANTLLSLLAVNYPANKLACYVSDDGC
SPLTYFSLKEASKFAKIWVPFCKKYNIKVRAPFRYFLNPPAATESSEFSKDWEITKREYEKLSRRVEDA
TGDSHWLDAEDDFEDFSNTKPNDHSTIVKVVWENKGGVGVENEVPHFVYISREKRPNYLHHYKAGA
MNFLVRVSGLMTNAPYMLNVDCDMYANEADVVRQAMCIFLQKSMNSNHCAFVQFPQEFYDSNADE
LTVLQSYLGRGIAGIQGPTYAGSGCFHTRRVMYGLSIDDLEDDGSLSSLATRKYLAEENLAREFGNSN
EMVTSVVEALQRKPNPQNTLANSLEAAQEVGHCHFEYQTSWGKTIGWLYESTAEDANTSIGIHSRG
WTSSYISPKPPAFLGAMPPGGPEAMLQQRRWATGLLEVLFNKQSPLIGMFCRKIRFRQSLAYLYIFT
WGLRSIPELIYCLLPAYCLLHNAALFPKGVYLGIVVTLVGMHCLYSLWEFMSLGFSVQSWFASQSFW
RIKTTCSWLFSIPDIILKLLGISKTVFIVTKKTMPKTMSGSGSEKSQREVDCPNQDSGKFEFDGSLYFLP
GTFILLVNLAALAGCSVGLQRHRGGGSGLAEACGCILVVILFLPFLKGMFEKGKYGIPWSTLSKAAFLA
VLFVVFSVGN
>CSLF_2BS_TGACv1_146146_AA0456710.1_Triticum_aestivum
MAAAVTRRANALRVEAPDGNTESGRASLAADSPVAKRAVDAKDDVWVAADEGEASGSIAGDGNRT
PLFRTFKVKGSILHPYRFMILVRLVAIVAFFAWRVKHKNHDGVWLWATSMVADVWFGFSWLLNQLPK
LNPVKRVPDLAALADHSGDANLPGIDIFVTTVDPVDEPLLYTVNTILSILATDYPVDKYACYLSDDGGTL
VHYEAMIEVANFAVLWVPFCRKYCVEPRSPENYFGMKTQPYAGSMAGEFMRDHRRVRREYDELKV
RVDSLSTTIRQRSDAYNSSTKGDGVRATWMADGTQWPGTWIEQVENHRRGQHAGIVQVILGHPSCK
PQLGSPASSDNPLDFSNVDTRLPMLVYMSREKRPGYNHQKKAGAMNVMLRVSALLSNAPFVVNFD
GDHYINNSQALCAPMCFMLDPRDGQNTAFVQFPQRFDDVDPTDRYANHNRVFFDGTMLSLNGLQG
PSYLGTGTMFRRVALYGMEPPRYRAENIKLAGKVNEFGSSTSFINSMPDGAIQERSITPVLVDEALSN
DLATLMTCAYEDGSSWGRDVGWVYNIATEDVVTGFRMHRQGWRSMYCSMEPAAFRGTAPINLTER
LYQVLRWSGGSLEMFFSHSNALMAGRRLHPLQRIAYLNMSTYPIVTVFILAYNLFPVLWLFSEQFYIQ
RPFAWGFFTDQARHVLLGMLFNVWILVLLYPFALGIMGKWGKRPVILFVMLVMAVGAVGLLYVAFHA
PYPADFSEVAASLGEASLTGPSG
>CSLH_2DL_TGACv1_158387_AA0517170.1_Triticum_aestivum
MAGGKKLQERVALGRTAWMLADFVILLLLLLALVARRAASLGERGGTWLAALVCEAWFAFVWILNMN
GKWSPVRFDTYPDNLSHRMEELPAVDMFVTTADPALEPPLITVNTVLSLLALDYPDVGRLACYVSDD
GCSPVTCYALREAAKFAGLWVPFCKRHDVGVRAPFMYFSSAPEVGNGTVDHEFLESWALMKSQYE
KLARRIENADEGTIMRDGGDEFAEFIDAERGNHPTIVKVLWDNSKSKAGEEFPHLVYLSREKSPRHRH
NFKAGAMNVLTRVSAVMTNAPIMLNVDCDMFANNPQVALHAMCLLLGFDDEIHSGFVQAPQKFYGG
LKDDPFGNQMQVITKKIGGGLAGIQGMFYGGTGCFHRRKVIYGVPPPDTVKHEMKGSPSYKELQAKF
GSSKELIESSRNIISGDLLARPTVDLSSRVEMAKQVGDCNYEAGTCWGQEIGWVYGSMTEDILTGLRI
HAAGWESALLDTEPPAFLGCAPTGGPASLTQFKRWATGLLEILISQNSPILGTIFRRLQLRQCLAYLIVE
AWPVRAPFELCYALLGPFCLLTNQSFLPTASDEGFRIPAALFLTCHIYHLMEYKECGLSVRAWWNNH
RMQRITSASAWLLAFLTVILKTLGLSETVFEVTRKESSTSSDGGAGTDEADPGLFTFDSAPVFIPVTVL
SMLNIVALAVAAWRAVVGAAAGVHGGPGIGEFVCCGWIVLCFWPFVRGLVSRGKYGIPWSVRVKAG
LIVAAFVHICTRN
>CSLH_2AL_TGACv1_094351_AA0296300.3_Triticum_aestivum
MAGGKKLHERVALGRTAWMLADFVILLLLLLALVARRAASLGERGGTWLAALVCEAWFAFVWILNMNG
KWSPVRFDTYPENLSHRLEELPAVDMFVTTADPALEPPLITVNTVLSLLALDYPDVGKLACYVSDDGC
SPVTCYALREAAKFASLWIPFCKRYDVGVRAPFMYFSSAPEVGTGTADHEFLESWALMKTEYEKLAS
RIENADEVSILRDGGEEFAEFIDAERGNHPTIVKVLWDNSKSKAGEGFPHLVYLSREKSPRHRHNFKA
GAMNVLTRVSAVMTNAPIMLNVDCDMFANNPQVALHAMCLLLGFDDEIHSGFVQAPQKFYGGLKDD
PFGNQMQVITKKIGGGLAGIQGTFYGGTGCFHRRKVIYGMPPPDTVKHETRGSPSYKELQAKFGSSK
ELIESSRNIISGDLLARPTVDISSRVEMAKQVGDCNYEAGTCWGQEIGWVYGSMTEDILTGQRIQAAG
WESALLDTDPPAFLGCAPTGGPASLTQFKRWATGLLEILISRNSPILGTIFKGLQLRQCLGYLIVDAWP
VRAPFELCYALLGPFCLLTNQSFLPTASDEGFHIPAALFLTYNIYHLMEYKECGLSVRAWWNNHRMQ
RITSASAWLLAFLTVILKTLGLSETVFEVTRKESSTSSDGGAGTDDADPGLFTFDSAPVFIPVTALSVLN
IVALTVAAWRAVVGTVAGVHGGPGVGEFVCCGWMVLCFWPFVRGLVSSGKYGIPWSVRVKAGLIVA
AFVHLCTRN
>CSLH_3B_TGACv1_222234_AA0760340.1_Triticum_aestivum
MSSAMKLQERVSVPRTAWKLADIFILCLLFALLSCRVASLREGGASVAALVCEAWFTFVWIINMNIKW
NPVRFNTYPENLSQRTDELPAVDMLVTTADPELEPPLMTVNTVLSLLAVDYPDVDKLACYVSDDGCS
PVTCYALREAAGFARLWVPFCKRHGVGVRAPFMYFASSRPEPELAGDWTFIKSEYDKLVSRIESADE

GSLLRHDDAADFTEFKEAKRGDHPAIVKVLWDNSKSSRTGSGDGFPNLVYVSREKTRKHDHHYKAG
AMNVLARVSAVMTNAPIILNMDCDMFVNNPQVVLHAMCLLLGFNDETCSGFVQVPQRFYAKLKDDPF
GNQIEVLREKLLGGLSGLQGIYYLGTGCFHRRKIIYGVAPPSFAAVKHERQGSLTYEDLRTKFGASVEL
AESARNIYSREIPLKPMIDISSRIQVAKQVSSCNYETGTHWGQEIGWSYGSMAEDILTGQRIHSAGWK
STSPDTNPPAFLGCAPTGGPASLTQYKRWATGLLEILLGPNTPIIATIFKRLQFRQYLGYLVFYVWSMR
APFELCYALLGPFCLFRNHSFLLKASNHGFSIQLALFLSYNIYNFVEYMECGLSARTWWNNMRMQRIV
SLSSWLLAFLSVVLKTIGLSKTVFEVTRKDKSTSDGDPSTHETDLGWFTFDSSPVFIPMTAVAILNIVTI
AIGVWRHAIFWMTTGNHDCQNIGEFLCCGLMILYFWPFIKGLVGRGRYGIPWNVKLKAWVIVVAFLYF
CRGD
>CSLC_3B_TGACv1_220758_AA0718310.2_Triticum_aestivum
MASSWWGDKEEHGTPVVVKMDNPYSLVEIDGPGMDSSEKARRSKNAKQFKWVLLLRAHRAVGCVA
WLAGGFWGLLGAVNRRVRRSRDADAEPDAEASGRGRHMLGFLRAFLLLSLAMLAFETAAYLKGWH
YFPRDLPEHYLRQLPEHLQNLPENLRHLPENLRHLPDGLRMPEQQEIQGWLHRAYVAWLAFRIDYIA
WAIEKLSGFCIVLFMVQSIDRILLCLGCFWIKLRGIKPGLKAAASKRGSKYADENDLEDGDDLGAYFPM
VLLQMPMCNEKEVYETSISHVCQIDWPRDRMLVQVLDDSDDETCQMLIRAEVTKWSQRGVNIIYRHR
LSRTGYKAGNLKSAMSCEYVKDYEFVAIFDADFQPNPDFLKLTVPHFKGNPELGLVQARWSFVNKDE
NLLTRLQNINLCFHFEVEQQVNGIYLNFFGFNGTAGVWRIEALEDSGGWMERTTVEDMDIAVRAHLQ
GWKFIYLNDVKVLCELPESYQAYRKQQHRWHSGPMQLFRLCLPAIIKSKIPLWKKANLVMLFFLLRKLI
LPFYSFTLFCVILPLTMFVPEAELPIWVICYVPMIMSVLNILPAPKSFPFVIPYLLFENTMSVTKFNAMVS
GLFQLGSSYEWVVTKKAGRTSSESDIFAMAEETNTATRPAPRLVRGVSEAGLEAWAKTHQLDNKDL
QLKAEAEEVTSLAAAIKKTSKAKPPNRIFKKELALAFLLLIAATRSLLSAQGLHFYFLLFQGVTFLVVGLD
LIGEQVS
>OS10G20090_CSLH1
MEAAARGNKKLQERVPIRRTAWRLADLAILFLLLALLLHRVLHDSGAPWRRAALACEAWFTFMWLLN
VNAKWSPVRFDTFPENLAERIDELPAVDMFVTTADPVLEPPLVTVNTVLSLLALDYPAAGEKLACYVS
DDGCSPLTCYALREAARFARTWVPFCRRHGVAVRAPFRYFSSTPEFGPADGKFLEDWTFMKSEYEK
LVHRIEDADEPSLLRHGGGEFAEFLDVERGNHPTIIKVLWDNNRSRTGDGFPRLIYVSREKSPNLHHH
YKAGAMNALTRVSALMTNAPFMLNLDCDMFVNNPRVVLHAMCLLLGFDDEISCAFVQTPQKFYGALK
DDPFGNQLEVSLMKVGRGIAGLQGIFYCGTGCFHRRKVIYGMRTGREGTTGYSSNKELHSKFGSSN
NFKESARDVIYGNLSTEPIVDISSCVDVAKEVAACNYEIGTCWGQEVGWVYGSLTEDVLTGQRIHAAG
WRSTLMEIEPPAFMGCAPNGGPACLTQLKRWASGFLEILISRNNPILTTTFKSLQFRQCLAYLHSYVW
PVRAPFELCYALLGPYCLLSNQSFLPKTSEDGFYIALALFIAYNTYMFMEFIECGQSARACWNNHRMQ
RITSASAWLLAFLTVILKTLGFSETVFEVTRKDKSTSDGDSNTDEPEPGRFTFDESTVFIPVTALAMLSV
IAIAVGAWRVVLVTTEGLPGGPGISEFISCGWLVLCFMPLLRGLVGSGRYGIPWSIKMKACLLVAIFLLF
CKRN
>CSLC_3AS_TGACv1_211225_AA0686890.2_Triticum_aestivum
MASSWWGDKEEHGTPVVVKMDNPYSLVEIDGPGMDSSEKARRSKNAKQFKWVLLLRAHRAVGCVA
WLAGGFWGLLGAVNRRVRRSRDADAEPDAEASGRGRHMLGFLRAFLLLSLAMLAFETAAYLKGWH
YFPRDLPEHYLRQLPEHLQNLPEHLRHLPENLRHLPDGLRMPEQQEIQGWLHRAYVAWLAFRIDYIA
WAIEKLSGFCIVLFMVQSIDRILLCLGCFWIKVRGIKPGLVATKKRGNKYADDNDLEDGDDLGAYFPMV
LLQMPMCNEKEVYETSISHVCQIDWPRDRMLVQVLDDSDDETCQMLIRAEVTKWNQRGVNIIYRHRL
SRTGYKAGNLKSAMSCEYVKDYEFVAIFDADFQPNPDFLKLTVPHFKGNPELGLVQARWSFVNKDE
NLLTRLQNINLCFHFEVEQQVNGIYLNFFGFNGTAGVWRIEALEDSGGWMERTTVEDMDIAVRAHLQ
GWKFIYLNDVKVLCELPESYQAYRKQQHRWHSGPMQLFRLCLPAIIKSKIPLWKKANLVMLFFLLRKLI
LPFYSFTLFCVILPLTMFVPEAELPIWVICYVPMIMSVLNILPAPKSFPFVIPYLLFENTMSVTKFNAMVS
GLFQLGSSYEWVVTKKAGRTSSESDIFAMAEKTDTATRPAPRLVRGVSEAGLEAWAKTHQLDNKDL
QLKAEAEEVTSLAAAIKKTSKAKPPNRIFKKELALAFLLLIAATRSLLSAQGLHFYFLLFQGVTFLVVGLD
LIGEQVS
>OS08G15420_OsCslC3
MAPPPNTYSESWWGGKEERGTPVVVKMDNPYSLVEIDGPGMAAPSEKARGKNAKQLTWVLLLRAH
RAVGCVAWLAAGFWAVLGAVNRRVRRSRDADAEPDAEASGRGRAMLRFLRGFLLLSLAMLAFETVA
HLKGWHFPRSAAGLPEKYLRRLPEHLQHLPEHLRRHLPEHLRMPEKEEIEGWLHRAYVAWLAFRIDY
IAWAIQKLSGFCIALFMVQSVDRLVLCLGCFWIKLRGIKPVADTSISNDDIEATAGDGGGYFPMVLIQM
PMCNEKEVYETSISHVCQIDWPRERMLVQVLDDSDDETCQMLIKAEVTKWSQRGVNIIYRHRLNRTG
YKAGNLKSAMSCDYVRDYEFVAIFDADFQPNPDFLKLTVPHFKGNPELGLVQARWSFVNKDENLLTR
LQNINLCFHFEVEQQVNGVYLSFFGFNGTAGVWRIKALEDSGGWMERTTVEDMDIAVRAHLNGWKFI
FLNDVKVLCELPESYQAYRKQQHRWHSGPMQLFRLCLPAVFKSKISTWKKANLVMLFFLLRKLILPFY
SFTLFCVILPLTMFVPEAELPIWVICYVPVIMSVLNILPAPKSFPFVIPYLLFENTMSVTKFNAMVSGLFQ
LGSSYEWVVTKKAGRTSSESDILALAEAADADARPPPAKLHRGVSEGGLKEWAKLHKEQEDATAAA
AAAAAPGTPVKKSKAAKAPNRIFKKELALAFLLLTAATRSLLSAQGLHFYFLLFQGVTFLAVGLDLIGEQ
VS
>GRMZM2G012044_P01

MAGSSVRGGSNCPPLFVTEKPTRMARYAYRLFASTVLAGVLLVWLYRATHVPPMSSGARWWAWLG
LSAAELWFGFYWVLTLSVRWSPVFRRAFPDQLLRRYKEEQLPGVDIFVCTADPTVEPPMLVISTVLSV
MAYDYPKEKLNIYLSDDAGSIITLYALYEASEFAKHWLPFCNKYQVEPRSPAAYFGTEASPPDACDRK
EWFSLKEMHKDLAARVNSVVNSGKIPEVSKCKLMGFSRWSENASFRDHPSIVQILIDGNKRKATDVD
GKVLPTLVYMAREKRPQEHHHFKAGSLNALIRVSSVISNSPVIMNVDCDMYSNNSGSIRDALCFFQDE
QLGQDIAFVQYPQNFENVVQNDIYGNPINTVNELDHPCLDGWGGMCYYGTGCFHRREALCGRIYSP
DYKEDWTRVARKTEDVIDLEGMAESLVTCTYEHNTLWGVEKGVIYGCPLEDVITGLQIQCRGWRSVY
HNPPRKGFLGMAPTSLGQILVQHKRWTEGFLQISLSKYSPFLLGHRKISLGLQMGYSVCGFWAANSF
PTLYYVTIPSLCFLNGISLFPEITSPWFVPFAYVAVAAYSCSLVESLQCGDTAVEWWNAQRMWLFRRI
TSYLLAAIDTIRRMLGVTESGFTLTAKVTDPRALERYKKGMMEFGSFSVMFAIITTVALLNLACMMLGV
AKVLLRKGAVSLGAMFVQAVLCALIVAINFPVYEAMFVRKDSGRLPASVSVVSLCIVLPFCILPTKL
>CSLJ_3DS_TGACv1_272297_AA0918580.1_Triticum_aestivum
MAAEPSQDAPLQLHTVQTDQPLATVNRHLAALHVALAAAAIAHRGAHVMLAADLVLLFLWALSQAPM
WRPVSRAAFPSRLSRAALPAVDVMVVTADPDKEPAAKVMNTVVSAMALDYPGGRLSVYLSDDAGSP
RTLLAARKAYAFARAWVPFCRKYGVRCPCPDRFFAGDDQLDLGGHHRQELDDDRLRIKNMYETFKE
GVEKVMNDAALSQSWTKADHDAHVEQDSSNSNSGDGEEDEDAMPLLVYVSREKRRSSAHHFKAGA
LNVLLRVSSLMSNSPYVMVLDCDMYCNSWSSVLEAMCFHLDGRRRADLAFVQFPQMFHNLSSSDIY
ANELRSIFWAGPTGLRDAVERRGRPPGPILSGTGFCVRRDAVYGAGPGSSQDHQSSGVEVGELKRR
FGVSNGHIASLRRSGTGSTIVARDGLPQPQEDAELVASCDYETGTEWGEEVGFLYQSVVEDYFTGY
RQLYCPGWTSVYCFPATGTRPPFLGSVPTNLNDALVQNKRWMSGMLAVGLSRHCPLASAAAVSVP
QSMGFAYYAFMALYAFPVLCYATVPQLCFLRGGTSFPGESALWFAAVLASSSLQHLVEVSFAKRGLA
ARAWWNEQRFWALNAVTGQLFACVSVALSLVGAGGRAVDFDLTSKASDDRLYRDSVFDFAGCSAL
LLPATTLCLLNTAALVGGVWKMVGRGGSVSGELFLLCYVAALSYPLLEGMFLRRDPARVPAWITAMS
VAMVATLLSLFG
>CSLE_6DL_TGACv1_526558_AA1687090.1_Triticum_aestivum
MAGSSVSGGGGGRPPLFATEKPKRVLAYRLYAGTIFAGILLIWFYRATHIPARGSSSLGWRAGLGLLV
AELWFGLYWVLTLSVRWNPVRRATFKDRLSERYDDDQLPGVDIFVCTADPALEPPMLVISTVLSVMA
YDYPPEKLNIYLSDDAGSAVTFYALHEASEFAKHWIPFCKNYKVEPRSPAAYFAKGATPHDACSPQEF
LRMKELYKDLTDRMNSVVHSGKIPEVPECNHRGFSEWNETITSGDHPSVVQILIDRNKRKAVDVDGN
ALPKLVYMAREKRPQEQHHFKAGSLNALIRVSSVISNSPVILNVDCDMYSNNSESIRDALCFFLDEEQ
GQDIGFVQYPQNFDNVVHNDSYGNPINVVNELDNPCLDGWGGMCYYGTGCFHRRETLSGQIYSKD
YKEDWARGVGIAENADELEETSKSLVTCTYEHNTPWGIEKGVRYGCPLEDVITGLQIQCHGWRSVYY
NPARKGFLGKAPTSLGQILVQHKRWSEGFLQMSLSNYSPFLLGHGKIKLGLQMGYSVCGFWALNSF
PTFYYVIIPSLCFLSGVSVFPEITSPWCIPFIYVVVASYSWSLMESLQCGDTAVEWWNAQRMWLMRR
TTSYLLAAIDTIGGMLGVSESGFELTVKVDESQALERYKKGKMEFGPISGMFVIITTIALFNLVCLVLGL
GRVLLRGGAEGLGPLFLQAVLCVAIVVINAPVYEALFIRRDSGGLPYFVTLVSLCFVSSLCLQAI
>CSLE_6AL_TGACv1_471004_AA1500600.1_Triticum_aestivum
MAGSSVSGGGGRPPLFATEKPKRVLAYRVYAGTIFAGILLIWFYRATHIPARGSSSLGWRAGLGLLVA
EILFGLYWVLTLSVRWNPVRRTTFKDRLSERYDDDQLPGVDIFVCTADPALEPPMLVISTVLSVMAYD
YPPEKLNIYLSDDAGSAVTFYALHEASEFAKHWIPFCKNYKVEPRSPAAYFAEGATPHDACSPQELLR
MKELYKDLTDRVNSVVHSGKIPEVPECNHRGSSEWNEMITSGDHPSIVQILIDRNKRKAVDVDGNALP
KLVYMSREKRPQEQHHFKAGSLNALIRVSSVISNSSVILNVDCDMYSNNSESIRDALCFFLDEEQGQD
IGFVQYPQNFDNVVHNDIYGNPINVVNELDNPCLDGWGGMCYYGTGCFHRRETLSGQIYSKDYKED
WDRGVGIAENADELEETSKSLVTCTYEHNTPWGIEKGVRYGCPLEDVITGLQIQCRGWRSVYYNPAR
KGFLGMAPTSLGQILVQHKRWSEGFLQISLSNYSPFLLGHGKIKLGLQMGYSVCGFWALNSFPTFYY
VIIPSLCFLSGVSVFPEITSPWCIPFIYVVVASYSWSLMESLQCGDTAVEWWNAQRMWLMRRTTSYLL
AAIDTIGGMLGVSESGFELTVKVDESQALERYKKGKMEFGPISGMFVIITTIALFNLVCLVVGLGRALLR
EGTAGLGPLFLQAVLCVAIVVINAPVYEALFIRRDSGSLPYFVTLVSLCFVSSLCLQAI
>CSLE_6BL_TGACv1_499967_AA1596110.2_Triticum_aestivum
MAGSSVSGGGRPPLFATEKPKRVLAYRLYAGTIFAGILLIWFYRATHIPERGDSSLGWRAGLGLLVAE
LLFGLYWVLTLSVRWNPVRRTTFKDRLSERYDDDQLPGVDIFVCTADPALEPPMLVISTVLSVMAYDY
PPEKLNIYLSDDAGSAVTFYALHEASEFAKHWIPFCKNYKVEPMSPAAYFAEGATPHDACSPQELLR
MKELYKDLTDRVNSVVHSGKIPEVPECNHRGFSVWNETITSGDHPSIVQILIDRNKRKAVDVDGNALP
KLVYMAREKRPQEQHHFKAGSLNALIRVSSVISSSPVILNVDCDMYSNNSESIRDALCFFLDEEQGQD
IGFVQYPQNFDNVVHNDIYGNPINVVNELDNTCLDGWGGMCYYGTGCFHRRETLSGQIYSKDYKED
WARGVGIAENADELEETSKSLVTCTYEHNTPWGIEKGVRYGCPLEDVITGLQIQCRGWRSVYYNPAR
KGFLGMAPTSLGQILVQHKRWSEGFLQISLSNYSPFLLGHGKIKLGLQMGYSVCGFWALNSFPTFYY
VIIPSLCFLSGVSVFPEITSPWCIPFIYVVAASYSWSLMESLQCGDTAVEWWNAQRMWLMRRTTSYLL
AAIDTIGGMLGVSESGFELTVKVDESQALERYKKGKMEFGPISGMFVIITTIALFNLVCLVLGLGRVVLR
EGAAGLGPLFLQAVLCVAIVVINAPVYEALFIRRDSGSLPYFVTLVSLCFVSSLCLQAI
>CSLJ_3DS_TGACv1_272756_AA0924850.1_Triticum_aestivum

MATKPSQDAPLPLHTVQTDQPLATVNRLLAAVHLALGAAAIAHRGAHVMLAADLVLLFLWALSQAPM
WRPVSRTAFPSRLSRAALPAVDVMVVTADPEKEPAAKVMNTVVSAMALDYPGGRLSVYLSDDAGSP
RTLLAARKAYAFARAWVPFCRKYGVRCPCPDRFFAGDDKLDLGSHHHHELADDRLRIKNMYETFNE
GVREVMSDADLSQSCTKADHDAHVEIITGDEQDSSNSNSGDGEEDEDAMPLLVYVSREKRRSSTHH
FKAGALNVLLRVSSLLSNSPYVMVLDCDMYCNSRSSILEAMCFHLDGRRRADLAFVQFPQMFHNLSS
SDIYANELRSIFWTRWKGLDGLRGPILSGTGFCARRDAIYGARPASSQDQFSGVEVGELKRRFGVSN
GHIASLRRSGTGSTIVARDALPQEDAELVASCAYETGTEWGEQVGFLYQSVVEDYFTGYRQLYCRG
WTSVYCFPAAASRPPFLGSVPTNLNDALVQNKRWMSGMLAVGLSRHCPLASAAICVPQSMGFAYYA
FMALYAFPVLCYATVPQLCFLRGGTSFPGAASTLWFAAVFASSSLQHLVEVSVAKRGLALRTWWNE
QRFWALNAVTGQLFACLGVALNLVGAGGRAVDFDLTSKASDDRLYRDGVFDFAGCTTLLLPATTLCL
LNAAALVGGVWKMVGRGGSVSGELFLLCYVAALSYPLLQGMFLRRDPARVPAPITAMSVAMVAALLS
LFG
>CSLJ_3B_TGACv1_221705_AA0747940.1_Triticum_aestivum
MAAKPSQDAPLQLHTVEVDQPIATVNRLLAVLHVALAAAAIAHRGAHVMLAADLVLLFLWALSQAPM
WRPVSRAAFPSRLSRAALPAVDVMVVTADPDKEPAAKVMNTVVSAMALNYPGGRLSVYLSDDAGSP
RTLLAARKAYAFARAWVPFCRKYGVRCPCPDRFFAGDDQLDLDGHHRQELDDDRLRIKKMYETFKE
GVEEVMSDAALSQSWTKADHDAHVEIITGDEQDSSNSNSGDGEEDEDATPLLVYVSRGKRRSSTHH
FKAGALNVLLRVSSLMSNSPYVMVLDCDMYCNSRSSILEAMCFHLDGRRRADLAFVQFPQMFHNLS
TSDIYANELRSIFWTRWKGLDGLRGPILSGTGFCARRDAIYGALPASSQDQFSGVEVGELKRRFGVS
NGHIASLRRPGTGSTIVARDALPQDAELVACCDYETGTEWGEEVGFLYQSVVEDYFTGYRQLYCRG
WTSVYCFPATGSRPPFLGSVPTNLNDALVQNKRWMSGMLAVGLSRHCPLASAAAISVPESMGFAYY
AFMALYAFPVLCYAIVPQLCFFRGGTSFPEASTLWFAAVFVSSSLQHLVEVSVAKRGLAARTCWNEQ
RFWALNAVTGQLFACLSVALNLVDGAGGRAVDFDLTSKASDDRLYRDGVFDFAGCSTLLLPATTLCL
LNAAALVGGVWKMVGRGGNMPGELFLLCYIAALSYPLLQGMFLRRDLARVPARITAMSVAMVATLLS
LFG
>CSLE_5BL_TGACv1_406235_AA1342600.1_Triticum_aestivum
MERSRRLFETETHGGRAVYRLHAVTVAAGILLLLYYRATRVPAAGEGRAAWLGMLAAELCYAAYWV
VTQSVRWSPLHRRPCRDRLAARYGERLPCVDIFVCTADPHSEPPSLVISTVLSLMAYNYPAEKISVYL
SDDGGSILTFYALWEASLFAKHWIPFCKRYNIEPRSPATYFSESDGHQDMCTPKEWSLIREMYEDMT
ERIDTAALSGKISEEVKENHKGFHEWDQENTSKNHQPIVQILIEGKDKNANDDEGNVLPTLVYMAREK
RPQHHHNFKAGAMNALIRVSSVISNSPIIMNVDCDMYSNNYDTIRDALCFFLDEEMGHKIGFVQFPQN
YNNLTKNNIYGNSHQVTNQVLMGGMDSVGGPMYVGTGCFHRREILCGRRFTKDYKEDWDGGIKDK
TQESIDEIEEKAKSLAASTYEHDTQWGDEIGIKYGYPAEDIVTGLEIHCRGWKSVHSNPPRPAFLGVAP
TTLAQTLLQHKRWSEGSFSIFLSKYCPFMFGHGKIKLRHQMGYSIYGLWAPNSIPTLYYVIIPSLALLKG
ISLFPEITSPWISPFIYVVCVKNMYSLYEALSCGDTLKGWWNEQRMWMVRRITSYLYGLTDTVRKLLG
LSKMTFAVTSKVSEENESKRYEQEIMEFGSSAPEYVIIATVALLNLICLVGGLSQILTGGWNILLNVFSP
QLILCGMLVITNIPFYEAMFVRKDKGRMPFSVTLASISFAMLALLVPIVYNSRYI
>AT4G23990_CslG1
MEPHRKHSVGDTTLHTCHPCRRTIPYRIYAVFHTCGIIALMYHHVHSLLTANTTLITSLLLLSDIVLAFM
WATTTSLRYKPVRRTEYPEKYAAEPEDFPKLDVFICTADPYKEPPMMVVNTALSVMAYEYPSDKISVY
VSDDGGSSLTLFALMEAAKFSKHWLPFCKKNNVQDRSPEVYFSSKLRSRSDEAENIKMMYEDMKSR
VEHVVESGKVETAFITCDQFRGVFDLWTDKFTRHDHPTIIQVLQNSENDMDDTKKYIMPNLIYVSREK
SKVSSHHFKAGALNTLLRVSGVMTNSPIILTLDCDMYSNDPATPVRALCYLTDPKIKTGLGFVQFPQTF
QGISKNDIYACAYKRLFEINMIGFDGLMGPNHVGTGCFFNRRGFYGAPSNLILPEIDELKPNRIVDKPIN
AQDVLALAHRVAGCIYELNTNWGSKIGFRYGSLVEDYYTGYRLHCEGWRSVFCRPKRAAFCGDSPK
SLIDVVSQQKRWAIGLLEVAISRYSPITYGVKSMGLVTGVGYCQYACWAFWSLPLIVYGFLPQLALLY
QSSVFPKSSDPWFWLYIVLFLGAYGQDLLDFVLEGGTYGGWWNDQRMWSIRGFSSHLFGFIEFTLK
TLNLSTHGFNVTSKANDDEEQSKRYEKEIFEFGPSSSMFLPLTTVAIVNLLAFVWGLYGLFAWGEGLV
LELMLASFAVVNCLPIYEAMVLRIDDGKLPKRVCFVAGILTFVLIVSGYVFLK
>AT1G55850_CslE1
MVNKDDRIRPVHEADGEPLFETRRRTGRVIAYRFFSASVFVCICLIWFYRIGEIGDNRTVLDRLIWFVM
FIVEIWFGLYWVVTQSSRWNPVWRFPFSDRLSRRYGSDLPRLDVFVCTADPVIEPPLLVVNTVLSVTA
LDYPPEKLAVYLSDDGGSELTFYALTEAAEFAKTWVPFCKKFNVEPTSPAAYLSSKANCLDSAAEEVA
KLYREMAARIETAARLGRIPEEARVKYGDGFSQWDADATRRNHGTILQVLVDGREGNTIAIPTLVYLS
REKRPQHHHNFKAGAMNALLRVSSKITCGKIILNLDCDMYANNSKSTRDALCILLDEKEGKEIAFVQFP
QCFDNVTRNDLYGSMMRVGIDVEFLGLDGNGGPLYIGTGCFHRRDVICGRKYGEEEEEEEESERIHEN
LEPEMIKALASCTYEENTQWGKEMGVKYGCPVEDVITGLTIQCRGWKSAYLNPEKQAFLGVAPTNLH
QMLVQQRRWSEGDFQIMLSKYSPVWYGKGKISLGLILGYCCYCLWAPSSLPVLIYSVLTSLCLFKGIP
LFPKVSSSWFIPFGYVTVAATAYSLAEFLWCGGTFRGWWNEQRMWLYRRTSSFLFGFMDTIKKLLG
VSESAFVITAKVAEEEAAERYKEEVMEFGVESPMFLVLGTLGMLNLFCFAAAVARLVSGDGGDLKTM
GMQFVITGVLVVINWPLYKGMLLRQDKGKMPMSVTVKSVVLALSACTCLAFL
>OS09G30130_OsCslE6

METTTTERRRLFATEKVGGRAVYRLQAATVAAGILLVLYYRATRVPAAGEGRAAWLGMAAAELWFAV
YWVITQSVRWCPVRRRTFKNRLAERYKENLPGVDVFVCTADPHAEPPSLVISTILSVMAYNYPSEKIS
VYLSDDGGSILTFYALWEASMFAKKWLPFCRRYNIEPRSPAAYFSESEGHHNLCSPKEWSFIKNLYE
EMRERIDSAVMSGKIPEEIKLKHKGFDEWNSEMTSKNHQPIVQVLIDGKSQNAVDDDGNVLPTLVYM
AREKSPQYHHNFKAGALNALIRVSALISDSPVILNVDCDMYSNNSDSIRDALCFFLDEEMSHKIGFVQY
PQNYNNMTKNNIYGNSLNVINHVEMRGLDSAGGCLYIGTGCFHRREILCGKKFSKDYKEDWGRGIKE
RGHENIDEIEEKAKSLATCTYELRTQWGNEIGVKYGCPVEDVITGLAIHCRGWESVYMEPQRAAFVG
VAPATLAQTILQHKRWSEGNFTIFLSKHNTFLFGHGKISLQLQMGYCIYGLWAANSLPTIYYVMIPALG
LVKGTPLFPEIMSPWATPFIYVFCVKTLYSLYEALLSGDTLKGWWNGQRMWMVKRITSYLYGFIDTIR
KLLGLSKMSFEITAKVSDGDEAKRYEQEILEFGSSSPEFVIIATVALLNFVCLVAGLSKIMAGVWNVFLP
QVILCGLIVITNIPIYEAMFVRKDKGRIPLPVTLASIGFVMLAFLLPIV
>CSLE_5DL_TGACv1_433536_AA1415830.1_Triticum_aestivum
MERTRRLFETETHGGRAAYRLHAVTVAAGILLLLYYRATHVPAAGEGRAAWLGMLAAELWYAAYWA
VTQSVRWSPVRRLPFIDRLAARYGERLPCVDIFVCTADPHSEPPSLVISTVLSLMAYNYPAEKISVYLS
DDGGSVLTFYALWEASLFAKHWIPFCKRYNIEPRSPAAYFSESDGHQDLCSPKEWSLIREMYEDMTE
RIDTAVLSGKISEEVKANHKGFHEWDQENTSKNHQPIVQILIEGKDKNANDDEGNVLPTLVYMAREKR
PQHHHNFKAGAMNALIRVSSVISNSPIIMNVDCDMYSNNCDTIRDALCFFLDEEMGHKIGFVQFPQNY
NNLTKNNIYGNSHQVTNQVLMGGMDSVGGPMYVGTGCFHRREILCGRRFTEDYKEDWNGGIKDKT
QESIVEIEEKAKSLAASTYEHDTQWGDEIGIKYGYPAEDIVTGLGIHCRGWKSVHSNPPRPAFLGVAP
TTLAQTLLQHKRWSEGSFSIFLSKYCPFMFGHGKIKLRHQMGYSIYGLWAPNSIPTLYYVIIPSLALLKG
ISLFPEITSPWMSPFIYVLCVKNMYSLYEALSCGDTLKGWWNEQRMWMVRRITSYLYGLTDTVRKLL
GLSKMTFAVTSKVSEESESKRYEQEIMEFGSSAPEYVIIATVALLNLNCLVVGLCQIMTGGWNILLNVF
SPQLILCGMLVITNIPFYEAMFVRKDKGRIPFSVTLASIGFAILALLVPIV
>CSLE_5BL_TGACv1_406235_AA1342610.1_Triticum_aestivum
MERSRRLFETETHGGRAAYRLHAVTVAAGILLVLYYRATHVPAAGEGRATWLGMLAAELWYAAYWV
VTQSVRWSPVRRRPFIDRLAARHGERLPCVDIFVCTADPYSEPPSLVVSTILSLMAYNYPPEKLSVYL
SDDGGSILTLYGMWEASLFAKHWLPFCKRYNIEPRSPAAYFSESDGHQELCTPKEWSLIKDMFDKMT
ERIDTAVMSGKVPEEIKARHKGFYEWNQEISSKNHQPIVQILIDGKDQNAVDNEGKVLPTLVYMAREK
RPQHHHNFKAGAMNALIRVSSVISNSPIIMNVDCDMYSNNNDAVRDALCFFLDEEMGHKIGFVQYPQ
NYNNLSKNNIYGNSLHVINEVEMGGMDSLGGPMYIGTGCFHRREILCGRKFTEDYQEDWNAGIKDKL
QESIDETEEKAKSLAACTYEHGTQWGDEIGVKYGCVVEDVNTGFAIHCRGWESVYNNPKKPAFMGV
GPTTLAQTILQHKRWSEGIFSIFLSKYNVFLFAHGKTKLQHQMGYHIYGLWAPNSLATLYYVIIPSLALL
KGTSLFPEITSPWIAPFVYVFCVKNMYSLYEALSSGDTLKGWWNGQRMWLVKRITSYLFGVLDNLRK
LLGLSKMNFVVTPKVSDEDESKRYEQEIMEFGSSDPEYVIIGTITLLNLVCLLGGLSKVMKGGWNEHL
DALFPQLILCGMVVITSIPFYEAMFLRKDKGRIPFPVTLASIGFVMLALLATIV
>CSLE_5AL_TGACv1_376126_AA1232370.2_Triticum_aestivum
MERTRLFETETHGGRAAYRLHAVTVAAGILLLLYYRATRVPAAGEGRAAWLGMLAAELWYAAYWVV
TQSVRWSPVRRRPFRDRLAARHGERLPSVDIFVCTADPYSEPPSLVVSTILSLMAYNYPPEKLSVYLS
DDGGSILTYYGMWEASLFAKHWLPFCKRYNIEPRSPAAYFSQSDGHQELCTPKEWSLIKDMFDEMT
ERIDTAVMSGKVPEEIKARQKGFHEWNQEITSKNHQPIVQILIDGKDQNAVDNEGNVLPTLVYMAREK
RPQHHHNFKAGAMNALIRVSSVISNSPIIMNVDCDMYSNNNDAVRDALCFFLDEEMGHKIGFVQYPQ
NYNNLSKNDIYGNSLQVINEVEMAGMDSLGGPLYIGTGCFHRREILCGRKFTKDYQEDWNAGIKDKL
QESIDETEEKAKSLAACTYEHGTQWGDEIGVKYGCAVEDVITGLAIHCRGWESVYNNPKKPAFMGVG
PTTLAQTILQHKRWSEGNLSIFLSKYNVFLFGHGKTKLRHQMGYHIYGLWAPNSLATLYYVIIPSLALLK
GTPLFPEITSPWIAPFVYVFCVKNMYSLYEALSSGDTLKGWWNGQRMWLVKRITSYLFGVLDNLWKL
LGLSKMNFVVSPKVSDEDESKRYEQEIMEFGSSDPEYVIIGTITLLNLVCLLGGLSKVMKVGWNNIHLD
ALFPQLILCGMVVITSIPFYEAMFLRKDKGRIPFPVTLASIGFVMLALLPAIV
>GRMZM2G014558_P01
MEERLFATEKHGGRALYRLHAVTVFLGICLLLCYRATHVPAAGSGGRAAWLGMLAAELWFGFYWVIT
QSVRWCPIRRRTFHDRLAARFGERLPCVDIFVCTADPRSEPPSLVVATVLSVMAYNYPPAKLNVYLS
DDGGSILTFYALWEASAFAKHWLPFCRRYGVEPRSPAAYFAQSDEKPRHDPPHALQEWTSVKNLYD
EMTERIDSAARTGNVPEETRAKHKGFSEWDTGITSKDHHPIVQILIDGKDKAVADNEGNVLPTLVYVA
REKRPQYHHNFKAGAMNALIRVSSVISNSPIILNVDCDMYSNNSDTIRDALCFFLDEETGHRIAFVQYP
QNYNNLTKNNIYGNSLNVINQVELSGLDAWGGPLYIGTGCFHRRETLCGRRFTEDYKEDWDRGTKE
QQQHRHRVDGETEAKAKSLATCAYEHDDDTRWGDEVGLKYGCSVEDVITGLAIHCRGWESVYSNP
ARAAFVGVAPTTLAQTILQHKRWSEGNFGIFVSRYCPFVFGRRGKTRLPHQMGYSIYGLWAPNSLPT
LYYAVVPSLCLLKGTPLFPELTSPWIAPFVYVAVAKNVYSAWEALWCGDTLRGWWNGQRMWLVRR
TTSYLYGFVDTVRDSLGLSKMGFVVSSKVSDEDEAKRYEQEMMEFGTASPEYVIVAAVALLNLVCLA
GMAAALDVFFVQVALCGVLVLLNVPVYEAMFVRKDRGRMPFPITLASVGFVTLALIVPFF
>CSLE_6DS_TGACv1_543277_AA1737920.1_Triticum_aestivum
MERRLFETVRHGGRALYRLHAVTVAASTLLVLYYRATRVPGSGGRRAAWLGMLAAELWYAAYWVVT
QSVRWSPVRRCTFRDRLTARYGDRLPGVDIFVCTADPLSEPPSLVISTILSVMAYNYLAEKLSVYLSD

DGGSVLTFYAMWEASLFAKHWLPFCKRYNIEPRSPAAYFSESYQDLCTPKEWSFIKDMYDEMTERID
TAVISRKIPEEIRSNHKGFYEWNPEITSKNHQPIVQVLIDGKDQKGVDSEGNVLPTLVYMAREKRPQH
HHNFKAGAMNALIRVSSVISNSPVIMNVDCDMYSNNNDTIRDALCFFLDEEMGHKIGFVQYPQNYNN
MTKNNLYGNSLHVINEVEMGGMDSLGGPLYVGTGCFHRREILRGKRFTKDYQEDWNGGIKGRIQDT
SIGEIEEKAKSLATCTYEHDTQWGDEIGLKYGCPVEDVITGLAIHCRGWESVYSNPTRPAFMGLGPTT
LAQTLLQHKRWSEGSFSIFLSKYCPFLFGHGKTNLRHQMGYCIYGLWAPNSLPTLYYVVIPSLALLKGI
PLFPKITSPWIAPFIYAFCVKNMYSLYEALSCGDTLKGWWNGQRMWLVKRITSFLFGAIDTVRKLLGL
SKMTFVVTPKVSIEDESKRYEEEIMEFGSSTPEYVIIATIALLNLVCLLGGLSQIMTGAWNIHLDAFSPQL
ILCGMLVITNMPFYEAMFLRNDKGKIPFTVTLASFGFVMLDFLVPIV
>AT4G24000_CslG2
MEPQRKHSTALHTCHPCRRTIPYRIYAVFHTCGIIALMYHHVHSIVNANNTLITCLLLLSDIVLAFMWAT
TTSLRLNPIHRTEYPEKYAAKPEDFPKLDVFICTADPYKEPPMMVVNTALSVMAYEYPSHKISVYVSD
DGGSSLTLFALMEAAKFSKHWLPFCKNNNVQDRSPEVYFSSKSHSSSDEAENLKMMYEDMKSRVE
HVVESGKVETAFIACDQFSCVFDLWTDKFTRHDHPTIIMVLQHNETEMMPNLIYVSREKSKVSPHHFK
AGALNTLLRVSAVMTNSPIILTLDCDMYSNNPTTPLHALCYLSDPKINFDLGFVQFPQKFQGVNKNDIY
ASELKRPFDINTVGFDGLMGPVHMGTGCFFNRRAFYGPPTTLILPEIETFGPNRIADKPIKAQDILALAH
DVAGCNYECNTNWGSKIGFRYGSLVEDYFTGFMLHCEGWRSIFCSPTKAAFYGDSPKCLTDVIGQQI
RWSVGLLEVAFSRYNPLTYGIKPLSLLMSLGYCHYAFWPFWCIPLVVYGILPQVALIHGVSVFPKASD
PWFWLYIILFLGGYAQDLSDFLLEGGTYRKWWNDQRMWMVRGLSSFFFGFTEFTLKTLNLSTQGYN
VTSKSNDDNEQMKRYEQEIFDFGPSSSMFLPITTVAIMNLLAFMRGLYGIFTWGEGPVLELMLASFAV
VNCLPIYEAMVLRIDDGKLPKRICFLAGLLSFVLTGSGYFFLK
>CSLH_3DS_TGACv1_271739_AA0907200.1_Triticum_aestivum
MSSAMKLQERVIVPRTAWKLADIFILCLLFALLSCRVLSLGEGGAGAASVAALVCEAWFTFVWILNMNI
RWNPVRFHTYPENLSQRMDGLPAVDMLVTTADPELEPPLMTVNTVLSLLAMDYPDVDKLACYVSDD
GCSPVTCYALHEAARFAGLWVPFCKRHGVGVRAPFMYFASRPEPELAGDNFSDEWTFIKSEYDKLV
SRIESADEGSLLRDDDAGEFTEFMEAKRGDHPGIVKVLWDNSKSSRTGEGFPNLVYVSREKSRKHD
HHYKAGAMNVLARVSAVMTNAPIILNVDCDMFVNNSQVVLHAMCLLLGFDDETCSGFVQVPQRFYG
KLKDDPFGNQMEVLREKLFGGLAGLQGIYYLGMGCFHRRKIIYGVAPSSSAAIKHEREGSRSYEDLRT
KFGASVELVESARNIYSGEIPPSPMIDISSRIQVAKQVSSCNYETDTHWGQEIGWSYGSMAEDILTGQ
RIHSSGWKSTLLDTNPPAFLGCAPTGGPASLTQYKRWATGLLEILLGQNSPIMATVFKRLQFRQSLAY
LVFYVWSMRAPFELCYALLGPFCLFRNQSFLLKASNHGFSIQLALFLSYNIYNFVEYMECGLSARTW
WNNMRMQRIVSISSWLLDFLSVVLKTIGLSKTVFEVTRKDKSTSDGDPSTHETDLGWFTFDSSPVFIL
VTTVAILNIATIAIGVWRHAIFWMITGNHDCQNIGELCVLDG
>CSLC_5BL_TGACv1_404820_AA1311790.1_Triticum_aestivum
MAPWTGLWGARAGAGAGAGAYRGTPVVVKMENPNWSISEISPEDAEDEDFLVSGAGAARRSRKGG
RGKNAKQITWVLLLKAHRAAGCLASLASAAVTLGAAARRRVADGRTDADAGAPGSAGESPVLRSRF
YAFIRAFLLLSLLLLAVELAARLHGWDLAASALALPIIGVESLYASWLRLRAAYLAPLLQFLTDACVVLFL
IQSADRLIQCLGSFYITVKRIKPTLKSPALPDAEDPDAGYYPMVLVQIPMCNEKEVYQQSIAAVCNLDW
PRSNFLVQVLDDSDDPTTQSLIREEVAKWQQTGARILYRHRVLRDGYKAGNLKSAMACSYVKDYEFV
AIFDADFQPNPDFLKRTVPHFKDNDELGLVQARWSFVNKDENLLTRLQNINLCFHFEVEQQVNGVFL
NFFGFNGTAGVWRIKALEESGGWMERTTVEDMDIAVRAHLHGWKFIFLNDVECQCELPESYEAYRK
QQHRWHSGPMQLFRLCIPDIIKSKISVWKKFNLIFLFFLLRKLILPFYSFTLFCIILPMTMFVPEAELPDW
VVCYIPALMSLLNILPSPKSFPFIIPHLLFENTMSVTKFNAMISGLFQLGSAYEWVVTKKSGRSSEGDLI
SLAAAAPPRELRHHPKTGSAPSLEALMVLKEQQPSPKKEGKKQQKKHNRIYKKELALSLLLLTAAARS
LLTKQGIHFYFLLFQGISFLLVGLDLIGEQVE
>GRMZM2G028286_P01
MAPWSGFWGGKPGGDAYRGTPVVVKMENPNWSISEISSPEDDDEDILAAAAAAGRRNKGGRTKNA
KQIRWVLLLKAHRAAGCLASLASAAVALGGAARRRVAAGRTDAEAGVVAAAGESPVVRSRFYAFIKA
FLVVSLLLLAVEVAAYFNGWDLAASALALPAIGLESLYASWLRFRATYVAPGIQFLTDACVVLFLIQSAD
RLIQCLGCFYIHIKRIKPKPRSLALPDAEDPDAGYYPMVLVQIPMCNEKEVYQQSIAAVCNLDWPKSNF
LVQVLDDSDDPLTQTLIREEVAKWQQQGARIVYRHRVLRDGYKAGNLKSAMSCSYVKEYEFVAIFDA
DFQPHPDFLKRTVPHFKDNDELGLVQARWSFVNKDENLLTRLQYINLCFHFEVEQQVNGVFLNFFGF
NGTAGVWRIKALEDSGGWLERTTVEDMDIAVRAHLHGWKFIFLNDVECQCELPESYEAYRKQQHRW
HSGPMQLFRLCLPDIIKCKIAFWKKANLIFLFFLLRKLILPFYSFTLFCIILPMTMFVPEAELPDWVVCYV
PALMSLLNILPSPKSFPFIIPYLLFENTMSVTKFNAMISGLFQLGSAYEWVVTKKSGRSSEGDLIALAPP
KEPVKHATRTGSAPNLDAVAKEEQQQQLAASRKDAAAKKKEKHNRIYKKELALSMLLLTAAARSLL
SKQGIHFYFLLFQGVSFLLVGLDLIGEQVE
>CSLC_5DL_TGACv1_435778_AA1454840.1_Triticum_aestivum
MAPWTGLWGARAGAGAGAYRGTPVVVKMENPNWSISEISPEDAEDEDFLVSGAGAARRRKGGRG
KNAKQITWVLLLKAHRAAGCLASLASAAVTLGAAARRRVADGRTDADAGATPGSAGESPVLRSRFYA
FIRAFLLLSLLLLAVELAARFHGWDLAASALALPIIGVESLYASWLRLRAAYLAPLLQFLTDACVVLFLIQ
SADRLIQCLGSFYITVKRIKPRLKSPALPDAEDPDAGYYPMVLVQIPMCNEKEVYQQSIAAVCNLDWP

RSNFLVQVLDDSDDPTTQSLIREEVAKWQQTGARILYRHRVLRDGYKAGNLKSAMACSYVKDYEFVA
IFDADFQPNPDFLKRTVPHFKDNDELGLVQARWSFVNKDENLLTRLQNINLCFHFEVEQQVNGVFLN
FFGFNGTAGVWRIKALEESGGWMERTTVEDMDIAVRAHLHGWKFIFLNDVECQCELPESYEAYRKQ
QHRWHSGPMQLFRLCIPDIIKSKISVWKKFNLIFLFFLLRKLILPFYSFTLFCIILPMTMFVPEAELPDWV
VCYIPALMSLLNILPSPKSFPFIIPYLLFENTMSVTKFNAMISGLFQLGSAYEWVVTKKSGRSSEGDLISL
AAAPPRELRHHPKTGSAPSMEALMVLKEQPSPKKEGKKQQKKHNRIYKKELALSLLLLTAAARSLLTK
QGIHFYFLLFQGISFLLVGLDLIGEQVE
>CSLC_3DL_TGACv1_251593_AA0882850.1_Triticum_aestivum
MAPWWGQEARGGVSGGVTGTPVVVKMQTPDWAISEVPPPGSPAAGGKDGRGKNARQITWVLLLK
AHRAAGKLTGAATAALSVAAAARRRVAAGRTDSDADNAPPGLGGSPALRTRLYGFIRASLLLSVLLLA
ADVAAHAQGWHLAALPDLEAVEGLFAAGYAAWMRARAAYLGPALQFLTNACVVLFMIQSADRLILCL
GCFWIKLRGIRPVPNAAAAAGNGNGKGSDDVEAGAQEEGDFPMVLVQIPMCNEKEVYQQSIGAVCN
LDWPRSNFLVQVLDDSDDAATSALIREEVEKWQREGVRVLYRHRVIRDGYKAGNLKSAMNCSYVKD
YEFVVIFDADFQPQEDFLKLTVPHFKGKEDVGLVQARWSFVNKDENLLTRLQNINLCFHFEVEQQVN
GAFLNFFGFNGTAGVWRIKALEDSGGWMERTTVEDMDIAVRAHLKGWKFLYLNDVECQCELPESYE
AYRKQQHRWHSGPMQLFRLCFVDIIKSKIGFWKKFNLIFLFFLLRKLILPFYSFTLFCVILPMTMFAPEA
ELPAWVVCYIPATMSLLNILPAPKSFPFIVPYLLFENTMSVTKFNAMISGLFQLGSAYEWVVTKKSGRS
SEGDLVALVENEKQSKQLRVGSAPNLDSLAAKEELYPKAEPKPKKKKHNRLYRKELALSFLLLTAAAR
SLLSVQGIHFYFLLFQGVSFLVVGLDLIGEQVE
>CSLC_3AL_TGACv1_197197_AA0665370.1_Triticum_aestivum
MAPWWGQEARGGVSGGVTGTPVVVKMQTPDWAISEVPPPGSPAAGGKDGRGKNARQITWVLLLK
AHRAAGKLTGAATAALSVAAAARRRVAAGRTDSDADADGAPPGPGAGRPALRTRLYGFIRASLLLSL
LLLAADVAAHAQGWHLAALPDLEAVEGLFAAGYAAWMRARAAYLGPALQFLTNACVVLFMIQSADRL
ILCLGCFWIKLRGIRPVPNAATAGNGKGSDDVEAGAQEEEEGEFPMVLVQIPMCNEKEVYQQSIGAV
CNLDWPRSNFLVQVLDDSDDAATSALIREEVEKWQREGVRILYRHRVIRDGYKAGNLKSAMNCSYVK
DYEFVVIFDADFQPQEDFLKLTVPHFKGKEDVGLVQARWSFVNKDENLLTRLQNINLCFHFEVEQQV
NGAFLNFFGFNGTAGVWRIKALEDSGGWMERTTVEDMDIAVRAHLKGWKFLYLNDVECQCELPESY
EAYRKQQHRWHSGPMQLFRLCFVDIIKSKIGFWKKFNLIFLFFLLRKLILPFYSFTLFCVILPMTMFAPE
AELPAWVVCYIPATMSLLNILPAPKSFPFIVPYLLFENTMSVTKFNAMISGLFQLGSAYEWVVTKKSGR
SSEGDLVALVENEKQSKQQRVGSAPNLDSLAAKEELYPKSEPKKKKHNRLYRKELALSFLLLTAAARS
LLSVQGIHFYFLLFQGVSFLVVGLDLIGEQVE
>CSLC_5AL_TGACv1_374268_AA1195590.3_Triticum_aestivum
MAPWTGLWGARAGAGAYRGTPVVVKMENPNWSISEISPEDAEDEDFLVSGAARRKGGRGKNAKQI
TWVLLLKAHRAAGCLASLASAAVTLGAAARRRVADGRTDADAGAPGPARESPVLRSRFYAFIRAFLLL
SLLLLAVELAARFHRWDLAASALALPIIGVESLYASWLRLRAAYLAPLLQFLTDACVVLFLIQSADRLIQ
CLGSFYITVKRIKPRLRSPALPDAEDPDAGYYPMVLVQIPMCNEKEVYQQSIAAVCNLDWPRSNFLVQ
VLDDSDDPTTQSLIREEVAKWQQTGARILYRHRVLRDGYKAGNLKSAMACSYVKDYEFVAIFDADFQ
PNPDFLKRTVPHFKDNDELGLVQARWSFVNKDENLLTRLQNINLCFHFEVEQQVNGVFLNFFGFNGT
AGVWRIKALEESGGWMERTTVEDMDIAVRAHLHGWKFIFLNDVECQCELPESYEAYRKQQHRWHS
GPMQLFRLCIPDIIKSKISVWKKFNLIFLFFLLRKLILPFYSFTLFCIILPMTMFVPEAELPDWVVCYIPALM
SLLNILPSPKSVPFIIPYLLFENTMSVTKFNAMISGLFQLGSAYEWVVTKKSGRSSEGDLISLAAAPPR
ELRQQQKTGSAPSLEALMVLKEEQASPRKEGKKQKKHNRIYKKELALSLLLLTAAARSLLTKQGIHFY
FLLFQGISFLLVGLDLIGEQVE
>CSLC_1DL_TGACv1_061928_AA0205730.1_Triticum_aestivum
MAPSFWGREARLSDGGGGTPVVVKMENPNWSISEMEQEAVPGSPAGLAAGKAGRGKNARQITWV
LLLKAHRAAGRLTGAASAALAVAAAARRRVAAGRTDGDAAPGESTALRARFYGCLRLFVVLSMLLLA
VEVAAYLQGWHLQMPEMPEMPGQLAMDGLLAVDGLAASAYAGWMRVRLQYIAPPLQFLTNSCVVL
FMIQSVDRLVLCLGCLWIKLRGIKPVPIAADKDDVEAGDEDFPMVLVQMPMCNEREVYQQSIGAICAL
DWPRSNFLVQVLDDSDDATTSALIKEEVEKWQREGVRIVYRHRVIRDGYKAGNLKSAMNCSYVKDY
EYVVIFDADFQPQADFLKRAMPHFKGKDDVGLVQARWSFVNNDENLLTRLQNINLCFHFEVEQQVN
GAFLNFFGFNGTAGVWRIKALEDSGGWMERTTVEDMDIAVRAHLKGWKFLYLNDVECQCELPESYE
AYRKQQHRWHSGPMQLFRLCFVDIIKSKIGFWKKCNLIFLFFLLRKLILPFYSFTLFCVILPMTMFVPEA
ELPAWVVCYIPATMSIMSILPSPKSFPFIVPYLLFENTMSVTKFNAMISGLFQLGSAYEWVVTKKSGRS
SEGDLVALVEKHTVQQQQRVGSAPDLAGLAAKDSSLPKKDAPKKKQKHNRIYRKELALSFLLLTAAA
RSVLSAQGIHFYFLLFQGVSFLVMGLDLIGEQVE
>CSLC_1BL_TGACv1_030750_AA0099830.1_Triticum_aestivum
MAPSFWGREARLSDGGGGTPVVVKMENPNWSISEMEQEAVPGSPAGLAAGKAGRGKNARQITWV
LLLKAHRAAGRLTGAASAALAVAAAARRRVAAGRTDGDAAPGESTALRARFYGCLRLFVVLSMLLLA
VEVAAYLQGWHLQMPEMPEMPGQLAMDGLLAVDGLAAAAYAGWMRVRLQYIAPPLQFLTNSCVVL
FMIQSVDRLILCLGCLWIKLRGIKPVPIAADKDDVEAGEEDFPMVLVQMPMCNEREVYQQSIGAICALD
WPRSNFLVQVLDDSDDATTSALIKEEVEKWQREGVRIVYRHRVIRDGYKAGNLKSAMNCSYVKDYEY
VVIFDADFQPQADFLKRAMPHFKGKDDVGLVQARWSFVNNDENLLTRLQNINLCFHFEVEQQVNGA

FLNFFGFNGTAGVWRIKALEDSGGWMERTTVEDMDIAVRAHLKGWKFLYLNDVECQCELPESYEAY
RKQQHRWHSGPMQLFRLCFVDIIKSKIGFWKKCNLIFLFFLLRKLILPFYSFTLFCVILPMTMFVPEAEL
PAWVVCYIPATMSIMSILPSPKSFPFIVPYLLFENTMSVTKFNAMISGLFQLGSAYEWVVTKKSGRSSE
GDLVALVEKHTVQQQQRVGSAPDLAGLAAKDSSLPKKDAPKKKQKHNRIYRKELALSFLLLTAAARS
VLSAQGIHFYFLLFQGVSFLVMGLDLIGEQVE
>CSLC_1AL_TGACv1_001272_AA0028090.1_Triticum_aestivum
MAPSFWGREARLSDGGGGTPVVVKMENPNWSISEMEQEPVPGSPAGLAAGKAGRGKNARQITWV
LLLKAHRAAGRLTGAASAALAVAAAARRRVAAGRTDGDAAPGESTALRARFYGCLRVFVVLSMLLLA
VEVAAYLQGWHLQMPQMPEMPGQLAMDGLLAVDGLAAAAYAGWMRVRLQYIAPPLQFLTNSCVVL
FMIQSVDRLVLCLGCLWIKLRGIKPVPIAADKDDVEAGDEDFPMVLVQMPMCNEREVYQQSIGAICAL
DWPRSNFLVQVLDDSDDATTSALIKEEVEKWQREGVRIVYRHRVIRDGYKAGNLKSAMNCSYVKDY
EYVVIFDADFQPQADFLKRAMPHFKGKDDVGLVQARWSFVNNDENLLTRLQNINLCFHFEVEQQVN
GAFLNFFGFNGTAGVWRIKALEDSGGWMERTTVEDMDIAVRAHLKGWKFLYLNDVECQCELPESYE
AYRKQQHRWHSGPMQLFRLCFVDIIKSKIGFWKKCNLIFLFFLLRKLILPFYSFTLFCVILPMTMFVPEA
ELPAWVVCYIPATMSIMSILPSPKSFPFIVPYLLFENTMSVTKFNAMISGLFQLGSAYEWVVTKKSGRS
SEGDLVALVEKHTAQQQQRVGSAPDLAGLAAKDSSLPKKDAPKKKQKHNRIYRKELALSFLLLTAAA
RSVLSAQGIHFYFLLFQGVSFLVMGLDLIGEQVE
>CSLF_2DS_TGACv1_178471_AA0596060.1_Triticum_aestivum
MIYTMNSIISILAADYPVDKHACYLSDDGGSIIHYDGLLETAKFAALWVPFCRKHSIEPRAPESYFSLNT
RPYTGNAPQDFVNDRRHMCREYDEFKERLDALFTLIPKRSDVYNHAAAKEGAKATWMADGTQWPG
TWIDPAENHKKGQHVGIVKVMLKHPSYEPELGLGASTNSPLDFSAVDVRLPMLVYISREKSPSCDHQ
KKAGAMNVQLRVSALLTNAPFIINFDGDHYVNNSKAFRAGICFMLDRREGDNTAFVQFPQRFDDVDP
TDRYCNHNRVFFDATLLGLNGIQGPSYVGTGCMFRRVALYGVDPPRWRPDDVKIVDSSTKFGSSAS
FISSILPAADQERSIMSPPALEESVMADLAHVMTCAYEDGTEWGSDVGWVYNIATEDVVTGFRLHRN
GWRSMYCRMEPDAFAGTAPINLTERLYQILRWSGGSLEMFFSRNCPLLAGRRLHPMQRIAYANMTA
YPVSSVFLVFYLLFPVIWIFRGQFYIQKPFPTYVLYLVIVIALTELIGMVEIKWAGLTLLDWIRNEQFYIIGA
TAVYPTAVFHIVLKLFGLKGVSFKLTAKQVASSTSDKFAELYAVQWAPMLIPTMVVIAVNVCAIGASIGK
AIVGGWSLMQMADAGLGLVFNAWILVLIYPFALGMIGRWSKRPYILFILFVIAFILIALVDIAIQAMRSGFV
RFHFKSSGGATFPTSWGL
>CSLF_2BS_TGACv1_148916_AA0495580.1_Triticum_aestivum
MIYTMNSIISILAADYPVDKHACYLSDDGGSIIHYDGLLETAKFAALWVPFCRKHSIEPRAPESYFSLNT
RPYTGNAPQDFVNDRRHMCREYDEFKERLDALFTLIPKRSDVYNHAAAKEGAKATWMADGTQWPG
TWIDPAENHKKGQHVGIVKVMLKHPSYEPELGLGASTNSPLDFSAVDVRLPMLVYISREKSPSCDHQ
KKAGAMNVQLRVSALLTNAPFIINFDGDHYVNNSKAFRAGICFMLDRREGDNTAFVQFPQRFDDVDP
TDRYCNHNRVFFDATLLGLNGIQGPSYVGTGCMFRRVALYGVDPPRWRPDDVKIVDSSTKFGSSAS
FISSILPAADQERSIMSPPALEEPVMADLAHVMTCAYEDGTEWGREVGWVYNIATEDVVTGFRLHRN
GWRSMYCRMEPDAFAGTAPINLTERLYQILRWSGGSLEMFFSRNCPLLAGRRLHPMQRIAYANMTA
YPVSSVFLVFYLLFPVIWIFRGQFYIQKPFPTYVLYLVIVIALTELIGMVEIKWAGLTLLDWIRNEQFYIIGA
TAVYPTAVFHIVLKLFGLKGVSFKLTAKQVASSTSDKFAELYAVQWAPMLIPTMVVIAVNVCAIGASIGK
AIVGGWSLMQMADAGLGLVFNAWILVLIYPFALGMIGRWSKRPYILFILFVIAFILIALVDIAIQAMRSGFV
RFHFKSSGGATFPTSWGL
>GRMZM2G173759_P01
MAPGLRLAYPWGRDVRRGTPVVVTMENPNYSVVEIDGPEAEALRAGVPPMDKGRGRGRSAKQFT
WVLLLRAHRAAGCLASLAAVTWALPSAVAKRFRRAAAAEGVGRGRGWLLYRFIKVLLALSLLALTVEL
AAYWKGWHFQRPNLTMPEFHVPEVEDIQGWLHTAYLAWMSFRADYIRRPIEFLSKVCILLFVVQSLD
RLVLCIACFWIKLKKIKPRLEGDPFREGSGYLHPMVLVQIPMCNEKEVYEQSISAVCQLDWPRDKFLIQ
VLDDSSDESIQMLIKAEVSKWNQQGVNIVYRHRVLRTGYKAGNLNSAMSCDYVKNFEFVAIFDADFQ
PSTDFLKKTIPHFDGNPELGLVQARWSFVNKDENLLTRLQNINLCFHFEVEQQVNGVFLNFFGFNGTA
GVWRIQALEESGGWLERTTVEDMDIAVRAHLNGWKFIFLNDVKVLCEVPESYEAYRKQQHRWHSGP
MHLFRLCFPDIITAKISSWKKANLILLFFLLRKLILPFYSFTLFCVILPLTMFVPEAELPVWVICYVPICMSF
LNILPSPRSFPFIVPYLLFENTMSVTKFNAMVSGLFKLGSSYEWVVTKKSGRSSELDLLTSEEKNRKCT
TLPQLQNQLPENSELVEINVRKEQREKVPNDAKRGNKIYKKELALSLLLLTAATRSLLSAQGIHFYFLLF
QGMSFLAVGLDLIGEQIS
>AT4G07960_CslC12
MAPKFEWWAKGNNNNTRKGTPVVVKMENPNNWSMVELESPSHDDFLVRTHEKSRNKNARQLTWV
LLLKAHRAAGCLTSLGSALFALGTAVRRRIAAGRTDIEISSSGVGSLQKQNHTKKSKLFYSCLKVFLWL
SLILLGFEIAAYFKGWSFGTSKLQLQFIFNKGFFDWVYTRWVLLRVEYLAPPLQFLANGCIVLFLVQSL
DRLILCLGCFWIRFKKIKPVPKPDSISDLESGDNGAFLPMVLVQIPMCNEKEVYQQSIAAVCNLDWPK
GKILIQILDDSDDPITQSLIKEEVHKWQKLGARIVYRHRVNREGYKAGNLKSAMNCSYVKDYEFVAIFD
ADFQPLPDFLKKTIPHFKDNEEIGLVQARWSFVNKEENLLTRLQNINLAFHFEVEQQVNSVFLNFFGF
NGTAGVWRIKALEDSGGWLERTTVEDMDIAVRAHLHGWKFVFLNDVECQCELPESYEAYRKQQHR
WHSGPMQLFRLCLPAVIKSKISIGKKFNLIFLFFLLRKLILPFYSFTLFCIILPMTMFVPEAELPAWVVCYI

PATMSFLNILPAPKSFPFIVPYLLFENTMSVTKFNAMVSGLFQLGSAYEWVVTKKSGRSSEGDLAALV
EKDEKTTKHQRGVSAPETEAEKKAEKTKRKKKKHNRIYMKELSLAFLLLTAATRSLLSAQGIHFYFLLF
QGISFLLVGLDLIGEQVE
>OS09G25900_OsCSLC2
MAPPGVGVGVAYLWGKGRGGRKGTPVVVTMESPNYSVVEVDGPDAEAELRTAAVAMDKGGGRGR
SRSRTARQLTWVLLLRARRAAGRLASFAAAAARRFRRSPADAADELGRGRGRLMYGFIRGFLALSLL
ALAVELAAYWNGWRLRRPELHVPEAVEIEGWAHSAYISWMSFRADYIRRPIEFLSKACILLFVIQSMD
RLVLCLGCFWIKLRKIKPRIEGDPFREGSGYQHPMVLVQIPMCNEKEVYEQSISAACQLDWPREKFLI
QVLDDSSDESIQLLIKAEVSKWSHQGVNIVYRHRVLRTGYKAGNLKSAMSCDYVKDYEFVAIFDADFQ
PTPDFLKKTIPHFEGNPELGLVQARWSFVNKDENLLTRLQNINLCFHFEVEQQVNGVFLNFFGFNGTA
GVWRIQALEESGGWLERTTVEDMDIAVRAHLNGWKFIFLNDVKVLCELPESYEAYRKQQHRWHSGP
MHLFRLCLPDILTAKISSWKKANLILLFFLLRKLILPFYSFTLFCVILPLTMFVPEAELPVWVICYVPVCMS
FLNILPSPRSFPFIVPYLLFENTMSVTKFNAMVSGLFKLGSSYEWIVTKKSGRSSESDLSTAAERDTKD
LTLPRLQKQISESELIELKMQKERQEKAPLGAKKANKVYKKELALSLLLLTAATRSLLSAQGIHFYFLLF
QGVSFLFVGLDLIGEQID
>AT4G31590_CslC5
MAPRLDFSDWWAKDTRKGTPVVVKMENPNYSVVEIDGPDSAFRPVEKSRGKNAKQVTWVLLLKAH
RAVGCLTWLATVFWSLLGAIKKRLSFTHPLGSEKLGRDRWLFTAIKLFLAVSLVILGFEIVAYFRGWHY
FQSPSLHIPTSTLEIQSLFHLVYVGWLTLRADYIAPPIKALSKFCIVLFLIQSVDRLVLCLGCFWIKYKKIK
PRFDEEPFRNDDAEGSGSEYPMVLVQIPMCNEREVYEQSISAVCQLDWPKDRILVQVLDDSNDESIQ
QLIKAEVAKWSQKGVNIIYRHRLVRTGYKAGNLKSAMSCDYVEAYEYVAIFDADFQPTPDFLKLTVPH
FKDNPELGLVQARWTFVNKDENLLTRLQNINLCFHFEVEQQVNGVFLNFFGFNGTAGVWRIKALEES
GGWLERTTVEDMDIAVRAHLHGWKFIYLNDVKVLCEVPESYEAYKKQQHRWHSGPMQLFRLCLGSI
LTSKIAIWKKANLILLFFLLRKLILPFYSFTLFCIILPLTMFVPEAELPVWVICYIPVFMSFLNLLPSPKSFPF
IVPYLLFENTMSVTKFNAMVSGLFQLGSSYEWIVTKKAGRSSESDLLSITEKETPTKKSQLLRGVSDSE
LLELSQLEEQKQAVSKKPVKKTNKIYHKELALAFLLLTAALRSLLAAQGVHFYFLLFQGVTFLLVGLDLI
GEQMS
>OS01G56130_OsCslC1
MARWWGGEGRGGSGTPVVVKMESPEWAISEVEAGAAAPGSPAAGGKAGRGKNARQITWVLLLKA
HRAAGKLTGAASAALSVAAAARRRVAAGRTDSDDAAAAPPGESPALRARFHGFLRAFLLLSVLLLAV
DVAAHAQGWHAVVPDLLAVEGLFAAAYASWLRVRLEYLAPGLQFLANACVVLFLIQSADRLILCLGCL
WIKLKGIKPVPKASGGGGGGKGSDDVEAGADEFPMVLVQIPMCNEKEVYQQSIGAVCNLDWPRSNF
LVQVLDDSDDAATSALIKEEVEKWQREGVRILYRHRVIRDGYKAGNLKSAMNCSYVKDYEFVVIFDAD
FQPQADFLKRTVPHFKGNEDVGLVQARWSFVNKDENLLTRLQNINLCFHFEVEQQVNGVFLNFFGF
NGTAGVWRIKALEDSGGWMERTTVEDMDIAVRAHLKGWKFLYINDVECQCELPESYEAYRKQQHR
WHSGPMQLFRLCFVDIIKSKIGVWKKFNLIFLFFLLRKLILPFYSFTLFCIILPMTMFVPEAELPAWVVCYI
PATMSLLNILPAPKSFPFIVPYLLFENTMSVTKFNAMISGLFQLGSAYEWVVTKKSGRSSEGDLVSLVE
KQPKQQRVGSAPNLDSLAKESHPKKDSKKKKHNRIYQKELALSFLLLTAAARSLLSVQGIHFYFLLFQ
GVSFLVVGLDLIGEQVE
>CSLC_1DL_TGACv1_062162_AA0209740.1_Triticum_aestivum
MAPWNGLWGGRAAIAGGNAYRDMPVIVKMENPNWSISEINGGGDNGEDFLARVGGQRRRVKNTKQ
ITWVFRLKAHRAAGCLARLTSAAVALGGAARRRVVAGRTDSDAADGECEDVEERDPASRRSRFYTLI
KACLMMSVFLLVVELAAYSNGRVNLAIFINSFNTSWIRFRATYVAPPLQLLADACVVLFLVQSADRLFQ
SLGCFYILVKRIKPKPLSPALADAEDPDAGYYPMVLVQIPMCNEKEVYRQSIAAVCNLDWPRSNFLVQ
VLDDSDDVATQALIKEEVEKWRHSGAHIVYRHRVLREGYKAGNLKSAMSCSYVKDYEYVAIFDADFQ
PYPDFLKRTVPHFKDNEDLGLVQARWSFVNKDENLLTRLQNINLCFHFEVEQQVNGVFINFFGFNGT
AGVWRIKAVEDSGGWMERTTVEDMDIAVRAHLKGWKFVFLNDVECQCELPESYEAYRKQQHRWHS
GPMQLFRLCLPDIIRCKIVFWKKANLIFLFFLLRKLILPFYSFTLFCIILPMTMFVPEAELPDWVVCYIPVL
MSFLNIAPAPKSFPFIIPYLLFENTMSVTKFNAMISGLFQLGSTYEWVVTKKSGRSLEGDLISLAPKGLK
QLKYGSVPAINVAIKEQSKAKKESKKYNRIYKKELAMSLLLLSAAARSLLSKQGIHFYFLLFQGISFLLV
GLDLIGQDIK
>AT2G24630_CslC8
MAPRFDFSDLWAKETRRGTPVVVKMENPNYSIVEVEEPDSAFQPMEKSRGKNAKQVTWVLLLKAHK
AVGCLTWVATVFWSLLGSVKRRLSFTHPLGSERLGRDGWLFSAIKLFLVASLAILAFELVAYYRGWHY
FKNPNLHIPTSKLEIQSLLHLFYVGWLSLRADYIAPPIKALSKFCIVLFLVQSVDRLILCLGCLWIKFKKIK
PRIDEEHFRNDDFEGSGSEYPMVLVQIPMCNEREVYEQSISAVCQLDWPKDRLLVQVLDDSDDESIQ
ELIRDEVTKWSQKGVNIIYRHRLVRTGYKAGNLKSAMSCDYVEAYEFVAIFDADFQPNSDFLKLTVPH
FKEKPELGLVQARWAFVNKDENLLTRLQNINLCFHFEVEQQVNGVFLNFFGFNGTAGVWRIKALEES
GGWLERTTVEDMDIAVRAHLHGWKFIYLNDVKVLCEVPESYEAYKKQQHRWHSGPMQLFRLCLRSI
LTSKIAMWKKANLILLFFLLRKLILPFYSFTLFCVILPITMFVPEAELPIWVICYVPIFMSLLNILPAPKSFPF
IVPYLLFENTMSVTKFNAMVSGLFQLGSSYEWIVTKKAGRSSESDLLALTDKESEKMPNQILRGVSDS

ELLEISQVEEQKKQPVSVKKTNKIFHKELALAFLLLTAAVRSLLASQGVHFYFLLFQGLTFLLVGLDLIG
EQMS
>OS05G43530_OsCslC7
MAPSWWGRSGGGGVGNGGGTPVVVKMENPNWSISEVEAAEVAPGSPAGAGKAGRGKNARQITW
VLLLKAHRAAGRLTGAASAALAVASAARRRVASGRTDADAAPGESTALRARSYGCIRVSLVLSLLLLA
VEVAAYLQGWHLEEVASLLAVDGLFAASYAGWMRLRLDYLAPPLQFLTNACVALFMVQSIDRLVLCL
GCFWIRFKGIKPVPQAAAAGKPDVEAGAGDYPMVLVQMPMCNEREVYQQSIGAVCNLDWPKSNFLV
QVLDDSDDATTSALIKEEVEKWQREGVRIIYRHRVIRDGYKAGNLKSAMNCSYVKDYEFVVIFDADFQ
PQADFLKRTVPHFKGKDDVGLVQARWSFVNKDENLLTRLQNVNLCFHFEVEQQVNGAFLNFFGFNG
TAGVWRIKALEDSGGWMERTTVEDMDIAVRAHLKGWKFVFLNDVECQCELPESYEAYRKQQHRWH
SGPMQLFRLCFVDIIKSKIGFWKKFNLIFLFFLLRKLILPFYSFTLFCVILPMTMFVPEAELPAWVVCYIPA
TMSILNILPAPKSFPFIVPYLLFENTMSVTKFNAMISGLFQLGSAYEWVVTKKSGRSSEGDLVGLVEKH
SKQQRVGSAPNLDALTKEESNPKKDSKKKKHNRIYRKELALSFLLLTAAARSLLSAQGIHFYFLLFQGV
SFLVVGLDLIGEQVE
>GRMZM2G027794_P01
MPQWWGREARGGGGGGGTSVVVKMESPDWAVREPEAARGKAGRGKNARQITWVLLLKAHRAAG
KLTGAASAALSVAAAARRRVAAGRTDADADAGAGAGAAPHGESPALRTRLYGFLRASLVLSMLLLAA
DVAAHLQGWHLLAVDVPDLLAVEGLFAAGYASWVRVRLQYLAPALQFLANACVLLFLVQSADRLILCL
GCLWIKLRGIKPVPNAAGKGPDDVEAGTGEFPMVLVQIPMCNEKEVYQQSIGAVCGLDWPRPNFLV
QVLDDSDDAATSALIKEEVERWQREGVRILYRHRVIRDGYKAGNLKSAMNCSYVKDYEFVVIFDADF
QPQPDFLKRTVPHFKGNEDVGLVQARWSFVNKDENLLTRLQNINLCFHFEVEQQVNGIFLNFFGFNG
TAGVWRIKALEDSGGWMERTTVEDMDIAVRAHLKGWKFLFLNDVECQCELPESYEAYRKQQHRWH
SGPMQLFRLCFVDIIKSKIGFWKKFNLIFLFFLLRKLILPFYSFTLFCIILPMTMFIPEAELPAWVVCYIPAT
MSLLNILPAPKSFPFIVPYLLFENTMSVTKFNAMVSGLFQLGSAYEWVVTKKSGRSSEGDLIALVEKQP
KQQRVGSAPNLESLTKESSSLKRDSKRKKHNRIYRKELALSFLLLTAAARSLLSVQGIHFYFLLFQGVS
FLVVGLDLIGEQVE
>OS07G03260_OsCslC10
MAPWSGFWAASRPALAAAAAGGTPVVVKMDNPNWSISEIDADGGEFLAGGRRRGRGKNAKQITWV
LLLKAHRAAGCLAWLASAAVALGAAARRRVAAGRTDDADAETPAPRSRLYAFIRASLLLSVFLLAVEL
AAHANGRGRVLAASVDSFHSSWVRFRAAYVAPPLQLLADACVVLFLVQSADRLVQCLGCLYIHLNRI
KPKPISSPAAAAAALPDLEDPDAGDYYPMVLVQIPMCNEKEVYQQSIAAVCNLDWPRSNILVQVLDDS
DDPITQSLIKEEVEKWRQNGARIVYRHRVLREGYKAGNLKSAMSCSYVKDYEYVAIFDADFQPYPDFL
KRTVPHFKDNEELGLVQARWSFVNKDENLLTRLQNINLCFHFEVEQQVNGIFINFFGFNGTAGVWRIK
ALEDSGGWMERTTVEDMDIAVRAHLNGWKFVFLNDVECQCELPESYEAYRKQQHRWHSGPMQLF
RLCLPDIIRCKIAFWKKANLIFLFFLLRKLILPFYSFTLFCIILPMTMFIPEAELPDWVVCYIPALMSFLNILP
APKSFPFIIPYLLFENTMSVTKFNAMISGLFQLGSAYEWVVTKKSGRSSEGDLIALAPKELKQQKILDLT
AIKEQSMLKQSSPRNEAKKKYNRIYKKELALSLLLLTAAARSLLSKQGIHFYFLMFQGLSFLLVGLDLIG
EDVK
>AC183932.3
MAPCSGLWGGKAGGDAYRGTPVVVKMENPNWSISEISSPEDDDDDEDILAAAGGRRRKGARTKNA
KQIRWVLLLKAHRAAGCLASLASAAVALGAAARRRVAAGRTDADADADAGVLAVAGESPVVRSGFY
AFIRAFLVVSLLLLAVEVAAYINGWDLAASALALPALGLESLYASWLRFRAAYVAPGIQFLTDACVVLFL
VQSADRLILCLGCFYIRVKRIKPEPKSPALPDAEDPDAGYYPMVLVQIPMCNEKEVYQQSIAAVCNLD
WPKSSFLVQVLDDSDDLLTQALIREEVAKWQQQGARIVYRHRVLRDGYKAGNLKSAMSCSYVKDYE
FVAIFDADFQPQPDFLKRTVLHFKDNDELGLVQARWSFVNKDENLLTRLQYINLCFHFEVEQQVNGV
FLNFFGFNGTAGVWRIKALEDSGGWMERTTVEDMDIAVRAHLHGWKFIFLNDVECQCELPESYEAY
RKQQHRWHSGPMQLFRLCLPDIIKCKMAFWKKGNLILLFFLLRKLILPLYSFTLFCIILPTAMFVPEAEL
PDWVVCYVPALMSLLNVLPSPRSFPFVIPYLLFENTMSVTKFNAMVSGLFQLGSAYEWVVTKKSGSG
PRSSEVAGDLVSLAAAAPTAKKKKKKKHNRIYKKELALSMLLLTAAARSLLSKQGIHFYFLLFQGVSFL
LVGLDLIGEQVE
>AT3G07330_CslC6
MSRSQNEEFQQWWNKQRDRNNHDVLYAGDDEAFLTVEIRTPATVDPDKDRIRTRTVRQLSRLYLLK
FKQLASSFLWIGNSFLYLVRTANRRIANDNPPSVSSSARLYRLIKGFLVVVVLLLCFELAAYFKGWHFT
PPSVASAEVAVEVVYAWWLEIRASYLAPPLQSLTNVCIVLFLIQSVDRLVLVLGCFWIKLRRIKPVASM
EYPTKLVGEGVRLEDYPMVIVQIPMCNEKEVYQQSIGAVCMLDWPRERMLVQVLDDSSELDVQQLIK
AEVQKWQQRGVRIVYRHRLIRTGYKAGNLKAAMNCEYVKDYEFVAIFDADFQPPADFLKKTVPHFKG
NEELALVQTRWAFVNKDENLLTRLQNINLSFHFEVEQQVNGVFINFFGFNGTAGVWRIKALEDCGGW
LERTTVEDMDIAVRAHLCGWKFIYLNDVKCLCELPESYEAYKKQQYRWHSGPMQLFRLCFFDILRSK
VSAAKKANMIFLFFLLRKLILPFYSFTLFCVILPLTMFFPEANLPSWVVCYIPGIMSILNIIPAPRSFPFIVP
YLLFENTMSVTKFGAMISGLFKFDSSYEWVVTKKLGRSSEADLVAYAESGSLVESTTIQRSSSDSGLT
ELSKLGAAKKKAGKTKRNRLYRTEIALAFILLAASVRSLLSAQGIHFYFLLFQGITFVIVGLDLIGEQVS
>AT3G28180_CslC4

MAPNSVAVTMEKPDNFSLLEINGSDPSSFPDKRKSISPKQFSWFLLLKAHRLISCLSWLVSSVKKRIAF
SAKNINEEEDPKSRGKQMYRFIKACLVISIIALSIEIVAHFKKWNLDLINRPSWEVYGLVEWSYMAWLSF
RSDYIAPLVISLSRFCTVLFLIQSLDRLVLCLGCFWIKFKKIEPKLTEESIDLEDPSSFPMVLIQIPMCNER
EVYEQSIGAASQLDWPKDRILIQVLDDSDDPNLQLLIKEEVSVWAEKGVNIIYRHRLIRTGYKAGNLKS
AMTCDYVKDYEFVTIFDADFTPNPDFLKKTVPHFKGNPELGLVQARWSFVNKDENLLTRLQNINLCFH
FEVEQQVNGVFLNFFGFNGTAGVWRIKALEESGGWLERTTVEDMDIAVRAHLNGWKFIYLNDVEVTC
ELPESYEAYKKQQHRWHSGPMQLFRLCLPSIIKSKISVWKKANLIFLFFLLRKLILPFYSFTLFCIILPLTM
FIPEAELPLWIICYVPIFISLLNILPSPKSFPFLVPYLLFENTMSITKFNAMISGLFQFGSAYEWVVTKKTG
RSSESDLLAFAEKEEKLHRRNSESGLELLSKLKEQETNLVGQETVKKSLGGLMRPKNKKKTNMVFKK
ELGLAFLLLTAAARSFLSAHGLHFYFLLFQGLSFLVVGLDLIGEQIS
>CSLF_2BL_TGACv1_130934_AA0420130.1_Triticum_aestivum
MVQVASFAALWVLFCRKHCVEPRSPESYFGMKTRSYAGGMAGEFMRDHRRVRREYEEFKVRIDSL
STTIRQRSDAYNSNNKGGVSATWMADGTQWPGTWVEQAENHRRGQHAGIVQVLLDHPSFKPQLGS
PASTDNPFDFSNVDTRLPMLVYISREKRPGYDNQKKAGAMNVMLRVSALLSNAPFVINFDCDHYINN
SQALRAPMCFMLDPHDGQNTAFVQFPQRFDDVDPTDRYANHNRVFFDGTMLALNGLQGPTYLGTG
TMFRRVALYGIEPPHYRAENTKLVCKTGEFGYSTSFINSVPDAAIQDRSITPVLVDERLSKDLATLMTC
AYEDGSSWGRDAGWVYNIATEDVVTGFRIHRQGWHSMYCSMEPAAFRGTAPINLTERLYQVLRWS
GGSLEVFFSHNNALIASRRLHPLQRITYLNMSTYPIVTVFILSYNFFPVMWLFSEQLYIQRPFGTYMAYL
VGIIAMVHLIGMFEVRWSGITLLDWFRNEQFYMIGATGVYPTAVLYMLLKLVTGKGIYFRLTSKQTEGC
SNDKFADLYTVRWVPLLIPTAAVIIVNVAAIGAAIGKAATWGFFTDEARHALLGMVFNMGILVLLYPFAL
GIMGKWGKRPIILFIVLVMAISVVGLLYVTLHAPYTGEWSQVAVSLGKASLTGPSGSG
>CSLC_1BL_TGACv1_030501_AA0092480.1_Triticum_aestivum
MENPNWSISEINIDDDNSEDFLARVGGQRRRVKNTKQITWVFRLKAHRAAGCLSWLTSAAFALGGAT
RRRVVAGRTDSNATDGECKDVEEWAPASRRSRFYTLIKACLMMFVCLLIVELAAYSNGKGNLAVFIN
SFNTSWIRFRAAYIAPPLQLLANACVVLFLVQSADRVFQSLGCFYILVKRIKPKPLFLALSDAEDPDAGY
YPMVLVQIPMCNEKEVYRQSIAAVCNLDWPRSNFLVQVLDDSDDVTTQALIKDEVEKWRHSGAHIVY
RHRVLREGYKAGNLKSAMSCSYVKDYEYVAIFDADFQPYPDFLKRTVPHFKDNEDLGLVQARWSFV
NKDENLLTRLQNINLCFHFEVEQQVNGVFINFFGFNGTAGVWRIKAVEDSGGWMERTTVEDMDIAG
WKFVFLNDVECQCELPETYEAYRKQQHRWHSGPMQLFRLCLPDIIRCKIVFWKKANLIFLFFLLRKLIL
PFYSFTLFCIILPMTMFVPEAELPDWVVCYIPVLMSFLNIAPAPKSFPFIIPYLLFENTMSVTKFNAMISG
LFQLGSTYEWVVTKKSGRSLEGDLISLAPKGLKQLKYGSVPAINVAIKEKLKAKKESKKYNRIYKKELA
MSLLLLSAAIRSLLSKQGIHFYFLLFQGISFLLVGLDLIGQDIK
>CSLF_7BL_TGACv1_580651_AA1914920.1_Triticum_aestivum
MATDTVADAAEGRRARDDVWVAAEEGDMPEASAGRPLLFRTMKVKGSILHPYRFLILVRLVAIVAFFA
WRVEHRNHDGVWLWATSMVADAWFGFSWLLNQLPKLNPIKRVPDLAALADRHGEAVLPGIDVFVTT
VDPVDEPVMYTVNTILSILAADYPVDKYACYLSDDGGTLVHYEAMLQVASFAALWVPFCRKHCVEPR
SPENYFGMKTRPYVGGMAGEFMSDHRRVRREYGEFKVRIDSLSSTIRRRSDAYNKGDDDVHATWM
ADGTQWPGTWIEQADNHRRGQHAGIVKVMLDHPSCKPQLGSSASTNKPVDLSNVDTRLPMLVYISR
EKRPGYDNQKKAGAMNVMLRVSALLSNAPFVINFDCDHYINNSQALRAPMCFMLDPRDGQNTAFVQ
FPQRFDDVDPTDRYSNHNRVFFDGTMLSLNGLQGPTYLGTGTMFHRVALYGMEPQRYRAENIKLVG
KGAELGKSTPFLNSIPDGAIQDRSITPVSVDEGLMSDLATLMTCAYEDRTSWGRDVGWVYNIATEDV
VTGFRIHRQGWRSMYCSMEPAAFRGTAPINLTERLYQVLGGQAAPSRCSSPTATLSSPAVGSTLYSA
SLTSTCRSTRSPRCSS
>OS09G30120_OsCslE1
MAYNYPSEKISVYLSDDGGSILTFYALWEASIFAKKWLPFCKRYNIEPRSPAAYFSESKVHHNLCIPKE
WALIKNLYEEMRERIDTATMSGKIPEEMKLKHKGFDEWNSDFTLKNHQPIVQILIDGKNRNAIDDDRNV
LPTMVYVAREKRPQYHHNFKAGALNALIRVSSVISDSPVILNVDCDMYSNNSDSIRDALCFFLDEEMG
QKIGFVQYPQIFNNMTQNDIYGNSFNVSYHVEMCGLDSVGGCLYIGTGCFHRREILCGRIFSKDYKEN
WNRGIKERGKENINEIEEKATSLVTCTYEHRTQWGNDIGVKYGFPAEDIITGLAIHCRGWESAFINPKR
AAFLGLAPSTLAQNILQHKRWSEGNLTIFLSKYCSFLFGHGKIKLQLQMGYCICGLWAANSLPTLYYVV
IPSLGLVKGTPLFPQIMSPWATPFIYVFCVKTLYGLYEALLSGDTLKGWWNGQRMWMVKSITSYLYGF
IDTIRKCVGMSKMSFEVTAKVSGHDEAKRYEQEILEFGSSSPEYVIIATVALLNFVCLVGGLSQIMAGV
WNMPWNVFLPQAILCGMIVIINMPIYEAMFLRKDNGRIPTAVTLASIGFVMLAFLVPIV
>DAA01745_OsCslA10
MEGQWGRWRLAAAAAASSSGDQIAAAWAVVRARAVAPVLQFAVWACMAMSVMLVLEVAYMSLVS
LVAVKLLRRVPERRYKWEPITTGSGGVGGGDGEDEEAATGGREAAAFPMVLVQIPMYNEKEVYKLSI
GAACALTWPPDRIIIQVLDDSTDPAIKDLVELECKDWARKEINIKYEIRDNRKGYKAGALKKGMEHIYTQ
QCDFVAIFDADFQPESDFLLKTIPFLVHNPKIGLVQTRWEFVNYDVCLMTRIQKMSLDYHFKVEQESG
SSMHSFFGFNGKIQYQNYLHELFMPGPNIADLPTGTAGVWRVSAINEAGGWKDRTTVEDMDLAVRA
SLKGWQFLYVGDIRVKSELPSTFKAYRHQQHRWTCGAANLFRKMXQRKLPKTRQAFFNXTEWNML
RDCHAIXFFTRLYVIQGVSVWKKLHLLYSFFFVRRVVAPILTFLFYCVVIPLSVMVPEVSIPVWGMVYIP

TAITIMNAIRNPGSIHLMPFWILFENVMAMHRMRAALTGLLETMNVNQWVVTEKVGDHVKDKLEVPLL
EPLKPTDCVERIYIPELMVAFYLLVCASYDLVLGAKHYYLYIYLQAFAFIALGFGFAGTSTPCS
>OS08G33740_OsCslA11
MSSSGGGGVAEEVARLWGELPVRVVWAAVAAQWAAAAAAARAAVVVPAVRALVAVSLAMTVMILA
EKLFVAAVCLAVRAFRLRPDRRYKWLPIGAAAAAASSEDDEESGLVAAAAAFPMVLVQIPMFNEREV
YKLSIGAACSLDWPSDRVVIQVLDDSTDLVVKVFIVIYFTDISSRIIRSTSSLVIKDLVEKECQKWQGKG
VNIKYEVRGNRKGYKAGALKEGLKHDYVKECEYIAMFDADFQPESDFLLRTVPFLVHNSEIALVQTRW
KFVNANECLLTRFQEMSLDYHFKYEQEAGSSVYSFFGFNGTAGVWRIAAIDDAGGWKDRTTVEDMD
LAVRATLQGWKFVYVGDVKVKSELPSTFKAYRFQQHRWSCGPANLFKKMMVEILENKKVSFWNKIH
LWYDFFFVGKIAAHTVTFIYYCFVIPVSVWLPEIEIPLWGVVYVPTVITLCKAVGTPSSFHLVILWVLFEN
VMSLHRIKAAVTGILEAGRVNEWVVTEKLGDANKTKPDTNGSDAVKVIDVELTTPLIPKLKKRRTRFW
DKYHYSEIFVGICIILSGFYDVLYAKKGYYIFLFIQGLAFLIVGFDYIGVCPP
>OS07G43710_OsCslA7
MVEAGEIGGAAVFALAAAAALSAASSLGAVDFRRPLAAVGGGGAFEWDGVVPWLIGVLGGGDEAAA
GGVSVGVAAWYEVWVRVRGGVIAPTLQVAVWVCMVMSVMLVVEATFNSAVSLGVKAIGWRPEWRF
KWEPLAGADEEKGRGEYPMVMVQIPMYNELEVYKLSIGAACELKWPKDKLIVQVLDDSTDPFIKNLVE
LECESWASKGVNIKYVTRSSRKGFKAGALKKGMECDYTKQCEYIAIFDADFQPEPNFLLRTVPFLMH
NPNVALVQARWAFVNDTTSLLTRVQKMFFDYHFKVEQEAGSATFAFFSFNGTAGVWRTTAINEAGG
WKDRTTVEDMDLAVRASLNGWKFIYVGDIRVKSELPSTYGAYCRQQFRWACGGANLFRKIAMDVLV
AKDISLLKKFYMLYSFFLVRRVVAPMVACVLYNIIVPLSVMIPELFIPIWGVAYIPMALLIITTIRNPRNLHI
MPFWILFESVMTVLRMRAALTGLMELSGFNKWTVTKKIGSSVEDTQVPLLPKTRKRLRDRINLPEIGF
SVFLIFCASYNLIFHGKTSYYFNLYLQGLAFLLLGFNFTGNFACCQ
>CSLA_2DS_TGACv1_177473_AA0578070.1_Triticum_aestivum
MEAAEIGGALLFALAAAAALFAAVSTGAVDFSHPPAVGGRVDFQEAISWFIGVFDGSSSSSSAAGGV
SLAEVYELWVRVRGRVIAPALQVAVWACMVMSVMLVVEALYNCVVSLGVKAVGWRPEWRFKWEPL
AGDDEEKGGAHYPMVLVQIPMYNELEVYKLSIGAACELQWPKDRIIVQVLDDSTDPFIKNLVELECES
WAVKGLNIKYATRSSRKGFKAGALKKGMECDYAKQCEYVAIFDADFQPEPDFLLRTVPFFVHNPEVA
LVQARWSFVNDTASLLTRVQKMFFDYHFKVEQEAGSATFSFFSFNGTAGVWRTAAIKEAGGWKDRT
TVEDMDLAVRATLKGWKFVYVGDIRVKSELPSTYKAYCRQQFRWSCGGAHLFRKVAKDILTAKDVSL
IKKFHMLYSFFLVRRVMAPTVACILYNIILPISVMIPELFLPVWGIAYIPTVLLVVTAIRHPKNLHILPFWILF
ESVMTMHRMRAALSGLFELSEFNEWVVTKKTGNNFEDNEVPLLQKTRKRLRDRVNFREIVFSAFLFF
CASYNLVFPGKTSYYFNLYLQGLAFVCLGLNFTGTCSCCQ
>CSLA_2BS_TGACv1_146583_AA0468630.1_Triticum_aestivum
MEAAEIGGALLFALAAAAALFSAVSTGAVDFSHPLAVGGRVDFQETISWFIGIFDGSSSSSSAAGGVS
LAEVYELWVRVRGRVIAPALQVAVWACMVMSVMLVVEALYNCVVSLGVKAVGWRPEWRFKWEPLA
GDDEEKGGAHYPMVLVQIPMYNELEVYKLSIGAACELQWPKDRIIVQVLDDSTDPFIKNLVELECESW
AVKGLNIKYATRSSRKGFKAGALKKGMECDYAKQCEYVAIFDADFQPEPDFLLRTVPFFVHNPEVALV
QARWSFVNDTASLLTRVQKMFFDYHFKVEQEAGSATFSFFSFNGTAGVWRTSAIKEAGGWKDRTTV
EDMDLAVRATLKGWKFVYVGDIRVKSELPSTYKAYCRQQFRWSCGGAHLFRKVAKDILTAKDVSLIK
KFHMLYSFFLVRRVVAPTVACILYNIILPISVMIPELFLPVWGIAYIPTVLLVVTAIRHPKNLHILPFWILFE
SVMTMHRMRAALSGLFELSEFNEWVVTKKTGNNFEDSEVPLLQKTRKRLRDRVNFREIVFSAFLFFC
ASYNLVFTGKTSYYFNLYLQGLAFVCLGLNFTGTCSCCQ
>CSLA_2AS_TGACv1_113418_AA0355820.2_Triticum_aestivum
MEAAEIGGALLFALAAAAALFAAVSTGAIDFSRPLAVGGRVDFQEAISWFIGVFDGSSSSSAAGGVSL
AEVYELWVRVRGRVIAPALQVAVWACMVMSVMLVVEALYNCVVSLGVKAVGWRPEWRFKWEPLAG
DDEEKGGAHYPVVLVQIPMYNELEVYKLSIGAACELQWPKDRIIVQVLDDSTDPFIKNLVELECESWS
VKGLNIKYATRSSRKGFKAGALKKGMEYDYAKQCEYVAIFDADFQPEPDFLLRTVPFFVHNPEVALVQ
ARWSFVNDNASLLTRVQKMFFDYHFKVEQEAGSATFSFFSFNGTAGVWRAAAIKEAGGWKDRTTVE
DMDLAVRATLKGWKFVYVGDIRVKSELPSTYKAYCRQQFRWSCGGAHLFRKVAKDILTAKDVSLIKK
FHMLYSFFLVRRVVAPTLACILYNIILPISVMIPELFLPIWGIAYIPTVLLVVTAIRHPKNLHILPFWILFESV
MTMHRMRAALSGLFELSEFNEWVVTKKTGNNFEDNEVPLLQKTRKRLRDRVNFREIVFSAFLFFCAS
YNLVFPGKTRYYFNLYLQGLAFVCLGLNFTGTCSCCQ
>CSLH_3DS_TGACv1_273502_AA0931770.1_Triticum_aestivum
MGSAMKLQERVILPRTAWKLADIFILCLLFALLSCRVASLGDGGAGAASVAALVCEAWFTFVWILNMNI
KWNPVRFHTYPENLSQRMDELPAVDMLVTTADPELEPPLMTVNTVLSLLAVDYPDVDKLACYVSDD
GCSPATCYALREAAWFARLWVPFCKRHDVRVRAPIIYFASRLEPELAGDTFSDEWTFIKSEYDKLVSR
IESADEGSLLRHDDAGEFTEFMEAERTDHPAIVKVLWDNSKSSRTGEAFPHLVYVSSEKSRKHHHHY
KAGAMNVLARVSAVMTNAPIILNVDCDMFVNNSQVVLHAMCLLLGFDDETCSGFVQVPQRFYGKLKD
DPFGNQMEVLREKLLGGLSGLQGIFYLGTGCFHRRKIIYGVAPSSFAAVKHEREGSLSYEDLRTKFGA
SVELVESTRNIYSREIPPKPMVNISSCIQVAKQVSSCNYETGTHWGQEIGWSYGSMAEDILTGQRIHS
AGWKSTLLDTNPPAFLGCAPTGGPASLTQYKRWATGVLEILLGQNCPIIATIFKRLQFRQCLAYLVFYV
WSMRAPFELCYALLGPFCLFRNHSFLLKHQTMVSASN

>CSLA_2AS_TGACv1_113300_AA0354190.1_Triticum_aestivum
MEAGEIGGALLFVLAAAAAVLAAVSTGAVDFSHPPAVGGQLDFQETISWFTGVYNGASYSSGAGAVS
LAEVHELWVRVRGRVIAPALQVTVWACMVMSVMLAVEALYNCVVSLGVKAIGWRPEWRFKWEPLA
GDEEKGSAHYPMVLVQIPMYNELEVYKLSIGAACELKWPKDRMIVQVLDNSTDPLIKNLVELECETWV
TKGLNIKYAPRSGQKGFKAGALKKGMECDYARQCEYVAIFDADFQPEPDFLLRTIPFFVHNPKVALVQ
ARWSFVNGTVSLLTRIQKMFFDYHFKVEQEAGSATFAFFSFNGTAGVWRTAAIKEAGGWKDRTTVE
DMDLAIRATLKGWKFIYVGDIRVKSELPSSYKAYCRQQFRWACGGANLFRKVAIDILTSKDVSVVKKF
YMLYSFLFVRRVVAPAVACILSNIIVPLSVMIPELYLPVWGVAYIPAVLLVVTAIRNPKNIHLLPFWILFES
VMTIHRTRAALVGLFEFSEFNEWVVTKKTGNNFEDNKVPLLQKTRKRLRDRVNFPEILFSAFLFFCAS
YNLVFPGKTSYYFNLYLQGLAFAFLGLNFSGTCTCFQ
>OS06G12460_OsCslA3
MAMAGADGPTAGAAAAVRWRGGESLLLLLLRWPSSAELVAAWGAARASAVAPALAAASAACLALSA
MLLADAVLMAAACFARRRPDRRYRATPLGAGAGADDDDDDEEAGRVAYPMVLVQIPMYNEREVYKL
SIGAACGLSWPSDRLIVQVLDDSTDPTVKTWYDRLRKTLVQQAHPAQADMDVHQSTKRKNKELMTR
VPILECDSNHGLASIISSYLIAVGLVELECKSWGNKGKNVKYEVRNTRKGYKAGALKEGLLRDYVQQC
NYVAIFDADFQPEPDFLLRTIPYLVRNPQIGLVQAHWEFGTAGVWRISALEEAGGWKDRTTVEDMDL
AVRAGLKGWKFVYLADVKVKSELPSNLKTYRHQQHRWTCGAANLFRKVGAEILFTKEVPFWWKFYL
LYSFFFVRKVVAHVVPFMLYCVVIPFSVLIPEVTVPVWGVVYVPTTITLLHAIRNTSSIHFIPFWILFENV
MSFHRTKAMFIGLLELGGVNEWVVTEKLGNGSNTKPASQILERPPCRFWDRWTMSEILFSIFLFFCAT
YNLAYGGDYYFVYIYLQAIAFLVVGIGFCGTISSNS
>OS03G26050_OsCslA5
MEAGEAAGAVLFLLAAAVSLLAAVSTGALDFTYLVTVVGEGSSTSPGSGGGAWWREAWVGARSRA
VAPALQVGVWACMVMSVMLVVEATYNSAVSVAARLVGWRPERWFKWEPLGGGAGAGDEEKGEAA
AAAYPMVMVQIPMYNELEVYKLSIGAVCGLKWPKERLIIQVLDDSTDAFIKNLVELECEDWASKGLNIK
YATRSGRKGFKAGALKKGMEWDYAKQCEYVAIFDADFQPEPDFLLRTVPFLMHNQNVALVQARWVF
VNDRVSLLTRIQKTFLDYHFKAEQEAGSATFAFFSFNGTAGVWRTEAINDAGGWKDRTTVEDMDLAV
RATLKGWKFIYLGDLRVKSELPSTYKAYCRQQFRWSCGGANLFRKMIWDVLVAKKVSSLKKIYILYSF
FLVRRVVAPAVAFILYNVIIPVSVMIPELFLPIWGVAYIPTALLIVTAIRNPENLHTVPLWILFESVMSMHR
LRAAVAGLLQLQEFNQWIVTKKVGNNAFDENNETPLLQKSRKRLINRVNLPEIGLSVFLIFCASYNLVF
HGKNSFYINLYLQGLAFFLLGLNCVGTLPDHCCF
>AAL25127_OsCslA6
MQGSSTSILHFVPSDPTSTSVLDFLSPTPRGTSPVHDRRLHAGDLALRAGGDRLLVADTVAAVVESLV
QAWRQVRMELLVPLLRGAVVACMVMSVIVLAEKVFLGVVSAVVKLLRRRPARLYRCDPVVVEDDDE
AGRASFPMVLVQIPMYNEKEVYQLSIGAACRLTWPADRLIVQVLDDSTDAIVKELVRKECERWGKKGI
NVKYETRKDRAGYKAGNLREGMRRGYVQGCEFVAMLDADFQPPPDFLLKTVPFLVHNPRLALVQTR
WEFVNANDCLLTRMQEMSMDYHFKVEQEAGSSLCNFFGYNGTAGVWRRQVIDESGGWEDRTTAE
DMDLALRAGLLGWEFVYVGSIKVKSELPSTLKAYRSQQHRWSCGPALLFKKMFWEILAAKKVSFWKK
LYMTYDFFIARRIISTFFTFFFFSVLLPMKVFFPEVQIPLWELILIPTAIILLHSVGTPRSIHLIILWFLFENV
MALHRLKATLIGFFEAGRANEWIVTQKLGNIQKLKSIVRVTKNCRFKDRFHCLELFIGGFLLTSACYDYL
YRDDIFYIFLLSQSIIYFAIGFEFMGVSVSS
>GRMZM2G010142_P02
MEAGEIGGALVFILAAAAAVAAAVSVGAVDFSRPLTAGAPFDFQAAVSWLIGILDGTSSAAADVDGAW
VAVRAGVIAPVLQVAVWACMVMSVMLVVEAVYNSVISLGVKAIGWRPEWRFKWKPLDSADEEKGTA
HFPMVLVQIPMYNELEVYKLSIAAACELQWPKDRIVIQVLDDSTDPFIKNLVELECEHWVNKGVNIKYA
TRTSRKGFKAGALKKGMECDYAWQSEYIAIFDADFQPEPDFLLQTVPFLLHNPEVALVQARWSFVND
TTSLLTRVQKMFYDYHFKVEQEAGSATFAFFSFNGTAGVWRTGAIRDAGGWKDRTTVEDMDLAVRA
TLKGWKFVYVGDVRVKSELPSTYKAYCRQQFRWSSGGANLFRKMAKDVLFAKDISLVKKFYMLYSF
FFVRRVVAPTAACILYNVIIPISVTIPELYLPVWGVAYIPMVLTVVTAIRHPKNLHILPFWILFESVMTLHR
MRAAMTGLLELEGFNQWIVTKKVGNDLEDTEVPLLQKTRKRLRDRVNLPEIGFSVFLFLCASYNLVFH
GKTSYYLYMYLQGLAFLLLGFNFTGNCSCYQ
>CSLA_3AL_TGACv1_197519_AA0666560.1_Triticum_aestivum
MAGAGEEFMASAAGVWAELPVRVDWAAVAAQCAWAGAQARAFLVVPAIRLLLVLSLTMTVMILLEKI
FVAAVCFAAKAFGHKPERRYQWRPIAASACKTGGVDEEASVGGGSSAFPVVLVQIPMYNEREVYKL
SIGAACALEWPSDRVVIQVLDDSTDPAVKDLVEIECQRWKGKGVNIKYEVRGNRKGYKAGALKEGLK
HDYVQECEFIAMFDADFQPESDFLLRTVPFLVHNPDIALVQTRWKFVNSDECLLTRFQEMSLDYHFKF
EQEAGSIVYSFFGFNGTAGVWRISAIDDAGGWKDRTTVEDMDLAVRTALKGWKFVYVGAVKVRSEL
PSTFKAYRFQQHRWSCGPANLFKKMLVEILENKKVSFWSKLHLLYDFFFVGKIAAHTVTFIYYCFAIPL
SVFFPEIQIPLWGVVYVPTVITLCKALGSPSSFHLVILWVLFENVMSLHRIKAAVTGLLDAGRVNEWVV
TEKLGDANKTKPAMEALDAVKVIDVELATPLVPKLKKRRIKLWDKYNCSEIFVGTSIIICGFYDLFYANK
GYYIYLFIQGLAFLVVGFEYIGTRPPTPSAE
>CSLA_3DL_TGACv1_249033_AA0835410.1_Triticum_aestivum

MAGAGEEFMAGVWAEVPVRVDWAAVAAQCAWAGAQARAFLVVPAIRLLVVLSLAMTVMILLEKIFVA
AVCFAAKAFGHRPERRYQWRPIAAGAAAAARGDEEAGLVGGGGGSAAFPVVLVQIPMYNEREVYKL
SIGAACALEWPSDRVVIQVLDDSTDPAVKDLVEIECQRWKGKGVNIKYEVRGNRKGYKAGALKEGLK
HDYVQECEFIAMFDADFQPESDFLLRTVPFLVHNPDIALVQTRWKFVNSDECLLTRFQEMSLDYHFKF
EQEAGSIVYSFFGFNGTAGVWRISAIDDAGGWKDRTTVEDMDLAVRTALKGWKFVYVGAVKVRSEL
PSTFKAYRFQQHRWSCGPANLFKKMLVEILESKKVSFWSKLHLLYDFFFVGKIAAHTVTFIYYCFAIPL
SVFFPEIQIPLWGVVYVPTVITLCKALGSPSSFHLVILWVLFENVMSLHRIRAAITGLLDAGRVNEWVVT
EKLGDANKTKPATEVLDAVKVIDVELTTPLVPKLKKRRIRLWDKYNCSEIFVGTCIVICGFYDLFYANKG
YYIYLFIQGLAFLVVGFEYIGTRPPTPSA
>CSLA_3B_TGACv1_221079_AA0729630.1_Triticum_aestivum
MAGAGEEFMAAVWAGLPVRVDWAAVAAQCAWAGMQARAFLVVPAIRLLVLLSLAMTVMILLEKVFV
AAVCFAAKAFGHRPERRYQWRPIAAGAAAAARGDEEAGVGGGGSAAFPVVLVQIPMYNEREVYKLS
IGAACALEWPAERVVIQVLDDSTDPVVKDLVEIECQRWKGKGVNIKYEVRGNRKGYKAGALKEGLKH
DYVQECEFIAMFDADFQPESDFLLRTVPFLVHNPDIALVQTRWKFVNSDECLLTRFQEMSLDYHFKFE
QEAGSIVYSFFGFNGTAGVWRIAAIDDAGGWKDRTTVEDMDLAVRTALKGWKFVYVGAVKVRSELP
STFKAYRFQQHRWSCGPANLFKKMLVEILESKKVSFWSKLHLLYDFFFVGKIAAHTVTFIYYCFAIPLS
VFFPEIQIPLWGVVYVPTVITLCKALGSPSSFHLVILWVLFENVMSLHRIRAAVTGLLDAGRVNEWVVT
EKLGDANKTKPAMEVLDAVKVIDVELTTPLVPKLKKRRIRLWDKYNCSEIFVGTCIIISGFYDLFYANKG
YYIYLFIQGLAFLVVGFEYIGTRPPTPSAG
>GRMZM2G108600_P03
MGGVRGHSSAAGVSEEAAAAAAWVWAAADLPVRVDWAGAAAQCASLWAHARALLVVPAVRLLVAL
SLAMTVMVLAEKLFVCAVCVAVRAFRLGPHRRYRWEPIAAAAAAVGYPVVLVQIPMYNEREVYKLSIG
AACALEWPPERFVIQVLDDSTDPVVKDLVETECQRWKSKGVNIKYEVRGNRKGYKAGALKEGLKHD
YVADCEYIAMFDADFQPDSDFLLRTIPFLVHNPEIALVQARWKFVNSDECLLTRFQEMSLDYHFKYEQ
EAGSSVYSFFGFNGTAGVWRISAIDDAGGWKDRTTVEDMDLAVRAMLQGWKFLYVGDIKVKSELPS
TFKAYRFQQHRWSCGPANLFKKMMVEILENKRVSLWSKIHLWYDFFFVGKVAAHTVTFIYYCFAIPVS
VLFPEIQIPLWGVVYVPTVITLLKALGTPSSFHLVILWVLFENVMSLHRIKAAVSGLLDAGGRVNEWVV
TEKLGDTSKAKPGTNGSDTAVKVIDVKLTEPLVPKLVKRRARFWERYHCSELFVGTCIILCGFYDLLFA
NKGYYIFLFLQGTAFLVVGFGYVGTLPPCTA
>CSLA_3DS_TGACv1_272005_AA0912960.1_Triticum_aestivum
MAATAWLWAEVPVPVRVDWAAVAAQCAWAGQQARALLVVPTVRLLVLLSLAMTVMILLEKLFVAAV
CYAAKAFGHRPESRYRWRPIAASACKAGGGDEEDGIVIVGSSSGSAAFPVVLVQIPMYNEREVYKVSI
GAACALEWPSDRMVIQVLDDSTDPVVKDLVKIECQRWKSKGVNIRYEVRENRKGYKAGALKQGLMR
DYVRECEFIAMFDADFQPESDFLLRTVPFLVHNPDIALVQTRWKFVNSDECLLTRFQEMSLDYHFKFE
QEAGSIVYSFFGFNGTAGVWRISAINDAGGWNDRTTVEDMDLAVRTALLGWKFVYVGDVKVRSELP
STFKAYRFQQHRWSCGPANLFKKMLVDILENKKVSFWSKLHLLYDFFFVGKIAAHTLTFIYYCFAIPVS
VFFPEIQIPLWGVVYVPTVITLCKSLGSPSSFHLVILWVLFENVMSLHRIKATITGLLDTRRVNEWVVTE
KLGDANKTEPAMEGLDDVQVIDVELSTPLVPKLEKRRTRLWDKYNCSEIPVGTCIIICGCYDVLYAKKG
YYIYLFIQGLAFLVVGFEYIGTRPPSAE
>CSLA_3B_TGACv1_220828_AA0720500.1_Triticum_aestivum
MAMAATAGLWAEVPVRLDWATVAAQCSLAGEQARAFLVVPALRLLVLLSLAMTVMILLEKLFVAAVC
YSAKAFRHRPESRYRWRPITASACKTGGDDEEDGIVVVGSGSGRAAFPVVLVQIPMYNEREVYKVSI
GAACALEWPSDRMVIQVLDDSTDPVVKELVKTECQRWKGKGVNIRYEVRGNRKGYKAGALKQGLM
RDYVRECEFIAMFDADFQPESDFLLRTVPFLVHNPDIALVQTRWKFVNSDKCLLTRFQEMSLDYHFKF
EQEAGSIVYSFFGFNGTAGVWRISAINDAGGWEDRTTVEDMDLAVRTSLLGWKFVYVGAVKVKSELP
STFKAYRFQQHRWSCGPANLFKKILLDILKNKKVSFWSKLHLLYDFFFVGKIAAHTVTFIYYCFAIPVSV
FFPEIQIPLWGVVYVPTVITLCKALGSPSSFHLVILWVLFDNVMSLHRIKATITGLLDTRRVNEWVVTEK
LGDANKTEPAMERLDDVQVIDVELSTPLVPKLEKRRTRLWHKYNCSEIFVGTFIIICGCYDVLYAKKGY
YIYLFIQGLAFLVIGFEYIGTRPPSTE
>CSLA_3DS_TGACv1_273022_AA0927600.1_Triticum_aestivum
MAATVGLREEVPVRLDWATVAAQCAWAGEQTRSFLVVPAVRLLVLLSLAMTVMILLEKLFVAAVCYA
AKAFGHRPESRYKWGPIAASACKTGGDDEEDGIVVVGSGSGSGAFPVVLVQIPMYNEREVYKVSIGA
ACALEWPSDRMVIQVLDDSTDPVVKELVKTECQRWKGKGVNIRYEVRGNRKGYKAGALKQGLMRD
YVRECKFIAMFDADFQPESDFLLRTVPFLVHNPDIALVQTRWKFVNSDKCLLTRFQEMSLDYHFKFEQ
EAGSIVYSFFGFNGTAGVWRISAINDAGGWEDRTTVEDMDLAVRTALLGWKFVYVGAVKVGSELPST
FKAYRFQQHRWSCGPANLFKKMLLDILRKKKVSFWSKLHLLYDFFFVGKIAAHMVTFIYYCFAIPVSVF
FPEIQIPLWGVVYVPTVITLCKALGSPSSFHLVILWVLFDNVMSLHRIKATITGLLDTRRVNEWVVTEKL
GDANKTEPAMEGLDDVQVIDVELSTPLVPKLEKRRTRLWDKYNCSEIFVGTCIIICGCYDVLYAKKGYY
IYLFIQGLAFLVIGFEYIGTRPPSIE
>CSLA_3AS_TGACv1_210508_AA0674280.1_Triticum_aestivum
MAMAATAWLWVEVPVRVDWPAVAAQCAWAGEQARAFLVVPAVRLLVLLSLAMTVMILLEKLFVAAV
CYAAKAFGHRPESRYQWRPIAASACKTGGDDEEDGIVVVGSAAFPVVLVQIPMYNEREVYKVSIGAA

CALEWPSDRMVIQVLDDSTDPVVKDLVKIECQRWKSKGVNIRYEVRQNRKGYKAGALKEGLMRDYV
RECEFIAMFDADFQPESDFLLRTVPFLVHNPDIALVQTRWKFVNSDECLLTRFQEMSLDYHFKFEQEA
GSIVYSFFGFNGTAGVWRISAINDAGGWKERTTVEDMDLAVRTALLGLKFVYVGAVKVKSELPSTFKA
YRFQQHRWSCGPANLFKKVLVEILHNKKVSFWCKLHLLYDFFFVGKITAHTVTFIYYCFAIPVSVFFPEI
QIPLWCVVYVPTVITLCKALGSPSSFHLVILWVLFDNVMSLHRIKATITGLLDARRVNEWVVTEKLGDA
NKTEPAVEGLNDVQVIDVELSTPLVPKLEKRRTRLWDKYNCSEIFVGTCIIICGCYDVLYANKGYYIYLF
IQGLAFLVIGFEYIGTRPPNTE
>AT2G35650_CslA7
MSPLPIFHRLPHATFSSFLLSLSQAGSSKTSVAFLNAFKSEDIIARIGLWWQLIRAVVVVPVFKFLVLLC
LVMSVMFFVEVMYMGIVVLYVKLFKRKPEKFYKWEAMEDDVECGSASYPMVLVQIPMYNEKEVCEQ
SIAAACKISWPSNRIIIQVLDDSTDPASKELVKKECDRWSKEGVNITFEIRDNRNGYKAGALREGMRHS
YVKQCDYVAIFDADFQPDPDFLHRTVPFLIHNPKLALVQGRWEFVNAGQCMMTRLQEMSLSYHFTIE
QQVGSSTFAFFGFNGTAGVWRISALNESGGWNDQTTVEDMDLAVRATLRGWKFLYIDDLKVKSELP
CSFKALRSQQHRWTCGPANLLRKMAGQIIRSENVSLWKKWYMLYSFFFMRKIVAHILTFCFYCVILPA
TVLFPEVTVPKWAAFYLPSLITLLIAIGRLRSIHLLAFWVLFENAMSLLRAKALVMGLFETGRVQEWVVT
EKLGDTLKTKLIPQVPNVRFRERVHLLELLVGAYLLFCGIYDIVYGKNTLYVYLLFQSVAFFVVGFGFVG
KYVPASSYLA
>AT1G23480_CslA3
MSPFLKFFLFLYDYLSPSSFFLVQRNTLGASLDTTDGVVRSGIIGEIIYIWKQTRIFVFIPILKCLVTICLVM
SLLLFIERVYMSIVVVFVKLLRRTPEKVHKWEPINDDDLELANTNYPMVLIQIPMYNEKEVCQLSIGAAC
RLSWPLDRMIVQVLDDSTDPASKELVNAECDKWARKGINIMSEIRDNRIGYKAGALKAGMMHNYVKQ
CEFVAIFDADFQPDPDFLERTIPFLIHNHEISLVQCRWKFVNANECLMTRMQEMSLNYHFVAEQESGS
SIHAFFGFNGTAGVWRIAALNEAGGWKDRTTVEDMDLAVRACLHGWKFVYVHDVEVKNELPSTFKA
YRFQQHRWSCGPANLWRKMTMEILQNKKVSAWKKLYLIYNFFFIRKIVVHIFTFVFYCLILPTTVLFPEL
QVPKWATVYFPTTITILNAIATPRSLHLLVFWILFENVMSMHRTKATFIGLLEAGRVNEWVVTEKLGDTL
KSKLIGKATTKLYTRFGQRLNWRELVVGLYIFFCGCYDFAYGGSYFYVYLFLQSCAFFVAGVGYIGTF
VPTV
>CSLA_7DL_TGACv1_602617_AA1962870.1_Triticum_aestivum
MEAAEQIAVVWKQVRGPVIVPLLRASVMVCLAMCVILFVEKVYMAVVIVAMRLIGRRPERQWRWEPL
RDDDPELGNAAYPMVLVQIPMYNEREVYKKSIGAACGLSWPSDRIVIQVLDDSTDPAIKELVQVECQR
WANKGVNIKYEIRDNRRGYKAGALKEGMKHGYVKDCDFVVIFDADFQPEPDYLSRAMPFLIHNPEIAL
IQARWVFVNANECLMTRMQEMSLDYHFKVEQEVGSSAYAFFGFNGTAGVWRISALNEAGGWKDRT
TVEDMDLAVRASLKGWKFVYLGDLRVKSELPSTFKAFRYQQHRWSCGPANLFRKMLMEIVKNQKVT
LWKKIYVIYNFFFVRKIIGHILTSVFYCLVIPATVFVPEIEIPRWGYFYIPTIITLLNAVGTPRSFHLVIFWVL
FENVMSLHRTKATFSGLLELGRVNEWVVTEKLGDVLKMKVQSKVTKKLRMRIRERYIIIIGLLMCISQLS
YLNFNMESGCSFWSLVLQPISSFVEVTTFCLAKDITISFSSCNPSLSLSSVWDLLGHLSPTDNALRSW
CNVVRKDSV
>AT1G24070_CslA10
MTTFLKSLIFLQDSCLAFLSLMFHRGSSEDAAEALKKLETSINGARISFDTTWTREFRSLFIVPLFKCLV
AFCLIISLLVFIEGIYMNLVVLYVKVFERKPEKVYRWEAMQEDIELGHETYPMVLVQIPMYNEKEVLQLS
IGAACRLIWPLDRLIVQVLDDSTDQTIKELVNTECAKWESKGVNIKCERRDNRNGYKAGALKEGMKHN
YVKLCNYVVIFDADFQPEPDYLQHSVPFLVHNPEVALVQARWRFMNANKCLMTRMQEMSLNYHFMA
EQESGSTRHAFFSFNGTAGVWRMAAMEEAGGWHDRTTVEDMDLAVRAGLLGWKFVFLNDLTVKSE
LPSKFKAFRFQQHRWSCGPANLFRKMIMEIIRNKRVTIWKKLYLVYSFFFLRKIIVHCFTFIFYCVILPTS

VFFPEVNIPAWSTFYIPSMITLCIVIATPRSFYLVIFWILFENVMSMHRTKGTFIGILERQRVNEWVVTEK
LGDALKTKLLPRIGKPSNMFLERVNSKEIMVGIYILCCACYGLFFGNTLLYLYLFMQAVAFLISGVGFVG
T
>CSLA_7AS_TGACv1_569190_AA1809650.1_Triticum_aestivum
MAGDGEGAGDGEGAAAFAAAKAEWLGGSGGLPLLRWWRASGGGELLRGWDAVRAGAVAPALAA
VSGACLAMSAMLLAEAVFMAAASLVRRRPERRYSAGPLGAQDGEDEERGLLGYPMVLVQIPMYNER
EVYKLSIGAACGLSWPSDRVIVQVLDDSTDPTIKDLVELECKIWAKKGKNVKYEVRNNREGYKAGALK
EGMLHAYVQQCDFLAVFDADFQPEPDFLMRTIPYLARNPQIALVQARWEFVNPNECLMTRIQKMTLD
YHFKVEQEAGSSTFAFFGFNGTAGVWRISAIKEAGGWDDRTTVEDMDLAVRAGLKGWKFVYVGDVK
VKSELPSNLKAYRRQQHRWTCGAANLFRKMGAEILLTKEVSLWWKLYLLYSFFLVRKVVAHVLPFVL
YCVVIPFSVLIPEIKIPAWGVVYIPTAITILYAVRNPSSIHFIPFWILFENVMSFHRTKATFIGLLELGSVNE
WVVTEKLGSVSNTKPVPQILERPRCRFWDRWTVSELLFAVFLFVCATYNLVYGSDFYFIYIYLQAITFII
VGTGFCGTSNS
>OS03G07350_OsCslA4
MEGQWGRWRLAAAAAASSSGDQIAAAWAVVRARAVAPVLQFAVWACMAMSVMLVLEVAYMSLVS
LVAVKLLRRVPERRYKWEPITTGSGGVGGGDGEDEEAATGGREAAAFPMVLVQIPMYNEKEVYKLSI
GAACALTWPPDRIIIQVLDDSTDPAIKDLVELECKDWARKEINIKYEIRDNRKGYKAGALKKGMEHIYTQ
QCDFVAIFDADFQPESDFLLKTIPFLVHNPKIGLVQTRWEFVNYDVCLMTRIQKMSLDYHFKVEQESG
SSMHSFFGFNGTAGVWRVSAINEAGGWKDRTTVEDMDLAVRASLKGWQFLYVGDIRVKSELPSTFK
AYRHQQHRWTCGAANLFRKMATEIAKNKGVSVWKKLHLLYSFFFVRRVVAPILTFLFYCVVIPLSVMV
PEVSIPVWGMVYIPTAITIMNAIRNPGSIHLMPFWILFENVMAMHRMRAALTGLLETMNVNQWVVTEK
VGDHVKDKLEVPLLEPLKPTDCVERIYIPELMVAFYLLVCASYDLVLGAKHYYLYIYLQAFAFIALGFGF
AGTSTPCS
>CSLA_2DL_TGACv1_160461_AA0550770.1_Triticum_aestivum
MEKKKRRSSISFLLSFGGGRRRMKGVSMLTMARAAWAVVRYAVVVPLLQLAVYLCAAMSLMLFAER
LYMGLVVAALWLRRRRRQRRSPSRNKGGDDDDLESGAAEDLPLVLVQIPMFNEKQVYRLSIGAACG
LWWPADKLVIQVLDDSTDAGIRAMVEAECRRWAGKGVQIRYENRSNRSGYKAGAMREGLKKGYAK
DCELVAVFDADFQPDADFLRRTVPVLQADPSVALVQARWRFVNADECILTRIQEMSLDYHFSVEQEV
GSACHGFFGFNGTAGVWRVQALADAGGWKDRTTVEDMDLAVRASMRGWRFVYAGDVQVRNELP
SSFKAYRYQQHRWSCGPANLMRKMFWEIVASRQVSAWKKVHVLYGFFFVRKVVAHLVTFLFYCVVI
PAYVLVGGQDVRLPKYVAMYVPAIITLLNAVCTPRSWHLLVFWILFENVMSMHRSKATIIGLVEASRAN
EWVVTEKLGSVTSSTPAATTTMATNKGATKKKKSQSSILAPEIVMGLCLLYCAVYDIVFGHDHFYVYLL
MQSAAAFVIGFGYVGSQ
>CSLA_7BS_TGACv1_592860_AA1945380.1_Triticum_aestivum
MAGDGEGAAAFAVAKAEWLAGSGGLPLLRWWRASGGGELLRGWDAVRAGAVAPALAAVSGACLA
MSAMLLAEAVFMAAASLVRRRPERRYSAGPLGAQDGEDEDEERGLLGYPMVLVQIPMYNEREVYKL
SIGAACGLSWPSDRVIVQVLDDSTDPTIKDLVELECKIWAKKGKNVKYEVRNNREGYKAGALKEGML
HAYVQQCDFLAVFDADFQPEPDFLMRTIPYLSRNPQIALVQARWEFVNPNECLMTRIQKMTLDYHFK
VEQEAGSSTFAFFGFNGTAGVWRISAIKEAGGWDDRTTVEDMDLAVRAGLKGWKFVYVGDVKVKSE
LPSNLKAYRRQQHRWTCGAANLFRKMGAEILLTKEVSLWWKLYLLYSFFLVRKVVAHVVPFVLYCVVI
PFSVLIPEIKIPAWGVVYIPTAITILYAVRNPSSIHFIPFWILFENVMSFHRTKATFIGLLELGSVNEWVVT

EKLGSVSNTKPVPQILEKPRCRFWDRWTVSELLFAVFLFVCATYNLVYGSDFYFIYIYLQAITFIIVGTG
FCGTSNS
>CSLA_7DS_TGACv1_623146_AA2050070.1_Triticum_aestivum
MAGDGEGAAAFAAAKAEWLDGSGGLPLLRWWRASGGGELLGRWDAVRAGAVAPALAAVSGACLA
MSAMLLAEAVFMAAASLVRRRPERRYSAGPLGAQDGEDEERGLLGYPMVLVQIPMYNEREVYKLSI
GAACGLSWPSDRVIVQVLDDSTDPTIKDLVELECKIWAKKGKNVKYEVRNNREGYKAGALKEGMLHA
YVQQCDFLAVFDADFQPEPDFLMRTIPYLARNPQIALVQARWEFVNPNECLMTRIQKMTLDYHFKVE
QEAGSSTFAFFGFNGTAGVWRISAIKEAGGWDDRTTVEDMDLAVRAGLKGWKFVYVGDVKVKSELP
SNLKAYRRQQHRWTCGAANLFRKMGAEILLTKEVSLWWKLYLLYSFFLVRKVVAHVVPFVLYCVVIPF
SVLIPEIKIPAWGVVYIPTAITVLYAVRNPSSIHFIPFWILFENVMSFHRTKATFIGLLELGSVNEWVVTEK
LGSASNTKPVPQILERPRCRFWDRWTVSELLFAVFLFVCATYNLVYGSDFYFIYIYLQAITFIIVGTGFC
GTSNS
>GRMZM2G405567_P02
MGAAAAGGHALRAVGDVVSFPATVAAFVEALLQGWAEARAGLLVPLLRAAVLLCTAMSLIVLAEKVFL
GAVSSVAKLRRRRPGRVCRCDPDEEAAAASQAYPMVLVQIPMYNEREVYQLSIEAACRLTWPVDRLI
VQVLDDSTDSVIKELVKGECERWATEEGINVKYETRKDRAGYKAGNLKEGMRHAYVRACEFVAMFD
ADFQPPPDFLVRTVPFLVHNPSLALVQTRWKFVNANDCLLTRMQEMSMDYHFKVEQEAGSSLCNFF
GYNGTAGVWRTQAIVESGGWEDRTTAEDMDLALRAGLLGWEFVYVGSIKVKSELPSTLKAYRSQQH
RWSCGPALLFKKMFWQILAAERVSVWKKWYMVYDFFIARRIVGTFYTFFFFSVLIPLNILLPEAQIPVW
ELIYIPIAITLLNSVGTPRSIHLVILWVLFENVMALHRFKAILIGFLEADRANEWIVTQKLGNLQKLKSIARL
TGSYRFKDRFHFLEVFIGLFLLASACFDYLYRDDYVYLFVLPQSIMYFAIGFQFVGLNVSED
>AT4G13410_CslA15
MFLLLKPLLSLHDLSLNLLSVMFHGETLKASVDGVGINMSTMWRETRNVFIVPLFKCIVVMCLIISLLVF
VESVYMNLVVLYVKLFNRKPEKVYKWEAMQEDMELGHQNYPMVLVQIPMYNEREVFELSIGAACRL
TWPSDRLIVQVLDDSTDPAIMELVSMECTKWASKDININYERRENRNGYKAGALKHGMRHSYVKQC
QYLAIFDADFQPEPDYLQRAIPFLIHNPEVALVQARWRFVNANTCLMTRMQEMSLNYHFMAEQQSGS
TRHAFFGFNGTAGVWRMVAMEEAGGWKDRTTVEDMDLAVRVGLLGWKFIFVNDLEVKSELPSQFK
AFRFQQHRWSCGPANLIRKMTMEIIHNKRVKIWKKFYVIYSFFFLRKIVVHFFTYFFYCVILPTSVFLPE
VNIPNWSTIYVPSVITLLSAIATPRSFYLVIFWVLFENVMAMHRTKGTLIGLFEGGRVNEWVVTEKLGD
TLNTKLLPQNGRLPKRVNLKEMMMGIYILCCACYDFAFGNAFLYLYLFMQATAFLISGVGFVGT
>AT3G56000_CslA14
MATLSDGLFDDMSVLGVIGYVLEQTRFIFLVPILKRLVNLCQVVSVLLFVDAAYMAIVVAIVKLLGRTPQ
KVLKWESFKSDDIELAPSSNHPMVLIQIPIFNEKEVCQLSIGAACKLSWPLDRMIIQVLDDSTEEESQKL
VRLECKKWESEGITIKSEVRGGFREGFKAGALTAGMKHSYVDEYKCEFVVIFDADFQPEPDFLERTV
PFLVHNPEIALVQAGWKYGNADECCMTRIQEMSLNYHFAVEQKSGSSILGFFGFNGTAGVWRIKALN
EAEGWKDRTIVEDMDLAVRAYLRGSKFVYVDDVKVKNELPSSFQAYRFQQHRWSCGPANLFKKIAM
EIIKNQNVSLWKKVYLIYNFFFLRKIVVHIFTFVFYCVILPATVIFPEIEVPKWTTIYIPATITILNAIATPKSF
YLILYWILFENVMAMHRSIGTLIGLLETSRVKEWIVTQKLGESNNLRENLIFPDHYSFPERLRWREIMVG
MYLFICGYYDFVFGRTYLYVYLFLQSIAFFVVGVGYVGMPVPSTPVQTSE
>AT5G22740_CslA2

MDGVSPKFVLPETFDGVRMEITGQLGMIWELVKAPVIVPLLQLAVYICLLMSVMLLCERVYMGIVIVLV
KLFWKKPDKRYKFEPIHDDEELGSSNFPVVLVQIPMFNEREVYKLSIGAACGLSWPSDRLVIQVLDDS
TDPTVKQMVEVECQRWASKGINIRYQIRENRVGYKAGALKEGLKRSYVKHCEYVVIFDADFQPEPDF
LRRSIPFLMHNPNIALVQARWRFVNSDECLLTRMQEMSLDYHFTVEQEVGSSTHAFFGFNGTAGIWR
IAAINEAGGWKDRTTVEDMDLAVRASLRGWKFLYLGDLQVKSELPSTFRAFRFQQHRWSCGPANLF
RKMVMEIVRNKKVRFWKKVYVIYSFFFVRKIIAHWVTFCFYCVVLPLTILVPEVKVPIWGSVYIPSIITILN
SVGTPRSIHLLFYWILFENVMSLHRTKATLIGLFEAGRANEWVVTAKLGSGQSAKGNTKGIKRFPRIFK
LPDRLNTLELGFAAFLFVCGCYDFVHGKNNYFIYLFLQTMSFFISGLGWIGTYVPS
>AT5G03760_CslA9
MELGDTTSVIPDSFMGYRDDITMQMSMVLDQIRAPLIVPALRLGVYICLTMSVMLFVERVYMGIVISLV
KLFGRKPDKRFKYEPIKDDIELGNSAYPMVLIQIPMFNEREVYQLSIGAACGLSWPSDRIVIQVLDDST
DPTIKDLVEMECSRWASKGVNIKYEIRDNRNGYKAGALKEGMKKSYVKSCDYVAIFDADFQPEADFL
WRTVPYLLHNPKLALVQARWKFVNSDECLMTRMQEMSLDYHFTVEQEVGSSTYAFFGFNGTAGIW
RISALNEAGGWKDRTTVEDMDLAVRASLKGWKFLYLGSLKVKNELPSTFKAYRYQQHRWSCGPANL
FRKMAFEIMTNKNVTLWKKVHVIYSFFVVRKLVAHIVTFIFYCVILPATVLVPEVTVPKWGAVYIPSVITL
LNAVGTPRSLHLMVFWILFENVMSLHRTKATFIGLLEGGRVNEWIVTEKLGDVKAKSATKTSKKVIRFR
FGDRIHVLELGVGMYLLFVGCYDAFFGKNHYYLYLFAQAIAFFIAGFGQIGTIVPNH
>CSLA_6DS_TGACv1_543811_AA1744360.1_Triticum_aestivum
MAPLGADAAAAAWAAVRARAVAPALTAAVWACLAMSAMLLLEAACMSLVSLVAVRLLRLRPERRFK
WEPMAGALEGGEADVEDPPASAGRREFPMVLVQIPMYNEKEVYKLSIGAVCALTWPPDRIIIQVLDDS
TDPIIKELVELECQEWASKKIDIKYEVRNNRKGYKAGALKKGMEHVYAQQCEFVAIFDADFQPESDFLL
KTIPFLVHNPKIALVQTRWKFVNYDACLMTRIQKMSLDYHFKVEQESGSFMHAFFGFNGTAGVWRVS
AINESGGWKDRTTVEDMDLAVRACLKEWEFLYVGDIRVKSELPSTFKAYRHQQHRWTCGAANLFRK
MGWEIVTNKGVSIWKKWHLLYSFLFVRRVIAPILTFLFYCVVIPLSAMVPEVHIPVWGLVYIPTAITVMN
AIRNPGSLHLMPFWILFENVMSMHRMRAALTGLLETAHVNDWVVTEKVGDLVKDDFDVPLLEPLKPT
ECVERIYIPELLLALYLLICASYDYVLGSQTYFMYIYLQALAFIVLGFGFVGMKTPCS
>GRMZM2G074792_P01
MRARLDYLAPPLQFLTNACVLLFLVQSVDRLVLCLGCFWIKLKGVRPVPPLPADKEDVEAGPDGVPM
VLVQMPMCNEREVYQQSIGAVCSLDWPRSNFLVQVLDDSDDATTSALIKEEVEKWQREGVRIVYRH
RVIRDGYKAGNLKSAMNCSYVKDYEFVVIFDADFQPQADFLKRTVPHFKGKDDVGLVQARWSFVNK
DENLLTRLQNINLCFHFEVEQQVNGAFLNFFGFNGTAGVWRIKALEESGGWMERTTVEDMDIAVRAH
LKGWKFLFLNDVECQCELPESYEAYRKQQHRWHSGPMQLFRLCFVDIIKSKIGFWKKFNLIFLFFLLR
KLILPFYSFTLFCVILPMTMFVPEAELPAWVVCYIPATMSILNILPSPKSFPFIVPYLLFENTMSVTKFNA
MVSGLFQLGSAYEWVVTKKSGRSSEGDLVALVEKHSKQQRVGSAPNLDALTKESKGTEEEKNKKKR
KKKHNRIYRKELALSFLLLTAAARSLLSAQGVHFYFLLFQGVSFLVVGLDLIGEQVD
>CSLA_6BS_TGACv1_513376_AA1639390.2_Triticum_aestivum
MAPLGADAAAAAWAAVRARAVSPALTAAVWACLAMSAMLLLEAACMSLVSLVAVRLLRLRPQRRFK
WEPMPGALPGAEADAEDPPGRREFPMVLVQIPMYNEKEVYKLSIGAVCALTWPPDRIIIQVLDDSTDP
IIKELVELECQEWASKKIDIKYEVRNNRKGYKAGALKKGMEHVYAQQCEFVAIFDADFQPESDFLLKTI
PFLVHNPKIALVQTRWKFVNYDACLMTRIQKMSLDYHFKVEQESGSFMHAFFGFNGTAGVWRVSAIN
ESGGWKDRTTVEDMDLAVRACLKEWEFLYVGDIRVKSELPSTFKAYRHQQHRWTCGAANLFRKMG

WEIVTNKGVSIWKKWHLLYSFLFVRRVIAPILTFLFYCVVIPLSAMVPEVHIPVWGLVYIPTAITIMNAIRN
PGSLHLMPFWILFENVMSMHRMRAALTGLLETAHVNDWVVTEKVGDLVKDDFDVPLLEPLKPTECVE
RIYIPELLLALYLLICASYDYVLGSQTYFMYIYLQALAFIVLGFGFVGTKTPCS
>CSLA_6AS_TGACv1_487286_AA1569690.1_Triticum_aestivum
MAPLSAGAAAAAWAAVRARAVAPALTAAVWACLAMSAMLLLEAACMSLVSLVAVRLLRLRPERRFK
WEPMTGALEGGEADVEDPAGRREFPMVLVQIPMYNEKEVYKLSIGAVCALTWPPDRIIIQVLDDSTDP
IIKELVELECQEWASKKIDIKYEVRNNRKGYKAGALKKGMEHVYAQQCEFVAIFDADFQPESDFLLKTI
PFLVHNPKIALVQTRWKFVNYDACLMTRIQKMSLDYHFKVEQESGSFMHAFFGFNGTAGVWRVSAIN
ESGGWKDRTTVEDMDLAVRACLKEWEFLYVGDIRVKSELPSTFKAYRHQQHRWTCGAANLFRKMG
WEIVTNKGVSIWKKWHLLYSFLFVRRVIAPILTFLFYCVVIPLSAMVPEVHIPVWGLVYIPTAITIMNAIRN
PGSLHLMPFWILFENVMSMHRMRAALTGLLETAHVNDWVVTEKVGDVVKDDFEVPLLEPLKPTECVE
RIYIPELLLALYLLICASYDYVLGSQTYFMYIYLQALAFIVLGFGFVGMKTPCS
>CSLA_2BL_TGACv1_129747_AA0394630.1_Triticum_aestivum
MRGVSMLTMARAAWAAVRYAVVVPLLQLAVYLCAAMSLMLFAERLYMGLVVAALWLRRRRRQRRN
PSRNKGDDDGGAGDLESGGGEDLPMVLVQIPMFNEKQVYRLSIGAACGLWWPADKLVIQVLDDSTD
AGIRALVEAECRRWAGKGVQIRYENRSNRSGYKAGAMREGLKKGYARDCELVAVFDADFQPDADFL
RRTVPVLQADPAVALVQARWRFVNADECILTRIQEMSLDYHFSVEQEVGSACHGFFGFNGTAGVWR
VQALADAGGWKDRTTVEDMDLAVRASMRGWRFVYAGDVQVRNELPSSFKAYRYQQHRWSCGPA
NLMRKMFWEIVASRQVSAWKKVHVLYGFFFVRKVVAHLVTFLFYCVVIPAYVLVGGQDVRLPKYVAM
YVPAIITLLNAVCTPRSWHLLVFWILFENVMSMHRSKATIIGLVEASRANEWVVTEKLGSVTSTPAAAT
TMAANKGAMKKKKSQSSILAPEIVMGLCLLYCAVYDIVFGHDHFYVYLLMQSAAAFVIGFGYVGSQ
>CSLA_2AL_TGACv1_093375_AA0278800.1_Triticum_aestivum
MKGVSMLTMARAAWAAVRHAVVVPLLQLAVYLCAAMSLMLFAERLYMGLVVAALWLRRRRRQRRN
PGRNKGGDDDVGDLESGAAEDLPVVLVQIPMFNEKQVYRLSIGAACGLWWPADKLVIQVLDDSTDA
GIRAMVEAECRRWAGKGVHIRYENRSNRSGYKAGAMREGLKKGYAKDCELVAVFDADFQPDADFL
RRTVPVLQADPAVALVQARWRFVNADECILTRIQEMSLDYHFSVEQEVGSACHGFFGFNGTAGVWR
VQALADAGGWKDRTTVEDMDLAVRASMRGWRFVYAGDVQVRNELPSSFKAYRYQQHRWSCGPA
NLMRKMFWEIVASRQVSAWKKVHVLYGFFFVRKVVAHLVTFLFYCVVIPAYVLVGGQDVRLPKYVAM
YVPAIITLLNAVCTPRSWHLLVFWILFENVMSMHRSKATIIGLVEASRANEWVVTEKLGSVTSTPAATT
TMATNKGAMKKKKSQSSILAPEIVMGLCLLYCAVYDIFFGHDHFYVYLLMQSAAAFVIGFGYVGSQ
>CSLA_U_TGACv1_642146_AA2112270.1_Triticum_aestivum
MAAALLPGTRITFSGAWQQVRGPVIVPLLRASVLLCVAMSAMLLAEKVYMAVVVLALRLLGRRPELQY
RWEPMRDGDDPELGSAAYPMVLVQIPMYNEREVYQLSIGAACGLSWPSDRIIVQVLDDSTDPVVKEL
VQVECQRWARKGVNIKYETRNNRRGYKAGALKEAMKHGYVKDCDLVAIFDADFQPEPDFLSRSVPF
LVHNPDIALVQARWKFVNADECLMTRMQEMSLDYHFKVEQEVGSSTYAFFGFNGTAGVWRISALNE
AGGWKDRTTVEDMDLAVRASLKGWKFVYLGDLKVKNELPSTFKAFRYQQHRWSCGPANLFRKMVM
EIIKNKKVTLWKKIHVVYNFFFLRKVVAHIMTFVFYCLVIPATVLVPEVEVPKWGCVYIPAIITLLSVVGTP
RSVHLVIFWALFENVMSLHRTKATFIGLLEAHTVNEWVVTEKLGDTVKTKMPSKALKKLRIGIGERLHL
WELGVAAYLFICGCYSISFGNNHYFIFLLMQSIAFFIVGVGYVGTFVTQ
>CSLA_7AL_TGACv1_558725_AA1795700.1_Triticum_aestivum

MSTLPGVWQIAAAWEQVRGPVIVPLLRASVLLCLAMSAMLFAEKVYMAVVVLAVRLLGRRPERQWK
WEPVGEDDPELGSAAYPMVLVQIPMYNEREVYQLSIGAACGLSWPSDRIVVQVLDDSTDPVIKELVQ
VECRRWARKGVNIKYEIRDNRRGYKAGALKEGMKHGYVKDCDLVAIFDADFQPEPDFLWRAVPFLV
HNPDIALVQARWKFVNADECLMTRMQEMSLDYHFRVEQEVGSSTYAFFGFNGTAGVWRISALNEAG
GWKDRTTVEDMDLAVRASLKGWKFVYLGDLKVKNELPSTFKAFRYQQHRWSCGPANLFRKMVMEII
RNKKVTLWKKIHVVYSFFLVRKVVAHIVTFVFYCLVIPATVLVPEVEVPKWGCVYIPTIITLLNAVGTPRS
VHLVVFWVLFENVMSLHRAKATFIGLLEAGTVNEWVVTEKLGDTLKAKMPSKALKKLRMRIGERLHL
WELGVAAYLFLCGCYDISFGNNRYFIFLFMQSIAFFIVGVGYVGTFVAQ
>CSLA_6BS_TGACv1_513375_AA1639370.1_Triticum_aestivum
MDAAVGLPDAWSQVRAPVIVPLLKLAVAVCLLMSVLLFLERLYMAVVIVGVKLLGRRPERRYKCDPIC
EDDDPELGSAAFPIVLVQIPMFNEREVYQLSIGAVCGLSWPSDRLVVQVLDDSTDPLIKEMVRMECER
WAHKGINITYQIREDRKGYKAGALKAGMKHGYVRECEYMVIFDADFQPDPDFLHRTIPYLHHNPEIAL
VQARWRFVNADECLMTRMQEMSLDYHFKVEQEVSSSVCAFFGFNGTAGVWRIAAVNEAGGWKDR
TTVEDMDLAIRASLKGWKFVYLGDVQVKSELPSTFKAFRFQQHRWSCGPANLFRKMLLEIVTNKKVTI
WKKFHVIYNFFLVRKIVAHIVTFTFYCIIIPTTIFVPEVHIPKWGCVYIPTIITLLNSVGTPRSFHLLFFWILF
ENVMSLHRTKATLIGLLEAGRANEWVVTEKLGSAMKMKSANKASARKSFMRMWERLNVPELGVGAF
LFSCGWYDVAFGKDNFFIYLFFQSMAFFVVGVGYVGTIVPPS
>CSLA_6AS_TGACv1_485966_AA1554960.1_Triticum_aestivum
MDAAVGLPDAWSQVRAPVIVPLLKLAVALCLLMSVLLFLERLYMAVVIVGVKLLGRRPERRYKCDPIC
EDDDPELGSAAFPVVLVQIPMFNEREVYQLSIGAVCGLSWPSDRLVVQVLDDSTDPLVKEMVRMECE
RWAHKGINITYQIREDRKGYKAGALKAGMKHGYVRECEYMVIFDADFQPDPDFLHRTIPYLHHNPEIA
LVQARWRFVNADECLMTRMQEMSLDYHFKVEQEVSSSVCAFFGFNGTAGVWRIAAVNEAGGWKD
RTTVEDMDLAIRASLKGWKFVYLGDVQVKSELPSTFKAFRFQQHRWSCGPANLFRKMLVEIVTNKKV
TIWKKFHVIYNFFLVRKIVAHIVTFTFYCIIIPTTIFVPEVHIPKWGCVYIPTIITLLNSVGTPRSFHLLFFWIL
FENVMSLHRTKATLIGLLEAGRANEWVVTEKLGSAMKMKSANKASARKSFMRMWERLNVPELGVGA
FLFSCGWYDVAFGKDNFFIYLFFQSMAFFVVGVGYVGTIVPQS
>OS02G09930_OsCslA1
MEVNGGGAAGLPEAWSQVRAPVIVPLLRLAVAVCLTMSVLLFLERMYMAVVISGVKILRRRPDRRYR
CDPIPDDDPELGTSAFPVVLIQIPMFNEREVYQLSIGAVCGLSWPSDRLVVQVLDDSTDPVIKEMVRIE
CERWAHKGVNITYQIRENRKGYKAGALKEGMKHGYVRECEYVAIFDADFQPDPDFLRRTIPFLVHNS
DIALVQARWRFVNADECLMTRMQEMSLDYHFTVEQEVSSSVCAFFGFNGTAGVWRVSAVNEAGGW
KDRTTVEDMDLAIRASLKGWKFVYLGDVQVKSELPSTFKAFRFQQHRWSCGPANLFRKMLMEIKVTI
WKKIHVIYNFFLIRKIIAHIVTFAFYCLIIPATIFVPEVRIPKWGCVYIPTIITLLNSVGTPRSFHLLFFWILFE
NVMSLHRTKATLIGLLEAGRANEWVVTEKLGNALKMKSSSKSSAKKSFMRVWDRLNVTELGVAAFLF
SCGWYDLAFGKDHFFIYLFFQGAAFFIVGIGYVGTIVPQS
>CSLA_7BL_TGACv1_578444_AA1895100.1_Triticum_aestivum
MEAAEQIAVVWKQVRGPVIVPLLRASVMVCLAMCVILFVEKVYMAVVIVAMRLIGRRPERQWRWEPL
RDDDPELGNAAYPMVLVQIPMYNEREVYKKSIGAVCGLSWPSDRIVIQVLDDSTEPAIKELVQVECQR
WANKGVNIKYEIRDNRRGYKAGALKEGMKHGYVKDCDFVVIFDADFQPEPDYLSRAMPFLIHNPEIAL
VQARWVFVNANECLMTRMQEMSLDYHFKVEQEVGSSAYAFFGFNGTAGVWRISALNEAGGWKDR
TTVEDMDLAVRASLKGWKFVYLGDLRVKSELPSTFKAFRYQQHRWSCGPANLFRKMLMEIVKNQKV

TLWKKIYVIYNFFFVRKIIGHILTSVFYCLVIPATVFVPEVEIPRWGYFYIPTVITLLNAVGTPRSFHLVIFW
VLFENVMSLHRTKATFSGLLELGRVNEWVVTEKLGDVLKMKVQSKVTKKLRMRIRERLQLLELGVAA
YIFFCGSYDLLFGKRYYYIFLFMQSIAFFVVGVGFVGTLVPN
>CSLA_7AL_TGACv1_557254_AA1778850.1_Triticum_aestivum
MEAAEQIAVVWKQVRGPVIAPLLRASVMVCLAMCVILFVEKVYMAVVIVAMRLIGRHPERQWRWEPL
RDDDPELGNAAYPMVLVQIPMYNEREVYKKSIGAACGLSWPSDRIVIQVLDDSTDPAIKELVQAECHR
WANKGVNIKYEIRDNRRGYKAGALKEGMKHGYVKDCDYVVIFDADFQPEPDYLSRAMPFLVHNPEIA
LVQARWVFVNANECLMTRMQEMSLDYHFKVEQEVGSSAYAFFGFNGTAGVWRISALNEAGGWKDR
TTVEDMDLAVRASLKGWKFVCLGDLRVKSELPSTFKAFRYQQHRWSCGPANLFRKMLMEIVKNQKV
TLWKKIYVIYNFFLVRKIIGHILTSVFYCLVIPATVFVPEVEIPRWGYFYIPTVITLLNAVGTPRSFHLVIFW
VLFENVMSLHRTKATFSGLLELGRVNEWVVTEKLGDILKMKVQSKVTKKLRMRIRERLQLLELGVAAY
IFFCGSYDLLFGKRYYYVFLFMQSIAFFVVGVGFVGTLVPN
>CSLA_1AS_TGACv1_019142_AA0061550.1_Triticum_aestivum
MSMLPMARAAWLVLRYAVVVPLLQLAIYLCVVMSLMLFADRLYMGLVVAVLWLYRRCRNRNQRNKG
DDDNLESDDADRPMVLVQIPMFNEKQVFRLSIGAACGLWWPADKLVIQVLDDSTDAGIRSLVEAECR
RWAGKGVHIRYENRSNRSGYKAGAMRDGLKKQYVKDCEFVAVFDADFQPDADFLRHTVPVLEADP
AVALVQARWRFVNADECILTRMQEMSLDYHFSVEQEVGSAFHGFFSFNGTAGVWRLHALADAGGW
KDRTTVEDMDLAVRASMRGWRFVYAGDVQVRNELPSSFKAYRYQQHRWSCGPPNLMRKMFWEIV
ANKQVSAWKKLHVLYGFFFVRKVVAHLATFLFCCVVIPVYVLVGGQDVWLPQYVPMYVAAVLTLLNA
VCTPRSCHLLVFWILFENVMSIHRCKATIIGLLEASRANEWVVTEKLGGSTTSTPAAATTTMVAKKKKS
SSSFLAPEIVMGLFLLYCALYDIVFGHDHFYVYLLMQSAAAFVIGFGYVGSQ
>GRMZM2G115772_P01
MAAWAVPALAAAAWALRAAVWACLAASAMLVAEAAYMGLASLASAAAMLWRRPDARYRWEPMPG
GCDVEAATGADFPMVLVQIPMYNEREVYKLSIDAACALTWPPDRIVIQVLDDSTDPIIKELVELECQDW
ATKKINIKYEVRNNRKGYKAGALKKGMEHIYAKQCEFVAIFDADFQPEPDFLLKTIPFLVHNPKIALVQA
RWEFVNYDVCLMTRIQKMSLDYHFKVEQESGSFVYSFFGFNGTAGVWRVSAINQSGGWKDRTTVE
DMDLAVRASLKGWEFLYVGDIRVKSELPSTFKAYRHQQHRWTCGAANLFRKMAWEIITNKEVSIWKK
HHLLYSFFFVRRVIAPLVTFLFYCVVIPLSAMVPGVSIPVWGLVYIPTAITCMNAIRNPWSLHLMPFWIL
FENVMSMHRMRAAVTGLLETARANDWVVTEKVGDLVKDDLDVPLLEPVKPTECVERIYFPELLLALH
LLICASYDFVLGSHKYYLYIYLQAFAYVVMGFGFVGTKTPCS
>GRMZM2G105631_P01
MDALPEAWSQVRAPVIVPLLRLAVAVCLTMSVLLFLERVYMAVVISGVRLLRLRPDRRYRCDPLPEDD
PELGSSAFPVVLVQIPMFNEREVYQLSIGAVCGLSWPADRLVVQVLDDSTDEVIKEMVRMECERWAR
KGINITYQIREDRKGYKAGALRAGMRHAYVRDCEYVAIFDADFQPDPDYLKRTIPYLVHNPEIALVQAR
WRFVNADECLMTRMQEMSLDYHFTVEQEVSSSVCAFFGFNGTAGVWRISAVNEAGGWKDRTTVED
MDLAIRASLKGWKFVYLGDVQVKSELPSTFKAFRFQQHRWSCGPANLFRKMLMEIVTNKKVTIWKKI
HVIYNFFLIRKIIAHIITFSFYCVIIPATIFVPEVRIPKWGCVYIPSAITLLNSVGTPRSFHLLFFWVAFENVM
SLHRTKATLIGLLEAGRANEWVVTAKLGSAMKMKSANKAGLRKQFMRIWERLHVTELGVAAFLFSCG
WYDLAYGRDHFFIYLFFQSVAFFIVGVGYVGTIVPQS
>OS06G42020_OsCslA9

MWEQVKAPVVVPLLRLSVAACLAMSVMLFVEKVYMSVVLVGVHLFGRRPDRRYRCDPIVAAGADND
DPELADANAAFPMVLIQIPMYNEREVYKLSIGAACGLSWPSDRVIVQVLDDSTDPVIKEMVQVECKRW
ESKGVRIKYEIRDNRVGYKAGALREGMKHGYVRDCDYVAIFDADFQPDPDFLARTIPFLVHNPDIALV
QARWKFVNANECLMTRMQEMSLDYHFKVEQEVGSSTHAFFGFNGTAGVWRISAMNEAGGWKDRT
TVEDMDLAVRAGLKGWKFVYLGDLMVKSELPSTFKAFRYQQHRWSCGPANLFRKMLVEIATNKKVT
LWKKIYVIYNFFLVRKIIGHIVTFVFYCLVVPATVLIPEVEIPRWGYVYLPSIVTILNSIGTPRSLHLLIFWVL
FENVMSLHRTKATLIGLLETGRVNEWVVTEKLGDALKLKLPGKAFRRPRMRIGDRVNALELGFSAYLS
FCGCYDIAYGKGYYSLFLFLQSITFFIIGVGYVGTIVPH
>CSLA_U_TGACv1_642146_AA2112290.1_Triticum_aestivum
MSTLPGAWHVAAAWEQVRGPVIVPLLRASVLLCLAMSAMLFAEKVYMAVVVLAVRLLGRRPERQYQ
WEPMGDDDPELGSAAYPMVLVQIPMYNEREVYQLSIGAACGLSWPSDRIVVQVLDDSTDPVIKELVR
VECRRWARKGVNIKYEIRDNRRGYKAGALKEGMKHGYVKDCDLVAIFDADFLWRAVPFLVHNPDIAL
VQARWKFVNADECLMTRMQEMSLDYHFKVEQEVGSSTYAFFGFNGTAGVWRISALNEAGGWKDRT
TVEDMDLAVRASLKGWKFVYLGDLKVKNELPSTFKAFRYQQHRWSCGPANLFRKMVMEIIRNKKVTL
WKKIHVVYSFFLVRKVVAHIVTFVFYCLVIPATVLVPEVEVPKWGCVYIPTIITLLNAVGTPRSVHLVVF
WVLFENVMSLHRAKATFIGLLEVGTVNEWVVTEKLGDTLKAKMPSKALRKLRMRIGERLHLWELGVA
AYLFLCGCYDISFGNNRYFIFLFMQSIAFFIVGVGYVGTFVAQ
>CSLF_7DL_TGACv1_607937_AA2011180.1_Triticum_aestivum
MVGRDSDTVAFVQFPQRFEGVDPTDLYANHNRIFFDGTLRALDGMQGPIYVGTGCLFRRITVYGFDP
PRINVGGPCFPRLAGLFAKTKYEKPGLEMTMAKAKAAPVPAKGKHGFLPLPKKTYGKSDAFVDSIPR
ASHPSPYAAAAEGIVADEATIVEAVNVTAAAFEKKTGWGKEIGWVYDTVTEDVVTGYRMHIKGWRSR
YCSIYPHAFIGTAPINLTERLFQVLRWSTGSLEIFFSKNNPLFGSTYLHPLQRVAYINITTYPFTAIFLIFY
TTVPALSFVTGHFIVQRPTTMFYVYLGIVLSTLLVIAVLEVKWAGVTVFEWFRNGQFWMTASCSAYLA
AVCQVLTKVIFRRDISFKLTSKLPSGDEKKDPYADLYVVRWTPLMITPIIIIFVNIIGSAVAFAKVLDGEWT
HWLKVAGGVFFNFWVLFHLYPFAKGILGKHGKTPVVVLVWWAFTFVITAVLYINIPHMHSSGGKHTTV
HGHHGKKFVDAGYYNWP
>OS10G26630_OsCslA2
MSAMLVAEAAWMGLASLAAAAARRLRGYGYRWEPMAAPPDVEAPAPAPAEFPMVLVQIPMYNEKE
VYKLSIGAACALTWPPDRIIIQVLDDSTDPFVKFSLVQELVELECKEWASKKINIKYEVRNNRKGYKAG
ALRKGMEHTYAQLCDFVAIFDADFEPESDFLLKTMPYLLHNPKIALVQTRWEFVNYNVCLMTRIQKMS
LDYHFKVEQESGSFMHAFFGFNGTAGVWRVSAINQSGGWKDRTTVEDMDLAVRASLKGWEFLYVG
DIRVKSELPSTFQAYRHQQHRWTCGAANLFRKMAWEIITNKEVSMWKKYHLLYSFFFVRRAIAPILTF
LFYCIVIPLSAMVPEVTIPVWGLVYIPTAITIMNAIRNPGSVHLMPFWILFENVMAMHRMRAALSGLLET
ARANDWVVTEKVGDQVKDELDVPLLEPLKPTECAERIYIPELLLALYLLICASYDFVLGNHKYYIYIYLQ
AVAFTVMGFGFVGTRTPCS
>CSLH_3B_TGACv1_221049_AA0728260.1_Triticum_aestivum
MSSAMKLQERVTVPRTAWKLADIFILCLLLVLLSCRVASLGEGGAGAAALVCEAWFTFVWILNMNIKW
NPVRFHTYPENLSQRMDELPAVDMLVTTADPELEPPLMTVNTVLSLLAVDYPDVDKLACYVSDDGCS
PVTCYALREAAGFARLWVPFCKRHGVGVRAPFIYFASSRPEPDLAGDKFSDDWIFIKSEYDKLVSLIE
SADEASLLRHDHAGEFTEFKGAECGDHPAIVKVLWDNSKSSGTGEGFPNLVYVSREKSRKHDHHYK
AGAMNVLARVSAVMTNAPIILNVDCDMFVNNPQVVLHATCLLLGFDDETCSGFVQVPQRFYGKLKDD

PFGNQMEVLRSLSYEDLLTKFGASMELVESSRNIYSVEIPPKPMIDITSRIQVAKQVSTCNYETGTHW
GEEASNHGFSIQLALFLSYNIYNFVEYKECGLSARTWWNNMRMRINLLLAPCFP
>OS03G56060_OsCslC9
MVLVQIPMCNEKEVYQQSIAAVCNLDWPRSNFLVQVLDDSDDPTTQTLIREEVLKWQQNGARIVYRH
RVLRDGYKAGNLKSAMSCSYVKDYEFVAIFDADFQPNPDFLKRTVPHFKDNDELGLVQARWSFVNK
DENLLTRLQNINLCFHFEVEQQVNGIFLNFFGFNGTAGVWRIKALDDSGGWMERTTVEDMDIAVRAH
LRGWKFIFLNDVECQCELPESYEAYRKQQHRWHSGPMQLFRLCLPDIIKCKIVFWKKANLIFLFFLLRK
LILPFYSFTLFCIILPMTMFVPEAELPDWVVCYIPALMSLLNILPSPKSFPFIIPYLLFENTMSVTKFNAMIS
GLFQLGNAYEWVVTKKSGRSSEGDLISLAPKELKHQKTESAPNLDAIAKEQSAPRKDVKKKHNRIYKK
ELALSLLLLTAAARSLLSKQGIHFYFLLFQGISFLLVGLDLIGEQIE
>GRMZM2G424486_P01
MSCVRNEASGQTGSVYRLSIGAACGMSWPSERLVIQVLDDSTNPAIRELVEVECARWARKGVRICYE
NRSNRNGYKAGAMREGLKKHYARDCEFVAIFDADFQPDSDFLRRTVPLLQRDPGVGLVQARWRYV
NADECILTRIQEMSLNYHFAVEQEVGSACHAFFGFNGTAGVWRVAALADAGGWKERTTVEDMDLAV
RASLRGWRFVYVGDLVVRNELPSTFKAYRYQQHRWSCGPANLFRKVLPEILRSDRVSLGKKFHLLYA
FFFVRKVVAHLVTFLFYCVVIPACVLVQGDVRLPKYVAMYVPALITLLNAACTPRSCHLLIFWILFENVM
SMHRSKAAIIGLLEASRANEWVVTDKLGSSKAAAAVVAKKKKQQLVRSRCCSTRREMHVLELAMGV
CLLYCAVYDIVFFGRDHYYMYLLLQSAAAFIVGFGYVGTTTPS
>AT5G16190_CslA11
MQEDLELGNQNFPMVLVQIPMYNEREVFKLSIGAACRLIWPLDRLIVQVLDDSTDPTIMEMVSTECGK
WATKGINIKCERRDNRNGYKAGALKQGMRHSYVKTCTYIAIFDADFQPEPDYLERTVPFLIHNPELAL
VQARWKFVNAKKCLMTRMQEMSLNYHFTAEQESGSTRHAFFGFNGTAGVWRLAAMEEAGGWKDR
TTVEDMDLAVRVGLHGWKFVFVNDVSVKSELPSQFKAFRFQQHRWSCGPANLFRKMTMEIIRNKRV
TIWKKLYVIYSFFFVRKIIVHFFTFFFYCFILPTSVFFPEVNIPTWSTVYFPFMITLFNAIATPRSFYLVIFW
VLFENVMAMHRTKGTFIGLLEGGRVNEWVVTEKLGDALETKLLPQVRKPRNGFLERINSKEMMVGIYI
LCCASYNLVFGKTVLYIYLYMQALAFIIAGIGFIGT
>Os09g39920_OsCslA8
MSIMLFLERLYMALVVAALWLIRRRRRRSNRREQDDDGAENDQLLQDPEAANSPMVLVQIPMFNEK
QVYRLSIGAACGMTWPSDKLVIQVLDDSTDPAIREMVEGECGRWAGKGVSIRYENRRNRSGYKAGA
MREGLRKAYARECELVAIFDADFQPDADFLLRTVPVLVADPGVALVQARWRFVNADECLLTRIQEMS
LDYHFRVEQEVGSACHGFFGFNGTAGVWRVRALEEAGGWKERTTVEDMDLALVAHLLTFSFYCVVI
PACVLAGSDHVRLPKYVALYVPAAITLLNAACTPRSCHLLIFWILFENVMSMHRTKATLIGLLEATRANE
WVVTDKRGNANPKHQQPANTTTRPGRKTTTSSSRTSFFNNDVHVAEILLGACLLYCALYDIAYGRDS
FYIYLLLQSAAAFIVGFGYVGT
>GRMZM2G178880_P03
MELEGPWARVLHLHDACEWGRAVRARAVAPALEAAAWACLAMSVMLVLEVCYMSVSSFVAVNLLR
RTPQRRYSWEPMPSGTARGDDEEAAVGDGGGEAYPMVLVQIPMYNEREVYKISIGAACALTWPPD
RIIIQVLDDSTDPFIKELVEFECKDWASKKINIKYEIRESRKGYKAGALKKGMEHSYAQECDFVAIFDAD
FQPDPDFLLRTIPFLVHNPKIALVQTRWEFVNYNICLLTRIQKMSLDYHFKVEQESGSSVHAFFGFNGT
AGVWRVSAIGEAGGWKDRTTVEDMDLAVRASLKGWQFLYVGDIRFFVQVKSELPSTFKAYRHQQH

RWTCGAANLFRKMAGDIVISKGATVWKKLHLLYSFFFVRRVIAPILTFLFYCVVIPLSVMVPEVSIPAW
GMFYIPTAITIMTAIRNPW
>GRMZM2G099088_P01
MAADLVETECQRWKSKGVNIKYEVRGNRKGYKAGALKEGLKHDYVEDCEYIAMFDADFQPEPDFLL
RAVPFLVHNPEIALVQARWKFVNSGECLLTRFQEMSLDYHFKYEQEAGSSLHSFFGFNGTAGVWRIA
AIDDAGGWKDRTTVEDMDLAVRAMLQGWKFVYVGDIKVKSELPSTFKAYRFQQHRWSCGPANLFK
KMMVEILENKKVSLWSKIYLWYNFFFVGKVAAHTVTFIYYCCAIPVSVLLPEIQIPLWGVVYIPTLITLLK
ALGTPSSFHLVILWVLFENVMSLHRIKAAASGLLDAGGRVNEWVVTEKLGDTSKAKPGANGSDDVKVI
DVKLTEPLVPKLVKRRARFWERYNCSELFVGTCILLCGFYDLLFAKKGYYIFLFLQGTAFLVVGFGYVG
TLPHCNA
>OS04G35030_OsCslH3
MAAASGEKEEEEKKLQERAPIRRTAWMLANFVVLFLLLALLVRRATAADAEERGVGGAAWRVAFACE
AWFAFVWLLNMNAKWSPARFDTYPENLAGRCGAAHRPRKSSCISGHLDLMRRQCALMQDRRAAG
GRHVRDDGGPGARAAGGDGEQGALAARRRLLPGRRRRRRRRLACYVSDDGCSPVTYYALREAA
GFARTWVPFCRRHGVAVRAPFRYFASAPEFGPADRKFLDDWTFMKSEYDKLVRRIEDADETTLLRQ
GGGEFAEFMDAKRTNHRAIVKVIWDNNSKNRIGEEGGFPHLIYVSREKSPGHHHHYKAGAMNALTRV
SAVMTNAPIMLNVDCDMFANDPQVVLHAMCLLLGFDDEISSGFVQVPQSFYGDLKDDPFGNKLEVIY
KKLLGGVAGI
>AT4G16590_CslA1
MVDLVVQVVDDFTDPAVREGVDVEIAKWQSQGINIRCERRDNRNGYKAGAMKEALTQSYVKQCDFV
AVFDADFQPEPDYLIRAVPFLVHNPDVALVQARWIFVNANKCLMTRMQEMSLNYHFKVEQESGSTR
HAFFGFNGTAGVWRISAMEAAGGWKSRTTVEDMDLAVRVGLHGWKFVYLNDLTVRNELPSKFKAY
RFQQHRWSCGPANLFRKMTMEIIFNKRVSIWKKFYVIYSFFFVRKVAVHFLTFFFYCIIVPTSVFFPEIHI
PSWSTIYVPSLISIFHTLATPRSFYLVIFWVLFENVMAMHRTKGTCIGLLEGGRVNEWVVTEKLGDALK
SKLLSRVVQRKSCYQRVNSKEVMVGVYILGCALYGLIYGHTWLHFYLFLQATAFFVSGFGFVGT
>CSLA_7BL_TGACv1_579090_AA1903960.1_Triticum_aestivum
MSTLPRVWQIAAAWEQVRGPVIVPLLRVSVLLCLAMSAMLFAEKVYMAVVVLAVRLLGRRPEQQYR
WEPVGDGNDPELGSAAYPMVLVQIPMYNEREVYQLSIGAACGLSWPSDRIIVQVLDDSTDPVIKELV
QVECRRWARKGVNIKYEIRDNRRGYKAGALKEGMKHGYVKDCDLVAIFDADFQPEPDFLSRSVPFLV
HNPDIALVQARWKFVNADECLMTRMQEMSLDYHFKVEQEVGSSTYAFFGFNGTAGVWRISALNEAG
GWKDRTTVEDMDLAVRASLKGWKFVYIGDLKVKNELPSTFKAFRYQQHRWSCGPANLFRKMVMEIV
RNKKVTLWKKIHVIYNFFLVRKVVAHIVTFVFYCVVERIDMVD
>CSLH_3AS_TGACv1_212952_AA0704280.1_Triticum_aestivum
MIDISSRIQVAKQVSSCNYETDTHWGQEIGWSYGSMAEDILTGQRIHSSGWKSTLLDTNPPAFLGCA
PTGGPASLTQYKRWATGLLEILLGQNSPIIATIFKRLQFRQFLAYLVFYVWSMRAPFELCYALLGPFCL
FRNQSFLLKASNHGFSIQLALFLSYNIYNFVEYMDCGLSARTWWNNMRMQRIVSISSWLLAFLSVVLK
TIGLSKTVFEVTREDKSTSDGDPSTHETDLGWFTFDSSLVFIPVTTVAILNIATIAIGVWRHAIFWMITGN
HDWQNIGEFICCGWAILYFWPFIKGLVGRGRYGIPWNVKLKAWVIVVAFLYFCRGD
>CSLD_1BS_TGACv1_049706_AA0160220.1_Triticum_aestivum
MSCKMRGCDMLALAATRPMICEECYMDCVAASGNCPGCKEAYSAGSDTDDSVDEDDDDAISSSEE
RDQMPMTSMSKRFSMVHSIKMPMSSSNDKPADFDHARWLFETKGTYSYGNALWPENEHGGGGNN

AGATFGFVGIEEPPNFATWMSDGSQWASTWLAGATDHARGNHAGIIQRPGYNHNKKAGAMNALVR
TSAIMSNGPFILNLDCDHYVHNSAALREGMCYMLDCRGDRVCYVQFPQRFEGIDPNDRYANHNLVFF
DVAMRAMDGLQGPMYVGGCIFRRIALYGFSPPRATKHHGWLGRKIKLFLRKPTMGKKTDRELVMAIL
QK
>GRMZM2G020742_P01
MTLDYHFKVEQEAGSSTFGFFGFNGTAGVWRTSSIKEAGGWEDRTTVEDMDLAVRAGLKGWKFIYV
GDVKVKSELPSNLKAYRRQQHRWTCGAANLFRKMGAEIILTKEVSLWRKLYLIYSFFFIRKVVAHVVP
FMLYCVIIPLSVLIPEVTVPVWGVVYIPTTITLLYAIRNPSSIHFIPFWILFENVMSFHRTKATFIGLLELGN
VNEWVVTEKLGRTKPVPQMLEKPRCRFWDRCTISEILVAIFLFFCATYNLVLGDDFYFVYIYLQEIAFLI
VGTGFCGT

**Fig. S19.** FASTA sequences of Csl proteins of ALO used for the phylogenetic analysis.

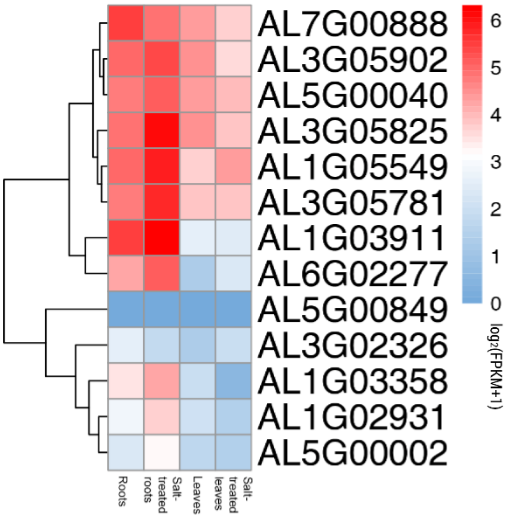

**Fig. S20.** Heat map showing hierarchical clustering of the ALO *callose* gene (*CalS*, *GSL*) families in roots, salt-treated roots, leaves and salt-treated leaves.

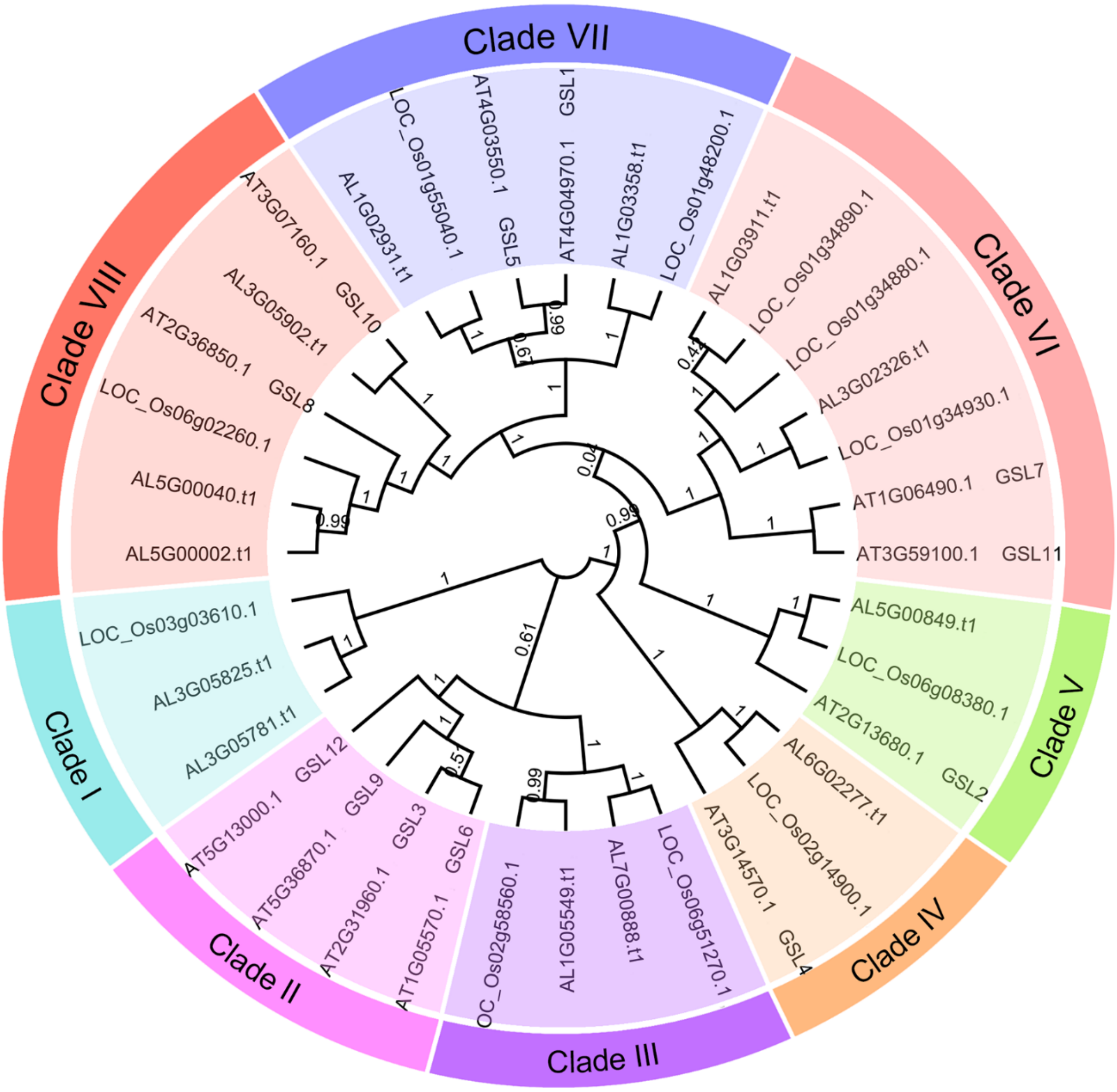

**Fig. S21.** The maximum likelihood phylogenetic tree constructed with CalS (GSL) proteins of ALO, *Arabidopsis thaliana* and *Oryza sativa*.

>AL7G00888.t1
MAAPGGRRPDFSSPGPSPAGPSSAGAAGGRRLLRTQTVGNMGESIFDSEVVPSSLVEIAPILRVANEVEA
GNPRVAYLCRFYAFEKAHRLDPTSSGRGVRQFKTALLQRLERENDPTLKGRVHQSDAREMQRFYREYY
KKYIQALQNAADKTDRALLTKAYQTAAVLFEVLKAVNVSQSVEVDQAILDTHHKIEEKKKLYVPYNILPLDPE
SNDQAIMQHSEIQAAVYALRNIRGLPWPKDKDKDKDKDKGNERKPEEKKTDKDLLDWLQAMFGFQKDNV
SNQREHLILLLANVHIRQIPKPEQQSKLDDRALDAVMKKLFKSYKMWCKYLGRKSSLWLPTIQQEVQQRK
LLYMGLYLLIWGEAANLRFMPECLCYIYHHMAFELYGMLAGNVSPMTGENVKPAYGGDNEAFLMKVVTPI
YKVIEEEALRSKTMKSKHSHWRNYDDLNEYFWKNDCFRLGWPMRDDADFFKTPKVAYPNHLNGEERSA
GSVHWMGKINFVEIRSFWHIFRSFDRMWIFLILSLQAMIIIAWNGGTPSDIFDAGVFKQVLSIFITAAVLKLGQ
ATLDIVFGWKARRSMSFARKLRYVLKLVSAAAWVVILPVTYAYSGGSPTGLTRIIKGWLGNGQNQSSLYIL
AVLIYLAPNMLVAMLFIFPFIRRSLESSNLKVVTFMMWWSQPRLFVGRGMHEGAFSLLKYTMFWIILLAMK
LTVSFYIEIKPLVQPTKDIMREPIRTFQWHEFFPNGNNNIGVVIALWAPIILVYFMDTQIWYAVFSTLVGGIYG
ACRRLGEIRTLGMLRSRFESLPSAFNKLLIPSDAHKRKGFRAAFASKLAKPSDDEQERDKIAARFAQMWN
LIITSFREEDLIDNREMDLLLVPYCKDRELKLFQWPPFLLASKIPIALDMAADSGGKDRDLNKRMGSDAYFS
YAIKECYASFKNIINTLVSGKREKIVMKEIFAVVDKHISEETVIKDLNMRNLPALSKKLIELLELLQKNKEEDLG
QVVILFQDMLEVVTRDIMEDQELGGVLDSMHGGNARKHETMTPLDQQDQLFTKAIKFPVEESNAWTEKIK
RLHLLLTVKESAMDVPTNLDARRRISFFANSLFMEMPNAPEVRNMLPFSVLTPYYKEDVLFSSHNLEEPNE
DGVSILFYLQKIYPDEWKNFLERVNRKTEEEVREDETLEDELRLWASYRGQTLTRTVRGMMYYRKALELQ
AFLDMAKDDDLMKGYKATELMSEDSPLMTQCKAIADMKFTYVVSCQQYGIQKRSGDPCAHDILRLMTTYP
SFRVAYIDEVEAPSQDRNKKIEKVYYSVLVKASVTKSNDPGQSLDQVIYKIKLPGNAILGEGKPENQNHAIIF
TRGECLQTIDMNQEHYMEEALKMRNLLEEFLEKHDGVRYPSILGVREHIFTGSVSSLAWFMSNQETSFVTI
GQRVLANPLRVRFHYGHPDIFDRLFHLTRGGISKASKIINLSEDIFAGFNSTLREGNVTHHEYMQVGKGRD
VGLNQISLFEAKIANGNGEQTLSRDIYRLGHRFDFFRMLSCYYTTIGFYFSTMITVWTVYVFLYGRLYLVLS
GLDYALATGRRFVHNESLQVALASESFVQLGFLMALPMMMEIGLERGFRTALSDFVLMQLQLASVFFTFS
LGTKTHYYGRTLLHGGAEYRATGRGFVVFHAKFAENYRLYSRSHFVKGIELMILLIVFEIFGQPYRGAITYIF
ITVSMWFMVGTWLFAPFLFNPSGFEWQKIVDDWADWNKWISNRGGIGVAPEKSWESWWDKEQGPLRH
SGKRGTILEILLALRFFIYQYGLVYHLNITKQYKQSVLVYGFSWVVILFMLLVMKTVSVGRRRFSAEFQLVFR
LIKGLIFITLITVIVTLTALAHMTVLDIFVCILAFMPTGWGLLLIAQAIKPAVEIAGLWGSVKALARGYEILMGLLL
FTPIAFLAWFPFVSEFQTRMLFNQAFSRGLQISRILGGHKKDKATRNKE
>AL1G05549.t1
MASSSGRRPGPGGGGGGGGGEPSSASPAPGGGGRRILRTQTAGNLGESSFDSEVVPSSLVEIAPILRVA
NEVENSNPRVAYLCRFYAFEKAHRLDPTSSGRGVRQFKTALLQRLERENDPTLKGRVKQSDAREMQSFY
QHYYKKYIQALQNAADKADRAQLTKAYQTAAVLFEVLKAVNVSQKIELDESIVETNKQVEEKKKLYLPFNIL
PLDPDSAHQPVMQFPEIQVAFYALRNTRGLPWPKDHEKKAGADLLDWLQAMFGFQKDNVSNQREHLILL
LANVHIRQIAKPDQQPKLDDHALDQVMKKLFKNYKRWCKYLGRKSSLWLPTIQQEVQQRKLLYMGLYLLI
WGEAANLRFMPECLCYIYHHMAFELYGMLAGNVSPTTGENVKPAYGGAEEAFLKLVVTPIYKIIEMEAERS
KTIKSKHSHWRNYDDLNEYFWSRDCFRLGWPMRADADFFKTPNFALARDQMSGENRPVGSDQWMGKV
NFVEIRSFWHIFRSFDRMWSFLILSLQAMIIIAWNGGTPSDIFDAGVFKQVLSIFITAAVMKFGQAILDVVLS
WKARKSMSLAVKLRYILKLLSGAAWVVIFPVTYAYSSENPTGLSRTIKSWFGDGRGQPPLYGLAVVIYLAP
NMLAATLFLFPVLRRALERSNLKVITFIMWWSQPRLFVGRGMHEGAFSLFKYTMFWVLLLATKLVVSFYVE
IKPLVQPTKDIMKVPITTFQWHEFFPHAKNNIGVVIALWAPIILVYFMDTQIWYAIFSTLVGGIYGACRRLGEI

RTLGMLRSRFESLPKAFNDHLIPNESKRRGFRSAFSSKHSQKPEDSKEEEKIAARFAQIWNLIITSFRDEDL
IDNREKDLLLVPYCKDREMDIIQWPPFLLASKIPIALDMAADSGGKDRDLKKRMKSDPYFTYAIKECYASFR
NIIYALVVRPREREFIEKIFKMVDDHIADDTLIKELHMSNLPTLSKKFIELLDILQKNNKEDQGQVIILFQDMLE
VVTRDIMEEQLSGLLETVHGGNNRKHEGITPLDQQEQEQLFTKAIEFPVKKSHAWSEKIKRLHLLLTVKES
AMDVPTNLDARRRISFFANSLFMEMPRAPKVRQMLPFSVLTPYYKEDVLFSSQALEDQNEDGVSILFYLQ
KIYPDEWKNFLERVECKEEELRETEQSEDELRLWASYRGQTLTRTVRGMMYYRQALVLQSFLDMAREDD
LMEGFRAADILSEESKLLTQCKAIADMKFTYVVSCQQYGIQKRSGDHRAQDILRLMTTYPSLRVAYIDEVE
EPTKDRSKKIEKVYYSALVKAAVTKPDDPGQKLDQDIYRIKLPGNAMLGEGKPENQNHAIIFTRGEGLQTID
MNQEHYMEETLKMRNLLQEFMKKHDGVRYPSILGVREHIFTGSVSSLAWFMSNQETSFVTIGQRVLANPL
RVRFHYGHPDIFDRLFHLTRGGVSKASKIINLSEDIFAGFNSTLREGNVTHHEYMQVGKGRDVGLNQISLF
EAKIAYGNGEQTLSRDVYRLGHRFDFFRMLSCYYTTIGFYFSTMITVWTVYVFLYGRLYLVLSGLDDQLAT
GKRFNHNNNLRVALASQSFVQLGFLMALPMMMEIGLERGFRTALSDFVLMQLQLASVFFTFSLGTKTHYY
GKTLLHGGAEYRATGRGFVVFHAKFAENYRLYSRSHFVKGIELMILLIVFEIFGQSYRGAIAYIFITFSMWFL
VFTWLFAPFLFNPSGFEWQKIVDDWTDWNKWISNRGGIGVSPEKSWESWWEKEHEPLKHSGKRGTVLE
IVLAARFFIYQYGLVYHLHITKKTQSVLVYCLSWVVIFVILMVMKGVSVGRLKFSAEFQLVFRLIKGLIFITFIT
VVIILIALAHMTVQDIFVCILAFMPTGWGLLLVAQAIKPAIVKLQLWGSIRALARGYEIIMGLLLFTPIAFLAWFP
FVSEFQTRMLFNQAFSRGLQISRILGGHKKDRAARNKE
>AL6G02277.t1
MELEIVEIGPSSRPLRYVPAPAPPQWRPPPPPPPLPPPSMAASSSARTSEQATASAVAQALQFDSEKLPQ
TLVSEIRPFLRVANQIEAESPRVAYLCRFHAFEKAHMMDARSTGRGVRQFKTALLQRLEQDEKSTFSKRK
ERNDAREIKSFYEKKRQANAHELMPVLAEVLKAVLIGTGLESVVASEDFADKSGLFRYNIIPLHPRSSQQPI
MLLQEIKVAVSAVFNVRSLPLANIKDEKTHTDIIRWLQSWFGFQKGNVANQREHLILLLANMHARMNPKSS
SAPMLDDRAVDELLAKTFENYLTWCKFLGRKSNIWLPSVKQEIQQHKLLYIALYLLIWGEASNLRLMPECL
CYIFHHMSYELYGVLSGAVSLITGEKVRPAYGGEDESFLNKVVTPIYAEISAEALKNKNGVSDHSTWRNYD
DLNEFFWSADCFKLGWPMRLNNDFFFTSTKNKMSPQTEIQYSQLPHGSSSAQSIANSEVPDQTQQQTAS
DTSQQRWLGKTNFVEVRSFWHLFRSFDRLWTLLVLGLQILIIMAWHGLESPLQLLDPNFFEDLLSIFITNAV
LRVIQVVLDIAFSWRTKGTMRSSQKLRFAIKLSLAITWAIILPIFYSSSQNYKACSARRPKNFLGMFCLSKYM
VVVELYLTSNVIGMALFFVPAVSSYIETSTWRICNILSWWCQPQSYVGRGMHEGQVPLLKYTSFWTILLSS
KFLFSYYFEIKPLVEPTKIIMNINVKTYEWHEFFPQVKSNAGAILAVWSPIVLVYFMDTQIWYSVFCTIFGGM
CGIVHHLGEIRTMGMVRSRFCTLPEVFNACLVPRSMLKEKKGILPSFLEKKIFKNLGKSERQDPTKFALVW
NQIINSFRSEDLISNREMDLMTMPMSLEYSSGSIRWPLFLLAKKFSKAVDMAANFTGKSEQLFWRIKKDNY
MFCAINDFYELTKGIFRFLIIGDVEKRVIAAIFAEIKKSIQNSTLLIDFNMDYLPLLVEKIERLAELLYKNKQGLR
YEVTILLQDIIDTLIQDMLVDAQRVLDQINSSDTLISDDDGVFDYYKPELFASISSISKIRFPFPDSGPLKEQVK
RLYLLLNTKEKAAEVPSNSEARRRISFFATSLFMDMPAAPKVRSMLSFSIVTPYFMEEVKFSDEELHSNQD
EASILSYMQKIYPDEWTNFLERLGLKVTNEDIRYWASFRGQTLSRTVRGMMYYRKALRLQAFLDRTNDQE
LYKGPAGTERGQNKKNIHQSLSTELDALADMKFSYVISCQKFGEQKSIGDPHAQDIIDLMARYPALRVAYIE
EKEIIVQNMPHKVYSSVLIKAENNLDQEIYRIKLPGPPIIGEGKPENQNHAIIFTRGEALQTIDMNQDNYLEEA
YKMRNVLQEFVRRPKDQSPTILGLREHIFTGSVSSLAGFMSYQETSFVTIGQRFLADPLRVRFHYGHPDIF
DRMFHLTRGGISKASKTINLSEDVFAGYNSILRRGHITYNEYIQVGKGRDVGLNQISKFEAKVANGNSEQTL
SRDIYRLGRRFDFFRMLSCYFTTVGFYFNSLISVVGIYVFLYGQLYLVLSGVQNVLMVKAKTLNMKSLETAL
ASQSFLQLGLLTGLPMVMELGLEKGFRAALSDFILMQLQVASVFFTFSLGTKAHYYGRTILHGGAKYRPTG

RKFVVFHASFTENYQLYSRSHFVKAFELLFLLIIYHIFRKSYGKIHVMVTYSTWFMAMTWLFAPFLFNPAGF
AWHKIVDDWADWNRWMRNQGGIGVQPEKSWESWWNAENAHLRYSVLSSRILEVLLCLRFFVYQYGLVY
HLKISHDNKNFLVYLLSWVVIIAIVGLVKLVNCASRQLSSKHQLVFRFIKLLTFLAVVTSFILLSCLCKLSIMDLI
ICCLAFIPTGWGLLLIVQVLRPKIEYYAIWEPIQVIAHAYDYGMGSLLFFPIAILAWMPIISAIQTRVLFNRAFS
RQLQIQPFIVGKTKRK
>AL5G00849.t1
MTTPRGGADEPSSAAAAAAAGNDPPTPTSAHASGLNRRGSRSAAMASFSMEVFDNEVVPSALNSIAPILR
VAAEIEHERPRVAYLCRFYAFEKAHRLDQNSVGRGVRQFKTALLQRLEKEWSELPCPNALINQRAQLGKA
YQTAGVLFEVLCAVNKNEKVEEMNPEIMRFHKEVQEKKDIFTPFNILPLDTASASQSIMQLEEIKAAVKALR
YIQGLTWPSAFEPERQKGGELDLLDWLRAMFGFQRDSVRNQREHLILLLANVHIRLEPKPDPLSKACSSID
IYFDSLTMDINLDDRAVDVLMNKLFNNYRKWCKFLSRKHSLRNPPGAQAQEVQQRRILYLGLYLLIWGESA
NIRFMPECLCYIFHNMAYELHGLLAGNVSIVTGENIRPSYGGDEESFLKKVVTPIYRVIRKEAEKSKHGKTP
HSAWCNYDDLNEYFWTSDCFSLGWPMRDDGEFFKSVHDSRRVTVAGSSSPKGSSKSTGKTNFVETRSF
WHIFRSFDRMWTFYLLALQAMLIFAWSGYSVTEIFRKDLLYSLSSIFVTAAFLQFLQSILDLILNFPGHIRCKF
IDIVRNVLKITVSAVWAVILPFFYISTAGNKVNLQLRGLEKLLAYVKGVPTLYILAVAVYLIPNMISAALFLFPM
FRRWIENSDWHIVRLLLWWSQKRIYVGRGMHESQAALLKYTLFWILLLCSKLSFSYFVQIEPLIKPTKDIMS
VHNIRYEWHEFFPNASYNIPAILSLWAPVLLVYFMDTQIWYAIFSTISGGMSGALGRLGEIRTLGMLRSRFH
SLPGAFNTFLVPSDKGRSRRFSLSKRFAEVSPNKRTEAAKFAQLWNEVICSFREEDFISDKEMDLLVVPYS
SDPSLKLMQWPLFLLASKIPIALDMAAQFRPKDSDLWKRICADEYMKCAVIECYESFKLVLNLVVVGENEK
RIIGIIIKEIEANIAKNTFLANFRMSTLPVLCNKFVELVSTLKEGDASKFDNVVLLLQDMLEVITRDMMVNEIKE
LAEFGHGNKDLVPRRQLFAGTGTKPAIVFPPPISAQWEEQIKRLYLLLTVKESAMDVPTNLEARRRISFFTN
SLFMDMPRAPRVRKMLSFSVMTPYYSEETVYSRNDLDLDNEDGVSIIFYLQKIFPDEWNNFMERINCKRE
SEVWVNEENVLQLRHWASLRGQTLCRTVRGMMYYRKALKLQAFLDMASESEILEGYKAVADPADEEKKS
QRSLSSQLEAIADMKFTYVATCQIYGNQKQSGDRHATDILNLMVNYPGLRVAYIDEVEEREGEKVQKVFY
SVLVKALDNHDQEIYRIKLPGPAKLGEGKPENQNHAIIFTRGEALQTIDMNQDNYLEEALKMRNLLEEFNES
HGVRPPTILGVREHIFTGSVSSLAWFMSNQETSFVTIGQRVLANPLKVRFHYGHPDVFDRIFHITRGGISKA
SCGINLSEDIFAGFNSTLRRGNVTHHEYIQVGKGRDVGLNQISLFEAKVACGNGEQVLSRDIYRLGHRFDF
FRMLSCYFTTVGFYVSSMMVVIIVYVFLYGRLYLALSGLEFAIMKQARMRGNRALQAAMGSQSIVQLGLLM
ALPMFMEIGLERGFRSALGDFIIMQLQLCAVFFTFSLGTKSHYFGRTILHGGAKYKATGRGFVVRHVKFAE
NYRMYSRSHFVKGLELMLLLVVYELYGDVATDSTAYVLLTSSMWFLVITWLFAPFLFNPSGFEWQKVVDD
WDDWNKWISSRGGIGVPATKAWESWWEEEQEHLQSTGLLGRIWEIILSFRFFIFQYGIMYHLNISNGNKSI
SVYGLSWLVIVAVVLLLKVVSMGRKKFSADFQLMFRLLKLFLFIGSVGTLAILFTVLHLTVGDIFASFLAFVPT
GWAILQISQASKPVVKALGLWGSVKALSRGYEYLMGIVIFVPVAVLAWFPFVSEFQTRLLFNQAFSRGLQI
SRILAGGKKQN
>AL5G00040.t1
MASSMRRRPSRGGGGPERVAENWERLVRAALKRDRAGGAGPGSSAAAGHAAGGGGVGLASAVPPSL
GRTTNIEQILQAADDIEDDDPNVARILCEQAYTMAQNLDPSSGGRGVLQFKTGLASVIKQKLAKKDGASID
RQNDIEVLWNFYLEYKRRRRVDDMQREQDMMRESGTFSTEMGARAVEMKKIYATLRALLDVLEILVGQA
PTDRLGRQILEEIKKIKRSDAALRGELMPYNIIPLDSASVGNVVGFFPEVRAAIAAIQNCEDLPRFPYDTPQL
RQKDIFDLLQYVFGFQDDNVRNQRENVTLTLANAQSRLSLPNETEPKIDERAVTEVFCKVLDNYIKWCRFL
GKRVAWTSLEAVNKNRKIILVALYFLIWGEAANVRFLPECLCYIFHNMAKELDGILDSSEAEPAKSCITSDG

STSYLEKIITPIYQTMSAEASSNNDGKAAHSAWRNYDDFNEYFWSRSCFELNWPPAEGSKFLRKPAKRKR
TGKTNFVEHRTFLHLYRSFHRLWIFLILMFQCLAIIAFHDGKINLSTIKVVLSAGPAFFILNFIECCLDVLLMFG
AYKTARGFAISRIVIRFIWLAAVSTFVTYLYVKVLDEENARNSDSTYFRIYVLVLGSYAAVRLVFALLAKIPAC
HRLSDFSDRLQFFQFFKWIYQERYYIGRGLYESISDYARYVIFWLVIFACKFTFTYFLQIRPLVVPTNIIVQLH
DLEYSWHDFVSKGNKNALTILSLWAPVVAIYLMDIHIWYTLLSALIGGVMGARDRLGEIRSIEMLHKRFESF
PEAFAKTLSPQRISNRPVAQDSEATKTYASIFSPFWNEIIKSLREEDYVSNREMDLLMMPSNCGNLRLVQ
WPLFLLTSKIMLANDYASDCKDSQYELWHRISKDEYMAYAVKECYYSAERILNSLVDAEGQRWVERLFRD
LNESIAQGSLLVTINLKKLQLVQSRLTGLTGLLIRDETADRAAGVTKALRELYEVVTHEFLAPNLREQFDTW
QLLLRARNDGRLFSNIFWPKDLEMKEQVKRLHLLLTVKDSAENIPKTLEATRRLQFFTNSLFMDMPEAKPV
SEMIPFCVFTPYYSETVLYSMSELCVDNEDGISILFYLQKIFPDEWANFLERIGRGESSEEDFKESSSDTLE
LRFWVSYRGQTLARTVRGMMYYRRALMLQSYLEKRYLGGIEDGYSAAEYIDTQGYQLSPDARAQADLKF
TYVVSCQIYGQQKQRKAPEAADIALLMQRNEALRVAFIHEEDSVSSDGHAIKEYHSKLVKADIHGKDQEIY
SIKLPGNPKLGEGKPENQNHAIIFTRGDAVQTIDMNQDNYLEEAMKMRNLLEEFRGSHGIRYPTILGVREH
VFTGSVSSLASFMSKQETSFVTLGQRVLAYLKVRMHYGHPDVFDRIFHITRGGISKASRVINISEDIYAGFN
STLRQGNITHHEYIQVGKGRDVGLNQIALFEGKVAGGNGEQVLSRDVYRLGQLFDFFRMLTFFYTTVGFY
VCTMMTVWTVYIFLYGRVYLALSGLDYSISRKARFLGNTALDAALNAQFLVQIGIFTAVPMIMGFILELGLMK
AIFSFITMQLQFCSVFFTFSLGTRTHYFGRTILHGGAKYRATGRGFVVRHIKFADNYRLYSRSHFVKALEVA
LLLIVYIAYGYTKGGSSSFILLTISSWFMVMSWLFAPYIFNPSGFEWQKTVEDFDDWTNWLFYKGGVGVR
GENSWESWWDEEQAHIQTFRGRILETILSLRFLLFQYGIVYKLKLTGDNTSLAIYGFSWIVLLVLVLLFKLFT
ATPKKSTALPTFVRFLQGLLALGIIAAIACLIGFTDFTIADLFASALAFLATGWCILCLSITWKRLVKTLGLWDS
VREIARMYDAGMGAIIFSPIVFFSWFPFVSTFQSRILFNQAFSRGLEISLILAGNKANQES

>AL3G05902.t1

MARAEANWERLVRAALRGERMGGGYGVPASGIAGNVPTSLGNNTHIDEVLRAADEIQDEDPTVARILCEH
AYALAQNLDPNSEGRGVLQFKTGLMSVIRQKLAKREGGAIDRSQDIAKLQEFYKLYREKHKVDELCEDEM
KLRESAVFSGNLGELERKTLKRKKVLATLKVLWSVIEDITREISPEDADKLISEEMKKVMQKDAARTEDVVA
YNIIPLDALSTTNAIVTFPEVRAAISALQYHRDLPRLPATISVPDARNSDMLDLLHCVFGFQKGNVSNQREHI
VHLLANEQSRLGKLSGNEPKIDEGAVHVVFSKSLDNYIKWCNYLPLRPVWNNTESLTKEKKLLYVCLYYLI
WGEAGNIRFLPECLCYIFHHLARELEENMRKQTAEPAESCISNDGVSFLGQVISPLYEIIAAEAANNNNGRA
AHSAWRNYDDFNEFFWSSKCFQLGWPWKLNSPFFSKPTRKEGLLHRNHHYGKTSFVEHRTFLHLYHSF
HRLWMFLIMMFQGLTIIAFNNGSFDTNTALELLSLGPTYVVMKFIESVLDILMMYGAYSTSRGSAITRVIWR
FFWFTMASFVICYLYIKALQDGAQSAFFKIYVVVISAYAGVQIIIGLLMSFPCCRGVTNACYSWSFVRLIQW
MHQEHNYVGRGMHERPLDYIKYVAFWLVILAAKFSFTYFLQESLIQPLVEPTRLITSFKGLQYQWHDFFSK
NNHNAFTILFLWAPVVSIYLLDIHVFYTIMSAIVGFLLGARERLGEIRSVEAVHRFFEKFPEAFMDKLHVPVP
KRKQLLSSGQDAELSKFDASRFAPFWNEIVRNLREEDYINNTELELLLMPKNKGGLPIVQWPLFLLASKVF
LAKDIAVDCKDSQDELWLRISKDEYMQYAVEECFHSIYHILTSILDKEGHLWVQRIFGGIRESISKKNIQSDI
HFSKLPNVIAKLVAVAGILKETESADMKKGAVNAIQDLYEVVHHEVLSVDMSGNIDDWNQINRARAEGRLF
SNLKWPNDPGLKDLIKRLHSLLTIKESAANVPKNLEACRRLEFFTNSLFMRMPLARPVAEMLSFSVFTPYY
SETVLYSIAELQKRNEDGISTLFYLQKIYPDEWKNFLTRINRDENAAESELFSSANDILELRLWASYRGQTL
ARTVRGMMYYRKALMLQSYLERMHSEDLESAFDMAGLADTHFEYSPEARAQADLKFTYVVTCQIYGLQK
GEGKQEAADIALLMQRNEALRIAYIDVVESIKNGKPSTEYYSKLVKADIHGKDKEIYSVKLPGNPKLGEGKP
ENQNHAVIFTRGNAVQTIDMNQDNYFEEALKMRNLLEEFSQDHGKFKPSILGVREHVFTGSVSSLASFMS

SQETSFVTLGQRVLSNPLKVRMHYGHPDVFDRVFHITRGGISKASRIINISEDIFAGFNSTLRQGNITHHEYI
QVGKGRDVGLNQIALFEGKVAGGNGEQVLSRDIYRLGQLFDFFRMLSFYVTTIGFYFCTMLTVLTVYIFLY
GKTYLALSGVGESIQNRADILQNKSLTVALNTQFLFQIGVFTAIPMILGLILEAGVLTAFVNFITMQFQLCSVF
FTFSLGTRTHYFGRTILHGGAKYRATGRGFVVRHIKFAENYRLYSRSHFVKGLEVALLLVIFLAYGFNDGG
AIGYILLSISSWFMALSWLFAPYIFNPSGFEWQKVVEDFRDWTNWLFYRGGIGVKGEESWEAWWDEELG
HIQTFRGRILETILSLRFFIFQYGVVYHMDASEPSTSLLVYWVSWAVLGGLFILLMVFSLNPKAMVHFQLLL
RLVKSIALLMVLAGLIVAIVSTRLSFTDVLASILAYVPTGWGILSIAVAWKPVVKRLGLWKTVRSLARLYDAG
MGMIIFVPIAICSWFPFISTFQTRLLFNQAFSRGLEISLILAGNNQNTSV
>AL1G03911.t1
MSRAITMRTDGFSGEEPGDIIESELVPSSLAPIVPILRAANEIEEENPRVAYLCRFTAFEKAHTMDPNSSGR
GVRQFKTYLLHRLEKDEQETKRRLASTDAKEIQRFYEQYCRKNLEEGLNMRKPDEMARYYQIASVLYDVL
KTVTPEKSHSEFDQYAKGVEKEKASFSHYNILPLNISGPTQPVMEIPEIKAAVDLLRGMESLPTPRPDMNT
VPEETDRPIVRDLLDWLWQTFGFQKGNVENQKEHLILLLANIDMRESGSVDQGRRQNHVIHSNTVKHLMN
KVFQNYISWCRYLHLESNIKIPTDASTQQPELLYIGLYFLIWGEASNVRFMPECLCYIFHHMARDLYDIISDR
REGSFDPPFRREGSDDAFLQLVIQPIYSVIHNEAMMSKKGTVSHSKWRNYDDLNEYFWSKKCFKQLGWP
MDPASDFFGDPTKIKDDIERHDRSISRRRMSKTNFVEVRTFFNLFRSFDRMWAFFILALQAMIIIAWSPSGS
LSAIFDPTVFRNVMTIFITAAFLNFLQATLEIILNWKAWRSLVCSQMIRHVLKFVVAIGWLIILPVTYASSIQNP
TGLIKFFSNWIGKFQSQSIYNFAVALYMLPNIFSALFFIFLPIRRALERSNSRIVRFLLWWTQPKLYVARGMY
EDTCSLLKYTTFWILLLICKLSFSYYVEISPLVGPTKIIMFLGRGTYVWHEFFPYLQHNLGVVFTIWAPIVMVY
FMDAQIWYAIFSTICGGVNGAFSRLGEIRTLGMLRSRFEAIPTAFGKNLVPRHGSQSKRREREEDKKLHID
KFSDIWNAFIISLRDEDLISNRERDLLIVPSSVGDTSVFQWPPFLLASKIPIALDMAKSVKKRDEELKKRIKQD
PYTYYAVIECYETLLNILYSLTEEASDRKVVDRIRDSLEDSIRRQSLVKEFRLDELPQLSAKFDKLLTLLLKTE
QEEHDTIKTQIANLLQDTMEIITQDIMKNGQGILKDENRNNQLFANLNLDSIKDEAWREKCVRLQLLLKTKE
SAIYVPTNLEARRRITFFANSLFMKMPRAPQVRSMMSFSVLTPYFKEEVLFSSEDLHKKNEDGISILFYLRKI
YPDEWKNCLERIKFKPKDEESLKLKMDEISPWASYRGQTLTRTVRGMMYYRRALEIQCIQDKIDIAKLDRQ
RTPASYQEGASIVDMALAIADIKFTYVVSCQVYGMQKISKDLKDKACYLNILNLMITYPSLRIAYIDEVEAPTR
NGTTEKTYYSVLVKGVGEKYDEEIYRIKLPGKPTDIGEGKPENQNHAIIFTRGEALQAIDMNQDNYLEEAFK
MRNVLEEFGSENYGKSKPTILGLREHIFTGSVSSLAWFMSNQETSFVTIGQRVLANPLKVRFHYGHPDIFD
RLFHITRGGISKASKTINLSEDIFSGFNSTMREGNITHHEYMQVGKGRDVGMNQISSFEAKVANGNGEQTL
SRDIYRLGRRFDFYRMLSFYFTTVGFYFSSMVTVLTVYVFLYGRLYLVMSGLEKSIFQDPRIQDNIKTLENA
LASQSVFQLGLLLVLPMVMEVGLEKGFRTALGEFIIMQLQLASVFFTFQLGTKTHYYGRTILHGGAKYRPT
GRGFVVYHAKFADNYRMYSRSHFVKGLELLILLVVYLAYGKSYRSSRLYLFVTFSIWFLVASWLFAPFIFNP
SCFEWQKTVDDWTDWRKWMGNRGGIGMSADQSWEAWWTSEQEHLRKTSIRALFLEILLSLRFLIYQYGI
VYHLNIARRNTSILVYGLSWLVMLTVLVVLKMVSIGRQKFGTDLQLMFRILKGLLFLGFVSVMAVLFVVCNL
TISDVFASILGFLPTGWCILLIGQACSPLIKKTTLWDSIMELGRAYDNIMGLVLFLPIGFLSWFPFVSEFQTRL
LFNQAFSRGLQISRILAGQKDIGEFE
>AL3G02326.t1
MSAMAVRRRRNVLAQTLASRRLPEGVAEPGERVPDALAPEVMPFVRAADEIEPLNPRVAFLCRKYAYKK
VQRMDPSSIQRGVRQFKTYMSLKLDQDDTHFLVNDAKEIQQFYKHYCDNLSRTSQRRNFDELARYYQVA
SALYEVLRDVTKNKVDSEVMKRAKVVEEKSGHFRSYRYNILPLSFPGSSEAVLELPEIKSAIDAIRNIYGLPK
PHMSSIPREGKSIQDLLDWLSLAFGFQNSNVENQRENIVLLLANISTRTSGQEGFPLVDAVNDLGEKIFGN

YESWCHYLHLSSRIMVNYDDALKKQQLMLLYIGLYLLIWGEASNVRFMPECLCYIFHHMAKQLNQMVEEN
YFQPPPGFEEEGSFLKVAIEPIYKVLQKESQRSKGGTAGHSAWRNYDDLNEQFWSEKCFMKLGWPWEL
SADFFHQAGRTGRKPKTNFVEVRTFLHLFRSFNRMWMFFILAFQAMLIVSWSSSGSLSGIADATVFRSILS
VFITAALLNFIKVTLDIMLTIQAWGNMEWTQIIRYLLKFFVAIAWIIILPVTYSSSIKNPSGAGKLLNSWTRNWY
DQSVYNFAIVIYMIPTILAAFLFLLPQLQNVMERSNSRAFVLLMWWIQPRLYVARGMHEDILSIFKYVFFWV
VLLTCKLAFSFYVEISPMIEPTKFILNQVVGNYEWHEVFPFLPHNLGVVITIWAPIVLVYFMDTQIWYAIFSTV
FGGVSGALSHVGEIRTLGMLRVRFKSMPDAFRKCHVATHKERALDVRSFFCVWNSFINSLREEDFISDRE
KDILMAPSSSNNLSVVPWPPFLLASKVPTALHMAMTSKEGDVHELIEKIKLDKDRYNAVIECYESLKNIVVS
LLLDYNDKKIVHDIDNIVQISMQNNTLLEDFEMAEIGKVSNTLAKLLQLLKSEPTDGSTERKIVNALQDFMEIL
TRDFMKDGHGILKDENERKQSFTNLNMNVIKEAFWREQFVRLHLLLTMKDSAMDVPTNLDARRRITFFAN
SLFMKMPRAPQVHDMISFSVLTPYYNEEVLYSSHELNRKNEDGISILFYLQKIYPDEWKNFLERIGCDPDN
EEAVKGCMDDIRIWASYRGQTLARTVKGMMYYRRALELQCYEDMKINQADLGGDESARSKAIADIKFTYV
VACQLYGMHKASKDSRERGLYENILNLMLTYPALRIAYIDEKEEPSSNGKLEKHYYSVLVKGDDEEIYRIKL
PGKPTEVGEGKPNNQNHAIIFTRGEALQAIDMNQDNYLEEAFKMRNLLEEFLIKHGKSKPTILGVREHIFTG
SVSSLAWFMSNQETSFVTIGQRVLANPLKVRFHYGHPDVFDRIFHITRGGISKASKVINLSEDIFAGFNSTL
RQGNVTHHEYIQLGKGRDVGMNQISNFEAKVANGNGEQTLCRDVYRLGHRFDFYRMFSMYFTTVGFYF
NSLVAVLTVYVFLYGRLYLVLSGLEKSILQDPQIKNIKPFENALATQSVFQLGTLLILPMIMEVGLEKGFGKA
FAEFVMMQLQLAPMFFTFHLGTKTHYYGRTILHGGAKYRGTGRGFVVRHAKFAENYRMYSRSHFVKALE
LLILLVVYLAYGSSYRSSSLYIYVTISMWFLVFCWLFAPFVFNPSCFEWHKTVDDWTDWWKWMNNRGGI
GLAPEQSWESWWVSEHEHLKNGTIRSLLLELILSLRLLIYQYGIVYHLHIVHENKSFMIYALSWLVIAIILVSL
KVVSLGREKFVTKFQLVFRILKGIVFLVLVGLMVLLFVGFDLAVSDVGASILAFIPTGWFILLIPQLCGPLFRR
LVIEPLRVLCCSCCPGGACRGPCCAKFRQRSGAVLRKIGPWDSIREMARMYEYTMGILIFLPIAVLSWFPF
MSEFQTRLLFNQAFSRGLQISRILAGQDGRGAKQE
>AL3G05781.t1
MQQPAPRRILRTQTAVNLGEPSFDSEVVPSSLVEIAPILRVANEVEAANPRVAYLCRFYAFEKAHRLDPTS
SGRGVRQFKTALLQRLERENVPTLTGRAQKSDAREIQTFYQHYYKKYIQALQNASDQVDRGQLIKAYQTA
NVLFEVLKAVTQQHSVKVDDEILETADKVKEKTKIYLPFNILPLDPDSGNQAVMKLPEIQAAATALRNTRGL
PLPKNYERKTDNVSNQREHLILLLANIHIRKNPKTDEHSKLDDNALNDVMKKLFKNYKKWCKYLGRKSSLW
LPTIQQEVQQRKLLYMGLYLLIWGEAANLRFMPECVCYIYHHMAFEMYGMLAGNVSALTGEYVKPAYGG
EKEAFLKRKLKGAKRKGNHSQWRNYDDLNEYFWSADCFQLGWPMRADADFFCQPINPPDERNESTVRA
DKQKGKINFVELRSFWHIFRSFDRMWNFFIIALQIMVILAWSEGGSLGNIFDPLVFKRILSIFITSAILNLGQAT
LDIIFNWRARRTMEFAVKLRYVLKFVLAAVWVILLRLPVIYLLPNMLGAVLFAFPSLRRILEGSDYKFMRLIM
WWSQPRLFVGRGMHESPFSLFMYTMFWVVLLLTKFVFSYYVEIKPLVDPTKDIMKFPINHFQWHEFFPRA
KGNIGVVIALWIPIILVYFMDMQIWYTIFSTLFGGIYGAFQRLGEIRTLGMLRSRFDSIPLAVHDCLVPSETSD
ARRKKGLKSNLQKRFKEMKHEDKEKVAARFAQMWNEIVTSFRDEDLIDNREKELLLVPYVADKGLRVVQ
WPPFLLASMVPIAVDMAKDSNGKDRDLKKRIENDYYFSCAIEECYTSCKNIINDLVQGEQEKRVINDIFVEV
EKFIDEDKVITDLNMQSLPDLYDKFVELVEYLKRNDDKDRVAVIKIFQDMLEVVTRDIMEDQLPSILESSHG
GSYRRPEGMTTWDQEYQLFQPSGAIKFPLPVSDAWTEKVKRLELLLTVKESAMDVPSNLEARRRLTFFTN
SLFMNMPVAPKVRNTLSFSSNLVISLELEEENEDGVSTLFYLQKIYPDEWKNFQERVCWDEESKENEELK
EELRLWASYRGQTLTRTVRGMMYYRKALVLEAFLDMAKHEDLMEGYKAAESISDEEEQYGNDKRSALPN
AQDILQLMRTYPSLRVAYIDQVEDRVGEKKIETAYYSTLVKVALTKDSESADPVQNLDQVIYRIKLPGPALL

GEGKPENQNHAIIFTRGEGLQTIDMNQDNYMEEALKMRNLLQEFLTVHGVRNPSILGVREHIFTGSVSSLA
WFMSNQEHSFVTIGQRLLANPLKVRFHYGHPDVFDRLFHLTRGGVSKASRSINLSEDIFAGYNSTLRGGN
VTHHEYMQVGKGRDVGLNQISKFEAKVANGNGEQTLSRDIYRLGHRFDFFRMLSCYFTTVGFYFSTLLTV
VTVYVFLYGRMYLALSGLEAGLSTQRKFSHNHALQVALASQSLVQLGFLMALPMMMEIGLEKGFGQALSE
FIMMNLQLAAVFFTFSLGTKTHYYGRMLLHGGAQYRSTGRGFVVFHAKFAENYRLYSRSHFVKGIELMILL
IIYEIFGQSSHSTIAYIFVMFSMWFLVLTWLFAPFLFNPSGFEWAKILDDWSDWNKWISNSGGIGVSPEKS
WESWWEKEQEHLKHSGVLGNLFEIILSLRFFLYQYGLVYHLNITDDKSILVYLFSWLVILGALFIMKVISVGR
RRFGASFQLFFRLIKFMIFVAFAAVLVVLILILHMTIRDIFVCILAFLPTGWGILLIAQACRPVLRVAGLWGSVR
ALARAYEVIMGVLLFTPITVLSWFPFVSEFQTRMLFNQAFSRGLQISRILGGQKERAASTKD
>AL1G02931.t1
MSHLRNRRPLAGAAAAASAEQPPVQASYNIVPIQDVVMHGQHPSLRFPEVRAAVAALVHVADLPPPPFAR
AWDPRRADLFDWLGLTFGFQRHNVRNQREHLVLLLANAQLRAGGTLALPRDHPPDVLHFSIARDIRKKLL
RNYDAWCAYLGQRPHVHAPRRTGGGADDTRRELLYTALYLLIWGEAANLRFMPECLCYVFHYMALDLNH
VTERSVDIETGRPAIPAVCGEDAFLSRVVTPIYNVLDAEVKASQNGTKPHSAWRNYDDVNEYFWSRRVFK
RLQWPLDSSRGFFVPPGKFGRIGKTGFVEQRSFWNLYRSFDRLWVMLILFFQAAMIIAWDGRPYPWQSL
SFRDIQVRVLSVFITWAGLRFMQALLDAGTQYSLVSRETKLIAVRMVLKAFVAAGWTITFSVLYVRMWDQR
WRDRRWSFAANTRVVNYLEAAAVFIIPQVLALVLFILPWIRNFTEKTNWRILYVLTWWFQTRTFVGRGLRE
GLIDNIKYSIFWISLLAAKFSFSYFLQIKPMVSPTKTIFSLHDIRRNWFEFMPHTERLAVIILWVPVVLIYLMDI
QIWYSVFSSLTGALIGLFSHLGEIRSVGQLRLRFQFFASAMQFNLMPEERLDRFHGGIRSKLYDAIHRLKLR
YGFGRPYRKIEANEVEAKRFALIWNEIIQTFREEDIVSDNEVELLELPPVVWKIRVVRWPCLLLNNELLLALS
QVKELVADDKTHWSRISTIEYRRCAVIEAYDSIRQLLLEIIEERSAEHIIVNQLFLAFDSEIENGRFSEEYRLNL
LPQIHSSVITLVELLLKANKDQTKIVNTLQTLYVLAVHDFPKNRKDIEQLRRDGLAPSRPTESGLLFEDAITC
PGEEDLSFYKQVRRLHTILTSRDSMNDIPKNPEARRRIAFFSNSLFMNMPRAPTVEKMVAFSVLTPYYNED
VLYNKDQLRRENEDGISILFYLQKIYEDDWGNFLERMRREGMVNDDEIWSGKFQELRLWASYRGQTLAR
TVRGMMYYYRALKMLAFLDTASEIDITEGTKRLAAYGSVRHENDVYPMNNGLQQRPQRRLNRGASTVSQ
LFKGMEDGAALMKYTYVVACQIYGNQKKGKDQRAEDILALMKKNEALRVAYVDEVRHEMGDMQYYSVLV
KFDQDLQKEVEIYRIRLPGPLKLGEGKPENQNHAIIFTRGEAVQTIDMNQDNYFEEALKMRNLLEQYSYNH
GSQKPTLLGVREHVFTGSVSSLAWFMSAQETSFVTLGQRVLANPLKVRMHYGHPDVFDRLWFLTRGGL
SKASRVINISEDIFAGFNCTLRGGNVSHHEYIQVGKGRDVGLNQISMFEAKVSSGNGEQTLSRDIYRLGHR
VDFFRMLSVFYTTIGFYFNTMLVVLTVYTFTWGRLYLALSGLEAGIQGSANATNNKALGAVLNQQFVIQLGI
FTALPMILENSLELGFLPAVWDFFTMQMNFSSVFYTFSMGTKSHYYGRTILHGGAKYRATGRGFVVQHKS
FAENYRLYSRSHFIKAIELGIILTVYAVHSVIARNTLVYIVMMLSSWFLVVSWIMAPFAFNPSGFDWLKTVYD
FEDFMTWIWFPGGIFSKAEHSWEVWWYEEQDHLRTTGLWGKILEILLDLRYFFFQYGVVYQLKIADGSRSI
AVYLLSWICVAVIFGIFVLMSYTRDQYAAKQHLYYRVVQCAIIILGVLVLILFLEFTEFQIIDIFTGLLAFIPTGW
GLISIAQVIRPFIESTVVWGSVISVARLYEILLGVIVMAPVALLSWLPGFQEMQTRVLFNEGFSRGLQISRILA
GKKTNAV
>AL1G03358.t1
MTTPRATAPRRVGSAAAAPAAGEPYNILPIHDLLAEHPSLRFPEVRAAAAALRAVGGLRPPPYSHWRADQ
DIMDWLGAFFGFQRDNVRNQREHLVLLLANAQMRLSSADFSDTLEPRIARSLRKKLLRNYTSWCGFLGR
RPNVYVPDDDPRADLLFAGLHLLVWGEAANLRFMPECICYIYHHMALELHRILEGYIDTATGRPANPAVNG
ENAFLTRVITPVYGVIRAEVESSHNGTVPHAAWRNYDDINEYFWRRDVFDRLGWPMEQSRQFFRTPPDR

SRVRKTGFVEVRSFWNIYRSFDRLWVMLVLYMQAAAIIAWNGETWPWQNFHREAQVHLLTVFITWAALR
FLQSLLDIGTQFRRAFRDGRMLAVRMVLKAIVAAVWVLVFAILYKRILNQRDSDRGWSRGTDTRIMSFLYA
AAVFVLPEVLAILLFIIPWVRNALEKTNWKICYALTWWFQSRSFVGRGLREGTWDNVKYSLFWVLLLAVKF
TFSYFLQIRPLVKPTQEIYKLGKITYAWHEFFGQSNRFAVLILWLPVVLIYLMDIQIWYAIFSALSGAFVGLFA
HLGEIRDMKQLRLRFQFFASAMSFNIMPEEQHVNERTFLPSRLRNFWQRLQLRYGFSRSFRKIESNQVEA
RRFALIWNEIITKFREEDIVSDLEVELLELPPELWNVRVIRWPCFLLCNELSLALGQAKEVQGNDRRLWRKI
CKNDYRRCAIIEVYDSAKYMLLEIIKERSEEHGIVTQLFREFDESMHLEKFTVEYKMSVMQNVHAKLAALLS
LLLKPNKDLTKIVNALQTLYDVVVRDFQAEKRSMEQLRNEGLAQSRPTSLLFVDTVVLPNEQNATFYKQVR
RMHTILTSRDSMVNVPKNLEARRRIAFFSNSLFMNIPRATQVEKMMAFSVLTPYYNEEVLYNKDQLYKQN
EDGISILYYLKKIYPDEWEFFIERMKREGMSDEEELYSEKQRLRDLRHWVSYRGQTLSRTVRGMMYYYDA
LKMLTFLDSASEHDLRTGSRELATMGSSRFGSSRRDGGAGGSGPYSRASSSRTLSRATSGVSSLFKGSE
YGTVLMKYTYVVACQIYGQQKAKNDPHAYEILELMKNYEALRVAYVDEKHSNGGETEYFSVLVKYDQQLQ
QEVEIYRVKLPGPLKLGEGKPENQNHALIFTRGDAVQTIDMNQDNYFEEALKMRNLLEEFNRHYGIRKPKI
LGVREHVFTGSVSSLAWFMSAQETSFVTLGQRVLADPLKVRMHYGHPDVFDRLWFLGRGGISKASRVINI
SEDIFAGFNCTLRGGNVTHHEYIQVGKGRDVGLNQVSMFEAKVASGNGEQTLSRDVYRLGHRLDFFRML
SFFYTTIGFYFNTMMVVLTVYAFVWGRFYLALSGLEDYISKNTSSTNNAALGAVLNQQFVIQLGLFTALPMII
ENSLEHGFLNAVWDFLKMQLQFASVFYTFSMGTKTHYYGRTLLHGGAKYRATGRGFVVEHKKFAENYRL
YARSHFLKAIELGVILVLYASYSSAAGNTLVYILLTLSSWFLVSSWILAPFIFNPSGFDWLKNFNDFEDFLDW
IWFQGGISVKSDQSWEKWWEEETDHLRTSGLWGSILEVIIDLRYFFFQYAIVYRLHIAGGSRSILVYLLSWT
CILLAFVALVTVAYFRDRYSAKKHIRYRFVQAVIVGFTVTAIVLLIEFTQFQFIDALTSLLALLPTGWGIISIALV
FKPYLRRSETVWKTIVTVARLYDIMFGVIVMTPVAVLSWLPGLQEMQTRILFNEAFSRGLHINQMLTGAKR
HGV
>AL3G05825.t1
MQQPAPRRILRTQTAVNLGEPSFDSEVVPSSLVEIAPILRVANEVEAANPRVAYLCRFYAFEKAHRLDPTS
SGRGVRQFKTALLQRLERENVPTLTGRAQKSDAREIQTFYQHYYKKYIQALQNASDQVDRGQLIKAYQTA
NVLFEVLKAVTQQHSVKVDDEILETADKVKEKTKIYLPFNILPLDPDSGNQAVMKLPEIQAAATALRNTRGL
PLPKNYERKTDNVSNQREHLILLLANIHIRKNPKTDEHSKLDDNALNDVMKKLFKNYKKWCKYLGRKSSLW
LPTIQQEVQQRKLLYMAGNVSALTGEYVKPAYGGEKEAFLKKVVTPIYSTIAKEAERSKKEKGNHSQWRN
YDDLNEYFWSADCFQLGWPMRADADFFCQPINPPDERNESTVRADKQKGKINFVELRSFWHIFRSFDRM
WNFFIIALQIMVILAWSEGGSLGNIFDPLVFKRILSIFITSAILNLGQATLDIIFNWRARRTMEFAVKLRYVLKF
VLAAVWVILLPVTYAYLFNNPAGIIRAIKNWFGNGRNQPSLFAIAAVIYLLPNMLGAVLFAFPSLRRILEGSD
YKFMRLIMWWSQPRLFVGRGMHESPFSLFMYTMFWVVLLLTKFVFSYYVEIKPLVDPTKDIMKFPINHFQ
WHEFFPRAKGNIGVVIALWIPIILVYFMDMQIWYTIFSTLFGGIYGAFQRLGEIRTLGMLRSRFDSIPLAVHD
CLVPSETSDARRKKGLKSNLQKRFKEMKHEDKEKVAARFAQMWNEIVTSFRDEDLIDNREKELLLVPYVA
DKGLRVVQWPPFLLASMVPIAVDMAKDSNGKDRDLKKRIENDYYFSCAIEECYTSCKNIINDLVQGEQEKR
VINDIFVEVEKFIDEDKVITDLNMQSLPDLYDKFVELVEYLKRNDDKDRVAVIKIFQDMLEVVTRDIMEDQLP
SILESSHGGSYRRPEGMTTWDQEYQLFQPSGAIKFPLPVSDAWTEKVKRLELLLTVKESAMDVPSNLEAR
RRLTFFTNSLFMNMPVAPKVRNTLSFSALTPYYNEHVLFSKKELEEENEDGVSTLFYLQKIYPDEWKNFQE
RVCWDEESKENEELKEELRLWASYRGQTLTRTVRGMMYYRKALVLEAFLDMAKHEDLMEGYKAAESISD
EEWKSLLAQCEALADMKFTYVVSCQQYGNDKRSALPNAQDILQLMRTYPSLRVAYIDQVEDRVGEKKIET
AYYSTLVKVALTKDSESADPVQNLDQVIYRIKLPGPALLGEGKPENQNHAIIFTRGEGLQTIDMNQDNYME

EALKMRNLLQEFLTVHGVRNPSILGVREHIFTGSVSSLAWFMSNQEHSFVTIGQRLLANPLKVRFHYGHP
DVFDRLFHLTRGGVSKASRSINLSEDIFAGYNSTLRGGNVTHHEYMQVGKGRDVGLNQISKFEAKVANGN
GEQTLSRDIYRLGHRFDFFRMLSCYFTTVGFYFSTLLTVVTVYVFLYGRMYLALSGLEAGLSTQRKFSHNH
ALQYGLVYHLNITDDKSILVYLFSWLVILGALFIMKVISVGRRRFGASFQLFFRLIKFMIFVAFAAVLVVLILILH
MTIRDIFVCILAFLPTGWGILLIAQACRPVLRVAGLWGSVRALARAYEVIMGVLLFTPITVLSWFPFVSEFQT
RMLFNQAFSRGLQISRILGGQKERAASTKD
>AL5G00002.t1
ILLAYFQMAKELDGILDSSEAELMSRSCFELNWPPAEGSKFLRKPAKRKRTGKTNFVEHLTFLHLYRSFHR
LLLRRILMFGAYKTARGFAISRIVIRFIWLAAVSTFVTYLYVKVLDEENARNSDSTYFRIYVLVLGSYAALGSD
LCLLAKIPACHRLSDFSDRLQFFQFFKWIYQGRYYIGRGLYEYVIFWLVIFACKFTFTYFLQIRPLVVPTNIIV
QLHDLEYSWHDFVSRIYLMDIHIWYTLLSLLLVAMGARDRLGEIRSIEMLHKRFESFPEAFAKTLSPQRISN
RPVAQDSEATKTYASIFSPFWNEIIKSLREEDYVSNREMDLLMMPSNCGNLRLVQWPLFLLTSKIMLANDY
ASDCKDSQYELWHRISKDEYMAYAVKECYYSAERILNSLVDAEGQRLNAFFRDLNESIAQGSLLVTINLKK
LQLVQSRLTGLTGLLIRDETADRAAGVTKALRELYEVVTHEFLAPNLREQFDTWQLLLRARNDGRLFSNIF
WPKDLEMKEQVKRLHLLLTVKDSAANIPKNLEAQRRLQFFTNSLFMDMPEAKPVSEMIPFCVFTPYYSET
VLYSMSELCVDNEDGISILFYLQKIFPDEWANFLERIGRGESSEEDFKESSSDTLELRFWVSYRGQTLART
VRGMMYYRRALMLQSYLEKRYLGGIEDGYSAAEYIDTQGYQLSPDARAQADLKFTYVVSCQIYGQQKQR
KAPEAADIALLMQRNEALRVAFIHEEDSVSSDGHAIKEYHSKLVKADIHGKDQEIYSIKLPGNPKLGEGKPE
NQNHAIIFTRGDAVQTIDMNQDNYLEEAMKMRNLLEEFRSSHGIRYPTILGKREHVFTGSVSSLTPFMSKQ
ETSFVTLGQRVLAYLKLQCCKNSQDMFCFKHLKTEAISGDRLVPAHSVSPKWNGFNSTLRQGNITHHEYI
QMTVWTVYIFLYGRVYLALSDSTIQYLAARFLGNTALDAALNAQFLAIFSFITMQLQFCSVFFTFSLGTRTH
YFGRTILHGGAKYRATGRGFVVRHIKFADNYRLYSRSHFVKALEVALLLIVYIAYGYTKGGSSSFILLTISSW
FMVMSWLFAPYIFNPSGFEWQKTVEDFDDWTNWLFYKGGVGVRGENSWESWWDEEQAHIQTFRGRIL
ETILSLRFLLFQYGIVYKLKLTGDNTSLAIYGFSWIVLLVLVLLFKVGYINSFYFHLWHEYTLSEPVCLSLFYC
ASAFTATPKKSTAPPTFVRFLQGLLALEIIAAIACLIGFTDFTIADLFASALAFLATGWCILCLSITWKRLVKTL
GLWDSVREIARMYDRMGAIIFSPIVFFSWFLRRPSSR
>AT3G14570.1 GSL4
MSHEIVPVDPIDVPSTSYSRPILGPREDSPERATEFTRSLTFREHVSSEPFDSERLPATLASEIQRFLRIANL
VESEEPRIAYLCRFHAFEIAHHMDRNSTGRGVRQFKTSLLQRLELDEEFTVRRRKEKSDVRELKRVYHAY
KEYIIRHGAAFNLDNSQREKLINARRIASVLYEVLKTVTSGAGPQAIADRESIRAKSEFYVPYNILPLDKGGV
HQAIMHLPEIKAAVAIVRNTRGLPPPEEFQRHQPFLDLFEFLQYAFGFQNGNVANQREHLILLLSNTIIRQP
QKQSSAPKSGDEAVDALMKKFFKNYTNWCKFLGRKNNIRLPYVKQEALQYKTLYIGLYLLIWGEASNLRF
MPECLCYIFHHMAYELHGVLTGAVSMITGEKVAPAYGGGHESFLADVVTPIYMVVQKEAEKNKNGTADHS
MWRNYDDLNEFFWSLECFEIGWPMRPEHDFFCVESSETSKPGRWRGMLRFRKQTKKTDEEIEDDEELG
VLSEEQPKPTSRWLGKTNFVETRSFWQIFRSFDRMWSFFVLSLQALIIMACHDVGSPLQVFNANIFEDVM
SIFITSAILKLIKGILDIIFKWKARNTMPINEKKKRLVKLGFAAMWTIILPVLYSHSRRKYICYFTNYKTWLGEW
CFSPYMVAVTIYLTGSAIELVLFFVPAISKYIETSNHGIFKTLSWWGQPRLYVGRGMQETQVSQFKYTFFWI
LVLLTKFAFSYAFEIKPLIEPTRLIMKVGVRNYEWHEIFPEVKSNAAAIVAVWAPIMVVYFMDTQIWYSVYCT
IFGGLYGVLHHLGEIRTLGMLRGRFHTLPSAFNASLIPHSTKDEKRRKQRGFFPFNLGRGSDGQKNSMAK
FVLVWNQVINSFRTEDLISNKELDLMTMPLSSEVLSGIIRWPIFLLANKFSTALSIAKDFVGKDEVLYRRIRK
DEYMYYAVKECYESLKYILQILVVGDLEKKIISGIINEIEESIRQSSLLEEFKMAELPALHDKCIELVQLLVEGS

AEQLQVEKSEELHGKLVKALQDIFELVTNDMMVHGDRILDLLQSREGSGEDTGIFMRVIEPQLFESYGEW
RCIHFPLPDSASLSEQIQRFLLLLTVKDSAMDIPENLDARRRLSFFATSLFMDMPDAPKVRNMMSFSVLTP
HYQEDINYSTNELHSTKSSVSIIFYMQKIFPDEWKNFLERMGCDNLDALKKEGKEEELRNWASFRGQTLS
RTVRGMMYCREALKLQAFLDMADDEDILEGYKDVERSNRPLAAQLDALADMKFTYVVSCQMFGAQKSS
GDPHAQDILDLMIKYPSLRVAYVEEREEIVLDVPKKVYYSILVKAVNGFDQEIYRVKLPGPPNIGEGKPENQ
NHAIVFTRGEALQTIDMNQDHYLEEAFKMRNLLQEFLRNRGRRPPTILGLREHIFTGSVSSLAWFMSYQET
SFVTIGQRLLANPLRVRFHYGHPDVFDRIFHITRGGISKSSRTINLSEDVFAGYNTTLRRGCITYNEYLQVG
KGRDVGLNQISKFEAKVANGNSEQTISRDIYRLGQRFDFFRMLSCYFTTIGFYFSSLISVIGIYIYLYGQLYLV
LSGLQKTLILEAKVKNIKSLETALASQSFIQLGLLTGLPMVMEIGLEKGFLIAFQDFILMQLQLAAFFFTFSLG
TKTHYFGRTILHGGAKYRPTGRKVVVFHANFSENYRLYSRSHFIKGFELMILLVVYELFKHTSQSNMAYSFI
TFSVWFMSFTWLCAPFLFNPSGFTWEIIVGDWRDWNRWIKEQGGIGIQQDKSWQSWWNDEQAHLRGS
GVGARCLEIILSLRFFVYQYGLVYHLDITQSNTNIIVYALSWVVILATFFTVKAVDLGRQLFSTRKHLVFRFFK
VFVFVSILTIIITLANICHLSVKDLLVSCLAFLPTGWGLILIAQAVRPKIEGTSLWEFTQVLARAYDYGMGVVLF
APMAILAWLPIISAFQTRFLFNEAFNRRLQIQPILAGKKKNR
>AT1G06490.1 GSL7
MASTSSGGRGEDGRPPQMQPVRSMSRKMTRAGTMMIEHPNEDERPIDSELVPSSLASIAPILRVANDIDQ
DNARVAYLCRFHAFEKAHRMDPTSSGRGVRQFKTYLLHKLEEEEEITEHMLAKSDPREIQLYYQTFYENNI
QDGEGKKTPEEMAKLYQIATVLYDVLKTVVPQARIDDKTLRYAKEVERKKEQYEHYNILPLYALGAKTAVM
ELPEIKAAILAVCNVDNLPRPRFHSASANLDEVDRERGRSFNDILEWLALVFGFQRGNVANQREHLILLLAN
IDVRKRDLENYVEIKPSTVRKLMEKYFKNYNSWCKYLRCDSYLRFPAGCDKQQLSLLYIGLYLLIWGEASN
VRFMPECLCYIFHNMANEVHGILFGNVYPVTGDTYEAGAPDEEAFLRNVITPIYQVLRKEVRRNKNGKASH
SKWRNYDDLNEYFWDKRCFRLKWPMNFKADFFIHTDEISQVPNQRHDQVSHGKRKPKTNFVEARTFWN
LYRSFDRMWMFLVLSLQTMIIVAWHPSGSILAIFTEDVFRNVLTIFITSAFLNLLQATLDLVLSFGAWKSLKF
SQIMRYITKFLMAAMWAIMLPITYSKSVQNPTGLIKFFSSWVGSWLHRSLYDYAIALYVLPNILAAVFFLLPP
LRRIMERSNMRIVTLIMWWAQPKLYIGRGMHEEMFALFKYTFFWVMLLLSKLAFSYYVEILPLVNPTKLIWD
MHVVNYEWHEFFPNATHNIGVIIAIWGPIVLVYFMDTQIWYAIFSTLFGGIYGAFSHLGEIRTLGMLRSRFKV
VPSAFCSKLTPLPLGHAKRKHLDETVDEKDIARFSQMWNKFIHTMRDEDLISDRERDLLLVPSSSGDVTVV
QWPPFLLASKIPIALDMAKDFKGKEDVDLFKKIKSEYYMHYAVVEAYETVRDIIYGLLQDESDKRIVREICYE
VDISIQQHRFLSEFRMTGMPLLSDKLEKFLKILLSDYEEDDYKSQIINVLQDIIEIITQDVMVNGHEILERAHLQ
SGDIESDKKEQRFEKIDLSLTQNISWREKVVRLLLLLTVKESAINIPQSLEARRRMTFFANSLFMNMPDAPR
VRDMLSFSVLTPYYKEDVLYSEEELNKENEDGITILFYLQRIYPEEWSNYCERVNDLKRNLSEKDKAEQLR
QWVSYRGQTLSRTVRGMMYYRVALELQCFQEYTEENATNGGYLPSESNEDDRKAFSDRARALADLKFT
YVVSCQVYGNQKKSSESRDRSCYNNILQLMLKYPSLRVAYIDEREETVNGKSQKVFYSVLLKGCDKLDEE
IYRIKLPGPPTEIGEGKPENQNHAIIFTRGEALQTIDMNQDNYFEECFKMRNVLQEFDEGRRGKRNPTILGL
REHIFTGSVSSLAWFMSNQETSFVTIGQRVLANPLRVRFHYGHPDIFDRIFHITRGGISKASKIINLSEDIFAG
YNSTLRGGYVTHHEYIQAGKGRDVGMNQISFFEAKVANGNGEQTLSRDVYRLGRRFDFYRMLSFYFTTV
GFYFSSMITVLTVYVFLYGRLYLVLSGLEKNILQSASVHESNALEQALAAQSVFQLGFLMVLPMVMEIGLEK
GFRTALGDFIIMQLQLASVFFTFQLGTKAHYFGRTILHGGSKYRATGRGFVVFHAKFAENYRLYSRSHFVK
GLELVILLVVYQVYGTSYRSSSTYMYITFSMWFLVTSWLFAPFIFNPSGFEWQKTVDDWTDWKRWMGNR
GGIGIVLDKSWESWWDIEQEHLKHTNLRGRVLEILLALRFLLYQYGIVYHLNIARRHTTFLVYGLSWAILLSV
LLVLKMVSMGRRKFGTDFQVMFRILKALLFLGFLSVMTVLFVVCGLTISDLFASILAFLPTGWAILLIGQALR

SVFKGLGFWDSVKELGRAYEYIMGLVIFTPIAVLSWFPFVSEFQTRLLFNQAFSRGLQISMILAGKKDKETP
STKYLGHTEESFGLEHDTNTFNHYYLWT
>AT5G13000.1 GSL12
MSATRGGPDQGPSQPQQRRIIRTQTAGNLGESFDSEVVPSSLVEIAPILRVANEVESSNPRVAYLCRFYAF
EKAHRLDPTSSGRGVRQFKTALLQRLEREHDPTLMGRVKKSDAREMQSFYQHYYKKYIQALHNAADKAD
RAQLTKAYQTANVLFEVLKAVNLTQSIEVDREILEAQDKVAEKTQLYVPYNILPLDPDSANQAIMRYPEIQA
AVLALRNTRGLPWPEGHKKKKDEDMLDWLQEMFGFQKDNVANQREHLILLLANVHIRQFPKPDQQPKLD
DQALTEVMKKLFKNYKKWCKYLGRKSSLWLPTIQQEMQQRKLLYMALYLLIWGEAANLRFMPECLCYIYH
HMAFELYGMLAGNVSPMTGENVKPAYGGEEDAFLRKVVTPIYEVIQMEAQRSKKGKSKHSQWRNYDDL
NEYFWSVDCFRLGWPMRADADFFCLPVAVPNTEKDGDNSKPIVARDRWVGKVNFVEIRSFWHVFRSFD
RMWSFYILCLQAMIIMAWDGGQPSSVFGADVFKKVLSVFITAAIMKLGQAVLDVILNFKAHQSMTLHVKLR
YILKVFSAAAWVIILPVTYAYSWKDPPAFARTIKSWFGSAMHSPSLFIIAVVSYLSPNMLAGVMFLFPLLRRF
LERSNYRIVMLMMWWSQPRLYVGRGMHESAFSLFKYTMFWVLLIATKLAFSYYIEIRPLVAPTQAIMKARV
TNFQWHEFFPRAKNNIGVVIALWAPIILVYFMDSQIWYAIFSTLFGGIYGAFRRLGEIRTLGMLRSRFESLPG
AFNDRLIPDGKNQQKKKGIRATLSHNFTEDKVPVNKEKEAARFAQLWNTIISSFREEDLISDREMDLLLVPY
WADRDLDLIQWPPFLLASKIPIALDMAKDSNGKDRELKKRIESDTYMKCAVRECYASFKNIIKFVVQGNRE
KEVIEIIFAEVDKHIDTGDLIQEYKMSALPSLYDHFVKLIKYLLDNKEEDRDHVVILFQDMLEVVTRDIMMEDY
NISSLVDSSHGGTWHGGMIPLEQQYQLFASSGAIRFPIEPVTEAWKEKIKRIYLLLTTKESAMDVPSNLEAR
RRISFFSNSLFMDMPMAPKVRNMLSFSVLTPYYTEEVLFSLRDLETPNEDGVSILFYLQKIFPDEWNNFLE
RVKCLSEEELKESDELEEELRLWASYRGQTLTRTVRGMMYYRKALELQAFLDMAMHEDLMEGYKAVELN
SENNSRGERSLWAQCQAVADMKFTYVVSCQQYGIHKRSGDPRAQDILRLMTRYPSLRVAYIDEVEEPVK
DKSKKGNQKVYYSVLVKVPKSTDHSTLAQNLDQVIYRIRLPGPAILGEGKPENQNHAIIFSRGEGLQTIDMN
QDNYMEEALKMRNLLQEFLTKHDGVRHPSILGLREHIFTGSVSSLAWFMSNQETSFVTIGQRLLANPLRV
RFHYGHPDVFDRLFHLTRGGVSKASKVINLSEDIFAGFNSTLREGNVTHHEYIQVGKGRDVGLNQISMFEA
KIANGNGEQTLSRDIYRLGHRFDFFRMMSCYFTTVGFYFSTLITVLTVYIFLYGRLYLVLSGLEQGLSTQKGI
RDNTPLQIALASQSFVQIGFLMALPMLMEIGLERGFRTALSEFVLMQLQLAPVFFTFSLGTKTHYYGRTLLH
GGAKYRSTGRGFVVFHAKFADNYRLYSRSHFVKGLEMMLLLVVYQIFGSAYRGVLAYLLITISMWFMVGT
WLFAPFLFNPSGFEWQKIVDDWTDWNKWINNIGGIGVPAEKSWESWWEEEQEHLRYSGKRGIVVEILLA
LRFFIYQYGLVYHLTITEKTKNFLVYGVSWLVIFLILFVMKTVSVGRRRFSASFQLMFRLIKGLIFMTFIAIIVILI
TLAHMTIQDIIVCILAFMPTGWGMLLIAQACKPVVHRAGFWGSVRTLARGYEIVMGLLLFTPVAFLAWFPFV
SEFQTRMLFNQAFSRGLQISRILGGHRKDRSSRNKE
>AT2G31960.1 GSL3
MAQRKGPDPPPPQRRILRTQTAGNLGEAMLDSEVVPSSLVEIAPILRVANEVEASNPRVAYLCRFYAFEKA
HRLDPTSSGRGVRQFKTALLQRLERENETTLAGRQKSDAREMQSFYQHYYKKYIQALQNAADKADRAQL
TKAYQTAAVLFEVLKAVNQTEDVEVADEILEAHTKVEEKSQIYVPYNILPLDPDSQNQAIMRFPEIQATVSAL
RNTRGLPWPAGHKKKLDEDMLDWLQTMFGFQKDNVSNQREHLILLLANVHIRQFPRPEQQPRLDDRALT
IVMKKLFKNYKKWCKYLGRKSSLWLPTIQQEVQQRKLLYMGLYLLIWGEAANLRFLPECLCYIYHHMAFEL
YGMLAGSVSPMTGEHVKPAYGGEDEAFLQKVVTPIYKTIAKEAKRSRGGKSKHSEWRNYDDLNEYFWSI
RCFRLGWPMRADADFFCQTAEELRLDRSENKPKTGDRWMGKVNFVEIRSFWHIFRSFDRMWSFYILSLQ
AMIIIAWNGSGKLSGIFQGDVFLKVLSIFITAAILKLAQAVLDIALSWKSRHSMSFHVKLRFIFKAVAAAIWVVL
MPLTYAYSWKTPSGFAETIKNWFGGHQNSSPSFFIIVILIYLSPNMLSTLLFAFPFIRRYLERSDYKIVMLMM

WWSQPRLYIGRGMHESALSLFKYTMFWVVLLISKLAFSFYAEIKPLVKPTKDIMRVHISVYRWHEFFPHAK
SNMGVVIALWSPVILVYFMDTQIWYAIVSTLVGGLNGAFRRLGEIRTLGMLRSRFQSLPEAFNACLVPNEK
SETPKKKGIMATFTRKFDQVPSSKDKEAARFAQMWNKIISSFREEDLISDREMELLLVPYWADRDLDLIRW
PPFLLASKIPIALDMAKDSNGKDRELTKRLSVDSYMTCAVRECYASFKNLINFLVVGEREGQVINEIFSRIDE
HIEKETLIKDLNLSALPDLYGQFVRLIEYLMENREEDKDQIVIVLLNMLEVVTRDIMDEEVPSMLESTHNGTY
VKYDVMTPLHQQRKYFSQLRFPVYSQTEAWKEKIKRLHLLLTVKESAMDVPSNLEARRRLTFFSNSLFME
MPDAPKIRNMLSFSVLTPYYSEDVLFSIFGLEKQNEDGVSILFYLQKIFPDEWTNFLERVKCGSEEELRARE
ELEEELRLWASYRGQTLTKTVRGMMYYRKALELQAFLDMAKDEELMKGYKALELTSEDASKSGTSLWAQ
CQALADMKFTFVVSCQQYSVQKRSGDQRAKDILRLMTTYPSLRVAYIDEVEQTHKESYKGADEKIYYSAL
VKAAPQTKSMDSSESVQTLDQVIYRIKLPGPAILGEGKPENQNHSIIFTRGEGLQTIDMNQDNYMEEAFKM
RNLLQEFLVKHGGVRTPTILGLREHIFTGSVSSLAWFMSNQENSFVTIGQRVLASPLKVRFHYGHPDVFDR
LFHLTRGGVCKASKVINLSEDIFAGFNSTLREGNVTHHEYIQVGKGRDVGLNQISMFEAKIANGNGEQTLS
RDLYRLGHRFDFFRMLSCYFTTIGFYFSTMLTVLTVYVFLYGRLYLVLSGLEEGLSNQKAFRSNMPLQAAL
ASQSFVQIGFLMALPMMMEIGLERGFHNALIDFVLMQLQLASVFFTFQLGTKTHYYGRTLFHGGAEYRGT
GRGFVVFHAKFAENYRFYSRSHFVKGIELMILLLVYQIFGHAYRGVVTYILITVSIWFMVVTWLFAPFLFNPS
GFEWQKIVDDWTDWNKWIYNRGGIGVPPEKSWESWWEKEIGHLRHSGKRGIILEIVLALRFFIFQYGLVY
QLSTFKQENQSLWIYGASWFVILFILLIVKGLGVGRQRFSTNFQLLFRIIKGFVFLTFLGLLITFLALRFLTPKD
IFLCMLAFMPTGWGMLLIAQACKPLIQRLGFWSSVRTLARGYEILMGLLLFTPVAFLAWFPFVSEFQTRML
FNQAFSRGLQISRILGGQRKDRSSKNKE
>AT1G05570.1 GSL6
MAQRREPDPPPPQRRILRTQTVGSLGEAMLDSEVVPSSLVEIAPILRVANEVEASNPRVAYLCRFYAFEKA
HRLDPTSSGRGVRQFKTALLQRLERENETTLAGRQKSDAREMQSFYQHYYKKYIQALLNAADKADRAQL
TKAYQTAAVLFEVLKAVNQTEDVEVADEILETHNKVEEKTQIYVPYNILPLDPDSQNQAIMRLPEIQAAVAAL
RNTRGLPWTAGHKKKLDEDILDWLQSMFGFQKDNVLNQREHLILLLANVHIRQFPKPDQQPKLDDRALTIV
MKKLFRNYKKWCKYLGRKSSLWLPTIQQEVQQRKLLYMGLYLLIWGEAANLRFMPECLCYIYHHMAFELY
GMLAGSVSPMTGEHVKPAYGGEDEAFLQKVVTPIYQTISKEAKRSRGGKSKHSVWRNYDDLNEYFWSIR
CFRLGWPMRADADFFCQTAEELRLERSEIKSNSGDRWMGKVNFVEIRSFWHIFRSFDRLWSFYILCLQA
MIVIAWNGSGELSAIFQGDVFLKVLSVFITAAILKLAQAVLDIALSWKARHSMSLYVKLRYVMKVGAAAVWV
VVMAVTYAYSWKNASGFSQTIKNWFGGHSHNSPSLFIVAILIYLSPNMLSALLFLFPFIRRYLERSDYKIMM
LMMWWSQPRLYIGRGMHESALSLFKYTMFWIVLLISKLAFSYYAEIKPLVGPTKDIMRIHISVYSWHEFFPH
AKNNLGVVIALWSPVILVYFMDTQIWYAIVSTLVGGLNGAFRRLGEIRTLGMLRSRFQSIPGAFNDCLVPQD
NSDDTKKKRFRATFSRKFDQLPSSKDKEAARFAQMWNKIISSFREEDLISDREMELLLVPYWSDPDLDLIR
WPPFLLASKIPIALDMAKDSNGKDRELKKRLAVDSYMTCAVRECYASFKNLINYLVVGEREGQVINDIFSKI
DEHIEKETLITELNLSALPDLYGQFVRLIEYLLENREEDKDQIVIVLLNMLELVTRDIMEEEVPSLLETAHNGS
YVKYDVMTPLHQQRKYFSQLRFPVYSQTEAWKEKIKRLHLLLTVKESAMDVPSNLEARRRLTFFSNSLFM
DMPPAPKIRNMLSFSVLTPYFSEDVLFSIFGLEQQNEDGVSILFYLQKIFPDEWTNFLERVKCGNEEELRA
REDLEEELRLWASYRGQTLTKTVRGMMYYRKALELQAFLDMAKDEELLKGYKALELTSEEASKSGGSLW
AQCQALADMKFTFVVSCQQYSIHKRSGDQRAKDILRLMTTYPSIRVAYIDEVEQTHKESYKGTEEKIYYSA
LVKAAPQTKPMDSSESVQTLDQLIYRIKLPGPAILGEGKPENQNHAIIFTRGEGLQTIDMNQDNYMEEAFK
MRNLLQEFLEKHGGVRCPTILGLREHIFTGSVSSLAWFMSNQENSFVTIGQRVLASPLKVRFHYGHPDIFD
RLFHLTRGGICKASKVINLSEDIFAGFNSTLREGNVTHHEYIQVGKGRDVGLNQISMFEAKIANGNGEQTLS

RDLYRLGHRFDFFRMLSCYFTTIGFYFSTMLTVLTVYVFLYGRLYLVLSGLEEGLSSQRAFRNNKPLEAAL
ASQSFVQIGFLMALPMMMEIGLERGFHNALIEFVLMQLQLASVFFTFQLGTKTHYYGRTLFHGGAEYRGT
GRGFVVFHAKFAENYRFYSRSHFVKGIELMILLLVYQIFGQSYRGVVTYILITVSIWFMVVTWLFAPFLFNPS
GFEWQKIVDDWTDWNKWIYNRGGIGVPPEKSWESWWEKELEHLRHSGVRGITLEIFLALRFFIFQYGLVY
HLSTFKGKNQSFWVYGASWFVILFILLIVKGLGVGRRRFSTNFQLLFRIIKGLVFLTFVAILITFLALPLITIKDL
FICMLAFMPTGWGMLLIAQACKPLIQQLGIWSSVRTLARGYEIVMGLLLFTPVAFLAWFPFVSEFQTRMLF
NQAFSRGLQISRILGGQRKDRSSKNKE
>AT2G13680.1 GSL2
MAQSSTSHDSGPQGLMRRPSRSAATTVSIEVFDHEVVPASLGTIAPILRVAAEIEHERPRVAYLCRFYAFE
KAHRLDPSSGGRGVRQFKTLLFQRLERDNASSLASRVKKTDGREVESFYQQYYEHYVRALDQGDQADR
AQLGKAYQTAGVLFEVLMAVNKSEKVEAVAPEIIAAARDVQEKNEIYAPYNILPLDSAGASQSVMQLEEVK
AAVAALGNTRGLNWPSGFEQHRKKTGNLDLLDWLRAMFGFQRDNVRNQREHLVCLFADNHIRLTPKPEP
LNKLDDRAVDTVMSKLFKNYKNWCKFLGRKHSLRLPQAAQDIQQRKILYMGLYLLIWGEAANIRFMPECL
CYIFHNMAYELHGLLAGNVSIVTGENIKPSYGGDDEAFLRKVITPIYRVVQTEANKNANGKAAHSDWSNYD
DLNEYFWTPDCFSLGWPMRDDGDLFKSTRDTTQGKKGSFRKAGRTGKSNFTETRTFWHIYHSFDRLWT
FYLLALQAMIILAFERVELREILRKDVLYALSSIFITAAFLRFLQSVLDVILNFPGFHRWKFTDVLRNILKIVVSL
AWCVVLPLCYAQSVSFAPGKLKQWLSFLPQVKGVPPLYIMAVALYLLPNVLAAIMFIFPMLRRWIENSDWH
IFRLLLWWSQPRIYVGRGMHESQIALIKYTIFWLLLFCCKFAFSYFLQVKLLVKPTNAIMSIRHVKYKWHEFF
PNAEHNYGAVVSLWLPVILVYFMDTQIWYAIFSTICGGVIGAFDRLGEIRTLGMLRSRFQSLPGAFNTYLVP
SDKTRRRGFSLSKRFAEVTAARRTEAAKFSQLWNEIISSFREEDLISDREMDLLLVPYTSDPSLKLIQWPPF
LLASKIPIALDMAAQFRTRDSDLWKRICADEYMKCAVIECYESFKHVLHTLVIGENEKRIIGIIIKEVESNISKN
SFLSNFRMAPLPALCSKFVELVGILKNADPAKRDTVVLLLQDMLEVVTRDMMQNENRELVELGHTNKESG
RQLFAGTDAKPAILFPPVATAQWHEQISRLHLLLTVKESAMDVPTNLEAQRRIAFFTNSLFMDMPRAPRVR
NMLSFSVLTPYYSEETVYSKNDLEMENEDGVSVVYYLQKIFPDEWTNFLERLDCKDETSVLESEENILQLR
HWVSLRGQTLFRTVRGMMYYRRALKLQAFLDMANETEILAGYKAISEPTEEDKKSQRSLYTQLEAVADLK
FTYVATCQNYGNQKRSGDRRATDILNLMVNNPSLRVAYIDEVEEREGGKVQKVFYSVLIKAVDNLDQEIYR
IKLPGPAKIGEGKPENQNHALIFTRGEALQAIDMNQDHYLEEALKMRNLLEEFNEDHGVRAPTILGFREHIF
TGSVSSLAWFMSNQETSFVTIGQRVLASPLKVRFHYGHPDVFDRIFHITRGGISKASRGINLSEDIFAGFNS
TLRRGNVTHHEYIQVGKGRDVGLNQISLFEAKVACGNGEQTLSRDLYRLGHRFDFFRMMSCYFTTVGFYI
SSMIVVLTVYAFLYGRLYLSLSGVEEAIVKFAAAKGDSSLKAAMASQSVVQLGLLMTLPMVMEIGLERGFR
TALSDLIIMQLQLAPVFFTFSLGTKVHYYGRTILHGGSKYRATGRGFVVKHEKFAENYRMYSRSHFVKGME
LMVLLICYRIYGKAAEDSVGYALVMGSTWFLVGSWLFAPFFFNPSGFEWQKIVDDWDDWNKWISSRGGI
GVPANKSWESWWEEEQEHLLHSGFFGKFWEIFLSLRYFIYQYGIVYQLNLTKESRMGKQHSIIVYGLSWL
VIVAVMIVLKIVSMGRKKFSADFQLMFRLLKLFLFIGSVVIVGMLFHFLKLTVGDIMQSLLAFLPTGWALLQIS
QVARPLMKTVGMWGSVKALARGYEYIMGVVIFMPVTVLAWFPFVSEFQTRLLFNQAFSRGLQIQRILAGG
KKQK
>AT3G59100.1 GSL11
MEASSSGTAELPRSLSRRAPSRATTMMIDRPNEDASAMDSELVPSSLASIAPILRVANEIEKDNPRVAYLC
RFHAFEKAHRMDATSSGRGVRQFKTYLLHRLEKEEEETKPQLAKNDPREIQAYYQNFYEKYIKEGETSRK
PEEMARLYQIASVLYDVLKTVVPSPKVDYETRRYAEEVERKRDRYEHYNILPLYAVGTKPAIVELPEVKAAF
SAVRNVRNLPRRRIHLPSNTPNEMRKARTKLNDILEWLASEFGFQRGNVANQREHIILLLANADIRKRNDE

EYDELKPSTVTELMDKTFKSYYSWCKYLHSTSNLKFPDDCDKQQLQLIYISLYLLIWGEASNVRFMPECIC
YIFHNMANDVYGILFSNVEAVSGETYETEEVIDEESFLRTVITPIYQVIRNEAKRNKGGTASHSQWRNYDDL
NEYFWSKKCFKIGWPLDLKADFFLNSDEITPQDERLNQVTYGKSKPKTNFVEVRTFWNLFRDFDRMWIFL
VMAFQAMVIVGWHGSGSLGDIFDKDVFKTVLTIFITSAYLTLLQAALDIILNFNAWKNFKFSQILRYLLKFAVA
FMWAVLLPIAYSKSVQRPTGVVKFFSTWTGDWKDQSFYTYAVSFYVLPNILAALLFLVPPFRRAMECSDM
RPIKVIMWWAQPKLYVGRGMHEDMFSLFKYTTFWIMLLISKLAFNYYVEILPLITPTKMIMNLHIGHYQWHE
FFPHATNNIGVVIAIWAPIVLVYLMDTQIWYAIFSTLFGGIHGAFSHLGEIRTLGMLRSRFESIPIAFSRTLMP
SEDAKRKHADDYVDQKNITNFSQVWNEFIYSMRSEDKISDRDRDLLLVPSSSGDVSVIQWPPFLLASKIPIA
VDMAKDFKGKEDAELFRKIKSDSYMYYAVIESYETLKKIIYALLEDEADRRVMNQVFLEVDMSMQQQRFIY
EFRMSGLPLLSDKLEKFLSILLSDYEDQGTYKSQLINVFQDVIEIITQDLLVNGHEILERARVHSPDIKNEKKE
QRFEKINIHLVRDRCWREKVIRLHLLLSVKESAINVPQNLEARRRITFFANSLFMNMPSAPRIRDMLSFSVL
TPYYKEDVLYSEEDLNKENEDGISILFYLQKIYPDEWTNYLDRLKDPKLPEKDKSEFLREWVSYRGQTLAR
TVRGMMYYRQALELQCYQEVAGEQAEFSVFRAMASNDENQKAFLERARALADLKFTYVVSCQVYGNQK
KSGDIHNRSCYTNILQLMLKYPSLRVAYVDEREETADAKSPKVFYSVLLKGGDKFDEEIYRIKLPGPPAEIG
EGKPENQNHAIIFTRGEALQTIDMNQDNYFEEAFKLRNVLEEFNKERVGRRKPTILGLREHIFTGSVSSLA
WFMSNQESSFVTIGQRILANPLRVRFHYGHPDIFDRIFHITRGGVSKASKVINLSEDIFGGFNSTLRGGYVT
HHEYIQVGKGRDVGLNPISIFEAKVANGNGEQTLSRDVYRLGHRFDFYRMLSFYFTTIGFYFSSMLTVLTV
YAFLYGRMYMVMSGLEKEILRLASPNQLEALEQALATQSIFQLGFLMVLPMVMEIGLEHGFRSAIVDFFIM
QLQLASVFFTFQLGTKSHYYGRTILHGGSKYRPTGRGFVVFHAKFAENYRLYSRSHFVKGLELLLLLVVYQ
IYGHSYRSSNLYLYITVSMWFMVGSWLFAPFIFNPSGFEWQKTVDDWTDWKRWLGDRGGIGIPVEKSWE
SWWNVEQEHLKHTSIRGRILEITLALRFFIYQYGIVYQLNISQRSKSFLVYGLSWVVLLTSLLVLKMVSMGR
RRFGTDFQLMFRILKALLFLGFLSVMTILFVVFKLTLTDLSASVLAFLPTGWAILLIGQVLRSPIKALGVWDSV
KELGRAYENIMGLVIFAPIAVLSWFPIVSEFQARLLFNQAFSRGLQISMILAGRKDKATSSHK
>LOC_Os06g08380.1
MTGPRGGGGEDPPRTSSGAAASASASSGVPNEPLTPTSMLMSGQQAGLNRRGSRSAAMATFSMEVFD
NEVVPSTLSSIAPILRVAAEIEPERPRVAYLCRFYAFEKAHRLDQNSVGRGVRQFKTALLQRLEKDNSPSL
AKRVKKTDAREIESFYQQYYENYVRALDKGEQADRAQLGKAYQTAGVLFEVLCAVNKNEKVEEVNPEIVR
LHRDVQEKKDIYTPFNILPLDAASASQSIMQMEEIKAAVAALRNTRGLTWPSTFEPERQKGGDLDLLDWLR
AMFGFQRDSVRNQREHLILLLANVHIRLEPKPEPLSKLDDRAVDEVMAKLFSNYRKWCNFLSRKHSLRSP
QGAQPQEIQQRNILFLGLYLLIWGEAANIRFMPECLCYIFHNMAYELNGLLAGNVSIVTGENIRPSYGGDEE
AFLKKVVTPIYRVIKKESGKSKHGKTPHSAWCNYDDLNEYFWTTDCFSLGWPMRDDGDFFKSVHDSRPV
TTAGSSSQKGSTKSTGKMNFVETRTFWHIFRSFDRMWTFYLLALQAMLIFAWSDYTLSQILQKDLLYSLSS
IFVTAAFLQFLQSILDFVLNFPGHHKCKFLDAMRNILKIIASAAWAVILPFFYISTASKVNLPIKDLDKWFQYV
KGVPPLYILAVAVYLIPNILSAALFLLPCFRRWIENSDWRIVRLLLWWSQKRIYVGRGMHESSVSLFKYTLF
WILLLCSKFAFSYFVQIKPLIKPTKDIMNVHNIHYEWHEFFPNASYNVGAVMSLWAPVLLVYLMDTQIWYAI
FSTISGGVSGALGRLGEVSPSKRTEAAKFAQLWNEVICSFREEDLISDKEMDLLVVPYSSDPSLKLMQWP
LFLLASKIPIALDMAAQFRPRDSDLWKRICADEYMKCAVLECYESFKLVLNLLVIGENEKRIIGIIIKEIEANIAK
NTFLANFRMSALPVLCKKFVELVSALKERDASKFDNVVLLLQDMLEVITRDMMVNEIRELAEFGHGNKDSV
PRRQLFAGTGTKPAIVFPPPISAQWDEQIKRLYLLLTVKESAMDVPTNLEARRRIAFFTNSLFMDMPRAPR
VRKMLSFSVMTPYYSEETVYSRNDLDLENEDGVSIIFYLQKIFPDEWNNFLERIGCQRESEVWGNEENVL
QLRHWASLRGQTLCRTVRGMMYYKRALKLQAFLDMASESEILEGYKAVADPAEEEKKSQRSLSSQLEAIA

DMKFTYVATCQIYGNQKQSGDRRATDILNLMVNYPGLRVAYIDEVEERDGEKVQKVFYSVLVKALDNHDQ
EIYRIKLPGPAKLGEGKPENQNHAIVFTRGEALQTIDMNQDNYLEEALKMRNLLEEFHENHGVRQPTILGV
REHIFTGSVSSLAWFMSNQETSFVTIGQRVLANPLKVRFHYGHPDVFDRIFHITRGGISKASCGINLSEDIF
AGFNSTLRRGNVTHHEYIQVGKGRDVGLNQISLFEAKVACGNGEQTLSRDIYRLGHRFDFFRMLSCYFTT
VGFYISSMMVVIIVYVFLYGRLYLALSGLELAIMKQARMRGNTALQAAMGSQSIVQLGLLMALPMFMEIGLE
RGFRSALGDFIIMQLQLCSVFFTFSLGTKSHYFGRTILHGGAKYKATGRGFVVRHVKFPENYRMYSRSHF
VKGLELMLLLVVYQMYGDVATDSTAYILLTSSMWFLVITWLFAPFLFNPSGFEWQKIVDDWDDWSKWISS
RGGIGVPANKAWESWWEEEQEHLQSTGFFGRLSEIILSLRFFIFQYGIMYHLNISAGNKSISVYGLSWLVIV
AVVMVLKVVSMGRKKFSADFQLMFRLLKLFLFIGSIGTLAILFTLLHLTVGDIFASFLAFAPTGWAILQISQAS
KPVVKAFGLWGSVKALSRGYEYLMGILIFVPVAVLAWFPFVSEFQTRLLFNQAFSRGLQISRILAGGKKQS
>LOC_Os03g03610.1
MTSLSRTLSRGGPMQPPGQRRILRTQTAVNLGEQIFDSEVVPSSLVEIAPILRVANEVEASNPRVAYLCRF
YAFEKAHRLDPTSSGRGVRQFKTALLQRLERENEPTLRGRARKSDAREIQAFYQHYYKKYIQALQNVSDQ
VDRAQLTKAYQTANVLFEVLKAVTQQHSVEVDHEILEAADKVKEKTKIYLPFNILPLDPDSGNQAVMKFPE
TDNVSNQREHLILLLANVHIRRNPKTDPQSKLDDNALNEVMKKLFKNYKKWCKYLDRKSSLWLPTIQQEV
QQRKLLYMGLYLLIWGEAANLRFMPECICYIYHHMAFEMYGMLVGNVSALTGEYVKPAYGGEKEAFLKKV
VTPIYLTIAKEAERSKREKGNHSEWRNYDDLNEYFWSAECFRLGWPMRADADFFCQHLNSPDQRNETTR
TEKQKGKVNFVELRSFWHIFRSFDRMWSFFILALQVMVILAWNGGSLGNIFDPVVFKKILSIFITSAILNLGQ
ATLDIIFNWKARRTMEFAVKLRYVLKFTLAALWVVLLPVTYAYTWENPTGIIRAIKGWFGNGQNHPSLFVLA
VVIYLSPSLLAAILFLLPFLRRILESSDYKFVRFVMWWSQPRLFVGRGMHESAFSLFMYTMFWIALLLIKFAF
SYYVEIKPLVEPTKDIMKLPIHTFQWHEFFPKANGNIGVVIALWAPIILVYFMDTQIWYTIFSTLLGGIYGAFQ
RLGEIRTLGMLRSRFGSIPLAFNACLIPAEESDAKRKKGLKSYLHSRFERKHTDKEKIAARFAQMWNEIITS
FREEDLINNKEKELLLVPYVADQALEIMQWPPFLLASKIPIAVDMAKDSNGKDRDLKKRLENDYYFKCAIEE
CYASFKNIIKDLVQGEPEKRVINTIFAEVEKYIADDKVITDLNMHALPDLYNKFVELVKYLEKNDKNDRDAVI
KIFQDMLEVVTRDIMEDQLSSILESSHGGSYQRPEGTTTWDQEYQLFQPAGAIKFPVQFTDAWIEKIKRLE
LLLTVKESAMDVPSNLEARRRLTFFTNSLFMDMPDAPKVRNMLSFSALTPYYNEPVLFSIKELQEENEDGV
STLFYLQKIYPDEWKNFQQRVEWDEELKENEDKNEELRLWASYRGQTLARTVRGMMYYRKALVLEAFLD
MAKHEDLMEGYKAVESTDEQWKLQRSLFAQCEAVADMKFTYVVSCQQYGNDKRAALPNAQDILQLMRT
YPSLRVAYIDQVEDRVEEKKMEPAYYSTLVKVALTKDSESTDPVQNLDQVIYRIKLPGPAMLGEGKPENQ
NHAIIFTRGEGLQTIDMNQDNYMEEALKMRNLLQEFLTEHGVRRPSILGVREHIFTGSVSSLAWFMSNQEH
SFVTIGQRLLANPLKVRFHYGHPDVFDRIFHLTRGGVSKASRSINLSEDIFAGYNSTLRGGNITHHEYVQVG
KGRDVGLNQISKFEAKVANGNGEQTLSRDIYRLGHRFDFFRMLSCYFTTVGFYFSTLLTVVTVYVFLYGRL
YLALSGLEEGLLTQRRYIHNHPLQVALASQSLVQLGFLMALPMMMEIGLEKGFGQALSEFIMMNLQLAAVF
FTFSLGTKTHYYGRMLLHGGAQYRATGRGFVVFHAKFAENYRLYSRSHFVKGIELLILLIIYQLFGQSYRST
IAYIFVTFSMWFLVLTWLFAPFLFNPSGFEWTKIVDDWSDWNKWISNRGGIGVSPDKSWESWWEIELEHL
KYSGTIGLFVEIILSLRFFIYQYGLVYHLNITGDKSILVYLISWLVILVVLLVMKTVSVGRRRFSADFQLFFRLIK
FMIFVSFIAILIVLIAILHMTLRDIFVCFLAFLPSGWGILLIAQACKPLARRAGLWGSVRALARAYEIIMGVLLFT
PITILAWFPFVSEFQTRMLFNQAFSRGLQISRILGGQKKERERSSRNKD
>AT2G36850.1 GSL8
MARVYSNWDRLVRATLRREQLRNTGQGHERVSSGLAGAVPPSLGRATNIDAILQAADEIQSEDPSVARIL
CEQAYSMAQNLDPNSDGRGVLQFKTGLMSVIKQKLAKRDGASIDRDRDIERLWEFYKLYKRRHRVDDIQK

EEQKWRESGTTFSSNVGEILKMRKVFATLRALIEVLEVLSRDADPNGVGRSIRDELGRIKKADATLSAELTP
YNIVPLEAQSMTNAIGVFPEVRGAVQAIRYTEHFPRLPVDFEISGQRDADMFDLLEYIFGFQRDNVRNQRE
HLVLTLSNAQSQLSIPGQNDPKIDENAVNEVFLKVLDNYIKWCKYLRIRVVYNKLEAIDRDRKLFLVSLYFLI
WGEAANVRFLPECICYIFHNMAKELDAKLDHGEAVRADSCLTGTDTGSVSFLERIICPIYETISAETVRNNG
GKAAHSEWRNYDDFNEYFWTPACFELSWPMKTESRFLSKPKGRKRTAKSSFVEHRTYLHLFRSFIRLWIF
MFIMFQSLTIIAFRNEHLNIETFKILLSAGPTYAIMNFIECLLDVVLMYGAYSMARGMAISRLVIRFLWWGLGS
AFVVYYYVKVLDERNKPNQNEFFFHLYILVLGCYAAVRLIFGLLVKLPACHALSEMSDQSFFQFFKWIYQE
RYFVGRGLFENLSDYCRYVAFWLVVLASKFTFAYFLQIKPLVKPTNTIIHLPPFQYSWHDIVSKSNDHALTIV
SLWAPVLAIYLMDIHIWYTLLSAIIGGVMGAKARLGEIRTIEMVHKRFESFPEAFAQNLVSPVVKRVPLGQH
ASQDGQDMNKAYAAMFSPFWNEIIKSLREEDYLSNREMDLLSIPSNTGSLRLVQWPLFLLCSKILVAIDLA
MECKETQEVLWRQICDDEYMAYAVQECYYSVEKILNSMVNDEGRRWVERIFLEISNSIEQGSLAITLNLKK
LQLVVSRFTALTGLLIRNETPDLAKGAAKAMFDFYEVVTHDLLSHDLREQLDTWNILARARNEGRLFSRIA
WPRDPEIIEQVKRLHLLLTVKDAAANVPKNLEARRRLEFFTNSLFMDMPQARPVAEMVPFSVFTPYYSETV
LYSSSELRSENEDGISILFYLQKIFPDEWENFLERIGRSESTGDADLQASSTDALELRFWVSYRGQTLART
VRGMMYYRRALMLQSFLERRGLGVDDASLTNMPRGFESSIEARAQADLKFTYVVSCQIYGQQKQQKKPE
ATDIGLLLQRYEALRVAFIHSEDVGNGDGGSGGKKEFYSKLVKADIHGKDEEIYSIKLPGDPKLGEGKPEN
QNHAIVFTRGEAIQTIDMNQDNYLEEAIKMRNLLEEFHGKHGIRRPTILGVREHVFTGSVSSLAWFMSNQE
TSFVTLGQRVLAYPLKVRMHYGHPDVFDRIFHITRGGISKASRVINISEDIYAGFNSTLRQGNITHHEYIQVG
KGRDVGLNQIALFEGKVAGGNGEQVLSRDVYRIGQLFDFFRMMSFYFTTVGFYVCTMMTVLTVYVFLYG
RVYLAFSGADRAISRVAKLSGNTALDAALNAQFLVQIGIFTAVPMVMGFILELGLLKAIFSFITMQFQLCSVF
FTFSLGTRTHYFGRTILHGGAKYRATGRGFVVQHIKFADNYRLYSRSHFVKAFEVALLLIIYIAYGYTDGGA
SSFVLLTISSWFLVISWLFAPYIFNPSGFEWQKTVEDFEDWVSWLMYKGGVGVKGELSWESWWEEEQA
HIQTLRGRILETILSLRFFMFQYGIVYKLDLTRKNTSLALYGYSWVVLVVIVFLFKLFWYSPRKSSNILLALRF
LQGVASITFIALIVVAIAMTDLSIPDMFACVLGFIPTGWALLSLAITWKQVLRVLGLWETVREFGRIYDAAMG
MLIFSPIALLSWFPFISTFQSRLLFNQAFSRGLEISIILAGNRANVET
>AT3G07160.1 GSL10
MSRAESSWERLVNAALRRDRTGGVAGGNQSSIVGYVPSSLSNNRDIDAILRAADEIQDEDPNIARILCEHG
YSLAQNLDPNSEGRGVLQFKTGLMSVIKQKLAKREVGTIDRSQDILRLQEFYRLYREKNNVDTLKEEEKQL
RESGAFTDELERKTVKRKRVFATLKVLGSVLEQLAKEIPEELKHVIDSDAAMSEDTIAYNIIPLDAPVTTNAT
TTFPEVQAAVAALKYFPGLPKLPPDFPIPATRTADMLDFLHYIFGFQKDSVSNQREHIVLLLANEQSRLNIP
EETEPKLDDAAVRKVFLKSLENYIKWCDYLCIQPAWSNLEAINGDKKLLFLSLYFLIWGEAANIRFLPECLC
YIFHHMVREMDEILRQQVARPAESCMPVDSRGSDDGVSFLDHVIAPLYGVVSAEAFNNDNGRAPHSAWR
NYDDFNEYFWSLHSFELGWPWRTSSSFFQKPIPRKKLKTGRAKHRGKTSFVEHRTFLHLYHSFHRLWIFL
AMMFQALAIIAFNKDDLTSRKTLLQILSLGPTFVVMKFSESVLEVIMMYGAYSTTRRLAVSRIFLRFIWFGLA
SVFISFLYVKSLKAPNSDSPIVQLYLIVIAIYGGVQFFFSILMRIPTCHNIANKCDRWPVIRFFKWMRQERHY
VGRGMYERTSDFIKYLLFWLVVLSAKFSFAYFLQIKPLVGPTRMIVKQNNIPYSWHDFVSRKNYNALTVAS
LWAPVVAIYLLDIHIFYTIFSAFLGFLLGARDRLGEIRSLEAIHKLFEEFPGAFMRALHVPLTNRTSDTSHQTV
DKKNKVDAAHFAPFWNQIIKSLREEDYITDFEMELLLMPKNSGRLELVQWPLFLLSSKILLAKEIAAESNSQ
EEILERIERDDYMKYAVEEVYHTLKLVLTETLEAEGRLWVERIYEDIQTSLKERNIHHDFQLNKLSLVITRVT
ALLGILKENETPEHAKGAIKALQDLYDVMRLDILTFNMRGHYETWNLLTQAWNEGRLFTKLKWPKDPELKA
LVKRLYSLFTIKDSAAHVPRNLEARRRLQFFTNSLFMDVPPPKSVRKMLSFSVFTPYYSEVVLYSMAELTK

RNEDGISILFYLQKIYPDEWKNFLARIGRDENALEGDLDNERDILELRFWASYRGQTLARTVRGMMYYRKA
LMLQSYLERKAGNDATDAEGFELSPEARAQADLKFTYVVTCQIYGRQKEDQKPEAVDIALLMQRNEALRI
AYIDVVDSPKEGKSHTEYYSKLVKADISGKDKEIYSIKLPGDPKLGEGKPENQNHAIVFTRGNAIQTIDMNQ
DNYFEEALKMRNLLEEFDRDHGIRPPTILGVREHVFTGSVSSLASFMSNQETSFVTLGQRVLAKPLKIRMH
YGHPDVFDRVFHITRGGISKASRVINISEDIFAGFNTTLRQGNVTHHEYIQVGKGRDVGLNQIALFEGKVAG
GNGEQVLSRDVYRLGQLLDFFRMMSFFFTTVGFYLCTMLTVLTVYIFLYGRAYLALSGVGATIRERAILLDD
TALSAALNAQFLFQIGVFTAVPMVLGFILEQGFLQAIVSFITMQFQLCTVFFTFSLGTRTHYFGRTILHGGAR
YQATGRGFVVKHIKFSENYRLYSRSHFVKAMEVILLLVVYLAYGNDEAGAVSYILLTVSSWFLAVSWLFAP
YLFNPAGFEWQKVVEDFKEWTNWLFYRGGIGVKGAESWEAWWEEELSHIRTLSGRIMETILSLRFFIFQY
GIVYKLKLQGSDTSFAVYGWSWVAFAMIIVLFKVFTFSQKISVNFQLLLRFIQGLSLLMALAGIIVAVVLTPLS
VTDIFACVLAFIPTGWGILSIACAWKPVLKRMGMWKSIRSLARLYDALMGMLIFLPVALCSWFPFVSTFQTR
MMFNQAFSRGLEISLILAGDNPNSGL
>AT5G36870.1 GSL9
MNQPNRGQILQTVFSHFFPVASPDSELVPSSLHEDITPILRVAKDVEDTNPRSLFLQDLDIKSVDDSINILSG
HSHALDKANELDPTSSGRDVRQFKNTILQWLEKNNESTLKARQKSSDAHEMQSFYQQYGDEGINDLLNA
GAGSSSSQRTKIYQTAVVLYDVLDAVHRKANIKVAAKILESHAEVEAKNKIYVPYNILPLDPDSKNHAMMRD
PKIVAVLKAIRYTSDLTWQIGHKINDDEDVLDWLKTMFRFQKDNVSNQREHLILLLANVQMRQTQRQPNLL
DDRALDTVMEKLLGNYNKWCNHVGLESSLRFPKDKQQKVVQQRKLLYTGLYLLIWGEAANLRFMPECLC
YIYHHMAFELFEMLESKGSKKKYKPKNPTYSGKDEDFLTKVVTPVYKTIAEEAKKSGEGKHSEWRNYDDL
NEYFWSKQYLDKLGWPMKANADFFCKTSQQLGLNKSEKKPDLGDGCVGKVNFVEIRTFWHLFRSFDRM
WSFYILSLQAMIIIAWNETSESGGAVFHKVLSVFITAAKLNLFQAFLDIALSWKARHSMSTHVRQRYIFKAVA
AAVWVLLMPLTYAYSHTSIFIVAILIYLSPNMLPEMLLLIPSIRRTLEKSDFRPVKLIMWWSQPELYIGRGMH
ESAWSIYKYMMFWIVLLTSKLAFSYYVEQIKPLMGPTKEIMSVPMPGYWLPEFFPHVKNNRGVVITLWSPV
ILVYFMDTQIWYAIVSTLVGGLYGAFRHIGEIQTLGMLRSRFQSLPGAFNACLIPNENTKEKGIKLAFSRKCH
KIPNTNGKEAKQFSQMWNTIINSFREEDLISNRELELLLMSCWAYPDLDFIRWPIFLLASKIPIAVDIAKKRN
GKHRELKNILAEDNCMSCAVRECYASIKKLLNTLVTGNSDLMLITTVFTIIDTHIEKDTLLTELNLSVLPDLHG
HFVKLTEYVLQNKDKDKIQIVNVLLKILEMVTKDILKEEIKRLHLLLTVKESAMDVPSNLEARRRLTFFSNSLF
MEMPGAPKIQNMLSFSALTPYYSEDVLFSTFDLEKENDGVSILFYLQKIFPDEWKNFLERVKCGTEEELDAI
DYLKEEIRLWASYRGQTLTKTVRGMMYYQKALELQAFFDLANERELMKGYKSAEASSSGSSLWAECQAL
ADIKFTYVVACQQYSIHKRSGDQRAKDILTLMTTYPSLRVAYIDEVEQTHIYSKGTSENFYYSALVKAAPQT
YSTDSSDSGHMLDQVIYQIKLPGPPIIGEGKPENQNNAIIFTRGEALQTIDMNQDYYIEEAFKMRNLLQEFLE
KNGGVRYPTILGLREHIFTRSVSCLAWFMSNQEHSFVTIGQRVLANPLKVRFHYGHPDVFDRVFHLTRGG
VSKASKVINLSEDIFAGFNSTLREGTVSHHEYIQVGKGRDVGLNQISMFEAKIANGSGEQTLSRDLYRLGH
QFDFFRMLSCYFTTVGFYFCSMLTVLTVYVFLYGRLYLVLSGVEKELGNKPMMMEIILASQSFVQIVFLMA
MPMIMEIGLERGFYDALFDFVLMQLQLASVFFTFQLGTKFHYYCKTLLHGGAEYRGTGRGFVVFHAKFAE
NYRFYSRSHFVKATELGILLLVYHIFGPTYIGLFTISIWFMVGTWLFAPFLFNPSGFEWHEIVEDWADWKK
WIEYDNGGIGVPPEKSWESWWEKDIEHLQHSGKWGIVVEIFFALRFFIFQYGLVYQLSAFKNKYSSLWVF
GASWLLILILLLTVTVLDYARRRLGTEFQLLFRIIKVSLFLAFMAIFITLMTCRLILPQDVFLCMLALIPTGWGLL
LIAQSCKPLIQQPGIWSWVMTLAWVYDLVMGSLLFIPIAFMAWFPFISEFQTRMLFNQAFSRGLHISRILSG
QRKHRSSKNKD
>LOC_Os06g51270.1

MAAPGGARRPDFSSAASPSPSPAGAAGRRLLRTQTVGNLGESIFDSEVVPSSLVEIAPILRVANEVEATNP
RVAYLCRFYAFEKAHRLDPTSNGRGVRQFKTALLQRLERENDPTLKGRVHQSDAREMQRFYREYYKKYI
QALQNAADKADRALLTKAYQTAAVLFEVLKAVNVSQSVEVDQAILDTHNKVEEKKKLYVPYNILPLDPEST
YQPIMQYPEIQAAVNALRNIRGLPWPKEHEKKPDEKKTGKDLLDWLQAMFGFQKDNVSNQREHLILLLAN
VHIRQSPKTEQQAKLDDRALDAVMKKLFKNYKKWCKYLGRKSSLWLPTIQQEVQQRKLLYMGLYLLIWGE
AANLRFMPECLCYIYHHIMEALLNSINYIEMAFELYGMLAGNVSPMTGENVKPAYGGDEEAFLMKVVTPIY
KVIEKEAERSKTIKSKHSHWRNYDDLNEYFWSVDCFRLGWPMRADADFFKTPEDAYPSRLNGIIGMDYIS
SLVLQCNAKKKKYLLGVLILWVPISRLVANAGEQIRRRFLESSNVKVITFIMWWSQPRLFVGRGMHEGAFS
LFKYTMFWVLLLAMKLTIKPLVQPTKDIMKEPIRDFQWHEFFPRANNNIGVVIALWAPIILVYFMDTQIWYAL
FSTLIGGIYGAYRRLGEIRTLGMLRSRFESLPEAFNEHLIPSDSHKSKGLRAAFTGKPSKTSGDEQEKEKIA
ARFAQMWNLIITSFREEDLIDNREMDLLLVPYCKDRELNIFQWPPFLLASKIPIALDMAADSGGKDRDLKKR
MGSDPYFSYAIRECYGSFKNIINTLVFGQREKIVIQQIFTIVDEHIEGGSLIKDLNMRSLPALSKKFIELLELLQ
KNKEEDLGQVVILFQDMLEVVTRDIMDEQDQLGGLLDSVHGGNRKHEGMTSLDQQDQLFTKAIRFPVEE
SNAWTEKIKRLHLLLTVKESAMDVPTNLDARRRISFFANSLFMEMPNAPKVRHMLPFSVLTPYYKEDVLFS
SHNLEEPNEDGVSILFYLQKIYPDEWKNFLDRVDRKSEEELREDETLEEELRLWASYRGQTLTRTVRGMM
YYRKALELQAFLDMAKDDDLMEGYRATELMSEDSQLMTQCKAIADMKFTYVVSCQQYGIQKRSGEACAH
DILRLMTVRAVVIVYPSLRVAYIDEVEAPSQDRNKKTDKVYYSALVKASVTKPNEPGQSLDQVIYKIKLPGN
AILGEGKPENQNHAIIFTRGECLQTIDMNQEHYMEEALKMRNLLDEFLKKHDGVRYPSILGVREHIFTGSVS
SLAWFMSNQETSFVTIGQRVLANPLRVRFHYGHPDIFDRLFHLTRGGVSKASKIINLSEDIFAGFNSTLREG
NVTHHEYMQVGKGRDVGLNQIALFEAKIANGNGEQTLSRDIYRLGHRFDFFRMLSCYYTTIGFYFSTMITV
WTVYVFLYGRLYLVLSGLDQALATGKKFVHNAPLQVALASESFVQLGFLMALPMMMEIGLERGFRTALSD
FVLMQLQLASVFFTFSLGTKTHYYGRTLLHGGAEYRATGRGFVVFHAKFADNYRLYSRSHFVKGIELMILL
VVYEIFGQSYRGAITYIFITVSMWFMVGTWLFAPFLFNPSGFEWQKIVDDWTDWNKWISNRGGIGVAPTK
SWESWWEKEQEPLRYSGKRGTILEILLALRFFVYQYGLVYHLNITKHTRSVLVYCFSWVVIFVILLVMKTVS
VGRRRFSAEFQLVFRLIKGLIFITFVAIVVILIAIPHMTVLDIFVCILAFMPTGWGLLLIAQAIKPAVQAIGLWGSI
KALARGYEILMGLLLFTPIAFLAWFPFVSEFQTRMLFNQAFSRGLQISRILGGHKKDRSTRNKE
>LOC_Os01g55040.1
MSLLRNRRAAAAAGSGEQTVVQAAYNIIPIQDVVMHGDHPSLQVPEVRAAVEALSHASDFPAPPLARVWD
PHRADIFDWLGATFGFQADNVRNQREHLVLLLANAQLRAAPGFPKDHPIDVLHLTVARGIRRKLLKNYTS
WCAYLGQKRHFRVPSGGGGGRRTGAATGNDVRMDLLYTALYLLIWGEAANLRFMPECLCYIFHYMALDL
HHVVEQSIDIETGRPAMPAVCGEDAFLIRVVTPIYNVLKNEVEASRNGTKPHSAWRNYDDVNEYFWSRRV
FKRLRWPLDPSRSFFVEPGKTGRIGKTGFVEQRSFWNVYRSFDRVWVMHILFFQAAMIVAWDGKTPWVS
LRFRDIQVRVLSVFITWGGLRFVQAMLDAGTQYSLVSRETKTVAVRMVLKVLVAAGWTITFSVLYKRMWD
QRWRDRRWSFAANTRVLNYLEAAAVFVIPQVLAIVLFIIPWIRNFLEKTNWKILYVLTWWFQTRTFVGRGL
REGLIDNIKYSIFWVCLLVSKFSFSYFLQIKPMVGPTKVIFKLHDIKRNWFEFMPHTERLAVIILWLPVIIIYLM
DIQIWYAVFSSLTGALIGLFSHLGEIRSVEQLRLRFQFFASAMQFNLMPEEHLDTVHGGIRSKFYDAINRLK
LRYGFGRPYRKIEANEVEAKRFALVWNEIIQTFREEDIISDKELGLLELPAVVWRIRVVRWPCLLLKNELLLA
LSQAAELVADDRTHWNKICNNEYRRCAVIEAYDSIRHLLLEIIKERTNEHIIVNQLFLAFDGAMEYGKFTEEY
RLTLLPQIHKYVISLVEQLLLKDKDQIKIVRTLQDLYDLAVHDFPKIKKDFEQLRREGLALSRPTESQLLFQD
AIKCPDDNDVSFYKQVRRLHTILTSRDSMDDVPKNPEARRRITFFSNSLFMNMPRAPTVQRMMAFSVLTP
YYNEDVLYNKDQLRRENEDGISILFYLQKIYEDDWKNFLERMQREGMASDDGIWAGKFQDLRLWASYRG

QTLARTVRGMMYYYRALKMLAFLDNASEVEITEGTKQLASFGSVQYENDVYPMNGGLSQRPRRRLERGT
STVSQLFKGQEDGAAIMKYTYVVACQIYGNQKKAKDQRAEDILTLMKKNDALRVAYVDEVHPEIGDTQYY
SVLVKFDPVLQREVEIYRIRLPGQLKLGEGKPENQNHAIIFTRGDAVQTIDMNQDNYFEEALKMRNLLEQY
DYYHGSQKPTLLGVREHVFTGSVSSLAWFMSAQETSFVTLGQRVLANPLKVRMHYGHPDVFDRLWFLT
RGGISKASRVINISEDIFAGFNCTLRGGNVSHHEYIQVGKGRDVGLNQISMFEAKVSSGNGEQTLSRDIYR
LGHRLDFFRSLSVFYTTVGFYFNTMMVVLTVYTFVWGRLYLALSGLEAGIQGSANATNNKALGAVLNQQF
VIQLGLFTALPMIIENSLEQGFLPAVWDFFTMQMMFSSVFYTFSMGTKSHYYGRTILHGGAKYRATGRGF
VVQHKSFAENYRLYARSHFIKAIELGIVLTVYAAHSVIARDTLVYIIMMISSWFLVVSWIMAPFAFNPSGFDW
LKTVYDFDDFMNWIWYPGSIFSKAEHSWEVWWFEEQDHLRTTGLWGKILEILLDLRYFFFQYGVVYQLKI
ANESRSIAVYLLSWICVAVIFGIFVLMSYARDKYAAKQHLYYRVIQSGVIILAVLVLIIFLKFTKFQIIDIFTSLLA
FIPTGWGLISIAQVIRPFIESTVVWASVVSVARLYEILLGVFVMAPVAFFSWLPGFQEMQTRVLFNEAFSRG
LQISRILAGKKTIAV
>AT4G03550.1 GSL5
MSLRHRTVPPQTGRPLAAEAVGIEEEPYNIIPVNNLLADHPSLRFPEVRAAAAALKTVGDLRRPPYVQWRS
HYDLLDWLALFFGFQKDNVRNQREHMVLHLANAQMRLSPPPDNIDSLDSAVVRRFRRKLLANYSSWCSY
LGKKSNIWISDRNPDSRRELLYVGLYLLIWGEAANLRFMPECICYIFHNMASELNKILEDCLDENTGQPYLP
SLSGENAFLTGVVKPIYDTIQAEIDESKNGTVAHCKWRNYDDINEYFWTDRCFSKLKWPLDLGSNFFKSR
GKSVGKTGFVERRTFFYLYRSFDRLWVMLALFLQAAIIVAWEEKPDTSSVTRQLWNALKARDVQVRLLTV
FLTWSGMRLLQAVLDAASQYPLVSRETKRHFFRMLMKVIAAAVWIVAFTVLYTNIWKQKRQDRQWSNAA
TTKIYQFLYAVGAFLVPEILALALFIIPWMRNFLEETNWKIFFALTWWFQGKSFVGRGLREGLVDNIKYSTF
WIFVLATKFTFSYFLQVKPMIKPSKLLWNLKDVDYEWHQFYGDSNRFSVALLWLPVVLIYLMDIQIWYAIYS
SIVGAVVGLFDHLGEIRDMGQLRLRFQFFASAIQFNLMPEEQLLNARGFGNKFKDGIHRLKLRYGFGRPFK
KLESNQVEANKFALIWNEIILAFREEDIVSDREVELLELPKNSWDVTVIRWPCFLLCNELLLALSQARELIDA
PDKWLWHKICKNEYRRCAVVEAYDSIKHLLLSIIKVDTEEHSIITVFFQIINQSIQSEQFTKTFRVDLLPKIYET
LQKLVGLVNDEETDSGRVVNVLQSLYEIATRQFFIEKKTTEQLSNEGLTPRDPASKLLFQNAIRLPDASNED
FYRQVRRLHTILTSRDSMHSVPVNLEARRRIAFFSNSLFMNMPHAPQVEKMMAFSVLTPYYSEEVVYSKE
QLRNETEDGISTLYYLQTIYADEWKNFKERMHREGIKTDSELWTTKLRDLRLWASYRGQTLARTVRGMM
YYYRALKMLAFLDSASEMDIREGAQELGSVRNLQGELGGQSDGFVSENDRSSLSRASSSVSTLYKGHEY
GTALMKFTYVVACQIYGSQKAKKEPQAEEILYLMKQNEALRIAYVDEVPAGRGETDYYSVLVKYDHQLEK
EVEIFRVKLPGPVKLGEGKPENQNHAMIFTRGDAVQTIDMNQDSYFEEALKMRNLLQEYNHYHGIRKPTIL
GVREHIFTGSVSSLAWFMSAQETSFVTLGQRVLANPLKVRMHYGHPDVFDRFWFLSRGGISKASRVINIS
EDIFAGFNCTLRGGNVTHHEYIQVGKGRDVGLNQISMFEAKVASGNGEQVLSRDVYRLGHRLDFFRMLS
FFYTTVGFFFNTMMVILTVYAFLWGRVYLALSGVEKSALADSTDTNAALGVILNQQFIIQLGLFTALPMIVE
WSLEEGFLLAIWNFIRMQIQLSAVFYTFSMGTRAHYFGRTILHGGAKYRATGRGFVVEHKGFTENYRLYA
RSHFVKAIELGLILIVYASHSPIAKDSLIYIAMTITSWFLVISWIMAPFVFNPSGFDWLKTVYDFEDFMNWIWY
QGRISTKSEQSWEKWWYEEQDHLRNTGKAGLFVEIILVLRFFFFQYGIVYQLKIANGSTSLFVYLFSWIYIF
AIFVLFLVIQYARDKYSAKAHIRYRLVQFLLIVLAILVIVALLEFTHFSFIDIFTSLLAFIPTGWGILLIAQTQRKW
LKNYTIFWNAVVSVARMYDILFGILIMVPVAFLSWMPGFQSMQTRILFNEAFSRGLRIMQIVTGKKSKGDV
>AT4G04970.1 GSL1
MRRQRPSVATARDAPSLEVYNIIPIHDFLTEHPSLRYPEVRAAAAALRIVGDLPKPPFADFTPRMDLMDWL
GLLFGFQIDNVRNQRENLVLHLANSQMRLQPPPRHPDGLDPTVLRRFRKKLLRNYTNWCSFLGVRCHVT

SPIQSRHQTNAVLNLRRELLYVALYLLIWGESANLRFMPECLCYIFHHMAMELNKVLAGEFDDMTGMPYW
PSFSGDCAFLKSVVMPIYKTVKTEVESSNNGTKPHSAWRNYDDINEYFWSKRALKSLKWPLDYTSNFFDT
TPKSSRVGKTGFVEQRSFWNVYRSFDRLWILLLLYLQAAIIVATSDVKFPWQDRDVEVALLTVFISWAGLR
LLQSVLDASTQYSLVSRETYWLFIRLTLKFVVAVAWTVLFSVFYARIWSQKNKDGVWSRAANERVVTFLK
VVFVYVIPELLALVLFIVPCIRNWVEELNLGVVYFLTWWFYSKTFVGRGMREGLVDNVKYTLFWIIVLATKFI
FSYFLQIRPLIAPTRALLNLKDATYNWHEFFGSTHRIAVGMLWLPVILVYLMDLQIWYSIYSSLVGATIGLFS
HLGEIRNIDQLRLRFQFFSSAMQFNLKPEEHLLSPKATMLKKARDAIHRLKLRYGIGQPFNKIESSQVEATW
FALIWNEIILTFREEDLISDREVELLELPPNCWNIRVIRWPCFLLCNELLLALSQANELCDAPDHWLWSKICS
SEYRRCAVMEAFDSIKFVILKIVKNGTEEESILNRLFMEIDENVENEKITEVYKLTVLLRIHEKLISLLERLMDP
EKKVFRIVNILQALYELCAWEFPKTRRSTPQLRQLGLAPISLEADTELLFVNAINLPPLDDVVFYRQIRRVHTI
LTSRDPMHNVPKNIEARERLAFFSNSLFMTMPQAPSVEKMMAFSVLTPYYDEEVMYRQEMLRAENEDGI
STLFYLQRIYEDEWVNFLERMRREGAENENDIWSKKVRDLRLWASYRGQTLSRTVRGMMYYYSALKKLA
FLDSASEMDIRMGTQIAPEARRSYYTNDGGDNTLQPTPSQEISRMASGITHLLKGSEYGSAMMKFTYVVA
CQVYGQHKARGDHRAEEILFLMKNHDALRIAYVDEVDLGRGEVEYYSVLVKFDQQLQREVEIYRIRLPGPL
KLGEGKPENQNHALIFTRGDAIQTIDMNQDNHFEEALKMRNLLESFKTYYGIRKPTILGVREKVFTGSVSSL
AWFMSAQETSFVTLGQRVLANPLKVRMHYGHPDVFDRFWFVPRGGISKASRVINISEDIFAGFNCTLRGG
NVTHHEYIQVGKGRDVGLNQISMFEAKVASGNGEQALSRDVYRLGHRLDFFRMLSFFYTTVGYYFNTMLI
VFTVYAFLWGRLYLALSGVEKIAKDRSSSNEALGAILNQQFIIQLGLFTALPMILENSLERGFLPAVWDFITM
QLQLASFFYTFSMGTRTHYFGRTILHGGAKYRATGRGFVVEHKKFAENYRLYARTHFIKAIELAIILLVYAAY
SPLAKSSFVYILMTISSWFLITSWIISPFLFNPSGFDWLKTVNDFDDFIAWLWSRGGLFTKADQSWFTWWN
EEQEHLKTTGVWGKLLEIILDLRFFFFQYSIVYHLRIAENRTSIGVYLISWGCIIGIVAIYITTIYAQKRYSVKEHI
KYRFIQFLVILLTVLVVVMMLQFTKLTVVDLLISLLAFVPTGWGLISIAQVLKPFLLSTVVWDTVISVARFYDLF
FGLIVMAPVALLSWLPGFQNMQTRILFNEAFSRGLQISIILAGKKST
>LOC_Os02g14900.1
MHARLNPKSSSETMLDDRAVDELLAKTFENYLTWCKFLGRKSNIWLPSVKQEIQQHKLLYISLYLLIWGEA
SNLRLMPECLCYIFHHESLKNKNGVSDHSTWRNYDDLNEFFWSADCFKLGWPMRLNNDFFFTSNKNKN
SRLPIVPPVQQTEQQINQLRTSQQTDQQNTQLRTSQQTEQRNTQLRTPNGSSSFQNMLNPEAPGQTQQ
QTTSDTSQQKWLGKTNFVEVRSFWHIFRSFDRMWTLLVLGLQFFRDIYLENLQYVSLVVSVKSNAGAILA
VWAPIILVYFMDTQIWYSVFCTIFGEMDLMTMPMSLEHRSGSIRWPMFLLAKKFSEAVDMVANFTGKSTR
LFCIIKKDNYMLCAINDFYELTKSILRHLVIGDVEKSFSSACPCEYYYDVLQILSRVIAAIYTEIEKSIQNASLLV
DFKMDHLPSLVAKFDRLAELLYTNKQELRYEVTILLQDIIDILVQDMLVDAQSVLGLINSSETLISDDDGTFEY
YKPELFASISSISNIRFPFPENGPLKEQVKRLYLLLNTKEKVVEVPSNLEARRRISFFATSLFMDMPSAPKVS
NEWRNFLERLGPKVTQEEIRYWASFHGQTLSRTVRGMMYYRKALRLQAFLDRTNDQELCKGPAANGRQ
TKNMHQSLSTELDALADMKFSYVISCQKFGEQKSSGNPHAQDIIDLMTRYPALRVAYIEEKEIIVDNRPHKV
YSSVLIKAENNLDQEIYRIKLPGPPLIGEGKPENQNHAIIFTRGEALQTIDMNQDNYLEEAYKMRNVLQEFV
RHPRGKAPTILGLREHIFTGSVSSLAGFMSYQETSFVTIGQRFLADPLRVRFHYGHPDIFDRMFHLTRGGI
SKASKTINLSEDVFAGYNSILRRGHITYNEYIQVGKGRDVGLNQISKFEAKVANGNSEQTLSRDIHRLGRRF
DFFRMLSCYFTTVGFYFNSLISVVGVYVFLYGQLYLVLSGLQRALLIEAETQNMKSLETALVSQSFLQLGLL
TGLPMVMELGLEKGFRVALSDFILMQLQLASVFFTFSLGTKAHYYGRTILHGGAKYRPTGRKFVAFHASFT
ENYQLYSRSHFVKGFELVFLLIIYHIFRRSYVSTVVHVMITYSTWFMAVTWLFAPFLFNPAGFAWRKIVEDW
ADWTIWMRNQGGIGVQPEKSWESWWNAENAHLRHSVLSSRILEVLLSLRFFMYQYGLVYHLKISQDNKN

FLVYLLSWVVIIAIVGLVKLVNCASRRLSSKHQLVFRLIKLLIFLSVMTSLILLSCLCQLSIMDLIICCLAFIPTG
WGLLLIVQVLRPKIEYYAIWEPIQVIAHAYDYGMGSLLFFPIAALAWMPVISAIQTRVLFNRAFSRQLQIQPFI
AGKTKRR
>LOC_Os01g34890.1
MPMGSRLGASPEMIERNMSLMLLLIVEKTGPFDIYAKEVEKEKASFSHYNILPLNISGQRQPVMEIPEIKAA
VDLLRKIDGLPMPRLDPVSAEKETDVPTVRDLFDWLWLTFGFQKGNVENQKEHLILLLANIDMRKGANAY
QSDRHNHVMHSDTVRSLMRKIFENYISWCRYLHLESNIKIPNDASTQQPEILYIGLYLLIWGEASNVRFMPE
CICYIFHHSHQYKNTIIPMCLFMEHVRQDFDPPFRREGSDDAFLQLVIQPIYSVMKQEAAMNKRGRTSHSK
WRNYDDLNEYFWSKRCFKQLKWPMDSAADFFAVPLKIKTEEHHDRVITRRRIPKTNFVEVRTFLHLFRSF
DRMWAFFILAFQAMVIVAWSPSGLPSAIFDPTVFRNVLTIFITAAFLNFLQATLEIILNWKAWRSLECSQMIR
YILKFVVAVAWLIILPTTYMSSIQNSTGLIKFFSSWIGNLQSESIYNFAVALYMLPNIFSALFFIFLPFRRVLER
SNSRIIRFFLWWTQPKLYVARGMYEDTCSLLKYTLFWILLLICKLAFSFYVEIYPLVGPTRTIMFLGRGQYA
WHEFFPYLQHNLGVVITVWAPIVMVYFMDTQIWYAIFSTICGGVNGAFSRLGEIRTLGMLRSRFEAIPIAFG
KHLVPGHDSQPKRHEHEEDKINKFSDIWNAFIHSLREEDLISNRERNLLIVPSSMGDTTVFQWPPFLLASKI
PIALDMANSVKKRDEELRKRINQDPYTYYAVVECYQTLFSILDSLIVEQSDKKVVDRIHDRIEDSIRRQSLVK
EFRLDELPQLSAKFDKLLNLLLRTDEDIEPIKTQIANLLQDIMEIITQDIMKNGQGILKDENRNNQLFANINLDS
VKDKTWKEKCVRLQLLLTTKESAIYVPTNLDARRRITFFANSLFMKMPKAPQPFCFCISVLTPYFKEEVLFS
AEDLYKKNEDGISILFYLRKIYPDEWKNFLERIEFQPTDEESLKTKMDEIRPWASYRGQTLTRTVKLEHRRT
VESSQQGWASFDMARAIADIKFTYVVSCQVYGMQKTSKDPKDKACYLNILNLMLMYPSLRVAYIDEVEAP
AGNGTTEKTYYSVLVKGGEKYDEEIYRIKLPGKPTDIGEGKPENQNHAIVFTRGEALQAIDMNQDNYLEEA
FKMRNVLEEFESEKYGKRKPTILGLREHIFTGSVSSLAWFMSNQETSFVTIGQRVLANPLNFYGPSFIDRH
H
>LOC_Os02g58560.1
MDVPTNLDARRRISFFANSLFMDMPSAPKVRHMLPFSVLTPYYKEDVLFSSQALEDQNEDGVSILFYLQKI
YPDEWKHFLQRVDCNTEEELRETEQLEDELRLWASYRGQTLTRTVRGMMYYRQALVLQAFLDMARDED
LREGFRAADLLNDESPLLTQCKAIADMKFTYVVSCQQYGIQKRSGDHRAQDILRLMTTYPSLRVAYIDEVE
EPSKDRNKKIEKVYYSALVKAAVTKPDDPGQKLDQDIYRIKLPGNAMLGEGKPENQNHAIIFTRGEGLQTID
MNQEHYMEETLKMRNLLQEFLKKHDGVRYPSILGVREHIFTGSVSSLAWFMSNQETSFVTIGQRVLANPL
RVRFHYGHPDIFDRLFHLTRGGVSKASKIINLSEDIFAGFNSTLREGNVTHHEYMQVGKGRDVGLNQISLF
EAKIANGNGEQTLSRDVYRLGHRFDFFRMLSCYYTTIGFYFSTMMTVWTVYVFLYGRLYLVLSGLDEALA
TGKRFIHNEPLQVALASQSFVQLGFLMALPMMMEIGLERGFRTALSDFVLMQLQLASVFFTFSLGTKTHYY
GTTLLHGGAEYRATGRGFVVFHAKFAENYRLYSRSHFVKGIELLILLIVYEIFGQSYRGAIAYIFITFSMWFM
VVTWLFAPFLFNPSGFEWQKIVDDWTDWNKWISNRGGIGVPPEKSWESWWEKEQEPIKYSGKRGIVLEI
VLALRFFIYQYGLVYHLNITKHTKSVLVYCLSWVVIFVILLVMKTVSVGRRKFSADFQLVFRLIKGLIFITFISIIII
LIAIPHMTVQDIFVCILAFMPTGWGLLLVAQAIKPVIVRIGLWGSIKALARGYEIIMGLLLFTPIAFLAWFPFVS
EFQTRMLFNQAFSRGLQISRILGGHKKDRATRNKE
>LOC_Os01g34930.1
MCIMQISFVLCSKLMVVLIKGFNSTLRQGNVTHHEYIQLGKGRDVGMNQISNFEAKVANGNGEQTLCRDIY
RLGHRFDFYRMLSLYFTTVGFYFNSMVAVLTVYVFLYGRLYLVLSGLEKSILQDPQIKNIKPFENALATQSIF
QLGMLLVLPMMIEVGLEKGFGRALGEFVIMQLQLASVFFTFHLGTKTHYYGRTILHGGAKYRGTGRGFVV
RHAKFAENYRMYSRSHFVKALELLILLVVYLAYGISYRSSSLYLYVTISIWFLVFCWLFAPFVFNPSCFEWH

KTVDDWTDWWHWMSNRGGIGLAPEQSWEAWWISEHDHLRNGTIRSLLLEFVLSLRFLIYQYGIVYHLHIV
HGNRSFMVYALSWLVIAIVLVSLKVVSMGREKFITNFQLVFRILKGIVFIVLISLVVILFVVFNLTVSDVGASIL
AFIPTGWFILQIAQLCGPLFRRLVTEPLCALFCSCCTGGTACKGRCCARFRLRSRDVLRKIGPWDSIQEMA
RMYEYTMGILIFFPIAVLSWFPFVSEFQTRLLFNQAFSRGLQISRILTGQNGSGSKRD
>LOC_Os01g34880.1
MREGNVTHHEYMQVGKGRDVGMNQISSFEAKVANGNGEQTLSRDIYRLGRRFDFYRMLSFYFTTVGFY
FSSMVTVLTVYVFLYGRLYLVMSGLERSILLDPRIEQNIKPLENALASQSFFQLGLLLVLPMVMEVGLEKGF
RTALGEFVIMQLQLASVFFTFQLGTKTHYYGRTILHGGAKYRPTGRGFVVYHAKFADNYRMYSRSHFVKG
LELLILLVVYLVYGSSYRSSSMYLFVTFSIWFLVASWLFAPFIFNPSCFEWQKTVDDWTDWRKWMGNRGG
IGMSVDQSWEAWWISEQEHLRKTSIRSLLLEIILSLRFLIYQYGIVYHLNIARRSKSILVYGLSWLVMLSVLVV
LKMVSIGRQKFGTDLQLMFRILKGLLFLGFVSVMAVLFVVCNLTISDVFASILGFMPTGWCILLIGQACSPLV
KKAMLWDSIMELGRSYENLMGLVLFLPIGLLSWFPFVSEFQTRLLFNQAFSRGLQISRILAGQKDIGEE
>LOC_Os01g48200.1
MFEAKVASGNGEQTLSRDVYRLGHRLDFFRMLSFFYTTIGFYFNTMMVVLTVYAFVWGRFYLALSGLEAF
ISSNTNSTNNAALGAVLNQQFVIQLGIFTALPMIIENSLEHGFLTAVWDFIKMQLQFASVFYTFSMGTKTHY
YGRTILHGGAKYRATGRGFVVEHKKFAENYRLYARSHFIKAIELGVILTLYASYGSSSGNTLVYILLTISSWF
LVLSWILAPFIFNPSGLDWLKNFNDFEDFLNWIWFRGGISVKSDQSWEKWWEEETDHLRTTGLFGSILEIIL
DLRFFFFQYAIVYRLHIAGTSKSILVYLLSWACVLLAFVALVTVAYFRDKYSAKKHIRYRLVQAIIVGATVAAI
VLLLEFTKFQFIDTFTSLLAFLPTGWGIISIALVFKPYLRRSEMVWRSVVTLARLYDIMFGVIVMAPVAVLSW
LPGLQEMQTRILFNEAFSRGLHISQIITGKKSHGV
>LOC_Os06g02260.1
MLTFFFTTVGYYVCTMMTVLTVYIFLYGRVYLALSGLDYEISRQFRFLGNTALDAALNAQFLVQIGIFTAVP
MIMGFILELGLLKAIFSFITMQLQFCSVFFTFSLGTRTHYFGRTILHGGAKYHATGRGFVVRHIKFAENYRLY
SRSHFVKALEVALLLIIYIAYGYTRGGSSSFILLTISSWFLVVSWLFAPYIFNPSGFEWQKTVEDFDDWTNW
LLYKGGVGVKGENSWESWWDEEQAHIQTLRGRILETILSLRFLIFQYGIVYKLKIASHNTSLAVYGFSWIVL
LVLVLLFKLFTATPKKSTALPTFVRFLQGLLAIGMIAGIALLIALTKFTIADLFASALAFVATGWCVLCLAVTWK
RLVKFVGLWDSVREIARMYDAGMGALIFVPIVFFSWFPFVSTFQSRFLFNQAFSRGLEISLILAGNKANQEA

**Fig. S22.** FASTA sequences of CalS (GSL) proteins of ALO used for the phylogenetic analysis.

**Genome assembly and evolutionary genomics of wild oats**

Qing Liu, Hongyu Yuan, Mingzhi Li, Ziwei Wang, Dongli Cui, Yushi Ye, Zongyi Sun, Xukai Tan, Trude Schwarzacher and John Seymour Heslop-Harrison

### Supplementary Tables

**Additional file 2: Table S1. Summary of sequencing libraries of *Avena* species included in the study. 3**

**Table S2. Deposited data of *Avena longiglumis* (ALO) and software used in the study. 6**

*a. Deposited data of genome assembly and annotation of ALO. 6*

*b. Software and algorithms. 6*

**Table S3. Summary of genome assembly and annotation of *Avena longiglumis* (ALO). 8**

*a. The statistics of ALO genome assembly. 8*

*b. Summary of Oxford Nanopore Technologies assembly metrics. 10*

**Table S4. Statistics of ALO genome assembly consistency. 11**

*a Statistics of illumina reads mapped to the ALO genome. 11*

*b Single base accuracy statistics of Illumina reads mapped to the ALO genome. 11*

*c Statistics of nanopore data alignment to the ALO genome. 11*

*d Statistics of nanopore data coverage of the ALO genome. 11*

**Table S5. Evaluation of gene space completeness in the ALO genome assembly. 12**

*a BUSCO analysis. 12*

*b CEGMA analysis. 12*

*c The Mapping rate of RNA-seq data to the ALO genome. 12*

**Table S6. Repetitive DNA composition comparison among genomes of ALO and six Poaceae species. 13**

*a Repetitive DNA composition of the ALO genome. 13*

*b Comparison of repetitive DNA composition of ALO and six Poaceae species genomes 14*

*c Genome percentage of top 10 abundant transposable element families in ALO and six Poaceae species genomes. 15*

**Table S7. Size and centromere localization of the ALO pseudomolecules. 16**

*a Centromere localization result from high abundance areas of repeat sequences 16*

*b Centromere localization result from centromeric retrotransposon Cereba 16*

*c Centromere localization from conserved regions between A. longiglumis and Oryza sativa genomes 16*

**Table S8. Gene characterization comparison between genomes of ALO and ten other plant species. 17**

**Table S9. Statistics of gene function annotation of the ALO genome. 18**

**Table S10. Statistics of annotated non-coding RNAs of the ALO genome. 19**

**Table S11. Identification of high-confidence (HC) and low-confidence (LC) protein-coding genes annotated in the ALO genome. 20**

*a HC genes supported by transcriptome data. 20*

*b HC genes supported by homology alignment results. 20*

*c Characteristics of HC and LC genes annotated in the ALO genome. 20*

**Table S12. Gene family categories in genomes of ALO and ten plant species. 21**

**Table S13. Gene family statistics of ALO and ten plant species genomes. 22**

**Table S14. Summary of orthologous gene clusters analyzed in the studied species. 23**

*a Expansion and contraction gene families in the studied species. 23*

*b Node number in the tree (Fig. 3a). 23*

**Table S15. GO enrichment analysis of expanded genes in the ALO genome. 24**

**Table S16. KEGG enrichment analysis of unique genes in the ALO genome. 34**

**Table S17. The gene pair statistics of SynVisio results between post-p ancestral grass karyotype (AGK) and grass species and between grass species. 35**

**Table S18. Statistics of up- and down-regulated DEGs in salt-treated roots and salt-treated leaves and expanded gene families of ALO. 36**

*a Gene number of up- and down-regulated DEGs in the analysed samples. 36*

*b Gene number of up- and down-regulated DEGs in expanded gene families of the analysed samples. 36*

*c Gene description of up- and down-regulated DEGs (≥5) in expanded gene families of the analysed samples. 36*
**Table S19. Characterization of biosynthetic gene clusters (BGCs) in the ALO genome. 68**
**Table S20. The *CYP450* gene copy number in the studied species and distribution along the ALO chromosomes. 72**
*a The copy number of CYP450 genes in the studied species. 72*
*b Distribution of CYP450 genes along the ALO chromosomes. 72*
**Table S21. FPKM value of *CYP450* genes in roots, leaves, salt-treated roots and salt-treated leaves of ALO. 73**
*a The transcript levels of CYP450 genes over 10-fold higher in salt-treated roots than in roots. 73*
*b The transcript levels of CYP450 genes over 10-fold higher in salt-treated leaves than in leaves. 74*
**Table S22. Description of 46 biosynthetic gene clusters (BGCs) containing 117 *CYP450* genes in the expression profiling heat map. 75**
**Table S23. The conserved synteny among *CYP450* genes within gene clusters in ALO and nine grass species. 88**
**Table S24. Beta-glucan encoding 11 *CesA* and 55 *Csl* genes corresponding to the expanded gene families in the ALO genome. 89**
**Table S25. Distribution of 11 *CesA* and 55 *Csl* genes in the ALO chromosomes. 92**
**Table S26. FPKM value of *CesA* and *Csl* genes in roots, leaves, salt-treated roots and salt-treated leaves of ALO. 93**
*a The transcript levels of CesA and Csl genes over 1-fold higher in salt-treated roots than in roots and FPKM value presence in salt-treated roots while zero in roots. 93*
*b The transcript levels of CesA and Csl genes higher in salt-treated roots than in roots. 93*
*c The transcript levels of CesA and Csl genes over 1-fold higher in salt-treated leaves than in leaves. 94*
*d The transcript levels of CesA and Csl genes higher in salt-treated leaves than in leaves. 94*
**Table S27. Gene description of 10 biosynthetic gene clusters (BGCs) containing *CesA* and *Csl* genes in the expression profiling heat map. 95**
**Table S28. The conserved synteny among *CesA* and *Csl* genes within gene clusters in ALO and nine grass species. 98**
**Table S29. Statistics of 13 *CalS* (*GSL*) genes corresponding the expanded gene families in the ALO genome. 99**
**Table S30. FPKM value of *CalS* genes in roots, leaves, salt-treated roots and salt-treated leaves of ALO. 100**
*a The transcript levels of CalS genes over 1-fold higher in salt-treated roots than in roots and FPKM value presence in salt-treated roots while zero in roots. 100*
*b The folder number of transcript levels of CalS genes higher in salt-treated leaves than in leaves. 100*

**Table S1. Summary of sequencing libraries of *Avena* species included in the study.**

| Species | Library | Type | Sample | Configuration | Insertion (bp) | Platform | Read count (Millions) | Total reads bases (bp) | Accession no. in Genome Sequence Archive or Genome Warehouse or NCBI |
|---|---|---|---|---|---|---|---|---|---|
| *A. longiglumis* | NDES00415_L4 | gDNA, Illumina | Leaf | 2 × 150bp, paired | 350 | Illumina HiSeq 2500 | 448.0 | 134,425,058,100 | PRJCA004488/SAMC344563/ |
| *A. longiglumis* | NDES00415_L5 | gDNA, Illumina | Leaf | 2 × 150bp, paired | 350 | Illumina HiSeq 2500 | 447.0 | 134,173,766,700 | CRR275326/CRR275326 |
| Subtotal | | | | | | | 895.0 | 268,598,824,800 | |
| *A. longiglumis* | 20190212-NPL0561-P1-E1-H1 | gDNA, Nanopore | Leaf | 20–30 kbp | > 23,000 | Nanopore PromethION | 3.2 | 48,220,684,593 | PRJCA004488/CRR275318 |
| *A. longiglumis* | 20190215-NPL0561-P1-E1-H1 | gDNA, Nanopore | Leaf | 20–30 kbp | > 23,000 | Nanopore PromethION | 2.7 | 51,028,783,483 | CRR275323 |
| *A. longiglumis* | 20190216-NPL0561-P1-A3-D3 | gDNA, Nanopore | Leaf | 20–30 kbp | > 23,000 | Nanopore PromethION | 1.2 | 21,004,088,941 | CRR275324 |
| *A. longiglumis* | 20190215-NPL0561-P1-A1-D1 | gDNA, Nanopore | Leaf | 20–30 kbp | > 23,000 | Nanopore PromethION | 4.6 | 84,191,503,177 | CRR275314 |
| *A. longiglumis* | 20190211-NPL0561-P1-E3-H3 | gDNA, Nanopore | Leaf | 20–30 kbp | > 23,000 | Nanopore PromethION | 3.6 | 49,715,012,344 | CRR275320 |
| *A. longiglumis* | 20190212-NPL0561-P2-A7-D7 | gDNA, Nanopore | Leaf | 20–30 kbp | > 23,000 | Nanopore PromethION | 3.7 | 56,313,401,650 | CRR275321 |
| *A. longiglumis* | 20190215-NPL0561-P1-A7-D7 | gDNA, Nanopore | Leaf | 20–30 kbp | > 23,000 | Nanopore PromethION | 1.4 | 24,947,902,936 | CRR275322 |
| *A. longiglumis* | 20190328-NPL0561-P2-E7-H7 | gDNA, Nanopore | Leaf | 20–30 kbp | > 23,000 | Nanopore PromethION | 2.1 | 38,735,683,870 | CRR275317 |
| *A. longiglumis* | 20190329-NPL0561-P1-E1-H1 | gDNA, Nanopore | Leaf | 20–30 kbp | > 23,000 | Nanopore PromethION | 2.4 | 45,977,659,069 | CRR275325 |
| *A. longiglumis* | 20190302-NPL0613-P1-A5-D5 | gDNA, Nanopore | Leaf | 20–30 kbp | > 23,000 | Nanopore PromethION | 2.5 | 38,218,681,407 | CRR275316 |
| *A. longiglumis* | 20190124-NPL0613-P2-A1-D1-ul | gDNA, Nanopore | Leaf | 20–30 kbp | > 23,000 | Nanopore PromethION | 2.1 | 22,158,343,522 | CRR275319 |

| | | | | | | | | | |
|---|---|---|---|---|---|---|---|---|---|
| *A. longiglumis* | 20190221-NPL0613-P2-E1-H1 | gDNA, Nanopore | Leaf | 20–30 kbp | > 23,000 | Nanopore PromethION | 1.7 | 30,848,177,162 | CRR275315 |
| Subtotal | | | | | | | 31.2 | 511,359,922,154.00 | |
| *A. longiglumis* | 447_18L3A026 | gDNA, HiC | Leaf | 2 × 150 bp, paired | 450 | Illumina Novaseq 6000 | 87.3 | 26,195,252,100 | PRJCA004488/CRR275304 |
| *A. longiglumis* | 447_19L3A027 | gDNA, HiC | Leaf | 2 × 150 bp, paired | 450 | Illumina Novaseq 6000 | 75.4 | 22,623,619,200 | CRR275310 |
| *A. longiglumis* | B447_5L4A005 | gDNA, HiC | Leaf | 2 × 150 bp, paired | 450 | Illumina Novaseq 6000 | 77.5 | 23,238,770,100 | CRR275307 |
| *A. longiglumis* | B447_6L4A006 | gDNA, HiC | Leaf | 2 × 150 bp, paired | 450 | Illumina Novaseq 6000 | 105.9 | 31,771,530,300 | CRR275308 |
| *A. longiglumis* | B447_16L4A024 | gDNA, HiC | Leaf | 2 × 150 bp, paired | 450 | Illumina Novaseq 6000 | 77.0 | 23,104,664,100 | CRR275306 |
| *A. longiglumis* | B447_18L3A026 | gDNA, HiC | Leaf | 2 × 150 bp, paired | 450 | Illumina Novaseq 6000 | 227.0 | 68,092,406,400 | CRR275309 |
| *A. longiglumis* | B447_19L3A027 | gDNA, HiC | Leaf | 2 × 150 bp, paired | 450 | Illumina Novaseq 6000 | 204.3 | 61,297,082,100 | CRR275305 |
| *A. longiglumis* | B447_5L3A005 | gDNA, HiC | Leaf | 2 × 150 bp, paired | 450 | Illumina Novaseq 6000 | 207.6 | 62,277,830,400 | CRR275312 |
| *A. longiglumis* | B447_6L3A006 | gDNA, HiC | Leaf | 2 × 150 bp, paired | 450 | Illumina Novaseq 6000 | 210.5 | 63,160,123,500 | CRR275313 |
| *A. longiglumis* | B447_16L3A024 | gDNA, HiC | Leaf | 2 × 150 bp, paired | 450 | Illumina Novaseq 6000 | 180.0 | 53,874,397,500 | CRR275311 |
| Subtotal | | | | | | | 1452.5 | 435,635,675,700 | |
| *A. longiglumis* | R1901811A-MJ | RNA-seq | Leaf | 2 × 150 bp, paired | 450 | Illumina Novaseq 6000 | 33.9 | 10,165,004,100 | CRR285670 |
| *A. longiglumis* | R1902606B-MJ | RNA-seq | Salt treated leaf | 2 × 150 bp, paired | 450 | Illumina Novaseq 6000 | 33.8 | 9,715,831,200 | CRR285672 |
| *A. longiglumis* | R1901812A-MJ | RNA-seq | Root | 2 × 150 bp, paired | 450 | Illumina Novaseq 6000 | 26.7 | 8,016,951,000 | CRR285671 |

| | | | | | | | | | |
|---|---|---|---|---|---|---|---|---|---|
| *A. longiglumis* | R1901790A-MJ | RNA-seq | Salt treated root | 2 × 150 bp, paired | 450 | Illumina Novaseq 6000 | 29.7 | 8,899,298,400 | CRR285673 |
| Subtotal | | | | | | | 124.1 | 36,797,084,700 | |
| *A. brevis* | Liu et al., 2019 | gDNA, Illumina | Leaf | 2 × 250 bp, paired | 300 | Illumina HiSeq 2500 | 60.4 | 30,183,435,500 | PRJNA407595/SRR6056491 |
| *A. hirtula* | Liu et al., 2019 | gDNA, Illumina | Leaf | 2 × 250 bp, paired | 300 | Illumina HiSeq 2500 | 61.5 | 30,738,036,000 | PRJNA407595/SRR6056492 |
| *A. sativa* | Liu et al., 2019 | gDNA, Illumina | Leaf | 2 × 250 bp, paired | 300 | Illumina HiSeq 2500 | 115.1 | 57,552,411,500 | PRJNA407595/SRR6056489 |
| *A. strigosa* | Liu et al., 2019 | gDNA, Illumina | Leaf | 2 × 250 bp, paired | 300 | Illumina HiSeq 2500 | 62.6 | 31,317,309,000 | PRJNA407595/SRR6056490 |
| *A. atlantica* | Liu et al., 2020 | gDNA, Illumina | Leaf | 2 × 250 bp, paired | 300 | Illumina HiSeq 2500 | 53.5 | 26,768,534,000 | GWHAOPC01000000 |
| *A. eriantha* | Liu et al., 2020 | gDNA, Illumina | Leaf | 2 × 250 bp, paired | 300 | Illumina HiSeq 2500 | 49.0 | 24,521,134,000 | GWHAOPE01000000 |
| *A. longiglumis* | Liu et al., 2020 | gDNA, Illumina | Leaf | 2 × 250 bp, paired | 300 | Illumina HiSeq 2500 | 48.6 | 24,297,359,000 | GWHAOPH01000000 |
| *A. murphyi* | Liu et al., 2020 | gDNA, Illumina | Leaf | 2 × 250 bp, paired | 300 | Illumina HiSeq 2500 | 46.0 | 22,981,249,000 | GWHAOPF01000000 |
| *A. nuda* | Liu et al., 2020 | gDNA, Illumina | Leaf | 2 × 250 bp, paired | 300 | Illumina HiSeq 2500 | 67.4 | 33,688,961,000 | GWHAOPD01000000 |
| *A. ventricosa* | Liu et al., 2020 | gDNA, Illumina | Leaf | 2 × 250 bp, paired | 300 | Illumina HiSeq 2500 | 28.5 | 14,267,461,000 | GWHAOPG01000000 |
| *A. wiestii* | Liu et al., 2020 | gDNA, Illumina | Leaf | 2 × 250 bp, paired | 300 | Illumina HiSeq 2500 | 25.4 | 12,685,761,500 | GWHAOPB01000000 |
| *A. hispanica* | GP-20191225-1698-15 | gDNA, Illumina | Leaf | 2 × 150 bp, paired | 300 | Illumina HiSeq 2500 | 72.8 | 21,829,345,000 | CRR285674 |
| Subtotal | | | | | | | 690.8 | 330,830,996,500 | |

**Table S2. Deposited data of *Avena longiglumis* (ALO) and software used in the study.**

**a. Deposited data of genome assembly and annotation of ALO.**

| Deposited data | | |
|---|---|---|
| Sequencing data for ALO | This study | National Genomic Data Center (https://bigd.big.ac.cn/bioproject/) under the accession no. PRJCA004488/CRR275304-CRR275326, CRR285670-285674, SRR605689-605692 |
| Genome assembly for ALO | This study | https://figshare.com/s/76392f11fc0c939302c7 or National Genomic Data Center (https://ngdc.cncb.ac.cn/search/?dbId=gsa&q=CRA004248) under the accession no. PRJCA003205/CRA004248 |

**b. Software and algorithms.**

| Software | Citation/Authors | Website |
|---|---|---|
| BUSCO v.4.0.5 | Simão et al., 2015 [29] | https://busco.ezlab.org/v3 |
| CEGMA v.2.5 | Parra et al., 2009 [30] | http://korflab.ucdavis.edu/datasets/cegma/#SCT8 |
| EDTA v.1.7.0 | Ou and Jiang, 2019 [36] | https://github.com/oushujun/EDTA |
| LTR_FINDER_parallel | Ou and Jiang, 2019 [36] | https://github.com/oushujun/LTR_FINDER_parallel |
| SynVisio | Bandi and Gutwin, 2020 [42] | https://synvisio.github.io/#/Dashboard/uploaded-source |
| eggNOG v.5.0 | Huerta-Cepas et al., 2019 [45] | http://eggnog5.embl.de/#/app/home |
| PlantTFDB v.5.0 | Tian et al., 2020 [48] | http://planttfdb.gao-lab.org/ |
| OrthoFinder v.2.3.14 | Emms and Kelly, 2019 [51] | https://github.com/davidemms/OrthoFinder |
| Jellyfish v.2.2.6 | Marçais and Kingsford, 2011 [99] | https://www.cbcb.umd.edu/software/jellyfish/ |
| GenomeScope | Vurture et al., 2017 [100] | http://qb.cshl.edu/genomescope/ |
| Guppy v.3.2.2 | Wick et al., 2019 [101] | https://github.com/nanoporetech/taiyaki |
| NextDenovo v.1.0 | Hu et al., 2020 [102] | https://github.com/Nextomics/NextDenovo |
| Wtdbg2.huge | Ruan and Li, 2020 [103] | https://github.com/ruanjue/wtdbg-2.huge |
| SMARTdenovo v.1.0.0 | Istace et al., 2017 [104] | https://github.com/ruanjue/smartdenovo |
| NextPolish v.1.01 | Hu et al., 2020 [102] | https://github.com/Nextomics/NextPolish |
| Fastp v.0.20.1 | Chen et al., 2018 [105] | https://github.com/OpenGene/fastp |
| Hi-C_Pro v.2.10.0 | Servant et al., 2015 [106] | https://github.com/nservant/HiC-Pro/releases |
| Bowtie2 v.2.3.2 | Langmead and Salzberg, 2012 [107] | https://github.com/BenLangmead/bowtie2 |
| Juicer v.1.5.7 | Durand et al., 2016 [108] | https://github.com/aidenlab/juicer |
| LACHESIS | Burton et al., 2013 [109] | https://github.com/shendurelab/LACHESIS |
| R v.3.6.3 | R core team, 2020 [111] | https://www.R-project.org/ |
| Kmerfreq_AR v.2.0.4 | Luo et al., 2012 [114] | https://github.com/fanagislab/kmerfreq |
| Minimap2 v.2.17 | Li, 2018 [115] | https://github.com/lh3/minimap2 |

| | | |
|---|---|---|
| Samtools | Li et al., 2009 [116] | http://samtools.sourceforge.net/ |
| LTRharvest | Ellinghaus et al., 2008 [118] | https://hajkd.github.io/LTRpred/reference/LTRharvest.html |
| LTR_retriever | Ou and Jiang, 2018 [117] | https://github.com/oushujun/LTR_retriever |
| Generic Repeat Finder | Shi and Liang, 2019 [119] | https://github.com/bioinfolabmu/GenericRepeatFinder |
| TIR-Learner | Su et al., 2016 [120] | https://github.com/WeijiaSu/TIR-Learner-Rice |
| HelitronScanner v.1.0 | Xiong et al., 2014 [121] | https://sourceforge.net/projects/helitronscanner/ |
| RepeatModeler v.2.0.2a | Flynn et al., 2020 [122] | http://www.repeatmasker.org/RepeatModeler/ |
| RepeatMasker v.4.1.1 | Smit et al, 2015 [123] | http://www.repeatmasker.org/ |
| TEsorter v.1.1.4 | Zhang et al., 2019 [124] | https://github.com/zhangrengang/TEsorter |
| Geneious Primer v.2021.1.1 | Biomatters Ltd. | https://www.geneious.com/ |
| Augustus v.3.3.2 | Stanke et al., 2006 [128] | http://bioinf.uni-greifswald.de/augustus/ |
| GeMoMa v.1.6.1 | Keilwagen et al., 2019 [129] | http://www.jstacs.de/index.php/GeMoMa |
| TransDecoder v.5.5.0 | Haas et al., 2013 [130] | https://github.com/TransDecoder/TransDecoder |
| EVidenceModeler v.1.1.1 | Haas et al., 2008 [131] | https://evidencemodeler.github.io/ |
| BLAST2GO v.1.44 | Conesa et al., 2005 [133] | https://www.blast2go.com/ |
| Rfam library v.11.0 | Griffiths-Jones et al., 2005 [134] | http://rfam.xfam.org/ |
| tRNAscan-SE v.2.0 | Chan and Lowe, 2019 [135] | http://lowelab.ucsc.edu/tRNAscan-SE/ tRNAscan-SE |
| miRanda v.3.0 | Betel et al., 2008 [136] | http://www.microrna.org |
| RNAmmer v.1.2 | Lagesen et al., 2007 [137] | https://services.healthtech.dtu.dk/service.php?RNAmmer-1.2 |
| STAR aligner | Dobin et al., 2013 [138] | http://code.google.com/p/rna-star/ |
| StringTie v.2.0.6 | Pertea et al., 2015 [139] | https://ccb.jhu.edu/software/stringtie/ |
| Phytozome v.13 | David et al., 2012 [140] | https://phytozome.jgi.doe.gov/pz/portal.html |
| Diamond v.0.9.24 | Buchfink et al., 2015 [141] | https://github.com/bbuchfink/diamond/releases |
| UpsetR v.1.4.0 | Conway et al., 2017 [142] | https://github.com/hms-dbmi/UpSetR/ |
| MAFFT v.7.48 | Katoh and Standley, 2013 [143] | https://mafft.cbrc.jp/alignment/software/ |
| RAxML v.8.1.17 | Stamatakis, 2014 [144] | https://cme.h-its.org/exelixis/web/software/ raxml/index.html |
| PAML v.4.9i | Yang, 2007 [145] | http://abacus.gene.ucl.ac.uk/software/paml.html |
| CAFE v.4.2 | De Bie et al., 2006 [146] | https://github.com/hahnlab/CAFE |
| TBtools v.1.092 | Chen et al., 2020 [147] | https://github.com/CJ-Chen/TBtools/releases |
| MCScanX | Wang et al., 2012 [148] | https://github.com/wyp1125/MCScanX |
| WGDI v.0.4.7 | Sun et al., 2021 [150] | https://github.com/SunPengChuan/wgdi |
| KaKs_calculator v.2.0 | Wang et al., 2010 [151] | https://bigd.big.ac.cn/tools/kaks |
| DESEQ2 v.3.11 | Love et al., 2014 [153] | https://bioconductor.org/packages/release/bioc/html/ DESeq2.html |
| plantiSMASH v.1.3 | Kautsar et al., 2017 [154] | http://plantismash.secondarymetabolites.org/help.html |
| FastTree v.2.1.10 | Price et al., 2010 [157] | http://www.microbesonline.org/fasttree/ |

**Table S3. Summary of genome assembly and annotation of *Avena longiglumis* (ALO).**

**a. The statistics of ALO genome assembly.**

| Genome features | Measures of index |
|---|---|
| Genome survey | |
| Illumina sequences | 267.54 Gbp (67.55×) |
| Genome survey contigs | 4,040,426 |
| *K*-mer | 17 |
| *K*-mer number | 208,165 |
| *K*-mer depth | 49.80 × |
| Estimated genome size | 3,965,670,000 bp |
| Heterozygous ratio (%) | 0.48 |
| Repeat (%) | 82.08 |
| Genome assembly | |
| Nanopore sequences | 252.78 Gbp (63.82×) |
| Nanopore contig number | 2,379 |
| Nanopore contig number correction | 2,381 |
| Assembled genome size | 3,960,768,570 bp |
| Assembly % of genome | 99.88% |
| Longest contigs | 99,445,397 bp |
| Contig N50 length | 12,682,464 bp |
| Contig N90 length | 662,566 bp |
| Contig L50 count | 74 |
| Contig L90 count | 806 |
| Number of scaffolds | 414 |
| Total size of scaffolds | 3,960,965,270 bp |
| Scaffold N50 length | 527,343,613 bp |
| Scaffold N90 length | 453,691,697 bp |
| Scaffold N50 count | 5 |
| Scaffold N90 count | 9 |
| Longest scaffold | 594,546,470 bp |
| GC content | 44.67% |
| Hi-C auxiliary assembly | |
| Library data | 435.64 Gb |
| Raw Paired-end reads | 2,904,215,238 |
| Clean Paired-end reads | 2,870,596,248 |
| Clean bases | 420,725,014,170 bp |
| Clean Q30 bases | 394,303,483,280 (99.55×) |
| Clean paired-end read rate | 98.84% |
| Clean Q30 base rate | 93.72% |
| Hi-C contig number | 2381 |

| | |
|---|---|
| Longest contig length | 99,445,397 |
| Contig N50 length | 12,682,464 |
| Contig N90 length | 661,603 |
| Total size of contigs | 3,960,768,570 |
| Gap length | 196,700 |
| Total size of scaffolds | 3,960,965,270 |
| Total length of pseudo-chromosomes (% assembly) | 3,847,578,604 (97.14%) |
| Hi-C scaffold number | 414 |
| Longest scaffold length | 594,546,470 |
| Scaffold N50 length | 527,343,613 |
| Scaffold N90 length | 6,968,329 |
| Gene models | |
| Number of predicted genes | 40,845 |
| Number of annotated genes | 39,491 |
| Number of HC genes (Proportion of genes) | 33,271 (81.46%) |
| Number of LC genes (Proportion of genes) | 7,574 (18.54%) |
| Mean gene density | 10 / Mbp |
| Total length of genes (bp) | 133,632,290 |
| Number of genes | 40,845 |
| Mean gene length (bp) | 3,271.69 |
| Total length of exons (bp) | 50,244,372 |
| Number of exons | 187,479 |
| Mean exon length (bp) | 268 |
| Mean exon number | 4.59 |
| Total length of HC genes (bp) | 115,042,270 |
| Total length of LC genes (bp) | 18,590,020 |
| Number of HC genes (Proportion of genes) | 33,271 (81.46%) |
| Number of LC genes (Proportion of genes) | 7,574 (18.54%) |
| Mean HC gene length (bp) | 3457.73 |
| Mean LC gene length (bp) | 2454.45 |
| Number of annotated genes | 39,491 |
| Number of mRNA | 15,446 |
| Mean exon number per HC gene | 5.02 |
| Mean exon length of HC genes | 1,306 bp |
| Mean intron length of HC genes | 2,152 bp |
| Nonprotein coding genes/elements | |
| Number of miRNA genes | 15,446 |
| Mean length of miRNA genes | 136.29 bp |
| miRNA genes share in genome | 0.0532% |
| Number of rRNA genes | 257 |
| Mean length of rRNA genes | 901.50 bp |
| rRNA genes share in genome | 0.0058% |
| Number of tRNA genes | 1,402 |

| | |
|---|---|
| Mean length of tRNA genes | 75.62 bp |
| tRNA gene share in genome | 0.0027% |
| Number of snRNA genes | 607 |
| Mean length of snRNA genes | 101.92 bp |
| snRNA genes share in genome | 0.0016% |
| Total repetitive DNAs | 3,479,237,283 bp |
| Repeat region % of assembly | 87.84% |
| Total transposable elements | 3,411,928,666 bp |
| Percentage of transposable element in genome | 86.14% |

**b. Summary of Oxford Nanopore Technologies assembly metrics.**

| | Contigs (after polish) | Scaffold (Nanodrop + Hi-C) |
|---|---|---|
| Total length | 3,960,768,570 bp | 3,960,965,270 bp |
| Number of contigs | 2,379 | 414 |
| Maximum contig length | 99,445,397 bp | 594,546,470 bp |
| N50 | 12,682,464 bp | 527,343,613 bp |
| N60 | 7,675,489 bp | 513,337,126 bp |
| N70 | 3,127,915 bp | 453,691,697 bp |
| N80 | 1,419,177 bp | 9,709,002 bp |
| N90 | 662,566 bp | 6,968,329 bp |

**Table S4. Statistics of ALO genome assembly consistency.**

**a Statistics of illumina reads mapped to the ALO genome.**

| Illumina sequences | Total reads | Map reads | Mapped ratio | Paired reads | Paired map reads | Properly paired reads | Properly map ratio |
|---|---|---|---|---|---|---|---|
| 267,537,826,727 (68×) | 1,349,168,075 | 1,348,307,165 | 99.94% | 1,346,023,478 | 1,345,121,368 | 1,304,341,354 | 96.90% |

**b Single base accuracy statistics of Illumina reads mapped to the ALO genome.**

| Type | Depth ≥ 1 × | Depth ≥ 5 × | Depth ≥ 10 × |
|---|---|---|---|
| Heterozygous SNPs | 3,914,721 | 3,895,495 | 3,830,731 |
| Heterozygous indels (bp) | 185,546 | 183,477 | 177,108 |
| Homozygous SNPs | 100,257 | 66,016 | 51,054 |
| Homozygous indels (bp) | 40,166 | 16,544 | 8,074 |
| Heterozygous SNP ratio | 0.0988% | 0.0984% | 0.0967% |
| Heterozygous indel ratio | 0.0047% | 0.0046% | 0.0045% |
| Homozygous SNP ratio | 0.0025% | 0.0017% | 0.0013% |
| Homozygous indel ratio | 0.0010% | 0.0004% | 0.0002% |
| Heterozygous variant ratio | 0.1035% | 0.1030% | 0.1012% |
| Homozygous variant ratio | 0.0035% | 0.0021% | 0.0015% |
| Genome accuracy ratio | 99.9965% | 99.9979% | 99.9985% |

**c Statistics of nanopore data alignment to the ALO genome.**

| Total reads | Mapped reads | Map ratio |
|---|---|---|
| 15,768,837 | 15,753,851 | 99.90% |

**d Statistics of nanopore data coverage of the ALO genome.**

| Depth (×) | Base number | Coverage ratio (%) |
|---|---|---|
| 1 | 3,960,157,317 | 99.98 |
| 5 | 3,956,158,830 | 99.88 |
| 10 | 3,950,188,188 | 99.73 |
| 20 | 3,925,074,004 | 99.10 |
| Nanopore coverage depth (×) | 74.02 × | |

**Table S5. Evaluation of gene space completeness in the ALO genome assembly.**

**a BUSCO analysis.**

| | Based on whole genome sequences | Based on annotated genome |
|---|---|---|
| Complete BUSCOs (C) | 1,348 (98.04%) | 1,311 (95.35%) |
| Complete and single-copy BUSCOs (S) | 1,125 (81.82%) | 1,114 (81.02%) |
| Complete and duplicated BUSCOs (D) | 223 (16.22%) | 197 (14.33%) |
| Fragmented BUSCOs (F) | 3 (0.22%) | 16 (1.16%) |
| Missing BUSCOs (M) | 24 (1.74%) | 48 (3.49%) |
| Total BUSCO groups searched | 1,375 | 1,375 |

**b CEGMA analysis.**

| Type | Complete | | Complete+ partial* | |
|---|---|---|---|---|
| | Prots | % Completeness | Prots | % Completeness |
| Group 1 | 60 | 90.91 | 65 | 98.48 |
| Group 2 | 48 | 85.71 | 52 | 92.86 |
| Group 3 | 57 | 93.44 | 60 | 98.36 |
| Group 4 | 64 | 98.46 | 64 | 98.46 |
| Total | 229 | 92.34 | 241 | 97.18 |

*The length of the gene assembled is less than 70%.

**c The Mapping rate of RNA-seq data to the ALO genome.**

| Samples | Libraries | Total bases | Q30 rate (%) | Mapping rate (%) | Multiple mapping rate (%) | Unique mapping rate (%) |
|---|---|---|---|---|---|---|
| Root | R1901812A-MJ | 8,016,951,000 | 93.98 | 97.24 | 6.72 | 90.53 |
| Salt-treated root | R1901790A-MJ | 8,899,298,400 | 93.88 | 74.26 | 5.45 | 68.79 |
| Leaf | R1901811A-MJ | 10,165,004,100 | 95.90 | 97.66 | 11.23 | 86.44 |
| Salt-treated leaf | R1902606B-MJ | 9,715,831,200 | 92.78 | 93.07 | 8.11 | 84.96 |

**Table S6. Repetitive DNA composition comparison among genomes of ALO and six Poaceae species.**

**a Repetitive DNA composition of the ALO genome.**

| Repeat type | | Super family | Family | Repeat sequences (bp) | Copy number | Repeat fraction | Genome fraction |
|---|---|---|---|---|---|---|---|
| Transposable elements | | | | | | | |
| Class I (Retrotransposons) | | | | | | | |
| LTR | | *Gypsy* | | 2,045,839,268 | 1,127,011 | 59.34% | 51.65% |
| | | *Copia* | | 1,035,647,971 | 575,382 | 30.04% | 26.15% |
| | | Unknown LTR | | 77,372,748 | 61,272 | 2.24% | 1.95% |
| | | Other LTR | | 213,586 | 628 | 0.01% | 0.01% |
| | Total LTR- Retrotransposons | | | 3,159,073,573 | 1,764,293 | 91.63% | 79.76% |
| Non-LTR | | LINE | L1 | 38,994,208 | 42,780 | 1.13% | 0.98% |
| | Total Class I retrotransposons | | | 3,198,067,781 | 1,807,073 | 92.76% | 80.74% |
| Class II (DNA transposons)-Subclass 1 | | | | | | | |
| | | Tc1_Mariner | | 21,858,891 | 69,956 | 0.63% | 0.55% |
| | | CACTA | | 29,566,991 | 75,699 | 0.86% | 0.75% |
| | | Mutator | | 25,388,681 | 82,292 | 0.74% | 0.64% |
| | | PIF_Harbinger | | 9,194,639 | 31,402 | 0.27% | 0.23% |
| | | hAT | | 7,600,354 | 22,266 | 0.22% | 0.19% |
| Class II (DNA transposons)-Subclass II | | | | | | | |
| | | Helitron | | 43,779,456 | 117,248 | 1.27% | 1.11% |
| | Total Class II DNA transposons | | | 137,389,012 | 398,863 | 3.99% | 3.47% |
| | Total transposable elements | | | 3,335,456,793 | 2,205,936 | 96.75% | 84.21% |
| | Tandem and simple sequence repeats | | | 11,144,119 | 162,888 | 0.32% | 0.28% |
| | Other repeats | | | 100,883,895 | 369,975 | 2.93% | 2.55% |
| | Total repetitive DNAs | | | 3,447,484,807 | 2,728,799 | 100% | 87.04% |

**b Comparison of repetitive DNA composition of ALO and six Poaceae species genomes**

| Repeat type | Super family | Family | *Avena longiglumis* (ALO) | *Aegilops tauschii* (ATA) | *Brachy-podium distachyon* (BDI) | *Oryza sativa* (OSA) | *Sorghum bicolor* (SBI) | *Setaria italica* (SIT) | *Zea mays* (ZMA) |
|---|---|---|---|---|---|---|---|---|---|
| LTR | *Gypsy* | *Retand* | 9.06% | 9.07% | 7.70% | 3.88% | 6.22% | 5.43% | 13.18% |
| | | *Ogre* | 1.62% | 0.07% | 0.00% | 1.18% | 5.27% | 3.45% | 0.82% |
| | | *Athila* | 8.10% | 3.48% | 0.00% | 0.45% | 7.27% | 0.63% | 0.00% |
| | | *Tekay* | 17.33% | 2.25% | 1.05% | 1.98% | 3.11% | 2.81% | 6.38% |
| | | *CRM* | 1.92% | 0.77% | 0.84% | 0.55% | 1.06% | 0.95% | 0.80% |
| | | *Reina* | 0.10% | 0.18% | 0.00% | 0.30% | 0.40% | 0.18% | 0.37% |
| | | *Galadriel* | 0.00% | 0.07% | 0.00% | 0.00% | 0.00% | 0.00% | 0.00% |
| | | Unclassified *Gypsy* | 13.51% | 13.33% | 3.32% | 7.41% | 13.50% | 5.86% | 17.67% |
| | | Total *Gypsy* | 51.65% | 29.70% | 12.91% | 15.75% | 37.53% | 19.31% | 39.22% |
| | *Copia* | *SIRE* | 1.40% | 2.00% | 0.75% | 0.75% | 2.35% | 0.42% | 5.06% |
| | | *Angela* | 17.39% | 5.33% | 0.48% | 0.07% | 0.07% | 2.11% | 0.25% |
| | | *Bianca* | 0.13% | 0.07% | 0.39% | 0.22% | 0.23% | 0.09% | 0.13% |
| | | *Ivana* | 0.12% | 0.07% | 0.40% | 0.22% | 0.37% | 0.41% | 0.15% |
| | | *TAR* | 0.50% | 0.20% | 0.30% | 0.53% | 0.22% | 0.20% | 0.41% |
| | | *Ale* | 0.60% | 0.63% | 0.41% | 0.30% | 0.60% | 0.80% | 0.51% |
| | | *Tork* | 0.08% | 0.04% | 0.18% | 0.17% | 0.04% | 0.10% | 0.02% |
| | | *Ikeros* | 0.15% | 0.07% | 0.18% | 0.18% | 0.13% | 0.23% | 0.06% |
| | | *Alesia* | 0.00% | 0.00% | 0.07% | 0.00% | 0.00% | 0.01% | 0.00% |
| | | Unclassified *Copia* | 5.77% | 5.25% | 0.30% | 0.34% | 1.81% | 1.44% | 4.89% |
| | | Total *Copia* | 26.15% | 13.66% | 3.46% | 2.78% | 5.82% | 5.81% | 11.48% |
| | *PARA-RT* | | 0.00% | 0.00% | 0.00% | 0.00% | 0.00% | 0.00% | 0.00% |
| | *Cassandra* | | 0.00% | 0.00% | 0.00% | 0.00% | 0.00% | 0.00% | 0.00% |
| | *Caulimoviridae* | | 0.00% | 0.00% | 0.06% | 0.00% | 0.00% | 0.00% | 0.00% |
| | Unclassified LTR | | 1.95% | 12.62% | 0.68% | 2.27% | 0.72% | 9.25% | 8.89% |

| Repeat type | Super family | Family | *Avena longiglumis* (ALO) | *Aegilops tauschii* (ATA) | *Brachy-podium distachyon* (BDI) | *Oryza sativa* (OSA) | *Sorghum bicolor* (SBI) | *Setaria italica* (SIT) | *Zea mays* (ZMA) |
|---|---|---|---|---|---|---|---|---|---|
| | Non-*Gypsy* and non-*Copia* | | 0.01% | 8.92% | 0.34% | 0.27% | 0.93% | 0.52% | 1.44% |
| Total LTR- Retrotransposons | | | 79.76% | 64.90% | 17.45% | 21.07% | 53.12% | 26.77% | 61.03% |
| Non-LTR | *LINE* | L1 | 0.98% | 0.00% | 0.00% | 0.02% | 0.08% | 0.08% | 0.09% |
| Total Class I retrotransposons | | | 80.74% | 64.90% | 17.45% | 21.09% | 53.20% | 26.89% | 61.12% |
| | *Tc1_Mariner* | | 0.55% | 2.95% | 3.96% | 7.01% | 1.73% | 1.66% | 0.30% |
| | *CACTA* | | 0.75% | 2.07% | 2.01% | 1.66% | 2.61% | 2.61% | 4.59% |
| | *Mutator* | | 0.64% | 4.06% | 1.55% | 3.58% | 1.16% | 1.16% | 8.76% |
| | *PIF_Harbinger* | | 0.23% | 1.19% | 0.85% | 1.18% | 2.55% | 2.55% | 0.74% |
| | *hAT* | | 0.19% | 0.22% | 0.38% | 1.01% | 0.61% | 0.61% | 0.99% |
| | *Helitron* | | 1.11% | 6.32% | 9.55% | 12.03% | 6.04% | 6.04% | 4.84% |
| Total DNA Class II transposons | | | 3.47% | 16.81% | 18.3% | 26.47% | 16.59% | 16.59% | 20.22% |
| Total Transposable elements | | | 84.21% | 81.71% | 35.75% | 47.56% | 67.83% | 43.48% | 81.34% |
| Other repeats | | | 2.83% | 0.01% | 0.00% | 0.31% | 0.52% | 1.76% | 1.69% |
| Total Repetitive DNA | | | 87.04% | 81.72% | 35.75% | 47.84% | 68.35% | 45.24% | 83.03% |

**c Genome percentage of top 10 abundant transposable element families in ALO and six Poaceae species genomes.**

| Top 10 abundant TE Families | *Avena longiglumis* (ALO) | *Aegilops tauschii* (ATA) | *Brachypodium distachyon* (BDI) | *Oryza sativa* (OSA) | *Sorghum bicolor* (SBI) | *Setaria italica* (SIT) | *Zea mays* (ZMA) |
|---|---|---|---|---|---|---|---|
| *Angela* | 17.39% | 5.33% | 0.48% | 0.07% | 2.11% | 0.07% | 0.25% |
| *Tekay* | 17.33% | 2.25% | 1.05% | 1.98% | 2.81% | 3.11% | 6.38% |
| *Retand* | 9.06% | 9.07% | 7.70% | 3.88% | 5.43% | 6.22% | 13.18% |
| *Athila* | 8.10% | 3.48% | 0.00% | 0.45% | 0.63% | 7.27% | 0.00% |
| *CRM* | 1.92% | 0.77% | 0.84% | 0.55% | 0.95% | 1.06% | 0.80% |
| *Ogre* | 1.62% | 0.07% | 0.00% | 1.18% | 3.45% | 5.27% | 0.82% |
| *SIRE* | 1.40% | 2.00% | 0.75% | 0.75% | 0.42% | 2.35% | 5.06% |
| *Helitron* | 1.11% | 6.32% | 9.55% | 12.03% | 6.95% | 6.04% | 4.84% |
| *LINE* | 0.98% | 0.00% | 0.00% | 0.02% | 0.12% | 0.08% | 0.09% |
| *CACTA* | 0.75% | 2.07% | 2.01% | 1.66% | 3.55% | 2.61% | 4.59% |

**Table S7. Size and centromere localization of the ALO pseudomolecules.**

**a Centromere localization result from high abundance areas of repeat sequences** on chromosome dotplots Using Geneious Primer v.2021.1.1.

| Chromosomes | Size (bp) | Suspected centromere | Start (bp) | End (bp) | Median (bp) |
|---|---|---|---|---|---|
| ALO01 | 594,546,470 | centormere1 | 315,061,249 | 324,147,201 | **319,604,225** |
| ALO02 | 587,543,788 | centormere1 | 323,004,417 | 335,723,521 | 329,363,969 |
| ALO03 | 587,190,583 | centormere1 | 280,098,446 | 290,268,161 | 285,183,304 |
| ALO04 | 583,925,327 | centormere1 | 336,345,261 | 345,614,337 | 340,979,799 |
| ALO05 | 527,343,613 | centormere1 | 305,957,889 | 318,402,561 | **312,180,225** |
| ALO06 | 513,337,126 | centormere1 | 245,726,549 | 250,604,796 | 248,165,673 |
| ALO06 | 513,337,126 | centormere2 | 252,557,313 | 253,143,041 | 252,850,177 |
| ALO07 | 453,691,697 | centormere1 | 288,095,587 | 297,553,921 | 292,824,754 |

**b Centromere localization result from centromeric retrotransposon *Cereba*** (KM948610 [40] Tomás et al., 2016) mapping result on ALO chromosomes by Geneious Primer v.2021.1.1.

| Chromosomes | Size (bp) | Start (bp) | End (bp) | Median (bp) |
|---|---|---|---|---|
| ALO01 | 594,546,470 | 302,554,936 | 358,293,671 | 330,424,304 |
| ALO02 | 587,543,788 | 269,301,057 | 360,462,979 | 314,882,018 |
| ALO03 | 587,190583 | 227,581,439 | 307,289,165 | 267,435,302 |
| ALO04 | 583,925,327 | 230,548,605 | 361,076,206 | **295,812,406** |
| ALO05 | 527,343,613 | 283,077,215 | 345,987,804 | 314,532,510 |
| ALO06 | 513,337,126 | 189,778,061 | 296,954,346 | **243,366,204** |
| ALO07 | 453,691,697 | 252,676,290 | 312,676,170 | 282,676,230 |

**c Centromere localization from conserved regions between *A. longiglumis* and *Oryza sativa* genomes** using SynVisio [41] result

| Chromosomes | Start alignment no. | End alignment no. | Start (bp) | End (bp) | Median (bp) |
|---|---|---|---|---|---|
| ALO01 | 73-139 | 41-3 | 218,336,850 | 456,530,974 | 337,433,912 |
| ALO02 | 148-8 | 186-8 | 218,434,180 | 372,761,714 | 295,597,947 |
| ALO03 | 303-3 | 297-67 | 215,567,169 | 380,159,203 | 297,863,186 |
| ALO04 | 417-428 | 341-2 | 209,204,572 | 325,956,657 | 267,580,615 |
| ALO05 | 506-1 | 485-14 | 214,260,721 | 370,096,114 | 292,178,418 |
| ALO06 | 564-6 | 585-11 | 213,285,659 | 282,988,269 | 248,136,964 |
| ALO07 | 687-39 | 663-0 | 262,540,264 | 403,854,365 | **333,197,315** |

**Table S8. Gene characterization comparison between genomes of ALO and ten other plant species.**

| Species | Total number of genes | Average gene length (bp) | Average CDS length (bp) | Average exon number per gene | Average exon length (bp) | Average intron length (bp) |
|---|---|---|---|---|---|---|
| AAT | 49,542 | 3115.68 | 1109.41 | 4.08 | 271.61 | 648.76 |
| AER | 47,361 | 3273.99 | 1040.83 | 4.11 | 253.26 | 715.46 |
| ALO | 40,845 | 3,271.69 | 1,231.98 | 4.59 | 268.33 | 567.96 |
| AST | 39,885 | 3720.99 | 1266.74 | 4.49 | 282.04 | 648.66 |
| ATA | 38,772 | 8095.90 | 1143.23 | 3.22 | 289.92 | 659.35 |
| ATH | 27,416 | 2203.02 | 1230.62 | 5.57 | 220.87 | 158.89 |
| BDI | 32,439 | 3353.90 | 1525.09 | 6.40 | 238.19 | 381.56 |
| OSA | 42,189 | 2871.77 | 1121.41 | 4.57 | 245.40 | 386.68 |
| SBI | 34,129 | 3713.16 | 1227.98 | 5.09 | 241.13 | 480.14 |
| SIT | 34,584 | 3180.36 | 1249.13 | 5.07 | 246.18 | 374.80 |
| ZMA | 39,498 | 4173.20 | 1447.53 | 7.03 | 205.83 | 572.53 |

**Table S9. Statistics of gene function annotation of the ALO genome.**

| Datebases (Abbreviation; Website) | Number | Percent (%) |
|---|---|---|
| NCBI non-redundant protein (NR; ftp://ftp.ncbi.nih.gov/blast/db) | 39,390 | 96.44% |
| NOG (Non-supervised Orthologous Groups) | 38,078 | 93.23% |
| Pfam (http://pfam.xfam.org/) | 35,573 | 87.09% |
| COG (Clusters of Orthologous Groups of proteins) | 35,460 | 86.82% |
| SwissProt (http://www.uniprot.org/uniprot/?query=*&fil=reviewed%3Ayes) | 32,099 | 78.59% |
| Gene ontology (GO; http://geneontology.org/) | 21,162 | 51.81% |
| Eukaryotic orthology groups (KOG; http://www.genome.jp/kaas-bin/kaas_main) | 20,756 | 50.82% |
| Kyoto Encyclopedia of Genes and Genomes (KEGG; http://muchong.com/html/201009/2325769.html) | 17,285 | 42.32% |
| PlantTFDB (http://planttfdb.gao-lab.org/) | 2,403 | 5.88% |
| CAZy (Carbohydrate-Active enZYmes) | 931 | 2.28% |
| Number of annotated genes | 39,558 | 96.85% |
| Number of predicted genes | 40,845 | 100% |

**Table S10. Statistics of annotated non-coding RNAs of the ALO genome.**

| Type | | Copy number | Mean length (bp) | Total length (bp) | Percentage of genome |
|---|---|---|---|---|---|
| miRNA | | 15,446 | 136.29 | 2,105,135 | 0.053150% |
| tRNA | | 1,402 | 75.61 | 106,012 | 0.002676% |
| | 18S | 43 | 1,859.86 | 79,974 | 0.002019% |
| | 28S | 33 | 3,957.88 | 130,610 | 0.003298% |
| | 5.8S | 2 | 151.00 | 302 | 0.000008% |
| | 5S | 179 | 116.20 | 20,799 | 0.000525% |
| rRNA | | 257 | 901.50 | 231,685 | 0.005850% |
| | CD-box & HACA-box, etc | 406 | 77.69 | 31,544 | 0.000796% |
| | Splicing | 201 | 150.84 | 30,319 | 0.000765% |
| snRNA | | 607 | 101.92 | 61,865 | 0.001562% |
| Other | | 185 | 135.03 | 24,981 | 0.000631% |
| ncRNA | | 16,439 | 135.19 | 2,222,342 | 0.056111% |
| Regulatory | | 165 | 39.52 | 6,521 | 0.000165% |

**Table S11. Identification of high-confidence (HC) and low-confidence (LC) protein-coding genes annotated in the ALO genome.**

**a HC genes supported by transcriptome data.**

| Sample | FPKM > 0 | Percent |
|---|---|---|
| Roots | 29,924 | 73.26% |
| Leaves | 24,600 | 60.23% |
| Salt-treated roots | 29,172 | 71.42% |
| Salt-treated leaves | 25,864 | 63.32% |
| Total expressed genes (HC) | 32,188 | 78.81% |

**b HC genes supported by homology alignment results.**

| Species | Gene number |
|---|---|
| *Triticum aestivum* | 940 |
| *Hordeum vulgare* | 690 |
| Oat A-genome | 1,612 |
| Oat C-genome | 919 |
| Total HC genes | 1,083 |

**c Characteristics of HC and LC genes annotated in the ALO genome.**

| Metric | HC | LC |
|---|---|---|
| Number of genes | 33,271 | 7,574 |
| Number of single-exon genes | 8,397 | 3,236 |
| Number of multi-exon genes | 24,874 | 4,338 |
| Mean gene length (bp) | 3457.73 | 2454.45 |
| Mean CDS length (bp) | 1,306.4 | 905.2 |
| Max CDS length (bp) | 16,176 | 5184 |
| Min CDS length (bp) | 150 | 163 |
| Mean exons per genes (no.) | 5.04 | 2.68 |

**Table S12. Gene family categories in genomes of ALO and ten plant species.**

| Species (Abbreviation) | Single-copy orthologs | Multiple-copy orthologs | Unique orthologs | Other orthologs | Unclustered genes | Genome size (Mbp)* |
|---|---|---|---|---|---|---|
| *Aegilops tauschii* (ATA) | 1,880 | 12,684 | 2,110 | 19,804 | 2,294 | 4,360 |
| *Arabidopsis thaliana* (ATH) | 1,880 | 13,123 | 5,965 | 3,453 | 2,995 | 125 |
| *Avena atlantica* (AAT) | 1,880 | 12,995 | 2,973 | 26,515 | 5,179 | 3,690 |
| *Avena eriantha* (AER) | 1,880 | 12,818 | 2,541 | 24,943 | 5,179 | 3,780 |
| *Avena longiglumis* (ALO) | 1,880 | 15,329 | 869 | 21,210 | 1,557 | 3,960 |
| *Avena strigose* (AST) | 1,880 | 12,570 | 1,012 | 22,112 | 2,311 | 4,100 |
| *Brachypodium distachyon* (BDI) | 1,880 | 11,716 | 840 | 13,815 | 4,188 | 260 |
| *Oryza sativa* (OSA) | 1,880 | 12,304 | 5,268 | 16,203 | 6,534 | 389 |
| *Sorghum bicolor* (SBI) | 1,880 | 12,235 | 1,495 | 14,070 | 4,449 | 730 |
| *Setaria italica* (SIT) | 1,880 | 12,615 | 1,391 | 15,390 | 3,308 | 423 |
| *Zea mays* (ZMA) | 1,880 | 15,414 | 5,625 | 14,075 | 2,504 | 2,300 |

* Genome size from literature: ATA [32], ATH [50], AAT and AER [12], AST [7], BDI [33], OSA [9], SBI [10], SIT [11] and ZMA [34].

**Table S13. Gene family statistics of ALO and ten plant species genomes.**

| Species (Abbreviation) | Gene number | Genes in families | Unclustered genes | Family number | Unique family number | Average gene number per family |
|---|---|---|---|---|---|---|
| *Aegilops tauschii* (ATA) | 38,772 | 36,478 | 2,294 | 19,247 | 556 | 1.90 |
| *Arabidopsis thaliana* (ATH) | 27,416 | 24,421 | 2,995 | 13,022 | 1,329 | 1.88 |
| *Avena atlantica* (AAT) | 49,542 | 44,363 | 5,179 | 22,813 | 845 | 1.94 |
| *Avena eriantha* (AER) | 47,361 | 42,182 | 5,179 | 21,750 | 818 | 1.94 |
| *Avena longiglumis* (ALO) | 40,845 | 39,288 | 1,557 | 19,954 | 259 | 1.97 |
| *Avena strigose* (AST) | 39,885 | 37,574 | 2,311 | 21,620 | 355 | 1.74 |
| *Brachypodium distachyon* (BDI) | 32,439 | 28,251 | 4,188 | 18,743 | 300 | 1.51 |
| *Oryza sativa* (OSA) | 42,189 | 35,655 | 6,534 | 19,870 | 1,351 | 1.79 |
| *Sorghum bicolor* (SBI) | 34,129 | 29,680 | 4,449 | 19,266 | 486 | 1.54 |
| *Setaria italica* (SIT) | 34,584 | 31,276 | 3,308 | 19,370 | 392 | 1.61 |
| *Zea mays* (ZMA) | 39,498 | 36,994 | 2,504 | 18,948 | 1,336 | 1.95 |

**Table S14. Summary of orthologous gene clusters analyzed in the studied species.**

**a Expansion and contraction gene families in the studied species.**

| Species name <node number> | Expansions | | Contractions | |
|---|---|---|---|---|
| | Families | Genes | Families | Genes |
| *Aegilops tauschii* <12> | 2,625 | 6,809 | 7,328 | 7,656 |
| *Arabidopsis thaliana* <0> | 779 | 1,126 | 16,619 | 18,396 |
| *Avena atlantica* <18> | 1,896 | 6,951 | 2,477 | 2,985 |
| *Avena eriantha* <14> | 2,421 | 7,514 | 4,187 | 4,379 |
| *Avena longiglumis* <16> | 4,329 | 7,236 | 4,942 | 6,327 |
| *Avena strigosa* <20> | 1,219 | 3,277 | 3,416 | 4,306 |
| *Brachypodium distachyon* <10> | 1,380 | 2,291 | 7,767 | 8,257 |
| *Oryza sativa* <8> | 2,302 | 4,591 | 8,264 | 8,525 |
| *Sorghum bicolor* <6> | 1,471 | 2,410 | 1,648 | 1,768 |
| *Setaria italica* <2> | 1,936 | 3,595 | 1,794 | 1,842 |
| *Zea mays* <4> | 4,774 | 7,700 | 2,954 | 3,868 |
| <3> | 1,002 | 1,214 | 6,676 | 6,774 |
| <5> | 492 | 587 | 1,163 | 1,332 |
| <7> | 484 | 837 | 2,441 | 2,770 |
| <9> | 512 | 813 | 1,139 | 1,159 |
| <11> | 769 | 1,230 | 1,127 | 1,213 |
| <13> | 1,047 | 1,965 | 566 | 580 |
| <15> | 1,440 | 2,649 | 962 | 1,096 |
| <17> | 1,395 | 2,003 | 729 | 913 |
| <19> | 491 | 849 | 869 | 961 |

**b Node number in the tree (Fig. 3a).**

CAFE labeled tree:

(ATH<0>,((SIT<2>,(ZMA<4>,SBI<6>)<5>)<3>,(OSA<8>,(BDI<10>,(ATA<12>,(AER<14>,(ALO<16>,(AAT<18>,AST<20>)<19>)<17>)<15>)<13>)<11>)<9>)<7>)<1>

**Table S15. GO enrichment analysis of expanded genes in the ALO genome.**

| GO.ID | Class | Term | Gene number | P value |
|---|---|---|---|---|
| GO:0016838 | Molecular function | carbon-oxygen lyase activity, acting on phosphates | 102 | 1.11E-16 |
| GO:0010333 | Molecular function | terpene synthase activity | 102 | 8.88E-16 |
| GO:0016747 | Molecular function | acyltransferase activity, transferring groups other than amino-acyl groups | 249 | 2.07E-13 |
| GO:0045735 | Molecular function | nutrient reservoir activity | 118 | 3.49E-13 |
| GO:0000287 | Molecular function | magnesium ion binding | 109 | 2.75E-11 |
| GO:0016835 | Molecular function | carbon-oxygen lyase activity | 109 | 9.94E-11 |
| GO:0016746 | Molecular function | acyltransferase activity | 263 | 1.56E-08 |
| GO:0020037 | Molecular function | heme binding | 489 | 1.81E-08 |
| GO:0046906 | Molecular function | tetrapyrrole binding | 489 | 1.81E-08 |
| GO:0016705 | Molecular function | oxidoreductase activity, acting on paired donors, with incorporation or reduction of molecular oxygen | 389 | 2.65E-08 |
| GO:0005506 | Molecular function | iron ion binding | 375 | 6.52E-08 |
| GO:0030246 | Molecular function | carbohydrate binding | 192 | 6.88E-08 |
| GO:0016829 | Molecular function | lyase activity | 186 | 1.13E-07 |
| GO:0016782 | Molecular function | transferase activity, transferring sulphur-containing groups | 53 | 1.34E-07 |
| GO:0008146 | Molecular function | sulfotransferase activity | 53 | 1.34E-07 |
| GO:0046872 | Molecular function | metal ion binding | 1003 | 4.21E-07 |
| GO:0043169 | Molecular function | cation binding | 1005 | 6.43E-07 |
| GO:0004867 | Molecular function | serine-type endopeptidase inhibitor activity | 48 | 1.84E-06 |
| GO:0030247 | Molecular function | polysaccharide binding | 116 | 6.26E-06 |
| GO:0016843 | Molecular | amine-lyase activity | 31 | 9.97E-06 |

| GO.ID | Class | Term | Gene number | P value |
|---|---|---|---|---|
| | function | | | |
| GO:0016844 | Molecular function | strictosidine synthase activity | 31 | 9.97E-06 |
| GO:0043531 | Molecular function | ADP binding | 323 | 2.03E-05 |
| GO:0016411 | Molecular function | acylglycerol O-acyltransferase activity | 21 | 6.42E-05 |
| GO:0004144 | Molecular function | diacylglycerol O-acyltransferase activity | 21 | 6.42E-05 |
| GO:0016840 | Molecular function | carbon-nitrogen lyase activity | 31 | 1.29E-04 |
| GO:0016491 | Molecular function | oxidoreductase activity | 1033 | 3.04E-04 |
| GO:0004857 | Molecular function | enzyme inhibitor activity | 66 | 4.58E-04 |
| GO:0004497 | Molecular function | monooxygenase activity | 53 | 8.24E-04 |
| GO:0008171 | Molecular function | O-methyltransferase activity | 46 | 8.40E-04 |
| GO:0016798 | Molecular function | hydrolase activity, acting on glycosyl bonds | 309 | 0.001226592 |
| GO:0140678 | Molecular function | molecular function inhibitor activity | 66 | 0.001296517 |
| GO:0060590 | Molecular function | ATPase regulator activity | 8 | 0.001617784 |
| GO:0000774 | Molecular function | adenyl-nucleotide exchange factor activity | 8 | 0.001617784 |
| GO:0042802 | Molecular function | identical protein binding | 8 | 0.001617784 |
| GO:0042803 | Molecular function | protein homodimerization activity | 8 | 0.001617784 |
| GO:0004499 | Molecular function | N,N-dimethylaniline monooxygenase activity | 38 | 0.001866047 |
| GO:0004553 | Molecular function | hydrolase activity, hydrolyzing O-glycosyl compounds | 284 | 0.001913315 |
| GO:0003690 | Molecular function | double-stranded DNA binding | 62 | 0.002263156 |
| GO:0030414 | Molecular function | peptidase inhibitor activity | 64 | 0.002352077 |
| GO:0061134 | Molecular function | peptidase regulator activity | 64 | 0.002352077 |
| GO:0061135 | Molecular function | endopeptidase regulator activity | 64 | 0.002352077 |

| GO.ID | Class | Term | Gene number | P value |
|---|---|---|---|---|
| GO:0004866 | Molecular function | endopeptidase inhibitor activity | 64 | 0.002352077 |
| GO:0016709 | Molecular function | oxidoreductase activity, acting on paired donors, with incorporation or reduction of molecular oxygen, NAD(P)H as one donor, and incorporation of one atom of oxygen | 38 | 0.002907005 |
| GO:0016641 | Molecular function | oxidoreductase activity, acting on the CH-NH2 group of donors, oxygen as acceptor | 7 | 0.003613307 |
| GO:0048038 | Molecular function | quinone binding | 7 | 0.003613307 |
| GO:0008131 | Molecular function | primary amine oxidase activity | 7 | 0.003613307 |
| GO:0004672 | Molecular function | protein kinase activity | 936 | 0.004055744 |
| GO:0016684 | Molecular function | oxidoreductase activity, acting on peroxide as acceptor | 155 | 0.004255612 |
| GO:0050661 | Molecular function | NADP binding | 49 | 0.004495565 |
| GO:0016209 | Molecular function | antioxidant activity | 158 | 0.004864995 |
| GO:0004601 | Molecular function | peroxidase activity | 153 | 0.005131033 |
| GO:0008061 | Molecular function | chitin binding | 22 | 0.005349513 |
| GO:0008308 | Molecular function | voltage-gated anion channel activity | 12 | 0.006028075 |
| GO:0005244 | Molecular function | voltage-gated ion channel activity | 12 | 0.006028075 |
| GO:0005247 | Molecular function | voltage-gated chloride channel activity | 12 | 0.006028075 |
| GO:0005253 | Molecular function | anion channel activity | 12 | 0.006028075 |
| GO:0005254 | Molecular function | chloride channel activity | 12 | 0.006028075 |
| GO:0022832 | Molecular function | voltage-gated channel activity | 12 | 0.006028075 |
| GO:0046914 | Molecular function | transition metal ion binding | 618 | 0.006474902 |
| GO:0030410 | Molecular function | nicotianamine synthase activity | 14 | 0.010381395 |
| GO:0016638 | Molecular | oxidoreductase activity, acting on the | 9 | 0.014267205 |

| GO.ID | Class | Term | Gene number | P value |
|---|---|---|---|---|
| | function | CH-NH2 group of donors | | |
| GO:0030976 | Molecular function | thiamine pyrophosphate binding | 9 | 0.014267205 |
| GO:0050997 | Molecular function | quaternary ammonium group binding | 9 | 0.014267205 |
| GO:0061733 | Molecular function | peptide-lysine-N-acetyltransferase activity | 12 | 0.014284422 |
| GO:0004402 | Molecular function | histone acetyltransferase activity | 12 | 0.014284422 |
| GO:0034212 | Molecular function | peptide N-acetyltransferase activity | 12 | 0.014284422 |
| GO:0050662 | Molecular function | obsolete coenzyme binding | 50 | 0.015380677 |
| GO:0008312 | Molecular function | 7S RNA binding | 13 | 0.017459427 |
| GO:0004345 | Molecular function | glucose-6-phosphate dehydrogenase activity | 7 | 0.017581963 |
| GO:0016435 | Molecular function | rRNA (guanine) methyltransferase activity | 5 | 0.018021383 |
| GO:0004014 | Molecular function | adenosylmethionine decarboxylase activity | 5 | 0.018021383 |
| GO:0098772 | Molecular function | molecular function regulator | 164 | 0.019435697 |
| GO:0016701 | Molecular function | oxidoreductase activity, acting on single donors with incorporation of molecular oxygen | 47 | 0.019872967 |
| GO:0005509 | Molecular function | calcium ion binding | 139 | 0.02365855 |
| GO:0016899 | Molecular function | oxidoreductase activity, acting on the CH-OH group of donors, oxygen as acceptor | 11 | 0.024469466 |
| GO:0015018 | Molecular function | galactosylgalactosylxylosylprotein 3-beta-glucuronosyltransferase activity | 11 | 0.024469466 |
| GO:0015020 | Molecular function | glucuronosyltransferase activity | 11 | 0.024469466 |
| GO:0003885 | Molecular function | D-arabinono-1,4-lactone oxidase activity | 11 | 0.024469466 |
| GO:0140102 | Molecular function | catalytic activity, acting on a rRNA | 23 | 0.025152301 |
| GO:0030234 | Molecular function | enzyme regulator activity | 154 | 0.025260809 |
| GO:0140096 | Molecular function | catalytic activity, acting on a protein | 1370 | 0.026354552 |

| GO.ID | Class | Term | Gene number | P value |
|---|---|---|---|---|
| GO:0010277 | Molecular function | chlorophyllide a oxygenase [overall] activity | 8 | 0.026530341 |
| GO:0016740 | Molecular function | transferase activity | 1887 | 0.027592357 |
| GO:0008080 | Molecular function | N-acetyltransferase activity | 12 | 0.028876938 |
| GO:0015035 | Molecular function | protein-disulfide reductase activity | 33 | 0.031866481 |
| GO:0015036 | Molecular function | disulfide oxidoreductase activity | 33 | 0.031866481 |
| GO:0004564 | Molecular function | beta-fructofuranosidase activity | 9 | 0.034401174 |
| GO:0004575 | Molecular function | sucrose alpha-glucosidase activity | 9 | 0.034401174 |
| GO:0090599 | Molecular function | alpha-glucosidase activity | 9 | 0.034401174 |
| GO:0015926 | Molecular function | glucosidase activity | 9 | 0.034401174 |
| GO:0004519 | Molecular function | endonuclease activity | 83 | 0.035754152 |
| GO:0008374 | Molecular function | O-acyltransferase activity | 25 | 0.03899017 |
| GO:0071949 | Molecular function | FAD binding | 16 | 0.041937051 |
| GO:0051213 | Molecular function | dioxygenase activity | 39 | 0.043054205 |
| GO:0016702 | Molecular function | oxidoreductase activity, acting on single donors with incorporation of molecular oxygen, incorporation of two atoms of oxygen | 39 | 0.043054205 |
| GO:0043167 | Molecular function | ion binding | 2497 | 0.047676929 |
| GO:0000786 | Cellular component | nucleosome | 78 | 7.66E-12 |
| GO:0044815 | Cellular component | DNA packaging complex | 78 | 1.57E-11 |
| GO:0032993 | Cellular component | protein-DNA complex | 78 | 1.57E-11 |
| GO:0000785 | Cellular component | chromatin | 78 | 3.15E-11 |
| GO:0005694 | Cellular component | chromosome | 92 | 6.48E-08 |
| GO:0070449 | Cellular | elongin complex | 8 | 4.48E-04 |

| GO.ID | Class | Term | Gene number | P value |
|---|---|---|---|---|
| | component | | | |
| GO:0005759 | Cellular component | mitochondrial matrix | 20 | 7.38E-04 |
| GO:0071944 | Cellular component | cell periphery | 73 | 0.003212858 |
| GO:0048500 | Cellular component | signal recognition particle | 13 | 0.003522842 |
| GO:0016459 | Cellular component | myosin complex | 12 | 0.006811914 |
| GO:0005956 | Cellular component | protein kinase CK2 complex | 5 | 0.008098839 |
| GO:0005938 | Cellular component | cell cortex | 32 | 0.011632134 |
| GO:0000145 | Cellular component | exocyst | 32 | 0.011632134 |
| GO:0070013 | Cellular component | intracellular organelle lumen | 45 | 0.013949672 |
| GO:0043233 | Cellular component | organelle lumen | 45 | 0.013949672 |
| GO:0031974 | Cellular component | membrane-enclosed lumen | 45 | 0.013949672 |
| GO:0005634 | Cellular component | nucleus | 263 | 0.021232121 |
| GO:0008023 | Cellular component | transcription elongation factor complex | 8 | 0.021614056 |
| GO:0099023 | Cellular component | vesicle tethering complex | 34 | 0.03025653 |
| GO:0000151 | Cellular component | ubiquitin ligase complex | 8 | 0.043339985 |
| GO:0043228 | Cellular component | non-membrane-bounded organelle | 209 | 0.045813137 |
| GO:0043232 | Cellular component | intracellular non-membrane-bounded organelle | 209 | 0.045813137 |
| GO:0042221 | Biological process | response to chemical | 133 | 0 |
| GO:0009719 | Biological process | response to endogenous stimulus | 120 | 0 |
| GO:0009725 | Biological process | response to hormone | 120 | 0 |
| GO:0009733 | Biological process | response to auxin | 96 | 0 |
| GO:0010033 | Biological process | response to organic substance | 120 | 0 |

| GO.ID | Class | Term | Gene number | P value |
|---|---|---|---|---|
| GO:0050896 | Biological process | response to stimulus | 498 | 1.11E-15 |
| GO:0000160 | Biological process | phosphorelay signal transduction system | 69 | 4.17E-09 |
| GO:0055114 | Biological process | obsolete oxidation-reduction process | 944 | 1.77E-07 |
| GO:0009611 | Biological process | response to wounding | 39 | 4.67E-07 |
| GO:0006468 | Biological process | protein phosphorylation | 934 | 2.70E-06 |
| GO:0080134 | Biological process | regulation of response to stress | 15 | 3.15E-06 |
| GO:0031347 | Biological process | regulation of defense response | 13 | 1.71E-05 |
| GO:0002682 | Biological process | regulation of immune system process | 13 | 1.71E-05 |
| GO:0002831 | Biological process | regulation of response to biotic stimulus | 13 | 1.71E-05 |
| GO:0032101 | Biological process | regulation of response to external stimulus | 13 | 1.71E-05 |
| GO:0009627 | Biological process | systemic acquired resistance | 13 | 1.71E-05 |
| GO:0050776 | Biological process | regulation of immune response | 13 | 1.71E-05 |
| GO:0010112 | Biological process | regulation of systemic acquired resistance | 13 | 1.71E-05 |
| GO:0035556 | Biological process | intracellular signal transduction | 78 | 2.06E-05 |
| GO:0009309 | Biological process | amine biosynthetic process | 23 | 2.54E-05 |
| GO:0042401 | Biological process | cellular biogenic amine biosynthetic process | 23 | 2.54E-05 |
| GO:0016310 | Biological process | phosphorylation | 943 | 3.87E-05 |
| GO:0006950 | Biological process | response to stress | 258 | 1.33E-04 |
| GO:0006464 | Biological process | cellular protein modification process | 1090 | 2.39E-04 |
| GO:0036211 | Biological process | protein modification process | 1090 | 2.39E-04 |
| GO:0006979 | Biological process | response to oxidative stress | 144 | 4.37E-04 |
| GO:0006595 | Biological | polyamine metabolic process | 9 | 5.01E-04 |

| GO.ID | Class | Term | Gene number | P value |
|---|---|---|---|---|
| | process | | | |
| GO:0006596 | Biological process | polyamine biosynthetic process | 9 | 5.01E-04 |
| GO:0043412 | Biological process | macromolecule modification | 1104 | 6.39E-04 |
| GO:0001101 | Biological process | response to acid chemical | 13 | 0.002092 |
| GO:0009415 | Biological process | response to water | 13 | 0.002092 |
| GO:0019538 | Biological process | protein metabolic process | 1576 | 0.003130113 |
| GO:0006508 | Biological process | proteolysis | 355 | 0.003789224 |
| GO:0006821 | Biological process | chloride transport | 12 | 0.004007357 |
| GO:0043207 | Biological process | response to external biotic stimulus | 16 | 0.004633848 |
| GO:0098542 | Biological process | defense response to other organism | 16 | 0.004633848 |
| GO:0002376 | Biological process | immune system process | 16 | 0.004633848 |
| GO:0051707 | Biological process | response to other organism | 16 | 0.004633848 |
| GO:0006955 | Biological process | immune response | 16 | 0.004633848 |
| GO:0044419 | Biological process | biological process involved in interspecies interaction between organisms | 16 | 0.004633848 |
| GO:0009607 | Biological process | response to biotic stimulus | 16 | 0.004633848 |
| GO:0032501 | Biological process | multicellular organismal process | 90 | 0.00546708 |
| GO:0072351 | Biological process | tricarboxylic acid biosynthetic process | 14 | 0.006725844 |
| GO:0072350 | Biological process | tricarboxylic acid metabolic process | 14 | 0.006725844 |
| GO:0030417 | Biological process | nicotianamine metabolic process | 14 | 0.006725844 |
| GO:0030418 | Biological process | nicotianamine biosynthetic process | 14 | 0.006725844 |
| GO:0045017 | Biological process | glycerolipid biosynthetic process | 32 | 0.008016394 |
| GO:0048583 | Biological | regulation of response to stimulus | 21 | 0.00821559 |

| GO.ID | Class | Term | Gene number | P value |
|---|---|---|---|---|
| | process | | | |
| GO:0007275 | Biological process | multicellular organism development | 42 | 0.008578517 |
| GO:0048856 | Biological process | anatomical structure development | 42 | 0.008578517 |
| GO:0032502 | Biological process | developmental process | 42 | 0.008578517 |
| GO:0007154 | Biological process | cell communication | 157 | 0.01040742 |
| GO:1901700 | Biological process | response to oxygen-containing compound | 13 | 0.011767453 |
| GO:0006006 | Biological process | glucose metabolic process | 14 | 0.013585462 |
| GO:0008215 | Biological process | spermine metabolic process | 5 | 0.014698742 |
| GO:0008216 | Biological process | spermidine metabolic process | 5 | 0.014698742 |
| GO:0008295 | Biological process | spermidine biosynthetic process | 5 | 0.014698742 |
| GO:0000154 | Biological process | rRNA modification | 5 | 0.014698742 |
| GO:0031167 | Biological process | rRNA methylation | 5 | 0.014698742 |
| GO:0006597 | Biological process | spermine biosynthetic process | 5 | 0.014698742 |
| GO:0070475 | Biological process | rRNA base methylation | 5 | 0.014698742 |
| GO:0070476 | Biological process | rRNA (guanine-N7)-methylation | 5 | 0.014698742 |
| GO:0036265 | Biological process | RNA (guanine-N7)-methylation | 5 | 0.014698742 |
| GO:0006612 | Biological process | protein targeting to membrane | 15 | 0.015200561 |
| GO:0006613 | Biological process | cotranslational protein targeting to membrane | 15 | 0.015200561 |
| GO:0006614 | Biological process | SRP-dependent cotranslational protein targeting to membrane | 15 | 0.015200561 |
| GO:0072599 | Biological process | establishment of protein localization to endoplasmic reticulum | 15 | 0.015200561 |
| GO:0045047 | Biological process | protein targeting to ER | 15 | 0.015200561 |
| GO:0006308 | Biological process | DNA catabolic process | 12 | 0.020240916 |

| GO.ID | Class | Term | Gene number | P value |
|---|---|---|---|---|
| GO:0006952 | Biological process | defense response | 50 | 0.023334809 |
| GO:0090150 | Biological process | establishment of protein localization to membrane | 15 | 0.02650164 |
| GO:0051668 | Biological process | localization within membrane | 15 | 0.02650164 |
| GO:0072657 | Biological process | protein localization to membrane | 15 | 0.02650164 |
| GO:0009605 | Biological process | response to external stimulus | 16 | 0.02794268 |
| GO:0023052 | Biological process | signaling | 109 | 0.028551197 |
| GO:0007165 | Biological process | signal transduction | 109 | 0.028551197 |
| GO:0042398 | Biological process | cellular modified amino acid biosynthetic process | 11 | 0.03415849 |
| GO:0000003 | Biological process | reproduction | 60 | 0.035565538 |
| GO:0022414 | Biological process | reproductive process | 60 | 0.035565538 |
| GO:0019319 | Biological process | hexose biosynthetic process | 7 | 0.038278903 |
| GO:0046364 | Biological process | monosaccharide biosynthetic process | 7 | 0.038278903 |
| GO:0006094 | Biological process | gluconeogenesis | 7 | 0.038278903 |
| GO:0009628 | Biological process | response to abiotic stimulus | 13 | 0.039601111 |
| GO:0006793 | Biological process | phosphorus metabolic process | 989 | 0.047310002 |
| GO:0006796 | Biological process | phosphate-containing compound metabolic process | 989 | 0.047310002 |
| GO:0032940 | Biological process | secretion by cell | 38 | 0.047643842 |
| GO:0006887 | Biological process | exocytosis | 38 | 0.047643842 |
| GO:0140352 | Biological process | export from cell | 38 | 0.047643842 |
| GO:0046903 | Biological process | secretion | 38 | 0.047643842 |

**Table S16. KEGG enrichment analysis of unique genes in the ALO genome.**

| Term name | Main class | Gene number | Rich factor | Corrected p value (Q value) |
|---|---|---|---|---|
| B 09102 Energy metabolism | A09100 Metabolism | 22 | 2.52 | 0.0023017 |
| 00592 alpha-linolenic acid metabolism | A09100 Metabolism | 10 | 4.84 | 0.0028007 |
| 00630 Glyoxylate and dicarboxylate metabolism | A09100 Metabolism | 10 | 4.61 | 0.0028417 |
| 00780 Biotin metabolism | A09100 Metabolism | 6 | 9.88 | 0.0036274 |
| 00190 Oxidative phosphorylation | A09100 Metabolism | 12 | 3.58 | 0.0037667 |
| 00966 Glucosinolate biosynthesis | A09100 Metabolism | 5 | 6.70 | 0.0160531 |
| 00020 Citrate cycle (TCA cycle) | A09100 Metabolism | 6 | 5.16 | 0.0175378 |
| 04145 Phagosome | A09140 Cellular Processes | 8 | 2.90 | 0.0794589 |
| 00710 Carbon fixation in photosynthetic organisms | A09100 Metabolism | 6 | 3.17 | 0.1200674 |
| B 09101 Carbohydrate metabolism | A09100 Metabolism | 32 | 1.46 | 0.1712943 |
| 02010 ABC transporters | A09130 Environmental Information Processing | 5 | 2.85 | 0.1964384 |
| B 09131 Membrane transport | A09130 Environmental Information Processing | 5 | 2.85 | 0.1964384 |
| 00970 Aminoacyl-tRNA biosynthesis | A09120 Genetic Information Processing | 5 | 2.69 | 0.2039866 |
| 04120 Ubiquitin mediated proteolysis | A09120 Genetic Information Processing | 12 | 1.80 | 0.2116835 |

For KEGG results of unique genes in AAT, AST and AER genomes can be requested from authors.

**Table S17. The gene pair statistics of SynVisio results between post-ρ ancestral grass karyotype (AGK) and grass species and between grass species.**

| Sample 1 | Sample 2 | Number of genome blocks | Number of gene pairs | Number of gene pairs between samples 1_1 | Number of gene pairs between samples 1_2 | Number of gene pairs between samples 2_2 |
|---|---|---|---|---|---|---|
| AGK and grass species | | | | | | |
| AGK | AAT | 505 | 14,400 | 1,013 | 11,579 | 1,808 |
| AGK | AER | 438 | 14,402 | 1,016 | 11,541 | 1,845 |
| AGK | ALO | 732 | 16,670 | 990 | 12,362 | 3,,318 |
| AGK | AST | 380 | 12,533 | 1,004 | 11,790 | 1,957 |
| AGK | ATA | 490 | 12,995 | 1,015 | 10,473 | 1,507 |
| AGK | BDI | 595 | 15,497 | 1,035 | 12,539 | 1,923 |
| AGK | OSA | 487 | 18,690 | 2,007 | 13,797 | 2,886 |
| Grass species AST-ALO-AER | | | | | | |
| ALO | AST | 670 | 29030 | 0 | 29,030 | 0 |
| ALO | AER | 824 | 27,116 | 0 | 27,116 | 0 |
| AER | AST | 172 | 21,536 | 0 | 21,536 | 0 |
| Grass species OSA_ALO_BDI | | | | | | |
| ALO | OSA | 697 | 19,178 | 0 | 19,178 | 0 |
| ALO | BDI | 778 | 22,004 | 0 | 22,004 | 0 |
| BDI | OSA | 582 | 20,460 | 0 | 20,460 | 0 |

**Table S18. Statistics of up- and down-regulated DEGs in salt-treated roots and salt-treated leaves and expanded gene families of ALO.**

**a Gene number of up- and down-regulated DEGs in the analysed samples.**

| Pairs | Up | Down | Total |
|---|---|---|---|
| Roots versus salt-treated roots | 1,569 | 1,627 | 3,196 |
| Leaves versus salt-treated leaves | 1,507 | 2,336 | 3,843 |
| Total | 3,076 | 3,963 | 7,139 |

**b Gene number of up- and down-regulated DEGs in expanded gene families of the analysed samples.**

| Regualtion | Roots | Leaves | All tissues |
|---|---|---|---|
| up-regulated DEGs | 599 | 451 | - |
| down-regulated DEGs | 753 | 642 | - |
| Total | 1,352 | 1,093 | 2,445 |

**c Gene description of up- and down-regulated DEGs (≥5) in expanded gene families of the analysed samples.**

| Gene ID | Description | KEGG_ko | PFAMs |
|---|---|---|---|
| 283 up-regulation DEGs (gene number ≥ 5 for each category) in expanded gene families of salt-treated roots of ALO | | | |
| AL3G05791 | Protein kinase | ko:K07198 | NAF,Pkinase |
| AL3G05815 | Protein kinase | ko:K07198 | NAF,Pkinase |
| AL6G05277 | Protein kinase | - | LRRNT_2,LRR_1,LRR_4,LRR_6,LRR_8 |
| AL2G05940 | Protein kinase | - | LRR_8,Malectin_like,Pkinase,Pkinase_Tyr |
| AL3G01132 | Protein kinase | - | EGF_CA,GUB_WAK_bind,Pkinase,Pkinase_Tyr |
| AL5G01007 | Protein kinase | ko:K04733 | Pkinase,Pkinase_Tyr |
| AL5G01009 | Protein kinase | ko:K04733 | Pkinase,Pkinase_Tyr |
| AL3G05281 | Protein kinase | ko:K04733 | Pkinase_Tyr |
| AL3G05293 | Protein kinase | ko:K04733 | Pkinase_Tyr |
| AL4G02186 | Protein kinase | - | Pkinase,Pkinase_Tyr |
| AL6G04702 | Protein kinase | - | Pkinase,Stress-antifung |
| AL3G05717 | Protein kinase | - | Pkinase_Tyr |
| AL3G05762 | Protein kinase | - | Pkinase_Tyr |
| AL4G00222 | Protein kinase | ko:K04733 | Pkinase,Pkinase_Tyr |
| AL2G01531 | Protein kinase | ko:K20603 | Pkinase |
| AL6G01333 | Protein kinase | - | LRRNT_2,LRR_1,LRR_6,LRR_8,Pkinase,Pkinase_Tyr |
| AL7G01113 | Protein kinase | - | LRRNT_2,LRR_1,LRR_6,LRR_8,Pkinase,Pkinase_Tyr |
| AL1G05289 | Protein kinase | - | LRRNT_2,LRR_1,LRR_8,Pkinase |
| AL4G01949 | Protein kinase | - | LRRNT_2,LRR_1,LRR_8,Pkinase |
| AL4G01959 | Protein kinase | - | LRRNT_2,LRR_1,LRR_8,Pkinase |
| AL7G01110 | Protein kinase | - | LRRNT_2,LRR_1,LRR_8,Pkinase |
| AL7G01134 | Protein kinase | - | LRRNT_2,LRR_1,LRR_8,Pkinase |
| AL6G01323 | Protein kinase | - | LRRNT_2,LRR_1,LRR_8,Pkinase,Pkinase_Tyr |
| AL6G01334 | Protein kinase | - | LRRNT_2,LRR_1,LRR_8,Pkinase,Pkinase_Tyr |

| Gene ID | Description | KEGG_ko | PFAMs |
|---|---|---|---|
| AL6G01336 | Protein kinase | - | LRRNT_2,LRR_1,LRR_8,Pkinase,Pkinase_Tyr |
| AL1G05907 | Protein kinase | - | DUF3403,Pkinase,Pkinase_Tyr,Stress-antifung |
| AL7G01104 | Protein kinase | - | Pkinase,Pkinase_Tyr |
| AL7G01105 | Protein kinase | - | Pkinase,Pkinase_Tyr |
| AL7G01143 | Protein kinase | - | Pkinase,Pkinase_Tyr |
| AL4G00475 | Protein kinase | - | B_lectin,PAN_2,Pkinase,Pkinase_Tyr,S_locus_glycop |
| AL4G00501 | Protein kinase | - | B_lectin,PAN_2,Pkinase,Pkinase_Tyr,S_locus_glycop |
| AL1G06007 | Protein kinase | - | GUB_WAK_bind,Pkinase,Pkinase_Tyr,WAK_assoc |
| AL1G06053 | Protein kinase | - | GUB_WAK_bind,Pkinase,Pkinase_Tyr,WAK_assoc |
| AL2G02169 | Protein kinase | - | Pkinase,Pkinase_Tyr,Stress-antifung |
| AL2G02291 | Protein kinase | - | Pkinase,Pkinase_Tyr,Stress-antifung |
| AL1G05798 | Protein kinase | ko:K04733 | Pkinase_Tyr |
| AL1G05815 | Protein kinase | ko:K04733 | Pkinase_Tyr |
| AL4G00502 | Protein kinase | ko:K04733 | Pkinase_Tyr |
| AL1G02505 | Protein kinase | - | B_lectin,DUF3403,PAN_2,Pkinase_Tyr,S_locus_glycop |
| AL1G06045 | Protein kinase | ko:K04733 | B_lectin,PAN_2,Pkinase,S_locus_glycop |
| AL2G04024 | Cytochrome P450 | - | p450 |
| AL4G00045 | Cytochrome P450 | - | p450 |
| AL4G00094 | Cytochrome P450 | - | p450 |
| AL4G01951 | Cytochrome P450 | - | p450 |
| AL6G01845 | Cytochrome P450 | - | p450 |
| AL6G01857 | Cytochrome P450 | - | p450 |
| AL7G00655 | Cytochrome P450 | - | p450 |
| AL7G00656 | Cytochrome P450 | - | p450 |
| AL1G05990 | Cytochrome P450 | ko:K00512 | p450 |
| AL1G05991 | Cytochrome P450 | ko:K00512 | p450 |
| AL1G05992 | Cytochrome P450 | ko:K00512 | p450 |
| AL1G05994 | Cytochrome P450 | ko:K00512 | p450 |
| AL1G06067 | Cytochrome P450 | ko:K00512 | p450 |
| AL1G06068 | Cytochrome P450 | ko:K00512 | p450 |
| AL1G06069 | Cytochrome P450 | ko:K00512 | p450 |
| AL1G06071 | Cytochrome P450 | ko:K00512 | p450 |
| AL7G02557 | Cytochrome P450 | ko:K00512 | p450 |
| AL7G02558 | Cytochrome P450 | ko:K00512 | p450 |
| AL7G04259 | Cytochrome P450 | ko:K00512 | p450 |
| AL7G05074 | Cytochrome P450 | ko:K00512 | p450 |
| AL4G00930 | Cytochrome P450 | ko:K04122,ko:K21719 | p450 |
| AL1G06294 | Cytochrome P450 | ko:K07418 | p450 |
| AL6G05230 | Cytochrome P450 | ko:K07426 | p450 |
| AL4G02594 | Cytochrome P450 | ko:K15639 | p450 |
| AL4G02610 | Cytochrome P450 | ko:K15639 | p450 |
| AL3G04730 | Cytochrome P450 | ko:K01723 | p450 |
| AL4G04383 | cytochrome p450 | - | p450 |
| AL1G04615 | Cupin | - | Cupin_1 |
| AL1G05438 | Cupin | - | Cupin_1 |

| Gene ID | Description | KEGG_ko | PFAMs |
|---|---|---|---|
| AL1G05439 | Cupin | - | Cupin_1 |
| AL1G05440 | Cupin | - | Cupin_1 |
| AL1G05442 | Cupin | - | Cupin_1 |
| AL1G05443 | Cupin | - | Cupin_1 |
| AL1G05445 | Cupin | - | Cupin_1 |
| AL1G05447 | Cupin | - | Cupin_1 |
| AL1G05448 | Cupin | - | Cupin_1 |
| AL1G05451 | Cupin | - | Cupin_1 |
| AL1G05481 | Cupin | - | Cupin_1 |
| AL1G05482 | Cupin | - | Cupin_1 |
| AL1G05484 | Cupin | - | Cupin_1 |
| AL1G05485 | Cupin | - | Cupin_1 |
| AL1G05486 | Cupin | - | Cupin_1 |
| AL1G05492 | Cupin | - | Cupin_1 |
| AL1G05493 | Cupin | - | Cupin_1 |
| AL4G00704 | Cupin | - | Cupin_1 |
| AL4G00705 | Cupin | - | Cupin_1 |
| AL4G00732 | Cupin | - | Cupin_1 |
| AL4G00733 | Cupin | - | Cupin_1 |
| AL4G04356 | Cupin | - | Cupin_1 |
| AL4G04357 | Cupin | - | Cupin_1 |
| AL5G02859 | Cupin | - | Cupin_1 |
| AL5G02979 | Cupin | - | Cupin_1 |
| AL5G02980 | Cupin | - | Cupin_1 |
| AL5G02981 | Cupin | - | Cupin_1 |
| AL3G03052 | Pathogenesis-related protein | - | Bet_v_1,Polyketide_cyc2 |
| AL3G03054 | Pathogenesis-related protein | - | Bet_v_1,Polyketide_cyc2 |
| AL3G03055 | Pathogenesis-related protein | - | Bet_v_1,Polyketide_cyc2 |
| AL4G02534 | Pathogenesis-related protein | - | Bet_v_1,Polyketide_cyc2 |
| AL4G02535 | Pathogenesis-related protein | - | Bet_v_1,Polyketide_cyc2 |
| AL4G02536 | Pathogenesis-related protein | - | Bet_v_1,Polyketide_cyc2 |
| AL4G02543 | Pathogenesis-related protein | - | Bet_v_1,Polyketide_cyc2 |
| AL4G02544 | Pathogenesis-related protein | - | Bet_v_1,Polyketide_cyc2 |
| AL4G02545 | Pathogenesis-related protein | - | Bet_v_1,Polyketide_cyc2 |
| AL4G02546 | Pathogenesis-related protein | - | Bet_v_1,Polyketide_cyc2 |
| AL4G02547 | Pathogenesis-related protein | - | Bet_v_1,Polyketide_cyc2 |
| AL4G02548 | Pathogenesis-related protein | - | Bet_v_1,Polyketide_cyc2 |
| AL4G02549 | Pathogenesis-related protein | - | Bet_v_1,Polyketide_cyc2 |
| AL4G02558 | Pathogenesis-related protein | - | Bet_v_1,Polyketide_cyc2 |
| AL4G02559 | Pathogenesis-related protein | - | Bet_v_1,Polyketide_cyc2 |
| AL4G02560 | Pathogenesis-related protein | - | Bet_v_1,Polyketide_cyc2 |
| AL5G00218 | Pathogenesis-related protein | - | Bet_v_1,Polyketide_cyc2 |
| AL1G00761 | Other | - | - |
| AL2G02061 | Other | - | - |
| AL3G03035 | Other | - | - |
| AL3G05845 | Other | - | - |

| Gene ID | Description | KEGG_ko | PFAMs |
|---|---|---|---|
| AL4G00386 | Other | - | - |
| AL4G00572 | Other | - | - |
| AL5G04165 | Other | - | - |
| AL6G04460 | Other | - | - |
| AL6G04502 | Other | - | - |
| AL1G05073 | Other | - | F-box,FBD |
| AL2G00803 | Other | - | F-box,FBD |
| AL5G00787 | Other | - | Transposase_28 |
| AL3G03856 | Other | - | zf-CW |
| AL3G02997 | Other | ko:K01183 | Chitin_bind_1,Glyco_hydro_19 |
| AL4G01261 | Receptor kinase | - | Lectin_legB,Pkinase,Pkinase_Tyr |
| AL3G05667 | Receptor kinase | ko:K04733 | Glyco_hydro_19,Pkinase |
| AL3G03448 | Receptor kinase | - | Glyco_hydro_19,Pkinase,Stress-antifung |
| AL3G03449 | Receptor kinase | - | Glyco_hydro_19,Pkinase,Stress-antifung |
| AL3G01333 | Receptor kinase | - | Pkinase,Pkinase_Tyr |
| AL3G01334 | Receptor kinase | - | Pkinase_Tyr |
| AL3G01338 | Receptor kinase | - | Pkinase_Tyr |
| AL5G02264 | Receptor kinase | - | B_lectin,PAN_2,Pkinase,Pkinase_Tyr,S_locus_glycop |
| AL5G02265 | Receptor kinase | - | B_lectin,PAN_2,Pkinase,Pkinase_Tyr,S_locus_glycop |
| AL5G00555 | Receptor kinase | - | EGF_CA,GUB_WAK_bind,Pkinase,Pkinase_Tyr |
| AL6G00238 | Receptor kinase | - | EGF_CA,GUB_WAK_bind,Pkinase,Pkinase_Tyr |
| AL2G00542 | Receptor kinase | - | GUB_WAK_bind,Pkinase_Tyr |
| AL2G00544 | Receptor kinase | - | GUB_WAK_bind,Pkinase_Tyr |
| AL2G00550 | Receptor kinase | - | GUB_WAK_bind,Pkinase_Tyr |
| AL2G01204 | Chalcone stilbene synthases | ko:K00660,ko:K12644,ko:K13232 | Chal_sti_synt_C,Chal_sti_synt_N |
| AL2G01205 | Chalcone stilbene synthases | ko:K00660,ko:K12644,ko:K13232 | Chal_sti_synt_C,Chal_sti_synt_N |
| AL2G01207 | Chalcone stilbene synthases | ko:K00660,ko:K12644,ko:K13232 | Chal_sti_synt_C,Chal_sti_synt_N |
| AL2G01208 | Chalcone stilbene synthases | ko:K00660,ko:K12644,ko:K13232 | Chal_sti_synt_C,Chal_sti_synt_N |
| AL3G06258 | Chalcone stilbene synthases | ko:K00660,ko:K12644,ko:K13232 | Chal_sti_synt_C,Chal_sti_synt_N |
| AL3G06259 | Chalcone stilbene synthases | ko:K00660,ko:K12644,ko:K13232 | Chal_sti_synt_C,Chal_sti_synt_N |
| AL3G06260 | Chalcone stilbene synthases | ko:K00660,ko:K12644,ko:K13232 | Chal_sti_synt_C,Chal_sti_synt_N |
| AL3G06261 | Chalcone stilbene synthases | ko:K00660,ko:K | Chal_sti_synt_C,Chal_sti_synt_N |

| Gene ID | Description | KEGG_ko | PFAMs |
|---|---|---|---|
| | | 12644,ko:K13232 | |
| AL3G06262 | Chalcone stilbene synthases | ko:K00660,ko:K12644,ko:K13232 | Chal_sti_synt_C,Chal_sti_synt_N |
| AL3G06263 | Chalcone stilbene synthases | ko:K00660,ko:K12644,ko:K13232 | Chal_sti_synt_C,Chal_sti_synt_N |
| AL3G06264 | Chalcone stilbene synthases | ko:K00660,ko:K12644,ko:K13232 | Chal_sti_synt_C,Chal_sti_synt_N |
| AL3G06265 | Chalcone stilbene synthases | ko:K00660,ko:K12644,ko:K13232 | Chal_sti_synt_C,Chal_sti_synt_N |
| AL3G06267 | Chalcone stilbene synthases | ko:K00660,ko:K12644,ko:K13232 | Chal_sti_synt_C,Chal_sti_synt_N |
| AL1G04944 | UDP-glycosyl transferase | - | UDPGT |
| AL2G00518 | UDP-glycosyl transferase | - | UDPGT |
| AL2G01098 | UDP-glycosyl transferase | - | UDPGT |
| AL5G03831 | UDP-glycosyl transferase | ko:K08237 | UDPGT |
| AL4G01548 | UDP-glycosyl transferase | ko:K13495 | UDPGT |
| AL4G01551 | UDP-glycosyl transferase | ko:K13495 | UDPGT |
| AL1G04938 | UDP-glycosyl transferase | ko:K13496 | UDPGT |
| AL1G04939 | UDP-glycosyl transferase | ko:K13496 | UDPGT |
| AL3G00572 | UDP-glycosyl transferase | ko:K13496 | UDPGT |
| AL4G05828 | UDP-glycosyl transferase | ko:K21374 | UDPGT |
| AL6G03850 | UDP-glycosyl transferase | ko:K21374 | UDPGT |
| AL2G00937 | Zinc finger protein | - | zf-C2H2_6 |
| AL2G00938 | Zinc finger protein | - | zf-C2H2_6 |
| AL2G00939 | Zinc finger protein | - | zf-C2H2_6 |
| AL2G01624 | Zinc finger protein | - | zf-C2H2_6 |
| AL2G01636 | Zinc finger protein | - | zf-C2H2_6 |
| AL2G01637 | Zinc finger protein | - | zf-C2H2_6 |
| AL7G01885 | Zinc finger protein | - | zf-C2H2_6 |
| AL2G04564 | Zinc finger protein | - | zf-RING_2 |
| AL2G04565 | Zinc finger protein | - | zf-RING_2 |
| AL3G00307 | Zinc finger protein | - | zf-RING_2 |
| AL3G00311 | Zinc finger protein | - | zf-RING_2 |
| AL6G03097 | Zinc finger protein | - | zf-FLZ |
| AL3G01790 | NB-ARC domain | - | LRR_1,LRR_4,LRR_8,NB-ARC,Pkinase_Tyr |
| AL4G03329 | NB-ARC domain | - | LRR_8,NB-ARC |
| AL7G05035 | NB-ARC domain | - | LRR_8,NB-ARC |
| AL1G00532 | NB-ARC domain | - | NB-ARC |
| AL1G01021 | NB-ARC domain | - | NB-ARC |
| AL1G01035 | NB-ARC domain | - | NB-ARC |

| Gene ID | Description | KEGG_ko | PFAMs |
|---|---|---|---|
| AL2G00462 | NB-ARC domain | - | NB-ARC |
| AL2G00464 | NB-ARC domain | - | NB-ARC |
| AL1G01348 | NB-ARC domain | ko:K02737 | NB-ARC |
| AL2G05821 | Disease resistance protein | - | NB-ARC |
| AL2G05832 | Disease resistance protein | - | NB-ARC |
| AL2G05834 | Disease resistance protein | - | NB-ARC |
| AL6G00103 | disease resistance protein | - | NB-ARC |
| AL6G00108 | disease resistance protein | - | NB-ARC |
| AL6G00716 | disease resistance protein | - | NB-ARC |
| AL6G00717 | disease resistance protein | - | NB-ARC |
| AL7G02427 | disease resistance protein | ko:K13457 | NB-ARC |
| AL1G00773 | disease resistance protein | - | NB-ARC |
| AL1G04830 | Glutathione S-transferase | ko:K00799 | GST_C,GST_C_2,GST_N |
| AL1G04831 | Glutathione S-transferase | ko:K00799 | GST_C,GST_C_2,GST_N,GST_N_3 |
| AL5G04168 | Glutathione S-transferase | ko:K00799 | GST_C,GST_C_2,GST_N,GST_N_3 |
| AL3G04311 | Glutathione S-transferase | ko:K00799 | GST_C_2,GST_N_3 |
| AL1G01205 | Glutathione S-transferase | - | GST_N_3 |
| AL1G01208 | Glutathione S-transferase | - | GST_N_3 |
| AL1G02068 | Glutathione S-transferase | ko:K00799 | GST_C,GST_C_2,GST_N |
| AL1G05641 | Glutathione S-transferase | ko:K00799 | GST_C,GST_C_2,GST_N,GST_N_3 |
| AL1G00734 | Leucine rich repeat | - | LRRNT_2,LRR_1,LRR_6,LRR_8 |
| AL6G03152 | Leucine rich repeat | - | LRRNT_2,LRR_1,LRR_6,LRR_8 |
| AL3G05593 | Leucine rich repeat | - | LRRNT_2,LRR_1,LRR_4,LRR_8,Pkinase,Pkinase_Tyr |
| AL3G05611 | Leucine rich repeat | - | LRRNT_2,LRR_1,LRR_4,LRR_8,Pkinase,Pkinase_Tyr |
| AL7G01132 | Leucine rich repeat | - | LRRNT_2,LRR_1,LRR_4,LRR_8,Pkinase |
| AL2G01327 | Leucine rich repeat | - | LRRNT_2,LRR_1,LRR_6,LRR_8 |
| AL2G01351 | Leucine rich repeat | - | LRRNT_2,LRR_1,LRR_6,LRR_8 |
| AL4G02990 | Leucine rich repeat | - | LRRNT_2,LRR_8,Pkinase,Pkinase_Tyr |
| AL1G06274 | No apical meristem (NAM) protein | - | NAM |
| AL1G06284 | No apical meristem (NAM) protein | - | NAM |
| AL2G04030 | No apical meristem (NAM) protein | - | NAM |
| AL4G00667 | No apical meristem (NAM) protein | - | NAM |
| AL4G00741 | No apical meristem (NAM) protein | - | NAM |
| AL4G05096 | No apical meristem (NAM) protein | - | NAM |
| AL7G00842 | No apical meristem (NAM) protein | - | NAM |
| AL2G04367 | Glycosyl hydrolase | ko:K01188,ko:K19964 | Glyco_hydro_1 |
| AL1G01324 | Glycosyl hydrolase | - | Glyco_hydro_17 |
| AL1G01345 | Glycosyl hydrolase | - | Glyco_hydro_17 |

| Gene ID | Description | KEGG_ko | PFAMs |
|---|---|---|---|
| AL1G01385 | Glycosyl hydrolase | - | Glyco_hydro_17 |
| AL1G01388 | Glycosyl hydrolase | - | Glyco_hydro_17 |
| AL4G01769 | Glycosyl hydrolase | ko:K01193 | DUF3357,Glyco_hydro_32C,Glyco_hydro_32N |
| AL4G01816 | Glycosyl hydrolase | ko:K01193 | DUF3357,Glyco_hydro_32C,Glyco_hydro_32N |
| AL5G00189 | Salt stress response/antifungal | - | Pkinase,Pkinase_Tyr,Stress-antifung |
| AL7G00705 | Salt stress response/antifungal | - | Pkinase,Pkinase_Tyr,Stress-antifung |
| AL2G01649 | Salt stress response/antifungal | - | Stress-antifung |
| AL2G01653 | Salt stress response/antifungal | - | Stress-antifung |
| AL2G01696 | Salt stress response/antifungal | - | Stress-antifung |
| AL2G01699 | Salt stress response/antifungal | - | Stress-antifung |
| AL3G03461 | Salt stress response/antifungal | ko:K04733 | Pkinase,Stress-antifung |
| AL2G00502 | ABC transporter | - | ABC2_membrane,ABC_tran |
| AL2G00507 | ABC transporter | - | ABC2_membrane,ABC_tran |
| AL1G03202 | ABC transporter | ko:K05658 | ABC_membrane,ABC_tran |
| AL2G01127 | ABC transporter | ko:K05665,ko:K05666 | ABC_membrane,ABC_tran |
| AL5G00651 | ABC transporter | ko:K05666 | ABC_membrane,ABC_tran |
| AL5G00657 | ABC transporter | ko:K05666 | ABC_membrane,ABC_tran |
| AL3G01096 | Glycosyl transferase | - | Cellulose_synt |
| AL5G00190 | Glycosyl transferase | ko:K20924 | Cellulose_synt,zf-RING_4 |
| AL7G02472 | Glycosyl transferase | - | Glyco_tranf_2_3,Glyco_trans_2_3 |
| AL4G04642 | Glycosyl transferase | ko:K09667 | Glyco_transf_41,LEA_5,TPR_1,TPR_11,TPR_16,TPR_7,TPR_8 |
| AL4G01940 | Glycosyl transferase | - | Glyco_transf_29 |
| AL4G01945 | Glycosyl transferase | - | Glyco_transf_29 |
| AL3G00043 | Oxidoreductase | - | 2OG-FeII_Oxy,DIOX_N |
| AL5G02139 | Oxidoreductase | - | 2OG-FeII_Oxy,DIOX_N |
| AL1G02944 | Oxidoreductase | ko:K04125 | 2OG-FeII_Oxy,DIOX_N |
| AL6G04259 | Oxidoreductase | ko:K05933 | 2OG-FeII_Oxy,DIOX_N |
| AL6G04267 | Oxidoreductase | ko:K05933 | 2OG-FeII_Oxy,DIOX_N |
| AL6G04273 | Oxidoreductase | ko:K05933 | 2OG-FeII_Oxy,DIOX_N |
| AL1G01307 | Amino acid permease | - | AA_permease_2 |
| AL1G01310 | Amino acid permease | - | AA_permease_2 |
| AL1G01314 | Amino acid permease | - | AA_permease_2 |
| AL1G05436 | Amino acid permease | - | AA_permease_2 |
| AL4G05076 | Amino acid permease | - | AA_permease_2 |
| AL4G02880 | Dirigent | - | Dirigent |
| AL4G02883 | Dirigent | - | Dirigent |
| AL4G02893 | Dirigent | - | Dirigent |
| AL4G02894 | Dirigent | - | Dirigent |
| AL3G00037 | Dirigent | - | Dirigent,Jacalin |
| AL3G04004 | DnaJ domain | - | DnaJ |
| AL4G02895 | DnaJ domain | - | DnaJ |
| AL6G04511 | DnaJ domain | ko:K09512 | DnaJ |
| AL6G05388 | DnaJ domain | ko:K09512 | DnaJ |
| AL5G00949 | DnaJ domain | ko:K09512 | DnaJ |

| Gene ID | Description | KEGG_ko | PFAMs |
|---|---|---|---|
| AL5G04145 | F-Box protein | - | Kelch_6 |
| AL5G04163 | F-Box protein | - | Kelch_6 |
| AL3G01691 | F-Box protein | ko:K10268,ko:K10632 | F-box,LRR_6 |
| AL3G01695 | F-Box protein | ko:K10268,ko:K10632 | F-box,LRR_6 |
| AL4G05029 | F-Box protein | ko:K10295 | F-box,F-box-like |
| AL3G05263 | Heat shock protein | ko:K08967,ko:K09419 | HSF_DNA-bind |
| AL6G02226 | Heat shock protein | ko:K09419 | HSF_DNA-bind |
| AL6G02229 | Heat shock protein | ko:K09419 | HSF_DNA-bind |
| AL2G04650 | Heat shock protein | ko:K03283 | HSP70,MreB_Mbl |
| AL3G04383 | Heat shock protein | ko:K03283 | HSP70,MreB_Mbl |
| AL4G04289 | Legume lectin domain | - | Lectin_legB,Pkinase |
| AL4G04504 | Legume lectin domain | - | Lectin_legB,Pkinase |
| AL2G02011 | Legume lectin domain | - | Lectin_legB,Pkinase,Pkinase_Tyr |
| AL4G01787 | Legume lectin domain | - | Lectin_legB,Pkinase,Pkinase_Tyr |
| AL4G03278 | Legume lectin domain | - | Lectin_legB,Pkinase,Pkinase_Tyr |
| AL2G02220 | MYB-like DNA-binding domain | ko:K09422 | Myb_DNA-binding |
| AL4G01103 | MYB-like DNA-binding domain | ko:K09422 | Myb_DNA-binding |
| AL4G01116 | MYB-like DNA-binding domain | ko:K09422 | Myb_DNA-binding |
| AL4G02022 | MYB-like DNA-binding domain | ko:K09422 | Myb_DNA-binding |
| AL6G03272 | MYB-like DNA-binding domain | ko:K09422 | Myb_DNA-binding |
| AL3G00043 | Oxidoreductase | - | 2OG-FeII_Oxy,DIOX_N |
| AL5G02139 | Oxidoreductase | - | 2OG-FeII_Oxy,DIOX_N |
| AL1G02944 | Oxidoreductase | ko:K04125 | 2OG-FeII_Oxy,DIOX_N |
| AL6G04259 | Oxidoreductase | ko:K05933 | 2OG-FeII_Oxy,DIOX_N |
| AL6G04267 | Oxidoreductase | ko:K05933 | 2OG-FeII_Oxy,DIOX_N |
| AL6G04273 | Oxidoreductase | ko:K05933 | 2OG-FeII_Oxy,DIOX_N |
| AL1G01307 | Amino acid permease | - | AA_permease_2 |
| AL1G01310 | Amino acid permease | - | AA_permease_2 |
| AL1G01314 | Amino acid permease | - | AA_permease_2 |
| AL1G05436 | Amino acid permease | - | AA_permease_2 |
| AL4G05076 | Amino acid permease | - | AA_permease_2 |
| AL4G02880 | Dirigent | - | Dirigent |
| AL4G02883 | Dirigent | - | Dirigent |
| AL4G02893 | Dirigent | - | Dirigent |
| AL4G02894 | Dirigent | - | Dirigent |
| AL3G00037 | Dirigent | - | Dirigent,Jacalin |
| AL3G04004 | DnaJ domain | - | DnaJ |
| AL4G02895 | DnaJ domain | - | DnaJ |

| Gene ID | Description | KEGG_ko | PFAMs |
|---|---|---|---|
| AL6G04511 | DnaJ domain | ko:K09512 | DnaJ |
| AL6G05388 | DnaJ domain | ko:K09512 | DnaJ |
| AL5G00949 | DnaJ domain | ko:K09512 | DnaJ |
| AL6G05013 | EXS family | - | EXS,SPX |
| AL5G04145 | F-Box protein | - | Kelch_6 |
| AL5G04163 | F-Box protein | - | Kelch_6 |
| AL3G01691 | F-Box protein | ko:K10268,ko:K10632 | F-box,LRR_6 |
| AL3G01695 | F-Box protein | ko:K10268,ko:K10632 | F-box,LRR_6 |
| AL4G05029 | F-Box protein | ko:K10295 | F-box,F-box-like |
| AL3G05263 | Heat shock protein | ko:K08967,ko:K09419 | HSF_DNA-bind |
| AL6G02226 | Heat shock protein | ko:K09419 | HSF_DNA-bind |
| AL6G02229 | Heat shock protein | ko:K09419 | HSF_DNA-bind |
| AL2G04650 | Heat shock protein | ko:K03283 | HSP70,MreB_Mbl |
| AL3G04383 | Heat shock protein | ko:K03283 | HSP70,MreB_Mbl |
| AL4G04289 | Legume lectin domain | - | Lectin_legB,Pkinase |
| AL4G04504 | Legume lectin domain | - | Lectin_legB,Pkinase |
| AL2G02011 | Legume lectin domain | - | Lectin_legB,Pkinase,Pkinase_Tyr |
| AL4G01787 | Legume lectin domain | - | Lectin_legB,Pkinase,Pkinase_Tyr |
| AL4G03278 | Legume lectin domain | - | Lectin_legB,Pkinase,Pkinase_Tyr |
| AL2G02220 | MYB-like DNA-binding domain | ko:K09422 | Myb_DNA-binding |
| AL4G01103 | MYB-like DNA-binding domain | ko:K09422 | Myb_DNA-binding |
| AL4G01116 | MYB-like DNA-binding domain | ko:K09422 | Myb_DNA-binding |
| AL4G02022 | MYB-like DNA-binding domain | ko:K09422 | Myb_DNA-binding |
| AL6G03272 | MYB-like DNA-binding domain | ko:K09422 | Myb_DNA-binding |
| AL5G04145 | F-Box protein | - | Kelch_6 |
| AL5G04163 | F-Box protein | - | Kelch_6 |
| AL3G01691 | F-Box protein | ko:K10268,ko:K10632 | F-box,LRR_6 |
| AL3G01695 | F-Box protein | ko:K10268,ko:K10632 | F-box,LRR_6 |
| AL4G05029 | F-Box protein | ko:K10295 | F-box,F-box-like |
| AL3G05263 | Heat shock protein | ko:K08967,ko:K09419 | HSF_DNA-bind |
| AL6G02226 | Heat shock protein | ko:K09419 | HSF_DNA-bind |
| AL6G02229 | Heat shock protein | ko:K09419 | HSF_DNA-bind |
| AL2G04650 | Heat shock protein | ko:K03283 | HSP70,MreB_Mbl |
| AL3G04383 | Heat shock protein | ko:K03283 | HSP70,MreB_Mbl |
| AL4G04289 | Legume lectin domain | - | Lectin_legB,Pkinase |

| Gene ID | Description | KEGG_ko | PFAMs |
|---|---|---|---|
| AL4G04504 | Legume lectin domain | - | Lectin_legB,Pkinase |
| AL2G02011 | Legume lectin domain | - | Lectin_legB,Pkinase,Pkinase_Tyr |
| AL4G01787 | Legume lectin domain | - | Lectin_legB,Pkinase,Pkinase_Tyr |
| AL4G03278 | Legume lectin domain | - | Lectin_legB,Pkinase,Pkinase_Tyr |
| AL2G02220 | MYB-like DNA-binding domain | ko:K09422 | Myb_DNA-binding |
| AL4G01103 | MYB-like DNA-binding domain | ko:K09422 | Myb_DNA-binding |
| AL4G01116 | MYB-like DNA-binding domain | ko:K09422 | Myb_DNA-binding |
| AL4G02022 | MYB-like DNA-binding domain | ko:K09422 | Myb_DNA-binding |
| AL6G03272 | MYB-like DNA-binding domain | ko:K09422 | Myb_DNA-binding |
| AL5G04145 | F-Box protein | - | Kelch_6 |
| AL5G04163 | F-Box protein | - | Kelch_6 |
| AL3G01691 | F-Box protein | ko:K10268,ko:K10632 | F-box,LRR_6 |
| AL3G01695 | F-Box protein | ko:K10268,ko:K10632 | F-box,LRR_6 |
| AL4G05029 | F-Box protein | ko:K10295 | F-box,F-box-like |
| AL3G05263 | Heat shock protein | ko:K08967,ko:K09419 | HSF_DNA-bind |
| AL6G02226 | Heat shock protein | ko:K09419 | HSF_DNA-bind |
| AL6G02229 | Heat shock protein | ko:K09419 | HSF_DNA-bind |
| AL2G04650 | Heat shock protein | ko:K03283 | HSP70,MreB_Mbl |
| AL3G04383 | Heat shock protein | ko:K03283 | HSP70,MreB_Mbl |
| AL4G04289 | Legume lectin domain | - | Lectin_legB,Pkinase |
| AL4G04504 | Legume lectin domain | - | Lectin_legB,Pkinase |
| AL2G02011 | Legume lectin domain | - | Lectin_legB,Pkinase,Pkinase_Tyr |
| AL4G01787 | Legume lectin domain | - | Lectin_legB,Pkinase,Pkinase_Tyr |
| AL4G03278 | Legume lectin domain | - | Lectin_legB,Pkinase,Pkinase_Tyr |
| AL2G02220 | MYB-like DNA-binding domain | ko:K09422 | Myb_DNA-binding |
| AL4G01103 | MYB-like DNA-binding domain | ko:K09422 | Myb_DNA-binding |
| AL4G01116 | MYB-like DNA-binding domain | ko:K09422 | Myb_DNA-binding |
| AL4G02022 | MYB-like DNA-binding domain | ko:K09422 | Myb_DNA-binding |
| AL6G03272 | MYB-like DNA-binding domain | ko:K09422 | Myb_DNA-binding |
| AL4G05029 | F-Box protein | ko:K10295 | F-box,F-box-like |
| AL3G05263 | Heat shock protein | ko:K08967,ko:K09419 | HSF_DNA-bind |
| AL6G02226 | Heat shock protein | ko:K09419 | HSF_DNA-bind |

| Gene ID | Description | KEGG_ko | PFAMs |
|---|---|---|---|
| AL6G02229 | Heat shock protein | ko:K09419 | HSF_DNA-bind |
| AL2G04650 | Heat shock protein | ko:K03283 | HSP70,MreB_Mbl |
| AL3G04383 | Heat shock protein | ko:K03283 | HSP70,MreB_Mbl |
| AL4G04289 | Legume lectin domain | - | Lectin_legB,Pkinase |
| AL4G04504 | Legume lectin domain | - | Lectin_legB,Pkinase |
| AL2G02011 | Legume lectin domain | - | Lectin_legB,Pkinase,Pkinase_Tyr |
| AL4G01787 | Legume lectin domain | - | Lectin_legB,Pkinase,Pkinase_Tyr |
| AL4G03278 | Legume lectin domain | - | Lectin_legB,Pkinase,Pkinase_Tyr |
| AL2G02220 | MYB-like DNA-binding domain | ko:K09422 | Myb_DNA-binding |
| AL4G01103 | MYB-like DNA-binding domain | ko:K09422 | Myb_DNA-binding |
| AL4G01116 | MYB-like DNA-binding domain | ko:K09422 | Myb_DNA-binding |
| AL4G02022 | MYB-like DNA-binding domain | ko:K09422 | Myb_DNA-binding |
| AL6G03272 | MYB-like DNA-binding domain | ko:K09422 | Myb_DNA-binding |
| 353 down-regulation DEGs (gene number ≥ 5 for each category) in expanded gene families of of salt-treated roots of ALO | | | |
| AL1G02385 | Nucleosome histone | ko:K11252 | Histone |
| AL2G04647 | Nucleosome histone | ko:K11252 | Histone |
| AL2G03646 | Nucleosome histone | ko:K11254 | CENP-T_C |
| AL2G03812 | Nucleosome histone | ko:K11254 | CENP-T_C |
| AL2G04590 | Nucleosome histone | ko:K11254 | CENP-T_C |
| AL2G04615 | Nucleosome histone | ko:K11254 | CENP-T_C |
| AL2G04765 | Nucleosome histone | ko:K11254 | CENP-T_C |
| AL3G00328 | Nucleosome histone | ko:K11254 | CENP-T_C |
| AL3G01397 | Nucleosome histone | ko:K11254 | CENP-T_C |
| AL3G01398 | Nucleosome histone | ko:K11254 | CENP-T_C |
| AL3G01401 | Nucleosome histone | ko:K11254 | CENP-T_C |
| AL3G04097 | Nucleosome histone | ko:K11254 | CENP-T_C |
| AL3G04480 | Nucleosome histone | ko:K11254 | CENP-T_C |
| AL3G04573 | Nucleosome histone | ko:K11254 | CENP-T_C |
| AL3G05826 | Nucleosome histone | ko:K11254 | CENP-T_C |
| AL4G01282 | Nucleosome histone | ko:K11254 | CENP-T_C |
| AL4G01283 | Nucleosome histone | ko:K11254 | CENP-T_C |
| AL4G01284 | Nucleosome histone | ko:K11254 | CENP-T_C |
| AL4G01295 | Nucleosome histone | ko:K11254 | CENP-T_C |
| AL4G01296 | Nucleosome histone | ko:K11254 | CENP-T_C |
| AL4G04367 | Nucleosome histone | ko:K11254 | CENP-T_C |
| AL6G01315 | Nucleosome histone | ko:K11254 | CENP-T_C |
| AL6G01624 | Nucleosome histone | ko:K11254 | CENP-T_C |
| AL6G02005 | Nucleosome histone | ko:K11254 | CENP-T_C |
| AL6G02341 | Nucleosome histone | ko:K11254 | CENP-T_C |
| AL6G03219 | Nucleosome histone | ko:K11254 | CENP-T_C |
| AL1G05225 | Nucleosome histone | ko:K11252 | Histone |

| Gene ID | Description | KEGG_ko | PFAMs |
|---|---|---|---|
| AL2G00950 | Nucleosome histone | ko:K11251 | Histone,Histone_H2A_C |
| AL1G00442 | Nucleosome histone | ko:K11251 | Histone,Histone_H2A_C |
| AL1G01007 | Nucleosome histone | ko:K11251 | Histone,Histone_H2A_C |
| AL1G01010 | Nucleosome histone | ko:K11251 | Histone,Histone_H2A_C |
| AL1G04238 | Nucleosome histone | ko:K11251 | Histone,Histone_H2A_C |
| AL1G05009 | Nucleosome histone | ko:K11251 | Histone,Histone_H2A_C |
| AL2G00951 | Nucleosome histone | ko:K11251 | Histone,Histone_H2A_C |
| AL2G05251 | Nucleosome histone | ko:K11251 | Histone,Histone_H2A_C |
| AL3G04354 | Nucleosome histone | ko:K11251 | Histone,Histone_H2A_C |
| AL3G04355 | Nucleosome histone | ko:K11251 | Histone,Histone_H2A_C |
| AL4G05052 | Nucleosome histone | ko:K11251 | Histone,Histone_H2A_C |
| AL5G01591 | Nucleosome histone | ko:K11251 | Histone,Histone_H2A_C |
| AL5G03167 | Nucleosome histone | ko:K11251 | Histone,Histone_H2A_C |
| AL5G03168 | Nucleosome histone | ko:K11251 | Histone,Histone_H2A_C |
| AL6G01705 | Nucleosome histone | ko:K11251 | Histone,Histone_H2A_C |
| AL6G01712 | Nucleosome histone | ko:K11251 | Histone,Histone_H2A_C |
| AL7G03974 | Nucleosome histone | ko:K11251 | Histone,Histone_H2A_C |
| AL7G04428 | Nucleosome histone | ko:K11251 | Histone,Histone_H2A_C |
| AL1G05222 | Nucleosome histone | ko:K11252 | Histone |
| AL1G05223 | Nucleosome histone | ko:K11252 | Histone |
| AL2G02000 | Nucleosome histone | ko:K11252 | Histone |
| AL2G02596 | Nucleosome histone | ko:K11252 | Histone |
| AL2G04362 | Nucleosome histone | ko:K11252 | Histone |
| AL3G04359 | Nucleosome histone | ko:K11252 | Histone |
| AL4G00784 | Nucleosome histone | ko:K11252 | Histone |
| AL4G00788 | Nucleosome histone | ko:K11252 | Histone |
| AL4G00789 | Nucleosome histone | ko:K11252 | Histone |
| AL4G00811 | Nucleosome histone | ko:K11252 | Histone |
| AL5G01496 | Nucleosome histone | ko:K11252 | Histone |
| AL5G01589 | Nucleosome histone | ko:K11252 | Histone |
| AL5G01737 | Nucleosome histone | ko:K11252 | Histone |
| AL7G01513 | Nucleosome histone | ko:K11252 | Histone |
| AL7G03854 | Nucleosome histone | ko:K11252 | Histone |
| AL3G00070 | Nucleosome histone | ko:K02326 | CBFD_NFYB_HMF |
| AL1G05258 | Nucleosome histone | ko:K11252 | Histone |
| AL1G02198 | Nucleosome histone | ko:K11253 | Histone |
| AL1G03295 | Nucleosome histone | ko:K11253 | Histone |
| AL2G00811 | Nucleosome histone | ko:K11253 | Histone |
| AL2G04193 | Nucleosome histone | ko:K11253 | Histone |
| AL2G04363 | Nucleosome histone | ko:K11253 | Histone |
| AL2G04764 | Nucleosome histone | ko:K11253 | Histone |
| AL2G05751 | Nucleosome histone | ko:K11253 | Histone |
| AL3G00687 | Nucleosome histone | ko:K11253 | Histone |
| AL3G04029 | Nucleosome histone | ko:K11253 | Histone |
| AL4G04146 | Nucleosome histone | ko:K11253 | Histone |
| AL4G04945 | Nucleosome histone | ko:K11253 | Histone |

| Gene ID | Description | KEGG_ko | PFAMs |
|---|---|---|---|
| AL5G00605 | Nucleosome histone | ko:K11253 | Histone |
| AL5G00649 | Nucleosome histone | ko:K11253 | Histone |
| AL5G01407 | Nucleosome histone | ko:K11253 | Histone |
| AL5G03418 | Nucleosome histone | ko:K11253 | Histone |
| AL6G04086 | Nucleosome histone | ko:K11253 | Histone |
| AL6G04105 | Nucleosome histone | ko:K11253 | Histone |
| AL7G02135 | Nucleosome histone | ko:K11253 | Histone |
| AL1G01009 | Nucleosome histone | ko:K11251 | Histone,Histone_H2A_C |
| AL1G00162 | Other | - | Other |
| AL1G00170 | Other | - | Other |
| AL1G00210 | Other | - | Other |
| AL1G01961 | Other | - | Other |
| AL1G01968 | Other | - | Other |
| AL1G06332 | Other | - | Other |
| AL1G06351 | Other | - | Other |
| AL2G00576 | Other | - | Other |
| AL2G01040 | Other | - | Other |
| AL2G05412 | Other | - | Other |
| AL3G00495 | Other | - | Other |
| AL3G04536 | Other | - | Other |
| AL3G05642 | Other | - | Other |
| AL3G05662 | Other | - | Other |
| AL4G00668 | Other | - | Other |
| AL4G00740 | Other | - | Other |
| AL4G01138 | Other | - | Other |
| AL6G00045 | Other | - | Other |
| AL6G00951 | Other | - | Other |
| AL6G01609 | Other | - | Other |
| AL6G01834 | Other | - | Other |
| AL6G04950 | Other | - | Other |
| AL7G00159 | Other | - | Other |
| AL7G00189 | Other | - | Other |
| AL2G00805 | Other | - | DUF295 |
| AL2G01202 | Other | - | LEA_2 |
| AL3G06269 | Other | - | LEA_2 |
| AL1G05866 | Other | - | NPR1_interact |
| AL1G05869 | Other | - | NPR1_interact |
| AL1G05874 | Other | - | NPR1_interact |
| AL1G05876 | Other | - | NPR1_interact |
| AL4G01899 | Other | - | TPR_8 |
| AL5G02493 | Other | ko:K06630 | 2014/3/3 |
| AL6G04835 | Other | - | PGG |
| AL6G04853 | Other | - | PGG |
| AL1G02055 | Other | - | LRRNT_2,LRR_1,LRR_8 |
| AL4G00361 | Other | - | RRM_1 |
| AL2G04316 | Other | ko:K13207 | RRM_1 |

| Gene ID | Description | KEGG_ko | PFAMs |
|---|---|---|---|
| AL2G02929 | Protein kinase | ko:K00924,ko:K14502 | Pkinase |
| AL2G03479 | Protein kinase | ko:K04733 | Pkinase |
| AL4G01372 | Protein kinase | - | LRRNT_2,LRR_1,LRR_4,LRR_6,LRR_8,Pkinase,Pkinase_Tyr |
| AL4G01373 | Protein kinase | - | LRRNT_2,LRR_1,LRR_4,LRR_6,LRR_8,Pkinase,Pkinase_Tyr |
| AL1G00089 | Protein kinase | - | LRRNT_2,LRR_1,LRR_6,LRR_8,Pkinase |
| AL2G05565 | Protein kinase | - | LRRNT_2,LRR_1,LRR_6,LRR_8,Pkinase |
| AL5G02756 | Protein kinase | - | LRRNT_2,LRR_1,LRR_6,LRR_8,Pkinase |
| AL4G00739 | Protein kinase | ko:K13420 | LRRNT_2,LRR_1,LRR_6,LRR_8,Pkinase,Pkinase_Tyr |
| AL1G05889 | Protein kinase | - | DUF3403,Pkinase,Pkinase_Tyr,Stress-antifung |
| AL4G00715 | Protein kinase | - | LRR_8,Malectin,Pkinase,Pkinase_Tyr |
| AL4G00722 | Protein kinase | - | LRR_8,Malectin,Pkinase,Pkinase_Tyr |
| AL2G05886 | Protein kinase | - | LRR_8,Malectin_like,Pkinase,Pkinase_Tyr |
| AL2G05898 | Protein kinase | - | LRR_8,Malectin_like,Pkinase,Pkinase_Tyr |
| AL2G05684 | Protein kinase | ko:K20716 | Pkinase |
| AL2G05685 | Protein kinase | ko:K20716 | Pkinase |
| AL2G05686 | Protein kinase | ko:K20716 | Pkinase |
| AL2G05687 | Protein kinase | ko:K20716 | Pkinase |
| AL2G05689 | Protein kinase | ko:K20716 | Pkinase |
| AL1G05577 | Protein kinase | - | Pkinase |
| AL7G03798 | Protein kinase | - | Pkinase,Pkinase_Tyr |
| AL3G03110 | Protein kinase | - | EGF_CA,GUB_WAK_bind,Pkinase,Pkinase_Tyr |
| AL4G04131 | Protein kinase | - | LRRNT_2,LRR_1,LRR_8,Pkinase,Pkinase_Tyr |
| AL7G03639 | Protein kinase | - | LRRNT_2,LRR_1,LRR_8,Pkinase,Pkinase_Tyr |
| AL5G00912 | Protein kinase | - | Pkinase,Pkinase_Tyr,Usp |
| AL1G05579 | Protein kinase | - | Pkinase,Stress-antifung |
| AL5G03781 | Protein kinase | ko:K04730 | LRRNT_2,LRR_1,LRR_8,Pkinase,Pkinase_Tyr |
| AL6G00920 | Protein kinase | ko:K06632 | Pkinase |
| AL6G00932 | Protein kinase | ko:K06632 | Pkinase |
| AL2G05690 | Protein kinase | ko:K20716 | Pkinase |
| AL3G00183 | Cytochrome P450 | - | p450 |
| AL4G00084 | Cytochrome P450 | - | p450 |
| AL4G00086 | Cytochrome P450 | - | p450 |
| AL4G00088 | Cytochrome P450 | - | p450 |
| AL6G04934 | Cytochrome P450 | - | p450 |
| AL3G02339 | Cytochrome P450 | - | p450,zf-RING_2 |
| AL1G00991 | Cytochrome P450 | ko:K00512 | p450 |
| AL3G06343 | Cytochrome P450 | ko:K00512 | p450 |
| AL6G04885 | Cytochrome P450 | ko:K00512 | p450 |
| AL2G01337 | Cytochrome P450 | ko:K00512,ko:K13029 | p450 |
| AL2G01343 | Cytochrome P450 | ko:K00512,ko:K13029 | p450 |
| AL5G00149 | Cytochrome P450 | ko:K00512,ko:K | p450 |

| Gene ID | Description | KEGG_ko | PFAMs |
|---|---|---|---|
| | | 13029 | |
| AL5G01310 | Cytochrome P450 | ko:K00512,ko:K13029 | p450 |
| AL5G00013 | Cytochrome P450 | ko:K05917 | p450 |
| AL5G00019 | Cytochrome P450 | ko:K05917 | p450 |
| AL1G04786 | Cytochrome P450 | ko:K07408,ko:K14985 | p450 |
| AL3G05629 | Cytochrome P450 | ko:K20619 | p450 |
| AL3G05648 | Cytochrome P450 | ko:K20619 | p450 |
| AL5G03203 | Cytochrome P450 | - | p450 |
| AL6G05296 | Cytochrome P450 | - | DUF1668,Sec63,p450 |
| AL1G05860 | Cytochrome P450 | ko:K00512,ko:K07408 | p450 |
| AL1G05871 | Cytochrome P450 | ko:K00512,ko:K07408 | p450 |
| AL1G01443 | Peptidase family | - | TAXi_C,TAXi_N |
| AL1G01445 | Peptidase family | - | TAXi_C,TAXi_N |
| AL1G01446 | Peptidase family | - | TAXi_C,TAXi_N |
| AL1G01448 | Peptidase family | - | TAXi_C,TAXi_N |
| AL1G01466 | Peptidase family | - | TAXi_C,TAXi_N |
| AL1G01467 | Peptidase family | - | TAXi_C,TAXi_N |
| AL1G01470 | Peptidase family | - | TAXi_C,TAXi_N |
| AL3G06280 | Peptidase family | - | TAXi_C,TAXi_N |
| AL4G04587 | Peptidase family | - | TAXi_C,TAXi_N |
| AL5G00313 | Peptidase family | ko:K01381 | TAXi_C,TAXi_N |
| AL5G00314 | Peptidase family | ko:K01381 | TAXi_C,TAXi_N |
| AL5G00315 | Peptidase family | ko:K01381 | TAXi_C,TAXi_N |
| AL5G00316 | Peptidase family | ko:K01381 | TAXi_C,TAXi_N |
| AL5G00317 | Peptidase family | ko:K01381 | TAXi_C,TAXi_N |
| AL2G01592 | Peptidase family | ko:K16290 | Inhibitor_I29,Peptidase_C1 |
| AL2G01614 | Peptidase family | ko:K16290 | Inhibitor_I29,Peptidase_C1 |
| AL4G01606 | Peptidase family | ko:K09756,ko:K16296 | Peptidase_S10 |
| AL3G00594 | Peptidase family | ko:K16296 | Peptidase_S10 |
| AL3G01196 | Peptidase family | ko:K16297 | Peptidase_S10 |
| AL4G02520 | Peptidase family | ko:K16297 | Peptidase_S10 |
| AL4G01756 | Peptidase family | ko:K01369 | Peptidase_C13 |
| AL2G00545 | Peroxidase family | ko:K00434 | peroxidase |
| AL2G00548 | Peroxidase family | ko:K00434 | peroxidase |
| AL2G00778 | Peroxidase family | ko:K00430 | peroxidase |
| AL2G00779 | Peroxidase family | ko:K00430 | peroxidase |
| AL2G00780 | Peroxidase family | ko:K00430 | peroxidase |
| AL2G00973 | Peroxidase family | ko:K00430 | peroxidase |
| AL2G00974 | Peroxidase family | ko:K00430 | peroxidase |
| AL2G00975 | Peroxidase family | ko:K00430 | peroxidase |
| AL2G01187 | Peroxidase family | ko:K00430 | peroxidase |

| Gene ID | Description | KEGG_ko | PFAMs |
|---|---|---|---|
| AL2G01196 | Peroxidase family | ko:K00430 | peroxidase |
| AL2G01197 | Peroxidase family | ko:K00430 | peroxidase |
| AL2G01198 | Peroxidase family | ko:K00430 | peroxidase |
| AL2G02229 | Peroxidase family | ko:K00430 | peroxidase |
| AL2G02256 | Peroxidase family | ko:K00430 | peroxidase |
| AL2G02915 | Peroxidase family | ko:K00430 | peroxidase |
| AL2G03631 | Peroxidase family | ko:K00430 | peroxidase |
| AL3G06275 | Peroxidase family | ko:K00430 | peroxidase |
| AL3G06284 | Peroxidase family | ko:K00430 | peroxidase |
| AL4G00985 | Peroxidase family | ko:K00430 | peroxidase |
| AL4G04450 | Peroxidase family | ko:K00430 | peroxidase |
| AL1G05842 | Methyltransferase | ko:K13259,ko:K13262,ko:K16040 | Dimerisation,Methyltransf_2 |
| AL1G05855 | Methyltransferase | ko:K13259,ko:K13262,ko:K16040 | Dimerisation,Methyltransf_2 |
| AL2G04955 | Methyltransferase | ko:K13259,ko:K13262,ko:K16040 | Dimerisation,Methyltransf_2 |
| AL3G03475 | Methyltransferase | ko:K13259,ko:K13262,ko:K16040 | Dimerisation,Methyltransf_2 |
| AL5G00223 | Methyltransferase | ko:K13259,ko:K13262,ko:K16040 | Dimerisation,Methyltransf_2 |
| AL1G00515 | Methyltransferase | ko:K16040 | Dimerisation,Methyltransf_2 |
| AL2G05971 | Methyltransferase | ko:K16040 | Dimerisation,Methyltransf_2 |
| AL2G05972 | Methyltransferase | ko:K16040 | Dimerisation,Methyltransf_2 |
| AL2G05973 | Methyltransferase | ko:K16040 | Dimerisation,Methyltransf_2 |
| AL2G03686 | Methyltransferase | ko:K08242 | Methyltransf_11,Sterol_MT_C |
| AL3G01217 | Methyltransferase | ko:K01887 | Arg_tRNA_synt_N,DALR_1,tRNA-synt_1d |
| AL5G02246 | Methyltransferase | ko:K00558 | BAH,Chromo,DNA_methylase |
| AL6G00268 | Methyltransferase | - | PHD,SET |
| AL6G00278 | Methyltransferase | - | PHD,SET |
| AL3G02700 | Methyltransferase | - | Methyltransf_29 |
| AL6G03106 | Methyltransferase | - | Methyltransf_29 |
| AL6G01682 | Methyltransferase | - | Methyltrans_SAM |
| AL5G01525 | Methyltransferase | ko:K21485 | Methyltransf_7 |
| AL2G05874 | UDP-glucosyl transferase | - | UDPGT |
| AL5G00006 | UDP-glucosyl transferase | - | UDPGT |
| AL5G00007 | UDP-glucosyl transferase | - | UDPGT |
| AL5G00044 | UDP-glucosyl transferase | - | UDPGT |
| AL6G05363 | UDP-glucosyl transferase | - | UDPGT |
| AL7G00016 | UDP-glucosyl transferase | - | UDPGT |
| AL7G00439 | UDP-glucosyl transferase | - | UDPGT |

| Gene ID | Description | KEGG_ko | PFAMs |
|---|---|---|---|
| AL7G02074 | UDP-glucosyl transferase | - | UDPGT |
| AL2G00546 | UDP-glucosyl transferase | ko:K13495 | UDPGT |
| AL5G00001 | UDP-glucosyl transferase | ko:K13496 | UDPGT |
| AL3G03588 | UDP-glucosyl transferase | ko:K21374 | UDPGT |
| AL5G02905 | UDP-glucosyl transferase | ko:K21374 | UDPGT |
| AL7G00723 | UDP-glucosyl transferase | - | UDPGT |
| AL7G02582 | UDP-glucosyl transferase | - | UDPGT |
| AL5G03366 | Transferase family | ko:K13667 | Glyco_transf_90 |
| AL1G02765 | Transferase family | - | Glyco_tranf_2_3,Glyco_trans_2_3 |
| AL2G00685 | Transferase family | - | MBOAT_2 |
| AL2G00686 | Transferase family | - | MBOAT_2 |
| AL4G05546 | Transferase family | - | MBOAT_2 |
| AL1G05862 | Transferase family | - | Transferase |
| AL2G02767 | Transferase family | - | Transferase |
| AL4G00933 | Transferase family | - | Transferase |
| AL5G00326 | Transferase family | - | Transferase |
| AL5G00608 | Transferase family | - | Transferase |
| AL7G04448 | Transferase family | - | Transferase |
| AL2G01672 | Transferase family | ko:K13065 | Transferase |
| AL7G04341 | Transferase family | ko:K21374 | UDPGT |
| AL1G05758 | Heavy-metal-associated domain | - | HMA |
| AL1G05773 | Heavy-metal-associated domain | - | HMA |
| AL2G00203 | Heavy-metal-associated domain | - | HMA |
| AL6G03768 | Heavy-metal-associated domain | - | HMA |
| AL6G03777 | Heavy-metal-associated domain | - | HMA |
| AL6G03778 | Heavy-metal-associated domain | - | HMA |
| AL6G03781 | Heavy-metal-associated domain | - | HMA |
| AL6G03788 | Heavy-metal-associated domain | - | HMA |
| AL6G03789 | Heavy-metal-associated domain | - | HMA |
| AL6G03790 | Heavy-metal-associated domain | - | HMA |
| AL6G03791 | Heavy-metal-associated domain | - | HMA |
| AL6G03796 | Heavy-metal-associated domain | - | HMA |
| AL3G06175 | Disease resistance protein | - | NB-ARC |
| AL6G00670 | Disease resistance protein | - | - |

| Gene ID | Description | KEGG_ko | PFAMs |
|---|---|---|---|
| AL2G01320 | Disease resistance protein | - | NB-ARC |
| AL2G01411 | Disease resistance protein | - | NB-ARC |
| AL2G01413 | Disease resistance protein | - | NB-ARC |
| AL6G00120 | Disease resistance protein | - | NB-ARC |
| AL6G00122 | Disease resistance protein | - | NB-ARC |
| AL6G00673 | Disease resistance protein | - | NB-ARC |
| AL7G02087 | Disease resistance protein | - | NB-ARC |
| AL7G00966 | Ribosomal protein | ko:K02895 | KOW,ribosomal_L24 |
| AL3G02943 | Ribosomal protein | ko:K02933 | Ribosomal_L6 |
| AL1G01266 | Ribosomal protein | ko:K02902 | Ribosomal_L28 |
| AL3G00069 | Ribosomal protein | ko:K02927 | Ribosomal_L40e,ubiquitin |
| AL6G05072 | Ribosomal protein | ko:K02922 | Ribosomal_L37e |
| AL6G05088 | Ribosomal protein | ko:K02922 | Ribosomal_L37e |
| AL3G00056 | Ribosomal protein | - | RIP |
| AL7G00404 | Ribosomal protein | ko:K02908 | Ribosomal_L7Ae |
| AL7G00443 | Ribosomal protein | ko:K02908 | Ribosomal_L7Ae |
| AL1G06107 | Aldo/keto reductase family | - | Aldo_ket_red |
| AL1G06108 | Aldo/keto reductase family | - | Aldo_ket_red |
| AL1G06109 | Aldo/keto reductase family | - | Aldo_ket_red |
| AL6G05121 | Aldo/keto reductase family | - | Aldo_ket_red |
| AL6G05138 | Aldo/keto reductase family | - | Aldo_ket_red |
| AL2G04677 | Aldo/keto reductase family | ko:K00002 | Aldo_ket_red |
| AL2G04681 | Aldo/keto reductase family | ko:K00002 | Aldo_ket_red |
| AL1G00870 | Root cap | - | Root_cap |
| AL1G00871 | Root cap | - | Root_cap |
| AL1G00873 | Root cap | - | Root_cap |
| AL1G02397 | Root cap | - | Root_cap |
| AL2G01387 | Root cap | - | Root_cap |
| AL2G01416 | Root cap | - | Root_cap |
| AL5G00908 | Root cap | - | Root_cap |
| AL3G02021 | Sulfotransferase | ko:K22312 | Sulfotransfer_1 |
| AL3G02022 | Sulfotransferase | ko:K22312 | Sulfotransfer_1 |
| AL5G01793 | Sulfotransferase | ko:K22312 | Sulfotransfer_1 |
| AL6G05117 | Sulfotransferase | ko:K22312 | Sulfotransfer_1 |
| AL6G05118 | Sulfotransferase | ko:K22312 | Sulfotransfer_1 |
| AL6G05135 | Sulfotransferase | ko:K22312 | Sulfotransfer_1 |
| AL3G02020 | Sulfotransferase | - | Sulfotransfer_1 |
| AL2G00503 | Alpha/beta hydrolase | - | Abhydrolase_1,Abhydrolase_6 |
| AL2G00510 | Alpha/beta hydrolase | - | Abhydrolase_1,Abhydrolase_6 |
| AL5G01053 | Alpha/beta hydrolase | - | Abhydrolase_3 |
| AL5G01057 | Alpha/beta hydrolase | - | Abhydrolase_3 |
| AL4G02649 | Alpha/beta hydrolase | ko:K07874,ko:K 14493 | Abhydrolase_3 |
| AL1G02524 | Alpha/beta hydrolase | ko:K20802 | Abhydrolase_1,Abhydrolase_6 |
| AL6G03106 | Methyltransferase | - | Methyltransf_29 |
| AL6G01682 | Methyltransferase | - | Methyltrans_SAM |

| Gene ID | Description | KEGG_ko | PFAMs |
|---|---|---|---|
| AL5G01525 | Methyltransferase | ko:K21485 | Methyltransf_7 |
| AL2G05874 | UDP-glucosyl transferase | - | UDPGT |
| AL5G00006 | UDP-glucosyl transferase | - | UDPGT |
| AL5G00007 | UDP-glucosyl transferase | - | UDPGT |
| AL5G00044 | UDP-glucosyl transferase | - | UDPGT |
| AL6G05363 | UDP-glucosyl transferase | - | UDPGT |
| AL7G00016 | UDP-glucosyl transferase | - | UDPGT |
| AL7G00439 | UDP-glucosyl transferase | - | UDPGT |
| AL7G02074 | UDP-glucosyl transferase | - | UDPGT |
| AL2G00546 | UDP-glucosyl transferase | ko:K13495 | UDPGT |
| AL5G00001 | UDP-glucosyl transferase | ko:K13496 | UDPGT |
| AL3G03588 | UDP-glucosyl transferase | ko:K21374 | UDPGT |
| AL5G02905 | UDP-glucosyl transferase | ko:K21374 | UDPGT |
| AL7G00723 | UDP-glucosyl transferase | - | UDPGT |
| AL7G02582 | UDP-glucosyl transferase | - | UDPGT |
| AL5G03366 | Transferase family | ko:K13667 | Glyco_transf_90 |
| AL1G02765 | Transferase family | - | Glyco_tranf_2_3,Glyco_trans_2_3 |
| AL2G00685 | Transferase family | - | MBOAT_2 |
| AL2G00686 | Transferase family | - | MBOAT_2 |
| AL4G05546 | Transferase family | - | MBOAT_2 |
| AL1G05862 | Transferase family | - | Transferase |
| AL2G02767 | Transferase family | - | Transferase |
| AL4G00933 | Transferase family | - | Transferase |
| AL5G00326 | Transferase family | - | Transferase |
| AL5G00608 | Transferase family | - | Transferase |
| AL7G04448 | Transferase family | - | Transferase |
| AL2G01672 | Transferase family | ko:K13065 | Transferase |
| AL7G04341 | Transferase family | ko:K21374 | UDPGT |
| AL1G05758 | Heavy-metal-associated domain | - | HMA |
| AL1G05773 | Heavy-metal-associated domain | - | HMA |
| AL2G00203 | Heavy-metal-associated domain | - | HMA |
| AL6G03768 | Heavy-metal-associated domain | - | HMA |
| AL6G03777 | Heavy-metal-associated domain | - | HMA |
| AL6G03778 | Heavy-metal-associated domain | - | HMA |
| AL6G03781 | Heavy-metal-associated domain | - | HMA |
| AL6G03788 | Heavy-metal-associated domain | - | HMA |
| AL6G03789 | Heavy-metal-associated domain | - | HMA |

| Gene ID | Description | KEGG_ko | PFAMs |
|---|---|---|---|
| AL6G03790 | Heavy-metal-associated domain | - | HMA |
| AL6G03791 | Heavy-metal-associated domain | - | HMA |
| AL6G03796 | Heavy-metal-associated domain | - | HMA |
| AL3G06175 | Disease resistance protein | - | NB-ARC |
| AL6G00670 | Disease resistance protein | - | - |
| AL2G01320 | Disease resistance protein | - | NB-ARC |
| AL2G01411 | Disease resistance protein | - | NB-ARC |
| AL2G01413 | Disease resistance protein | - | NB-ARC |
| AL6G00120 | Disease resistance protein | - | NB-ARC |
| AL6G00122 | Disease resistance protein | - | NB-ARC |
| AL6G00673 | Disease resistance protein | - | NB-ARC |
| AL7G02087 | Disease resistance protein | - | NB-ARC |
| AL7G00966 | Ribosomal protein | ko:K02895 | KOW,ribosomal_L24 |
| AL3G02943 | Ribosomal protein | ko:K02933 | Ribosomal_L6 |
| AL1G01266 | Ribosomal protein | ko:K02902 | Ribosomal_L28 |
| AL3G00069 | Ribosomal protein | ko:K02927 | Ribosomal_L40e,ubiquitin |
| AL6G05072 | Ribosomal protein | ko:K02922 | Ribosomal_L37e |
| AL6G05088 | Ribosomal protein | ko:K02922 | Ribosomal_L37e |
| AL3G00056 | Ribosomal protein | - | RIP |
| AL7G00404 | Ribosomal protein | ko:K02908 | Ribosomal_L7Ae |
| AL7G00443 | Ribosomal protein | ko:K02908 | Ribosomal_L7Ae |
| AL1G06107 | Aldo/keto reductase family | - | Aldo_ket_red |
| AL1G06108 | Aldo/keto reductase family | - | Aldo_ket_red |
| AL1G06109 | Aldo/keto reductase family | - | Aldo_ket_red |
| AL6G05121 | Aldo/keto reductase family | - | Aldo_ket_red |
| AL6G05138 | Aldo/keto reductase family | - | Aldo_ket_red |
| AL2G04677 | Aldo/keto reductase family | ko:K00002 | Aldo_ket_red |
| AL2G04681 | Aldo/keto reductase family | ko:K00002 | Aldo_ket_red |
| AL1G00870 | Root cap | - | Root_cap |
| AL1G00871 | Root cap | - | Root_cap |
| AL1G00873 | Root cap | - | Root_cap |
| AL1G02397 | Root cap | - | Root_cap |
| AL2G01387 | Root cap | - | Root_cap |
| AL2G01416 | Root cap | - | Root_cap |
| AL5G00908 | Root cap | - | Root_cap |
| AL3G02021 | Sulfotransferase | ko:K22312 | Sulfotransfer_1 |
| AL3G02022 | Sulfotransferase | ko:K22312 | Sulfotransfer_1 |
| AL5G01793 | Sulfotransferase | ko:K22312 | Sulfotransfer_1 |
| AL6G05117 | Sulfotransferase | ko:K22312 | Sulfotransfer_1 |
| AL6G05118 | Sulfotransferase | ko:K22312 | Sulfotransfer_1 |
| AL6G05135 | Sulfotransferase | ko:K22312 | Sulfotransfer_1 |
| AL3G02020 | Sulfotransferase | - | Sulfotransfer_1 |
| AL2G00503 | Alpha/beta hydrolase | - | Abhydrolase_1,Abhydrolase_6 |

| Gene ID | Description | KEGG_ko | PFAMs |
|---|---|---|---|
| AL2G00510 | Alpha/beta hydrolase | - | Abhydrolase_1,Abhydrolase_6 |
| AL5G01053 | Alpha/beta hydrolase | - | Abhydrolase_3 |
| AL5G01057 | Alpha/beta hydrolase | - | Abhydrolase_3 |
| AL4G02649 | Alpha/beta hydrolase | ko:K07874,ko:K14493 | Abhydrolase_3 |
| AL1G02524 | Alpha/beta hydrolase | ko:K20802 | Abhydrolase_1,Abhydrolase_6 |
| AL5G02037 | DnaJ domain | - | DnaJ |
| AL3G01667 | DnaJ domain | - | DnaJ |
| AL3G01668 | DnaJ domain | - | DnaJ |
| AL3G01669 | DnaJ domain | - | DnaJ |
| AL7G00522 | DnaJ domain | - | DnaJ |
| AL7G00524 | DnaJ domain | - | DnaJ |
| AL1G02998 | GST superfamily | ko:K00799 | GST_C,GST_N |
| AL3G05623 | GST superfamily | ko:K00799 | GST_C,GST_N |
| AL4G00481 | GST superfamily | ko:K00799 | GST_C,GST_N |
| AL4G00507 | GST superfamily | ko:K00799 | GST_C,GST_N |
| AL5G00613 | GST superfamily | ko:K00799 | GST_C,GST_N |
| AL6G05320 | GST superfamily | ko:K00799 | GST_C,GST_N |
| AL2G00216 | Kinesin family | - | FERM_M,FERM_N,Kinesin,MyTH4 |
| AL2G00270 | Kinesin family | - | FERM_M,FERM_N,Kinesin,MyTH4 |
| AL6G04981 | Kinesin family | ko:K10398 | Kinesin |
| AL6G05003 | Kinesin family | ko:K10398 | Kinesin |
| AL4G01738 | Kinesin family | ko:K11498 | DUF3490,Kinesin |
| AL4G01751 | Kinesin family | ko:K11498 | DUF3490,Kinesin |
| AL1G00743 | NB-ARC domain | - | NB-ARC |
| AL1G01030 | NB-ARC domain | - | NB-ARC |
| AL1G01032 | NB-ARC domain | - | NB-ARC |
| AL1G01038 | NB-ARC domain | - | NB-ARC |
| AL1G01039 | NB-ARC domain | - | NB-ARC |
| AL6G00629 | NB-ARC domain | - | NB-ARC |
| AL2G02247 | Oxidation of toxic reductants | ko:K00430 | peroxidase |
| AL2G02562 | Oxidation of toxic reductants | ko:K00430 | peroxidase |
| AL4G02709 | Oxidation of toxic reductants | ko:K00430 | peroxidase |
| AL4G02723 | Oxidation of toxic reductants | ko:K00430 | peroxidase |
| AL4G03255 | Oxidation of toxic reductants | ko:K00430 | peroxidase |
| AL4G02708 | Oxidation of toxic reductants | ko:K00430 | peroxidase |
| AL4G03435 | Armadillo/beta-catenin-like repeats | ko:K08332 | Arm |
| AL4G03438 | Armadillo/beta-catenin-like repeats | ko:K08332 | Arm |
| AL3G01341 | Armadillo/beta-catenin-like repeats | ko:K06228 | HEAT,HEAT_2,Pkinase |
| AL4G02052 | Armadillo/beta-catenin-like repeats | ko:K06228 | HEAT,HEAT_2,Pkinase |
| AL4G02066 | Armadillo/beta-catenin-like repeats | ko:K06228 | HEAT,HEAT_2,Pkinase |

| Gene ID | Description | KEGG_ko | PFAMs |
|---|---|---|---|
| AL4G01632 | AP2 domain | - | AP2 |
| AL6G03215 | AP2 domain | - | AP2 |
| AL3G05083 | AP2 domain | ko:K09286 | AP2 |
| AL3G05084 | AP2 domain | ko:K09286 | AP2 |
| AL2G02436 | AP2 domain | ko:K14516 | AP2 |
| AL3G02021 | Sulfotransferase | ko:K22312 | Sulfotransfer_1 |
| AL3G02022 | Sulfotransferase | ko:K22312 | Sulfotransfer_1 |
| AL5G01793 | Sulfotransferase | ko:K22312 | Sulfotransfer_1 |
| AL6G05117 | Sulfotransferase | ko:K22312 | Sulfotransfer_1 |
| AL6G05118 | Sulfotransferase | ko:K22312 | Sulfotransfer_1 |
| AL6G05135 | Sulfotransferase | ko:K22312 | Sulfotransfer_1 |
| AL3G02020 | Sulfotransferase | - | Sulfotransfer_1 |
| AL2G00503 | Alpha/beta hydrolase | - | Abhydrolase_1,Abhydrolase_6 |
| AL2G00510 | Alpha/beta hydrolase | - | Abhydrolase_1,Abhydrolase_6 |
| AL5G01053 | Alpha/beta hydrolase | - | Abhydrolase_3 |
| AL5G01057 | Alpha/beta hydrolase | - | Abhydrolase_3 |
| AL4G02649 | Alpha/beta hydrolase | ko:K07874,ko:K14493 | Abhydrolase_3 |
| AL1G02524 | Alpha/beta hydrolase | ko:K20802 | Abhydrolase_1,Abhydrolase_6 |
| AL5G02037 | DnaJ domain | - | DnaJ |
| AL3G01667 | DnaJ domain | - | DnaJ |
| AL3G01668 | DnaJ domain | - | DnaJ |
| AL3G01669 | DnaJ domain | - | DnaJ |
| AL7G00522 | DnaJ domain | - | DnaJ |
| AL7G00524 | DnaJ domain | - | DnaJ |
| AL1G02998 | GST superfamily | ko:K00799 | GST_C,GST_N |
| AL3G05623 | GST superfamily | ko:K00799 | GST_C,GST_N |
| AL4G00481 | GST superfamily | ko:K00799 | GST_C,GST_N |
| AL4G00507 | GST superfamily | ko:K00799 | GST_C,GST_N |
| AL5G00613 | GST superfamily | ko:K00799 | GST_C,GST_N |
| AL6G05320 | GST superfamily | ko:K00799 | GST_C,GST_N |
| AL2G00216 | Kinesin family | - | FERM_M,FERM_N,Kinesin,MyTH4 |
| AL2G00270 | Kinesin family | - | FERM_M,FERM_N,Kinesin,MyTH4 |
| AL6G04981 | Kinesin family | ko:K10398 | Kinesin |
| AL6G05003 | Kinesin family | ko:K10398 | Kinesin |
| AL4G01738 | Kinesin family | ko:K11498 | DUF3490,Kinesin |
| AL4G01751 | Kinesin family | ko:K11498 | DUF3490,Kinesin |
| AL1G00743 | NB-ARC domain | - | NB-ARC |
| AL1G01030 | NB-ARC domain | - | NB-ARC |
| AL1G01032 | NB-ARC domain | - | NB-ARC |
| AL1G01038 | NB-ARC domain | - | NB-ARC |
| AL1G01039 | NB-ARC domain | - | NB-ARC |
| AL6G00629 | NB-ARC domain | - | NB-ARC |
| AL2G02247 | Oxidation of toxic reductants | ko:K00430 | peroxidase |
| AL2G02562 | Oxidation of toxic reductants | ko:K00430 | peroxidase |
| AL4G02709 | Oxidation of toxic reductants | ko:K00430 | peroxidase |

| Gene ID | Description | KEGG_ko | PFAMs |
|---|---|---|---|
| AL4G02723 | Oxidation of toxic reductants | ko:K00430 | peroxidase |
| AL4G03255 | Oxidation of toxic reductants | ko:K00430 | peroxidase |
| AL4G02708 | Oxidation of toxic reductants | ko:K00430 | peroxidase |
| AL4G03435 | Armadillo/beta-catenin-like repeats | ko:K08332 | Arm |
| AL4G03438 | Armadillo/beta-catenin-like repeats | ko:K08332 | Arm |
| AL3G01341 | Armadillo/beta-catenin-like repeats | ko:K06228 | HEAT,HEAT_2,Pkinase |
| AL4G02052 | Armadillo/beta-catenin-like repeats | ko:K06228 | HEAT,HEAT_2,Pkinase |
| AL4G02066 | Armadillo/beta-catenin-like repeats | ko:K06228 | HEAT,HEAT_2,Pkinase |
| AL4G01632 | AP2 domain | - | AP2 |
| AL6G03215 | AP2 domain | - | AP2 |
| AL3G05083 | AP2 domain | ko:K09286 | AP2 |
| AL3G05084 | AP2 domain | ko:K09286 | AP2 |
| AL2G02436 | AP2 domain | ko:K14516 | AP2 |
| 119 up-regulation DEGs (gene number ≥ 5 for each category) in expanded gene families of salt-treated leaves of ALO | | | |
| AL2G00501 | Zinc finger protein | - | zf-UBR |
| AL7G00708 | Zinc finger protein | - | zf-RING_2 |
| AL7G00740 | Zinc finger protein | - | zf-RING_2 |
| AL6G03437 | Zinc finger protein | ko:K10666 | zf-C3HC4,zf-C3HC4_2,zf-C3HC4_3 |
| AL6G03446 | Zinc finger protein | ko:K10666 | zf-C3HC4,zf-C3HC4_2,zf-C3HC4_3 |
| AL1G01511 | Zinc finger protein | ko:K16284 | zf-RING_2 |
| AL6G01184 | Zinc finger protein | ko:K06874,ko:K09025 | zf-ZPR1 |
| AL4G04841 | Zinc finger protein | ko:K19042 | zf-C3HC4_3 |
| AL6G03610 | Zinc finger protein | - | 4F5,zf-met |
| AL6G03112 | Zinc finger protein | - | zf-Dof |
| AL6G03114 | Zinc finger protein | - | zf-Dof |
| AL6G01478 | Zinc finger protein | - | zf-CCCH |
| AL6G01485 | Zinc finger protein | - | zf-CCCH |
| AL4G01212 | Zinc finger protein | - | zf-FLZ |
| AL4G01219 | Zinc finger protein | - | zf-FLZ |
| AL4G01221 | Zinc finger protein | - | zf-FLZ |
| AL4G01230 | Zinc finger protein | - | zf-FLZ |
| AL4G01239 | Zinc finger protein | - | zf-FLZ |
| AL4G01243 | Zinc finger protein | - | zf-FLZ |
| AL1G00162 | Other | - | Other |
| AL1G01153 | Other | - | Other |
| AL4G00494 | Other | - | Other |
| AL4G00521 | Other | - | Other |
| AL6G04860 | Other | - | Other |
| AL6G04876 | Other | - | Other |
| AL6G04892 | Other | - | Other |

| Gene ID | Description | KEGG_ko | PFAMs |
|---|---|---|---|
| AL7G02160 | Other | - | Other |
| AL7G00979 | Other | - | DUF223 |
| AL1G04911 | Other | - | F-box |
| AL2G05605 | Other | - | F-box |
| AL7G00881 | Other | - | Hemopexin |
| AL7G00050 | Other | - | LRR_6 |
| AL1G05385 | Other | - | NB-ARC |
| AL5G00787 | Other | - | Transposase_28 |
| AL6G04651 | Other | - | Transposase_28 |
| AL1G02094 | Other | - | ubiquitin |
| AL1G02106 | Other | - | ubiquitin |
| AL4G05035 | Protein kinase | ko:K14498 | Pkinase |
| AL4G05046 | Protein kinase | ko:K14498 | Pkinase |
| AL1G05216 | Protein kinase | - | LRRNT_2,LRR_1,LRR_4,LRR_8,Pkinase |
| AL6G00931 | Protein kinase | ko:K20538 | Pkinase |
| AL3G03953 | Protein kinase | ko:K04688 | Pkinase,Pkinase_C |
| AL1G00127 | Protein kinase | - | Pkinase |
| AL1G06336 | Protein kinase | - | Pkinase,Stress-antifung |
| AL1G05675 | Protein kinase | ko:K04733 | Pkinase |
| AL4G01721 | Protein kinase | ko:K04733 | Pkinase |
| AL4G01727 | Protein kinase | ko:K04733 | Pkinase |
| AL3G06323 | Protein kinase | ko:K08269 | Pkinase |
| AL1G03759 | Protein kinase | ko:K08287 | Pkinase |
| AL3G00360 | Protein kinase | - | B_lectin,PAN_2,Pkinase,Pkinase_Tyr,S_locus_glycop |
| AL6G04702 | Protein kinase | - | Pkinase,Stress-antifung |
| AL4G00222 | Protein kinase | ko:K04733 | Pkinase,Pkinase_Tyr |
| AL5G03021 | Protein kinase | ko:K04733 | Pkinase,Pkinase_Tyr |
| AL7G00155 | Protein kinase | ko:K04733 | Pkinase,Stress-antifung |
| AL1G02668 | Glycosyl hydrolase | ko:K01188,ko:K19964 | Glyco_hydro_1 |
| AL6G04035 | Glycosyl hydrolase | ko:K01188,ko:K19964 | Glyco_hydro_1 |
| AL4G02538 | Glycosyl hydrolase | ko:K05350 | Glyco_hydro_1 |
| AL4G01862 | Glycosyl hydrolase | ko:K05350 | Glyco_hydro_1,Pectinesterase |
| AL3G04784 | Glycosyl hydrolase | ko:K01176 | Alpha-amyl_C2,Alpha-amylase |
| AL1G02228 | Glycosyl hydrolase | - | Glyco_hydro_17 |
| AL1G06032 | Glycosyl hydrolase | ko:K01193 | Glyco_hydro_32C,Glyco_hydro_32N |
| AL4G00122 | Glycosyl hydrolase | ko:K01193 | Glyco_hydro_32C,Glyco_hydro_32N |
| AL1G00848 | Glycosyl hydrolase | ko:K01193,ko:K20848 | Glyco_hydro_32C,Glyco_hydro_32N |
| AL5G00116 | Glycosyl hydrolase | ko:K21351 | DUF3357,Glyco_hydro_32C,Glyco_hydro_32N |
| AL5G00117 | Glycosyl hydrolase | ko:K21351 | DUF3357,Glyco_hydro_32C,Glyco_hydro_32N |
| AL7G02632 | Glycosyl hydrolase | ko:K01188,ko:K05349 | Glyco_hydro_3,Glyco_hydro_3_C |
| AL6G00982 | Peptidase | ko:K01369 | Peptidase_C13 |
| AL6G01283 | Peptidase | ko:K01369 | Peptidase_C13 |

| Gene ID | Description | KEGG_ko | PFAMs |
|---|---|---|---|
| AL2G00726 | Peptidase | - | Peptidase_C65 |
| AL1G05839 | Peptidase | - | TAXi_C,TAXi_N |
| AL1G05850 | Peptidase | - | TAXi_C,TAXi_N |
| AL3G06247 | Peptidase | ko:K01363,ko:K01365,ko:K16290 | Granulin,Inhibitor_I29,Peptidase_C1 |
| AL3G03426 | Peptidase | ko:K09756,ko:K16296 | Peptidase_S10 |
| AL4G01608 | Peptidase | ko:K09756,ko:K16296 | Peptidase_S10 |
| AL4G03768 | Peptidase | ko:K09756,ko:K16296 | Peptidase_S10 |
| AL7G03749 | Peptidase | ko:K09756,ko:K16296 | Peptidase_S10 |
| AL3G01119 | Peptidase | ko:K16297 | Peptidase_S10 |
| AL3G05880 | Peptidase | ko:K00430 | peroxidase |
| AL2G05413 | RNA recognition motif | ko:K11294 | RRM_1 |
| AL1G01867 | RNA recognition motif | ko:K12741 | RRM_1 |
| AL4G00900 | RNA recognition motif | ko:K14787 | RRM_1 |
| AL1G00052 | RNA recognition motif | - | RRM_1 |
| AL1G01970 | RNA recognition motif | - | RRM_1 |
| AL1G01983 | RNA recognition motif | - | RRM_1 |
| AL6G00269 | RNA recognition motif | ko:K12893 | RRM_1,RrnaAD |
| AL6G05147 | Methyltransferase | ko:K21806 | Methyltransf_16 |
| AL3G00348 | Methyltransferase | - | Dimerisation,Methyltransf_2 |
| AL2G05970 | Methyltransferase | ko:K13259,ko:K13262,ko:K16040 | Dimerisation,Methyltransf_2 |
| AL6G00277 | Methyltransferase | - | RrnaAD |
| AL5G00056 | Methyltransferase | ko:K08241,ko:K21482,ko:K21483,ko:K21484 | Methyltransf_7 |
| AL5G01262 | Methyltransferase | ko:K08241,ko:K21482,ko:K21483,ko:K21484,ko:K21485 | Methyltransf_7 |
| AL5G01264 | Methyltransferase | ko:K08241,ko:K21482,ko:K21483,ko:K21484,ko:K21485 | Methyltransf_7 |
| AL4G01027 | EF hand family protein | - | - |
| AL4G01049 | EF hand family protein | - | - |
| AL7G02413 | EF hand family protein | ko:K13448,ko:K16465 | EF-hand_1,EF-hand_5,EF-hand_6,EF-hand_7 |
| AL4G05780 | EF hand family protein | - | EF-hand_1,EF-hand_5,EF-hand_8 |
| AL6G00251 | EF hand family protein | ko:K02183,ko:K | EF-hand_8 |

| Gene ID | Description | KEGG_ko | PFAMs |
|---|---|---|---|
| | | 10840,ko:K16465 | |
| AL6G00282 | EF hand family protein | ko:K02183,ko:K10840,ko:K16465 | EF-hand_8 |
| AL2G02603 | EamA-like transporter | - | EamA |
| AL5G00062 | EamA-like transporter | - | EamA |
| AL6G01126 | EamA-like transporter | - | EamA |
| AL6G01155 | EamA-like transporter | - | EamA |
| AL6G01125 | EamA-like transporter | ko:K02940 | EamA,Ribosomal_L6 |
| AL6G01156 | EamA-like transporter | ko:K02940 | EamA,Ribosomal_L6 |
| AL1G01674 | Heavy-metal-associated domain | - | HMA |
| AL1G01703 | Heavy-metal-associated domain | - | HMA |
| AL2G00627 | Heavy-metal-associated domain | - | HMA |
| AL3G05421 | Heavy-metal-associated domain | - | HMA |
| AL3G06027 | Heavy-metal-associated domain | - | HMA |
| AL1G01272 | No apical meristem (NAM) protein | - | NAM |
| AL1G01812 | No apical meristem (NAM) protein | - | NAM |
| AL1G01825 | No apical meristem (NAM) protein | - | NAM |
| AL2G04030 | No apical meristem (NAM) protein | - | NAM |
| AL6G01508 | No apical meristem (NAM) protein | - | NAM |
| AL3G02922 | Cytochrome P450 | ko:K00512 | p450 |
| AL7G05063 | Cytochrome P450 | ko:K00512 | p450 |
| AL4G02874 | Cytochrome P450 | ko:K10717,ko:K20660 | p450 |
| AL2G01841 | Cytochrome P450 | ko:K20556 | p450 |
| AL4G01386 | Cytochrome P450 | - | p450 |
| 252 down-regulation DEGs (gene number ≥ 5 for each category) in expanded gene families of of salt-treated leaves of ALO | | | |
| AL3G01135 | Protein kinase | - | EGF_CA,GUB_WAK_bind,Pkinase,Pkinase_Tyr |
| AL3G01147 | Protein kinase | - | EGF_CA,GUB_WAK_bind,Pkinase,Pkinase_Tyr |
| AL2G02268 | Protein kinase | - | Pkinase,VWA_3 |
| AL6G03364 | Protein kinase | - | LRRNT_2,LRR_1,LRR_4,LRR_6,LRR_8,Pkinase |
| AL6G01321 | Protein kinase | - | LRRNT_2,LRR_1,LRR_8,Pkinase,Pkinase_Tyr |
| AL1G00584 | Protein kinase | - | Pkinase_Tyr,Stress-antifung |
| AL3G02244 | Protein kinase | - | LRRNT_2,LRR_1,LRR_4,LRR_8,Pkinase,Pkinase_Tyr |
| AL6G00994 | Protein kinase | - | LRRNT_2,LRR_1,LRR_4,LRR_8,Pkinase,Pkinase_Tyr |

| Gene ID | Description | KEGG_ko | PFAMs |
|---|---|---|---|
| AL4G00715 | Protein kinase | - | LRR_8,Malectin,Pkinase,Pkinase_Tyr |
| AL2G05824 | Protein kinase | ko:K04733 | LRRNT_2,LRR_1,LRR_4,LRR_8,Pkinase,Pkinase_Tyr |
| AL1G03017 | Protein kinase | ko:K07198 | KA1,Pkinase,UBA |
| AL2G00785 | Protein kinase | ko:K07198,ko:K12761 | NAF,Pkinase |
| AL2G00786 | Protein kinase | ko:K07198,ko:K12761 | NAF,Pkinase |
| AL1G00226 | Protein kinase | - | LRRNT_2,LRR_1,LRR_6,LRR_8 |
| AL4G00501 | Protein kinase | - | B_lectin,PAN_2,Pkinase,Pkinase_Tyr,S_locus_glycop |
| AL5G00188 | Protein kinase | - | DUF3403,Pkinase,Pkinase_Tyr,Stress-antifung |
| AL7G01202 | Protein kinase | - | EGF_CA,GUB_WAK_bind,Pkinase |
| AL3G06318 | Protein kinase | - | GUB_WAK_bind,Pkinase |
| AL5G04122 | Protein kinase | - | GUB_WAK_bind,Pkinase |
| AL5G04123 | Protein kinase | - | GUB_WAK_bind,Pkinase |
| AL5G04128 | Protein kinase | - | GUB_WAK_bind,Pkinase |
| AL5G04129 | Protein kinase | - | GUB_WAK_bind,Pkinase |
| AL5G04131 | Protein kinase | - | GUB_WAK_bind,Pkinase,Pkinase_Tyr |
| AL6G03380 | Protein kinase | - | GUB_WAK_bind,Pkinase,Pkinase_Tyr |
| AL4G04131 | Protein kinase | - | LRRNT_2,LRR_1,LRR_8,Pkinase,Pkinase_Tyr |
| AL7G01812 | Protein kinase | - | Pkinase,Pkinase_Tyr |
| AL7G01810 | Protein kinase | ko:K04730 | Pkinase,Pkinase_Tyr |
| AL7G00389 | Protein kinase | ko:K04733 | LRRNT_2,LRR_8,Pkinase_Tyr |
| AL4G05615 | Protein kinase | ko:K20714 | Pkinase |
| AL4G05624 | Protein kinase | ko:K20714 | Pkinase |
| AL4G00459 | Protein kinase | - | B_lectin,DUF3403,PAN_2,Pkinase_Tyr,S_locus_glycop |
| AL5G02955 | Protein kinase | - | B_lectin,PAN_2,Pkinase_Tyr,S_locus_glycop |
| AL2G05878 | Protein kinase | ko:K04733 | B_lectin,PAN_2,Pkinase,S_locus_glycop |
| AL4G02960 | Protein kinase | ko:K00924 | Pkinase |
| AL4G02971 | Protein kinase | ko:K00924 | Pkinase |
| AL4G05616 | Protein kinase | ko:K08818 | Pkinase |
| AL7G03798 | Protein kinase | - | Pkinase,Pkinase_Tyr |
| AL1G00530 | Cytochrome P450 | - | p450 |
| AL6G01662 | Cytochrome P450 | - | p450 |
| AL6G01857 | Cytochrome P450 | - | p450 |
| AL6G01858 | Cytochrome P450 | - | p450 |
| AL7G00655 | Cytochrome P450 | - | p450 |
| AL7G00656 | Cytochrome P450 | - | p450 |
| AL1G01054 | Cytochrome P450 | ko:K00512 | p450 |
| AL1G02436 | Cytochrome P450 | ko:K00512 | p450 |
| AL1G05995 | Cytochrome P450 | ko:K00512 | p450 |
| AL1G06071 | Cytochrome P450 | ko:K00512 | p450 |
| AL2G04963 | Cytochrome P450 | ko:K00512 | p450 |
| AL2G04965 | Cytochrome P450 | ko:K00512 | p450 |
| AL4G01671 | Cytochrome P450 | ko:K00512 | p450 |
| AL2G04267 | Cytochrome P450 | ko:K00512,ko:K07408 | p450 |

| Gene ID | Description | KEGG_ko | PFAMs |
|---|---|---|---|
| AL5G00395 | Cytochrome P450 | ko:K00512,ko:K07410 | p450 |
| AL3G04730 | Cytochrome P450 | ko:K01723 | p450 |
| AL3G04731 | Cytochrome P450 | ko:K01723 | p450 |
| AL1G02016 | Other | - | Other |
| AL2G01567 | Other | - | Other |
| AL2G01582 | Other | - | Other |
| AL3G02097 | Other | - | Other |
| AL3G05801 | Other | - | Other |
| AL3G05845 | Other | - | Other |
| AL4G04150 | Other | - | Other |
| AL4G05039 | Other | - | Other |
| AL6G00767 | Other | - | Other |
| AL6G00871 | Other | - | Other |
| AL6G01853 | Other | - | Other |
| AL6G02314 | Other | - | Other |
| AL6G05075 | Other | - | Other |
| AL1G05869 | Other | - | NPR1_interact |
| AL4G01899 | Other | - | TPR_8 |
| AL6G03658 | Other | ko:K20478 | Other |
| AL4G01411 | Other | - | DUF810 |
| AL4G00816 | Other | - | DUF2358 |
| AL4G01338 | Other | - | DUF2358 |
| AL4G01354 | Other | - | DUF2358 |
| AL5G00074 | Peptidase family | - | TAXi_C,TAXi_N |
| AL6G00569 | Peptidase family | - | TAXi_C,TAXi_N |
| AL2G05235 | Peptidase family | ko:K01381 | TAXi_C,TAXi_N |
| AL3G06242 | Peptidase family | ko:K01363,ko:K01365,ko:K16290 | Granulin,Inhibitor_I29,Peptidase_C1 |
| AL2G05385 | Peptidase family | ko:K01365 | Inhibitor_I29,Peptidase_C1 |
| AL2G05386 | Peptidase family | ko:K01365 | Inhibitor_I29,Peptidase_C1 |
| AL2G05387 | Peptidase family | ko:K01365 | Inhibitor_I29,Peptidase_C1 |
| AL2G05389 | Peptidase family | ko:K01365 | Inhibitor_I29,Peptidase_C1 |
| AL4G02414 | Peptidase family | ko:K11841 | UCH |
| AL4G03348 | Peptidase family | ko:K01408 | Peptidase_M16,Peptidase_M16_C,Peptidase_M16_M |
| AL3G00594 | Peptidase family | ko:K16296 | Peptidase_S10 |
| AL2G00781 | Peptidase family | ko:K00430 | peroxidase |
| AL4G00949 | Peptidase family | ko:K00430 | peroxidase |
| AL4G05382 | Peptidase family | ko:K00430 | peroxidase |
| AL4G01264 | Receptor kinase | - | Lectin_legB,Pkinase,Pkinase_Tyr |
| AL3G05667 | Receptor kinase | ko:K04733 | Glyco_hydro_19,Pkinase |
| AL4G04542 | Receptor kinase | - | GUB_WAK_bind,Pkinase |
| AL4G04545 | Receptor kinase | - | GUB_WAK_bind,Pkinase |
| AL4G04552 | Receptor kinase | - | GUB_WAK_bind,Pkinase |
| AL4G04553 | Receptor kinase | - | GUB_WAK_bind,Pkinase |

| Gene ID | Description | KEGG_ko | PFAMs |
|---|---|---|---|
| AL4G04555 | Receptor kinase | - | GUB_WAK_bind,Pkinase |
| AL6G00142 | Receptor kinase | ko:K04733 | GUB_WAK_bind,Pkinase |
| AL6G00146 | Receptor kinase | ko:K04733 | GUB_WAK_bind,Pkinase |
| AL1G03251 | Receptor kinase | - | GUB_WAK_bind,WAK_assoc |
| AL1G03254 | Receptor kinase | - | GUB_WAK_bind,WAK_assoc |
| AL6G00383 | Receptor kinase | - | EGF_CA,GUB_WAK_bind,Pkinase,Pkinase_Tyr,WAK |
| AL6G00388 | Receptor kinase | - | EGF_CA,GUB_WAK_bind,Pkinase,Pkinase_Tyr,WAK |
| AL1G05980 | Receptor kinase | - | GUB_WAK_bind,Pkinase |
| AL1G06002 | Receptor kinase | - | GUB_WAK_bind,Pkinase,Pkinase_Tyr,WAK_assoc |
| AL2G02088 | Light-harvesting complex (LHC) | ko:K08912 | Chloroa_b-bind |
| AL2G02130 | Light-harvesting complex (LHC) | ko:K08912 | Chloroa_b-bind |
| AL2G02132 | Light-harvesting complex (LHC) | ko:K08912 | Chloroa_b-bind |
| AL3G00362 | Light-harvesting complex (LHC) | ko:K08912 | Chloroa_b-bind |
| AL3G03238 | Light-harvesting complex (LHC) | ko:K08912 | Chloroa_b-bind |
| AL5G02082 | Light-harvesting complex (LHC) | ko:K08912 | Chloroa_b-bind |
| AL6G01876 | Light-harvesting complex (LHC) | ko:K08912 | Chloroa_b-bind |
| AL6G01880 | Light-harvesting complex (LHC) | ko:K08912 | Chloroa_b-bind |
| AL6G01882 | Light-harvesting complex (LHC) | ko:K08912 | Chloroa_b-bind |
| AL2G05460 | Light-harvesting complex (LHC) | ko:K08912,ko:K08913 | Chloroa_b-bind |
| AL2G05461 | Light-harvesting complex (LHC) | ko:K08912,ko:K08913 | Chloroa_b-bind |
| AL6G01875 | Light-harvesting complex (LHC) | ko:K08912,ko:K08913 | Chloroa_b-bind |
| AL6G01878 | Light-harvesting complex (LHC) | ko:K08912,ko:K08913 | Chloroa_b-bind |
| AL4G05649 | Zinc finger protein | - | zf-B_box |
| AL4G02245 | Zinc finger protein | - | zf-CW |
| AL4G02256 | Zinc finger protein | - | zf-CW |
| AL1G04255 | Zinc finger protein | - | zf-Dof |
| AL2G00860 | Zinc finger protein | - | zf-Dof |
| AL4G01987 | Zinc finger protein | - | zf-Dof |
| AL6G04216 | Zinc finger protein | ko:K11380 | PHD_2,zf-HC5HC2H_2 |
| AL6G03896 | Zinc finger protein | ko:K10664,ko:K19041 | zf-RING_2 |
| AL6G03907 | Zinc finger protein | ko:K10664,ko:K19041 | zf-RING_2 |

| Gene ID | Description | KEGG_ko | PFAMs |
|---|---|---|---|
| AL2G02085 | Zinc finger protein | - | Ank_2,zf-CCCH |
| AL2G02125 | Zinc finger protein | - | Ank_2,zf-CCCH |
| AL1G02021 | ABC transporter | ko:K05656 | ABC_membrane,ABC_tran |
| AL1G02034 | ABC transporter | ko:K05656 | ABC_membrane,ABC_tran |
| AL5G00652 | ABC transporter | ko:K05666 | ABC_membrane,ABC_tran |
| AL4G01186 | ABC transporter | ko:K05665,ko:K05666,ko:K05670 | ABC_membrane,ABC_tran |
| AL6G04805 | ABC transporter | ko:K05665 | ABC_membrane,ABC_tran |
| AL6G04822 | ABC transporter | ko:K05665 | ABC_membrane,ABC_tran |
| AL2G01118 | ABC transporter | ko:K05665,ko:K05666 | ABC_membrane,ABC_tran |
| AL5G00657 | ABC transporter | ko:K05666 | ABC_membrane,ABC_tran |
| AL3G05315 | ABC transporter | - | ABC2_membrane,ABC_tran |
| AL3G05368 | ABC transporter | - | ABC2_membrane,ABC_tran |
| AL2G00258 | Disease resistance protein | - | LRR_8,NB-ARC |
| AL7G05033 | Disease resistance protein | - | LRR_8,NB-ARC |
| AL7G05036 | Disease resistance protein | - | LRR_8,NB-ARC |
| AL7G05037 | Disease resistance protein | - | LRR_8,NB-ARC |
| AL7G05038 | Disease resistance protein | - | LRR_8,NB-ARC |
| AL7G05040 | Disease resistance protein | - | LRR_8,NB-ARC |
| AL2G01224 | Disease resistance protein | - | NB-ARC |
| AL2G01305 | Disease resistance protein | - | NB-ARC |
| AL6G00107 | Disease resistance protein | - | NB-ARC |
| AL6G00113 | Disease resistance protein | - | NB-ARC |
| AL2G00359 | Oxidoreductase family | - | 2OG-FeII_Oxy,DIOX_N |
| AL3G01479 | Oxidoreductase family | - | 2OG-FeII_Oxy,DIOX_N |
| AL3G01482 | Oxidoreductase family | - | 2OG-FeII_Oxy,DIOX_N |
| AL3G04344 | Oxidoreductase family | - | 2OG-FeII_Oxy,DIOX_N |
| AL3G04345 | Oxidoreductase family | - | 2OG-FeII_Oxy,DIOX_N |
| AL3G05833 | Oxidoreductase family | - | 2OG-FeII_Oxy,DIOX_N |
| AL4G04537 | Oxidoreductase family | - | 2OG-FeII_Oxy,DIOX_N |
| AL4G04560 | Oxidoreductase family | - | 2OG-FeII_Oxy,DIOX_N |
| AL5G00418 | Oxidoreductase family | - | 2OG-FeII_Oxy,DIOX_N |
| AL6G04273 | Oxidoreductase family | ko:K05933 | 2OG-FeII_Oxy,DIOX_N |
| AL2G02136 | Legume lectin domain | - | Lectin_legB,Pkinase |
| AL4G04504 | Legume lectin domain | - | Lectin_legB,Pkinase |
| AL6G00940 | Legume lectin domain | - | Lectin_legB,Pkinase |
| AL2G01227 | Legume lectin domain | - | Lectin_legB,Pkinase,Pkinase_Tyr |
| AL2G01228 | Legume lectin domain | - | Lectin_legB,Pkinase,Pkinase_Tyr |
| AL2G01245 | Legume lectin domain | - | Lectin_legB,Pkinase,Pkinase_Tyr |
| AL2G02097 | Legume lectin domain | - | Lectin_legB,Pkinase,Pkinase_Tyr |
| AL4G04295 | Legume lectin domain | - | Lectin_legB,Pkinase,Pkinase_Tyr |
| AL6G00937 | Legume lectin domain | - | Lectin_legB,Pkinase,Pkinase_Tyr |
| AL4G01500 | MIP aquaporin family | ko:K09872 | MIP |
| AL4G03849 | MIP aquaporin family | ko:K09872 | MIP |

| Gene ID | Description | KEGG_ko | PFAMs |
|---|---|---|---|
| AL6G03347 | MIP aquaporin family | ko:K09872 | MIP |
| AL6G03348 | MIP aquaporin family | ko:K09872 | MIP |
| AL6G03361 | MIP aquaporin family | ko:K09872 | MIP |
| AL6G05247 | MIP aquaporin family | ko:K09872 | MIP |
| AL3G05484 | MIP aquaporin family | ko:K09873 | MIP |
| AL3G05485 | MIP aquaporin family | ko:K09873 | MIP |
| AL3G05493 | MIP aquaporin family | ko:K09873 | MIP |
| AL7G02586 | UDP-glucosyl transferase | - | UDPGT |
| AL2G01097 | UDP-glucosyl transferase | - | UDPGT |
| AL2G01147 | UDP-glucosyl transferase | - | UDPGT |
| AL6G01905 | UDP-glucosyl transferase | ko:K08236 | UDPGT |
| AL6G01912 | UDP-glucosyl transferase | ko:K08236 | UDPGT |
| AL2G00547 | UDP-glucosyl transferase | ko:K13495 | UDPGT |
| AL4G01547 | UDP-glucosyl transferase | ko:K13495 | UDPGT |
| AL2G00341 | UDP-glucosyl transferase | ko:K21374 | UDPGT |
| AL6G02593 | UDP-glucosyl transferase | ko:K21374 | UDPGT |
| AL3G01095 | Glycosyl transferase family | - | Cellulose_synt |
| AL3G01096 | Glycosyl transferase family | - | Cellulose_synt |
| AL5G00190 | Glycosyl transferase family | ko:K20924 | Cellulose_synt,zf-RING_4 |
| AL2G00834 | Glycosyl transferase family | ko:K10999 | Cellulose_synt,zf-UDP |
| AL2G00843 | Glycosyl transferase family | ko:K10999 | Cellulose_synt,zf-UDP |
| AL4G04642 | Glycosyl transferase family | ko:K09667 | Glyco_transf_41,LEA_5,TPR_1,TPR_11,TPR_16,TPR_7,TPR_8 |
| AL5G03371 | Glycosyl transferase family | ko:K13667 | Glyco_transf_90 |
| AL1G02765 | Glycosyl transferase family | - | Glyco_tranf_2_3,Glyco_trans_2_3 |
| AL1G05948 | EF-hand domain | ko:K13448 | EF-hand_1,EF-hand_5,EF-hand_6,EF-hand_7 |
| AL1G05958 | EF-hand domain | ko:K13448 | EF-hand_1,EF-hand_5,EF-hand_6,EF-hand_7 |
| AL1G05959 | EF-hand domain | ko:K13448 | EF-hand_1,EF-hand_5,EF-hand_6,EF-hand_7 |
| AL1G01220 | EF-hand domain | ko:K13448 | EF-hand_1,EF-hand_5,EF-hand_7 |
| AL1G01249 | EF-hand domain | ko:K13448 | EF-hand_1,EF-hand_5,EF-hand_7 |
| AL1G05417 | EF-hand domain | ko:K13448 | EF-hand_1,EF-hand_5,EF-hand_7 |
| AL3G04066 | EF-hand domain | ko:K02183,ko:K16465 | EF-hand_1,EF-hand_5,EF-hand_6,EF-hand_7 |
| AL2G00704 | Leucine rich repeat | - | LRRNT_2,LRR_1,LRR_4,LRR_6,LRR_8 |
| AL1G00228 | Leucine rich repeat | - | LRRNT_2,LRR_1,LRR_6,LRR_8 |
| AL1G00734 | Leucine rich repeat | - | LRRNT_2,LRR_1,LRR_6,LRR_8 |
| AL1G05885 | Leucine rich repeat | - | LRRNT_2,LRR_1,LRR_6,LRR_8 |
| AL1G05898 | Leucine rich repeat | - | LRRNT_2,LRR_1,LRR_6,LRR_8 |
| AL1G05092 | Leucine rich repeat | - | LRRNT_2,LRR_1,LRR_6,LRR_8 |
| AL6G01351 | Leucine rich repeat | - | LRRNT_2,LRR_1,LRR_6,LRR_8,Pkinase |
| AL3G03052 | Pathogenesis-related protein | - | Bet_v_1,Polyketide_cyc2 |
| AL3G03054 | Pathogenesis-related protein | - | Bet_v_1,Polyketide_cyc2 |
| AL3G03055 | Pathogenesis-related protein | - | Bet_v_1,Polyketide_cyc2 |
| AL4G02535 | Pathogenesis-related protein | - | Bet_v_1,Polyketide_cyc2 |
| AL4G02544 | Pathogenesis-related protein | - | Bet_v_1,Polyketide_cyc2 |
| AL4G02545 | Pathogenesis-related protein | - | Bet_v_1,Polyketide_cyc2 |

| Gene ID | Description | KEGG_ko | PFAMs |
|---|---|---|---|
| AL5G00218 | Pathogenesis-related protein | - | Bet_v_1,Polyketide_cyc2 |
| AL1G04895 | Transferase family | - | Transferase |
| AL5G00265 | Transferase family | - | Transferase |
| AL5G02926 | Transferase family | - | Transferase |
| AL5G02927 | Transferase family | - | Transferase |
| AL5G02410 | Transferase family | ko:K13065 | Transferase |
| AL2G03442 | Transferase family | ko:K19861 | Transferase |
| AL3G00727 | Transferase family | ko:K19861 | Transferase |
| AL1G01780 | Auxin responsive protein | ko:K14486 | AUX_IAA,Auxin_resp,B3 |
| AL4G05128 | Auxin responsive protein | ko:K14486 | AUX_IAA,Auxin_resp,B3 |
| AL4G05148 | Auxin responsive protein | ko:K14486 | AUX_IAA,Auxin_resp,B3 |
| AL4G00710 | Auxin responsive protein | ko:K14488 | Auxin_inducible |
| AL4G00726 | Auxin responsive protein | ko:K14488 | Auxin_inducible |
| AL7G01250 | Auxin responsive protein | ko:K14488 | Auxin_inducible |
| AL1G01288 | Glycosyl hydrolase family | - | Glyco_hydro_17 |
| AL1G01289 | Glycosyl hydrolase family | - | Glyco_hydro_17 |
| AL1G01290 | Glycosyl hydrolase family | - | Glyco_hydro_17 |
| AL1G01335 | Glycosyl hydrolase family | - | Glyco_hydro_17 |
| AL1G00849 | Glycosyl hydrolase family | ko:K01193,ko:K20848 | Glyco_hydro_32C,Glyco_hydro_32N |
| AL3G05930 | Glycosyl hydrolase family | - | Glyco_hydro_16 |
| AL5G03937 | Methyltransferase | ko:K13259,ko:K13262,ko:K16040 | Dimerisation,Methyltransf_2 |
| AL2G00363 | Methyltransferase | ko:K15332 | RRM_1,tRNA_U5-meth_tr,zf-CCCH |
| AL3G05929 | Methyltransferase | ko:K08241,ko:K21483 | Methyltransf_7 |
| AL2G01329 | Methyltransferase | ko:K08247 | Aminotran_1_2,MTS,Methyltransf_31 |
| AL2G01511 | Methyltransferase | ko:K08247 | Aminotran_1_2,MTS,Methyltransf_31 |
| AL6G02904 | Methyltransferase | ko:K21483,ko:K21485 | Methyltransf_7 |
| AL4G05217 | Pentatricopeptide repeat | - | PPR,PPR_1,PPR_2,PPR_3 |
| AL6G00393 | Pentatricopeptide repeat | - | PPR,PPR_1,PPR_2,PPR_3 |
| AL6G02707 | Pentatricopeptide repeat | ko:K17710 | PPR,PPR_1,PPR_2,PPR_3 |
| AL4G01288 | Pentatricopeptide repeat | - | PPR,PPR_2 |
| AL4G01300 | Pentatricopeptide repeat | - | PPR,PPR_2 |
| AL6G03622 | Pentatricopeptide repeat | - | PPR,PPR_2,PPR_3 |
| AL4G05219 | Plant lipoxygenase | ko:K00454 | Lipoxygenase,PLAT |
| AL6G01432 | Plant lipoxygenase | ko:K15718 | Lipoxygenase,PLAT |
| AL6G01434 | Plant lipoxygenase | ko:K15718 | Lipoxygenase,PLAT |
| AL6G01440 | Plant lipoxygenase | ko:K15718 | Lipoxygenase,PLAT |
| AL6G01443 | Plant lipoxygenase | ko:K15718 | Lipoxygenase,PLAT |

**Table S19. Characterization of biosynthetic gene clusters (BGCs) in the ALO genome.**

| Chromosome | Cluster number | Cluster type | Start position | End position | Core domain |
|---|---|---|---|---|---|
| ALO01 | Cluster1 | Saccharide | 42494028 | 42660011 | UDPGT_2, p450 |
| ALO01 | Cluster2 | Polyketide | 48240112 | 48811452 | COesterase, Chal_sti_synt_C, Chal_sti_synt_N, adh_short, p450 |
| ALO01 | Cluster3 | Saccharide | 123192768 | 123822687 | UDPGT_2, p450 |
| ALO01 | Cluster4 | Lignan | 553374586 | 553576950 | Dirigent, FA_desaturase, Transferase |
| ALO01 | Cluster5 | Terpene | 556126286 | 559744128 | SQHop_cyclase_C, SQHop_cyclase_N, p450 |
| ALO01 | Cluster6 | Saccharide | 566927392 | 567452204 | Methyltransf_2, UDPGT_2, p450 |
| ALO01 | Cluster7 | Saccharide-Polyketide | 567903409 | 568539773 | Chal_sti_synt_C, Chal_sti_synt_N, Methyltransf_2, Transferase, UDPGT_2, p450 |
| ALO01 | Cluster8 | Terpene | 570910888 | 571520560 | 2OG-FeII_Oxy, DIOX_N, Terpene_synth, Terpene_synth_C, p450 |
| ALO01 | Cluster9 | Saccharide-Terpene | 578834584 | 579031152 | Glycos_transf_1, Terpene_synth, Terpene_synth_C, p450 |
| ALO01 | Cluster10 | Saccharide-Terpene | 590248266 | 590876095 | Prenyltransf, Terpene_synth, Terpene_synth_C, UDPGT_2, p450 |
| ALO01 | Cluster11 | Terpene | 591110087 | 591350542 | Terpene_synth, Terpene_synth_C, p450 |
| ALO02 | Cluster12 | Lignan | 26190381 | 26753946 | 2OG-FeII_Oxy,Cellulose_synt,DIOX_N,Dirigent |
| ALO02 | Cluster13 | Lignan-Terpene | 26888043 | 27398024 | Cellulose_synt,DIOX_N,Dirigent,Terpene_synth,Terpene_synth_C |
| ALO02 | Cluster14 | Lignan-Polyketide | 28776812 | 29510398 | COesterase,Chal_sti_synt_C,Chal_sti_synt_N,Dirigent |
| ALO02 | Cluster15 | Lignan-Polyketide | 29752852 | 30430197 | 2OG-FeII_Oxy,COesterase,Chal_sti_synt_C,Chal_sti_synt_N,Dirigent |
| ALO02 | Cluster16 | Putative | 32099951 | 32294373 | 2OG-FeII_Oxy,DIOX_N,p450 |
| ALO02 | Cluster17 | Putative | 39987864 | 40333723 | 2OG-FeII_Oxy,Cellulose_synt,DIOX_N,NAD_binding_1,p450 |
| ALO02 | Cluster18 | Saccharide | 55139391 | 55313404 | Aminotran_1_2,Transferase,UDPGT_2 |
| ALO02 | Cluster19 | Saccharide | 57673820 | 58010344 | Aminotran_1_2,Transferase,UDPGT_2 |
| ALO02 | Cluster20 | Terpene | 70581320 | 71433642 | Methyltransf_7,Terpene_synth,Terpene_synth_C,p450 |
| ALO02 | Cluster21 | Terpene | 72140023 | 72417686 | Methyltransf_7,Peptidase_S10,Terpene_synth,Terpene_synth_C |
| ALO02 | Cluster22 | Saccharide | 76044812 | 76444221 | UDPGT_2,p450 |
| ALO02 | Cluster23 | Saccharide | 86139798 | 86237684 | Transferase,UDPGT_2 |
| ALO02 | Cluster24 | Lignan | 128865222 | 130141912 | Dirigent,GMC_oxred_C,GMC_oxred_N,Transferase |
| ALO02 | Cluster25 | Putative | 476168412 | 476593694 | AMP-binding,Amino_oxidase,Cellulose_synt,Methyltransf_11,p450 |

| Chromosome | Cluster number | Cluster type | Start position | End position | Core domain |
|---|---|---|---|---|---|
| ALO02 | Cluster26 | Putative | 544152235 | 544604500 | 2OG-FeII_Oxy,DIOX_N,Methyltransf_2,p450 |
| ALO02 | Cluster27 | Lignan | 546111627 | 546966190 | Dirigent,Epimerase |
| ALO02 | Cluster28 | Polyketide | 559319181 | 559547156 | Chal_sti_synt_C,Chal_sti_synt_N,Peptidase_S10,polyprenyl_synt |
| ALO03 | Cluster29 | Putative | 11066972 | 11567638 | Epimerase,NAD_binding_1,Transferase,p450 |
| ALO03 | Cluster30 | Putative | 55676950 | 56394933 | COesterase,adh_short |
| ALO03 | Cluster31 | Alkaloid | 169464495 | 170732104 | HMGL-like,Str_synth,p450 |
| ALO03 | Cluster32 | Alkaloid | 350295775 | 351189819 | Bet_v_1,Cellulose_synt |
| ALO03 | Cluster33 | Saccharide | 453017043 | 453438810 | Glycos_transf_1,Peptidase_S10 |
| ALO03 | Cluster34 | Lignan | 485718253 | 486148556 | Dirigent,Epimerase |
| ALO03 | Cluster35 | Terpene | 533508172 | 533742914 | 2OG-FeII_Oxy,Epimerase,Terpene_synth,Terpene_synth_C |
| ALO03 | Cluster36 | Saccharide-Polyketide | 536404827 | 537018053 | Chal_sti_synt_C,DIOX_N,UDPGT_2 |
| ALO03 | Cluster37 | Putative | 554782655 | 556137340 | AMP-binding,Aminotran_1_2,Epimerase,Transferase,p450 |
| ALO03 | Cluster38 | Polyketide | 571704299 | 572114406 | Chal_sti_synt_C,Chal_sti_synt_N,Methyltransf_2,p450 |
| ALO03 | Cluster39 | Terpene | 581303471 | 581566688 | Terpene_synth,Terpene_synth_C,p450 |
| ALO03 | Cluster40 | Terpene | 583725071 | 583884706 | Terpene_synth,Terpene_synth_C,p450 |
| ALO04 | Cluster41 | Saccharide | 584803 | 900953 | Glyco_transf_28,Transferase,UDPGT,p450 |
| ALO04 | Cluster42 | Putative | 1115900 | 1352008 | 2OG-FeII_Oxy,Transferase,p450 |
| ALO04 | Cluster43 | Lignan | 1797952 | 2057491 | Dirigent,Methyltransf_7,p450 |
| ALO04 | Cluster44 | Terpene | 2221799 | 2679970 | SQHop_cyclase_C,SQHop_cyclase_N,Terpene_synth,Terpene_synth_C,p450 |
| ALO04 | Cluster45 | Lignan-Terpene | 3063465 | 3237939 | 2OG-FeII_Oxy,DIOX_N,Dirigent,Terpene_synth,Terpene_synth_C |
| ALO04 | Cluster46 | Terpene | 3618856 | 4450463 | Terpene_synth,Terpene_synth_C,p450 |
| ALO04 | Cluster47 | Saccharide-Terpene | 19302657 | 19403033 | Terpene_synth,Terpene_synth_C,UDPGT_2 |
| ALO04 | Cluster48 | Terpene | 32474544 | 32570247 | Terpene_synth,Terpene_synth_C,p450 |
| ALO04 | Cluster49 | Terpene | 33405279 | 33570612 | Terpene_synth,Terpene_synth_C,p450 |
| ALO04 | Cluster50 | Terpene | 35311885 | 35576862 | Terpene_synth,Terpene_synth_C,Transferase,p450 |
| ALO04 | Cluster51 | Terpene | 36153938 | 36543946 | Terpene_synth,Terpene_synth_C,Transferase,p450 |
| ALO04 | Cluster52 | Saccharide | 46032283 | 46280232 | Epimerase,Glycos_transf_1,Transferase |
| ALO04 | Cluster53 | Polyketide | 76973489 | 77115266 | Chal_sti_synt_C,Chal_sti_synt_N,FA_desaturase |
| ALO04 | Cluster54 | Saccharide-Alkaloid | 91691952 | 91897096 | Bet_v_1,Methyltransf_11,UDPGT_2 |

| Chromosome | Cluster number | Cluster type | Start position | End position | Core domain |
|---|---|---|---|---|---|
| ALO04 | Cluster55 | Lignan | 102083403 | 102292760 | Dirigent,adh_short_C2 |
| ALO04 | Cluster56 | Lignan | 102393144 | 102610665 | Dirigent,adh_short_C2 |
| ALO04 | Cluster57 | Terpene | 111809090 | 112281106 | Methyltransf_11,Peptidase_S10,Terpene_synth,Terpene_synth_C |
| ALO04 | Cluster58 | Saccharide | 130556458 | 130802842 | 2OG-FeII_Oxy,DIOX_N,Glycos_transf_2 |
| ALO04 | Cluster59 | Lignan | 141698083 | 142117077 | Aminotran_1_2,Dirigent |
| ALO04 | Cluster60 | Polyketide | 147445808 | 148403757 | COesterase,Chal_sti_synt_C,Chal_sti_synt_N,Methyltransf_2 |
| ALO04 | Cluster61 | Saccharide | 411982519 | 414263928 | 2OG-FeII_Oxy,DIOX_N,UDPGT_2,adh_short_C2 |
| ALO04 | Cluster62 | Putative | 418015627 | 422620277 | COesterase,Methyltransf_11,adh_short_C2 |
| ALO04 | Cluster63 | Saccharide | 476688333 | 477787820 | AMP-binding,Peptidase_S10,UDPGT_2 |
| ALO04 | Cluster64 | Polyketide | 540996740 | 542295538 | Chal_sti_synt_C,Chal_sti_synt_N,Lipoxygenase,Methyltransf_2,p450 |
| ALO04 | Cluster65 | Alkaloid | 554560760 | 554827289 | Acetyltransf_1,Bet_v_1,Transferase |
| ALO04 | Cluster66 | Alkaloid | 555811081 | 556183079 | Acetyltransf_1,Bet_v_1,Transferase |
| ALO05 | Cluster67 | Saccharide-Terpene | 62875 | 2213952 | Aminotran_1_2,Methyltransf_2,Peptidase_S10,Prenyltrans,SQHop_cyclase_C,SQHop_cyclase_N,UDPGT_2,p450 |
| ALO05 | Cluster68 | Lignan-Saccharide | 3048813 | 3436280 | Dirigent,UDPGT_2,adh_short_C2,p450 |
| ALO05 | Cluster69 | Alkaloid | 3528145 | 3561828 | Bet_v_1,Cellulose_synt,Methyltransf_7 |
| ALO05 | Cluster70 | Putative | 5515019 | 5854552 | 2OG-FeII_Oxy,DIOX_N,Methyltransf_11,SQS_PSY,p450 |
| ALO05 | Cluster71 | Terpene | 6412245 | 6754187 | DIOX_N,FA_desaturase_2,Terpene_synth,Terpene_synth_C,p450 |
| ALO05 | Cluster72 | Saccharide-Terpene-Alkaloid | 8395566 | 9590354 | 2OG-FeII_Oxy,AMP-binding,Bet_v_1,Terpene_synth,Terpene_synth_C,UDPGT_2,p450 |
| ALO05 | Cluster73 | Polyketide | 9734044 | 10039623 | 2OG-FeII_Oxy,Chal_sti_synt_C,Chal_sti_synt_N,DIOX_N,Methyltransf_2 |
| ALO05 | Cluster74 | Terpene | 10401767 | 10627488 | Terpene_synth,Terpene_synth_C,p450 |
| ALO05 | Cluster75 | Terpene-Polyketide | 11567098 | 11715971 | Aminotran_1_2,Chal_sti_synt_C,Chal_sti_synt_N,Terpene_synth_C |
| ALO05 | Cluster76 | Saccharide | 16950522 | 17270435 | DIOX_N,Glycos_transf_1,p450 |
| ALO05 | Cluster77 | Putative | 17931357 | 18449217 | 2OG-FeII_Oxy,DIOX_N,Transferase,p450 |
| ALO05 | Cluster78 | Putative | 20595733 | 20867963 | Epimerase,Methyltransf_11,Transferase,p450 |
| ALO05 | Cluster79 | Saccharide | 26243750 | 26449647 | Glycos_transf_1,Transferase,p450 |
| ALO05 | Cluster80 | Saccharide-Polyketide | 28753425 | 29365649 | Chal_sti_synt_C,Epimerase,Methyltransf_11,Methyltransf_3,UDPGT_2 |

| Chromosome | Cluster number | Cluster type | Start position | End position | Core domain |
|---|---|---|---|---|---|
| ALO05 | Cluster81 | Terpene-Polyketide | 29686589 | 30183996 | Aminotran_1_2,Chal_sti_synt_C,Chal_sti_synt_N,Prenyltransf,Terpene_synth,Terpene_synth_C |
| ALO05 | Cluster82 | Putative | 32821558 | 33035296 | 2OG-FeII_Oxy,DIOX_N,Transferase,adh_short_C2,p450 |
| ALO05 | Cluster83 | Terpene | 34808873 | 35728583 | Prenyltrans,SQHop_cyclase_C,SQHop_cyclase_N,p450 |
| ALO05 | Cluster84 | Putative | 36727519 | 37963819 | Epimerase,Peptidase_S10,Transferase,adh_short_C2 |
| ALO05 | Cluster85 | Saccharide | 39212749 | 39654637 | 2OG-FeII_Oxy,DIOX_N,UDPGT,p450 |
| ALO05 | Cluster86 | Polyketide | 49655790 | 50017347 | Chal_sti_synt_C,Chal_sti_synt_N,p450 |
| ALO05 | Cluster87 | Lignan-Saccharide | 54880325 | 55298502 | Dirigent,Glycos_transf_1,Glycos_transf_2,UDPGT_2 |
| ALO05 | Cluster88 | Polyketide | 62229293 | 63719482 | Chal_sti_synt_C,ERG4_ERG24,p450 |
| ALO05 | Cluster89 | Putative | 153492475 | 156921470 | COesterase,adh_short,p450 |
| ALO05 | Cluster90 | Saccharide | 506473502 | 507671313 | UDPGT_2,adh_short_C2 |
| ALO05 | Cluster91 | Saccharide | 508375523 | 509506788 | Transferase,UDPGT_2 |
| ALO05 | Cluster92 | Lignan | 513477472 | 513742044 | 2OG-FeII_Oxy,DIOX_N,Dirigent |
| ALO05 | Cluster93 | Terpene | 517558826 | 517792552 | 2OG-FeII_Oxy,DIOX_N,Epimerase,Terpene_synth,Terpene_synth_C |
| ALO06 | Cluster94 | Saccharide | 25126332 | 25726598 | UDPGT_2,p450 |
| ALO06 | Cluster95 | Saccharide-Polyketide | 148576741 | 149527530 | Chal_sti_synt_C,UDPGT_2,p450 |
| ALO06 | Cluster96 | Saccharide | 287182672 | 289489225 | Amino_oxidase,Methyltransf_11,UDPGT_2 |
| ALO06 | Cluster97 | Putative | 479319285 | 479653041 | AMP-binding,adh_short,adh_short_C2,p450 |
| ALO06 | Cluster98 | Saccharide | 491235826 | 491443724 | UDPGT_2,p450 |
| ALO06 | Cluster99 | Putative | 491595358 | 491959958 | Acetyltransf_1,Methyltransf_7,Transferase,p450 |
| ALO06 | Cluster100 | Terpene-Polyketide | 494166248 | 494677580 | Chal_sti_synt_C,Chal_sti_synt_N,Epimerase,Terpene_synth,Terpene_synth_C,p450 |
| ALO06 | Cluster101 | Polyketide | 495239257 | 495487792 | Chal_sti_synt_C,Chal_sti_synt_N,p450 |
| ALO06 | Cluster102 | Saccharide | 508892932 | 509034736 | 2OG-FeII_Oxy,Cellulose_synt,UDPGT_2 |
| ALO07 | Cluster103 | Polyketide | 23082657 | 23190634 | Chal_sti_synt_C,Chal_sti_synt_N |
| ALO07 | Cluster104 | Saccharide | 85408254 | 85624976 | COesterase,Glycos_transf_2,UDPGT_2 |
| ALO07 | Cluster105 | Saccharide | 95090499 | 95432866 | SQS_PSY,UDPGT_2,adh_short |
| ALO07 | Cluster106 | Lignan | 171903467 | 172711624 | Dirigent,Methyltransf_2,p450 |
| ALO07 | Cluster107 | Saccharide-Alkaloid | 223109538 | 224257506 | AMP-binding,Str_synth,UDPGT_2 |
| ALO07 | Cluster108 | Terpene | 248357317 | 249456339 | SQS_PSY,Terpene_synth,Terpene_synth_C,p450 |
| ALO07 | Cluster109 | Polyketide | 449606117 | 452060832 | AMP-binding,Chal_sti_synt_C,adh_short |

**Table S20. The *CYP450* gene copy number in the studied species and distribution along the ALO chromosomes.**

**a The copy number of *CYP450* genes in the studied species.**

| Species | Copy number of CYP450 |
|---|---|
| ALO | 557 |
| AAT | 454 |
| AER | 444 |
| ATA | 413 |
| AST | 389 |
| SIT | 340 |
| OSA | 330 |
| SBI | 318 |
| ZMA | 287 |
| BDI | 251 |
| ATH | 246 |

**b Distribution of *CYP450* genes along the ALO chromosomes.**

| Chromosome no. | Number of CYP450 genes | Percentage of total CYP450 genes |
|---|---|---|
| ALO01 | 100 | 17.95% |
| ALO02 | 73 | 13.11% |
| ALO03 | 69 | 12.39% |
| ALO04 | 93 | 16.70% |
| ALO05 | 85 | 15.26% |
| ALO06 | 83 | 14.90% |
| ALO07 | 84 | 15.08% |
| Total | 557 | 100% |

**Table S21. FPKM value of *CYP450* genes in roots, leaves, salt-treated roots and salt-treated leaves of ALO.**

**a The transcript levels of *CYP450* genes over 10-fold higher in salt-treated roots than in roots.**

| Gene ID | Roots | Salt-treated roots | Fold number (Salt-treated roots / roots) | Leaves | Salt-treated leaves |
|---|---|---|---|---|---|
| AL2G04509 | 0.03 | 28.52 | 912.82 | 0.00 | 0.01 |
| AL2G04508 | 0.06 | 37.58 | 655.34 | 0.00 | 0.00 |
| AL2G02902 | 0.21 | 53.01 | 248.18 | 33.08 | 26.94 |
| AL1G00509 | 0.03 | 4.40 | 150.68 | 0.00 | 0.00 |
| AL1G05993 | 0.07 | 6.72 | 98.75 | 0.00 | 0.00 |
| AL7G04259 | 1.72 | 166.02 | 96.33 | 1.93 | 5.47 |
| AL4G00094 | 0.92 | 84.55 | 92.38 | 0.00 | 0.00 |
| AL5G00194 | 0.02 | 1.25 | 79.53 | 0.05 | 0.27 |
| AL7G01060 | 0.05 | 1.58 | 29.46 | 0.00 | 0.00 |
| AL1G03034 | 0.18 | 4.40 | 24.09 | 0.00 | 0.00 |
| AL1G06069 | 1.46 | 31.45 | 21.58 | 0.23 | 0.24 |
| AL1G06068 | 2.78 | 52.91 | 19.04 | 0.29 | 0.31 |
| AL1G06067 | 3.29 | 59.10 | 17.96 | 0.57 | 0.05 |
| AL5G00207 | 0.22 | 3.66 | 16.27 | 0.30 | 0.97 |
| AL1G05990 | 4.08 | 62.09 | 15.24 | 1.24 | 0.16 |
| AL7G05075 | 0.19 | 2.63 | 13.95 | 0.00 | 0.00 |
| AL5G02319 | 0.18 | 2.30 | 12.73 | 6.00 | 0.63 |
| AL1G05994 | 1.73 | 21.79 | 12.58 | 0.00 | 0.23 |
| AL6G03023 | 11.64 | 144.54 | 12.42 | 97.57 | 37.87 |
| AL7G04461 | 2.14 | 26.57 | 12.40 | 0.32 | 1.52 |
| AL4G00908 | 18.12 | 216.02 | 11.92 | 17.00 | 3.01 |
| AL5G00200 | 0.17 | 1.98 | 11.61 | 0.02 | 0.08 |
| AL5G02497 | 1.43 | 15.97 | 11.13 | 0.00 | 0.00 |
| AL1G05992 | 4.91 | 54.52 | 11.10 | 0.51 | 0.00 |
| AL4G00821 | 1.66 | 18.24 | 11.01 | 0.00 | 0.00 |
| AL1G05991 | 10.60 | 115.48 | 10.89 | 1.16 | 1.96 |
| AL3G01291 | 0.58 | 6.19 | 10.67 | 0.00 | 0.00 |
| AL7G00656 | 2.82 | 29.46 | 10.44 | 152.46 | 12.60 |

**b The transcript levels of *CYP450* genes over 10-fold higher in salt-treated leaves than in leaves.**

| Gene ID | Leaves | Salt-treated leaves | Fold number (Salt-treated leaves / leaves) | Roots | Salt-treated leaves |
|---|---|---|---|---|---|
| AL7G05074 | 0.06 | 25.41 | 408.62 | 1.84 | 11.80 |
| AL7G05062 | 0.26 | 42.95 | 163.41 | 0.00 | 0.00 |
| AL3G02922 | 0.65 | 42.49 | 65.48 | 0.60 | 0.11 |
| AL2G01343 | 0.01 | 0.83 | 56.59 | 5.22 | 0.76 |
| AL2G01337 | 0.01 | 0.61 | 48.75 | 5.33 | 0.69 |
| AL6G02127 | 0.02 | 0.78 | 42.46 | 50.53 | 68.41 |
| AL3G03732 | 0.16 | 5.25 | 33.47 | 0.00 | 1.18 |
| AL3G03970 | 0.29 | 9.37 | 32.72 | 12.43 | 5.38 |
| AL6G01819 | 0.76 | 13.38 | 17.60 | 1.35 | 0.09 |
| AL5G03860 | 0.27 | 3.93 | 14.82 | 11.05 | 22.11 |
| AL3G05558 | 0.52 | 7.41 | 14.25 | 3.19 | 1.07 |
| AL4G03387 | 0.15 | 1.92 | 12.68 | 0.19 | 0.52 |
| AL4G02776 | 0.06 | 0.68 | 12.24 | 2.20 | 1.44 |
| AL7G05064 | 0.21 | 2.29 | 11.14 | 0.00 | 0.00 |
| AL5G00242 | 0.09 | 0.90 | 10.32 | 0.72 | 1.58 |

**Table S22. Description of 46 biosynthetic gene clusters (BGCs) containing 117 *CYP450* genes in the expression profiling heat map.**

| Cluster number | Gene ID | Start position | End position | Gene description |
|---|---|---|---|---|
| Cluster01 | AL1G00968 | 42522450 | 42524161 | Cytochrome P450 |
| Cluster01 | AL1G00969 | 42534440 | 42535963 | Cytokinin-O-glucosyltransferase |
| Cluster01 | AL1G00970 | 42538434 | 42542769 | Mate efflux family |
| Cluster01 | AL1G00971 | 42571300 | 42572984 | Cytochrome P450 |
| Cluster01 | AL1G00972 | 42584427 | 42586053 | Cytochrome P450 |
| Cluster01 | AL1G00973 | 42629901 | 42630482 | Cytochrome P450 |
| Cluster01 | AL1G00974 | 42630624 | 42631589 | Cytochrome P450 |
| Cluster02 | AL1G01100 | 48321887 | 48323230 | Short-chain dehydrogenase/Reductase |
| Cluster02 | AL1G01101 | 48382053 | 48385127 | Wax synthase |
| Cluster02 | AL1G01102 | 48505941 | 48507485 | Cytochrome P450 |
| Cluster02 | AL1G01103 | 48523279 | 48524553 | Chalcone and stilbene synthases |
| Cluster02 | AL1G01104 | 48695715 | 48696452 | Transposon protein |
| Cluster02 | AL1G01105 | 48728301 | 48729677 | Gibberellin receptor |
| Cluster02 | AL1G01106 | 48759261 | 48760249 | MYB Family transcription factor |
| Cluster03 | AL1G02598 | 123211411 | 123212341 | Cysteine proteinase |
| Cluster03 | AL1G02599 | 123239317 | 123240104 | Cysteine proteinase |
| Cluster03 | AL1G02600 | 123242491 | 123244111 | Cytochrome P450 |
| Cluster03 | AL1G02601 | 123291972 | 123293610 | Auxin-induced protein |
| Cluster03 | AL1G02602 | 123382899 | 123384455 | Cytochrome P450 |
| Cluster03 | AL1G02603 | 123454915 | 123456090 | Cytochrome P450 |
| Cluster03 | AL1G02604 | 123491816 | 123493336 | Cytochrome P450 |
| Cluster03 | AL1G02605 | 123586162 | 123587697 | Cytochrome P450 |
| Cluster03 | AL1G02606 | 123588223 | 123590136 | 50S ribosomal protein |
| Cluster03 | AL1G02607 | 123662219 | 123663278 | Other |
| Cluster03 | AL1G02608 | 123663555 | 123664569 | Thylakoid lumenal 20 kda protein |
| Cluster03 | AL1G02609 | 123672533 | 123674029 | Cytokinin-N-glucosyltransferase |
| Cluster03 | AL1G02610 | 123705568 | 123707174 | Cytokinin-N-glucosyltransferase |
| Cluster03 | AL1G02611 | 123770645 | 123772964 | Indole-3-acetate beta-glucosyltransferase |
| Cluster03 | AL1G02612 | 123774944 | 123777342 | Cyclin |
| Cluster03 | AL1G02613 | 123780797 | 123787406 | Proline-rich family protein |
| Cluster05 | AL1G05701 | 556127208 | 556132172 | Other |
| Cluster05 | AL1G05702 | 556132240 | 556133163 | Other |
| Cluster05 | AL1G05703 | 556162137 | 556163165 | Ubiquitin |
| Cluster05 | AL1G05704 | 556199372 | 556200379 | Absentia protein family |
| Cluster05 | AL1G05705 | 556435215 | 556445600 | Cytochrome P450 |
| Cluster05 | AL1G05706 | 556469174 | 556475788 | Cytochrome P450 |
| Cluster05 | AL1G05707 | 556727156 | 556728202 | O-acyltransferase |
| Cluster05 | AL1G05708 | 557257863 | 557267929 | Cytochrome P450 |

| Cluster number | Gene ID | Start position | End position | Gene description |
|---|---|---|---|---|
| Cluster05 | AL1G05709 | 557540267 | 557550659 | Cytochrome P450 |
| Cluster05 | AL1G05710 | 557565247 | 557575034 | Cytochrome P450 |
| Cluster05 | AL1G05711 | 558167269 | 558177483 | Cytochrome P450 |
| Cluster05 | AL1G05712 | 559399882 | 559435199 | Cycloartenol synthase |
| Cluster05 | AL1G05713 | 559610756 | 559611550 | Other |
| Cluster06 | AL1G05842 | 567005319 | 567006518 | O-methyltransferase |
| Cluster06 | AL1G05843 | 567013972 | 567015306 | Anthocyanidin 5,3-O-glucosyltransferase |
| Cluster06 | AL1G05844 | 567036664 | 567038034 | UDP-glucoronosyl |
| Cluster06 | AL1G05845 | 567119718 | 567126593 | C2 domain containing protein |
| Cluster06 | AL1G05846 | 567127885 | 567132693 | Tetratricopeptide-like helical |
| Cluster06 | AL1G05847 | 567136048 | 567139145 | Nuclease Pa3 |
| Cluster06 | AL1G05848 | 567315715 | 567318453 | Other |
| Cluster06 | AL1G05849 | 567372588 | 567374277 | Cytochrome P450 |
| Cluster08 | AL1G05922 | 570980891 | 570995028 | Ent-Kaurene synthase |
| Cluster08 | AL1G05923 | 571102630 | 571110665 | Ent-Kaurene synthase |
| Cluster08 | AL1G05924 | 571368489 | 571370003 | Cytochrome P450 |
| Cluster08 | AL1G05925 | 571404126 | 571409747 | Other |
| Cluster08 | AL1G05926 | 571423749 | 571426276 | Other |
| Cluster08 | AL1G05927 | 571449385 | 571450557 | Oxidoreductase |
| Cluster08 | AL1G05928 | 571482035 | 571483735 | Transposon protein |
| Cluster08 | AL1G05929 | 571492766 | 571496315 | Fatty acid hydroxylase |
| Cluster08 | AL1G05930 | 571497584 | 571501524 | Zinc finger protein |
| Cluster09 | AL1G06096 | 578860058 | 578863127 | Glycosyl Transferase |
| Cluster09 | AL1G06097 | 578864727 | 578866871 | Tp53Rk-binding protein |
| Cluster09 | AL1G06098 | 578874033 | 578875304 | F-box domain containing protein |
| Cluster09 | AL1G06099 | 578905071 | 578914582 | Terpene synthase |
| Cluster09 | AL1G06100 | 578942112 | 578946254 | Peptidase M50 family protein |
| Cluster09 | AL1G06101 | 578946951 | 578951826 | Retrotransposon protein |
| Cluster09 | AL1G06102 | 578972022 | 578975710 | G-beta repeat domain |
| Cluster09 | AL1G06103 | 578976319 | 578980493 | DUF647 domain |
| Cluster09 | AL1G06104 | 579004179 | 579005678 | Cytochrome P450 |
| Cluster10 | AL1G06292 | 590322194 | 590324142 | Cytochrome P450 |
| Cluster10 | AL1G06293 | 590471170 | 590491613 | Ent-kaurene synthase |
| Cluster10 | AL1G06294 | 590496528 | 590498360 | Cytochrome P450 |
| Cluster10 | AL1G06295 | 590520612 | 590524544 | Cytochrome P450 |
| Cluster10 | AL1G06296 | 590626162 | 590631663 | Ent-kaurene synthase |
| Cluster10 | AL1G06297 | 590761025 | 590762095 | Dehydrodolichyl diphosphate synthase |
| Cluster10 | AL1G06298 | 590781270 | 590781575 | Dehydrodolichyl diphosphate synthase |
| Cluster10 | AL1G06299 | 590781683 | 590786330 | Dehydrodolichyl diphosphate synthase |
| Cluster10 | AL1G06300 | 590800800 | 590802167 | UDP-glucoronosyl and UDP-glucosyl transferase domain |

| Cluster number | Gene ID | Start position | End position | Gene description |
|---|---|---|---|---|
| Cluster10 | AL1G06301 | 590831004 | 590833018 | ABC-2 Type transporter domain |
| Cluster10 | AL1G06302 | 590863617 | 590864619 | Phosphatidylethanolamine-binding protein |
| Cluster10 | AL1G06303 | 590875206 | 590891171 | Protein kinase domain |
| Cluster11 | AL1G06307 | 591149538 | 591150002 | Phosphatase family protein |
| Cluster11 | AL1G06308 | 591157430 | 591159236 | Cytochrome P450 |
| Cluster11 | AL1G06309 | 591193794 | 591194225 | Cytochrome P450 |
| Cluster11 | AL1G06310 | 591199949 | 591208248 | Cytochrome P450 |
| Cluster16 | AL2G00672 | 32132056 | 32132964 | F-Box domain |
| Cluster16 | AL2G00673 | 32145902 | 32156418 | Cytochrome P450 |
| Cluster16 | AL2G00674 | 32156460 | 32156999 | Cytochrome P450 |
| Cluster16 | AL2G00675 | 32247248 | 32247844 | Gibberellin 3-beta-dioxygenase |
| Cluster16 | AL2G00676 | 32248051 | 32248422 | Gibberellin 3-beta-dioxygenase |
| Cluster17 | AL2G00827 | 40006436 | 40014451 | Helicase |
| Cluster17 | AL2G00828 | 40020980 | 40024677 | Other |
| Cluster17 | AL2G00829 | 40028288 | 40030518 | Cytochrome P450 |
| Cluster17 | AL2G00830 | 40057039 | 40061721 | Glycosyl transferase |
| Cluster17 | AL2G00831 | 40086692 | 40094026 | Carboxylate xxidase homolog 1 |
| Cluster17 | AL2G00832 | 40209332 | 40210758 | Proline-rich protein |
| Cluster17 | AL2G00833 | 40233295 | 40235511 | CESA8 |
| Cluster17 | AL2G00834 | 40235553 | 40238348 | CESA2 |
| Cluster17 | AL2G00835 | 40268987 | 40269741 | Protease inhibitor/Seed Storage |
| Cluster17 | AL2G00836 | 40292412 | 40293299 | Fruit protein |
| Cluster17 | AL2G00837 | 40301453 | 40304261 | Zinc finger protein |
| Cluster20 | AL2G01337 | 70655675 | 70657303 | Cytochrome P450 |
| Cluster20 | AL2G01338 | 70875160 | 70877044 | Terpene synthase |
| Cluster20 | AL2G01339 | 70887321 | 70887834 | Tigr01627 family protein |
| Cluster20 | AL2G01340 | 70891669 | 70895004 | Zinc finger protein |
| Cluster20 | AL2G01341 | 70895232 | 70895882 | Retrotransposon protein |
| Cluster20 | AL2G01342 | 70900022 | 70900894 | Tigr01627 family protein |
| Cluster20 | AL2G01343 | 71055412 | 71056806 | Cytochrome P450 |
| Cluster20 | AL2G01344 | 71284303 | 71286309 | Terpene synthase |
| Cluster20 | AL2G01345 | 71289947 | 71292827 | SAM dependent carboxyl methyltransferase |
| Cluster20 | AL2G01346 | 71293791 | 71294657 | Tigr01627 family protein |
| Cluster20 | AL2G01347 | 71358129 | 71359287 | SAM dependent carboxyl methyltransferase |
| Cluster20 | AL2G01348 | 71433452 | 71435377 | Glutelin |
| Cluster22 | AL2G01455 | 76099854 | 76101323 | UDP-glucosyl transferase |
| Cluster22 | AL2G01456 | 76149766 | 76151313 | Cytokinin-O-glucosyltransferase |
| Cluster22 | AL2G01457 | 76188680 | 76190176 | Cytochrome P450 |
| Cluster22 | AL2G01458 | 76386632 | 76389179 | Cytochrome P450 |

| Cluster number | Gene ID | Start position | End position | Gene description |
|---|---|---|---|---|
| Cluster25 | AL2G03967 | 476203060 | 476205454 | NADH dehydrogenase |
| Cluster25 | AL2G03968 | 476206378 | 476209811 | Transporter family protein |
| Cluster25 | AL2G03969 | 476211346 | 476213605 | Cytochrome P450 |
| Cluster25 | AL2G03970 | 476264534 | 476265775 | Fad dependent oxidoreductase |
| Cluster25 | AL2G03971 | 476290692 | 476295659 | Dehydration protein |
| Cluster25 | AL2G03972 | 476338372 | 476346659 | AMP-binding enzyme |
| Cluster25 | AL2G03973 | 476348985 | 476351033 | Cytidylyltransferase domain |
| Cluster25 | AL2G03974 | 476351749 | 476354027 | Lrgb-like family protein |
| Cluster25 | AL2G03975 | 476443872 | 476452762 | Lrgb-like family protein |
| Cluster25 | AL2G03976 | 476481954 | 476482456 | Plastocyanin-like domain |
| Cluster25 | AL2G03977 | 476488799 | 476489947 | Other |
| Cluster25 | AL2G03978 | 476539185 | 476541947 | Proteasome subunit |
| Cluster25 | AL2G03979 | 476543359 | 476545236 | PPR Repeat Domain |
| Cluster25 | AL2G03980 | 476546530 | 476550760 | CslD1 |
| Cluster26 | AL2G04955 | 544186163 | 544188904 | O-Methyltransferase |
| Cluster26 | AL2G04956 | 544249328 | 544253314 | O-Methyltransferase |
| Cluster26 | AL2G04957 | 544266464 | 544267786 | 60S ribosomal protein |
| Cluster26 | AL2G04958 | 544268333 | 544270304 | Chch domain containing protein |
| Cluster26 | AL2G04959 | 544275385 | 544275813 | Mitosis pProtein dim1 |
| Cluster26 | AL2G04960 | 544276330 | 544281989 | Strubbelig-receptor family 6 precursor |
| Cluster26 | AL2G04961 | 544380283 | 544384549 | O-Methyltransferase |
| Cluster26 | AL2G04962 | 544385069 | 544389637 | Auxin response factor 14 |
| Cluster29 | AL3G00227 | 11074923 | 11078723 | Other |
| Cluster29 | AL3G00228 | 11096432 | 11100238 | Male sterility protein |
| Cluster29 | AL3G00229 | 11150909 | 11152428 | Hydrolase |
| Cluster29 | AL3G00230 | 11184488 | 11186163 | Transferase family protein |
| Cluster29 | AL3G00231 | 11189913 | 11190645 | Transferase family protein |
| Cluster29 | AL3G00232 | 11208485 | 11210254 | Hydrolase |
| Cluster29 | AL3G00233 | 11263875 | 11265871 | Retrotransposon protein |
| Cluster29 | AL3G00234 | 11299954 | 11301588 | Cytochrome P450 |
| Cluster29 | AL3G00235 | 11389267 | 11389644 | Transferase family protein |
| Cluster29 | AL3G00236 | 11420454 | 11420810 | Other |
| Cluster29 | AL3G00237 | 11441724 | 11442892 | Other |
| Cluster29 | AL3G00238 | 11444399 | 11448983 | Other |
| Cluster29 | AL3G00239 | 11491906 | 11493693 | Other |
| Cluster29 | AL3G00240 | 11499490 | 11504434 | Pattern formation protein |
| Cluster29 | AL3G00241 | 11533264 | 11538178 | NADPH reductase |
| Cluster29 | AL3G00242 | 11543239 | 11543727 | Zinc finger protein |
| Cluster29 | AL3G00243 | 11552683 | 11553180 | Zinc finger protein |
| Cluster31 | AL3G02460 | 169468616 | 169471000 | Broad complex BTB domain |
| Cluster31 | AL3G02461 | 169638958 | 169643115 | Strictosidine synthase |
| Cluster31 | AL3G02462 | 169845303 | 169850687 | 2-Isopropylmalate synthase |

| Cluster number | Gene ID | Start position | End position | Gene description |
|---|---|---|---|---|
| Cluster31 | AL3G02463 | 170014852 | 170017241 | Zinc finger protein |
| Cluster31 | AL3G02464 | 170287939 | 170293881 | Other |
| Cluster31 | AL3G02465 | 170554976 | 170557641 | Cytochrome P450 |
| Cluster37 | AL3G05629 | 554830749 | 554832409 | Cytochrome P450 |
| Cluster37 | AL3G05630 | 554833561 | 554839216 | ATP phosphoribosyltransferase |
| Cluster37 | AL3G05631 | 554851724 | 554856649 | Serine acetyltransferase |
| Cluster37 | AL3G05632 | 554885153 | 554885733 | Aldehyde oxidase |
| Cluster37 | AL3G05633 | 554886465 | 554890124 | Aldehyde oxidase |
| Cluster37 | AL3G05634 | 554893368 | 554896390 | Leucoanthocyanidin reductase |
| Cluster37 | AL3G05635 | 555023764 | 555025922 | Mate efflux family protein |
| Cluster37 | AL3G05636 | 555057628 | 555059115 | Cytochrome P450 |
| Cluster37 | AL3G05637 | 555185508 | 555189948 | Serine palmitoyltransferase |
| Cluster37 | AL3G05638 | 555322818 | 555324467 | AMP-binding enzyme |
| Cluster37 | AL3G05639 | 555326207 | 555328617 | AMP-binding enzyme |
| Cluster37 | AL3G05640 | 555395500 | 555396333 | Hydrolase |
| Cluster37 | AL3G05641 | 555409479 | 555412939 | Anchored protein precursor |
| Cluster37 | AL3G05642 | 555415010 | 555419696 | Other |
| Cluster37 | AL3G05643 | 555455321 | 555455698 | Other |
| Cluster37 | AL3G05644 | 555458614 | 555459870 | Glutathione S-transferase |
| Cluster37 | AL3G05645 | 555464258 | 555465339 | Glutathione S-transferase |
| Cluster37 | AL3G05646 | 555469489 | 555470514 | Transferase family protein |
| Cluster37 | AL3G05647 | 555473252 | 555474370 | Calmodulin-binding motif family protein |
| Cluster37 | AL3G05648 | 555549443 | 555551100 | Cytochrome P450 |
| Cluster37 | AL3G05649 | 555552256 | 555557480 | ATP phosphoribosyltransferase |
| Cluster37 | AL3G05650 | 555602631 | 555607567 | Serine acetyltransferase protein |
| Cluster37 | AL3G05651 | 555614216 | 555619248 | Aldehyde oxidase |
| Cluster37 | AL3G05652 | 555622777 | 555625418 | Leucoanthocyanidin reductase |
| Cluster37 | AL3G05653 | 555793716 | 555798252 | Serine palmitoyltransferase |
| Cluster37 | AL3G05654 | 555805973 | 555806455 | AMP-binding enzyme |
| Cluster37 | AL3G05655 | 555913812 | 555915238 | Cytochrome P450 |
| Cluster37 | AL3G05656 | 555928888 | 555930814 | Mate efflux family protein |
| Cluster37 | AL3G05657 | 556018705 | 556019088 | Other |
| Cluster37 | AL3G05658 | 556081444 | 556083093 | AMP-binding enzyme |
| Cluster37 | AL3G05659 | 556086784 | 556089246 | AMP-binding enzyme |
| Cluster39 | AL3G06283 | 581313593 | 581314710 | Peroxidase precursor |
| Cluster39 | AL3G06284 | 581325281 | 581326570 | Peroxidase precursor |
| Cluster39 | AL3G06285 | 581336552 | 581338256 | Cytochrome P450 |
| Cluster39 | AL3G06286 | 581376949 | 581378593 | Cytochrome P450 |
| Cluster39 | AL3G06287 | 581380124 | 581383701 | Pheophorbide a oxygenase |
| Cluster39 | AL3G06288 | 581388621 | 581389259 | Cytochrome P450 |
| Cluster39 | AL3G06289 | 581412681 | 581413193 | Retrotransposon protein |

| Cluster number | Gene ID | Start position | End position | Gene description |
|---|---|---|---|---|
| Cluster39 | AL3G06290 | 581457177 | 581458833 | Cytochrome P450 |
| Cluster39 | AL3G06291 | 581476823 | 581477311 | Cytochrome P450 |
| Cluster39 | AL3G06292 | 581516455 | 581519686 | Terpene synthase |
| Cluster39 | AL3G06293 | 581531901 | 581533607 | Cytochrome P450 |
| Cluster39 | AL3G06294 | 581549854 | 581600343 | Cysteine-rich receptor-like protein |
| Cluster40 | AL3G06343 | 583759237 | 583760709 | Cytochrome P450 |
| Cluster40 | AL3G06344 | 583817908 | 583819816 | Cytochrome P450 |
| Cluster40 | AL3G06345 | 583822926 | 583823264 | Other |
| Cluster40 | AL3G06346 | 583845519 | 583850540 | ENT-kaurene synthase |
| Cluster40 | AL3G06347 | 583854054 | 583855316 | Peptidase |
| Cluster41 | AL4G00006 | 611046 | 613295 | Receptor-like protein kKinase |
| Cluster41 | AL4G00007 | 628315 | 631290 | Receptor-like protein kKinase |
| Cluster41 | AL4G00008 | 632487 | 642725 | Glucosyltransferase |
| Cluster41 | AL4G00009 | 686183 | 697315 | Glucosyltransferase |
| Cluster41 | AL4G00010 | 711306 | 713436 | Other |
| Cluster41 | AL4G00011 | 730350 | 731792 | Cytochrome P450 |
| Cluster41 | AL4G00012 | 786263 | 792039 | Other |
| Cluster41 | AL4G00013 | 829941 | 831449 | Transferase family protein |
| Cluster41 | AL4G00014 | 834236 | 837689 | Other |
| Cluster41 | AL4G00015 | 851962 | 853269 | Transferase family protein |
| Cluster41 | AL4G00016 | 896033 | 899812 | Other |
| Cluster42 | AL4G00017 | 1139560 | 1143256 | Receptor-like protein kinase |
| Cluster42 | AL4G00018 | 1157228 | 1158205 | Oxidoreductase |
| Cluster42 | AL4G00019 | 1179520 | 1181586 | Cytochrome P450 |
| Cluster42 | AL4G00020 | 1186040 | 1193797 | Pfkb family |
| Cluster42 | AL4G00021 | 1202168 | 1208002 | Cytochrome P450 |
| Cluster42 | AL4G00022 | 1214795 | 1219934 | Protein kinase domain |
| Cluster42 | AL4G00023 | 1243186 | 1244169 | Transferase family protein |
| Cluster42 | AL4G00024 | 1246209 | 1248486 | CBS domain |
| Cluster42 | AL4G00025 | 1271036 | 1283253 | Protein kinase domain |
| Cluster42 | AL4G00026 | 1309405 | 1310680 | Transferase family protein |
| Cluster43 | AL4G00041 | 1855642 | 1856424 | Dirigent |
| Cluster43 | AL4G00042 | 1927038 | 1928785 | Cytochrome P450 |
| Cluster43 | AL4G00043 | 1936800 | 1938851 | Other |
| Cluster43 | AL4G00044 | 1982029 | 1996402 | SAM dependent carboxyl methyltransferase |
| Cluster43 | AL4G00045 | 1997735 | 1999801 | Cytochrome P450 |
| Cluster44 | AL4G00048 | 2304285 | 2307484 | Cytochrome P450 |
| Cluster44 | AL4G00049 | 2329723 | 2332896 | Terpene synthase |
| Cluster44 | AL4G00050 | 2490478 | 2492526 | ATO binding protein |
| Cluster44 | AL4G00051 | 2523515 | 2536685 | Cycloartenol synthase |
| Cluster44 | AL4G00052 | 2595441 | 2597484 | Cytochrome P450 |

| Cluster number | Gene ID | Start position | End position | Gene description |
|---|---|---|---|---|
| Cluster44 | AL4G00053 | 2601532 | 2602752 | Other |
| Cluster44 | AL4G00054 | 2604525 | 2608607 | Other |
| Cluster46 | AL4G00080 | 3670394 | 3674385 | Terpene synthase |
| Cluster46 | AL4G00081 | 3715055 | 3718622 | Cytochrome P450 |
| Cluster46 | AL4G00082 | 3719097 | 3719468 | Pollen Allergen |
| Cluster46 | AL4G00083 | 3746443 | 3746862 | Cytochrome P450 |
| Cluster46 | AL4G00084 | 3746966 | 3748344 | Cytochrome P450 |
| Cluster46 | AL4G00085 | 3774913 | 3778195 | Terpene synthase |
| Cluster46 | AL4G00086 | 3779800 | 3781723 | Cytochrome P450 |
| Cluster46 | AL4G00087 | 3832937 | 3836188 | Terpene synthase |
| Cluster46 | AL4G00088 | 3837914 | 3840035 | Cytochrome P450 |
| Cluster46 | AL4G00089 | 3927223 | 3928875 | Cytochrome P450 |
| Cluster46 | AL4G00090 | 3930528 | 3933814 | Terpene synthase |
| Cluster46 | AL4G00091 | 3994994 | 3996568 | Decarboxylase |
| Cluster46 | AL4G00092 | 4066835 | 4067683 | Wax synthase |
| Cluster46 | AL4G00093 | 4074856 | 4079105 | Wax synthase |
| Cluster46 | AL4G00094 | 4125631 | 4128378 | Cytochrome P450 |
| Cluster46 | AL4G00095 | 4261389 | 4264895 | Terpene synthase |
| Cluster46 | AL4G00096 | 4394691 | 4398925 | Cytochrome P450 |
| Cluster46 | AL4G00097 | 4435859 | 4438431 | Hexokinase |
| Cluster46 | AL4G00098 | 4448123 | 4451393 | Other |
| Cluster48 | AL4G00820 | 32495835 | 32498998 | Terpene synthase |
| Cluster48 | AL4G00821 | 32507525 | 32509134 | Cytochrome P450 |
| Cluster48 | AL4G00822 | 32534010 | 32548956 | Ent-Kaurene synthase |
| Cluster48 | AL4G00823 | 32551858 | 32553126 | Retrotransposon protein |
| Cluster48 | AL4G00824 | 32554543 | 32557670 | Shwachman-Bodian-Diamond syndrome protein |
| Cluster49 | AL4G00855 | 33419001 | 33422165 | Shwachman-Bodian-Diamond syndrome protein |
| Cluster49 | AL4G00856 | 33423584 | 33424880 | Retrotransposon protein |
| Cluster49 | AL4G00857 | 33429365 | 33443678 | Ent-Kaurene synthase |
| Cluster49 | AL4G00858 | 33540407 | 33541944 | Cytochrome P450 |
| Cluster49 | AL4G00859 | 33543335 | 33546526 | Terpene synthase |
| Cluster50 | AL4G00905 | 35341513 | 35342823 | Other |
| Cluster50 | AL4G00906 | 35343534 | 35345471 | Other |
| Cluster50 | AL4G00907 | 35346492 | 35351076 | Cytochrome P450 |
| Cluster50 | AL4G00908 | 35354760 | 35357339 | Cytochrome P450 |
| Cluster50 | AL4G00909 | 35380012 | 35389559 | Cytochrome P450 |
| Cluster50 | AL4G00910 | 35407261 | 35410486 | Cytochrome P450 |
| Cluster50 | AL4G00911 | 35415371 | 35422601 | Ent-Kaurene synthase |
| Cluster50 | AL4G00912 | 35470867 | 35472546 | Transferase family domain |
| Cluster50 | AL4G00913 | 35483018 | 35542255 | Transferase family domain |

| Cluster number | Gene ID | Start position | End position | Gene description |
|---|---|---|---|---|
| Cluster50 | AL4G00914 | 35563026 | 35566140 | Protein phosphatase |
| Cluster51 | AL4G00927 | 36189120 | 36191057 | Other |
| Cluster51 | AL4G00928 | 36191450 | 36211650 | Cytochrome P450 |
| Cluster51 | AL4G00929 | 36229253 | 36231855 | Cytochrome P450 |
| Cluster51 | AL4G00930 | 36301014 | 36308242 | Cytochrome P450 |
| Cluster51 | AL4G00931 | 36333148 | 36336075 | Cytochrome P450 |
| Cluster51 | AL4G00932 | 36340915 | 36348259 | Ent-Kaurene synthase |
| Cluster51 | AL4G00933 | 36352063 | 36353748 | Transferase family domain |
| Cluster51 | AL4G00934 | 36438001 | 36439795 | Transferase family domain |
| Cluster51 | AL4G00935 | 36504576 | 36506434 | Transferase family domain |
| Cluster51 | AL4G00936 | 36509590 | 36512590 | Protein phosphatase |
| Cluster51 | AL4G00937 | 36537668 | 36540098 | Transporter family protein |
| Cluster64 | AL4G05218 | 541136376 | 541136885 | Lipoxygenase protein |
| Cluster64 | AL4G05219 | 541398217 | 541402001 | Lipoxygenase protein |
| Cluster64 | AL4G05220 | 541590755 | 541596314 | Caleosin related protein |
| Cluster64 | AL4G05221 | 541603641 | 541605756 | Glutamate decarboxylase |
| Cluster64 | AL4G05222 | 541729345 | 541733739 | O-Methyltransferase |
| Cluster64 | AL4G05223 | 541809358 | 541811386 | Chalcone synthase |
| Cluster64 | AL4G05224 | 542154382 | 542155902 | Cytochrome P450 |
| Cluster64 | AL4G05225 | 542246291 | 542250046 | Oxidoreductase |
| Cluster64 | AL4G05226 | 542250758 | 542251998 | Oxidoreductase |
| Cluster64 | AL4G05227 | 542281988 | 542283709 | Oxidoreductase |
| Cluster64 | AL4G05228 | 542284553 | 542286506 | Glutamate decarboxylase |
| Cluster72 | AL5G00192 | 8421548 | 8423554 | Ubiquitin family protein |
| Cluster72 | AL5G00193 | 8459596 | 8461982 | Cytochrome P450 |
| Cluster72 | AL5G00194 | 8480731 | 8482339 | Cytochrome P450 |
| Cluster72 | AL5G00195 | 8487280 | 8488224 | Cytochrome P450 |
| Cluster72 | AL5G00196 | 8500901 | 8503770 | Cytochrome P450 |
| Cluster72 | AL5G00197 | 8505840 | 8506585 | Cytochrome P450 |
| Cluster72 | AL5G00198 | 8508657 | 8511641 | Cytochrome P450 |
| Cluster72 | AL5G00199 | 8512301 | 8513543 | Oxidoreductase |
| Cluster72 | AL5G00200 | 8895832 | 8897446 | Cytochrome P450 |
| Cluster72 | AL5G00201 | 8904939 | 8931216 | Cytochrome P450 |
| Cluster72 | AL5G00202 | 8970356 | 8971436 | Oxidoreductase |
| Cluster72 | AL5G00203 | 8977047 | 8977892 | Cytochrome P450 |
| Cluster72 | AL5G00204 | 9010331 | 9011374 | Cytochrome P450 |
| Cluster72 | AL5G00205 | 9018734 | 9020483 | Other |
| Cluster72 | AL5G00206 | 9022072 | 9023445 | DNDj heat shock N-terminal domain |
| Cluster72 | AL5G00207 | 9023933 | 9026304 | Cytochrome P450 |
| Cluster72 | AL5G00208 | 9028162 | 9029646 | Anthocyanin 5-O-glucosyltransferase |
| Cluster72 | AL5G00209 | 9340536 | 9343326 | Terpene synthase |
| Cluster72 | AL5G00210 | 9344152 | 9345717 | Cytochrome P450 |

| Cluster number | Gene ID | Start position | End position | Gene description |
|---|---|---|---|---|
| Cluster72 | AL5G00211 | 9349775 | 9351664 | Terpene synthase |
| Cluster72 | AL5G00212 | 9398513 | 9401744 | Terpene synthase |
| Cluster72 | AL5G00213 | 9402838 | 9412276 | AMP-binding enzyme family |
| Cluster72 | AL5G00214 | 9412951 | 9414351 | 6-Phosphogluconate dehydrogenase |
| Cluster72 | AL5G00215 | 9497354 | 9501523 | G-Protein alpha subunit |
| Cluster72 | AL5G00216 | 9502486 | 9506054 | PPR repeat domain |
| Cluster72 | AL5G00217 | 9519883 | 9523105 | Potassium transporter |
| Cluster72 | AL5G00218 | 9525761 | 9526324 | Pathogenesis-related Bet VI family |
| Cluster72 | AL5G00219 | 9526959 | 9531547 | Osfbx182-F-Box domain |
| Cluster72 | AL5G00220 | 9535842 | 9537844 | Other |
| Cluster72 | AL5G00221 | 9569604 | 9583130 | Resistance protein |
| Cluster74 | AL5G00237 | 10423065 | 10426753 | Terpene synthase |
| Cluster74 | AL5G00238 | 10436918 | 10437733 | Other |
| Cluster74 | AL5G00239 | 10440795 | 10442050 | Regulatory protein |
| Cluster74 | AL5G00240 | 10446001 | 10454819 | Pectinacetylesterase domain |
| Cluster74 | AL5G00241 | 10460981 | 10462627 | Cytochrome P450 |
| Cluster74 | AL5G00242 | 10469439 | 10471699 | Cytochrome P450 |
| Cluster74 | AL5G00243 | 10515322 | 10518489 | Terpene synthase |
| Cluster74 | AL5G00244 | 10533299 | 10533637 | Serine/Threonine-protein kinase |
| Cluster74 | AL5G00245 | 10556059 | 10560816 | Terpene synthase |
| Cluster74 | AL5G00246 | 10560910 | 10562873 | Terpene synthase |
| Cluster74 | AL5G00247 | 10599438 | 10606190 | Terpene synthase |
| Cluster74 | AL5G00248 | 10609283 | 10615224 | Pinhead |
| Cluster76 | AL5G00412 | 16991818 | 16994625 | Starch synthase |
| Cluster76 | AL5G00413 | 17031619 | 17033258 | Cytochrome P450 |
| Cluster76 | AL5G00414 | 17035477 | 17038165 | Other |
| Cluster76 | AL5G00415 | 17098066 | 17100018 | Putative DNA polymerase delta complex subunit |
| Cluster76 | AL5G00416 | 17122332 | 17125801 | Glycosyl hydrolase family |
| Cluster76 | AL5G00417 | 17202527 | 17205997 | Glycosyl hydrolase family |
| Cluster76 | AL5G00418 | 17228180 | 17229139 | 1-Aminocyclopropane-1-Carboxylate Oxidase Homolog 1 |
| Cluster78 | AL5G00481 | 20601639 | 20601998 | Ossaur24-Auxin-responsive Saur gene family |
| Cluster78 | AL5G00482 | 20606904 | 20612681 | Brix domain containing protein |
| Cluster78 | AL5G00483 | 20614542 | 20615874 | Ribosome-binding factor A |
| Cluster78 | AL5G00484 | 20620494 | 20621606 | Gdp-Mannose 4,6 Dehydratase |
| Cluster78 | AL5G00485 | 20691351 | 20692475 | Gdp-Mannose 4,6 Dehydratase |
| Cluster78 | AL5G00486 | 20699832 | 20705685 | Transferase family protein |
| Cluster78 | AL5G00487 | 20722122 | 20723432 | Osfbx254-F-Box domain |
| Cluster78 | AL5G00488 | 20726901 | 20736526 | Methyltransferase |
| Cluster78 | AL5G00489 | 20800227 | 20802984 | Osfbx361-F-Box domain |

| Cluster number | Gene ID | Start position | End position | Gene description |
|---|---|---|---|---|
| Cluster78 | AL5G00490 | 20810846 | 20814384 | Duf630/Duf632 Domain |
| Cluster78 | AL5G00491 | 20842240 | 20843202 | Cytochrome P450 |
| Cluster78 | AL5G00492 | 20866969 | 20870072 | Bromodomain domain |
| Cluster79 | AL5G00635 | 26267628 | 26272806 | RNA recognition motif |
| Cluster79 | AL5G00636 | 26273756 | 26275180 | Transferase family domain |
| Cluster79 | AL5G00637 | 26280138 | 26282742 | Gdsl-Like lipase |
| Cluster79 | AL5G00638 | 26283827 | 26284141 | Transcription elongation factor 1 |
| Cluster79 | AL5G00639 | 26352859 | 26355274 | Cytochrome P450 |
| Cluster79 | AL5G00640 | 26373586 | 26374860 | G-Patch domain containing protein |
| Cluster79 | AL5G00641 | 26394633 | 26398026 | Xylosyltransferase |
| Cluster79 | AL5G00642 | 26413324 | 26419641 | Starch synthase |
| Cluster79 | AL5G00643 | 26420924 | 26422506 | Other |
| Cluster83 | AL5G00839 | 34898595 | 34908149 | Cycloartenol synthase |
| Cluster83 | AL5G00840 | 35005090 | 35006614 | Cytochrome P450 |
| Cluster83 | AL5G00841 | 35096873 | 35101978 | Cytochrome P450 |
| Cluster83 | AL5G00842 | 35221162 | 35221778 | AP2 domain containing protein |
| Cluster83 | AL5G00843 | 35310065 | 35312683 | Cytochrome P450 |
| Cluster83 | AL5G00844 | 35428024 | 35429328 | Cytochrome P450 |
| Cluster83 | AL5G00845 | 35605044 | 35605262 | Transcription initiation factor |
| Cluster83 | AL5G00846 | 35637326 | 35638861 | Cytochrome P450 |
| Cluster83 | AL5G00847 | 35663853 | 35664890 | Other |
| Cluster85 | AL5G00909 | 39254269 | 39256992 | Flavonol synthase |
| Cluster85 | AL5G00910 | 39277798 | 39278850 | Serine threonine kinase |
| Cluster85 | AL5G00911 | 39278880 | 39279584 | Serine threonine kinase |
| Cluster85 | AL5G00912 | 39293045 | 39295571 | Serine threonine kinase |
| Cluster85 | AL5G00913 | 39318641 | 39320342 | Cytochrome P450 |
| Cluster85 | AL5G00914 | 39353726 | 39354184 | Retrotransposon protein |
| Cluster85 | AL5G00915 | 39404488 | 39406162 | Cytochrome P450 |
| Cluster85 | AL5G00916 | 39485008 | 39486697 | Cytochrome P450 |
| Cluster85 | AL5G00917 | 39518791 | 39520538 | Cytochrome P450 |
| Cluster85 | AL5G00918 | 39521587 | 39522606 | Osmkk5 |
| Cluster85 | AL5G00919 | 39526050 | 39528756 | Zinc ion binding protein |
| Cluster85 | AL5G00920 | 39554965 | 39559457 | Glutamate receptor |
| Cluster85 | AL5G00921 | 39571691 | 39573001 | Anthocyanidin 3-O-glucosyltransferase |
| Cluster85 | AL5G00922 | 39611489 | 39613117 | Cytochrome P450 |
| Cluster85 | AL5G00923 | 39624449 | 39624991 | 60S ribosomal protein |
| Cluster85 | AL5G00924 | 39626493 | 39631985 | DNA binding protein |
| Cluster85 | AL5G00925 | 39632749 | 39633814 | Hypro1 |
| Cluster86 | AL5G01102 | 49731331 | 49732361 | Chalcone synthase |
| Cluster86 | AL5G01103 | 49798206 | 49799914 | Cytochrome P450 |
| Cluster86 | AL5G01104 | 49940498 | 49940902 | Chalcone synthase |
| Cluster86 | AL5G01105 | 49940929 | 49941806 | Chalcone synthase |

| Cluster number | Gene ID | Start position | End position | Gene description |
|---|---|---|---|---|
| Cluster88 | AL5G01347 | 62228905 | 62230082 | Odorant1 |
| Cluster88 | AL5G01348 | 62333613 | 62337119 | Delta14-sterol reductase |
| Cluster88 | AL5G01349 | 62426773 | 62442441 | NAD binding domain |
| Cluster88 | AL5G01350 | 62443588 | 62450032 | Phosphatidylinositol-4-phosphate 5-kinase family protein |
| Cluster88 | AL5G01351 | 62607974 | 62611021 | Other |
| Cluster88 | AL5G01352 | 62734205 | 62735677 | 3-Ketoacyl-Coa synthase |
| Cluster88 | AL5G01353 | 62887117 | 62888583 | 3-Ketoacyl-Coa synthase |
| Cluster88 | AL5G01354 | 62966534 | 62967982 | 3-Ketoacyl-Coa synthase |
| Cluster88 | AL5G01355 | 62988521 | 62989804 | 3-Ketoacyl-Coa synthase |
| Cluster88 | AL5G01356 | 63072693 | 63074141 | 3-Ketoacyl-Coa synthase |
| Cluster88 | AL5G01357 | 63204659 | 63205498 | 3-Ketoacyl-Coa synthase |
| Cluster88 | AL5G01358 | 63228508 | 63229464 | 3-Ketoacyl-Coa synthase |
| Cluster88 | AL5G01359 | 63391485 | 63391718 | Cytochrome P450 |
| Cluster88 | AL5G01360 | 63570208 | 63572352 | Retrotransposon protein |
| Cluster88 | AL5G01361 | 63598833 | 63599363 | 3-Ketoacyl-Coa synthase |
| Cluster88 | AL5G01362 | 63599390 | 63600199 | 3-Ketoacyl-Coa synthase |
| Cluster88 | AL5G01363 | 63613798 | 63615162 | 3-Ketoacyl-Coa synthase |
| Cluster89 | AL5G02309 | 153794488 | 153795513 | Oxidoreductase |
| Cluster89 | AL5G02310 | 153961016 | 153967298 | Retrotransposon protein |
| Cluster89 | AL5G02311 | 154026031 | 154027435 | Other |
| Cluster89 | AL5G02312 | 154683768 | 154684754 | Gibberellin receptor Gid1L2 |
| Cluster89 | AL5G02313 | 154685405 | 154686367 | Gibberellin receptor Gid1L2 |
| Cluster89 | AL5G02314 | 154808980 | 154810035 | Gibberellin receptor Gid1L2 |
| Cluster89 | AL5G02315 | 155143169 | 155144263 | Gibberellin receptor Gid1L2 |
| Cluster89 | AL5G02316 | 155715060 | 155726124 | Protein transport protein |
| Cluster89 | AL5G02317 | 155977462 | 155978259 | AP2 Domain |
| Cluster89 | AL5G02318 | 155986033 | 155987599 | Alpha-Amylase precursor |
| Cluster89 | AL5G02319 | 156615714 | 156619457 | Cytochrome P450 |
| Cluster89 | AL5G02320 | 156694613 | 156705798 | Glycoprotein 3-Alpha-L-fucosyltransferase |
| Cluster94 | AL6G00557 | 25144547 | 25149254 | Nodulin |
| Cluster94 | AL6G00558 | 25211688 | 25212964 | Dj-1 family protein |
| Cluster94 | AL6G00559 | 25216197 | 25218013 | Cytochrome P450 |
| Cluster94 | AL6G00560 | 25415865 | 25417673 | Cytochrome P450 |
| Cluster94 | AL6G00561 | 25635119 | 25636733 | Cytokinin-O-glucosyltransferase |
| Cluster94 | AL6G00562 | 25668980 | 25671408 | Erythronate-4-Phosphate dehydrogenase |
| Cluster94 | AL6G00563 | 25672290 | 25674763 | Skp1-like protein |
| Cluster95 | AL6G02059 | 148575339 | 148578763 | FAD binding domain |
| Cluster95 | AL6G02060 | 148615117 | 148617211 | Agenet domain |
| Cluster95 | AL6G02061 | 148682501 | 148684916 | Cytochrome P450 |

| Cluster number | Gene ID | Start position | End position | Gene description |
|---|---|---|---|---|
| Cluster95 | AL6G02062 | 149002792 | 149004114 | Other |
| Cluster95 | AL6G02063 | 149019642 | 149023230 | 26S protease regulatory subunit |
| Cluster95 | AL6G02064 | 149208939 | 149210513 | 3-Ketoacyl-Coa synthase |
| Cluster95 | AL6G02065 | 149261503 | 149268621 | Putative lysosomal pro-X carboxypeptidase |
| Cluster95 | AL6G02066 | 149270730 | 149272373 | UDP-glucoronosyl and UDP-glucosyl transferase |
| Cluster95 | AL6G02067 | 149347005 | 149348564 | UDP-glucoronosyl and UDP-glucosyl transferase |
| Cluster95 | AL6G02068 | 149420250 | 149421770 | UDP-glucoronosyl and UDP-glucosyl transferase |
| Cluster97 | AL6G04699 | 479358999 | 479360969 | Oxoacyl-reductase |
| Cluster97 | AL6G04700 | 479370344 | 479372366 | Pheophorbide A oxygenase |
| Cluster97 | AL6G04701 | 479374034 | 479376723 | Tumour differentially expressed protein, Putative, Expresse |
| Cluster97 | AL6G04702 | 479382590 | 479387785 | Other |
| Cluster97 | AL6G04703 | 479389580 | 479393029 | Zinc finger protein |
| Cluster97 | AL6G04704 | 479412173 | 479416958 | Cytochrome P450 |
| Cluster97 | AL6G04705 | 479454935 | 479456606 | AMP-binding domain containing protein |
| Cluster97 | AL6G04706 | 479514754 | 479526827 | Cytochrome P450 |
| Cluster97 | AL6G04707 | 479575459 | 479579751 | Cytochrome P450 |
| Cluster97 | AL6G04708 | 479583606 | 479587665 | Cytochrome P450 |
| Cluster97 | AL6G04709 | 479611297 | 479613327 | Transposon protein |
| Cluster98 | AL6G04927 | 491257735 | 491262739 | Ribonucleoside-diphosphate reductase large subunit |
| Cluster98 | AL6G04928 | 491266296 | 491268027 | Cytokinin-O-glucosyltransferase |
| Cluster98 | AL6G04929 | 491297092 | 491298633 | Putative subtilisin homologue |
| Cluster98 | AL6G04930 | 491303675 | 491310141 | Putative subtilisin homologue |
| Cluster98 | AL6G04931 | 491323381 | 491332024 | Other |
| Cluster98 | AL6G04932 | 491370258 | 491370710 | Cytochrome P450 |
| Cluster98 | AL6G04933 | 491370753 | 491371598 | Cytochrome P450 |
| Cluster98 | AL6G04934 | 491411656 | 491413254 | Cytochrome P450 |
| Cluster98 | AL6G04935 | 491417681 | 491419553 | Osfbx120-F-box domain |
| Cluster99 | AL6G04941 | 491619275 | 491621094 | Dehydrogenase |
| Cluster99 | AL6G04942 | 491628938 | 491630685 | Cytochrome P450 |
| Cluster99 | AL6G04943 | 491736666 | 491738764 | Sam dependent carboxyl methyltransferase |
| Cluster99 | AL6G04944 | 491775357 | 491777168 | Dehydrogenase |
| Cluster99 | AL6G04945 | 491782394 | 491795861 | Elmo/Ced-12 family protein |
| Cluster99 | AL6G04946 | 491816473 | 491820499 | Hydrolase |
| Cluster99 | AL6G04947 | 491820812 | 491825150 | Acetyltransferase |

| Cluster number | Gene ID | Start position | End position | Gene description |
|---|---|---|---|---|
| Cluster99 | AL6G04948 | 491865063 | 491867314 | Zinc finger domain |
| Cluster99 | AL6G04949 | 491925170 | 491926378 | Transferase family protein |
| Cluster99 | AL6G04950 | 491952688 | 491955572 | Other |
| Cluster100 | AL6G05030 | 494199611 | 494201299 | Pb1 domain |
| Cluster100 | AL6G05031 | 494205326 | 494206772 | Dehydrogenase |
| Cluster100 | AL6G05032 | 494210564 | 494211957 | Dehydrogenase |
| Cluster100 | AL6G05033 | 494237886 | 494240840 | G-beta repeat domain |
| Cluster100 | AL6G05034 | 494273477 | 494274657 | Chalcone synthase |
| Cluster100 | AL6G05035 | 494332135 | 494342437 | Oxidoreductase |
| Cluster100 | AL6G05036 | 494351187 | 494358309 | G-beta repeat domain |
| Cluster100 | AL6G05037 | 494363390 | 494365536 | Dehydrogenase |
| Cluster100 | AL6G05038 | 494444458 | 494445805 | Dehydrogenase |
| Cluster100 | AL6G05039 | 494476294 | 494477759 | Dehydrogenase |
| Cluster100 | AL6G05040 | 494502611 | 494508811 | Translation iInitiation factor |
| Cluster100 | AL6G05041 | 494513412 | 494515396 | Osfbx66-F-box domain |
| Cluster100 | AL6G05042 | 494543411 | 494544547 | Osfbx66-F-box domain |
| Cluster100 | AL6G05043 | 494551072 | 494554505 | Disease resistance protein |
| Cluster100 | AL6G05044 | 494578146 | 494587586 | Terpene synthase |
| Cluster100 | AL6G05045 | 494636064 | 494638502 | Cytochrome P450 |
| Cluster106 | AL7G03628 | 171954645 | 171957979 | G-beta repeat domain |
| Cluster106 | AL7G03629 | 171994764 | 171996099 | O-methyltransferase |
| Cluster106 | AL7G03630 | 172088255 | 172089610 | O-methyltransferase |
| Cluster106 | AL7G03631 | 172212662 | 172213234 | Dirigent, Putative, Expressed |
| Cluster106 | AL7G03632 | 172348632 | 172353710 | Cytochrome P450, Putative, Expressed |
| Cluster106 | AL7G03633 | 172468607 | 172469857 | O-methyltransferase |
| Cluster106 | AL7G03634 | 172487791 | 172489267 | Other |
| Cluster106 | AL7G03635 | 172538724 | 172560648 | O-methyltransferase |
| Cluster106 | AL7G03636 | 172619165 | 172620327 | O-methyltransferase |
| Cluster106 | AL7G03637 | 172626559 | 172632403 | Other |
| Cluster106 | AL7G03638 | 172644135 | 172649387 | Cycloeucalenol cycloisomerase |
| Cluster108 | AL7G04458 | 248472425 | 248479225 | Other |
| Cluster108 | AL7G04459 | 248483839 | 248486422 | Phytoene synthase |
| Cluster108 | AL7G04460 | 248497627 | 248499006 | Osfbx110-F-box domain containing protein |
| Cluster108 | AL7G04461 | 248627319 | 248629047 | Cytochrome P450 |
| Cluster108 | AL7G04462 | 248706261 | 248706981 | Late embryogenesis abundant protein |
| Cluster108 | AL7G04463 | 248722695 | 248723108 | Other |
| Cluster108 | AL7G04464 | 248924971 | 248928890 | Leucine rich repeat family protein |
| Cluster108 | AL7G04465 | 248959397 | 248964400 | Terpene synthase |
| Cluster108 | AL7G04466 | 249324985 | 249329817 | Terpene synthase |

**Table S23. The conserved synteny among *CYP450* genes within gene clusters in ALO and nine grass species.**

**a** Presence (1) of 24 *CYP450* genes from the synteny relationships with ALO in four *Avena* species and absence (0) of the genes from the synteny relationships with ALO and six plant species.

| Gene ID | AAT | AER | AST | ATA | BDI | OSA | SBI | SIT | ZMA |
|---|---|---|---|---|---|---|---|---|---|
| AL1G00972 | 1 | 1 | 1 | 0 | 0 | 0 | 0 | 0 | 0 |
| AL1G02602 | 1 | 1 | 1 | 1 | 1 | 1 | 0 | 1 | 0 |
| AL1G06104 | 1 | 1 | 1 | 0 | 0 | 0 | 0 | 0 | 0 |
| AL1G06294 | 1 | 1 | 1 | 0 | 0 | 0 | 0 | 0 | 0 |
| AL2G03969 | 1 | 1 | 1 | 0 | 0 | 0 | 0 | 0 | 0 |
| AL3G05636 | 1 | 1 | 1 | 0 | 0 | 0 | 0 | 0 | 0 |
| AL3G05655 | 1 | 1 | 1 | 0 | 0 | 0 | 0 | 0 | 0 |
| AL3G06285 | 1 | 1 | 1 | 1 | 0 | 0 | 0 | 0 | 0 |
| AL3G06293 | 1 | 1 | 1 | 1 | 0 | 0 | 0 | 0 | 0 |
| AL3G06343 | 1 | 1 | 1 | 0 | 0 | 0 | 0 | 0 | 0 |
| AL4G00019 | 1 | 1 | 1 | 0 | 0 | 0 | 0 | 0 | 0 |
| AL4G00048 | 1 | 1 | 1 | 1 | 0 | 0 | 0 | 0 | 0 |
| AL4G00052 | 1 | 1 | 1 | 0 | 0 | 0 | 0 | 0 | 0 |
| AL4G00858 | 1 | 1 | 1 | 1 | 1 | 0 | 0 | 1 | 0 |
| AL4G00907 | 1 | 1 | 1 | 1 | 0 | 0 | 0 | 0 | 0 |
| AL4G00928 | 1 | 1 | 1 | 1 | 0 | 0 | 0 | 0 | 0 |
| AL5G00241 | 1 | 1 | 1 | 0 | 0 | 0 | 0 | 0 | 0 |
| AL5G00242 | 1 | 1 | 1 | 0 | 0 | 0 | 0 | 0 | 0 |
| AL5G00413 | 1 | 1 | 1 | 0 | 0 | 0 | 0 | 0 | 0 |
| AL5G00913 | 1 | 1 | 1 | 1 | 1 | 0 | 1 | 1 | 1 |
| AL6G00559 | 1 | 1 | 1 | 0 | 0 | 0 | 0 | 0 | 0 |
| AL6G00560 | 1 | 1 | 1 | 0 | 0 | 0 | 0 | 0 | 0 |
| AL6G04704 | 1 | 1 | 1 | 0 | 0 | 0 | 0 | 0 | 0 |
| AL7G04461 | 1 | 1 | 1 | 0 | 0 | 0 | 0 | 0 | 0 |

**b** Presence (1) of six *CYP450* genes from the synteny relationships with ALO and nine grass species.

| Gene ID | AAT | AER | AST | ATA | BDI | OSA | SBI | SIT | ZMA |
|---|---|---|---|---|---|---|---|---|---|
| AL1G02600 | 1 | 1 | 1 | 1 | 1 | 1 | 1 | 1 | 1 |
| AL3G02465 | 1 | 1 | 1 | 1 | 1 | 1 | 1 | 1 | 1 |
| AL3G05629 | 1 | 1 | 1 | 1 | 1 | 1 | 1 | 1 | 1 |
| AL3G05648 | 1 | 1 | 1 | 1 | 1 | 1 | 1 | 1 | 1 |
| AL5G02319 | 1 | 1 | 1 | 1 | 1 | 1 | 1 | 1 | 1 |
| AL6G02061 | 1 | 1 | 1 | 1 | 1 | 1 | 1 | 1 | 1 |

**Table S24. Beta-glucan encoding 11 *CesA* and 55 *Csl* genes corresponding to the expanded gene families in the ALO genome.**

| Csl genes | Orthogroup | Expanded gene family in ALO | KEGG annotation |
|---|---|---|---|
| AL1G01201 | OG0008043 | OG0008043 | K13680 CSLA; beta-mannan synthase [EC:2.4.1.32] |
| AL1G02765 | OG0000702 | OG0000702 | None (CSLC in Fig. 5c) |
| AL1G02887 | OG0013205 | None | K10999 CESA; cellulose synthase A [EC:2.4.1.12] |
| AL1G05017 | None | None | None (non-group in Fig. 5c) |
| AL1G05019 | OG0018481 | None | None (CSLJ in Fig. 5c) |
| AL1G05406 | OG0008043 | None | K13680 CSLA; beta-mannan synthase [EC:2.4.1.32] |
| AL2G05226 | OG0009492 | None | K00721 DPM1; dolichol-phosphate mannosyltransferase [EC:2.4.1.83] |
| AL2G00588 | OG0025675 | None | None (CSLF in Fig. 5c) |
| AL2G00589 | OG0029677 | None | K20923 CSLF; mixed-linked glucan synthase [EC:2.4.1.-] |
| AL2G00609 | OG0025675 | None | K20923 CSLF; mixed-linked glucan synthase [EC:2.4.1.-] |
| AL2G00833 | None | None | K10999 CESA; cellulose synthase A [EC:2.4.1.12] |
| AL2G00834 | OG0002979 | OG0002979 | K10999 CESA; cellulose synthase A [EC:2.4.1.12] |
| AL2G00843 | OG0002979 | OG0002979 | K10999 CESA; cellulose synthase A [EC:2.4.1.12] |
| AL2G00965 | OG0004752 | None | K10999 CESA; cellulose synthase A [EC:2.4.1.12] |
| AL2G03166 | OG0013661 | None | K10999 CESA; cellulose synthase A [EC:2.4.1.12] |
| AL2G03980 | OG0001194 | OG0001194 | K20924 CSLD; cellulose synthase-like protein [EC:2.4.1.-] |
| AL2G04987 | OG0000702 | OG0000702 | None (CSLC in Fig. 5c) |
| AL3G01091 | OG0001474 | OG0001474 | None (CSLE in Fig. 5c) |
| AL3G01092 | OG0001474 | OG0001474 | None (CSLE in Fig. 5c) |
| AL3G01093 | OG0030718 | None | None (CSLE in Fig. 5c) |
| AL3G01094 | OG0001474 | OG0001474 | None (CSLE in Fig. 5c) |
| AL3G01095 | OG0001474 | OG0001474 | None (CSLE in Fig. 5c) |
| AL3G01096 | OG0001474 | OG0001474 | None (CSLE in Fig. 5c) |
| AL3G01097 | OG0001474 | OG0001474 | None (CSLE in Fig. 5c) |
| AL3G01441 | OG0009955 | None | K10999 CESA; cellulose synthase A [EC:2.4.1.12] |

| Csl genes | Orthogroup | Expanded gene family in ALO | KEGG annotation |
|---|---|---|---|
| AL3G03056 | OG0012597 | None | K00770 E2.4.2.24; 1,4-beta-D-xylan synthase [EC:2.4.2.24] |
| AL3G03394 | OG0007337 | None | K13680 CSLA; beta-mannan synthase [EC:2.4.1.32] |
| AL4G00954 | OG0030791 | None | K20924 CSLD; cellulose synthase-like protein [EC:2.4.1.-] |
| AL4G00955 | OG0001194 | OG0001194 | K20924 CSLD; cellulose synthase-like protein [EC:2.4.1.-] |
| AL4G00978 | OG0001194 | OG0001194 | K20924 CSLD; cellulose synthase-like protein [EC:2.4.1.-] |
| AL4G00979 | OG0030791 | None | K20924 CSLD; cellulose synthase-like protein [EC:2.4.1.-] |
| AL4G02826 | OG0014581 | None | None (CSLA in Fig. 5c) |
| AL4G03483 | OG0016973 | None | K20923 CSLF; mixed-linked glucan synthase [EC:2.4.1.-] |
| AL4G03484 | OG0009781 | None | K20923 CSLF; mixed-linked glucan synthase [EC:2.4.1.-] |
| AL4G03485 | OG0003005 | None | K20923 CSLF; mixed-linked glucan synthase [EC:2.4.1.-] |
| AL4G04310 | OG0000702 | OG0000702 | K20923 CSLF; mixed-linked glucan synthase [EC:2.4.1.-] |
| AL4G04312 | OG0000702 | OG0000702 | None (CSLC in Fig. 5c) |
| AL4G05011 | OG0006430 | None | None (CSLH in Fig. 5c) |
| AL5G00055 | OG0003005 | None | K20923 CSLF; mixed-linked glucan synthase [EC:2.4.1.-] |
| AL5G00190 | OG0001194 | None | K20924 CSLD; cellulose synthase-like protein [EC:2.4.1.-] |
| AL5G01190 | OG0001102 | None | K13680 CSLA; beta-mannan synthase [EC:2.4.1.32] |
| AL5G02454 | OG0008988 | None | K13680 CSLA; beta-mannan synthase [EC:2.4.1.32] |
| AL5G02634 | OG0004286 | None | None (CSLC in Fig. 5c) |
| AL5G02870 | OG0014528 | None | K20923 CSLF; mixed-linked glucan synthase [EC:2.4.1.-] |
| AL5G03480 | OG0015451 | None | K20924 CSLD; cellulose synthase-like protein [EC:2.4.1.-] |
| AL5G03868 | OG0012659 | None | K10999 CESA; cellulose synthase A [EC:2.4.1.12] |
| AL5G04027 | None | None | K13680 CSLA; beta-mannan synthase [EC:2.4.1.32] |

| Csl genes | Orthogroup | Expanded gene family in ALO | KEGG annotation |
|---|---|---|---|
| AL6G01955 | OG0001102 | None | K13680 CSLA; beta-mannan synthase [EC:2.4.1.32] |
| AL6G02856 | OG0011300 | None | None (CSLE in Fig. 5c) |
| AL6G02857 | OG0011300 | None | None (CSLE in Fig. 5c) |
| AL6G02863 | OG0011300 | None | None (CSLE in Fig. 5c) |
| AL6G04010 | OG0007969 | OG0007969 | K13680 CSLA; beta-mannan synthase [EC:2.4.1.32] |
| AL6G04018 | OG0007219 | None | K20923 CSLF; mixed-linked glucan synthase [EC:2.4.1.-] |
| AL6G04028 | OG0007969 | None | K13680 CSLA; beta-mannan synthase [EC:2.4.1.32] |
| AL6G04043 | None | None | K20923 CSLF; mixed-linked glucan synthase [EC:2.4.1.-] |
| AL6G04044 | OG0030937 | None | K20923 CSLF; mixed-linked glucan synthase [EC:2.4.1.-] |
| AL6G04053 | OG0007219 | None | K20923 CSLF; mixed-linked glucan synthase [EC:2.4.1.-] |
| AL6G04059 | OG0007219 | None | K20923 CSLF; mixed-linked glucan synthase [EC:2.4.1.-] |
| AL6G04060 | OG0028265 | None | K20923 CSLF; mixed-linked glucan synthase [EC:2.4.1.-] |
| AL6G05360 | OG0001474 | None | None (CSLE in Fig. 5c) |
| AL7G01647 | OG0001300 | None | K10999 CESA; cellulose synthase A [EC:2.4.1.12] |
| AL7G01648 | OG0001300 | None | K10999 CESA; cellulose synthase A [EC:2.4.1.12] |
| AL7G01826 | OG0009492 | None | K00721 DPM1; dolichol-phosphate mannosyltransferase [EC:2.4.1.83] |
| AL7G01858 | OG0016501 | None | K00729 ALG5; dolichyl-phosphate beta-glucosyltransferase [EC:2.4.1.117] |
| AL7G02010 | OG0002979 | None | K10999 CESA; cellulose synthase A [EC:2.4.1.12] |
| AL7G02472 | OG0000702 | None | None (CSLC in Fig. 5c) |

**Table S25. Distribution of 11 *CesA* and 55 *Csl* genes in the ALO chromosomes.**

| Chromosome no. | Number of CESA and Csl genes | Percentage of total CESA and Csl genes |
| --- | --- | --- |
| ALO01 | 6 | 9.09% |
| ALO02 | 11 | 16.67% |
| ALO03 | 10 | 15.15% |
| ALO04 | 11 | 16.67% |
| ALO05 | 9 | 13.64% |
| ALO06 | 13 | 19.70% |
| ALO07 | 6 | 9.09% |
| Total | 66 | 100% |

**Table S26. FPKM value of *CesA* and *Csl* genes in roots, leaves, salt-treated roots and salt-treated leaves of ALO.**

**a The transcript levels of *CesA* and *Csl* genes over 1-fold higher in salt-treated roots than in roots and FPKM value presence in salt-treated roots while zero in roots.**

| Gene ID | Roots | Salt-treated roots | Fold number (Salt-treated roots / roots) | Leaves | Salt-treated leaves |
|---|---|---|---|---|---|
| AL4G03484 | 17.36 | 123.25 | 7.10 | 0.06 | 0.26 |
| AL4G05011 | 1.83 | 12.27 | 6.70 | 18.73 | 2.47 |
| AL5G00190 | 45.23 | 195.60 | 4.32 | 18.78 | 7.17 |
| AL3G01096 | 11.51 | 43.69 | 3.79 | 46.96 | 6.54 |
| AL7G02472 | 26.54 | 71.55 | 2.70 | 4.93 | 5.51 |
| AL2G03980 | 0.69 | 1.52 | 2.21 | 0.75 | 1.38 |
| AL7G02010 | 3.66 | 5.73 | 1.56 | 19.93 | 12.95 |
| AL1G05019 | 42.83 | 51.83 | 1.21 | 20.79 | 12.45 |
| AL2G00589 | 0.72 | 0.81 | 1.13 | 5.93 | 5.15 |
| AL6G02863 | 6.76 | 6.38 | 0.94 | 4.58 | 2.64 |
| AL4G00954 | 0.00 | 0.18 | None | 0.10 | 0.75 |
| AL4G00979 | 0.00 | 0.18 | None | 0.10 | 0.75 |

**b The transcript levels of *CesA* and *Csl* genes higher in salt-treated roots than in roots.**

| Gene ID | Roots | Salt-treated roots | Difference of FPKM value in salt-treated roots and roots | Leaves | Salt-treated leaves |
|---|---|---|---|---|---|
| AL5G00190 | 45.23 | 195.60 | 150.37 | 4.32 | 18.78 |
| AL4G03484 | 17.36 | 123.25 | 105.89 | 7.10 | 0.06 |
| AL7G02472 | 26.54 | 71.55 | 45.01 | 2.70 | 4.93 |
| AL3G01096 | 11.51 | 43.69 | 32.18 | 3.79 | 46.96 |
| AL4G05011 | 1.83 | 12.27 | 10.44 | 6.70 | 18.73 |
| AL1G05019 | 42.83 | 51.83 | 8.99 | 1.21 | 20.79 |
| AL7G02010 | 3.66 | 5.73 | 2.07 | 1.56 | 19.93 |
| AL2G03980 | 0.69 | 1.52 | 0.84 | 2.21 | 0.75 |
| AL4G00954 | 0.00 | 0.18 | 0.18 | None | 0.10 |
| AL4G00979 | 0.00 | 0.18 | 0.18 | None | 0.10 |
| AL2G00589 | 0.72 | 0.81 | 0.10 | 1.13 | 5.93 |

**c The transcript levels of *CesA* and *Csl* genes over 1-fold higher in salt-treated leaves than in leaves.**

| Gene ID | Leaves | Salt-treated leaves | Fold number (Salt-treated leaves / leaves) | Roots | Salt-treated roots |
|---|---|---|---|---|---|
| AL3G01097 | 0.01 | 0.42 | 33.26 | 4.33 | 2.76 |
| AL4G02826 | 0.59 | 7.44 | 12.64 | 14.96 | 3.86 |
| AL4G00954 | 0.10 | 0.75 | 7.27 | 0.00 | 0.18 |
| AL4G00979 | 0.10 | 0.75 | 7.27 | 0.00 | 0.18 |
| AL4G03484 | 0.06 | 0.26 | 4.10 | 17.36 | 123.25 |
| AL5G04027 | 0.38 | 1.00 | 2.61 | 0.73 | 0.28 |
| AL7G01826 | 8.97 | 22.76 | 2.54 | 28.25 | 21.27 |
| AL6G02856 | 1.43 | 3.09 | 2.15 | 3.61 | 1.88 |
| AL5G02870 | 0.15 | 0.31 | 2.00 | 56.60 | 15.74 |
| AL2G03980 | 0.75 | 1.38 | 1.83 | 0.69 | 1.52 |
| AL7G01858 | 13.96 | 18.68 | 1.34 | 21.85 | 17.57 |
| AL2G05226 | 5.16 | 6.18 | 1.20 | 16.07 | 14.60 |
| AL4G03485 | 0.72 | 0.85 | 1.19 | 11.08 | 0.04 |
| AL7G02472 | 4.93 | 5.51 | 1.12 | 26.54 | 71.55 |
| AL2G03166 | 1.73 | 1.73 | 1.00 | 196.57 | 17.33 |

**d The transcript levels of *CesA* and *Csl* genes higher in salt-treated leaves than in leaves.**

| Gene ID | Roots | Salt-treated roots | Difference of FPKM value in salt-treated roots and roots | Leaves | Salt-treated leaves |
|---|---|---|---|---|---|
| AL7G01826 | 8.97 | 22.76 | 13.79 | 2.54 | 28.25 |
| AL4G02826 | 0.59 | 7.44 | 6.85 | 12.64 | 14.96 |
| AL7G01858 | 13.96 | 18.68 | 4.72 | 1.34 | 21.85 |
| AL6G02856 | 1.43 | 3.09 | 1.65 | 2.15 | 3.61 |
| AL2G05226 | 5.16 | 6.18 | 1.02 | 1.20 | 16.07 |
| AL4G00954 | 0.10 | 0.75 | 0.65 | 7.27 | 0.00 |
| AL4G00979 | 0.10 | 0.75 | 0.65 | 7.27 | 0.00 |
| AL2G03980 | 0.75 | 1.38 | 0.63 | 1.83 | 0.69 |
| AL5G04027 | 0.38 | 1.00 | 0.62 | 2.61 | 0.73 |
| AL7G02472 | 4.93 | 5.51 | 0.59 | 1.12 | 26.54 |
| AL3G01097 | 0.01 | 0.42 | 0.41 | 33.26 | 4.33 |
| AL4G03484 | 0.06 | 0.26 | 0.20 | 4.10 | 17.36 |
| AL5G02870 | 0.15 | 0.31 | 0.15 | 2.00 | 56.60 |
| AL4G03485 | 0.72 | 0.85 | 0.13 | 1.19 | 11.08 |

**Table S27. Gene description of 10 biosynthetic gene clusters (BGCs) containing *CesA* and *Csl* genes in the expression profiling heat map.**

| Cluster number | Gene ID | Start position | End position | Gene description |
|---|---|---|---|---|
| Cluster12 | AL2G00585 | 26210658 | 26213897 | YL1 nuclear protein C-terminal domain |
| Cluster12 | AL2G00586 | 26216226 | 26217323 | Other |
| Cluster12 | AL2G00587 | 26234303 | 26241947 | Dirigent |
| Cluster12 | AL2G00588 | 26355391 | 26356605 | CslF9 |
| Cluster12 | AL2G00589 | 26356772 | 26385660 | CslF8 |
| Cluster12 | AL2G00590 | 26389327 | 26392885 | Glyoxalase |
| Cluster12 | AL2G00591 | 26396267 | 26401540 | Oxidoreductase |
| Cluster12 | AL2G00592 | 26403383 | 26404554 | Dehydrogenase |
| Cluster12 | AL2G00593 | 26404958 | 26406401 | Dioxygenase |
| Cluster12 | AL2G00594 | 26421140 | 26421484 | NB-ARC domain |
| Cluster12 | AL2G00595 | 26423883 | 26424380 | NB-ARC domain |
| Cluster12 | AL2G00596 | 26485034 | 26492699 | MYB family transcription factor |
| Cluster12 | AL2G00597 | 26510311 | 26511742 | Dioxygenase |
| Cluster12 | AL2G00598 | 26512895 | 26514479 | G-patch domain |
| Cluster12 | AL2G00599 | 26594352 | 26595282 | Aquaporin protein |
| Cluster12 | AL2G00600 | 26663374 | 26665463 | Dioxygenase |
| Cluster12 | AL2G00601 | 26707705 | 26710024 | Dioxygenase |
| Cluster12 | AL2G00602 | 26717480 | 26717779 | Other |
| Cluster12 | AL2G00603 | 26721214 | 26721558 | Zinc finger protein |
| Cluster13 | AL2G00604 | 26927219 | 26930450 | YL1 nuclear protein C-terminal domain |
| Cluster13 | AL2G00605 | 26932778 | 26933881 | Other genes |
| Cluster13 | AL2G00606 | 26939207 | 26940131 | Dirigent |
| Cluster13 | AL2G00607 | 26945581 | 26946462 | Dirigent |
| Cluster13 | AL2G00608 | 26953007 | 26953905 | Dirigent |
| Cluster13 | AL2G00609 | 27037845 | 27038870 | CslF9 |
| Cluster13 | AL2G00610 | 27052749 | 27056306 | Glyoxalase |
| Cluster13 | AL2G00611 | 27057891 | 27062195 | Oxidoreductase |
| Cluster13 | AL2G00612 | 27064961 | 27066176 | Dehydrogenase |
| Cluster13 | AL2G00613 | 27068907 | 27080995 | Terpene synthase |
| Cluster13 | AL2G00614 | 27308103 | 27319978 | Terpene synthase |
| Cluster13 | AL2G00615 | 27345969 | 27346860 | Dioxygenase |
| Cluster13 | AL2G00616 | 27396290 | 27397138 | Other |
| Cluster17 | AL2G00827 | 40006436 | 40014451 | Helicase |
| Cluster17 | AL2G00828 | 40020980 | 40024677 | Other |
| Cluster17 | AL2G00829 | 40028288 | 40030518 | Cytochrome P450 |
| Cluster17 | AL2G00830 | 40057039 | 40061721 | Glycosyl transferase |
| Cluster17 | AL2G00831 | 40086692 | 40094026 | Dioxygenase |
| Cluster17 | AL2G00832 | 40209332 | 40210758 | Proline-rich protein |
| Cluster17 | AL2G00833 | 40233295 | 40235511 | CESA8-cellulose synthase |

| Cluster number | Gene ID | Start position | End position | Gene description |
|---|---|---|---|---|
| Cluster17 | AL2G00834 | 40235553 | 40238348 | CESA2-cellulose synthase |
| Cluster17 | AL2G00835 | 40268987 | 40269741 | LTPL79 Protease inhibitor |
| Cluster17 | AL2G00836 | 40292412 | 40293299 | Oxidoreductase |
| Cluster17 | AL2G00837 | 40301453 | 40304261 | Zinc finger protein |
| Cluster25 | AL2G03967 | 476203060 | 476205454 | Dehydrogenase |
| Cluster25 | AL2G03968 | 476206378 | 476209811 | Transporter family protein |
| Cluster25 | AL2G03969 | 476211346 | 476213605 | Cytochrome 450 |
| Cluster25 | AL2G03970 | 476264534 | 476265775 | Oxidoreductase |
| Cluster25 | AL2G03971 | 476290692 | 476295659 | Dehydration protein-related |
| Cluster25 | AL2G03972 | 476338372 | 476346659 | CoA-ligase |
| Cluster25 | AL2G03973 | 476348985 | 476351033 | Cytidylyl transferase |
| Cluster25 | AL2G03974 | 476351749 | 476354027 | lrgB-like family protein |
| Cluster25 | AL2G03975 | 476443872 | 476452762 | lrgB-like family protein |
| Cluster25 | AL2G03976 | 476481954 | 476482456 | Plastocyanin-like domain |
| Cluster25 | AL2G03977 | 476488799 | 476489947 | Methyltransferase |
| Cluster25 | AL2G03978 | 476539185 | 476541947 | Proteasome |
| Cluster25 | AL2G03979 | 476543359 | 476545236 | PPR repeat domain |
| Cluster25 | AL2G03980 | 476546530 | 476550760 | CslLD1 |
| Cluster32 | AL3G03047 | 350293759 | 350296963 | Polygalacturonase |
| Cluster32 | AL3G03048 | 350349790 | 350359264 | Retrotransposon protein |
| Cluster32 | AL3G03049 | 350420954 | 350421412 | StAR-related lipid-transfer |
| Cluster32 | AL3G03050 | 350592696 | 350593025 | StAR-related lipid-transfer |
| Cluster32 | AL3G03051 | 350632489 | 350632932 | StAR-related lipid-transfer |
| Cluster32 | AL3G03052 | 350766648 | 350767133 | StAR-related lipid-transfer |
| Cluster32 | AL3G03053 | 350788915 | 350789232 | Retrotransposon protein |
| Cluster32 | AL3G03054 | 351041877 | 351042356 | StAR-related lipid-transfer |
| Cluster32 | AL3G03055 | 351052667 | 351053158 | StAR-related lipid-transfer |
| Cluster32 | AL3G03056 | 351060796 | 351064640 | CslD4 |
| Cluster58 | AL4G02824 | 130570852 | 130571397 | Zinc finger protein |
| Cluster58 | AL4G02825 | 130589022 | 130593803 | Other |
| Cluster58 | AL4G02826 | 130594694 | 130599387 | CslA7 |
| Cluster58 | AL4G02827 | 130696059 | 130700455 | Other |
| Cluster58 | AL4G02828 | 130709001 | 130709841 | Ribonuclease T2 family |
| Cluster58 | AL4G02829 | 130721037 | 130721375 | Other |
| Cluster58 | AL4G02830 | 130754314 | 130754811 | Gibberellin 2-beta-dioxygenase |
| Cluster58 | AL4G02831 | 130763032 | 130764606 | Gibberellin 2-beta-dioxygenase |
| Cluster69 | AL5G00054 | 3525796 | 3532036 | Methyltransferase |
| Cluster69 | AL5G00055 | 3536261 | 3541931 | CslF3 |
| Cluster69 | AL5G00056 | 3542358 | 3544874 | Methyltransferase |
| Cluster69 | AL5G00057 | 3547297 | 3551156 | Other |
| Cluster69 | AL5G00058 | 3553154 | 3553712 | StAR-related lipid-transfer |
| Cluster69 | AL5G00059 | 3555689 | 3560271 | ATPase |

| Cluster number | Gene ID | Start position | End position | Gene description |
|---|---|---|---|---|
| Cluster87 | AL5G01187 | 54882771 | 54883953 | Lipase/acylhydrolase |
| Cluster87 | AL5G01188 | 54935083 | 54940122 | Glycosyl transferase |
| Cluster87 | AL5G01189 | 54944496 | 54948839 | Other |
| Cluster87 | AL5G01190 | 55006809 | 55010398 | CslA3 |
| Cluster87 | AL5G01191 | 55121165 | 55132111 | Myosin |
| Cluster87 | AL5G01192 | 55156384 | 55157306 | Dirigent |
| Cluster87 | AL5G01193 | 55234023 | 55235567 | Pentatricopeptide |
| Cluster87 | AL5G01194 | 55242155 | 55243744 | Glycosyl transferase |
| Cluster87 | AL5G01195 | 55272180 | 55274403 | Other |
| Cluster87 | AL5G01196 | 55275803 | 55277602 | Other |
| Cluster102 | AL6G05359 | 508898953 | 508905454 | Dirigent |
| Cluster102 | AL6G05360 | 508922653 | 508927715 | CslE6 |
| Cluster102 | AL6G05361 | 508939332 | 508940429 | Glycosyl transferase |
| Cluster102 | AL6G05362 | 508969560 | 508971606 | Oxidoreductase |
| Cluster102 | AL6G05363 | 509003567 | 509005015 | Glycosyl transferase |
| Cluster104 | AL7G01822 | 85408689 | 85412518 | Protein kinase family |
| Cluster104 | AL7G01823 | 85445144 | 85446088 | Gibberellin receptor GID1L2 |
| Cluster104 | AL7G01824 | 85457834 | 85459301 | Glucosyl transferase |
| Cluster104 | AL7G01825 | 85537824 | 85545367 | Membrane protein |
| Cluster104 | AL7G01826 | 85583649 | 85588086 | Glycosyl transferase |

**Table S28. The conserved synteny among *CesA* and *Csl* genes within gene clusters in ALO and nine grass species.**

**a** Presence (1) of one *CesA* and three *Csl* genes from the synteny relationships with ALO and four *Avena* species and absence (0) of the genes from the synteny relationships with ALO and six plant species.

| Gene ID | AAT | AER | AST | ATA | BDI | OSA | SBI | SIT | ZMA |
|---|---|---|---|---|---|---|---|---|---|
| AL2G00833 | 1 | 1 | 1 | 1 | 0 | 0 | 0 | 0 | 0 |
| AL5G00055 | 1 | 1 | 1 | 0 | 0 | 1 | 0 | 0 | 0 |
| AL5G01190 | 1 | 1 | 1 | 0 | 1 | 1 | 1 | 1 | 1 |
| AL7G01826 | 1 | 1 | 1 | 1 | 0 | 1 | 1 | 1 | 1 |

**b** Presence (1) of two *Csl* genes from the synteny relationships with ALO and nine grass species.

| Gene ID | AAT | AER | AST | ATA | BDI | OSA | SBI | SIT | ZMA |
|---|---|---|---|---|---|---|---|---|---|
| AL3G03056 | 1 | 1 | 1 | 1 | 1 | 1 | 1 | 1 | 1 |
| AL4G02826 | 1 | 1 | 1 | 1 | 1 | 1 | 1 | 1 | 1 |

**Table S29. Statistics of 13 *CalS* (*GSL*) genes corresponding the expanded gene families in the ALO genome.**

| Cals genes | Orthogroup | Expanded gene family in ALO | KEGG annotation |
|---|---|---|---|
| AL1G02931 | OG0002915 | None | K11000CALS; callosesynthase [EC:2.4.1] |
| AL1G03358 | OG0002915 | None | K11000CALS; callosesynthase [EC:2.4.1] |
| AL1G03911 | OG0009427 | None | K11000CALS; callosesynthase [EC:2.4.1] |
| AL1G05549 | OG0001280 | OG0001280 | K11000CALS; callosesynthase [EC:2.4.1] |
| AL3G02326 | OG0018416 | None | K11000CALS; callosesynthase [EC:2.4.1] |
| AL3G05781 | OG0001280 | OG0001280 | K11000CALS; callosesynthase [EC:2.4.1] |
| AL3G05825 | OG0001280 | OG0001280 | K11000CALS; callosesynthase [EC:2.4.1] |
| AL3G05902 | OG0003707 | OG0003707 | K11000CALS; callosesynthase [EC:2.4.1] |
| AL5G00002 | OG0003707 | OG0003707 | K11000CALS; callosesynthase [EC:2.4.1] |
| AL5G00040 | OG0003707 | None | K11000CALS; callosesynthase [EC:2.4.1] |
| AL5G00849 | OG0010884 | None | K11000CALS; callosesynthase [EC:2.4.1] |
| AL6G02277 | OG0014473 | None | K11000CALS; callosesynthase [EC:2.4.1] |
| AL7G00888 | OG0001280 | None | K11000CALS; callosesynthase [EC:2.4.1] |

#### Table S30. FPKM value of *CalS* genes in roots, leaves, salt-treated roots and salt-treated leaves of ALO.

**a The transcript levels of *CalS* genes over 1-fold higher in salt-treated roots than in roots and FPKM value presence in salt-treated roots while zero in roots.**

| Gene ID | Roots | Salt-treated roots | Fold number (Salt-treated roots / roots) | Leaves | Salt-treated leaves |
|---|---|---|---|---|---|
| AL5G00849 | 0.00 | 0.02 | None | 0.00 | 0.00 |
| AL6G02277 | 1.58 | 17.42 | 11.04 | 33.46 | 3.84 |
| AL1G03911 | 4.83 | 43.14 | 8.93 | 78.85 | 4.76 |
| AL1G03358 | 2.85 | 10.47 | 3.68 | 17.96 | 0.48 |
| AL3G02326 | 1.58 | 4.78 | 3.03 | 2.48 | 2.95 |
| AL1G05549 | 11.91 | 30.64 | 2.57 | 55.56 | 19.96 |
| AL7G00888 | 19.42 | 43.52 | 2.24 | 27.87 | 12.61 |
| AL3G05781 | 14.00 | 26.68 | 1.91 | 55.21 | 13.36 |
| AL1G02931 | 3.42 | 6.34 | 1.85 | 12.52 | 1.69 |
| AL5G00002 | 2.35 | 4.17 | 1.77 | 7.99 | 1.64 |
| AL3G05825 | 20.59 | 29.19 | 1.42 | 68.39 | 13.24 |
| AL5G00040 | 19.75 | 26.29 | 1.33 | 34.27 | 14.25 |

**b The folder number of transcript levels of *CalS* genes higher in salt-treated leaves than in leaves.**

| Gene ID | Leaves | Salt-treated leaves | Fold number (Salt-treated leaves / leaves) | Roots | Salt-treated leaves |
|---|---|---|---|---|---|
| AL5G00849 | 0.00 | 0.00 | None | 0.00 | 0.02 |
| AL3G02326 | 2.48 | 2.95 | 1.19 | 1.58 | 4.78 |
| AL7G00888 | 27.87 | 12.61 | 0.45 | 19.42 | 43.52 |
| AL5G00040 | 34.27 | 14.25 | 0.42 | 19.75 | 26.29 |
| AL1G05549 | 55.56 | 19.96 | 0.36 | 11.91 | 30.64 |
| AL3G05902 | 40.17 | 11.62 | 0.29 | 22.26 | 29.53 |
| AL3G05781 | 55.21 | 13.36 | 0.24 | 14.00 | 26.68 |
| AL5G00002 | 7.99 | 1.64 | 0.21 | 2.35 | 4.17 |
| AL3G05825 | 68.39 | 13.24 | 0.19 | 20.59 | 29.19 |
| AL1G02931 | 12.52 | 1.69 | 0.14 | 3.42 | 6.34 |
| AL6G02277 | 33.46 | 3.84 | 0.11 | 1.58 | 17.42 |
| AL1G03911 | 78.85 | 4.76 | 0.06 | 4.83 | 43.14 |
| AL1G03358 | 17.96 | 0.48 | 0.03 | 2.85 | 10.47 |
